# Genetic mapping and genomic prediction for agronomic, grain compositional, and sensing-enabled traits in a cowpea MAGIC population along an environmental gradient

**DOI:** 10.64898/2026.08.04.742818

**Authors:** Jonathan M. Berlingeri, Sassoum Lo, Margaret Riggs, Heesup Yun, Hamid Kamangir, Earl Ranario, Isaac Kazuo Uyehara, Ismael Mayanja, Astrid Lao, Isaac Onziga Dramadri, Patrick Obia Ongom, Ousmane Boukar, Antonia Palkovic, Brian N. Bailey, J. Mason Earles, Bao-Lam Huynh, Christine Diepenbrock

## Abstract

Cowpea (*Vigna unguiculata* [L.] Walp.) is a resilient grain legume and an important global source of dietary protein, yet the genetic and environmental basis of phenological and canopy development, as well as grain composition, remains incompletely characterized across production environments. In this study, we evaluated a cowpea multi-parent advanced generation intercross (MAGIC) population along an environmental gradient in California (with contrasting daylengths, temperatures, and soil types) using agronomic, grain compositional, and uncrewed aerial vehicle (UAV) and rover-enabled phenotyping. Near-infrared spectroscopy (NIRS) enabled assessment of grain compositional traits, while sensing-enabled time-series imaging captured canopy and reproductive dynamics. Quantitative trait locus (QTL) mapping identified 267 QTL, and genome-wide association studies (GWAS) detected 1,973 marker-trait associations. Integrating QTL mapping and GWAS results identified two major genomic hotspots affecting multiple traits. A chromosome 9 hotspot (5.8–6.0 Mb) was associated with flowering time and co-localized with sensing-enabled measures of flower and pod counts, plant height, and vegetation fraction, indicating broad effects on phenological and canopy development. A chromosome 8 hotspot (37.3–37.9 Mb) contained co-localized signals for seed weight, protein, starch, phytate, and moisture. A total of 22 prioritized candidate genes were identified within these and other loci with multi-environment QTL and GWAS support. Genomic predictive abilities were moderate to high for most traits and scenarios, with multi-trait MegaLMM outperforming RR-BLUP. Together, these results define major genomic regions controlling cowpea phenology, canopy development, and grain composition, and provide targets and strategies for breeding cowpea cultivars with favorable and environmentally resilient productivity and grain composition.

**Significance Statement:** To dissect the genetic basis of cowpea productivity, adaptation, and grain composition, and how performance for these traits varies and can be predicted across environments, we combined multi-environment phenotyping, including sensing of canopy and reproductive traits, with quantitative genetic analyses in a multi-parental population. We identified genomic hotspots for seed size/composition and reproductive phenology and an across-environment predictive advantage for multi-trait vs. single-trait genomic prediction. Overall, these findings support the comprehensive improvement of cowpea.

## Introduction

Cowpea (*Vigna unguiculata* [L.] Walp.) is a major grain legume cultivated in tropical, subtropical, and semi-arid regions of the world. Approximately 95% of global production occurs in Sub-Saharan Africa (*FAOSTAT*), where cowpea is a major component of smallholder farming systems. Cowpea is also produced in the United States, particularly in California and Texas, with a five-year average U.S. farmgate value of USD $9.2 million (*USDA/NASS QuickStats*). Cowpea is recognized for its ability to tolerate challenging environmental conditions (Carvalho et al., 2017; Timko & Singh, 2008). Nonetheless, high temperatures, limited or erratic rainfall, and low soil fertility are ongoing abiotic production constraints for cowpea and are targets for further stress tolerance improvement (A. E. Hall, 2012; Boukar et al., 2018; Omomowo & Babalola, 2021; Mekonnen et al., 2022). The reproductive stages (particularly flowering and pod filling) of cowpea production are more sensitive to heat and drought, with greater impacts on pod set, seed size, grain yield, and grain quality than comparable stresses imposed during vegetative stages (Hall, 1980; Ismail & Hall, 1998; Nevhulaudzi et al., 2020; Poudel et al., 2025). Nighttime temperature is a key factor influencing reproductive heat stress in cowpea, with elevated nighttime temperatures reducing both pod set and grain yield by approximately 4-14% for each degree above 16 °C during reproductive development (Ismail & Hall, 1998).

Reproductive phenology is a key determinant of adaptation in cowpea and strongly influences the environmental conditions experienced during reproductive development (Ehlers & Hall, 1997; Hall et al., 1997). Cowpea is generally considered a short-day species, although photoperiod sensitivity varies among genotypes and can strongly influence adaptation across environments (Craufurd, et al., 1996a; Craufurd, et al., 1996b; Ehlers & Hall, 1996). Flowering time varies widely across cowpea germplasm, ranging from approximately 30 to more than 100 days after planting (DAP), and is regulated by photoperiod and temperature (Huynh et al., 2018; Paudel et al., 2021). Numerous studies have documented significant effects of genotype, environment, and genotype-by-environment interactions on flowering time, and marker-trait associations with days to flowering have been identified across much of the cowpea genome in the University of California, Riverside (UCR) mini-core (Muñoz-Amatriaín et al., 2021).

Genomic prediction studies of growth habit, maturity, and flowering time traits in cowpea have resulted in moderate accuracies (Ravelombola et al., 2021). Variation in phenology may indirectly influence yield components and grain composition. Recent advances in field-based sensing technologies such as uncrewed aerial vehicle (UAV) and ground-based rover platforms, high-resolution imaging, and computer vision-based object detection models enable quantification of reproductive development, yield components, and canopy dynamics at high temporal resolution within scalable field workflows (Ninomiya, 2022; Berlingeri et al., 2025). Cowpea architecture is well suited for such approaches because the majority of flowers and pods are positioned above the leaf canopy (i.e., less occluded than in other grain legume species) and can be reliably detected using downward-facing imaging systems. Time-series measurements of reproductive traits including flower and pod counts (Kamangir et al., 2026), together with canopy traits such as green pixel fraction and canopy height, enable detailed characterization of developmental trajectories and yield formation. Integrating these dynamic phenotypes with genetic analyses enables the identification of loci influencing phenology, yield components, and grain composition, as well as the development of GP models within and across these traits.

Cowpea is considered a nutritious grain legume and an important source of dietary protein (Timko & Singh, 2008; Jayathilake et al., 2018; Kim et al., 2025). Raw cowpea grains contain approximately 60% carbohydrates and 20–30% protein, and are also a source of dietary fiber and essential minerals such as iron (Fe), zinc (Zn), magnesium (Mg), and calcium (Ca) (USDA, 2018). Considerable variation in grain composition exists among cowpea genotypes, and improvement of grain nutritional quality has been a priority in cowpea breeding programs (Timko & Singh, 2008; Boukar et al., 2011; Dhanasekar et al., 2021). This variation is further shaped by environmental conditions and management practices. Previous studies have demonstrated significant genotype-by-environment (GxE), genotype-by-management (GxM), and three-way interaction (G×E×M) effects on nutritional traits including protein and mineral content (Gerrano et al., 2019), indicating that the expression of favorable nutritional alleles is at times environmentally contingent. Characterizing the genetic determinants of these traits, and the stability of genetic effects across environments, is therefore essential for designing breeding strategies that reliably improve nutritional outcomes across diverse production contexts and shifting environmental conditions. Unfortunately, progress toward this goal has been constrained by the labor and costs associated with traditional wet-chemistry phenotyping, which limit the scale of germplasm evaluation and the number of environments that can be surveyed in a single study.

Near-infrared spectroscopy (NIRS) calibrations developed for cowpea enable rapid prediction of grain protein and other macronutrients, making large-scale grain compositional phenotyping of diverse germplasm across multiple environments feasible (Muranaka et al., 2015; Padhi et al., 2022). When combined with multi-environment trials (MET), NIRS-based phenotyping can generate the data density required to estimate variance components, quantify G×E, and conduct quantitative genetic analyses with adequate statistical resolution. Genetic mapping studies have identified loci associated with protein and mineral accumulation across multiple regions of the cowpea genome (Huynh et al., 2024; Akinmade et al., 2026), and genomic prediction (GP) approaches have been applied and yielded moderate predictive accuracy (Olatoye et al., 2019). Nevertheless, the stability of quantitative trait loci (QTL) for grain compositional traits across production environments and the cross-environment generalizability of GP models for grain composition alongside agronomics remain largely unresolved in cowpea.

In this study, we conducted intensive phenotyping of a cowpea multi-parent advanced generation intercross (MAGIC) population across multiple environments in California to investigate the genetic architecture of grain compositional traits, canopy dynamics, and reproductive development. This 305-line MAGIC population was developed by Huynh et al. (2018) using seven parents representing the west and southeast African gene pools and one high-temperature tolerant California blackeye cultivar. We field-evaluated all 305 lines under short-day conditions in the Coachella Valley of southern California (Thermal, CA) for two years. Additionally, we evaluated the 168-line photoperiod-insensitive subset under long-day conditions in the Sacramento Valley of northern California (Davis, CA) and the central San Joaquin Valley of California (Parlier, CA), each for two years. Agronomic traits (stand count, flowering time, plant height, grain yield, and 100-seed weight) were collected manually. Sensing was conducted in Davis and Parlier at weekly intervals, and twice per week during the peak flowering window.

Traits extracted from sensing data were rover-enabled flower and pod counts (Kamangir et al., 2026) and UAV-enabled vegetation fraction and plant height. Seed compositional traits (protein, starch, fat, ash, moisture, and phytate percentages) were predicted via NIRS and accompanying wet-chemistry assays on a subset of samples per location-year to build custom calibrations.

Spatial model fitting, QTL mapping, conditional scans as a screen for pleiotropy, GWAS, and single- and multi-trait GP were then conducted for agronomic, sensing-enabled, and seed compositional traits. By combining MET and genetic analyses, we aimed to quantify genotypic, environmental, and G×E effects, identify genomic loci associated with key traits, and assess the potential for genomic prediction. Together, this work seeks to advance understanding of how agronomics, phenology, grain composition, and environmental variation intersect in cowpea, informing breeding strategies for productive, resilient, and nutritionally-enhanced cultivars.

## Results

### Genomic characterization of the MAGIC population and the photoperiod-insensitive subset

Founder haplotype mosaics confirmed representation of all eight founders across all 11 chromosomes (Figure 1). Linkage disequilibrium (LD) decayed to half of its maximum *r*² at 1.24 Mb in the full population and 1.30 Mb in the photoperiod-insensitive subset. Genomic PCA showed no indication of population structure, with PC1 and PC2 explaining 3.4% and 2.9% of genomic variance, respectively.

**Figure 1.**
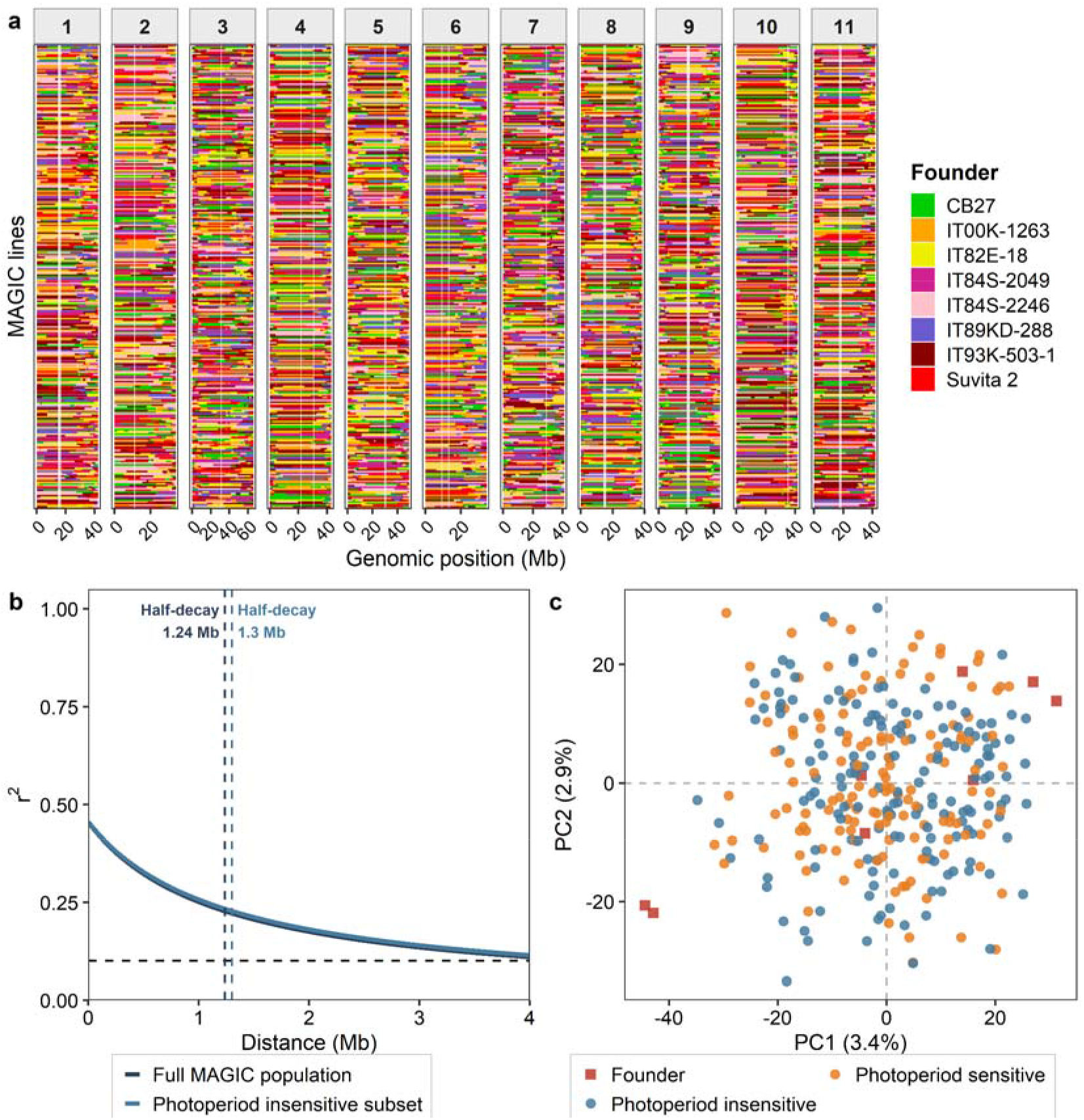
Genomic characterization of the cowpea MAGIC population and subsets. **(a)** Founder haplotype mosaic showing the inferred ancestral composition of 305 recombinant inbred lines across the eleven cowpea chromosomes. Each row represents a single MAGIC line, and each column represents a marker position along the physical map (Mb). Colors indicate the most probable founder of origin at each locus based on genotype probability estimates from qtl2. **(b)** Linkage disequilibrium (LD) decay curves for the full MAGIC population and the photoperiod-insensitive subset (n = 168; teal), estimated using the Hill-Weir model. The dashed black horizontal line indicates an r² threshold of 0.1. Vertical dashed lines indicate the half-decay distance — the physical distance at which r² declines to half its maximum value — for each population. **(c)** Principal component analysis (PCA) biplot of genome-wide SNP data for the full MAGIC population. Each round point represents a single recombinant inbred line colored by photoperiod sensitivity, while red squares represent founders. The first two principal components explain 3.4% and 2.9% of total genomic variance, respectively, indicating broad genetic diversity with no strong population stratification.

### Phenotypic data and correlation structure

The cowpea MAGIC population exhibited substantial phenotypic variation across all trait classes evaluated in the multi-environment trial as evidenced by descriptive statistics of best linear unbiased estimates (BLUEs; Table 1). Grain yield showed the greatest variation (883 ± 368 kg ha ¹ overall), with mean yield nearly five-fold higher at Parlier 2023 (2,426 ± 866 kg ha ¹) than at Thermal in both 2022 (557 ± 316 kg ha ¹) and 2023 (541 ± 330 kg ha ¹). Flowering time averaged 53.5 ± 5.8 DAP overall, ranging from 43.0 ± 5.0 DAP at Thermal 2022 to 61.5 ± 5.8 DAP at Davis 2022. Daylengths from sowing until the DAP two standard deviations above the within-environment flowering time mean (Table 1) were >14 h in Davis, >13 h in Parlier, and declined from ∼12.5 h (at sowing) to 11 h in Thermal. Plant height averaged 54.9 ± 7.6 cm across all trials with Davis averaging approximately 61.6 cm compared with 53.1 cm at Parlier and 47.9 cm at Thermal. Hundred-seed weight averaged 18.6 ± 4.4 g, with lower seed weight observed at Thermal.

**Table 1.** Descriptive statistics of spatially corrected BLUEs for agronomic, grain compositional, and rover-enabled sensing traits across six field environments. A “—” indicates a trait not measured in that environment, or that across-environment BLUEs and a grand mean were not calculated for traits measured in fewer than three environments.

| Trait (unit) | n<br>† | Grand<br>mean<br>± SD | Davis ‘22<br>mean<br>± SD | Davis ‘23<br>mean<br>± SD | Parlier ‘22<br>mean<br>± SD | Parlier ‘23<br>mean<br>± SD | Thermal‡<br>‘22 mean<br>± SD | Thermal‡<br>‘23 mean<br>± SD |
| --- | --- | --- | --- | --- | --- | --- | --- | --- |
| **Agronomic** |  |  |  |  |  |  |  |  |
| Stand count<br>(plants plot<sup>−1</sup>) | 316 | 38.7 ± 4.9 | 21.4 ± 8.0 | 32.7 ± 10.8 | 30.1 ± 7.1 | — | 53.6 ± 7.4 | 54.4 ± 4.9 |
| Flowering time<br>(days after<br>planting)§ | 315 | 53.5 ± 5.8 | 61.5 ± 5.8 | 59.6 ± 6.0 | 54.1 ± 6.3 | 51.9 ± 6.4 | 43.0 ± 5.0 | 47.0 ± 6.5 |
| Plant height (cm) | 315 | 54.9 ± 7.6 | 61.2 ± 9.9 | 62.0 ± 9.4 | 53.1 ± 9.7 | — | 45.2 ± 7.9 | 50.5 ± 8.9 |
| Grain yield<br>(kg ha<sup>−1</sup>)§ | 312 | 883 ± 368 | 505 ± 335 | 854 ± 449 | 1337 ± 602 | 2426 ± 866 | 557 ± 316 | 541 ± 330 |
| Hundred-seed<br>weight (g)§ | 305 | 18.6 ± 4.4 | 21.0 ± 4.2 | 22.3 ± 3.8 | 20.8 ± 4.6 | 21.3 ± 4.8 | 13.8 ± 4.0 | 15.5 ± 4.6 |
| **Grain compositional** |  |  |  |  |  |  |  |  |
| Moisture (% FW) | 299 | 8.75 ± 0.31 | 7.73 ± 0.32 | 7.55 ± 0.29 | 8.83 ± 0.44 | 9.33 ± 0.20 | 9.28 ± 0.69 | 10.9 ± 0.5 |
| Protein (% FW)§ | 299 | 23.7 ± 2.1 | 25.1 ± 1.9 | 25.2 ± 1.6 | 25.1 ± 1.8 | 24.0 ± 1.4 | 20.8 ± 2.7 | 20.7 ± 2.9 |
| Fat (% FW) | 299 | 1.12 ± 0.15 | 1.07 ± 0.15 | 1.15 ± 0.14 | 1.17 ± 0.15 | 1.04 ± 0.14 | 1.18 ± 0.19 | 1.15 ± 0.18 |
| Starch (% FW) | 299 | 33.1 ± 1.8 | 32.1 ± 1.6 | 31.8 ± 1.4 | 29.9 ± 1.5 | 33.7 ± 1.4 | 34.9 ± 2.7 | 37.1 ± 2.4 |
| Ash (% FW)§ | 298 | 3.64 ± 0.26 | 3.86 ± 0.26 | 3.64 ± 0.18 | 3.87 ± 0.19 | 3.60 ± 0.19 | 3.45 ± 0.32 | 3.48 ± 0.29 |
| Phytate (% FW) | 299 | 0.78 ± 0.13 | 1.22 ± 0.17 | 1.15 ± 0.14 | 0.98 ± 0.17 | 0.63 ± 0.12 | 0.38 ± 0.19 | 0.32 ± 0.14 |
| **Sensing (rover) ¶** |  |  |  |  |  |  |  |  |
| Flower count (T1)§<br>(flowers per plot) | 180 | 0.16 ± 0.07 | 0.40 ± 0.57 | 0.07 ± 0.24 | 0.03 ± 0.12 | 0.01 ± 0.08 | — | — |
| Flower count (T2)§ | 180 | 0.99 ± 1.14 | 2.24 ± 3.43 | 0.69 ± 1.69 | 0.47 ± 1.27 | 0.59 ± 1.19 | — | — |
| Flower count (T3)§ | — | — | 2.54 ± 3.89 | 3.28 ± 5.21 | — | — | — | — |
| Flower count (T4)§ | — | — | 7.37 ± 8.74 | 6.38 ± 7.87 | — | — | — | — |
| Pod count (T1)§<br>(pods per plot) | 180 | 4.07 ± 1.34 | 3.73 ± 1.87 | 19.47 ± 8.13 | 0.35 ± 0.50 | 0.14 ± 0.32 | — | — |
| Pod count (T2)§ | 180 | 6.22 ± 2.35 | 6.64 ± 4.60 | 12.96 ± 6.22 | 1.76 ± 1.61 | 6.65 ± 9.03 | — | — |
| Pod count (T3)§ | — | — | 11.04 ± 8.64 | 16.65 ± 14.28 | — | — | — | — |
| Pod count (T4)§ | — | — | 13.84 ± 14.73 | 29.99 ± 24.97 | — | — | — | — |

† Indicates the number of genotypes for which across-environment BLUEs were estimated for a given trait.
‡ The Thermal environments were unreplicated trials; estimates reflect spatial models without replicate effects.
§ Trait required Box-Cox transformation in at least one environment prior to spatial modeling; back-transformed values are shown.
¶ Rover/UAV-enabled sensing traits were collected only at Davis and Parlier; Thermal cells are empty (—).

The custom NIRS calibration developed for grain compositional traits, using partial least squares regression with standard normal variate spectral pretreatment, had a coefficient of determination (*R^2^*) of 0.93 for protein, 0.77 for fat, 0.75 for moisture, 0.63 for ash, 0.27 for phytate, and 0.20 for starch (Figure S1, Table S1 contains additional performance metrics). Among NIRS-predicted grain compositional traits, protein averaged 23.7 ± 2.1% fresh weight and was lowest in Thermal 2023 (20.8 ± 2.7%). Phytate showed the steepest environmental gradient, averaging 1.22 ± 0.17% at Davis 2022 and declining to 0.38 ± 0.19% at Thermal 2023. Moisture showed the least variation across environments (8.75 ± 0.31% overall). Rover-derived reproductive count traits showed substantial variation across environments and time points. The first two rover sensing time points (T1 and T2) were collected during the first to third weeks of flowering (from onset to early flowering) at both Parlier and Davis, whereas the later time points (T3 and T4) were collected only in Davis during weeks four to six of flowering (Figure S2). Flower counts were low at T1 and T2, with Davis 2022 showing the highest counts at both time points and increasing to 7.37 ± 8.74 flowers per plot by T4. Davis 2023 showed the highest pod counts at T1 and T2, with an increase to 29.99 ± 24.97 pods per plot by T4. Descriptive statistics of UAV-enabled traits can be found in Table S2.

The correlation structures of agronomic and grain compositional traits scored in the cowpea MAGIC MET were analyzed using pairwise phenotypic correlations and PCA (Figure 2a, Figure S3), revealing strong negative correlations between protein and starch (*r* = -0.80), starch and phytate (*r* = -0.68), and phytate and moisture (*r* = -0.84). A strong positive correlation was observed between protein and phytate (*r* = 0.68). Seed weight was moderately positively correlated with protein (*r* = 0.41) and moderately negatively correlated with starch (*r* = -0.40).

**Figure 2.**
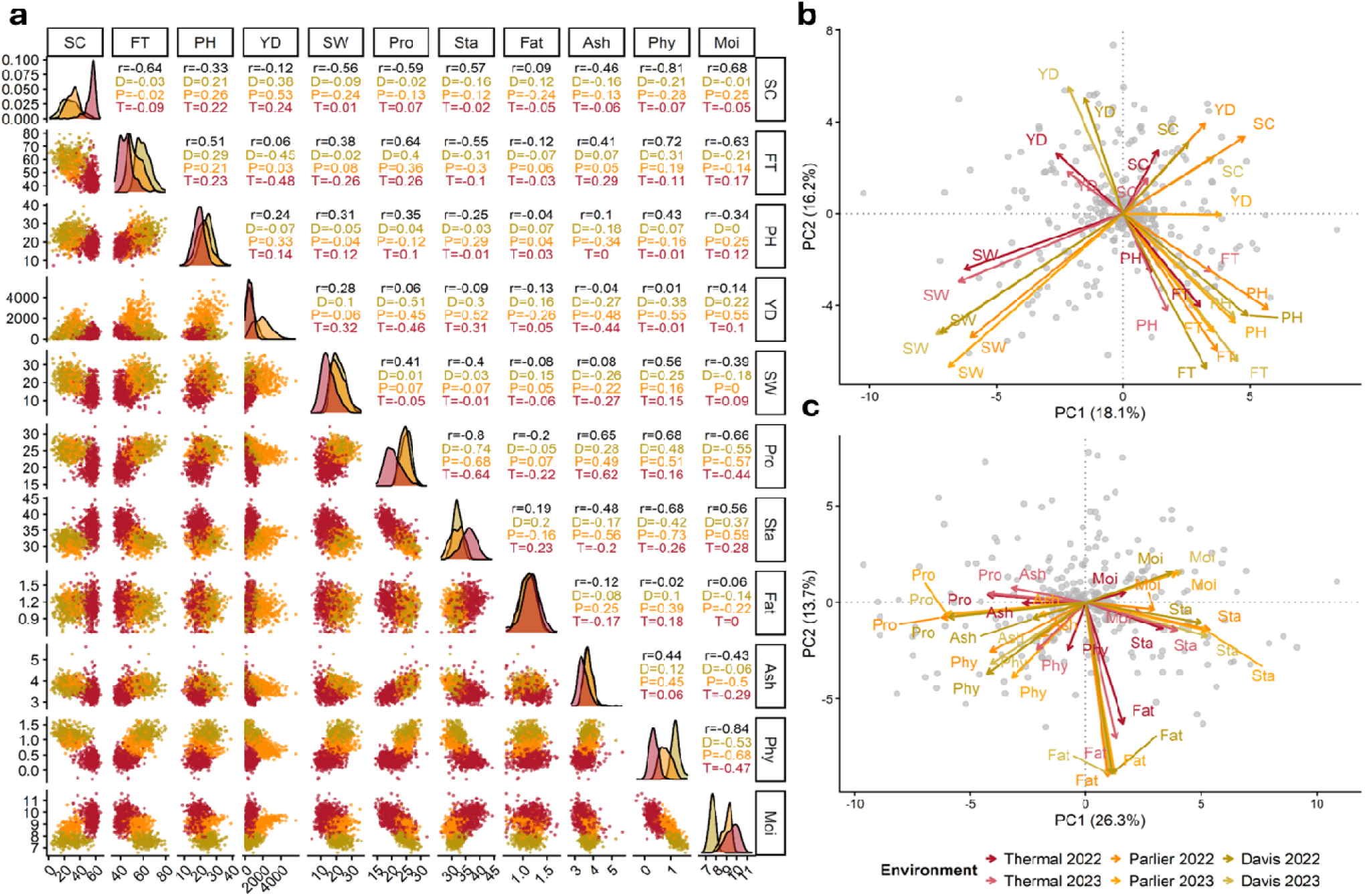
Phenotypic correlations, distributions, and principal components in the cowpea MAGIC multi-environment trial across agronomic and grain compositional traits. **(a)** Pairwise scatterplots, density distributions, and Pearson correlation coefficients for agronomic (SC, FT, PH, YD, SW) and grain compositional (Pro, Sta, Fat, Ash, Phy, Moi) traits. Overall correlations (*r*) are shown in black; location-specific correlations are shown in color (D = Davis, CA; P = Parlier, CA; T = Thermal, CA). Each point represents a genotype mean averaged across replicates within each environment. **(b, c)** Principal component analysis biplots of agronomic **(b)** and grain compositional **(c)** traits. Each grey point represents a single recombinant inbred line averaged across environments. Colored arrows indicate trait loadings with arrow color denoting environment-year combination (see legend); labels show trait abbreviations only. Abbreviations: SC, stand count; FT, flowering time; PH, plant height; YD, yield; SW, hundred-seed weight; Pro, protein; Sta, starch; Fat, fat; Ash, ash; Phy, phytate; Moi, moisture.

Agronomic trait correlations showed evidence of being environment-dependent. For example, the flowering time–yield relationship ranged from no correlation in Parlier (*r* = 0.03) to moderately negative in Davis (*r* = −0.45) and Thermal (*r* = -0.48). Flowering time also showed strong correlations with several grain compositional traits including protein, starch, and phytate, as did yield with negative correlation to protein within individual environments (Davis, *r* = -0.51; Parlier, *r* = -0.45, Thermal, *r* = -0.46). In phenotypic PCAs, the first two PCs explained 18.1% and 16.2% of the variance among agronomic traits, and 26.3% and 13.7% among grain compositional traits (Figure 2bc). In the agronomic traits biplot (Figure 2b), no traits loaded strongly onto either PC1 or PC2. Stand count and yield showed variable relationships ranging from loading together, particularly in Parlier (where an *r* of 0.53 was observed), to orthogonal relationships in Davis and Thermal. Seed weight loaded in opposition to stand count across all environments and to yield in Parlier, but no strong correlations were observed between these traits within any individual location. Environments showed clear evidence of divergence in the agronomic biplot with Thermal being the least aligned with Davis and Parlier. The grain compositional traits biplot (Figure 2c) showed protein, ash, and phytate loaded in opposition to moisture and starch along PC1, while fat loaded independently on PC2 orthogonal to all remaining traits. Environments appeared to diverge in the grain compositional biplot, with Thermal environments separating from Davis and Parlier along PC1, driven primarily by starch and protein loadings.

Phenotypic and genotypic correlation matrices showed agreement in trait relationship structure, with genotypic correlations generally amplifying patterns observed at the phenotypic level (Figure 3). The most striking trends occurred within sensing-enabled traits. The UAV-derived time-series canopy traits, plant height (PtHt) and vegetation fraction (VgFr), were strongly positively correlated both across time points (T1–T9) within each trait and between traits, particularly during the later-season time points. Vegetation fraction time points shifted substantially in their genotypic correlations between T5 and T6, with earlier time points correlating positively with stand count and later time points showing positive correlations with flowering time and strong negative correlations with flower and pod counts. Maximum growth rates and area-under-the-curve traits for PtHt and VgFr showed positive phenotypic and genotypic correlations with their respective time points. Days to canopy closure (VgFrCl_0.8_DAP) showed strong negative phenotypic and genotypic correlations with late-season vegetation fraction time points, reflecting that earlier canopy closure corresponded with greater late-season cover. Similarly, the days-to-maximum vegetation fraction growth rate (VgFr_Max_Rate_DAP) showed strong negative phenotypic and genotypic correlations with early-season vegetation fraction.

**Figure 3.**
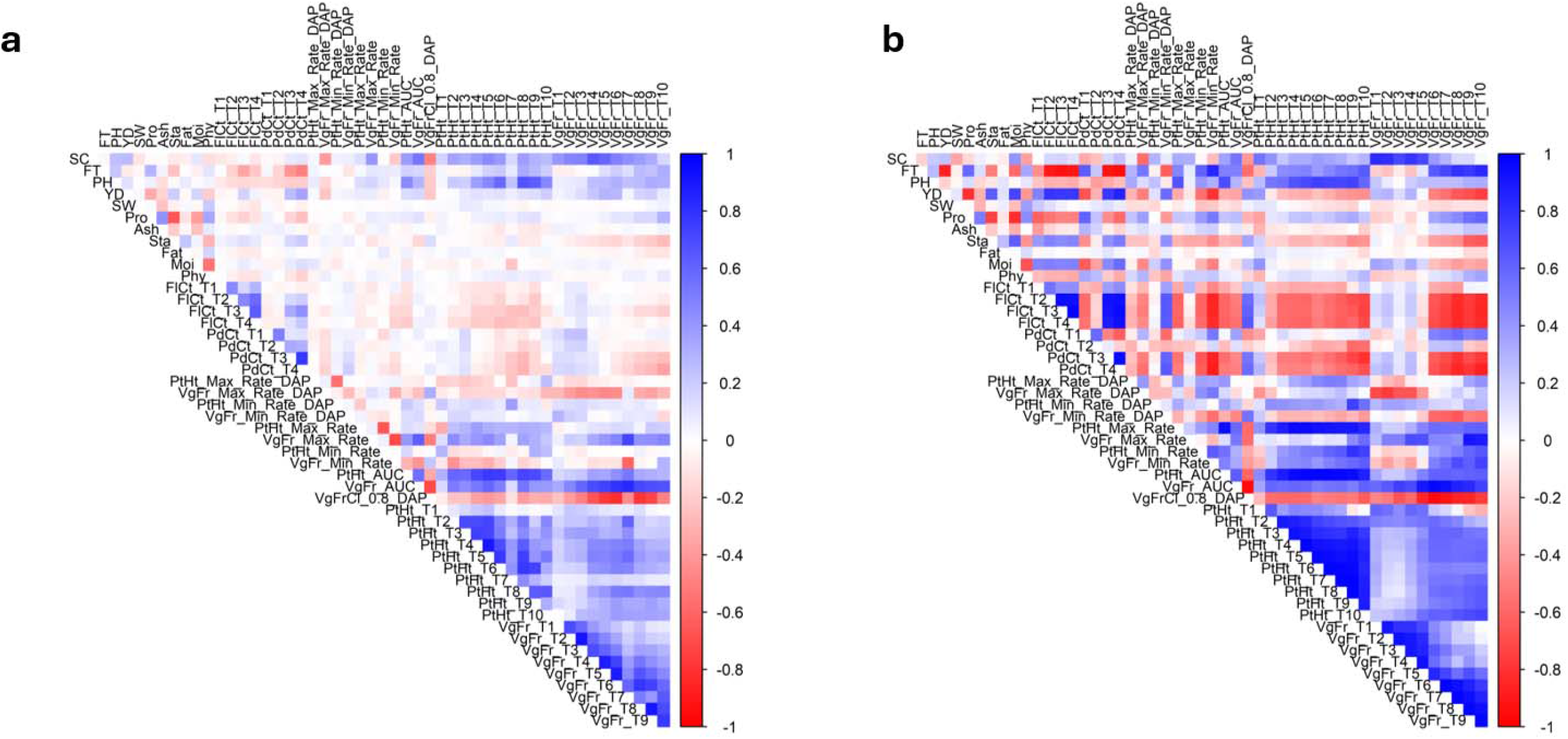
Pairwise **(a)** phenotypic and **(b)** genotypic correlation matrices for agronomic, grain compositional, and sensing-enabled traits in the cowpea MAGIC population. Color indicates correlation direction and magnitude (blue = positive, red = negative, white ≈ 0). Traits are ordered by class: agronomic (SC, FT, PH, YD, SW), grain compositional (Pro, Ash, Sta, Fat, Moi, Phy), reproductive counts (FlCt, PdCt), canopy dynamics parameters from spline modeling (PtHt and VgFr growth curve statistics), and time-series UAV-enabled traits (PtHt_T1–T9, VgFr_T1–T8). **(a)** Phenotypic correlations estimated from plot-level observations averaged across all environments with phenotypic records. Canopy dynamics and reproductive counts were not scored in Thermal; correlations with these traits were based solely on the Davis and Parlier environments. Additionally, stand count and plant height were not scored in Parlier 2023; correlations with these traits are based on the other five environments. **(b)** Genotypic correlations derived from MegaLMM posterior means of genomic-estimated breeding values from the across-environment analysis.

Rover-enabled flower and pod counts (FlCt, PdCt) were positively correlated, both genotypically and phenotypically, across time points (T1–T4) within each trait; the two traits were also correlated with each other, but only at later time points. Additionally, rover-enabled flower count showed negative genotypic correlations with UAV-enabled plant height and vegetation fraction. Manually scored flowering time showed positive genotypic correlations with UAV-enabled plant height time points that increased in strength at later-seasonal time points, and negative correlations with reproductive count traits, consistent with later-flowering lines having taller canopies but fewer flowers and pods. Manually scored flowering time also showed strong positive genotypic correlations with grain protein and phytate and negative correlations with grain starch, suggesting that phenological timing influences grain composition. Seed weight showed strong positive genotypic correlations with grain protein and strong negative genotypic correlations with grain starch, consistent with coordinated genetic controls for seed size and macronutrient composition. The grain compositional traits had broadly consistent genotypic and phenotypic correlation structure, with the protein-starch negative correlation and the protein-phytate positive correlation among the strongest signals. Protein and yield also showed a strong negative relationship, while yield and starch showed a positive relationship. Notably, agronomic and sensing-enabled traits showed relatively weak genotypic correlations with grain compositional traits, suggesting largely independent genetic control of canopy development and grain composition.

### Heritability and repeatability

Broad-sense heritability and within-environment repeatability varied substantially across agronomic, grain compositional, and rover-enabled sensing trait classes (Table 2).

**Table 2.** Broad-sense heritability (*H*²) and within-environment repeatability for agronomic, grain compositional, and sensing-enabled traits evaluated across the cowpea MAGIC multi-environment trial (2022–2023). *H*² was estimated on an entry-mean basis using the Cullis average standard error of difference (avSED) method. Repeatability was estimated analogously from single-trial models. The (—) symbol indicates that a given trait was not measured in that environment (in columns reporting repeatability), or that heritability was not calculated for traits measured in fewer than three environments. The highest repeatability and heritability were bolded within each trait class. Abbreviations: rept., within-environment repeatability; T1, time point 1; T2, time point 2; and so on for T3 and T4.

| Trait | $H^2$ | Davis '22<br>rept. | Davis '23<br>rept. | Parlier '22<br>rept. | Parlier '23<br>rept. |
| --- | --- | --- | --- | --- | --- |
| **Agronomic** |  |  |  |  |  |
| Stand count | 0.59 | 0.71 | 0.81 | 0.74 | — |
| Flowering time | 0.82 | 0.87 | **0.91** | **0.93** | 0.94 |
| Plant height | 0.81 | 0.88 | 0.88 | 0.91 | — |
| Grain yield | 0.70 | 0.78 | 0.82 | 0.84 | 0.82 |
| Hundred-seed weight | **0.89** | **0.92** | **0.91** | 0.85 | **0.99** |
| **Grain compositional** |  |  |  |  |  |
| Protein | 0.82 | **0.90** | 0.89 | 0.94 | **0.96** |
| Fat | **0.85** | **0.90** | **0.93** | **0.96** | **0.96** |
| Starch | 0.70 | 0.78 | 0.79 | 0.85 | 0.93 |
| Ash | 0.71 | 0.62 | 0.73 | 0.89 | 0.92 |
| Phytate | 0.72 | 0.77 | 0.77 | 0.83 | 0.92 |
| Moisture | 0.49 | 0.34 | 0.34 | 0.26 | 0.70 |
| **Sensing (rover)** |  |  |  |  |  |
| Flower count (T1) | 0.18 | 0.00 | 0.41 | 0.29 | 0.00 |
| Flower count (T2) | **0.74** | 0.48 | 0.76 | **0.78** | 0.34 |
| Flower count (T3) | — | 0.61 | **0.88** | — | — |
| Flower count (T4) | — | 0.59 | 0.84 | — | — |
| Pod count (T1) | 0.55 | 0.20 | 0.46 | 0.45 | 0.17 |
| Pod count (T2) | 0.23 | 0.54 | 0.37 | 0.56 | **0.86** |
| Pod count (T3) | — | 0.68 | 0.76 | — | — |
| Pod count (T4) | — | **0.72** | 0.81 | — | — |

Repeatabilities and heritabilities for the 29 UAV-enabled sensing traits can be found in Table S3. Among agronomic traits, hundred-seed weight showed the highest heritability (*H*² = 0.88), followed by flowering time and plant height (both with *H*² = 0.85) and grain yield (*H*² = 0.78). Within-environment repeatability for these traits was consistently high among environments, with flowering time repeatability exceeding 0.87 in all environments. Among grain compositional traits, fat and protein showed the highest heritability (*H*² = 0.85 and 0.82, respectively), with repeatability values reaching 0.96 at Parlier 2023 for both traits. Starch, ash, and phytate showed moderate heritability (*H*² = 0.71–0.75), while moisture was the least heritable grain compositional trait (*H*² = 0.51). Rover-enabled reproductive count traits showed variable heritability. At T1, flower and pod counts were poorly to moderately heritable (*H*² = 0.22 and 0.54, respectively), while at T2 they were substantially more heritable (*H*² = 0.85 and 0.67, respectively). Within-environment repeatability for reproductive counts was similarly variable, with near-zero values at T1 in some environments.

### Genotype-by-environment variance partitioning

Variance partitioning revealed substantial differences in the contributions of genotype, environment, and their interaction across trait classes (Figure 4). In the model across all environments (Figure 4a), fat was most underlain by genetic variance (V_G_ = 65%), followed by hundred-seed weight (V_G_ = 45%), while environmental effects dominated for most of the other traits. G×E interactions were generally modest, contributing 14% or less across most traits, with plant height (V_GE_ = 14%) and flowering time (V_GE_ = 13%) among the highest values. Flower count was dominated by residual and replicate-within-environment variance.

**Figure 4.**
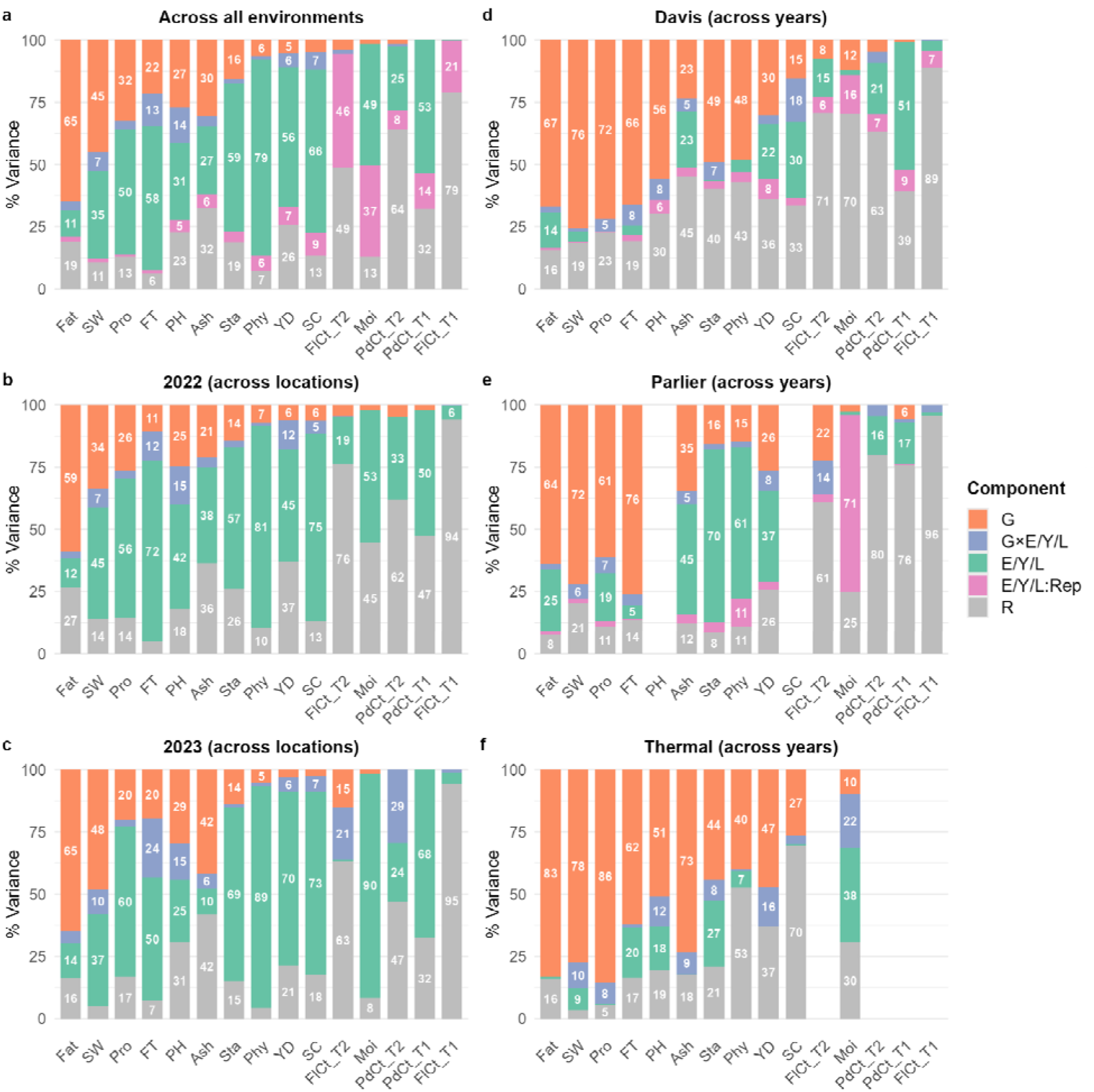
Partitioning of phenotypic variance across six analytical contexts for agronomic, grain compositional, and reproductive count traits in the cowpea MAGIC multi-environment trial. Variance components were estimated using linear mixed-effects models fit in ASReml-R. Panel **(a)** shows the variance decomposition across all environments, with components for genotype (G), environment (E), genotype × environment interaction (G×E), replicate nested within environment (E:Rep), and residual (R). Panels **(b–c)** show variance decomposition across locations within a single year, with components for G, location (L), G×L, (L:Rep), and R. Panels **(d–f)** show variance decomposition across years within a single location, with components for G, year (Y), G×Y, (Y:Rep), and R. Traits are ordered left to right by decreasing mean G% across all panels. Abbreviations: SW, seed weight; Pro, protein; FT, flowering time; PH, plant height; YD, yield; Sta, starch; FlCt, flower count; SC, stand count; PdCt, pod count; Ash, ash; Fat, fat; Moi, moisture; Phy, phytate.

Similar trends were observed in the across-year and across-location models where fat, seed weight, and (in certain analytical contexts) protein and flowering time showed the largest genetic signal, although G×E patterns differed across contexts. Variance partitioned to effects of location (L) (Figure 4b,c) was markedly higher than variance partitioned to effects of year (Y) (Figure 4d–f). Comparing the across-location, within-year models, G and G×L were higher in 2023 than in 2022 for flowering time (V_G_ = 20% vs. 11%, and V_GE_ = 24% vs. 12%, respectively). In the across-year, within-location models (Figure 4d–f), pod count at T1 showed the highest variance partitioned to effects of Y (51%) in Davis, whereas starch, phytate, ash, and yield had the largest variance partitioned to Y in Kearney (37–70%), and moisture in Thermal (38%).

### QTL mapping and GWAS

QTL mapping across all trait-environment combinations identified 267 significant QTL distributed across 10 of the 11 cowpea chromosomes, with the highest density on chromosome 9 (111 QTL) followed by chromosomes 1 (38), 8 (34), 5 (22), and 7 (22; Figure 5, Figure S4, Table S4). Genome-wide association studies identified 1,973 significant marker-trait associations across all 11 chromosomes, with chromosomes 9 (807 associations), 6 (367), and 8 (240) containing the largest number of significantly associated markers (Figure 5, Figure S5, Table S5). QTL mapping and GWAS results using within- and across-environment BLUEs are presented in Figures S3 and S4, respectively. The QTL and GWAS results were integrated using a tiered framework that prioritized agreement between GWAS and QTL methods (signals within LD at 1.24 Mb) and multi-environment support (signals detected in at least four weighted environments where individual environments had a weight of 1 and across environments signals had a weight of 2). We identified 129 Tier 1 (method agreement and multi-environment) QTL, 43 Tier 2 QTL, and 27 Tier 3 QTL, while an additional 40 loci (defined based on LD and physical overlap) only detected in GWAS met the multi-environment threshold (Tier 4). The remaining single-environment, non-concordant associations (Tier 5) totaled 68 QTL. Tier 1 QTL were most concentrated on chromosomes 9 (65 loci) and 8 (29 loci), with smaller clusters on chromosomes 1 (16 loci), 6 (9 loci), 5 (6 loci), and 7 (3 loci). The majority of significant marker-trait associations in GWAS were Tier 1 (1,163), followed by Tier 5 (514), Tier 2 (256), and Tier 4 (40; Tier 3 was not defined for GWAS-only associations). Tier 1 associations were most concentrated on chromosomes 9 (643), 6 (253), 8 (178), and 1 (79) consistent with the QTL distribution.

**Figure 5.**
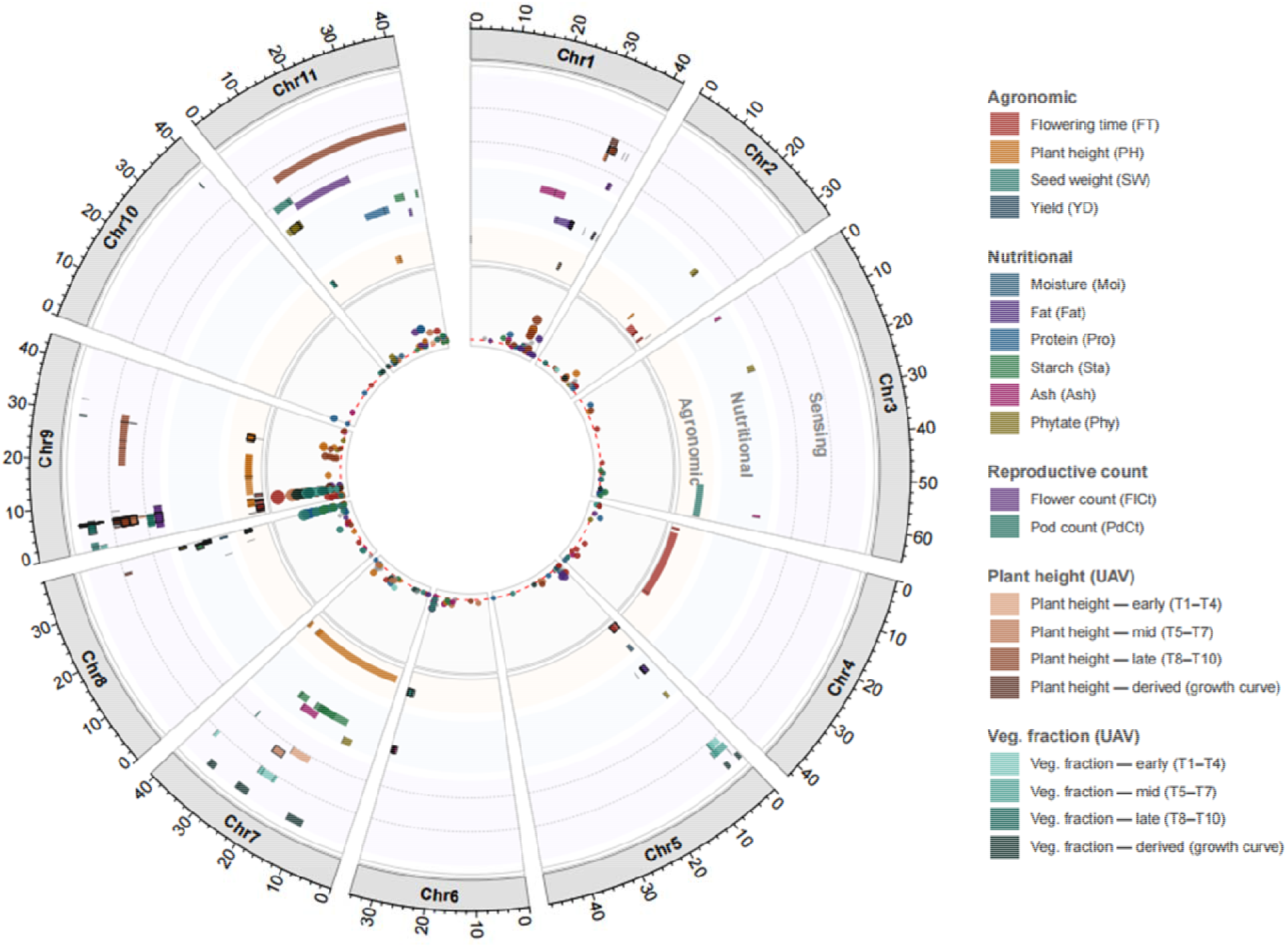
Circos plot showing the genomic distribution of quantitative trait loci (QTL) and genome-wide association study (GWAS) marker-trait associations across the cowpea genome. The outermost ring displays the physical map in megabases (Mb) based on the *V. unguiculata* v1.1 reference genome assembly. The middle ring displays QTL confidence intervals identified through composite interval mapping across multiple environments, organized into three trait groups: agronomic traits (flowering time [FT], plant height [PH], seed weight [SW], and yield [YD]), grain compositional traits (moisture [Moi], fat [Fat], protein [Pro], starch [Sta], ash [Ash], and phytate [Phy]), and sensing-enabled traits comprising reproductive counts (flower count [FlCt] and pod count [PdCt], collapsed across time points) and canopy dynamics derived from UAV-enabled remote sensing (plant height [PtHt] and vegetation fraction [VgFr], each grouped into early [T1–T4], mid [T5–T7], and late [T8–T10] sensing periods, with spline-derived growth curve parameters represented as a single derived row). Within the sensing group, dashed grey lines demarcate rover-enabled reproductive count traits, UAV-enabled plant height traits, and UAV-enabled vegetation fraction traits. QTL intervals are colored by trait (see legend), and Tier 1 loci (QTL-GWAS concordant, reproduced across multiple environments) are additionally distinguished by a thin black border. The innermost ring displays GWAS peak positions for SNPs significant in at least two of four methods (TASSEL_MLM, MVP_MLM, BLINK, FarmCPU), with point size and position proportional to -logLL(*p*). The dashed red line indicates the Bonferroni-corrected significance threshold (-logLL[0.05/32,130]). QTL, quantitative trait locus; GWAS, genome-wide association study; UAV, uncrewed aerial vehicle; SNP, single-nucleotide polymorphism.

A dense cluster on chromosome 9 near 5.8–6.0 Mb was detected as a Tier 1 QTL across six weighted environments (Nenv = 6) for flowering time (PVE = 27.5–54.5%), flower count T2 (PVE = 19.3–19.7%, Nenv = 4), UAV-derived PtHt T4–T9 (PVE = 13.0–53.4%, Nenv = 4–6), VgFr T5–T9 (PVE = 16.6–40.6%, Nenv = 4), VgFr AUC (PVE = 17.6–32.9%, Nenv = 5), and days to canopy closure (PVE = 19.1–38.3%, Nenv = 5). Genetic signal from sensing-enabled traits appeared to increase at later seasonal timepoints (Figure S6). Beyond the Tier 1 associations, this region also showed signal for flower count (T3, T4) and pod count (T1, T2, T4), predominantly as Tier 2 QTL and GWAS associations, along with isolated Tier 2/3/5 signal for manually measured plant height, UAV-enabled PtHt time points and AUC, VgFr growth rate metrics, and grain protein and starch.

Chromosome 8 contained a dense cluster of Tier 1 QTL near 37.3–37.9 Mb detected for seed weight (PVE = 10.3–35.6%, Nenv = 8), protein (PVE = 17.7–26.2%, Nenv = 8), starch (PVE = 11.7–20.3%, Nenv = 7), phytate (PVE = 9.4–24.3%, Nenv = 5), and moisture (PVE = 16.8%, Nenv = 1, Parlier 2023).

Three additional regions contained Tier 1 QTL clusters. On chromosome 1 near 33.0–35.4 Mb, signal for fat was the most stable, detected as Tier 1 (PVE = 14.2–27.0%, Nenv = 8). A co-localized cluster of late-season UAV-enabled plant height was detected at different environments per timepoint: T8 (PVE = 24.5%, Nenv = 1, Davis 2023) and T9 (PVE = 13.9–18.8%, Nenv = 3, Parlier 2023 and Across-environment). On chromosome 5, Tier 1 QTL were detected for fat near 4.3–4.4 Mb (PVE = 14.2–15.0%, Nenv = 5) and for flowering time near 0.4–1.5 Mb (PVE = 7.9–9.8%, Nenv = 4). On chromosome 6 near 32.7–34.4 Mb, Tier 1 QTL for ash (PVE = 10.7–24.2%, Nenv = 7) co-localized with Tier 1 seed weight QTL (PVE = 10.3–16.7%, Nenv = 5).

### Pleiotropy

To evaluate potential pleiotropy within each of the two identified hotspots, we used conditional scans to test whether co-localized signals were explained by the most abundant trait signal and compared founder-effect patterns at peak markers across environments. At the chromosome 8 hotspot, conditional scans using seed weight (the most abundant signal) as a covariate found that genetic signal for the remaining traits at the hotspot (starch, protein, moisture, phytate) was largely preserved across most environments (Figure S7 and S8). Strikingly, the protein conditional scans produced stronger signals than the corresponding unconditioned scans in most environments. Founder-effect profiles at this hotspot also showed consistent allelic-effect patterns among founder genotypes at peak positions for most traits and environments (Figure S9).

At the chromosome 9 hotspot, conditional scans using flowering time (the most abundant signal) as a covariate for the remaining traits found that signals were greatly reduced and generally became non-significant across most traits and environments (Figures S10 and S11). A small number of UAV-derived plant height and vegetation-fraction traits retained significance, although their signals were also substantially reduced relative to the corresponding unconditioned scans. Founder-effect profiles at this hotspot showed coordinated allelic-effect patterns among founder genotypes for flowering time and the sensing-enabled measures of flower count, pod count, plant height, and vegetation fraction across several peak positions and environments (Figure S12). Together, these results suggest that much of the chromosome 9 hotspot reflects a common genetic basis associated with flowering time, while the residual signals for some UAV-derived traits may indicate additional linked genetic effects.

### Genomic prediction

Genomic predictive ability varied by method, trait, and prediction scenario. Under baseline Within-Environment Fold-Out (WEFO), RR-BLUP and MegaLMM performed equivalently across all traits, with mean predictive ability ranging from *r* = 0.13 (stand count) to *r* = 0.46 (seed weight) and negligible differences between methods (|Δ| ≤ 0.005; Figure S13, Table S6).

For agronomic traits, Within-Location Year-Out (WLYO) predictive ability increased substantially relative to WEFO, with flowering time reaching *r* = 0.82 and plant height *r* = 0.69 under RR-BLUP (Figures 6 and 7). Comparisons of MegaLMM to RR-BLUP (Δ = *r_MegaLMM_ - r_RR-BLUP_*) in WYLO showed modest gains attributable to MegaLMM for yield (Δ = 0.06) particularly in Parlier 2023, but a modest loss for flowering time (Δ = -0.06), with otherwise negligible differences (|Δ| ≤ 0.01). Under Leave-One-Location-Out (LOLO), RR-BLUP predictive ability was modest for yield (*r* = 0.10) and stand count (*r* = 0.11), while MegaLMM recovered gains for both traits (*r* = 0.39 and 0.33, respectively). Plant height remained highly predictable under MegaLMM in LOLO (*r* = 0.65), representing gains of Δ = 0.12 over RR-BLUP, but gains of only Δ = 0.03 were observed for flowering time (Figure 6, Table S6).

**Figure 6.**
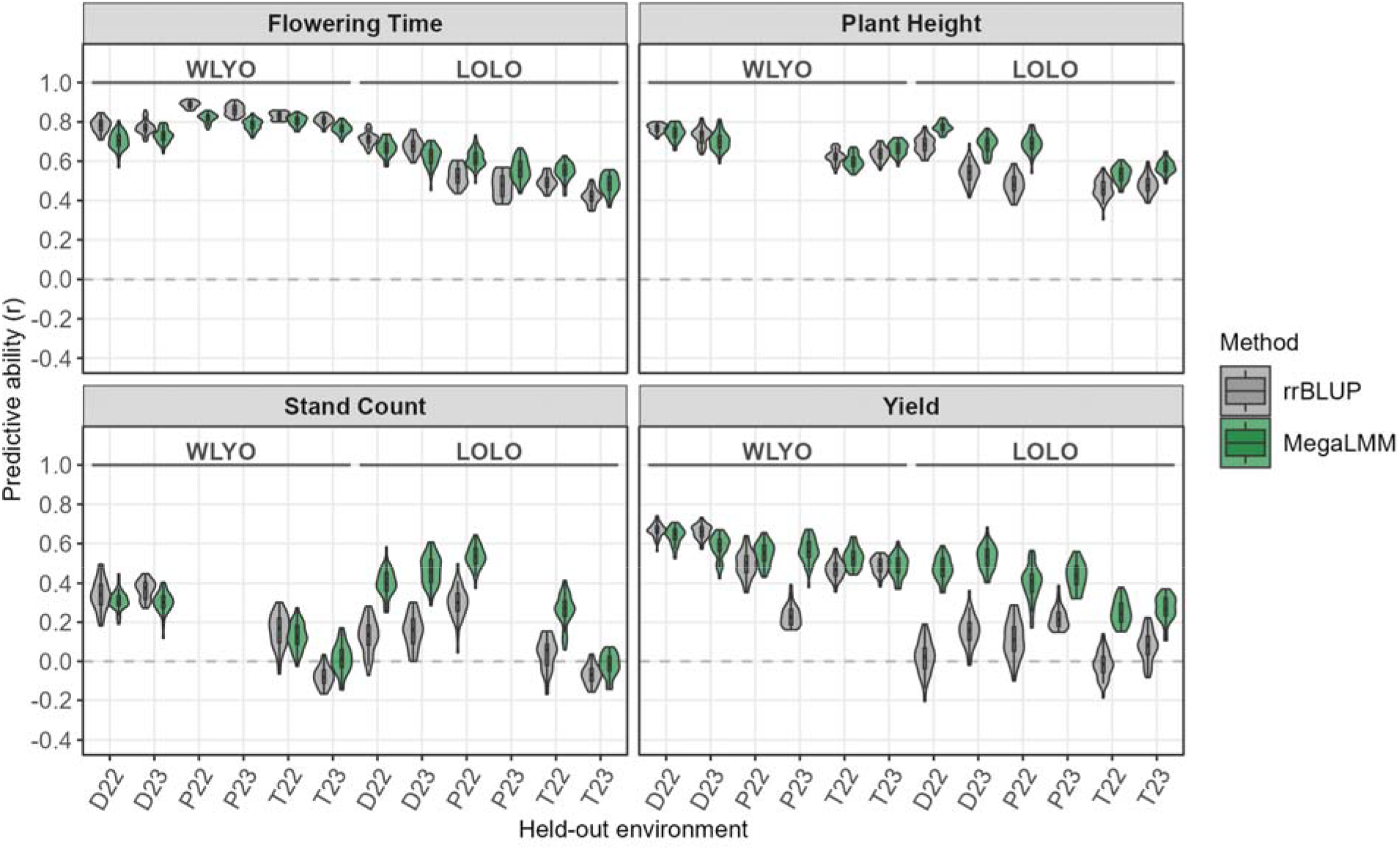
Predictive ability (Pearson correlation, *r*) of RR-BLUP and MegaLMM for agronomic traits in two across-environment genomic prediction schemes. In the WLYO (within-location year-out) scheme, models were trained on one year and evaluated in the alternate year at the same location. In the LOLO (leave-one-location-out) scheme, models were trained using both years at the remaining two locations and evaluated in the held-out location in each year. Distributions are based on 40 replicates of a 50/50 train/test partition. Stand count and plant height were not measured at Parlier in 2023 and were therefore excluded from scenarios involving that environment. D, Davis; P, Parlier; T, Thermal.

**Figure 7.**
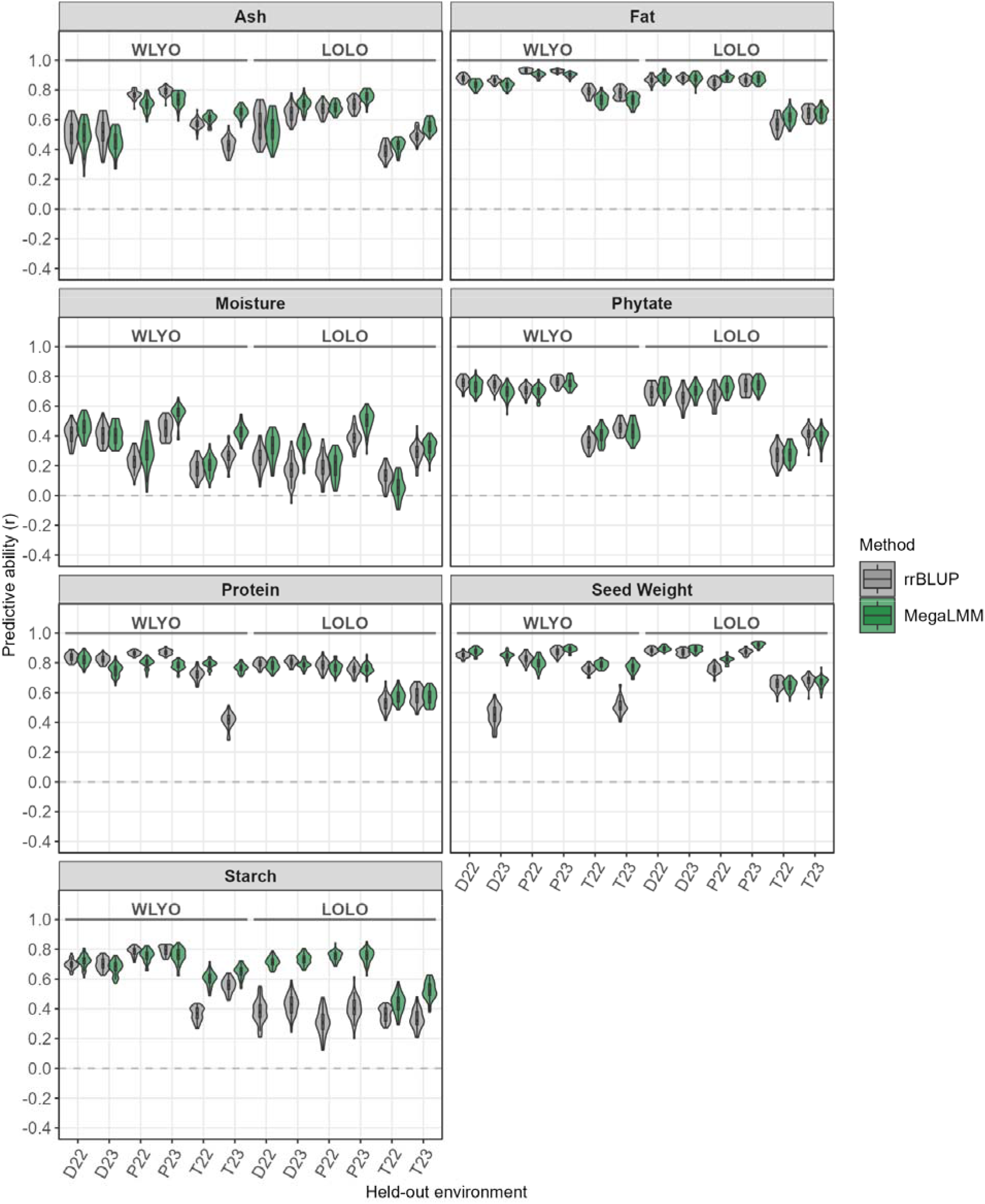
Predictive ability (Pearson correlation, *r*) of RR-BLUP and MegaLMM for seed traits in two across-environment genomic prediction schemes. In the WLYO (within-location year-out) scheme, models were trained for one year and evaluated in the alternate year at the same location. In the LOLO (leave-one-location-out) scheme, models were trained using both years at the remaining two locations and evaluated in the held-out location in each year. Distributions are based on 40 replicates of a 50/50 train/test partition. D, Davis; P, Parlier; T, Thermal.

For seed traits, WLYO predictive ability was high across both methods except for moisture, with seed weight (*r* = 0.83), fat (*r* = 0.82), and protein (*r* = 0.79) showing the strongest performance under MegaLMM. MegaLMM provided modest WLYO gains for seed weight (Δ = 0.12), starch (*r* = 0.69; Δ = 0.04), and moisture (*r* = 0.39; Δ = 0.07) and a modest loss for fat (Δ = -0.04). Under LOLO, RR-BLUP predictive ability declined for starch (*r* = 0.37) and moisture (*r* = 0.24), while MegaLMM maintained high predictive ability for starch (*r* = 0.65, Δ = 0.28). Seed weight, protein, and fat remained highly predictable under both methods in LOLO (MegaLMM *r* = 0.81, 0.71, and 0.80; Figure 7, Table S6).

### Candidate genes

To focus candidate gene discovery on the most robust genetic signals, candidate gene analysis was restricted to Tier 1 loci, which represented QTL-GWAS concordant regions with multi-environment support. This filtering retained 35 Tier 1 trait-by-chromosome loci distributed across seven chromosomes. The largest number of Tier 1 loci occurred on chromosome 9 (17 loci), followed by chromosome 8 (6), chromosome 1 (4), chromosome 7 (3), chromosomes 5 and 6 (2 each), and chromosome 2 (1). Concordance between QTL and GWAS peak positions was generally high: GWAS peaks fell within the corresponding QTL confidence interval for 28 of 35 loci, and QTL and GWAS peaks were separated by ≤100 kb for 16 loci and ≤500 kb for 25 loci.

Across these Tier 1 regions, 1,677 candidate genes were identified within ±250 kb of a QTL or GWAS peak, with the largest numbers on chromosomes 1 (373 genes), 9 (343), 5 (294), 8 (221), 6 (208), and 7 (182), and a smaller set on chromosome 2 (56, Table S7). Most candidate genes had support for cross-species annotation transfer: 1,621 genes (96.7%) had a protein BLAST hit to a common bean (*Phaseolus vulgaris*) ortholog, with a mean percent identity of 88.3%.

Expression in seed (based on Yao et al., 2016) was used to further prioritize the candidate set for seed weight and compositional traits, with 1,198 genes (71.4%) in Tier 1 regions for these traits expressed in at least one seed developmental stage. With the LD observed in this study, the specific candidate genes described in the text below were identified as putative prioritized candidates and are not being claimed as candidate causal genes.

Candidate gene associations were concentrated in the major multi-trait hotspots identified by QTL mapping and GWAS. On chromosome 8, the seed size and grain composition hotspot contained 221 candidate genes, of which 166 (75.1%) were expressed in seed. Most genes in this region were associated with multiple traits, including seed weight (198 genes), starch (189), phytate (158), protein (155), and moisture (88). Notably, 146 genes on chromosome 8 were in the search space for four or more traits, and 88 for all five of these traits. Candidate genes in this region were annotated as encoding a glucose-1-phosphate adenylyltransferase family protein (*Vigun08g214900*) and a late embryogenesis abundant (LEA) hydroxyproline-rich glycoprotein family (*Vigun08g213200*).

The chromosome 9 phenology and canopy development hotspot contained 343 candidate genes. Flowering time was the most commonly associated trait within this region (with 210 genes in the respective search spaces), followed by flower count at T2 (155), plant height (117), plant height maximum growth rate (112), and late-season UAV-derived plant height traits including PtHt_T9 (112), PtHt_T8 (109), PtHt_T7 (108), PtHt_T6 (108), and PtHt_T5 (107). This locus also contained candidate genes associated with vegetation fraction traits, including VgFr_T9 (62), canopy closure DAP (59), VgFr_T8 (38), and VgFr_AUC (29). A total of 110 genes were associated with four or more traits; of those, 59 were associated with ten or more traits. Candidate genes in this region were annotated as encoding two TCP family transcription factors (*Vigun09g062200* and *Vigun09g073600*) and a MADS-box transcription factor 6 (*Vigun09g059700*).

Additional candidate gene clusters were identified on chromosomes 1, 2, 5, 6, and 7. Chromosome 1 contained 373 candidate genes, primarily associated with fat (310 genes), plant height (254), and late-season UAV-enabled plant height traits, with *TERMINAL FLOWER (TFL) 1-LIKE* (*Vigun01g173000*) identified as a candidate. Chromosome 2 contained 56 candidate genes, all of which were associated with flowering time, with genes annotated as encoding a *WUSCHEL*-related protein (*Vigun02g123400*) and MADS-box transcription factor family protein (*Vigun02g125600*) identified as candidates. Chromosome 5 contained 294 candidate genes split between flowering time (224) and fat (70), with four consecutive homologs annotated as encoding *FT* (*Vigun05g004000*, *100*, *200*, and *300*) identified as candidates. Chromosome 6 contained 208 genes associated with seed weight and ash, including 141 genes shared by both traits; genes annotated as encoding a SWEET transporter (*Vigun06g228900*), magnesium transporter (*Vigun06g236100*), and zinc finger protein (*Vigun06g236600*) were identified as candidates. Chromosome 7 contained 182 genes associated with fat, flowering time, and PtHt_T6, but no candidates with characterized relevance were identified.

## Discussion

### Environmental drivers of phenotypic variation

The cowpea MAGIC MET spanned a gradient of environmental conditions and varied widely in edaphic and climatic factors (Figure S14 shows temperature and solar radiation trends). The Thermal location stood out as most distinctive, showing notably lower seed protein and phytate, as well as reduced seed weight and plant height (Table 1, Figure 2a). Thermal is characterized by desert sandy soils that generally have poorer nutrient-holding capacity and are less fertile than the clay loam soils found at Davis and Parlier. This lower soil fertility may have limited plant-available N and P, key nutrients influencing seed protein and phytate content, potentially explaining the reductions observed in these traits. Taken together, this broader nutrient limitation may have also contributed to smaller seed size and reduced plant height but is confounded with planting date among other factors such as high temperatures during vegetative stages.

Conversely, the Parlier and Davis locations were closer latitudinally and shared more similar soils and climatic conditions, although Parlier featured higher minimum and maximum temperatures (Figure S10). Grain yield was highest in Parlier particularly in 2023, which showed a three- to five-fold increase over other environments (Table 1). The 2023 season was also more favorable in Davis, which showed about a two-fold increase over 2022. Yields at Thermal were stable across years and closely aligned with Davis 2022. The trend for increased yield in Davis and Parlier could represent a regional trend for more favorable growing conditions in the 2023 season. Additionally, examination of flowering time across this gradient showed a trend towards early flowering in Thermal, which likely reflected the shorter photoperiod and warm pre-flowering temperatures, and in Parlier, which featured overall warmer temperatures than in Davis. Dareus et al. (2021) identified positive phenotypic correlations for plant height with each of flowering time and destructively measured plant biomass (at flowering and at pod maturity) in the UCR mini-core across two years of field trials in Florida. Our results for manual and sensed plant height, manual flowering time, and sensed vegetative fraction at later time points were consistent with this directionality, and were indicative of genotypic correlations being more pronounced than phenotypic correlations, in this MAGIC MET.

Flowering time QTL shifted in genomic position between the short-day (Thermal) and long-day (Davis, Parlier) locations, recovering loci previously reported as genetic controls of cowpea flowering time (Lo et al., 2018; Huynh et al., 2018; Muñoz-Amatriaín et al., 2021). This contrast could be partly structural in the cowpea MET, as the long-day trials (>14 h in Davis and >13 h in Parlier) evaluated the photoperiod-insensitive subset, while the short-day trials (∼12 h to 10.5 h) enabled by fall planting in Thermal included the full population. The pattern is nonetheless consistent with distinct genetic controls (and potentially also differential epistasis as evidenced in *P. vulgaris*; González et al., 2021) being engaged under contrasting photothermal regimes, potentially reflecting components of the photothermal perception and signaling machinery whose effects are resolvable only under the daylength and temperature conditions that activate them, as also hypothesized for high-temperature tolerance loci identified under long days in cowpea (Hall et al., 2002; Hall, 2012).

The pronounced effects of environment on seed composition, yield, and phenology traits highlight an important consideration for cowpea breeding: the definition of the target population of environments (TPE) for cultivar development. The distinctive edaphic and climatic conditions at Thermal suggest that it may represent a distinctly different production environment than California’s Central Valley, where the Davis and Parlier field sites and most commercial cowpea farms are located in the state. However, Thermal’s hot, arid conditions enabled a fall planting window and thereby a discriminating environment that may better approximate short-day production environments in sub-Saharan Africa than the more temperate conditions of California’s Central Valley (Huynh et al., 2018).

### Genetic architecture of grain compositional and phenological traits

Large QTL intervals and broad LD (1.24 and 1.30 Mb in the full population and photoperiod-insensitive subset, respectively) presented a fundamental challenge for candidate gene identification, as a substantial number of genes initially satisfied proximity criteria, necessitating iterative refinement through the integration of QTL-GWAS concordance, expression data, and functional keyword searches. Despite these refinements, many candidate genes in search spaces encoded proteins of unknown function, and causal genes could not be definitively called. Fine mapping and functional validation will ultimately be required. This limitation is shared broadly across GWAS and QTL studies conducted in populations with large LD blocks and highlights the value of continued investment in cowpea functional genomics resources.

The co-localization of genetic signals for seed weight and composition on chromosome 8 suggests that a genetic factor in this region could control their coupling, with larger seeds tending towards less starch but higher protein. Breeding programs selecting for smaller seeds may thus benefit from also monitoring grain protein content to ensure its maintenance or tandem improvement, in spite of a negative genetic correlation potentially mediated by this hotspot.

Previous studies have also identified signal in this chromosome 8 region for seed traits. Akinmade et al. (2025) identified a seed length association (at ∼37.63 Mb). Lo et al. (2019) identified a seed density association (at ∼36.98 Mb) and a seed width meta-QTL (at ∼37.71 Mb), and proposed *Vigun08g217000*, which encodes a histidine kinase 2. Lo et al. (2018) found the chromosome 8 region identified in this study to affect not only seed weight but also pod length, leaf length, and leaf width, suggesting broader pleiotropic control of organ size at this locus indicative of gigantism. Wu et al. (2024) identified this chromosome 8 region as a candidate selective sweep in a study of both vegetable and grain cowpea, and found colocalized associations with pod crude protein, seed starch, and strikingly pod shattering, raising the possibility that this locus has been an indirect target of domestication or breeding selection.

Taken together, the convergence of signals in the present study and in the literature at this region across multiple populations underscores the distinct possibility of a pleiotropic genetic control or a cluster of linked causal genes that regulates several fundamental processes related to the breadth of associated traits.

One of two candidate genes in this chromosome 8 region was annotated as encoding a glucose-1-phosphate adenylyltransferase family protein (also known as AGPase), specifically a small subunit thereof based on BLASTp results (Camacho et al., 2009). A non-synonymous mutant allele of a *Lotus japonicus* homolog exhibited reduced starch content in embryos alongside leaves and roots (Vriet et al., 2010). This candidate gene was reported to be expressed in multiple tissues in cowpea (Yao et al., 2016), as was the other candidate gene in this region, annotated as encoding an LEA protein. LEA proteins have a role in desiccation tolerance in seeds and protection of proteins and other constituents from dehydration (Gilles et al., 2007; Dam et al., 2009; Chatelain et al., 2012), alongside other roles in stress tolerance in legumes (Battaglia & Covarrubias, 2013). Homologs in *P. vulgaris* were found to be upregulated in earlier stages of seed development, suggesting activation during reserve accumulation rather than solely during final maturation (Lopes et al., 2025).

A separate Tier 1 region identified for grain ash and seed weight on chromosome 6 near 32.7–34.4 Mb co-localized with mineral (including Ca, Mg, Mn, N, and K) and seed weight QTL reported in this population by Huynh et al. (2024) across two years of field trials. Given that ash represents total mineral content, this convergence across independent populations and mineral-specific assays lends support to the relevance of the region for grain mineral accumulation.

Within the chromosome 9 hotspot with QTL for flowering time, vegetation fraction, plant height, and minor signals for grain protein and starch, candidate genes included two TCP family TFs (*Vigun09g062200* was annotated as TCP18-like, and *Vigun09g073600* could be TCP14-like based on BLASTp) and one MADS-box TF 6. A TCP14 homolog in *Lupinus luteus* was a candidate for flowering onset under short days and the end of flowering under long days (Dogra et al., 2026). In cowpea, the same MADS-box TF 6 identified herein was also a candidate for flowering time under long days in the UCR minicore (Muñoz-Amatriaín et al., 2021), with co-localized signal under long days in the presently studied MAGIC population (Huynh et al., 2018) and under short days in a cultivated-by-wild cross (Lo et al., 2018). BLASTp supported homology to *AGAMOUS-LIKE (AGL) 8*, which was a candidate for photoperiod sensitivity in *Phaseolus coccineus* (Rendón-Anaya et al., 2025), flowering time in *P. vulgaris* (González et al., 2021), and reproductive period length in *Glycine max* (L. Wang et al., 2024). *AGL8* is relevant for inflorescence identity and floral development and negatively regulated by *APETALA1 (AP1)* (Mandel & Yanofsky, 1995). BLASTp also supported homology with truncated *CAULIFLOWER A (CAL)*, an earlier-stage regulator of flowering initiation (Mandel & Yanofsky, 1995). *CAL* has high sequence identity with *AP1* (and both are also MADS-box family TFs) but interacts with only some of the same TFs due to a difference in the C-terminal domain (Saini & Yadav, 2020) and has differentiated expression attributed to evolutionary gains/losses of TF binding sites (Ye et al., 2016). Multi-omic investigations of flowering-related gene regulatory networks and TF complexes (Serrano-Mislata et al., 2017) in cowpea, with integration of evolutionary genetics and mathematical modeling (Duk et al., 2024), would be informative.

Candidates on other chromosomes included, for flowering time, another MADS-box TF with homology to *AGL* and *DEFICIENS* (also known as *AP3-LIKE*), a *WUSCHEL*-related protein (relevant for floral initiation in *G. max* and flowering in *Medicago sativa*; Wong et al., 2011; Jha et al., 2020; Wang et al., 2025), and four homologs of *FT* (which binds *AP1* and was also a candidate in the UCR minicore; Paudel et al., 2021). A candidate for manual and sensed plant height was *TFL1-LIKE*, which also interacts with flowering-related TFs (Serrano-Mislata et al., 2017; Goretti et al., 2020; Zhu et al., 2020) and is a major controller of determinacy in cowpea among other legumes (Benlloch et al., 2015; Dhanasekar & Reddy, 2015; Su et al., 2026). This finding suggests that plant height may have been an indirect measure of determinacy or that *TFL* could have otherwise contributed to vertical growth. Notably, *FT* and *TFL* are differentiated by subtle molecular changes (Dhanasekar & Reddy, 2015) that merit detailed examination (for *AGL* and *CAL* as well) in a broader survey of allelic variation in cowpea, and interactions of TCP family TFs with FT/TFL1-like proteins have also been noted (Colleoni et al., 2024). Finally, three candidates were identified for seed traits: a SWEET transporter (N3-like based on BLASTp) involved in carbohydrate metabolism for which respective homologs were upregulated in ovules after fertilization in *Cicer arietinum* (Singh et al., 2023) and proximal to a seed protein QTL in *Pisum sativum* (Burstin et al., 2007); a magnesium transporter, of which a family was recently characterized in soy, with magnesium being an important macronutrient for seed development (Akter et al., 2025); and a zinc finger protein, of which homologs were upregulated in pod and seed in *L. japonicus* and *M. trunculata* (Bhattacharjee et al., 2015). Given the mapping resolution in this population, the detailed pursuit (e.g., in a functional genetics context) of lesser characterized candidate genes or TFs would not be recommended until multi-omic examinations have been conducted in the relevant tissues in cowpea and/or until genetic signal in a given region has been more finely mapped.

### Phenotyping methodologies

The subset of RIL and founder genotypes with indeterminate or sprawling growth habits produced extensive lateral vegetative growth into alleys and furrows, which precluded rover-enabled sensing at later reproductive stages due to the risk of damage to plants and yield loss. This limited our ability to fully evaluate genetic mapping and GP of flowering and pod formation dynamics, and meant that our results could reflect phenology (differences in flowering time) and did not capture cumulative reproductive outputs. Future studies could address this limitation through the selection of lines with upright type I or II growth habits or a narrower flowering-time distribution, modified planting arrangements to reduce lateral spread (e.g., super-blocking), or the use of alternative sensing platforms such as low-altitude UAV flights that avoid ground-level contact with the canopy, to enable a more complete time series.

Predicted traits, whether NIRS- or image-based, contain sources of error absent from direct quantifications that may propagate into downstream analyses. The NIRS calibration performed poorly for starch and phytate (*R^2^*of 0.20 and 0.27, respectively), though these traits had repeatabilities of 0.77–0.93, heritabilities of 0.70 and 0.72 (respectively), and genetic signal observed in mapping. Further work on the chromosome 8 hotspot, detected as pleiotropic for those and other seed traits, would benefit from detailed wet chemistry analyses for macronutrient content and composition. As pertains to image-based phenotyping, pod detection was less accurate than flower detection (Kamangir et al., 2026), likely because of greater structural variability, occlusion, and visual similarity to stems, petioles, and peduncles. UAV-derived canopy traits were also influenced by image processing (stitching, vegetation segmentation, ground-surface estimation) and the temporal density of flights; derived features such as maximum growth rate are also sensitive to spline modeling choices. Additionally, at Parlier, narrow and elongated plot dimensions caused pairs of adjacent plots to be captured within single images during rover operation, requiring stitching at the two-plot level and additional processing steps to obtain plot-level values, which may have reduced the precision of rover-enabled sensing trait estimates at that location. Certain types of error are not necessarily random: Zhou et al. (2021) demonstrated that measurement error in image-based phenotyping can be partly heritable and produce different associations than those obtained with ground-truth values. Careful alignment of field trial design with the operational constraints of sensing platforms and projected canopy development is recommended for future studies.

### Multi-trait genomic prediction advantage

Our comparison of MegaLMM, a Bayesian multi-trait factor-analytic model, against single-trait RR-BLUP found a consistent, albeit variable, advantage of MegaLMM. In the WEFO scenario, which compared prediction of unseen genotypes in a seen environment, RR-BLUP and MegaLMM performed equivalently across all traits (|Δ*r*| ≤ 0.005). This similarity in performance was expected as the ability of the multi-trait model to leverage trait correlations, perhaps its key advantage, was removed by only inputting a single trait to both models. When additional traits were input to the models in the cross-environment prediction scenarios (LOLO and WLYO), the two GP methods diverged particularly in the LOLO scenario. In that scenario, which requires prediction into an unobserved location, single-trait RR-BLUP collapsed for yield, stand count, starch, and moisture while MegaLMM recovered substantial predictive ability for each of these traits (Δ*r* = 0.178–0.467). This improvement indicates that borrowing information from correlated traits measured in the training environments partially compensated for the absence of any observation in the target location. Testing which correlated traits are most beneficial to multi-trait genomic predictive ability for priority traits in cowpea breeding pipelines would be an important next step.

## Conclusion

In this study, we evaluated the cowpea MAGIC population in six field environments for agronomic, phenological, canopy development, and grain compositional traits; dissected their genetic basis via GWAS and QTL mapping, including conditional scans as a screen for pleiotropy; and assessed predictive ability via single- and multi-trait GP. Hotspots were identified for seed traits (on chromosome 8) and for phenology and canopy development (on chromosome 9), though conditioning on flowering time reduced other significant trait signals within the latter; these and other loci identified herein await further examination in functional mechanistic work. Finally, multi-trait GP conferred across-environment predictive advantages and is recommended for use in cowpea METs. The findings and methodologies from this study can be used towards the comprehensive improvement of cowpea productivity, adaptation, and quality.

## Materials and Methods

### Plant materials and field trials

The germplasm under study herein was an eight-founder cowpea MAGIC population developed by Huynh et al. (2018). This cowpea MAGIC population was planted in a multi-environment field trial conducted in California over two years (2022 and 2023). The full population of 305 recombinant inbred lines and eight founder genotypes was evaluated at the Coachella Valley Agricultural Research Station (CVARS) in Thermal, CA (33.52° N, 116.15° W) using an unreplicated randomized complete block design (RCBD) during the fall-winter season under short-day conditions. A subset of 168 photoperiod-insensitive lines and the eight founders (when available) were evaluated in thrice-replicated RCBDs at the University of California Davis Field Station in Davis, CA (38.54° N, 121.78° W) and at the Kearney Agricultural Research and Extension Center in Parlier, CA (36.60° N, 119.50° W) during the spring-summer season under long-day conditions. The three field trial locations differed primarily in temperature and photoperiod and were selected as relevant environment types for Californian cowpea production or, in the case of Thermal, for population evaluation under short-day conditions. According to the Köppen-Geiger climate classification, Davis is categorized as hot-summer Mediterranean (Csa), Parlier as cold winter semi-arid (BSk), and Thermal as hot desert (BWh; Peel et al., 2007). Davis had Reiff very fine sandy loam soil, Parlier had Hanford sandy loam, and Thermal had Carsitas gravelly sand. At Davis, planting occurred on May 24, 2022, and May 23, 2023. At Parlier, planting occurred on June 24, 2022, and June 21, 2023. At Thermal, planting occurred on September 8, 2022, and September 12, 2023. Davis used plots measuring 1.52 m × 3.05 m in both years, Parlier used plots measuring 0.76 m × 4.57 m in 2022 and 0.76 m × 6.1 m in 2023, and Thermal used plots measuring 0.76 m × 4.57 m in both years. All three trials were conventionally managed and irrigated, via subsurface drip in Davis and Thermal and furrow in Parlier. Daylengths were recorded from the Astronomical Applications Department of the U.S. Naval Observatory (https://aa.usno.navy.mil/data/Dur_OneYear) with specification of each year in which a trial was conducted, using the ‘Need USA Location?’ functionality to specify Davis, CA, and Parlier, CA, and inputting the Location coordinates of 33.5201, -116.1515 in the case of Thermal.

### Phenotyping

An intensive phenotyping strategy was employed to characterize variation across agronomic, sensing-enabled, and grain compositional traits. Traits were collected using a combination of manual field measurements, rover and UAV sensing platforms, and benchtop NIRS, enabling extensive coverage of plant performance from early vegetative development through grain maturity. Raw phenotypic data underwent environment-specific spatial correction as described below prior to use in genetic analyses. All computational analyses in the phenotyping and remaining sections were performed in R version 4.3.0 unless otherwise noted.

#### Agronomic traits

Grain yield, plant height, stand count, and flowering time were evaluated as key agronomic traits. Grain yield, i.e., the weight of clean seed, was standardized to kg per ha to enable comparisons across different plot sizes. In 2022, plant height was measured as the vertical distance from the soil surface to the highest point of the canopy (excluding pods). In 2023, plant height was measured using two measuring poles connected by a taut string. Two measurements, the height at each measuring pole after adjusting the string to best approximate the height of the canopy, were collected per plot and averaged; measurements were taken during late pod fill stages to minimize phenological effects. Flowering time, defined as the number of days after planting (DAP) when 50% of plants in a plot had at least one open flower, was scored weekly except during peak-flowering periods, when it was scored twice per week. Hundred-seed weight was collected using a digital scale and 3D-printed 100-well plates.

#### Sensing-enabled traits

RGB imagery was collected from rover and UAV platforms in Davis and Parlier during the 2022 and 2023 field seasons to support automated phenotyping. Rover image acquisition was performed using a robotic sensing platform (Mineral, Mountain View, CA, USA) equipped with a downward-facing Basler acA2500-20gc RGB camera mounted 1.6 m above the ground (Basler AG, Ahrensburg, Germany). Rover operations were conducted at weekly intervals at both locations from vegetative through mid-reproductive stages, and twice during the week of peak flowering in Davis (Figure S2). Rover images acquired from flowering through early pod fill were used to estimate visible flower and pod counts (Kamangir et al., 2026). Images were stitched into orthomosaics using AgRowStitch (Uyehara et al., 2026), cropped to 640 × 640 pixels, and annotated by plant-breeding experts using bounding boxes in Roboflow (Roboflow, Inc., 2020). Across the four environments, the annotated dataset contained 992 images with 6,926 flower instances and 702 images with 4,998 pod instances (Kamangir et al., 2026). The number of represented genotypes varied among environments because reproductive expression and image availability differed across genotypes, locations, and years. Flower and pod object-detection models were developed using YOLOv11x initialized from COCO-pretrained weights (Common Objects in Context; Lin et al., 2014) as described in Kamangir et al. (2026). Briefly, a consistent model architecture and training configuration were used, with separate models fitted for each environment. Flower detection achieved 76.3% mean average precision at 50% intersection over union (mAP@50), 80.8% precision, and 71.5% recall, and pod detection achieved 65.1% mAP@50, 76.2% precision, and 59.2% recall, when evaluated on represented genotypes and environments (Kamangir et al., 2026). Model performance was relatively stable when applied to unseen genotypes within represented environments (but declined more substantially across locations and years, in other scenarios evaluated in Kamangir et al., 2026). These traits represent estimates of visible reproductive organs at specific sensing dates rather than direct measurements of cumulative flower or pod production.

UAV imagery was collected using a Phantom 4 Pro V2.0 with an RGB camera (DJI, Shenzhen, China), flown at 3 m s^−1^ at an altitude of 10 m, with 80% overlap and 70% sidelap. UAV images were acquired from early vegetative through seed fill stages at a weekly time step (Figure S2) and used for assessing canopy dynamics. UAV imagery was stitched using Metashape (Agisoft LLC) with eight ground control points per field. The resulting orthomosaics were filtered to remove sensing dates with image stitching failures and excessively weedy periods which occurred briefly prior to mechanical cultivation or hand-weeding.

Two traits were extracted from UAV imagery for canopy dynamics modeling: 1) vegetation fraction (VgFr), the proportion of the plot occupied by thresholded green pixels over total pixel count, and 2) predicted plant height (PtHt), the 95th percentile of a canopy height model (CHM). RGB orthomosaics and digital elevation models (DEMs) were generated using Metashape v2.1.3. The RGB orthomosaics were cropped using plot boundaries defined in a GeoJSON file, and the Excess Green (ExG) vegetation index was calculated in Python 3.8.11 using the GDAL (version 3.6.2), GeoPandas (version 0.12.2), and OpenCV (version 4.11.0) packages. The resulting vegetation index images were classified using a binary mask with Otsu thresholding (Otsu, 1975). After the binary mask images were processed, vegetation fraction (VgFr) values were calculated for each plot boundary by dividing the number of vegetation pixels by the total number of pixels within the boundary. Plant height (PtHt) was calculated by taking the 95th percentile of a CHM formed by subtracting the baseline ground elevation from the DEM values for vegetation pixels (sampled using the ExG mask).

PtHt and VgFr were each modeled as temporal growth trajectories, with splines fit independently for each plot using the gam() function from the ‘mgcv’ R package (Wood, 2011), with smoothing parameters estimated by restricted maximum likelihood (REML). For each plot, the response variable (PtHt or VgFr) was modeled as a smooth function of the calendar date using a one-dimensional spline with basis dimension k = min(6, n − 1), where n is the number of unique flight dates for that plot. A value of k = 6 was used for the majority of plots, which had eight to 10 repeated measurements.

Predicted values were generated on a dense grid of 500 equally spaced points spanning the observed date range to support numerical estimation of derivatives, extrema, threshold crossings, and area under the curve; this interpolation did not increase the temporal resolution of the underlying weekly observations. Instantaneous growth rates were approximated using finite differences of the predicted values. From these smoothed trajectories, we extracted extrema, minimum and maximum values and minimum and maximum rates of change (growth rates), as well as the DAP at which these extrema occurred. Additionally, we extracted the final area under the curve (AUC) using trapezoidal numerical integration of the spline-predicted values across the observed date window. Finally we estimated days to canopy closure as the first DAP where VgFr > 0.8 as predicted using the spline models.

#### Near-infrared spectroscopy data acquisition and calibration development

Grain macronutrients were predicted using NIRS and a reference wet chemistry training/validation dataset. Cowpea grain samples were first ground into a fine powder using an IKA tube mill programmed to oscillate up to 10,000 RPM for 1.3 min and stainless steel grinding chambers (IKA, Breisgau, Germany). Ground samples were subsequently scanned using a FOSS DS2500 (FOSS, Hillerød, Denmark) with a pre-existing manufacturer calibration for Vegetal Protein Meals. Scans were conducted over the visible through near-infrared range (400–2500 nm) at a spectral resolution of 0.5 nm. Pseudoabsorbance spectra log_10_(R^−1^) were exported from FossManager™ as .nir files, which were converted into .csv format using Spectragryph software (Menges, 2024).

Reference wet chemistry data collection was stratified across environments to capture variation attributable to both genotype and growing conditions. A total of 40 samples per environment was selected via Kennard-Stone algorithm using the kenStone() function from the ‘prospectr’ package (Stevens & Ramirez-Lopez, 2026) based on Mahalanobis distances, resulting in 240 wet chemistry samples across the six study environments. Two additional environments (Riverside 2023 and Davis 2024) were incorporated to expand environmental representation, yielding a final total training/validation set of 320 samples.

Reference wet chemistry analyses on these 320 samples were also conducted on ground seed, namely a subset of the aliquot scanned via NIRS. Protein content was determined using the Dumas combustion method (*AOAC 990.03*). Moisture content was measured via standard oven drying (*AOAC 925.10*). Crude fat was quantified using the Randall modification of Soxhlet extraction (*AOAC 2003.05*), ash content by gravimetric loss at 600 °C (*AOAC 942.05*2), and starch concentration using a Megazyme total starch assay kit (Latimer, 2023). Phytate concentration was measured using a Megazyme phytate assay kit (McKie & McCleary, 2016). Aside from starch and phytate quantifications, all wet chemistry analyses were conducted at the UC Davis Analytical Laboratory.

Spectral preprocessing (spectral transformation) was performed using the ‘prospectr’ package (Stevens & Ramirez-Lopez, 2026). Twelve spectral pretreatments were evaluated, including untreated spectra, baseline correction, detrending, standard normal variate (SNV), multiplicative scatter correction, Savitzky-Golay (SG) smoothing, and first- and second-order SG derivatives. Additionally, combinations of pretreatments were also evaluated. Combinations included SNV (applied first) and SG methods and detrending transformations (Figure S15).

Outlier detection was performed for both spectral pretreatments and wet chemistry data (Figures S16 and S17, respectively). Wet chemistry outliers were detected individually for each trait as points that exceeded 2.5 times the interquartile range below the 1^st^ and above the 3^rd^ quartiles.

This resulted in the removal of three data points, two for ash and one for starch, representing <1% of the wet chemistry data. Spectral outlier detection was performed independently for each spectral pretreatment to identify local outliers, which were then pooled to identify global outliers for removal. Local spectral outliers were detected as points that exceeded the 99.5^th^ percentile of robust Mahalanobis distances. Robust Mahalanobis distances were calculated using the Mahalanobis() function from the ‘stats’ package (R Core Team, 2025) and the covMcd() function from the ‘robustbase’ package (Maechler et al., 2026) on the first five PCs of the spectral data after each pretreatment was applied. The center and covariance matrix from the minimum covariance determinant were input into the Mahalanobis call to avoid excessive leveraging of extreme values. Global spectral outliers were defined as data points detected as local outliers in ≥75% of pretreatments. Six global spectral outliers were identified and removed from the analysis, representing <1% of spectra.

Subsequently, partial least squares regression (PLSR) was implemented using the ‘pls’ R package (Liland et al., 2026). Separate models were fitted for each combination of macronutrient compositional trait and pretreatment, and performance was assessed via five-fold cross-validation using the normalized root mean square error of prediction (NRMSEP, the average RMSE of held-out folds divided by the trait’s standard deviation). The optimal number of latent components was initially determined using the one-sigma criterion, with later visual inspection of root mean square error of prediction (RMSEP) in validation plots used to assess model adequacy. When the one-sigma criterion selected overspecified or overly parsimonious models, the number of components was adjusted by manually selecting an optimum that minimized both the number of components and RMSEP. Pretreatments were evaluated using the RMSEP, with model parsimony also considered. The SNV transformation was selected as the optimal pretreatment and applied globally using the optimal number of components identified in five-fold cross validation to obtain final predicted macronutrient compositional traits. These predicted values were used in subsequent spatial modeling and quantitative genetic analyses.

### Spatial modeling

All plot-level phenotypic data were analyzed using linear mixed-effects models implemented in ASReml-R 4.2 (VSNi, Hemel Hempstead, United Kingdom; Butler et al., 2023). Phenotypic data were analyzed using a two-stage framework consisting of single-trial analyses (STA) followed by multi-environment trial (MET) analyses. The STA modeled within-environment spatial variation, whereas the MET modeled genetic covariance and genotype-by-environment (G×E) interactions. Prior to model fitting, residuals versus fitted values plots were examined for heteroskedasticity. Traits exhibiting heteroskedastic residuals were Box-Cox transformed by evaluating λ values from -2 to 2 in 0.5-unit increments and selecting the transformation that maximized the log-likelihood. Outlier detection was conducted in the MET using the ASReml alternative outlier method (AOM).

For the STA, phenotypic data were analyzed in each environment using the following linear mixed-effects model:

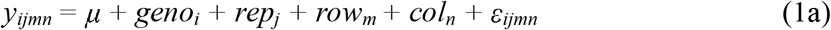

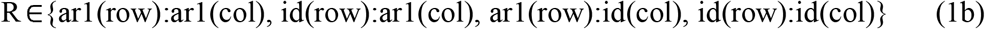

where *y_ijmn_* is the observed phenotype of genotype *i* in replicate *j* at spatial position *mn*; μ is the overall mean; *geno_i_* is the genotype effect; *rep_j_* is the replicate effect; and *row_m_* and *col_n_* represent larger-scale spatial effects associated with field rows and columns, respectively. The residual term ε*_ijmn_* captures unexplained variation including local spatial correlation.

Spatial correlation in residuals was modeled using alternative variance-covariance structures to account for field trends along rows and columns. The residual variance-covariance matrix R was specified as one of the structures shown in equation 1b where ‘ar1’ denotes a first-order autoregressive process and ‘id’ denotes an identity (i.e., independent and identically distributed) structure. For each trait-by-environment combination, the optimal residual structure was selected via likelihood ratio testing (LRT), beginning with the most parameterized residual model and sequentially comparing reduced models using LRT.

Because stand establishment can directly impact yield, stand count was included as a fixed covariate in STA yield models. Additionally for the Thermal location, which consisted of a single replication, model selection frequently resulted in boundary, convergence, and overspecification issues. To ensure model stability and maintain a consistent spatial correction, we simplified the Thermal STA models by removing the random effects of row, column, and replicate.

To estimate within-environment BLUEs, replicate, row, and column were treated as random effects (except at Thermal, where these effects were removed), and genotype effects were treated as fixed. These within-environment BLUEs were used for downstream quantitative genetic analyses described below.

The MET analysis was conducted using a second-order factor analytic (FA) mixed model framework:

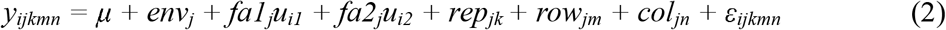

where *y_ijkmn_* is the observed phenotype of genotype *i* in environment *j*, replicate *k*, and spatial position *mn*; μ is the overall mean; and *env_j_* is the fixed effect of environment *j*. The terms *fa1_j_* and *fa2_j_*are environment-specific loadings for the first and second latent factors, respectively, while *u_i1_* and *u_i2_* are the corresponding genotype effects (also known as scores) for the genotype responses to those factors. The terms rep_jk_, row_jm_, and col_jn_ are the environment-specific random effects of replicate, row, and column, respectively. The LRT-determined environment-specific residual structures from STA (Equation 1b) were incorporated into the MET model to account for within-environment spatial variation. Together, these terms modeled the genetic covariance structure among environments, thereby capturing the major patterns of G×E interaction.

An FA2 variance-covariance structure was used because it provided an appropriate balance between model flexibility and parsimony for the number of environments evaluated. There were six cases of model convergence failure in the FA2 modeling, which required defaulting to first-order FA models (Table S8). Replicate effects and spatial row and column effects were modeled in the MET as environment-specific random effects for all environments except for Thermal, which excluded these terms. Residual diagnostics for all final MET models are shown in Figure S18. Outliers were detected in the MET models using the AOM with a threshold of four standard deviations. Phenotypes with AOM Studentized conditional residuals exceeding four standard deviations were removed from the analysis before refitting the final model to estimate BLUEs.

Across-environment BLUEs were then derived using a two-step procedure to avoid bias associated with reusing predicted genetic effects (Holland & Piepho, 2024). In the first MET model, genotype was specified as a random effect to estimate genetic and residual variance components under the FA structure. The model was then refit with genotype specified as a fixed effect using the ASReml update() function, while retaining the previously estimated variance-covariance and residual structures, to obtain across-environment BLUEs.

#### Heritability and repeatability

Broad-sense heritability across environments was quantified using the average standard error of the difference (avSED) as described by Cullis et al. (2006). This approach is appropriate for unbalanced multi-environment mixed models because it accounts for heterogeneous prediction error among genotypes. For each trait, an across-environment random effects model was fit in ASReml:

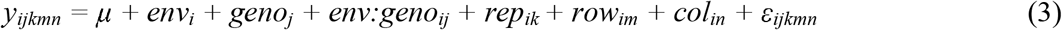

where *y_ijkmn_*is the observed phenotype of genotype *j* in environment *i*, replicate *k*, and spatial position defined by row *m* and column *n*, μ is the overall intercept (grand mean), *env_i_* is the random effect of environment *i*, *geno_j_* is the random effect of genotype *j* with *g_j_*∼*N(0,*σ*_g_^2^)*, *geno:env_ij_* is the random genotype-by-environment interaction for environment *i* and genotype *j*, *rep_kl_* is the random effect of replicate *k* nested within environment *i*, *row_im_*is the random effect of field row *m* within environment *i*, *col_in_*is the random effect of field column *n* within environment *i*, and ε*_ijklmn_* is the residual error term, modeled using the environment-specific spatial variance-covariance structure selected in the STA.

The genetic variance component σ*_g_^2^* and the avSED among genotype predictions were extracted from this model. Using these values, Cullis heritability was calculated as follows:

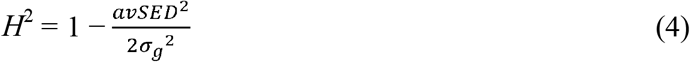

where the numerator represents the average prediction error variance of pairwise genotype differences (i.e., the square of the avSED). Heritability was estimated separately for each trait using all available environments. In cases where there were fewer than three environments with phenotypic records the calculation was skipped. Repeatability within environments was estimated analogously from a corresponding STA model for each trait × environment combination, using the same expression.

### Genetic and phenotypic correlation analyses

We estimated genetic and phenotypic correlations using principal component analysis (PCA), correlation matrices, and the ggpairs() function from the GGally R package (Schloerke et al., 2025). Correlation matrices were estimated using trait values averaged per genotype and mean-centered. Pairwise phenotypic correlations were computed across all environments using the cor() function in R. The averaged and mean-centered traits were also used to summarize the phenotypic space in PCA using the prcomp() function with scale=TRUE. Biplots were constructed to visualize relationships among traits, environments, and genotypes.

Genotypic correlations among traits were estimated using the MegaLMM R package (Runcie et al., 2021). After fitting a MegaLMM model, the additive genetic posterior mean matrix (termed U_train) was used in the estimate_gcor() function to estimate genetic correlations.

### QTL mapping

QTL mapping analyses were conducted using pedigree and genomic data from Huynh et al. (2018). The genomic dataset consisted of 32,130 single nucleotide polymorphisms (SNPs) genotyped on the iSelect Cowpea Consortium SNP array (Muñoz-Amatriaín et al., 2017). Prior to analysis, individuals lacking pedigree, genotypic, or phenotypic records were excluded, and SNPs were filtered to retain only those with a minor allele frequency ≥ 0.05. All QTL mapping was performed in R using the qtl2 package (Broman et al., 2019).

Genotype probabilities were calculated for observed markers and for pseudo-markers inserted at 0.1-centiMorgan intervals across the genetic map. We conservatively assumed a genotyping error rate of 1%, which is common for SNP arrays (J. Wang, 2018). Kinship matrices were estimated from the genotype probability grid using the leave-one-chromosome-out (LOCO) method to control for genome-wide relatedness while preserving ability to detect QTLs within each chromosome. Three models were initially compared including Haley-Knott regression (H-K), a linear mixed model with genome-wide kinship (LMM), and LOCO (Figure S19). All models performed well, and the LOCO model was used as the primary basis for QTL inference. Whole genome scans were performed iteratively for within- and across-environment BLUEs for each trait using the scan1() function and LOCO kinship matrix.

Permutation-based significance thresholds were determined for each trait, for within- and across-environment BLUEs, via 1,000 permutations of phenotypes using the scan1perm() function.

QTL peaks were identified using the resulting trait-specific LOD threshold (α = 0.05) with a minimum peak drop of 2 LOD units between adjacent peaks. Confidence intervals (CIs) for each peak were estimated using two complementary approaches: a 1.5-LOD drop interval and a 95% Bayes credible interval. The narrower of the two intervals was retained as the final CI for each QTL (Figure S20). The proportion of phenotypic variance explained (PVE) by each QTL was estimated using the residual sum of squares (RSS) and total sum of squares (TSS) of a single-QTL model fit at the peak position using fit1() with the chromosome-specific LOCO kinship matrix.

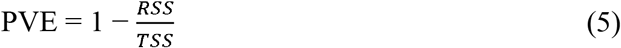

### Genome-wide association studies

Genome-wide association studies (GWAS) were conducted using four complementary methods to maximize robustness of inference: Mixed Linear Models (MLM) implemented in both rMVP (Yin et al., 2021) and rTASSEL (Monier et al., 2022), FarmCPU (Liu et al., 2016) implemented in rMVP, and BLINK (Huang et al., 2019). All models included the first two principal components (PCs) as fixed covariates to account for population structure, and MLM models additionally incorporated a kinship matrix estimated from marker data. Significance was assessed using a Bonferroni-corrected threshold (with a desired α of 0.05, divided by 32,130 markers).

### Pleiotropy

To investigate potential mechanisms underlying QTL co-localization at trait hotspots, conditional genome scans and founder allele effect comparisons were performed. For each hotspot, the trait exhibiting the highest LOD score at that locus was included as an additive covariate in genome-wide scans (using LOCO kinship) of all other traits for which a significant QTL was identified in the region, using the scan1() function with the addcovar argument. To assess whether signal remained after conditioning on the highest-LOD score trait, and/or whether signal was displaced to an alternative locus elsewhere in the genome, whole-genome and single-chromosome scan profiles (restricted to the chromosome containing the hotspot) were compared between the unconditioned and conditional models for each trait. Founder allele effects at each hotspot were estimated using the scan1coef() function with the chromosome-specific LOCO kinship matrix, and effects at the peak marker were compared across all traits with a significant QTL at the hotspot to evaluate the degree of similarity in founder allele effect patterns among co-localized traits.

### Candidate gene search

To determine the physical distance from peak signals to use as a candidate gene search space, we examined linkage disequilibrium (LD) in the cowpea MAGIC population. The genome-wide intrachromosomal LD half-decay distance was calculated by inputting 32,130 markers to PLINK (Purcell et al., 2007) to calculate *r*^2^ values. Resulting *r*^2^ values for all pairwise comparisons were input to a Hill-Weir model to determine LD half-decay distance (the genomic distance at which LD decays to half of its maximum *r*^2^; Hill & Weir, 1988). Ultimately, LD decayed over too long of a distance (1.24 Mb) to be useful for defining candidate gene search spaces.

QTL intervals and GWAS marker-trait associations were integrated to assess concordance between mapping approaches. To define GWAS loci, all significant SNPs (exceeding the Bonferroni threshold in any of the four methods) were expanded by ± 1.24 Mb to approximate LD decay, and any physically overlapping expanded intervals within the same trait, chromosome, and environment were merged into consensus GWAS loci. QTL-GWAS concordance was determined by testing whether any merged GWAS locus physically overlapped (by any amount) with QTL confidence intervals for the same trait, chromosome, and environment. Additionally, environmental reproducibility was assessed independently for both QTL and GWAS loci: a locus was considered multi-environment if the weighted sum of environments with an overlapping interval (≥ 25% overlap of the smaller region) reached a threshold of 4, with individual environments weighted as 1 and the across-environment (AE) BLUE weighted as 2. Following the evaluation of method concordance and multi-environmental support, all loci were assigned to a five-tier prioritization hierarchy: Tier 1 (QTL and GWAS concordant, with multi-environment support in either mapping approach), Tier 2 (QTL and GWAS concordant, without multi-environment support), Tier 3 (QTL only, with multi-environment support), Tier 4 (GWAS only, with multi-environment support) and Tier 5 (all remaining significant loci). We performed candidate gene searches only for Tier 1 to prioritize the most consistent results.

To conduct the candidate search, we first discarded QTL intervals that exceeded 5 Mb as poor-quality associations likely reflecting low statistical resolution rather than true locus boundaries. Genes within 250 kb of QTL and/or GWAS peaks were retained as gene candidates. Next, we expanded ± 250 kb from the peak signal at Tier 1 loci and intersected these regions with the *Vigna unguiculata* v1.1 reference genome (Lonardi et al., 2019) using the rtracklayer and GenomicRanges packages in R (Lawrence et al., 2013). Candidate genes were annotated using the *V. unguiculata* v1.1 annotation file (Phytozome 14, Goodstein et al., 2012). Ortholog relationships between cowpea and common bean were inferred using OrthoFinder v3.1.5 (Emms et al., 2026) with primary transcript protein sequences from *V. unguiculata* v1.1 and *P. vulgaris* v2.1 (Phytozome v14). Sequence similarity between cowpea candidate genes and their inferred common bean orthologs was confirmed by protein BLASTp (BLAST+ v2.16.0; Camacho et al., 2009; e-value ≤ 1×10); 97% of candidate genes had a BLASTp hit with a mean percent identity of at least 88%, supporting reliable annotation transfer. Seed expression profiles were obtained from the *V. unguiculata* Gene Expression Atlas (VuGEA; Yao et al., 2016) accessed through the Legume Information System (Dash et al., 2016), comprising RNA-seq data across seven tissue types including four seed developmental stages (8, 10, 14, and 18 days after pollination). Transcripts per million (TPM) values were averaged across biological replicates within each developmental stage, and genes with mean TPM > 1 in at least one seed developmental stage were classified as being expressed in seed.

Final candidates were selected from the Tier 1 gene set using functional annotations, tissue-specific expression (seed-expressed genes for seed traits), QTL-GWAS peak proximity, and BLASTp comparison against characterized homologs in the literature. Genes lacking direct evidence of putative relevance to the associated trait(s) were filtered out. All final gene candidates were examined in detailed BLASTp analysis for comparison with existing literature on proteins of related structure, upon which further filtering was conducted if there was not direct evidence of putative relevance to the trait(s) with significant signal in the relevant region.

### Genomic prediction

Genomic prediction was performed for agronomic and seed traits to assess predictive ability in the cowpea multi-environment trial and to determine whether multi-trait GP improved performance relative to a single-trait baseline. The baseline model was ridge regression–best linear unbiased prediction implemented with the rrBLUP R package (Endelman, 2011), while the multi-trait model was MegaLMM, a Bayesian factor-analytic mixed model that jointly models correlated traits (Runcie et al., 2021).

Models were evaluated using a common cross-validation framework consisting of 20 replicates of two-fold cross-validation, with identical fold assignments between methods to enable direct comparisons. Three prediction scenarios were evaluated: Within-Environment Fold-Out (WEFO), in which models were trained and evaluated independently within each environment using reciprocal two-fold (50/50) genotype partitions; Within-Location Year-Out (WLYO), in which models were trained using one year at a location and evaluated on the alternate year at that location; and Leave-One-Location-Out (LOLO), in which models were trained using data from two locations and evaluated on the held-out location. For LOLO, training environments comprised all environments aside from the left-out location (e.g., train on D22, D23, P22, P23, and evaluate against T22 and T23). For WLYO, training was restricted to the alternate year from the same location (e.g., train on D22 and predict D23).

For WLYO and LOLO, phenotype masking followed Scenario 1 of Laporte et al. (2025). Within the target environment, all non-focal traits were fully masked while the focal trait was split into two folds that were reciprocally masked. Both models were trained on the unmasked half of the focal trait in the target environment, but differed in the information available from the non-target environments: the multi-trait model was trained on all traits in non-target environments, whereas the single-trait model, which cannot use information from correlated traits, was trained on the focal trait alone. In the multi-trait model, this design prevents leakage of predictive power from phenotypic correlations within the target environment by excluding them from model training, while still leveraging trait correlations measured in the non-target environment (Bernardo, 2010; Runcie & Cheng, 2019).

Predictive ability was quantified as the Pearson correlation between observed BLUEs and predicted additive genetic values. For RR-BLUP, predictions were genomic estimated breeding values (GEBVs), whereas for MegaLMM they were posterior mean genetic values (U_Train). Predictive ability was summarized as the mean correlation across 40 fold estimates (20 replicates × 2 folds) for each trait-scenario combination. Analyses included 11 traits measured across six environments (D22, D23, P22, P23, T22, and T23), yielding 66 possible trait-by-environment prediction targets per prediction scenario. Stand count and plant height were excluded from the WLYO and LOLO scenarios involving Parlier 2023 because these traits were not measured in that environment.

### Genotype-by-environment analyses

To partition phenotypic variance into genotypic (G), environmental (E), genotype-by-environment interaction (G×E), replicate, and residual (R) components, a linear mixed-effects model was fit for each trait using ASReml-R 4.2 (Butler et al., 2017). The model included environment, genotype, and genotype-by-environment interaction as random effects with a homogeneous residual variance structure:

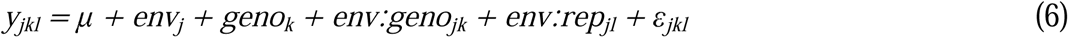

where *y* is the plot-level phenotypic observation, μ is the overall mean, *env_j_* is the random effect of environment *j*, *geno_k_* is the random effect of genotype *k*, *env:geno_jk_*is the random genotype-by-environment interaction for genotype *k* in environment *j*, *env:rep_jl_* is the random replicate-within-environment effect for environment *j* and replicate *l*, and ε*_jkl_* is the residual error modeled using the environment-specific spatial variance-covariance structure selected in the STA (Equation 1b).

This model was applied in six analytical contexts to decompose variance at different scales of the trial structure: (1) across all six environments; (2) across locations within 2022; (3) across locations within 2023; (4) across years within Davis; (5) across years within Parlier; and (6) across years within Thermal. In contexts (2) and (3), the grouping factor E represents location; in contexts (4)–(6), it represents year. Traits with phenotypic records in fewer than three levels of the grouping factor were excluded from that analytical context. Variance components were extracted from each fitted model and expressed as a percentage of total phenotypic variance to facilitate interpretation of the relative contributions of each component.

### Data Statement

Data and scripts are provided as supplemental materials for the review process. Data will be published as supplemental materials, with BLUEs and master phenotypic records provided as Data S1 and Data S2, respectively. Scripts will be published in a public GitHub repository with DOI generated via Zenodo at time of manuscript acceptance.

### Disclosure of AI Use

AI was used to estimate visible flower and pod counts as described in the ‘Sensing-enabled traits’ subsection of Materials and Methods. Partial least squares regression was used to develop a custom NIRS calibration as described in the ‘Near-infrared spectroscopy data acquisition and calibration development’ subsection of Materials and Methods. Additionally, browser-based Claude (Anthropic) Opus 4.6, 4.7, 4.8 and Sonnet 4.6 generative AI models were used during January 2026 to August 2026 as a coding assistant to help write, debug, and review R scripts for some portions of the analytical pipeline. This included pleiotropy testing, GWAS, genomic prediction, GxE variance decomposition, and figure generation. All code suggested with AI assistance was run line by line and carefully reviewed, tested, and in all cases debugged and modified via human intervention. Outputs were verified against the underlying data by the authors before use, and all analyses are reproducible from the code provided as supplemental materials for the review process that will also be made available in a public GitHub repository at time of manuscript acceptance, as noted in our Data Statement above. This study did not use any human-subject, personal, or proprietary information, and no data was shared directly with generative AI models, so no additional privacy or compliance measures were taken. The same models were at times consulted for rewording sections of manuscript text (namely, suggested rephrasing of one or a couple of sentences most commonly, and rarely up to a paragraph) but were not relied on for drafting, interpretation of results, or scientific conclusions, which remained the responsibility of the authors.

## Supporting information

Large supplemental tables and figures, uploaded separately

## Acknowledgments

The authors gratefully acknowledge Alison Kelly and Clayton Forknall for their tutorials in AsREML-R and spatial modeling. We also gratefully acknowledge the teams at each of the three field stations who managed these field trials. We also acknowledge the High Performance Computing Core Facility (https://hpc.ucdavis.edu) at the University of California, Davis, for providing computational resources that have contributed to the research results reported in this paper.

## Supporting Information

**Figure S1.** Custom near-infrared spectroscopy calibration for predicting cowpea grain composition.

**Figure S2.** A timeline illustrating rover and drone sensing dates of the cowpea MAGIC MET within developmental periods based on flowering time data collected in each environment.

**Figure S3.** Pairwise phenotypic correlations among agronomic and grain composition traits across environments in the cowpea MAGIC population.

**Figure S4.** Within- and across-environment QTL profiles in the cowpea MET.

**Figure S5.** Manhattan plots and quantile-quantile (Q–Q) plots from GWAS in the cowpea MET.

**Figure S6.** QTL profiles for sensing-enabled traits in the Davis and Parlier environments (2022–2023).

**Figure S7.** Within- and across-environment QTL profiles for chromosome 8, conditional on seed weight vs. unconditioned, in the cowpea MET, for seed traits that had genetic signal in the chromosome 8 hotspot.

**Figure S8.** Within- and across-environment QTL profiles for all 11 chromosomes, conditional on seed weight vs. unconditioned, in the cowpea MET, for seed traits that had genetic signal in the chromosome 8 hotspot.

**Figure S9.** Heat map of founder allele effects at the chromosome 8 hotspot for seed traits (moisture, Moi; phytate, Phy; protein, Pro; starch, Sta; hundred-seed weight, SW).

**Figure S10.** Within- and across-environment QTL profiles for chromosome 9, conditional on flowering time (in days after planting) vs. unconditioned, in the cowpea MET, for several traits that had genetic signal in the chromosome 9 hotspot.

**Figure S11.** Within- and across-environment QTL profiles for all 11 chromosomes, conditional on flowering time (in days after planting) vs. unconditioned, in the cowpea MET, for several traits that had genetic signal in the chromosome 9 hotspot.

**Figure S12.** Heat map of founder allele effects at the chromosome 9 hotspot for flowering time and reproductive sensing traits (flowering time, FT; pod count, PdCt; flower count, FlCt).

**Figure S13.** Genomic predictive ability for agronomic and seed traits under the Within-Environment Fold-Out (WEFO) cross-validation scenario.

**Figure S14.** Temperature and solar radiation trends across the cowpea MAGIC multi-environment trial.

**Figure S15.** Spectral pretreatments used in partial least squares regression.

**Figure S16.** Principal component analysis (PCA) biplots showing spectral outlier detection using robust Mahalanobis distances calculated from the first five principal components of the spectral data after a given pretreatment was applied.

**Figure S17.** Wet chemistry outlier detection shows four outliers were detected across the training and validation set (n=320).

**Figure S18.** Diagnostic plots from spatial model fitting in the cowpea MET.

**Figure S19.** Within- and across-environment QTL profiles using the Haley-Knott (H-K) regression, linear mixed model (LMM), and leave-one-chromosome-out (LOCO) methods from the R/qtl2 package.

**Figure S20.** Comparison of QTL confidence interval widths under three interval estimation methods.

**Table S1.** Performance metrics of custom near-infrared spectroscopy calibrations of cowpea grain composition using partial least squares regression.

**Table S2.** Descriptive statistics of spatially corrected BLUEs for drone sensing traits across Davis and Parlier environments (2022–2023).

**Table S3.** Broad-sense heritability (H²) and within-environment repeatability (rept.) for drone sensing traits across Davis and Parlier environments (2022–2023).

**Table S4.** All discovered QTL loci.

**Table S5.** All discovered significant marker-trait associations.

**Table S6.** Genomic predictive ability by trait, method, and cross-validation scheme.

**Table S7.** Candidate genes within 250 kb of a Tier 1 QTL or GWAS peak.

**Table S8.** Factor analytic order used in ASReml multi-environment spatial modeling.

**Data S1.** Final BLUEs from the cowpea MAGIC MET.

**Data S2.** Master phenotypic records from the cowpea MAGIC MET.

