## Supplemental Tables and Figures for "Genetic mapping and genomic prediction for agronomic, grain compositional, and sensing-enabled traits in a cowpea MAGIC population along an environmental gradient"

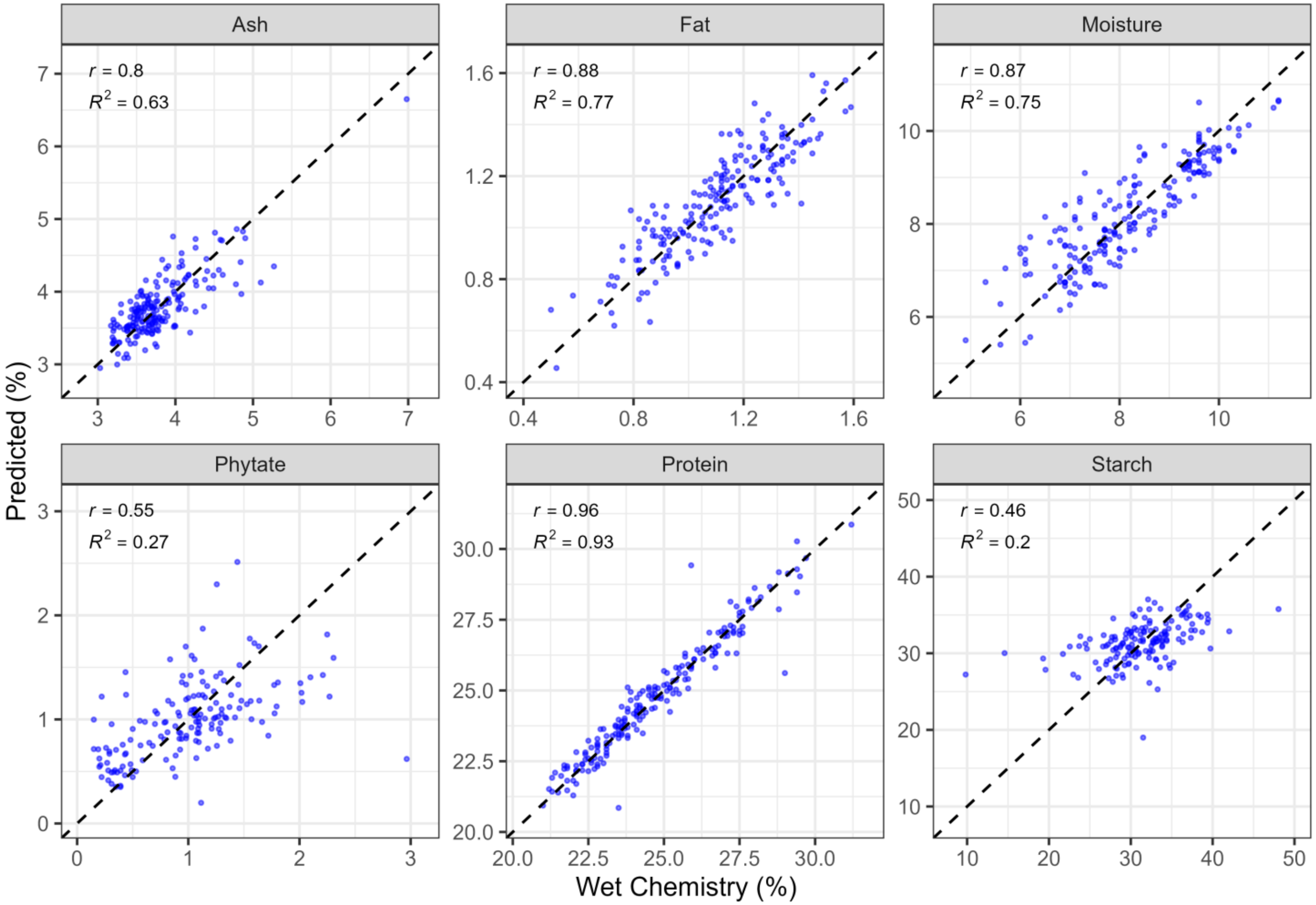


**Figure S1.** Custom near-infrared spectroscopy calibration for predicting cowpea grain composition. Each point represents a single field plot characterized spectrally with NIRS and for grain composition in wet chemistry bioassays. Each panel compares the percent composition (fresh weight) of macronutrient and antinutrient components predicted from the held-out prediction from the custom calibration with the wet chemistry values for the full 320 training and validation points collected across the cowpea MAGIC multi-environment trial. Pearson's *r* and the coefficient of determination ($R^2$) calculated by comparing the held-out predictions and wet chemistry values are shown in the top left of each panel.

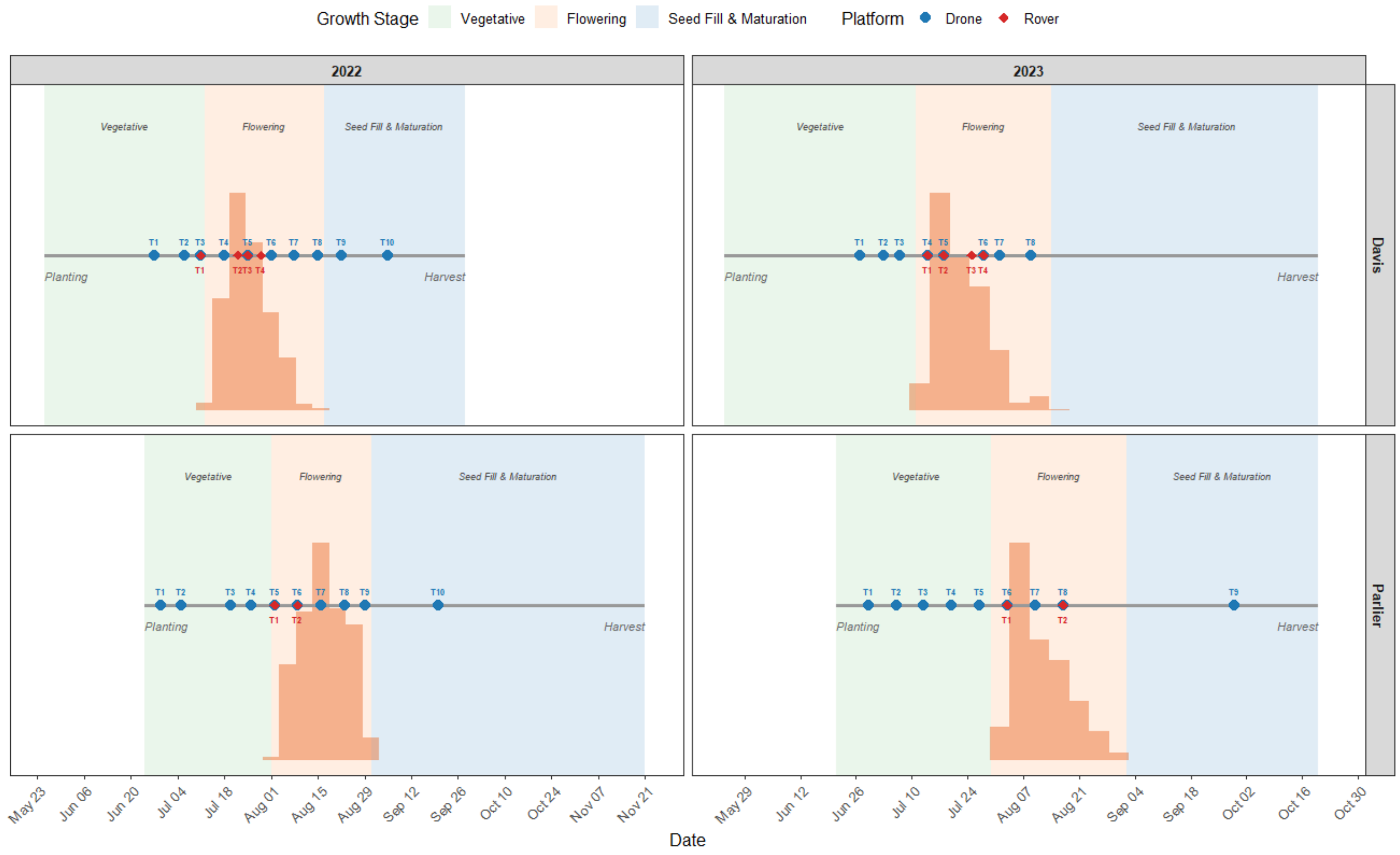


**Figure S2.** A timeline illustrating rover and drone sensing dates of the cowpea MAGIC MET within developmental periods based on flowering time data collected in each environment. Orange histograms show flowering time distributions. Due to equipment availability and logistical constraints, sensing was not performed in the Thermal location. Abbreviations: T1: Time point 1; T2 Time point 2; and so on, for subsequent time points.

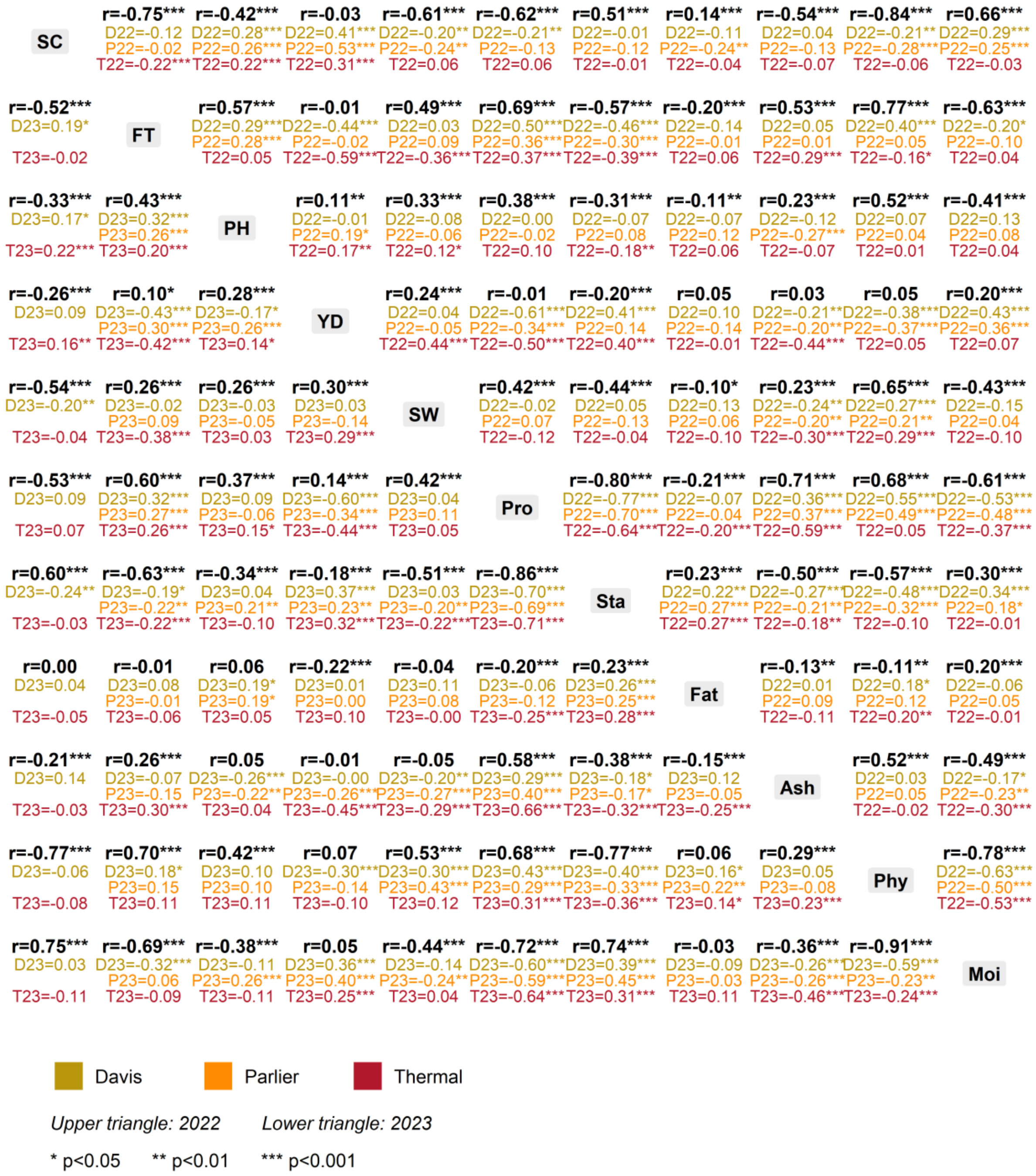


**Figure S3.** Pairwise phenotypic correlations among agronomic and grain composition traits across environments in the cowpea MAGIC population. Pearson correlation coefficients (*r*) were computed among replicate-averaged lines with significance testing. The upper triangle summarizes the 2022 environments, and the lower triangle, the 2023 environments. Within each cell, the bold value is the correlation pooled across all locations scored that year, and the three colored values below are the corresponding within-location correlations. Asterisks denote significance: *$P < 0.05$, **$P < 0.01$, ***$P < 0.001$. Blank entries indicate that the trait was not scored for that environment. Abbreviations: SC, stand count; FT, flowering time; PH, plant height; YD, yield; SW, 100-seed weight; Pro, protein; Sta, starch; Fat, fat; Ash, ash; Phy, phytate; and Moi, moisture

**Figure S4.** Within- and across-environment QTL profiles in the cowpea MET. Uploaded separately due to file size.

**Figure S5.** Manhattan plots and quantile-quantile (Q–Q) plots from GWAS in the cowpea MET. Uploaded separately due to file size.

**Figure S6.** QTL profiles for sensing-enabled traits in the Davis and Parlier environments (2022–2023). Uploaded separately due to file size.

**Figure S7.** Within- and across-environment QTL profiles for chromosome 8, conditional on seed weight vs. unconditioned, in the cowpea MET, for seed traits that had genetic signal in the chromosome 8 hotspot. Uploaded separately due to file size.

**Figure S8.** Within- and across-environment QTL profiles for all 11 chromosomes, conditional on seed weight vs. unconditioned, in the cowpea MET, for seed traits that had genetic signal in the chromosome 8 hotspot. Uploaded separately due to file size.

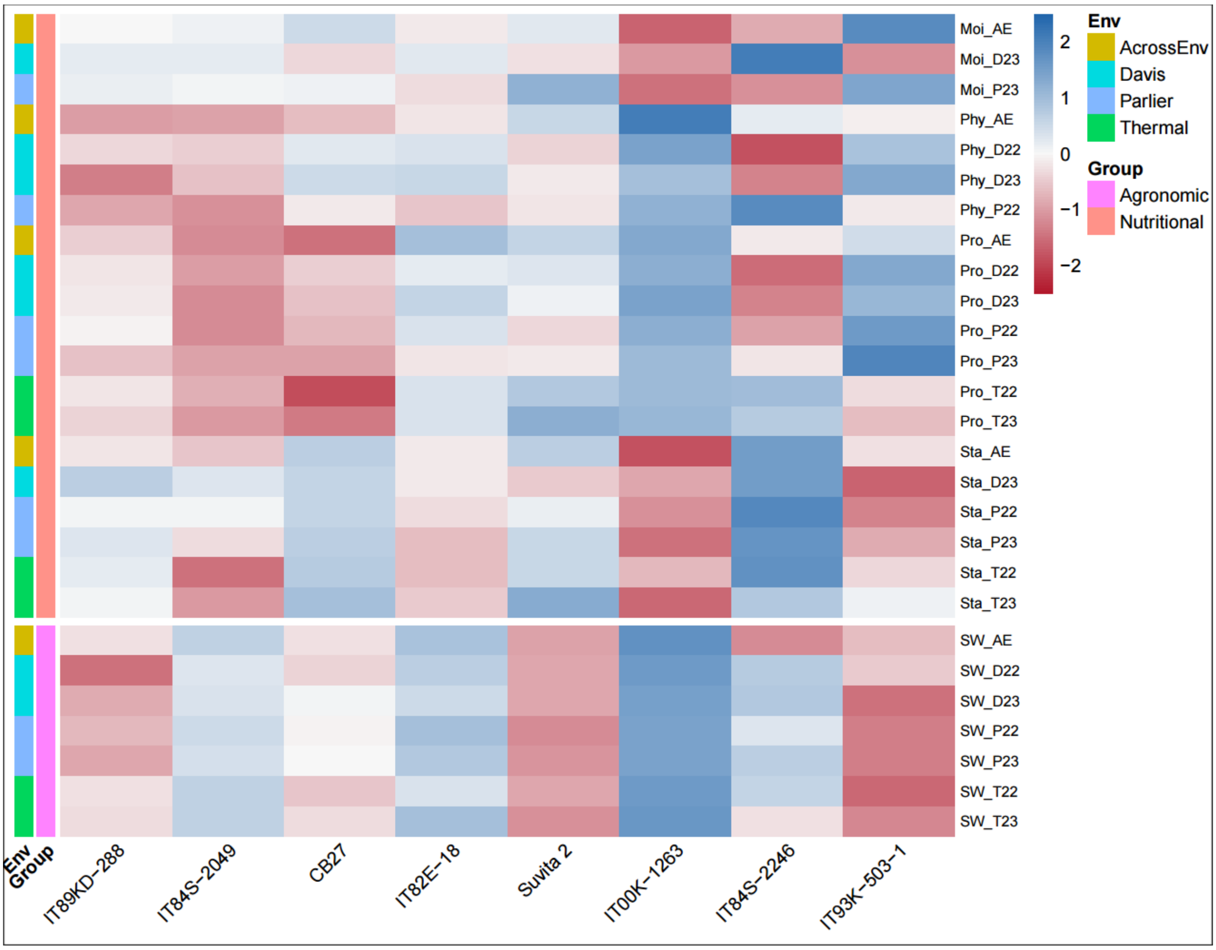


**Figure S9.** Heat map of founder allele effects at the chromosome 8 hotspot for seed traits (moisture, Moi; phytate, Phy; protein, Pro; starch, Sta; hundred-seed weight, SW). Rows show trait-environment combinations for all traits with a significant QTL at this locus; columns show the eight MAGIC founders. Founder effects were estimated at the peak marker position and z-scored within each trait-environment combination, such that color intensity reflects the number of standard deviations a given founder's effect deviates from the trait mean (blue = higher than average, red = lower than average, white = near average). Row annotations indicate trait group (Nutritional and Agronomic) and environment (AcrossEnv, Davis, Parlier, Thermal); AE denotes the across-environment BLUE, and D22/D23, P22/P23, and T22/T23 denote year-specific BLUEs for Davis, Parlier, and Thermal, respectively.

**Figure S10.** Within- and across-environment QTL profiles for chromosome 9, conditional on flowering time (in days after planting) vs. unconditioned, in the cowpea MET, for several traits that had genetic signal in the chromosome 9 hotspot. Uploaded separately due to file size.

**Figure S11.** Within- and across-environment QTL profiles for all 11 chromosomes, conditional on flowering time (in days after planting) vs. unconditioned, in the cowpea MET, for several traits that had genetic signal in the chromosome 9 hotspot. Uploaded separately due to file size.

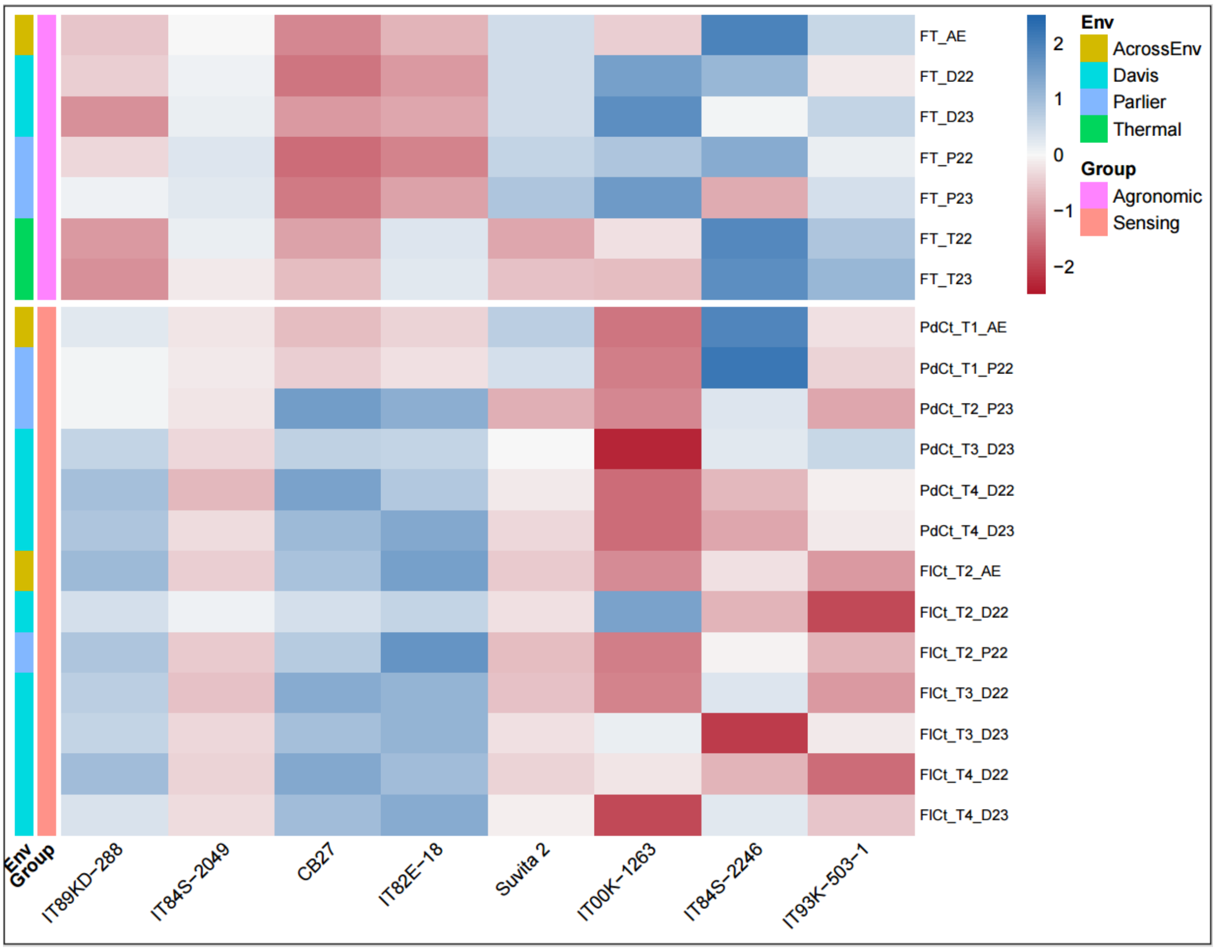


**Figure S12.** Heat map of founder allele effects at the chromosome 9 hotspot for flowering time and reproductive sensing traits (flowering time, FT; pod count, PdCt; flower count, FlCt). Rows show trait-environment-timepoint combinations for all traits with a significant QTL at this locus; columns show the eight MAGIC founders. Founder effects were estimated at the peak marker position and z-scored within each trait-environment combination, such that color intensity reflects the number of standard deviations a given founder's effect deviates from the trait mean (blue = higher than average, red = lower than average). Row annotations indicate trait group (Agronomic and Sensing and environment (AcrossEnv, Davis, Parlier, Thermal); AE denotes the across-environment BLUE, and D22/D23, P22/P23, and T22/T23 denote year-specific BLUEs for Davis, Parlier, and Thermal, respectively. T1–T4 time point suffixes indicate sequential rover imaging dates for pod and flower count.

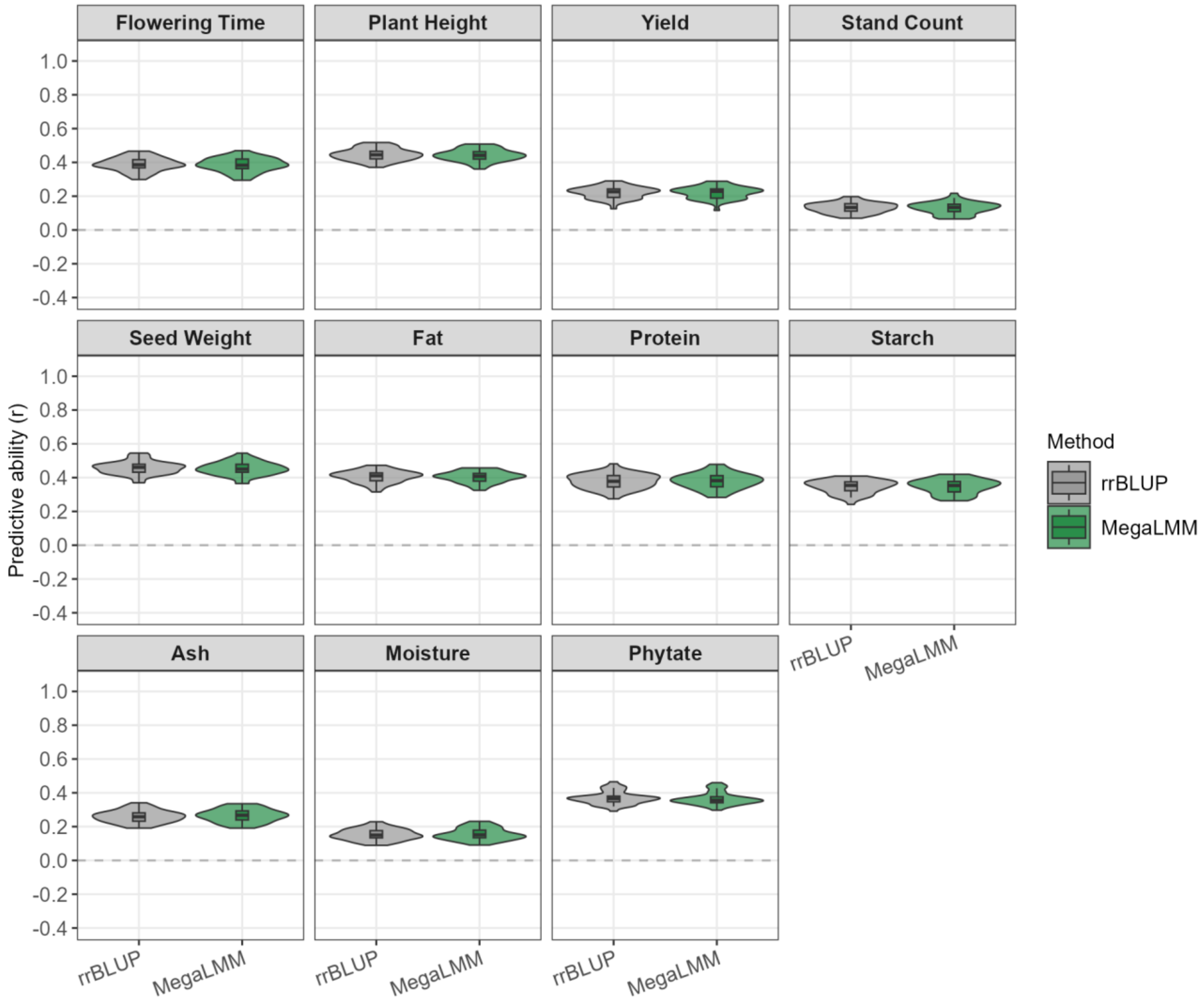


**Figure S13.** Genomic predictive ability for agronomic and seed traits under the Within-Environment Fold-Out (WEFO) cross-validation scenario. Predictive ability (Pearson correlation between predicted and observed values, *r*) is shown for RR-BLUP (gray) and MegaLMM (green) across 11 traits spanning agronomic (Flowering Time, Plant Height, Yield, Stand Count) and seed (Seed weight, Fat, Protein, Starch, Ash, Moisture, Phytate) trait categories. Violins represent the distribution of predictive ability across 20 replicates of 2-fold cross-validation (40 fold estimates in total) performed in each environment; internal box plots show the median and interquartile range.

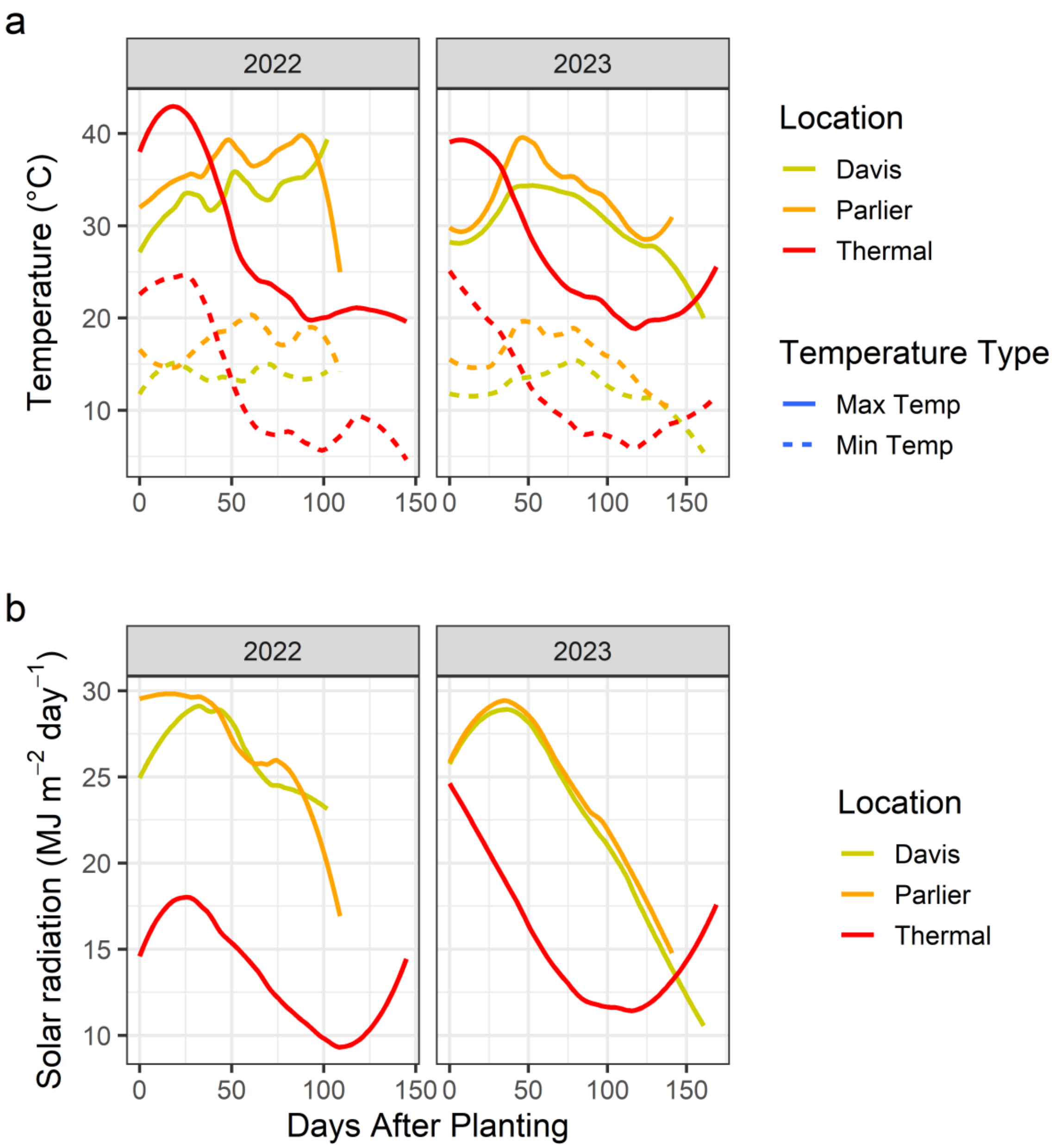


**Figure S14.** Temperature and solar radiation trends across the cowpea MAGIC multi-environment trial.

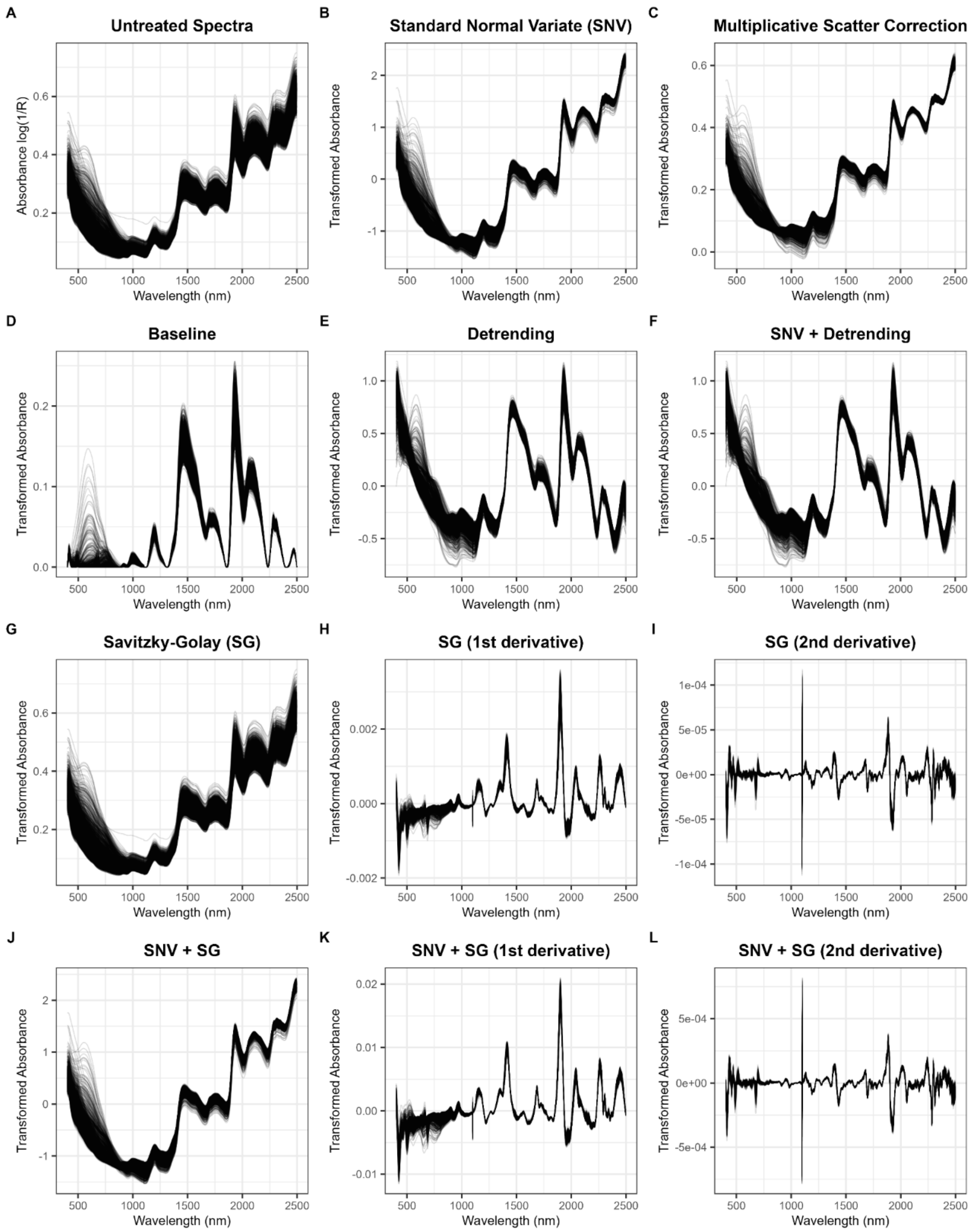


**Figure S15.** Spectral pretreatments used in partial least squares regression. All pretreatments are single transformations except those in panels F and J–L, which contain two transformations starting with standard normal variate (SNV).

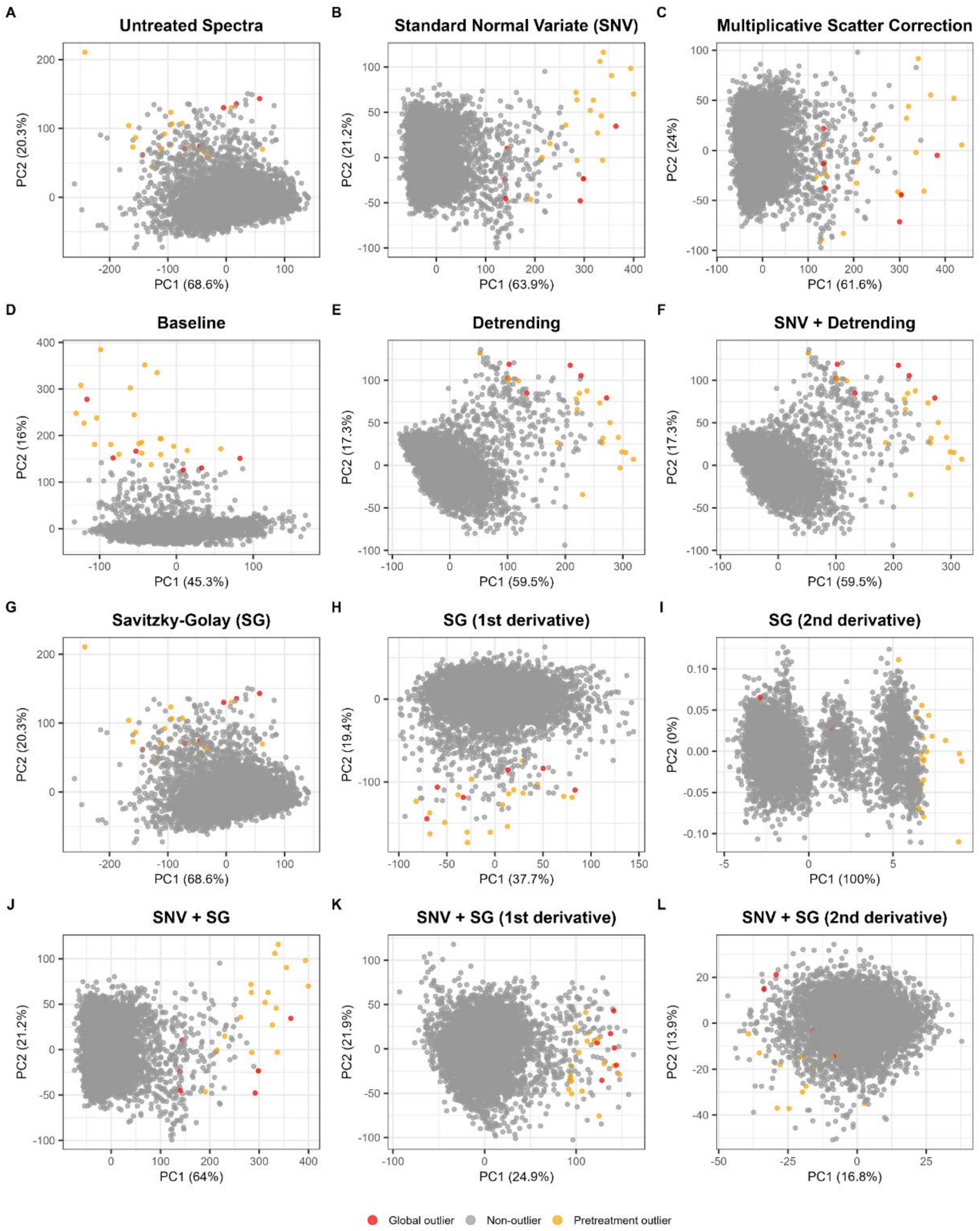


**Figure S16.** Principal component analysis (PCA) biplots showing spectral outlier detection using robust Mahalanobis distances calculated from the first five principal components of the spectral data after a given pretreatment was applied. Samples identified as global outliers (red) were removed prior to fitting partial least squares regression models used to develop custom near-infrared spectroscopy calibrations.

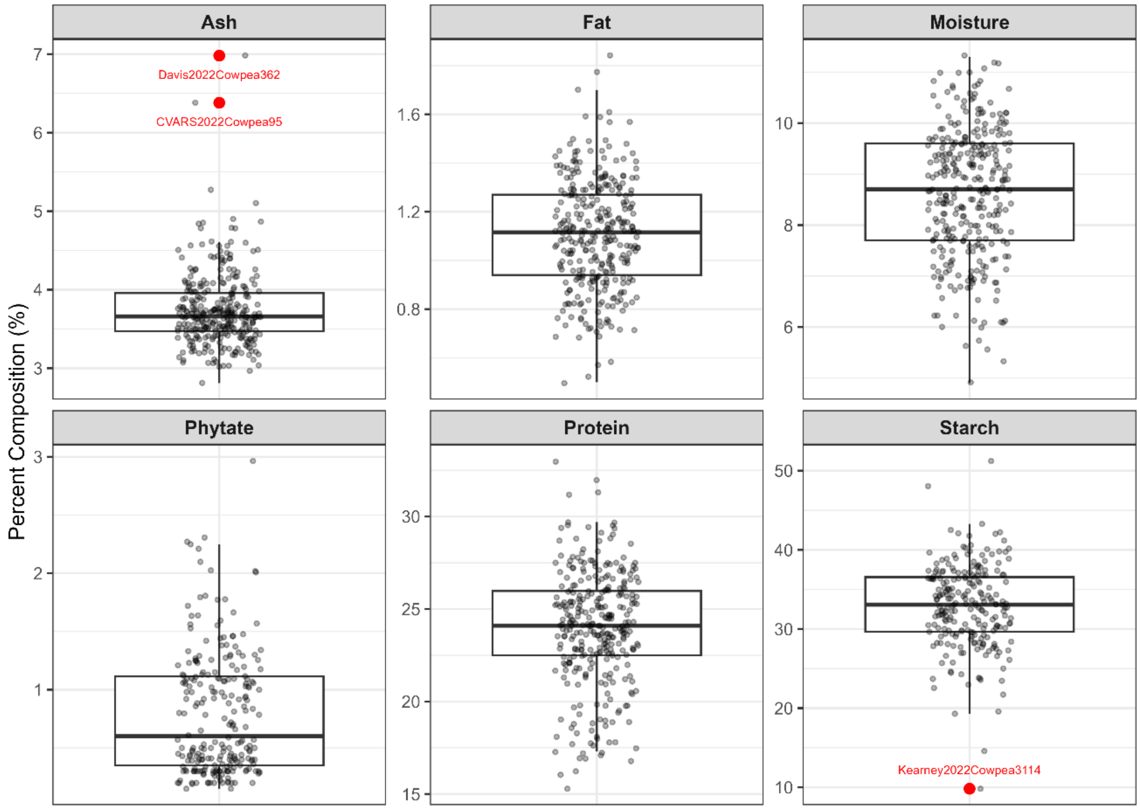


**Figure S17.** Wet chemistry outlier detection shows four outliers were detected across the training and validation set (n=320). Points in red are outliers and were removed from analysis.

**Figure S18.** Diagnostic plots from spatial model fitting in the cowpea MET. Uploaded separately due to file size.

**Figure S19.** Within- and across-environment QTL profiles using the Haley-Knott (H-K) regression, linear mixed model (LMM), and leave-one-chromosome-out (LOCO) methods from the R/qtl2 package. Uploaded separately due to file size.

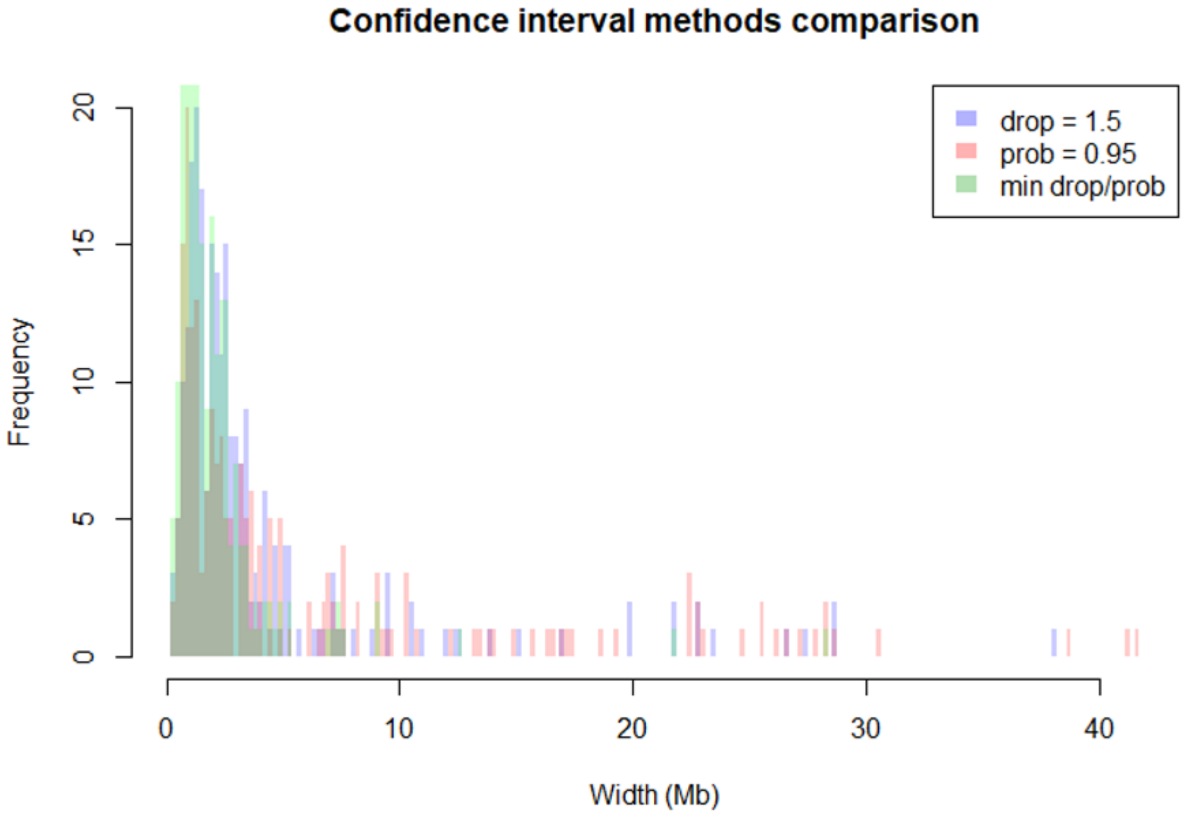


**Figure S20.** Comparison of QTL confidence interval widths under three interval estimation methods. Histograms show the distribution of confidence interval widths (Mb) across all detected QTL peaks for the 1.5-LOD drop interval (blue), 95% Bayes credible interval (pink), and the minimum of the two methods (green), the latter of which was used as the final interval estimate in this study. The minimum method consistently produced narrower intervals than the 95% Bayes credible interval alone, reducing the number of wide intervals while retaining the precision of the LOD drop method where applicable.

**Table S1.** Performance metrics of custom near-infrared spectroscopy calibrations of cowpea grain composition using partial least squares regression. Models were fitted to standard normal variate pretreated spectra and evaluated by five-fold cross-validation; all statistics in this table were derived from held-out predictions, with each sample predicted by a model fitted without it. *n*, samples with paired predictions and reference values; *ncomp*, latent variables retained; *r*, Pearson correlation of predicted with reference values; $R^2$, coefficient of determination relative to the 1:1 line; *RMSECV*, root mean squared error of cross-validation; *bias*, mean signed deviation of predicted from reference values, with positive indicating overprediction; *slope*, from ordinary least squares regression of predicted on reference values, with below 1 indicating compression of the predicted range. Traits are % fresh weight. Starch and phytate were not assayed in the Riverside 2023 and Davis 2024 environments that were used to supplement custom calibration development for the study environments.

| Trait | mean | SD | *r* | $R^2$ | RMSECV | bias | slope | n | ncomp |
|---|---|---|---|---|---|---|---|---|---|
| **Ash** | 3.73 | 0.49 | 0.80 | 0.63 | 0.28 | 0.01 | 0.73 | 196 | 28 |
| **Fat** | 1.12 | 0.23 | 0.88 | 0.77 | 0.10 | 0.00 | 0.84 | 196 | 30 |
| **Moisture** | 8.59 | 1.31 | 0.87 | 0.75 | 0.63 | 0.02 | 0.76 | 196 | 8 |
| **Phytate** | 0.79 | 0.52 | 0.55 | 0.27 | 0.44 | -0.01 | 0.40 | 158 | 19 |
| **Protein** | 23.76 | 3.20 | 0.96 | 0.93 | 0.57 | 0.04 | 0.95 | 196 | 21 |
| **Starch** | 32.86 | 5.27 | 0.46 | 0.20 | 4.28 | 0.13 | 0.25 | 153 | 12 |

**Table S2.** Descriptive statistics of spatially corrected BLUEs for drone sensing traits across Davis and Parlier environments (2022–2023).

| Trait (unit) | n † | Grand mean ± SD | Davis '22 mean ± SD | Davis '23 mean ± SD | Parlier '22 mean ± SD | Parlier '23 mean ± SD |
|---|---|---|---|---|---|---|
| **Sensing (Drone)** | | | | | | |
| PtHt T1 (m)‡ | 180 | 0.213 ± 0.007 | 0.279 ± 0.034 | 0.373 ± 0.052 | 0.130 ± 0.005 | 0.154 ± 0.013 |
| PtHt T2 (m)‡ | 180 | 0.354 ± 0.010 | 0.440 ± 0.059 | 0.447 ± 0.040 | 0.095 ± 0.008 | 0.378 ± 0.018 |
| PtHt T3 (m)‡ | 180 | 0.450 ± 0.027 | 0.487 ± 0.068 | 0.547 ± 0.049 | 0.199 ± 0.035 | 0.530 ± 0.031 |
| PtHt T4 (m) | 180 | 0.540 ± 0.033 | 0.651 ± 0.076 | 0.603 ± 0.074 | 0.263 ± 0.033 | 0.627 ± 0.034 |
| PtHt T5 (m) | 180 | 0.647 ± 0.049 | 0.783 ± 0.091 | 0.694 ± 0.085 | 0.373 ± 0.049 | 0.725 ± 0.048 |
| PtHt T6 (m) | 180 | 0.607 ± 0.061 | 0.419 ± 0.094 | 0.724 ± 0.095 | 0.493 ± 0.073 | 0.784 ± 0.057 |
| PtHt T7 (m) | 180 | 0.680 ± 0.081 | 0.565 ± 0.107 | 0.711 ± 0.093 | 0.476 ± 0.097 | 0.958 ± 0.090 |
| PtHt T8 (m) | 180 | 0.795 ± 0.087 | 0.915 ± 0.112 | 0.731 ± 0.105 | 0.573 ± 0.093 | 0.954 ± 0.097 |
| PtHt T9 (m) | 176 | 0.544 ± 0.086 | 0.114 ± 0.111 | — | 0.570 ± 0.092 | 0.936 ± 0.102 |
| VgFr T1 (proportion)‡ | 144 | 0.061 ± 0.005 | 0.070 ± 0.033 | 0.212 ± 0.044 | 0.004 ± 0.002 | 0.003 ± 0.002 |
| VgFr T2 (proportion)‡ | 180 | 0.133 ± 0.010 | 0.207 ± 0.085 | 0.394 ± 0.069 | 0.020 ± 0.006 | 0.036 ± 0.011 |
| VgFr T3 (proportion)‡ | 180 | 0.264 ± 0.041 | 0.347 ± 0.117 | 0.652 ± 0.081 | 0.164 ± 0.048 | 0.113 ± 0.028 |
| VgFr T4 (proportion)‡ | 180 | 0.408 ± 0.058 | 0.555 ± 0.142 | 0.651 ± 0.099 | 0.313 ± 0.073 | 0.209 ± 0.038 |
| VgFr T5 (proportion)‡ | 180 | 0.562 ± 0.073 | 0.725 ± 0.131 | 0.677 ± 0.101 | 0.425 ± 0.109 | 0.427 ± 0.069 |
| VgFr T6 (proportion)‡ | 180 | 0.747 ± 0.071 | 0.808 ± 0.129 | 0.780 ± 0.102 | 0.805 ± 0.109 | 0.515 ± 0.087 |
| VgFr T7 (proportion)‡ | 180 | 0.784 ± 0.067 | 0.888 ± 0.102 | 0.805 ± 0.103 | 0.877 ± 0.055 | 0.531 ± 0.089 |
| VgFr T8 (proportion)‡ | 180 | 0.926 ± 0.039 | 0.927 ± 0.076 | 0.880 ± 0.078 | 0.944 ± 0.038 | 0.925 ± 0.045 |
| VgFr T9 (proportion)‡ | 176 | 0.936 ± 0.037 | 0.934 ± 0.071 | — | 0.924 ± 0.054 | 0.940 ± 0.041 |
| PtHt max rate (DAP)‡ | 180 | 23.1 ± 2.5 | 33.0 ± 6.3 | 31.1 ± 7.4 | 34.5 ± 9.6 | 8.65 ± 0.63 |
| VgFr max rate (DAP) | 180 | 42.6 ± 2.7 | 52.1 ± 5.3 | 38.3 ± 5.4 | 40.6 ± 3.2 | 38.7 ± 7.9 |
| PtHt min rate (DAP)‡ | 180 | 84.9 ± 1.7 | 87.1 ± 4.9 | 70.6 ± 8.7 | 76.7 ± 11.9 | 98.8 ± 3.9 |
| VgFr min rate (DAP)‡ | 180 | 71.7 ± 5.3 | 61.5 ± 26.1 | 44.6 ± 16.2 | 77.0 ± 8.3 | 97.5 ± 6.0 |
| PtHt max growth rate (m day$^{-1}$)‡ | 180 | 0.014 ± 0.002 | 0.013 ± 0.003 | 0.015 ± 0.003 | 0.011 ± 0.003 | 0.022 ± 0.003 |
| VgFr max growth rate (day$^{-1}$) | 180§ | 0.028 ± 0.006 | 0.029 ± 0.004 | 0.028 ± 0.006 | 0.033 ± 0.003 | 0.022 ± 0.003 |
| PtHt min growth rate (m day$^{-1}$)‡ | 180 | -0.003 ± 0.002 | -0.006 ± 0.005 | -0.004 ± 0.009 | -0.000 ± 0.003 | -0.003 ± 0.003 |
| VgFr min growth rate (day$^{-1}$)‡ | 179 | -0.001 ± 0.001 | -0.003 ± 0.003 | 0.002 ± 0.004 | -0.003 ± 0.002 | -0.001 ± 0.002 |
| PtHt AUC (m·day) | 180 | 44.1 ± 3.4 | 34.9 ± 4.5 | 33.7 ± 3.7 | 32.3 ± 4.0 | 74.0 ± 5.8 |
| VgFr AUC (day)‡ | 180 | 46.1 ± 3.0 | 48.3 ± 6.1 | 32.3 ± 3.9 | 48.2 ± 3.5 | 56.6 ± 3.5 |
| Canopy closure (DAP)‡ | 134 | 56.8 ± 3.9 | 66.6 ± 7.5 | 63.5 ± 6.5 | 49.7 ± 4.3 | 52.5 ± 4.1 |

† Indicates the number of genotypes for which across-environment BLUEs were estimated for a given trait.
‡ Trait required Box-Cox transformation in at least one environment prior to spatial modeling; back-transformed values are shown.
§ Denotes that the value of n for VgFr max growth rate was determined based on BLUPs and the maximum number of genotypes assayed for this trait within a given environment.

**Table S3.** Broad-sense heritability ($H^2$) and within-environment repeatability (rept.) for drone sensing traits across Davis and Parlier environments (2022–2023).

| **Trait** | **$H^2$** | **N envs** | **Davis '22 rept.** | **Davis '23 rept.** | **Parlier '22 rept.** | **Parlier '23 rept.** |
|---|---|---|---|---|---|---|
| **Sensing (Drone)** | | | | | | |
| PtHt T1 | 0.29 | 4 | 0.56 | 0.79 | 0.35 | 0.40 |
| PtHt T2 | 0.70 | 4 | 0.70 | 0.81 | 0.62 | 0.73 |
| PtHt T3 | 0.75 | 4 | 0.69 | 0.37 | 0.75 | 0.67 |
| PtHt T4 | 0.79 | 4 | 0.76 | **0.90** | 0.79 | 0.82 |
| PtHt T5 | **0.88** | 4 | 0.82 | **0.90** | 0.92 | 0.93 |
| PtHt T6 | 0.85 | 4 | 0.86 | 0.88 | **0.94** | **0.94** |
| PtHt T7 | 0.81 | 4 | 0.89 | 0.87 | 0.24 | 0.88 |
| PtHt T8 | 0.88 | 4 | **0.90** | 0.88 | 0.90 | 0.93 |
| PtHt T9 | 0.83 | 3 | 0.86 | — | 0.87 | 0.86 |
| VgFr T1 | 0.81 | 4 | 0.68 | 0.44 | 0.88 | 0.82 |
| VgFr T2 | 0.72 | 4 | 0.74 | 0.56 | 0.77 | 0.87 |
| VgFr T3 | 0.79 | 4 | 0.72 | 0.44 | 0.85 | 0.86 |
| VgFr T4 | 0.83 | 4 | 0.79 | 0.69 | 0.81 | 0.83 |
| VgFr T5 | 0.83 | 4 | 0.79 | 0.72 | 0.85 | 0.89 |
| VgFr T6 | 0.83 | 4 | 0.81 | 0.78 | 0.79 | 0.87 |
| VgFr T7 | 0.76 | 4 | 0.81 | 0.78 | 0.46 | 0.51 |
| VgFr T8 | 0.76 | 4 | 0.76 | 0.79 | 0.46 | 0.76 |
| VgFr T9 | 0.64 | 3 | 0.74 | — | 0.56 | 0.50 |
| PtHt max rate DAP | 0.81 | 4 | 0.17 | 0.42 | 0.31 | 0.11 |
| VgFr max rate DAP | 0.72 | 4 | 0.69 | 0.15 | 0.52 | 0.48 |
| PtHt min rate DAP | 0.76 | 4 | 0.36 | 0.14 | 0.35 | 0.28 |
| VgFr min rate DAP | 0.05 | 4 | 0.28 | 0.00 | 0.19 | 0.01 |
| PtHt max growth rate | 0.79 | 4 | 0.62 | 0.52 | 0.58 | 0.71 |
| VgFr max growth rate | 0.83 | 4 | 0.66 | 0.46 | 0.62 | 0.23 |
| PtHt min growth rate | 0.64 | 4 | 0.60 | 0.19 | 0.30 | 0.82 |
| VgFr min growth rate | 0.05 | 4 | 0.26 | 0.29 | 0.45 | 0.25 |
| PtHt AUC | 0.86 | 4 | 0.77 | 0.83 | 0.85 | 0.92 |
| VgFr AUC | 0.81 | 4 | 0.79 | 0.74 | 0.83 | 0.73 |
| Canopy closure | 0.80 | 4 | 0.77 | 0.48 | 0.81 | 0.76 |

**Table S4.** All discovered QTL loci. Uploaded separately due to file size.

**Table S5.** All discovered significant marker-trait associations. Uploaded separately due to file size.

**Table S6.** Genomic predictive ability by trait, method, and cross-validation scheme. Uploaded separately due to file size.

**Table S7.** Candidate genes within 250 kb of a Tier 1 QTL or GWAS peak. Uploaded separately due to file size.

**Table S8.** Factor analytic order used in ASReml multi-environment spatial modeling.

| Trait Class | Trait | Factor Analytic Order |
|---|---|---|
| Agronomic | Stand_count | 2 |
| | Flowerting Time | 2 |
| | Plant Height | 2 |
| | Yield | 2 |
| Seed Trait | Hundred Seed Weight | 2 |
| | Moisture | 2 |
| | Fat | 2 |
| | Starch | 2 |
| | Ash | 2 |
| | Protein | 2 |
| | Phytate | 2 |
| Sensing | flower_count_T1 | 2 |
| | flower_count_T2 | 2 |
| | pod_count_T1 | 2 |
| | pod_count_T2 | 2 |
| | max_value_DAP_Plant_Height | 2 |
| | max_value_DAP_Vegetation_Fraction | 2 |
| | max_rate_DAP_Plant_Height | 2 |
| | max_rate_DAP_Vegetation_Fraction | 2 |
| | min_rate_DAP_Plant_Height | 2 |
| | min_rate_DAP_Vegetation_Fraction | 2 |
| | max_rate_Plant_Height | 2 |
| | max_rate_Vegetation_Fraction | **1** |
| | min_rate_Plant_Height | 2 |
| | min_rate_Vegetation_Fraction | **1** |
| | AUC_Plant_Height | 2 |
| | AUC_Vegetation_Fraction | 2 |
| | VgFrCl_0.8_DAP | 2 |
| | PtHt_T1 | **1** |
| | PtHt_T2 | 2 |
| | PtHt_T3 | 2 |
| | PtHt_T4 | 2 |
| | PtHt_T5 | 2 |
| | PtHt_T6 | 2 |
| | PtHt_T7 | 2 |
| | PtHt_T8 | 2 |
| | PtHt_T9 | 2 |
| | VgFr_T1 | 2 |
| | VgFr_T2 | **1** |
| | VgFr_T3 | **1** |
| | VgFr_T4 | 2 |
| | VgFr_T5 | 2 |
| | VgFr_T6 | 2 |
| | VgFr_T7 | 2 |
| | VgFr_T8 | 2 |
| | VgFr_T9 | **1** |

Table only includes traits for which there were at least three unique environments with phenotypic records. Abbreviations: DAP, days after planting; AUC, area under the curve; PtHt, Plant height; VgFr, Vegetation fraction.
