## Supplementary material for "Genetic mapping and genomic prediction for agronomic, grain compositional, and sensing-enabled traits in a cowpea MAGIC population along an environmental gradient": Large supplemental tables and figures, uploaded separately: SuppFig_05_MET_Manhattan.pdf

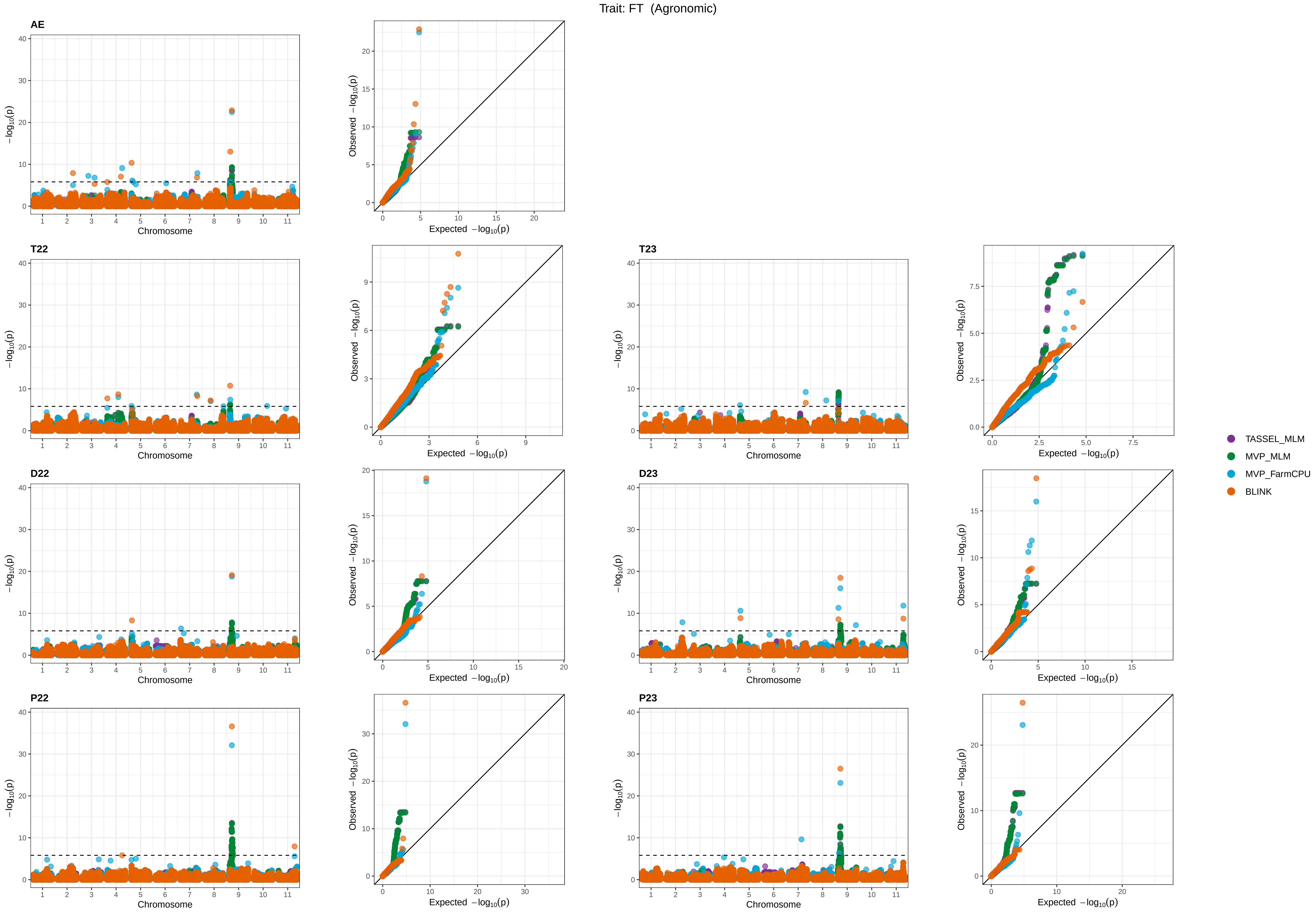
Trait: FT (Agronomic)
AE
T22
T23
D22
D23
P22
P23
Chromosome
Expected −log10(p)
Observed −log10(p)
−log10(p)
TASSEL_MLM
MVP_MLM
MVP_FarmCPU
BLINK

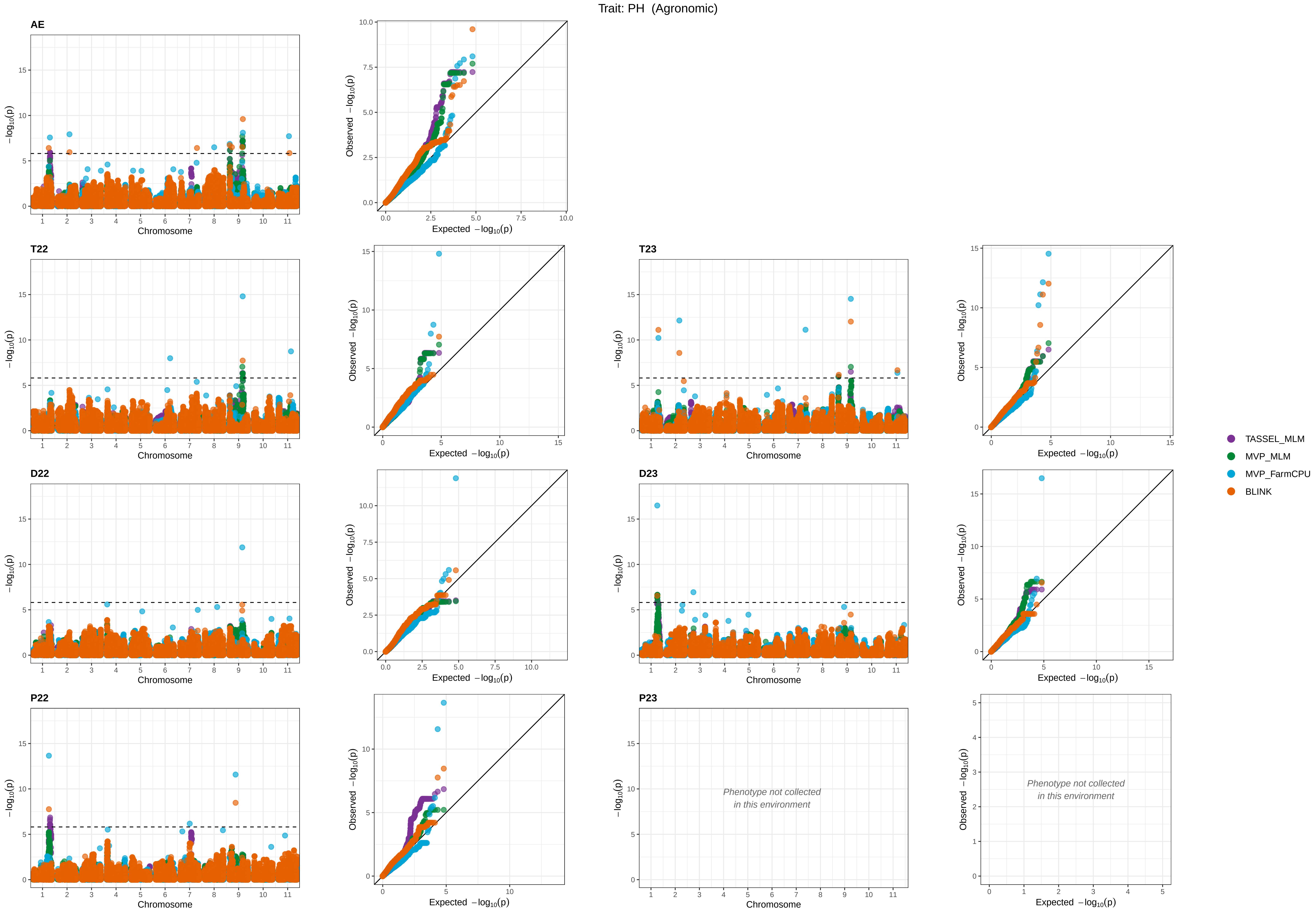

Trait: PH (Agronomic)
AE
T22
T23
D22
D23
P22
P23
Chromosome
−log10(p)
Expected −log10(p)
Observed −log10(p)
Phenotype not collected
in this environment
TASSEL_MLM
MVP_MLM
MVP_FarmCPU
BLINK

Trait: SC (Agronomic)

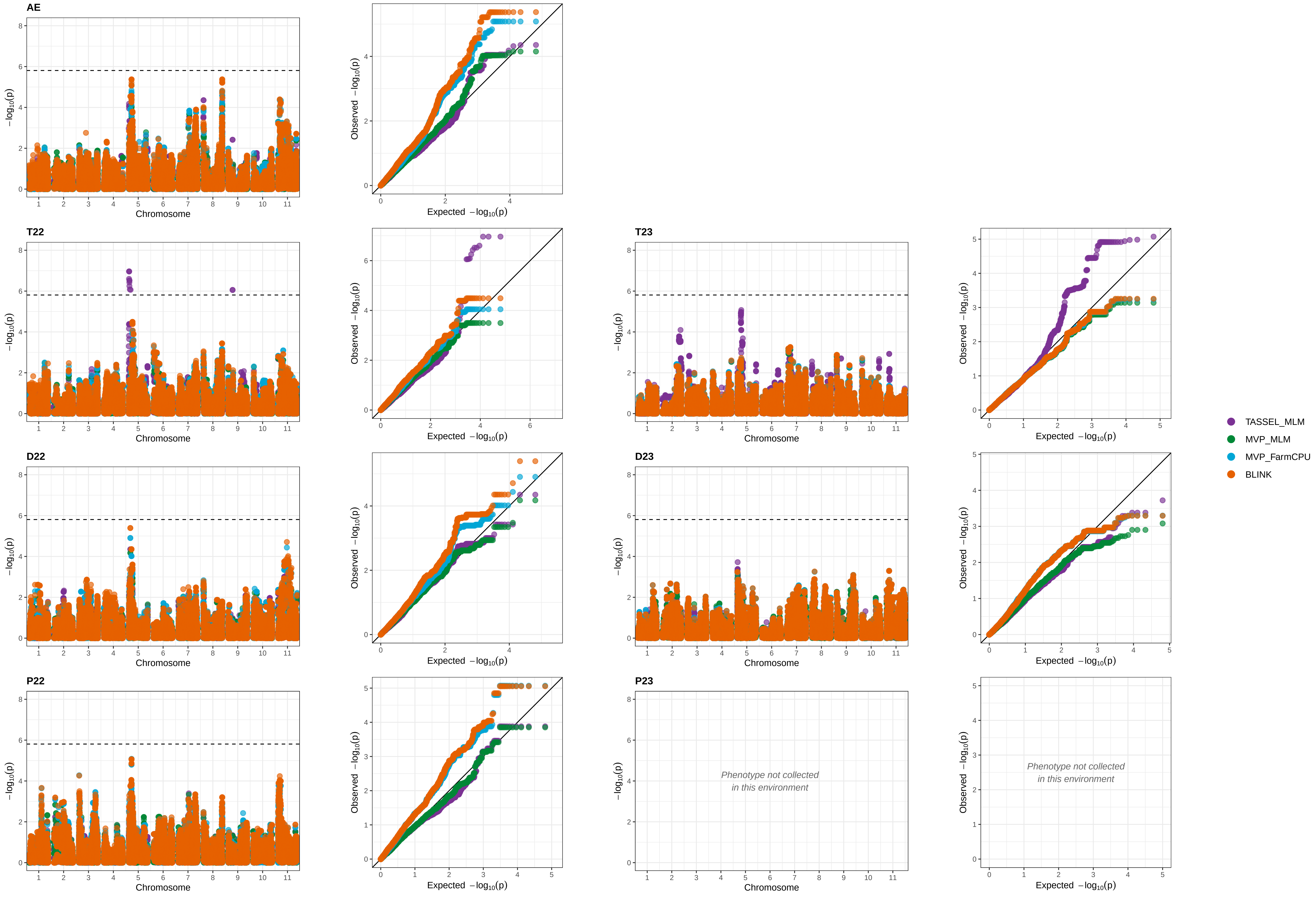

Trait: SW (Agronomic)

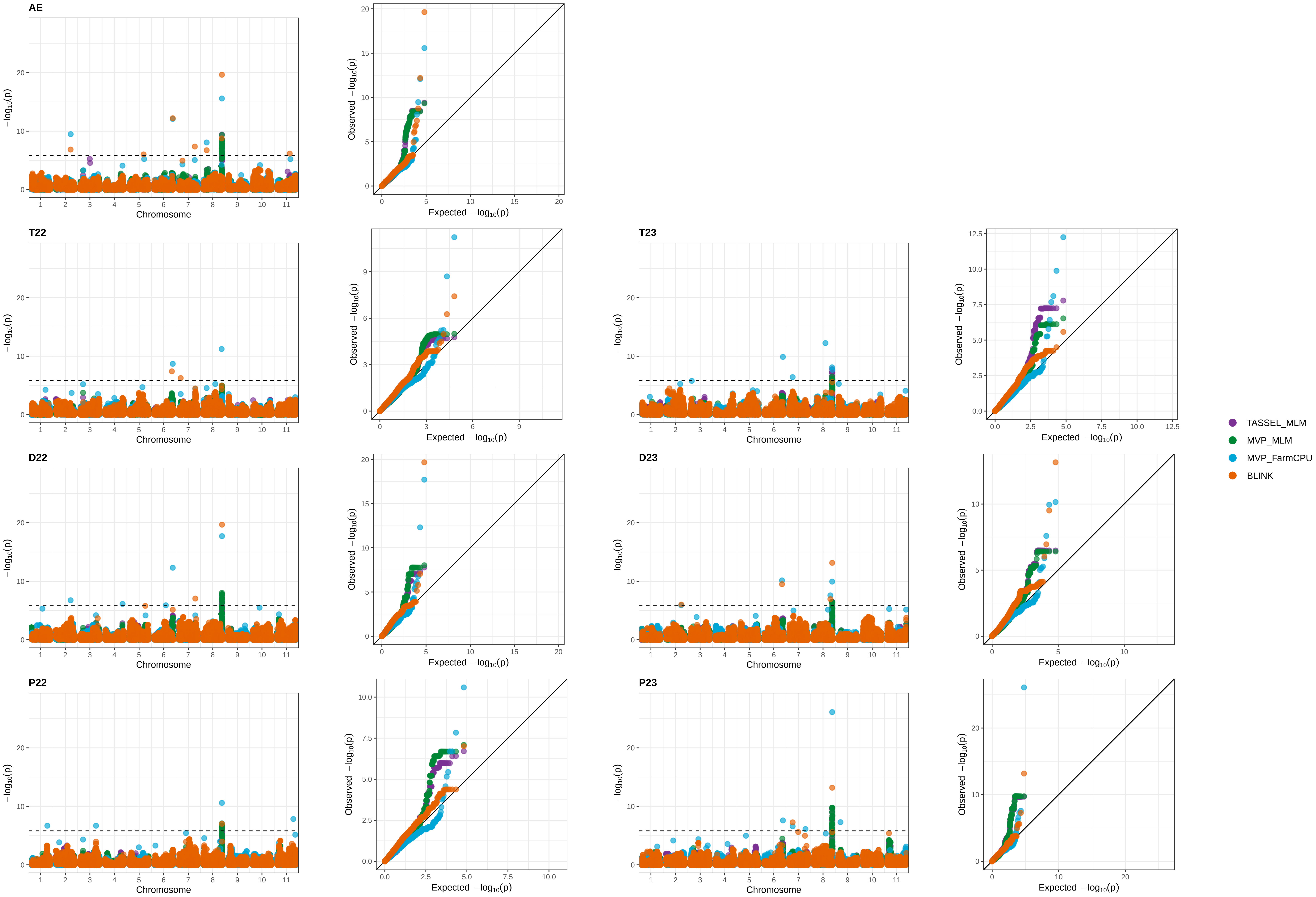

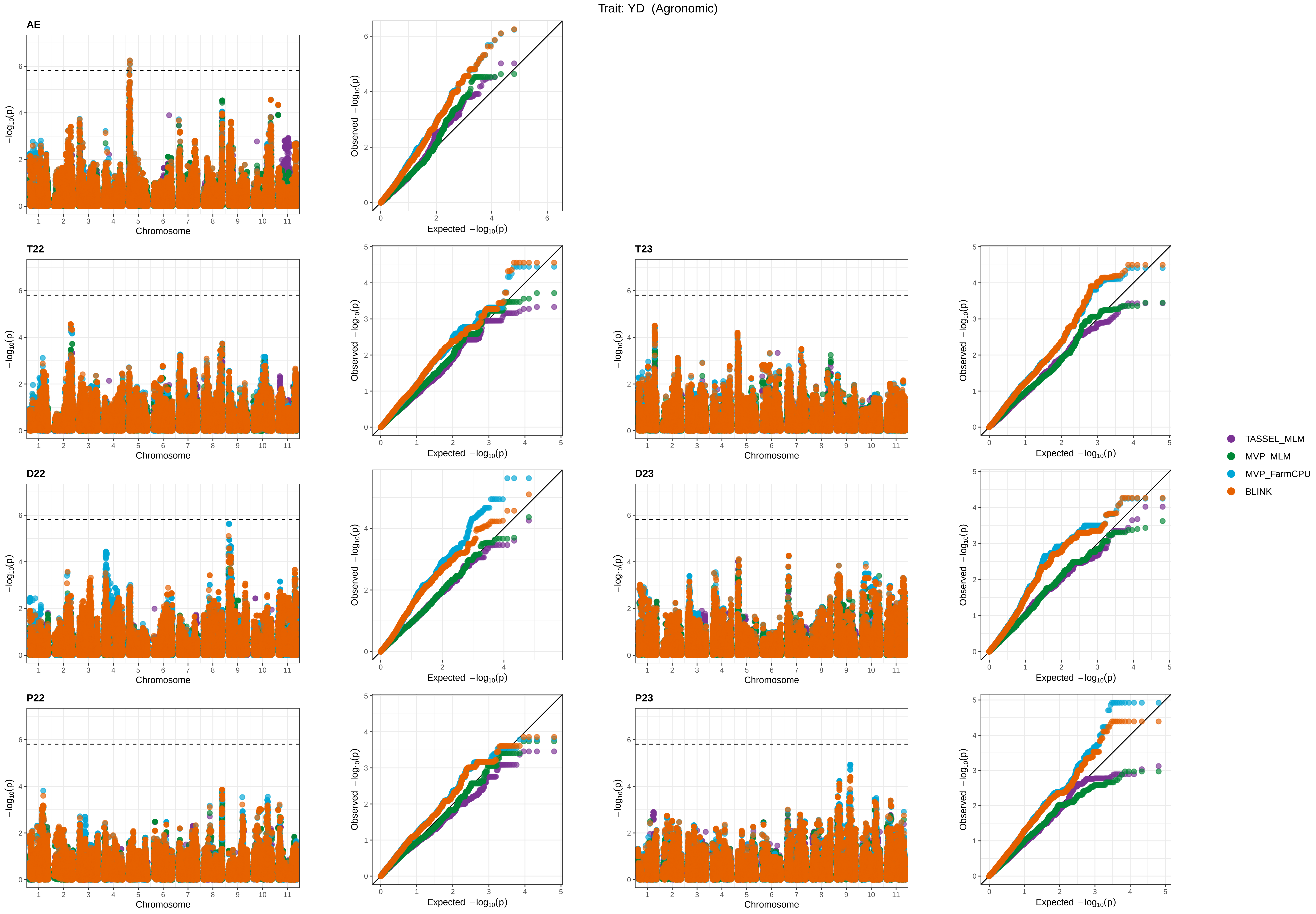

Trait: YD (Agronomic)
AE
T22
T23
D22
D23
P22
P23
Chromosome
$-\log_{10}(p)$
Observed $-\log_{10}(p)$
Expected $-\log_{10}(p)$
TASSEL_MLM
MVP_MLM
MVP_FarmCPU
BLINK

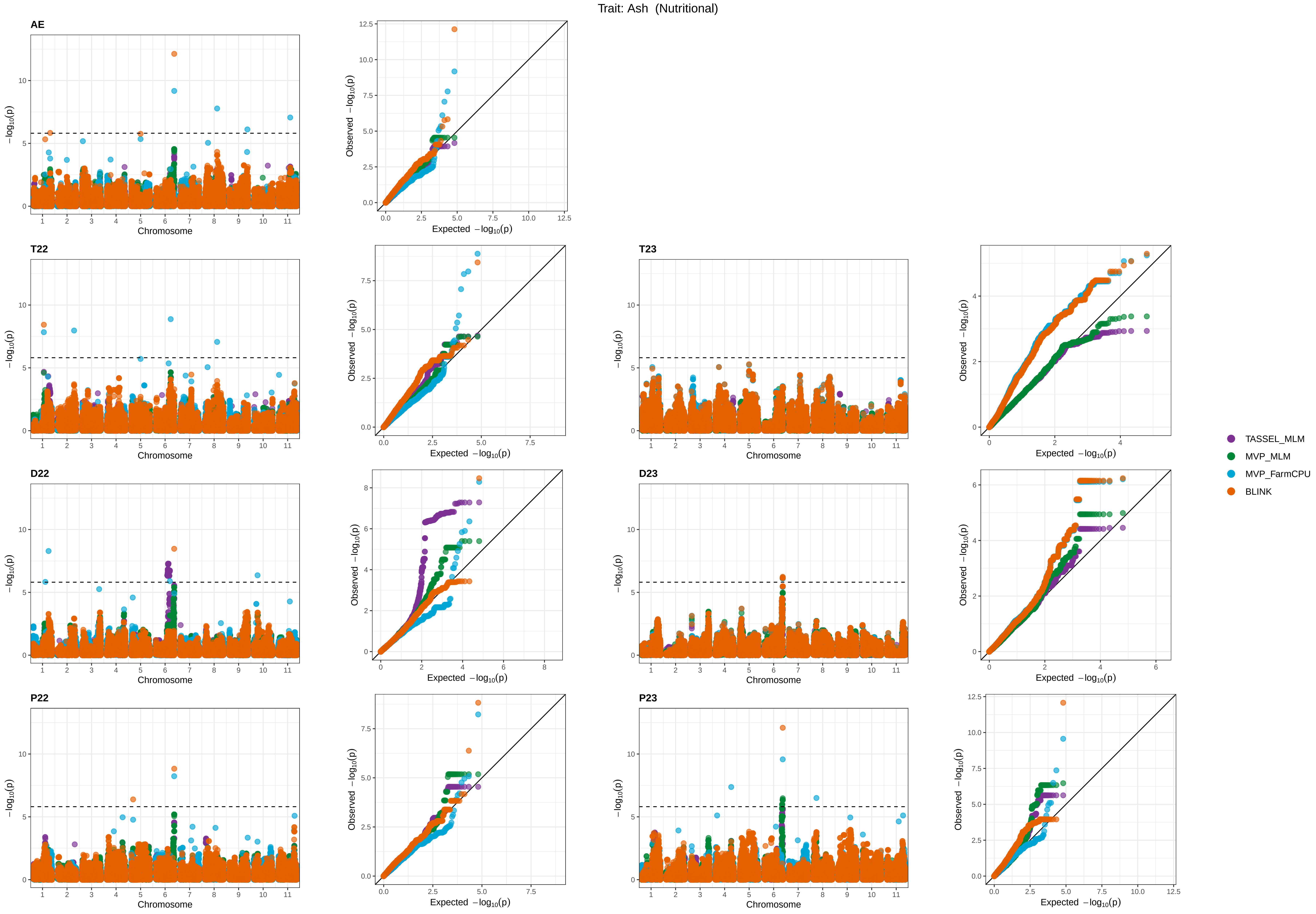

Trait: Ash (Nutritional)
AE
T22
T23
D22
D23
P22
P23
Chromosome
$-\log_{10}(p)$
Expected $-\log_{10}(p)$
Observed $-\log_{10}(p)$
TASSEL_MLM
MVP_MLM
MVP_FarmCPU
BLINK

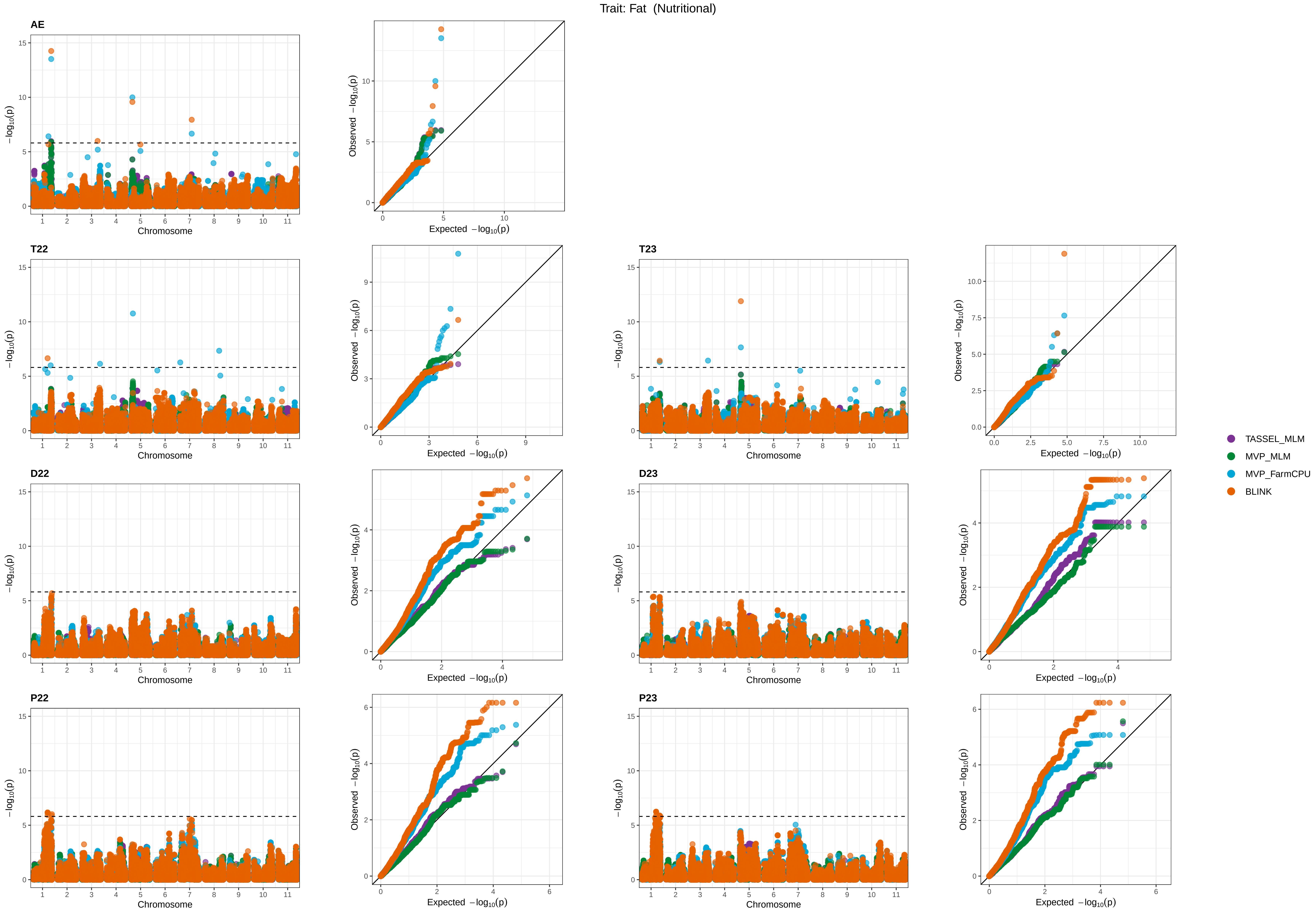

Trait: Fat (Nutritional)
AE
T22
T23
D22
D23
P22
P23
−log10(p)
Chromosome
Observed −log10(p)
Expected −log10(p)
TASSEL_MLM
MVP_MLM
MVP_FarmCPU
BLINK

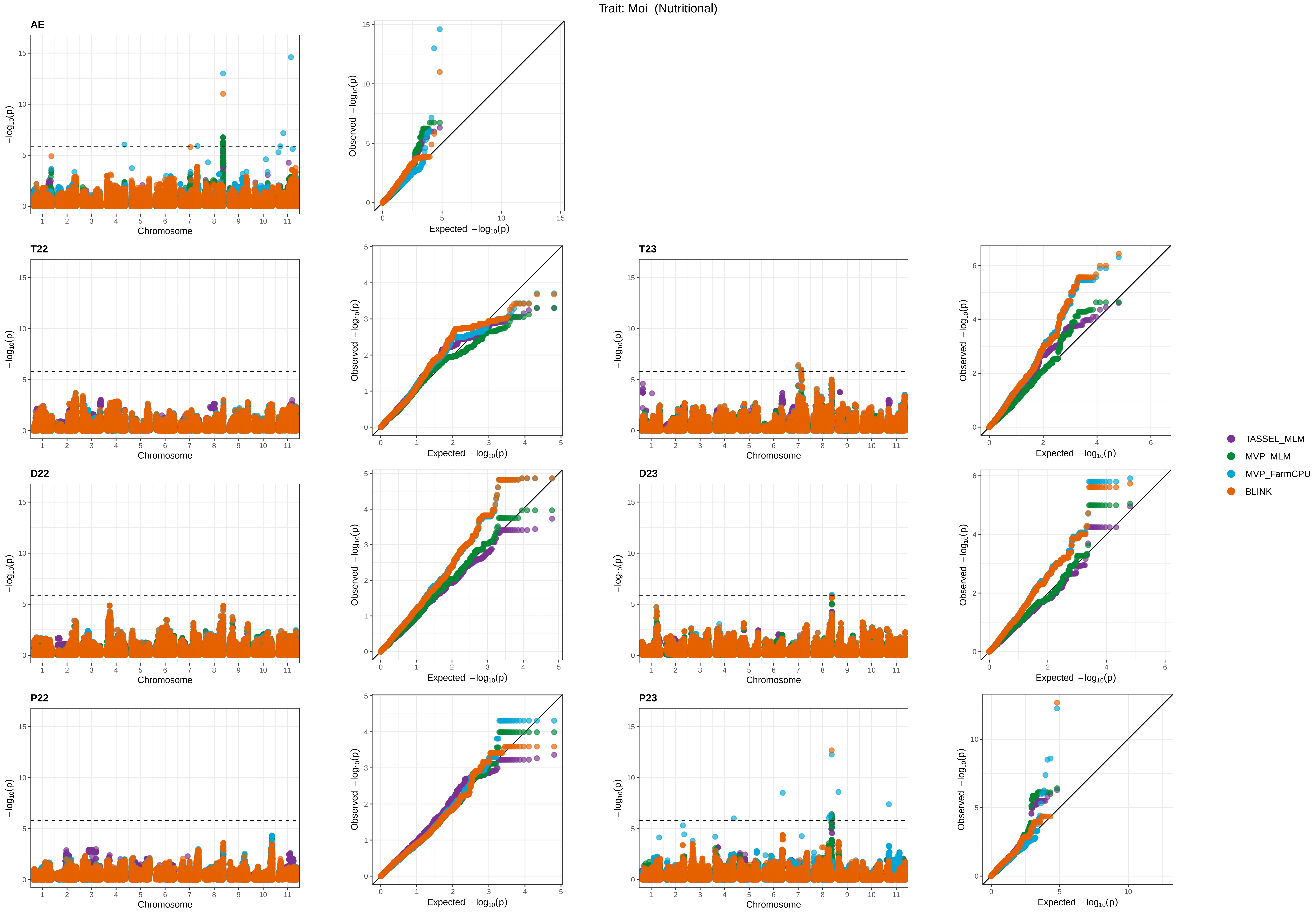
Trait: Moi (Nutritional)
AE
T22
T23
D22
D23
P22
P23
−log10(p)
Chromosome
Observed −log10(p)
Expected −log10(p)
TASSEL_MLM
MVP_MLM
MVP_FarmCPU
BLINK

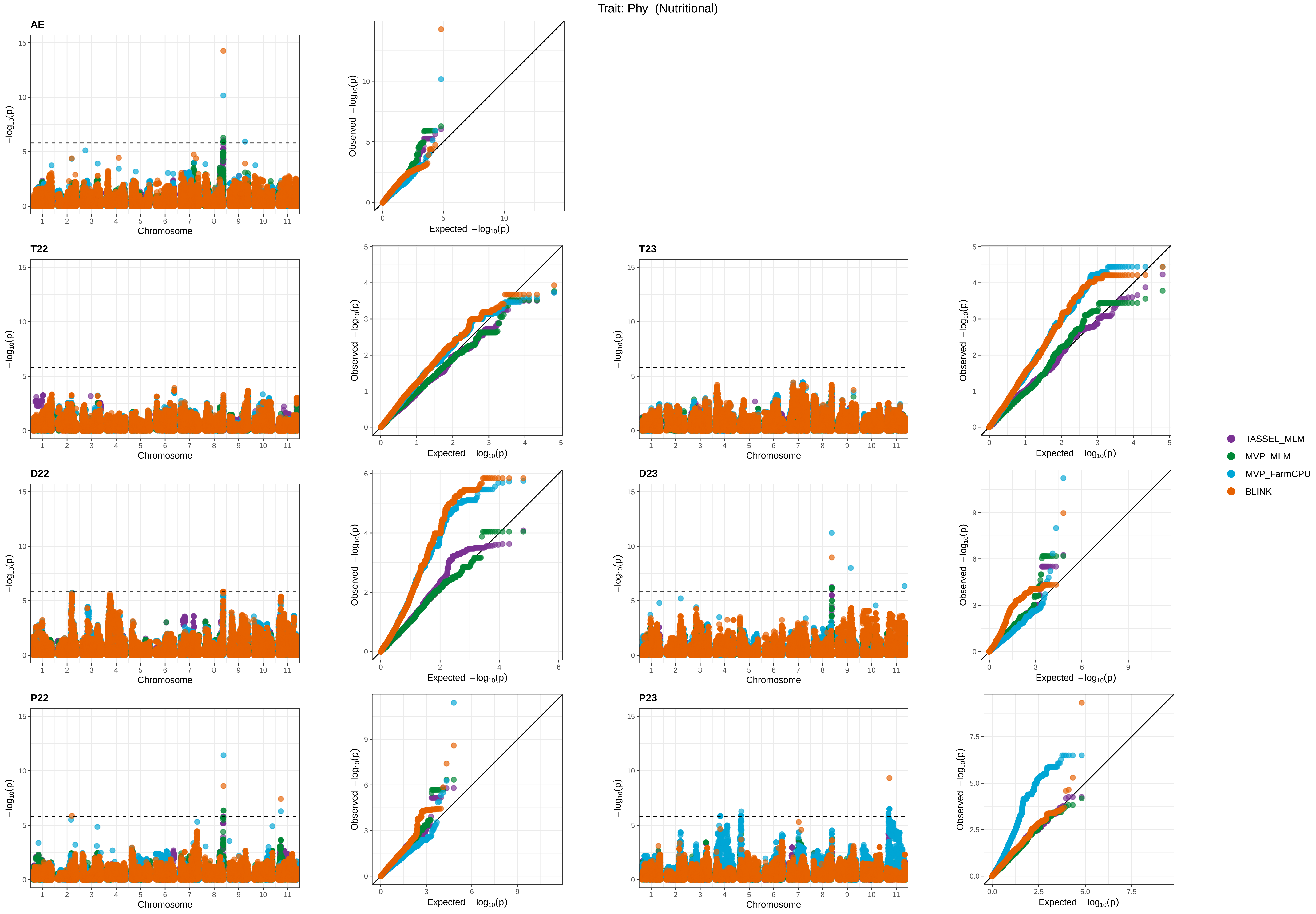
Trait: Phy (Nutritional)
AE
T22
T23
D22
D23
P22
P23
Chromosome
Expected −log10(p)
Observed −log10(p)
TASSEL_MLM
MVP_MLM
MVP_FarmCPU
BLINK

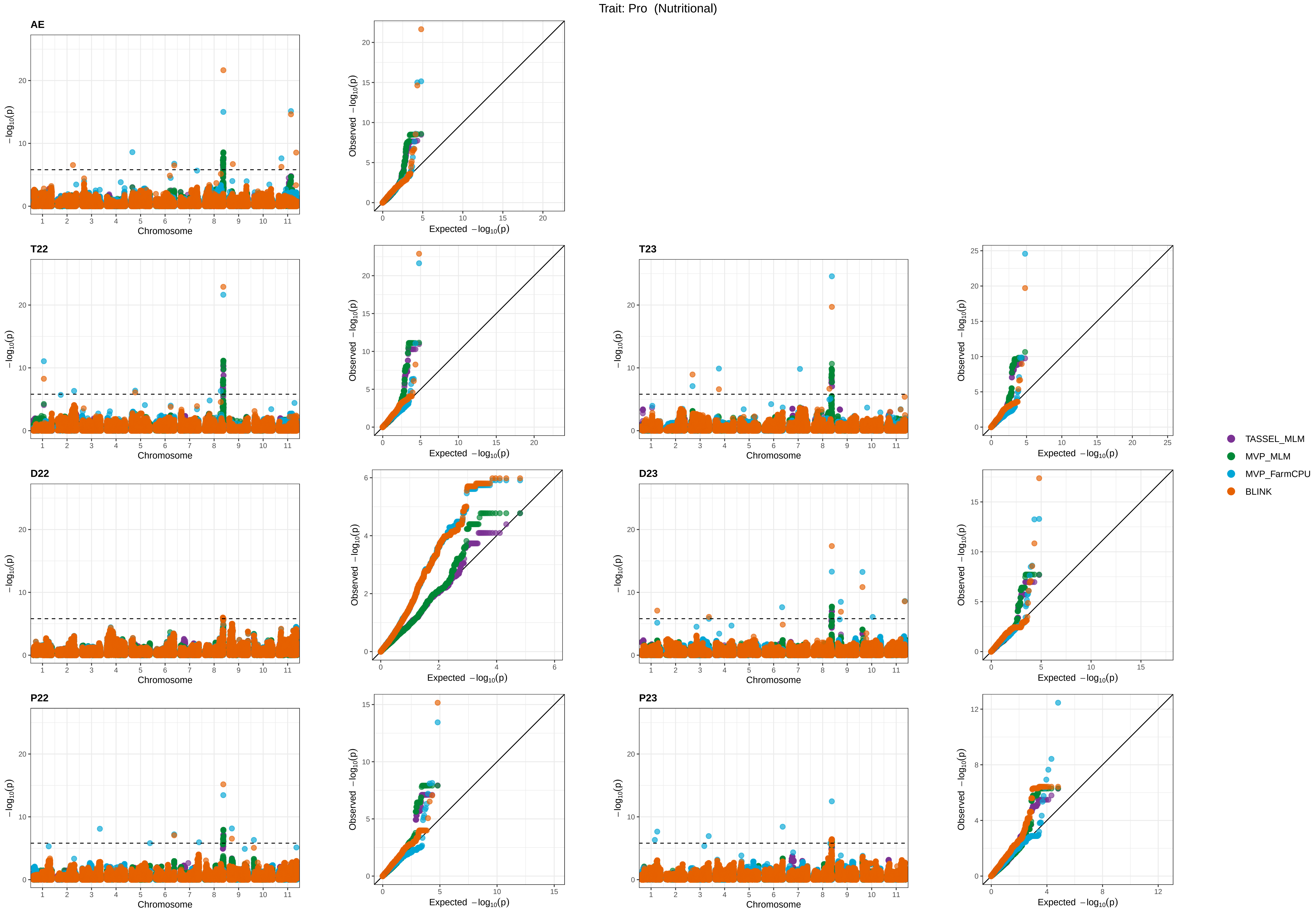

Trait: Pro (Nutritional)
AE
T22
T23
D22
D23
P22
P23
Chromosome
$-\log_{10}(p)$
Expected $-\log_{10}(p)$
Observed $-\log_{10}(p)$
TASSEL_MLM
MVP_MLM
MVP_FarmCPU
BLINK

Trait: Sta (Nutritional)

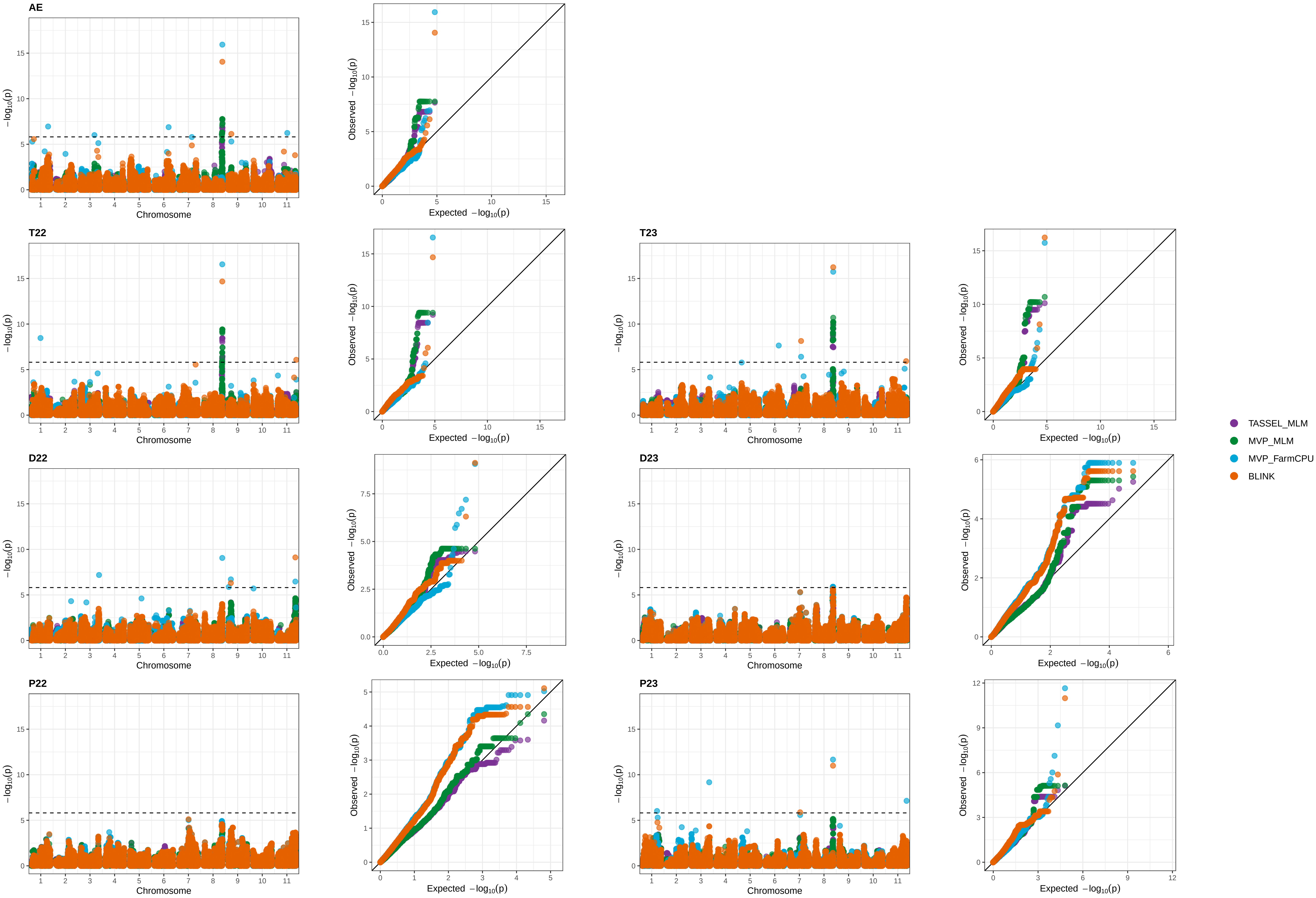

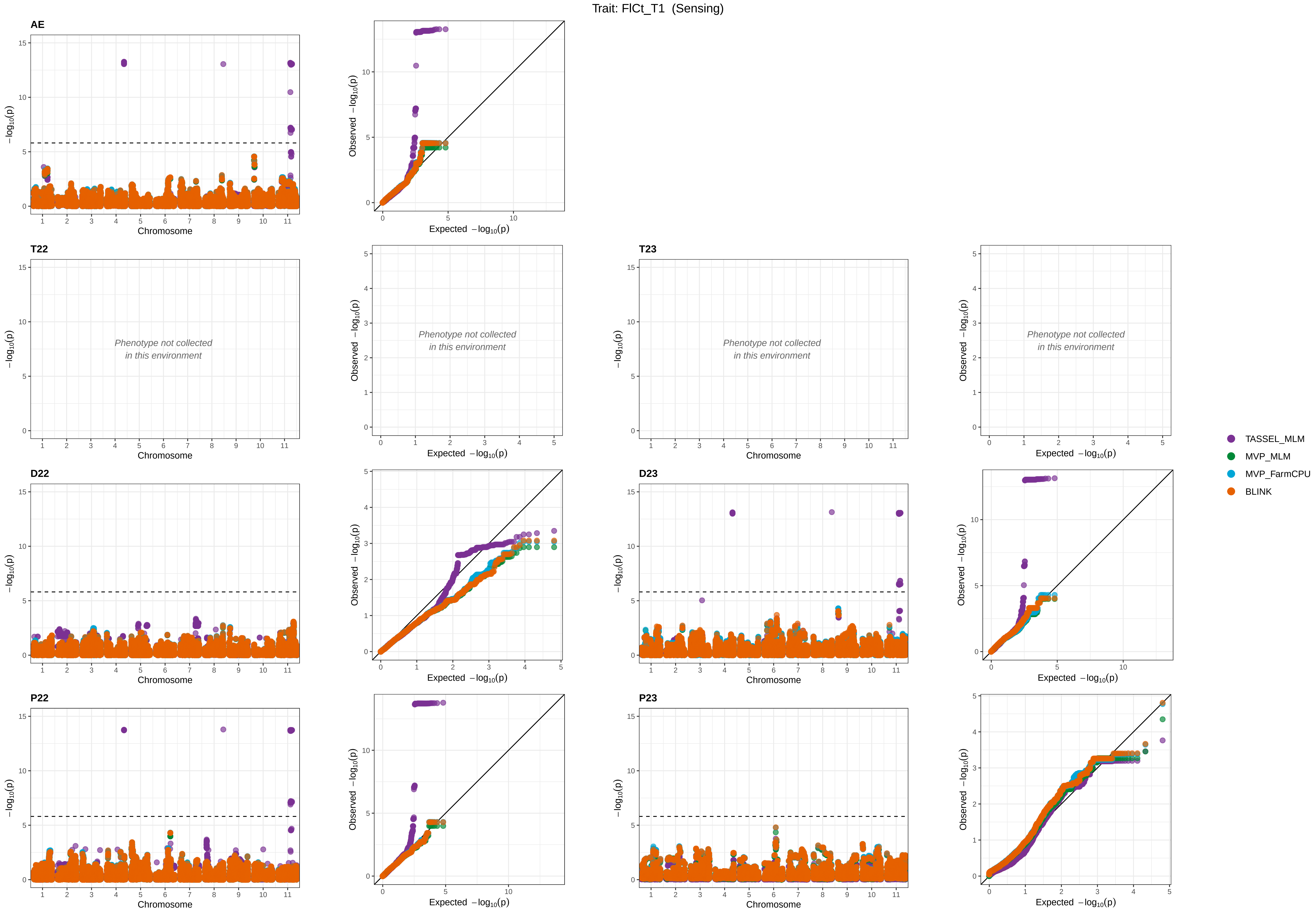
Trait: FlCt_T1 (Sensing)
AE
T22
T23
D22
D23
P22
P23
−log10(p)
Chromosome
Observed −log10(p)
Expected −log10(p)
Phenotype not collected
in this environment
TASSEL_MLM
MVP_MLM
MVP_FarmCPU
BLINK

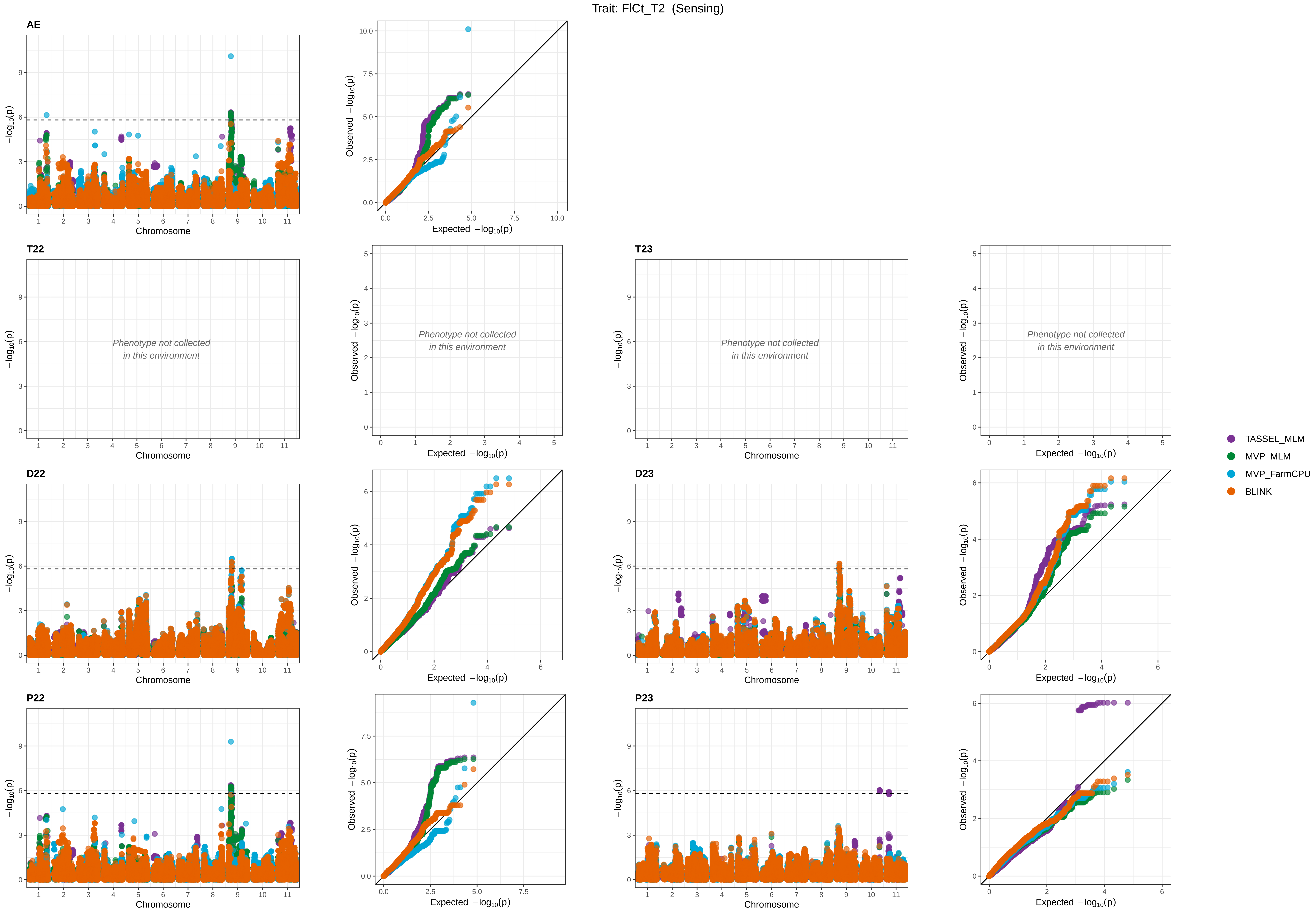
Trait: FlCt_T2 (Sensing)
AE
T22
T23
D22
D23
P22
P23
Phenotype not collected in this environment
Chromosome
−log10(p)
Expected −log10(p)
Observed −log10(p)
TASSEL_MLM
MVP_MLM
MVP_FarmCPU
BLINK

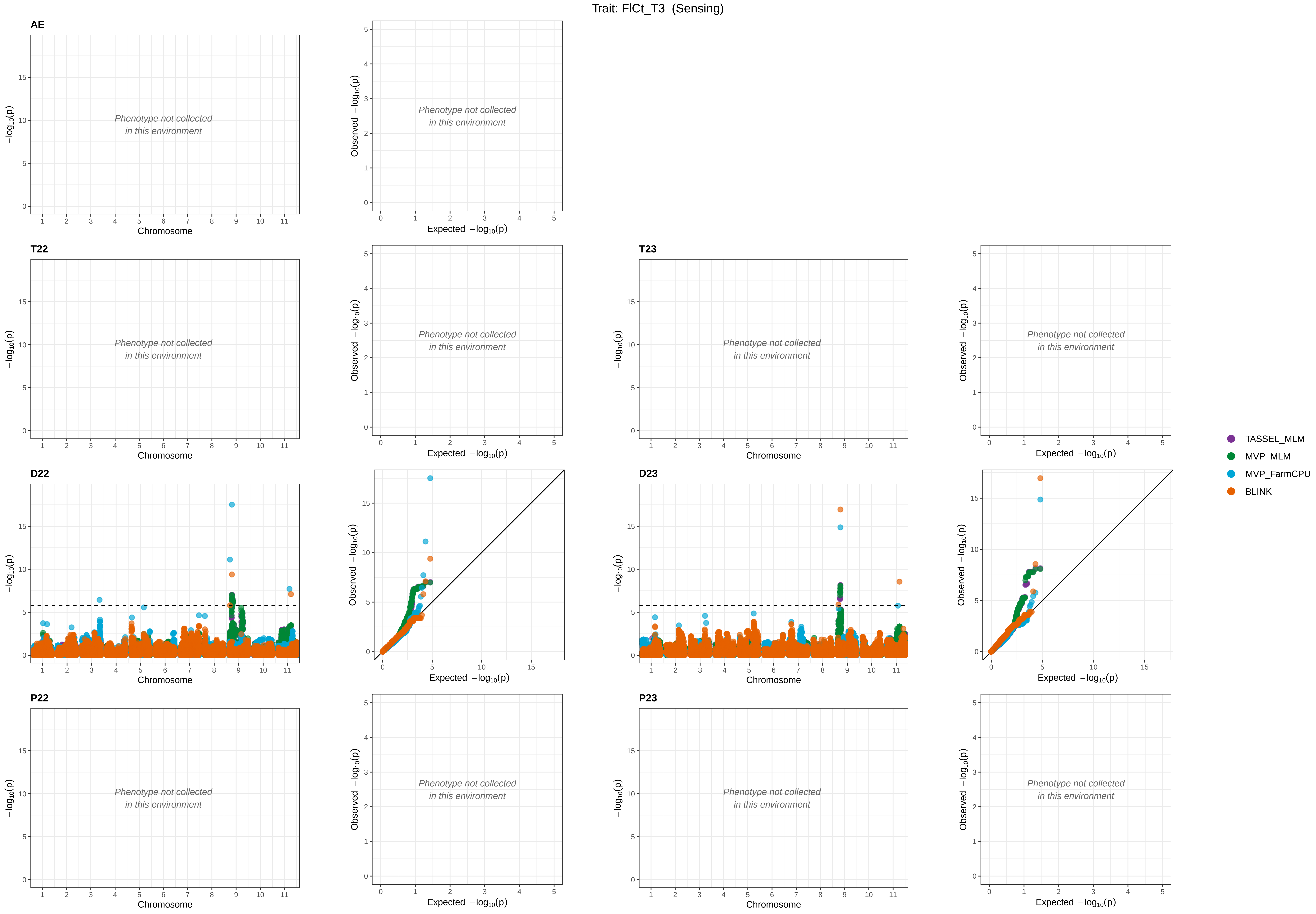

Trait: FlCt_T3 (Sensing)
AE
T22
T23
D22
D23
P22
P23
Phenotype not collected in this environment
Chromosome
Expected −log10(p)
Observed −log10(p)
−log10(p)
TASSEL_MLM
MVP_MLM
MVP_FarmCPU
BLINK

Trait: FlCt_T4 (Sensing)

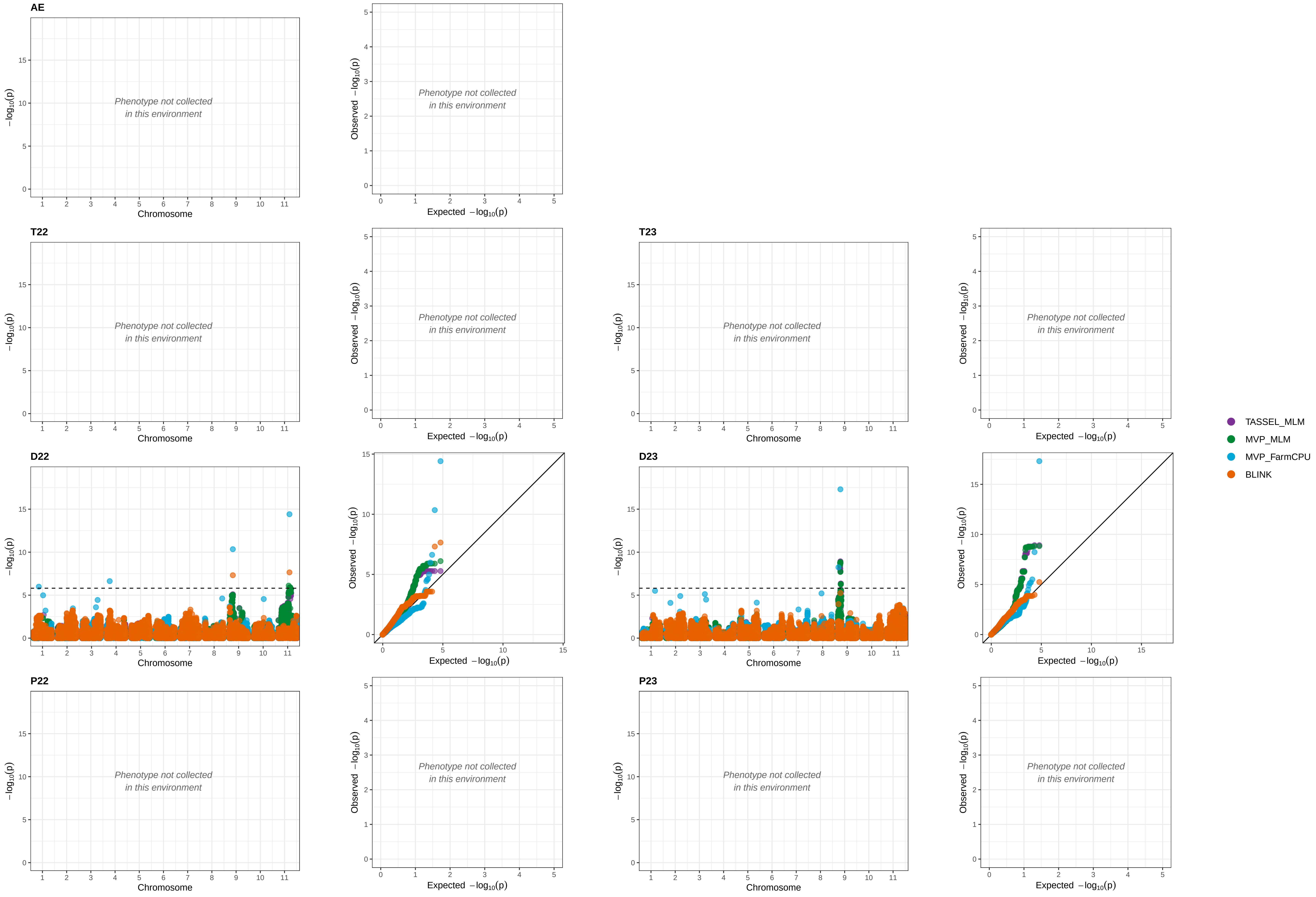

Trait: PdCt_T1  (Sensing)

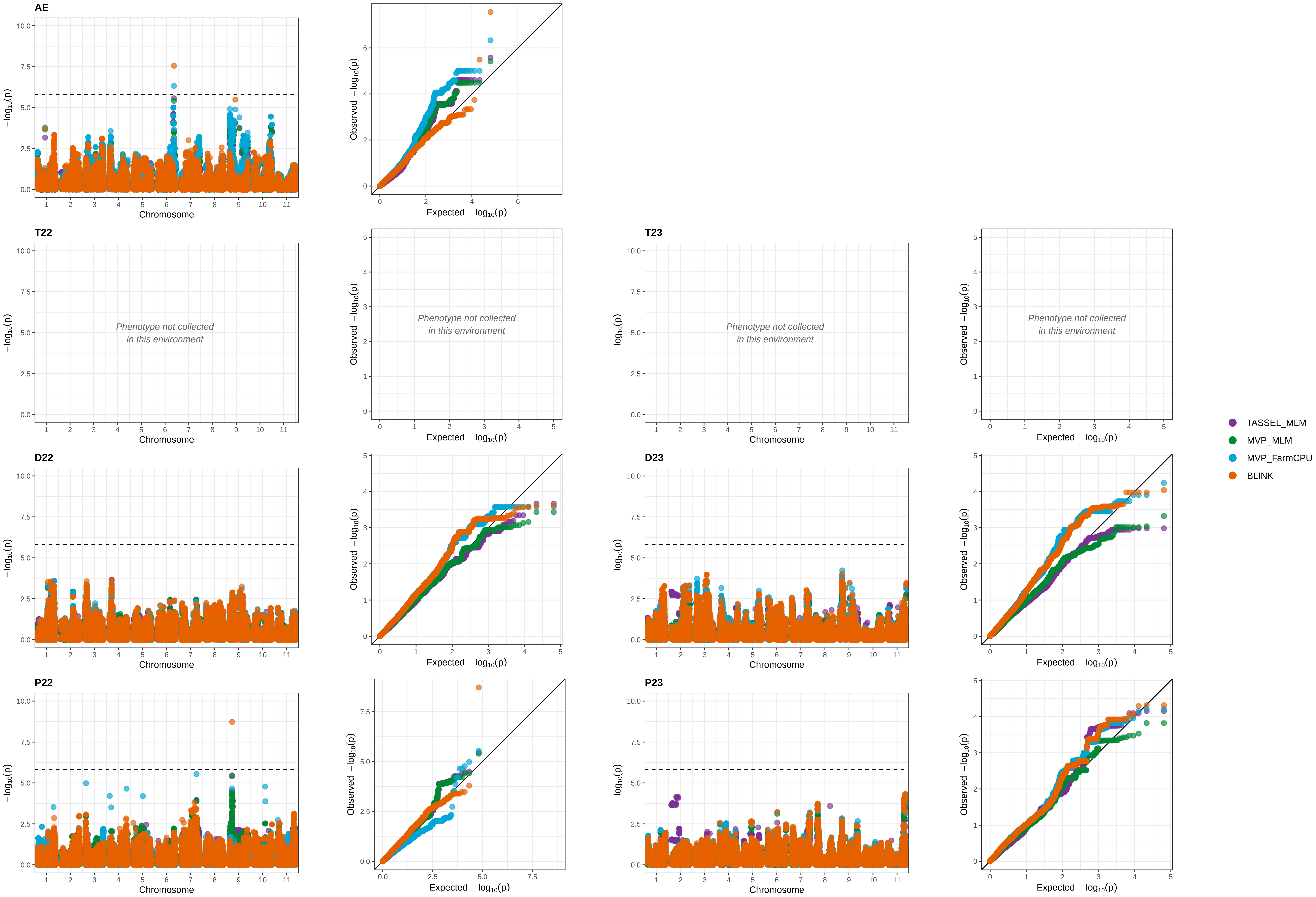

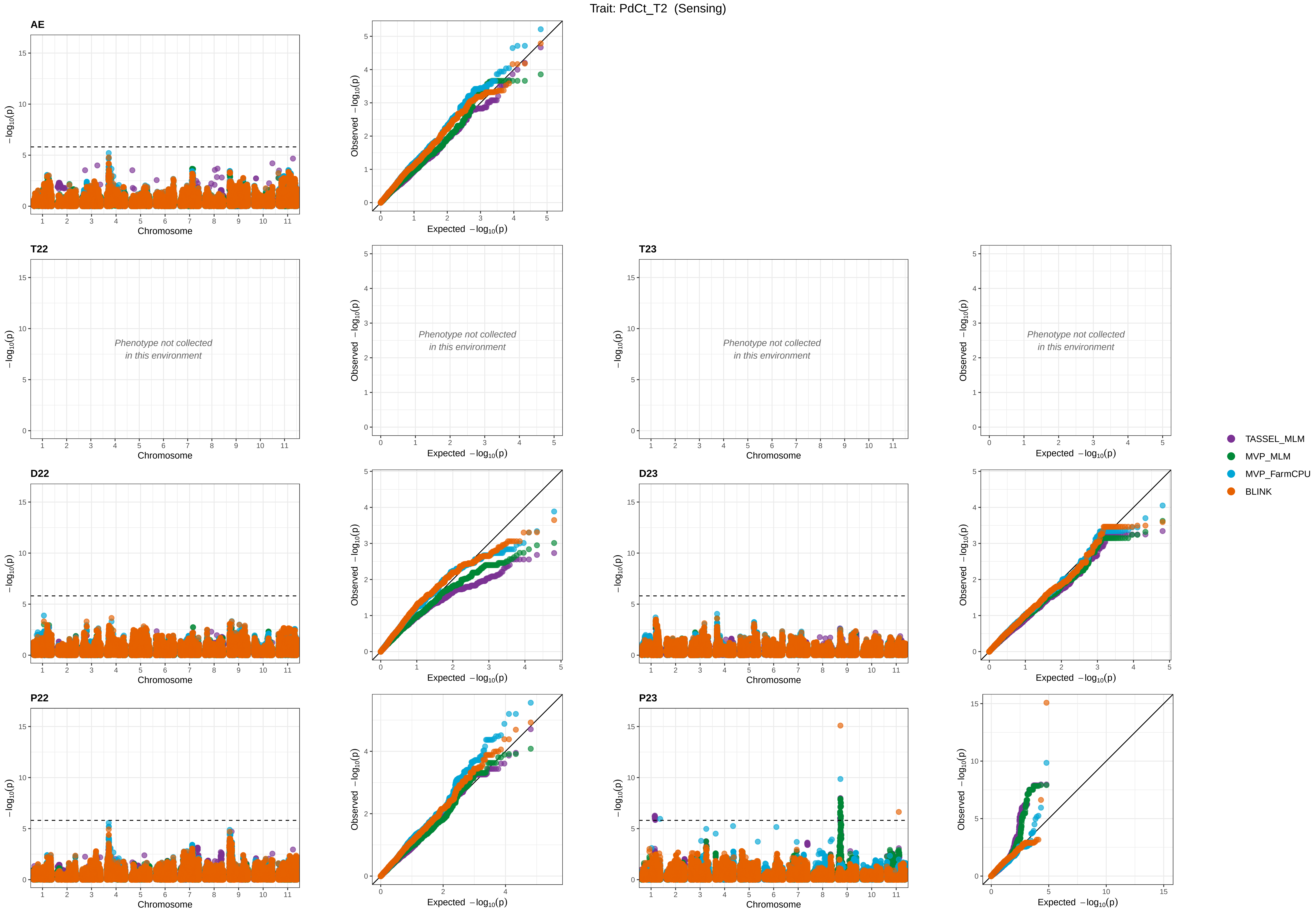

Trait: PdCt_T2 (Sensing)
AE
T22
T23
D22
D23
P22
P23
Phenotype not collected in this environment
Chromosome
Expected −log10(p)
Observed −log10(p)
−log10(p)
TASSEL_MLM
MVP_MLM
MVP_FarmCPU
BLINK

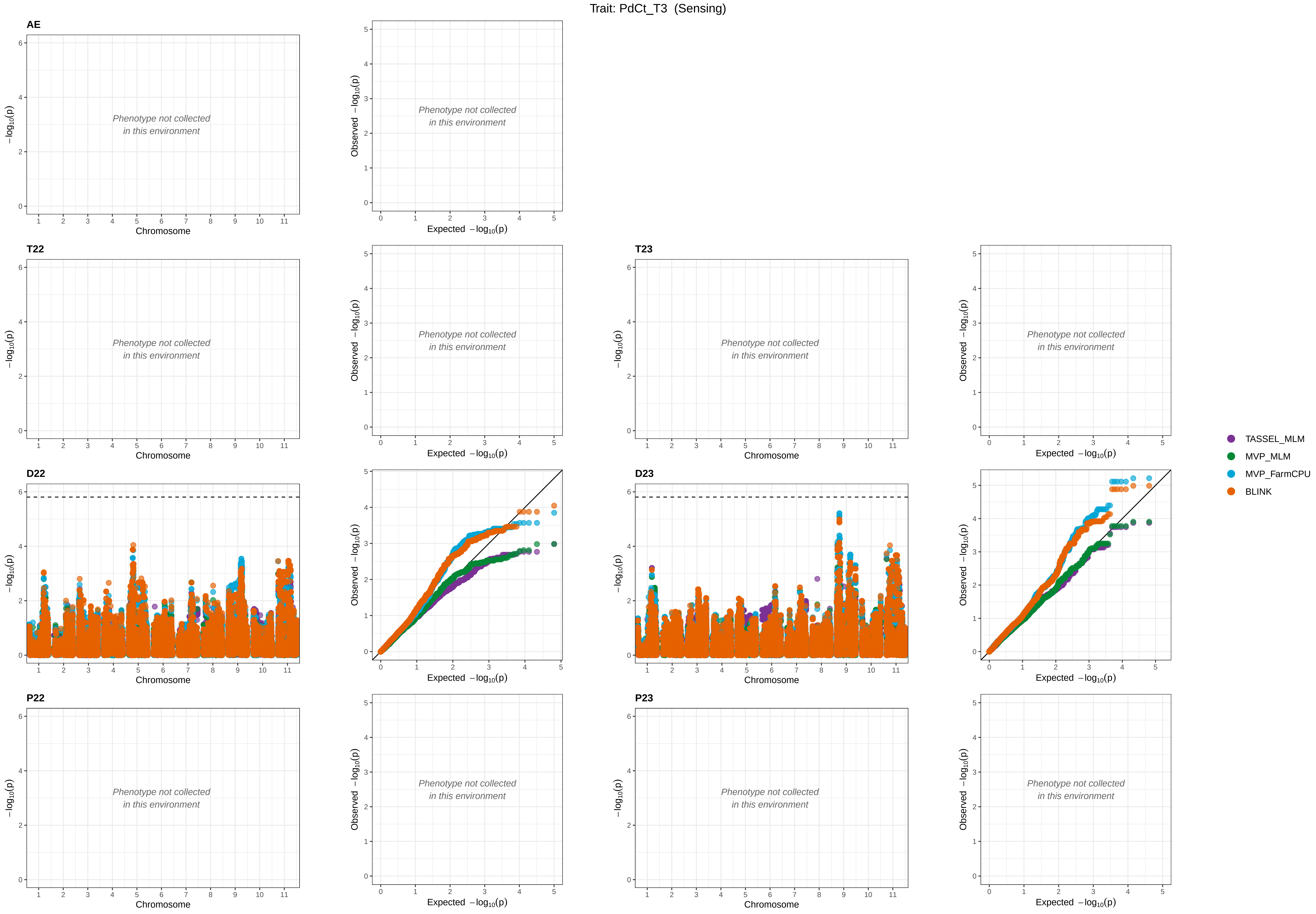
Trait: PdCt_T3 (Sensing)
AE
T22
T23
D22
D23
P22
P23
Phenotype not collected in this environment
Chromosome
Expected −log10(p)
Observed −log10(p)
−log10(p)
TASSEL_MLM
MVP_MLM
MVP_FarmCPU
BLINK

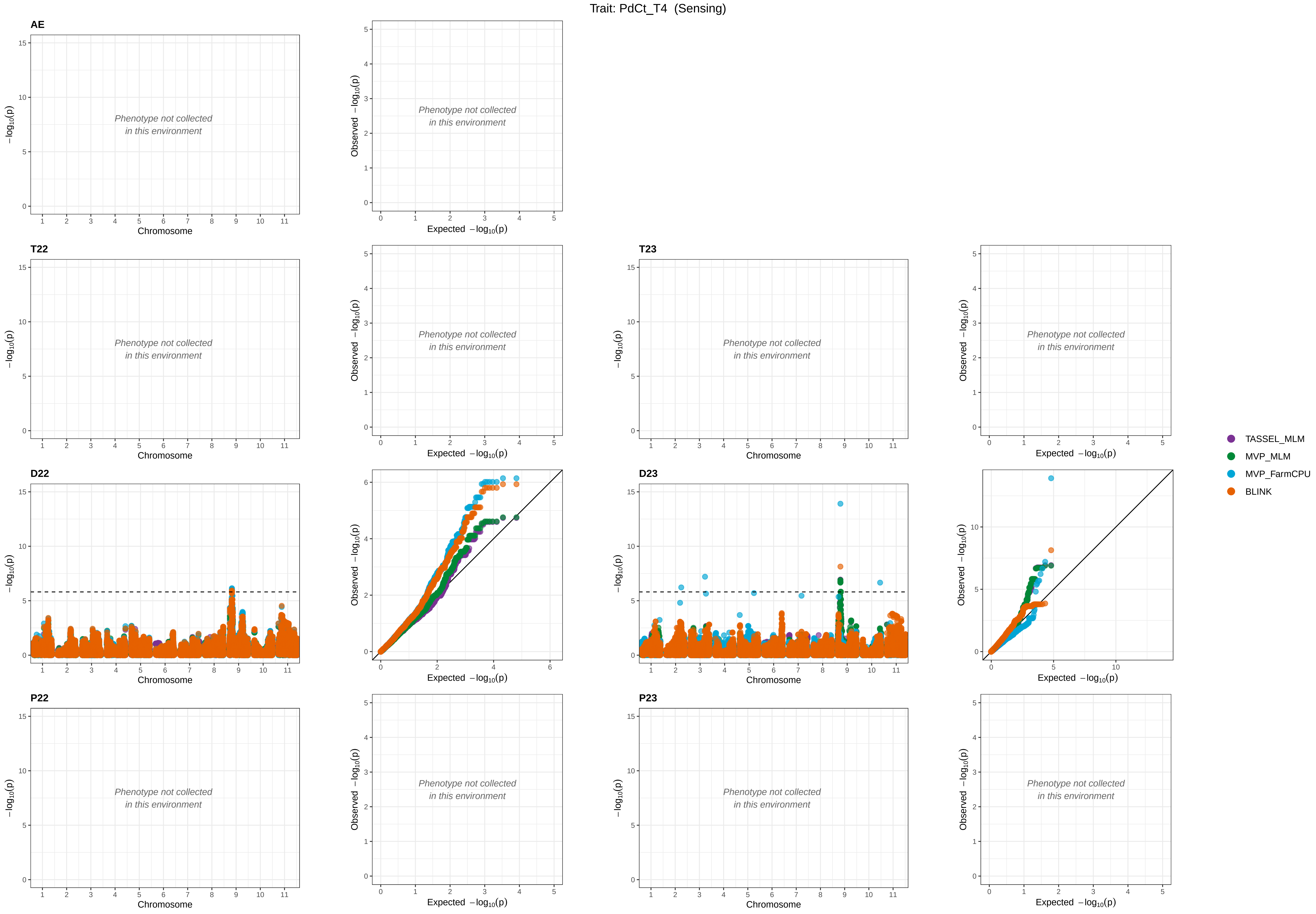

Trait: PdCt_T4 (Sensing)
AE
T22
T23
D22
D23
P22
P23
Phenotype not collected
in this environment
Chromosome
Expected −log10(p)
Observed −log10(p)
−log10(p)
TASSEL_MLM
MVP_MLM
MVP_FarmCPU
BLINK

Trait: PtHt_T1  (Sensing)

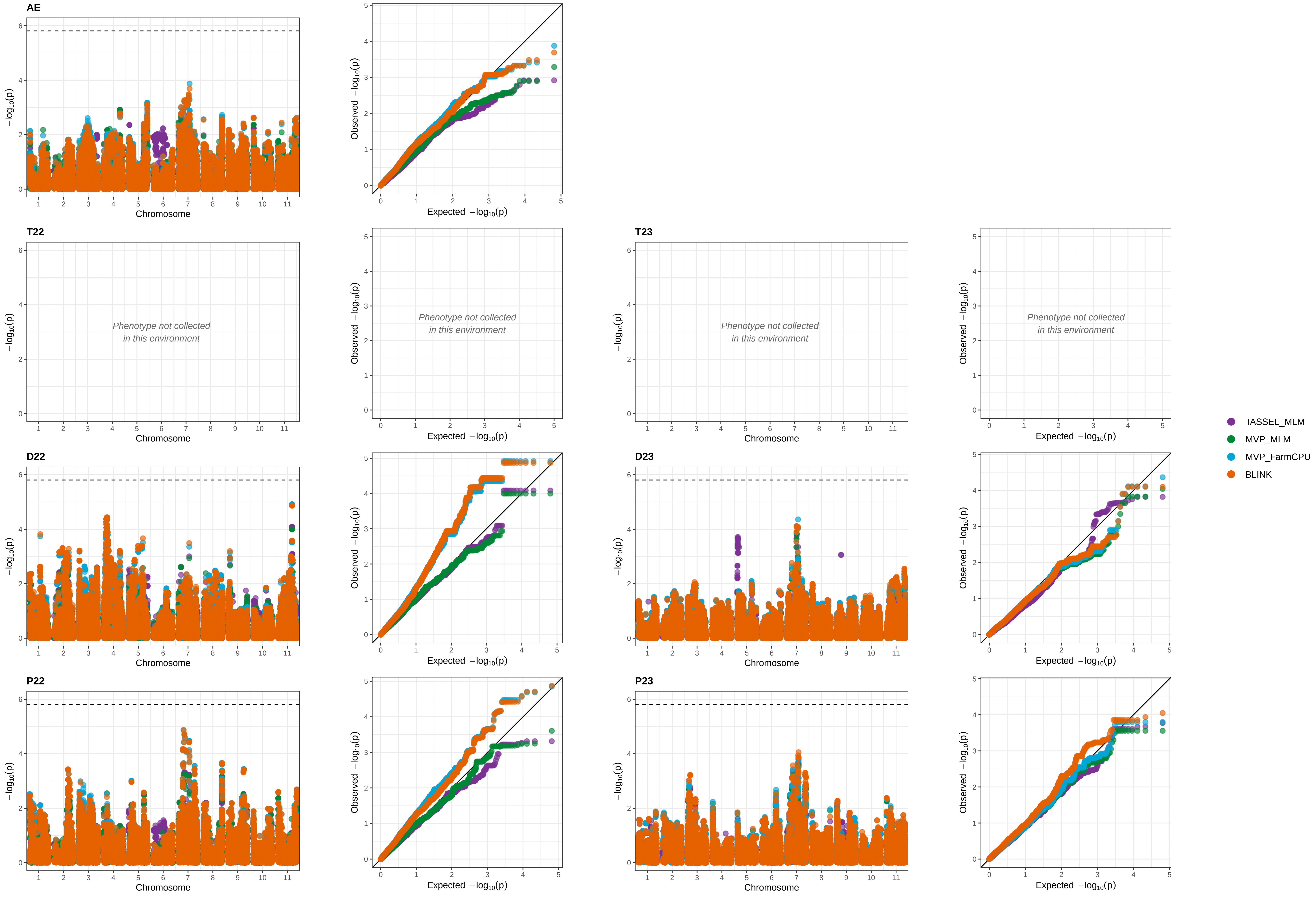

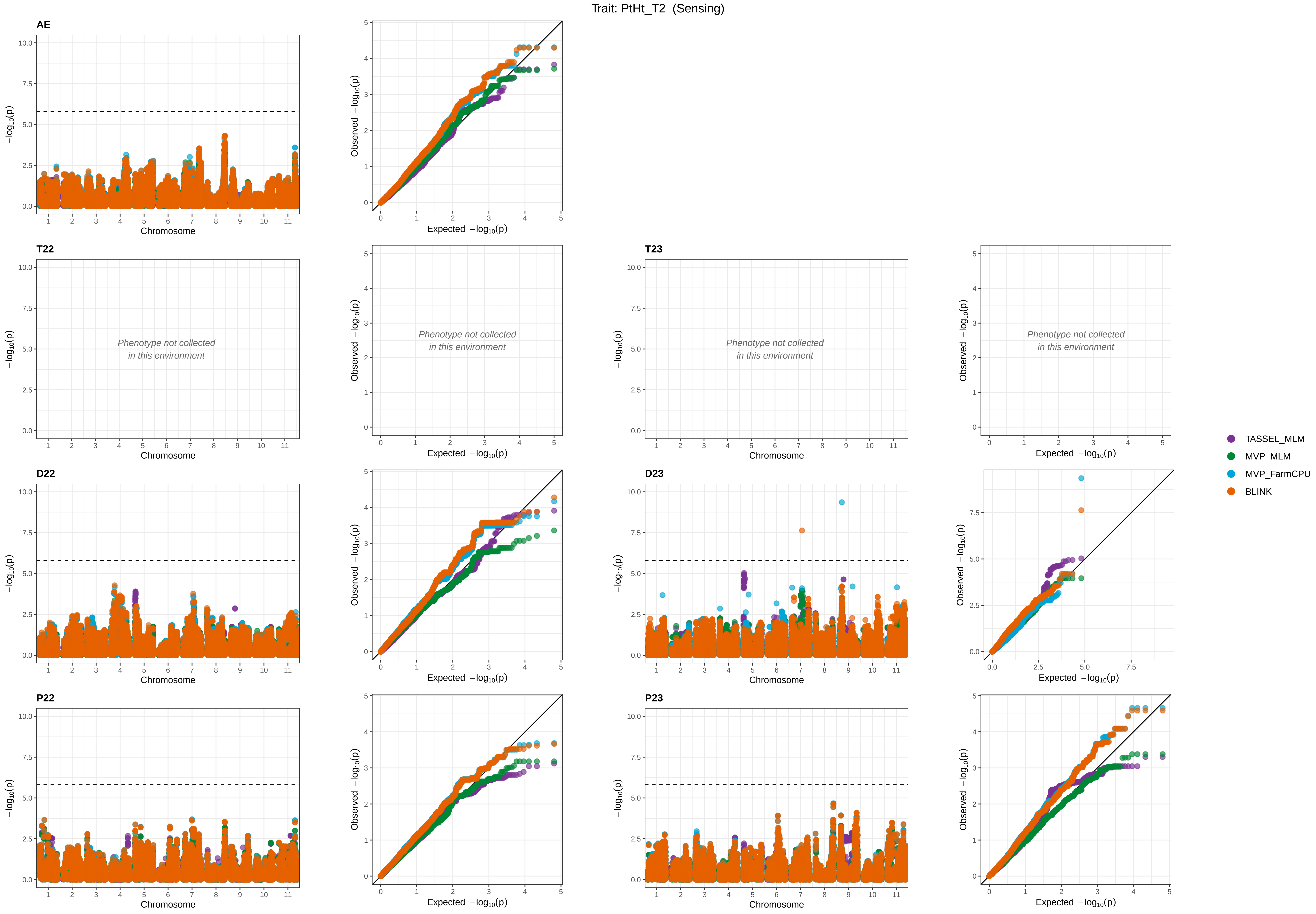

Trait: PtHt_T2 (Sensing)
AE
T22
T23
D22
D23
P22
P23
Phenotype not collected in this environment
Chromosome
Expected −log10(p)
Observed −log10(p)
−log10(p)
TASSEL_MLM
MVP_MLM
MVP_FarmCPU
BLINK

Trait: PtHt_T3  (Sensing)

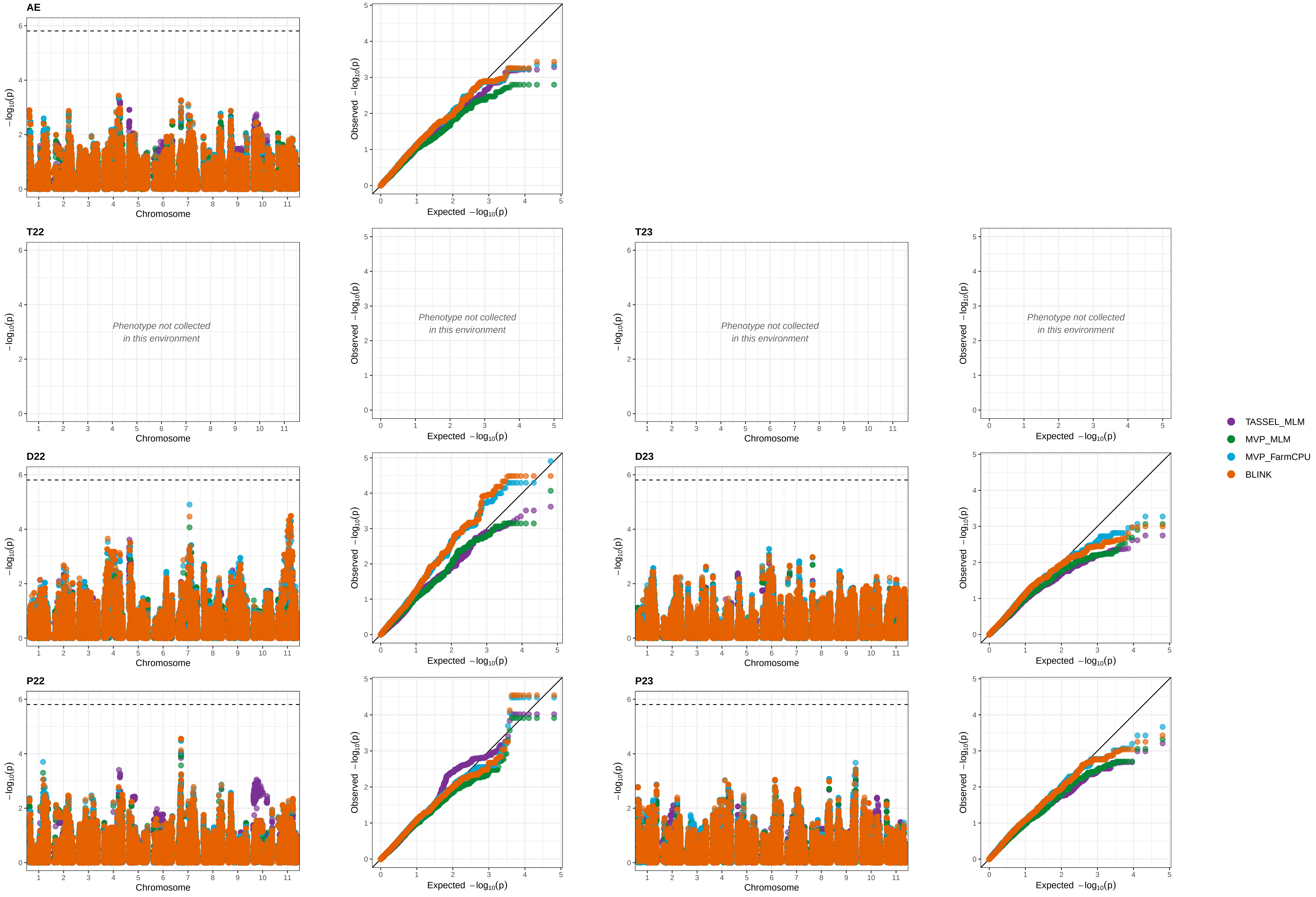

Trait: PtHt_T4 (Sensing)

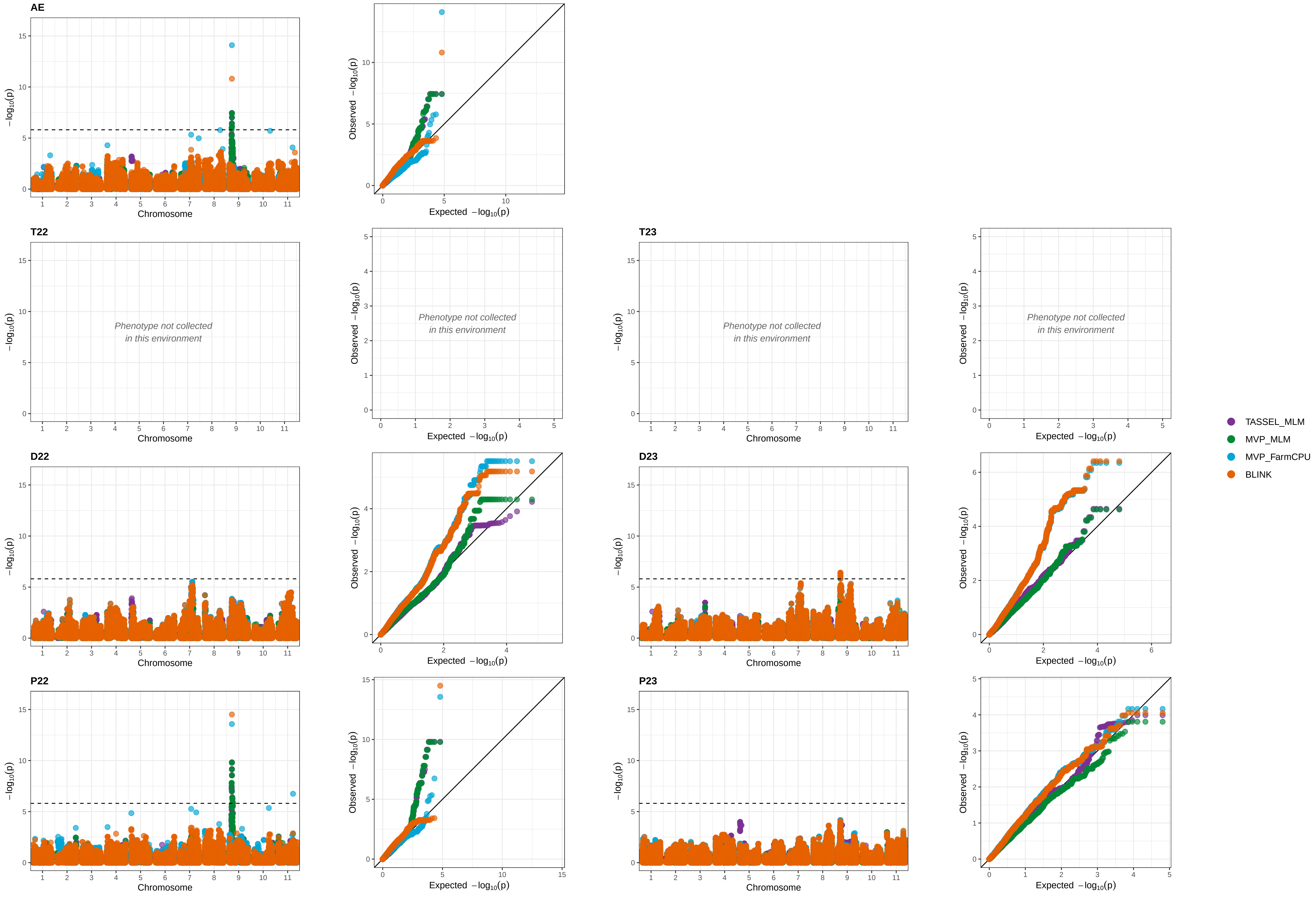

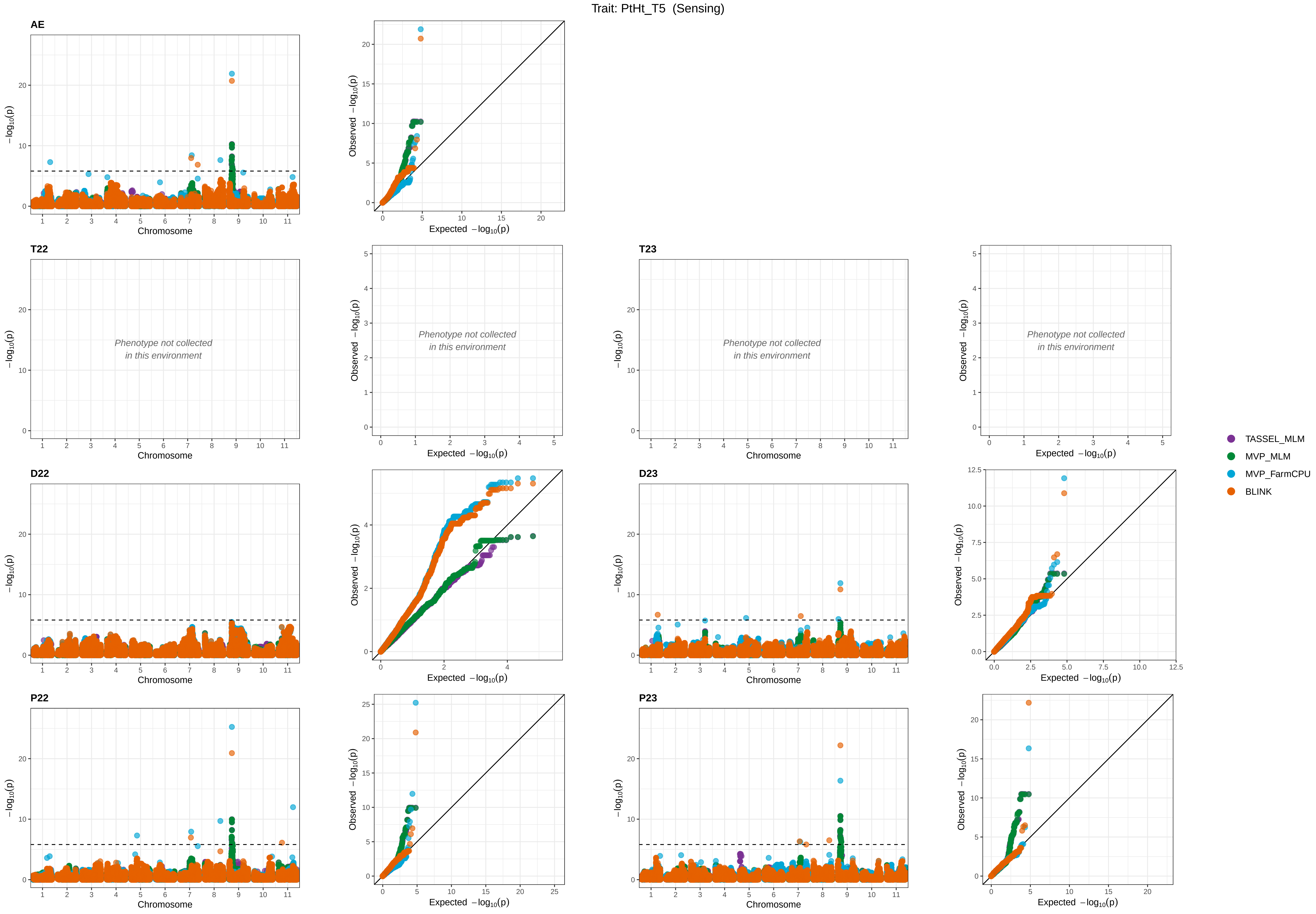

Trait: PtHt_T5 (Sensing)
AE
T22
T23
D22
D23
P22
P23
Chromosome
Expected −log10(p)
Observed −log10(p)
−log10(p)
Phenotype not collected
in this environment
TASSEL_MLM
MVP_MLM
MVP_FarmCPU
BLINK

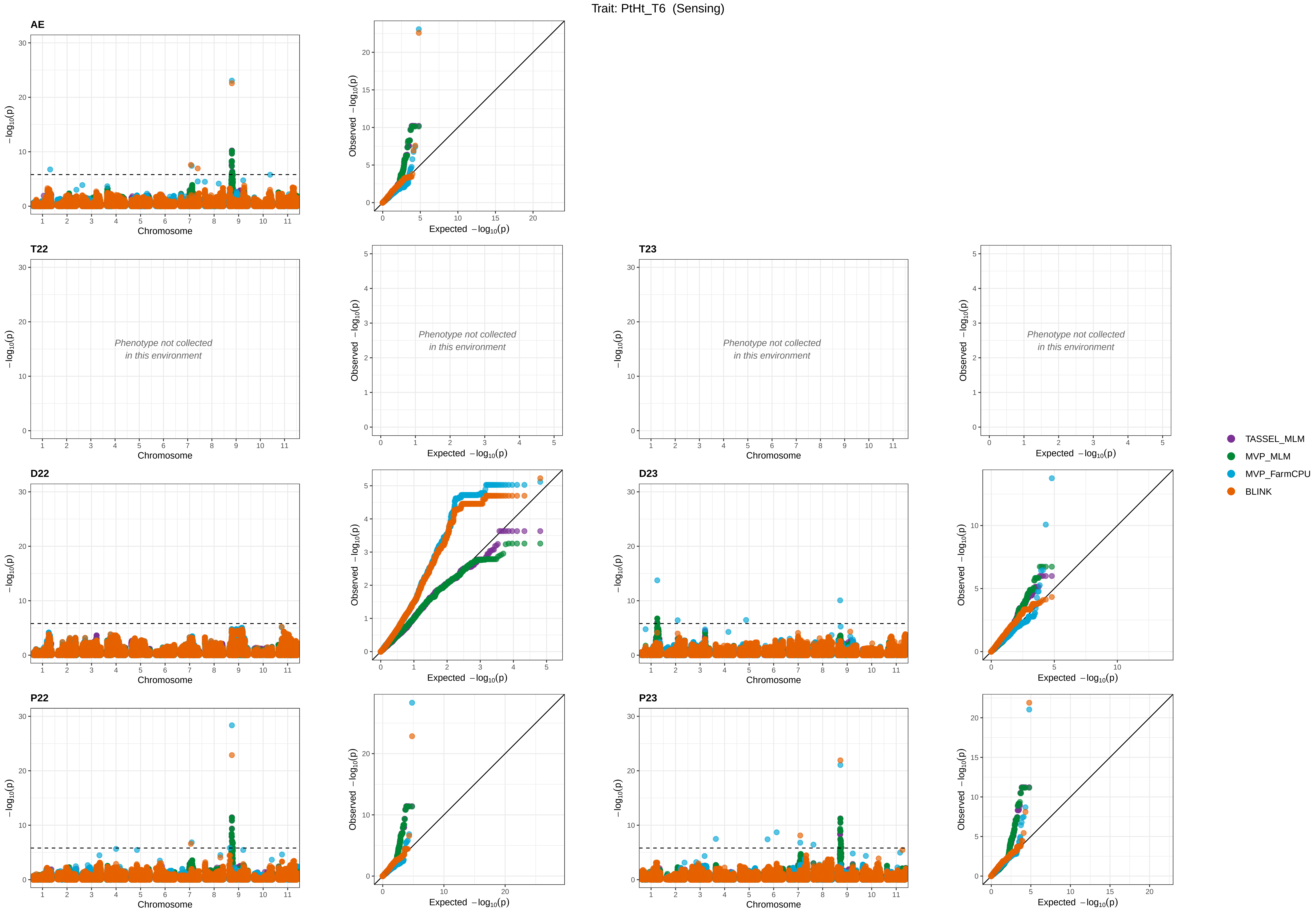
Trait: PtHt_T6 (Sensing)
AE
T22
T23
D22
D23
P22
P23
Phenotype not collected
in this environment
−log10(p)
Chromosome
Observed −log10(p)
Expected −log10(p)
TASSEL_MLM
MVP_MLM
MVP_FarmCPU
BLINK

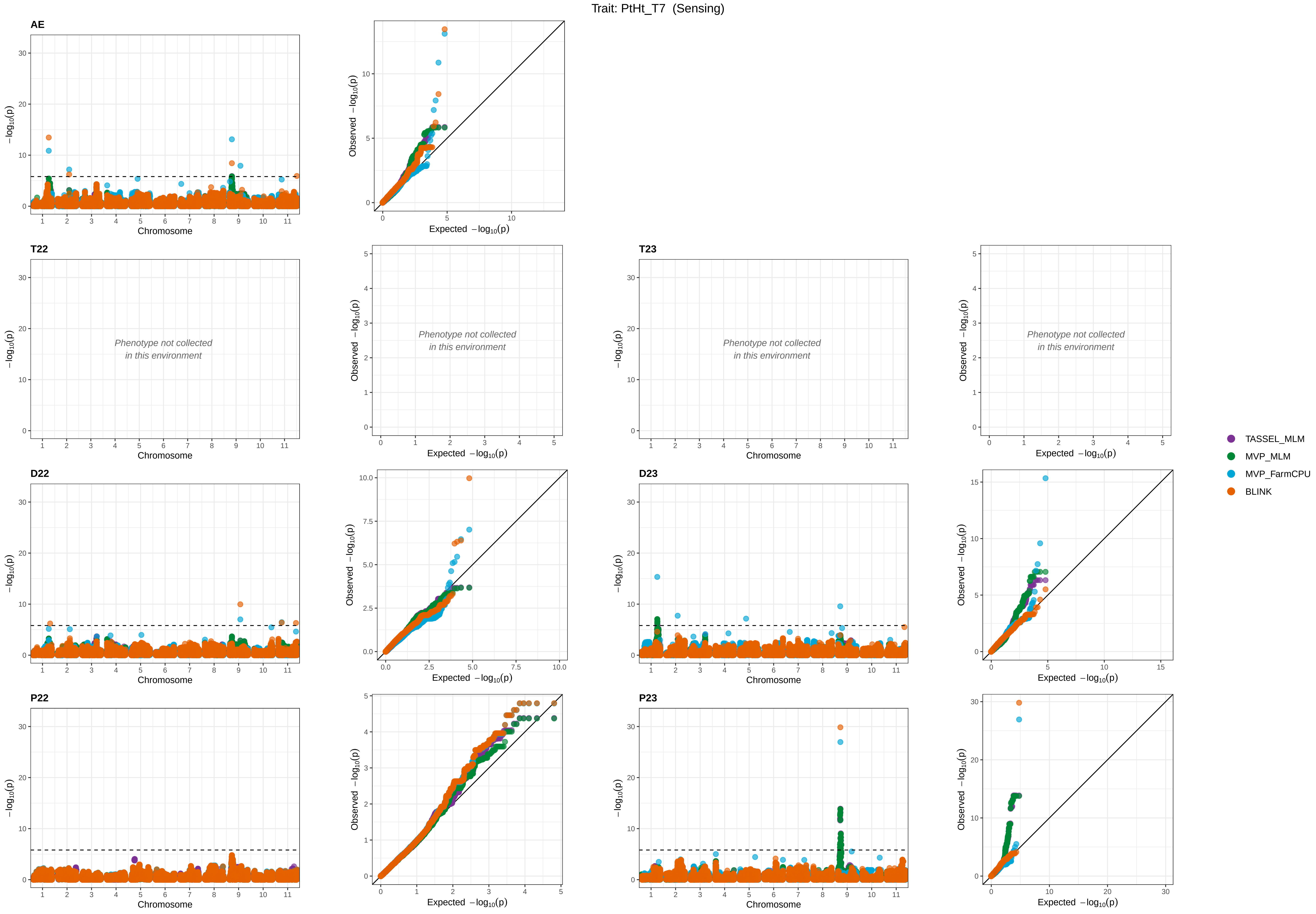

Trait: PtHt_T7 (Sensing)
AE
T22
T23
D22
D23
P22
P23
Phenotype not collected in this environment
−log10(p)
Chromosome
Observed −log10(p)
Expected −log10(p)
TASSEL_MLM
MVP_MLM
MVP_FarmCPU
BLINK

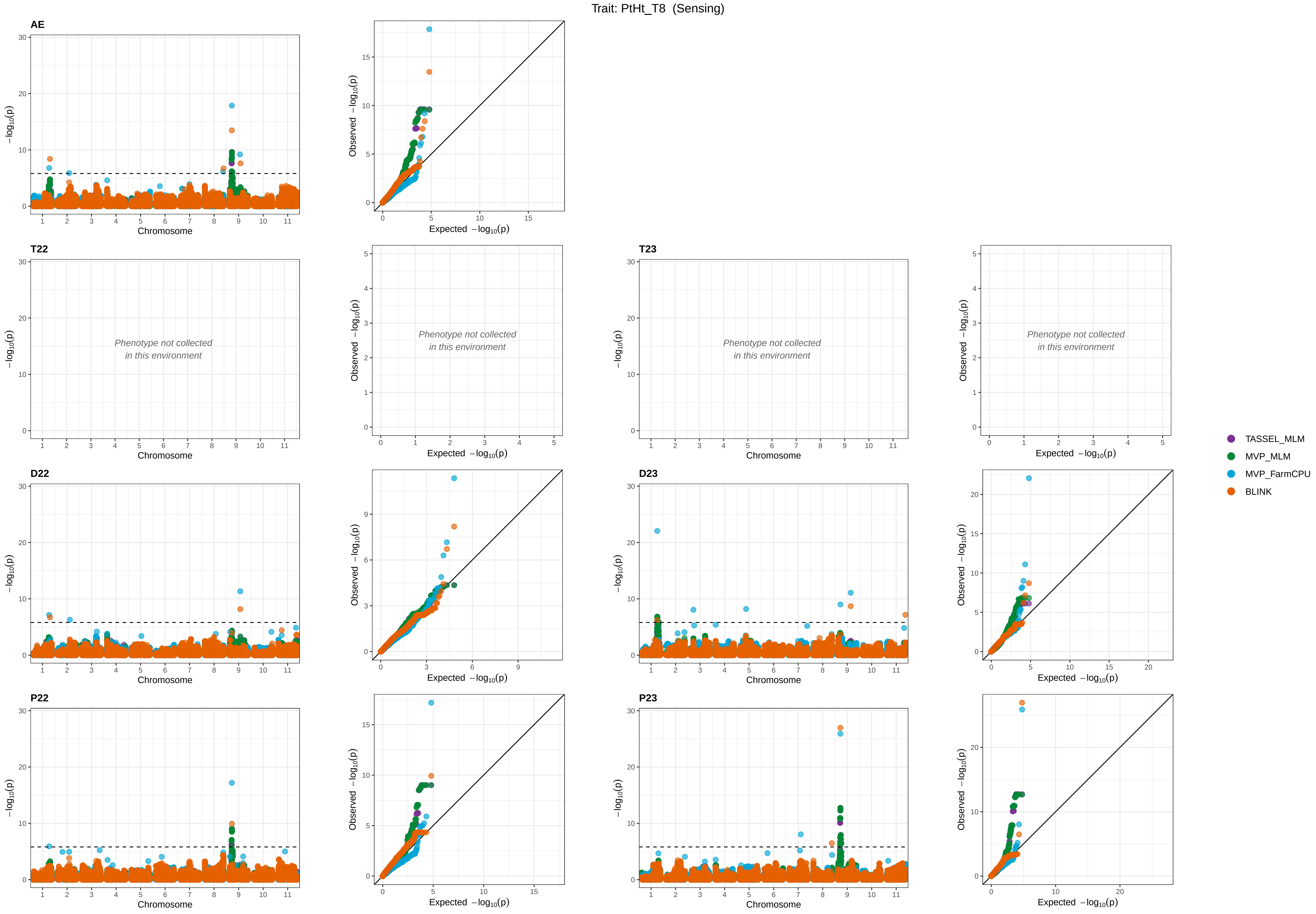

Trait: PtHt_T8 (Sensing)
AE
T22
T23
D22
D23
P22
P23
Phenotype not collected in this environment
Chromosome
−log10(p)
Observed −log10(p)
Expected −log10(p)
TASSEL_MLM
MVP_MLM
MVP_FarmCPU
BLINK

Trait: PtHt_T9 (Sensing)

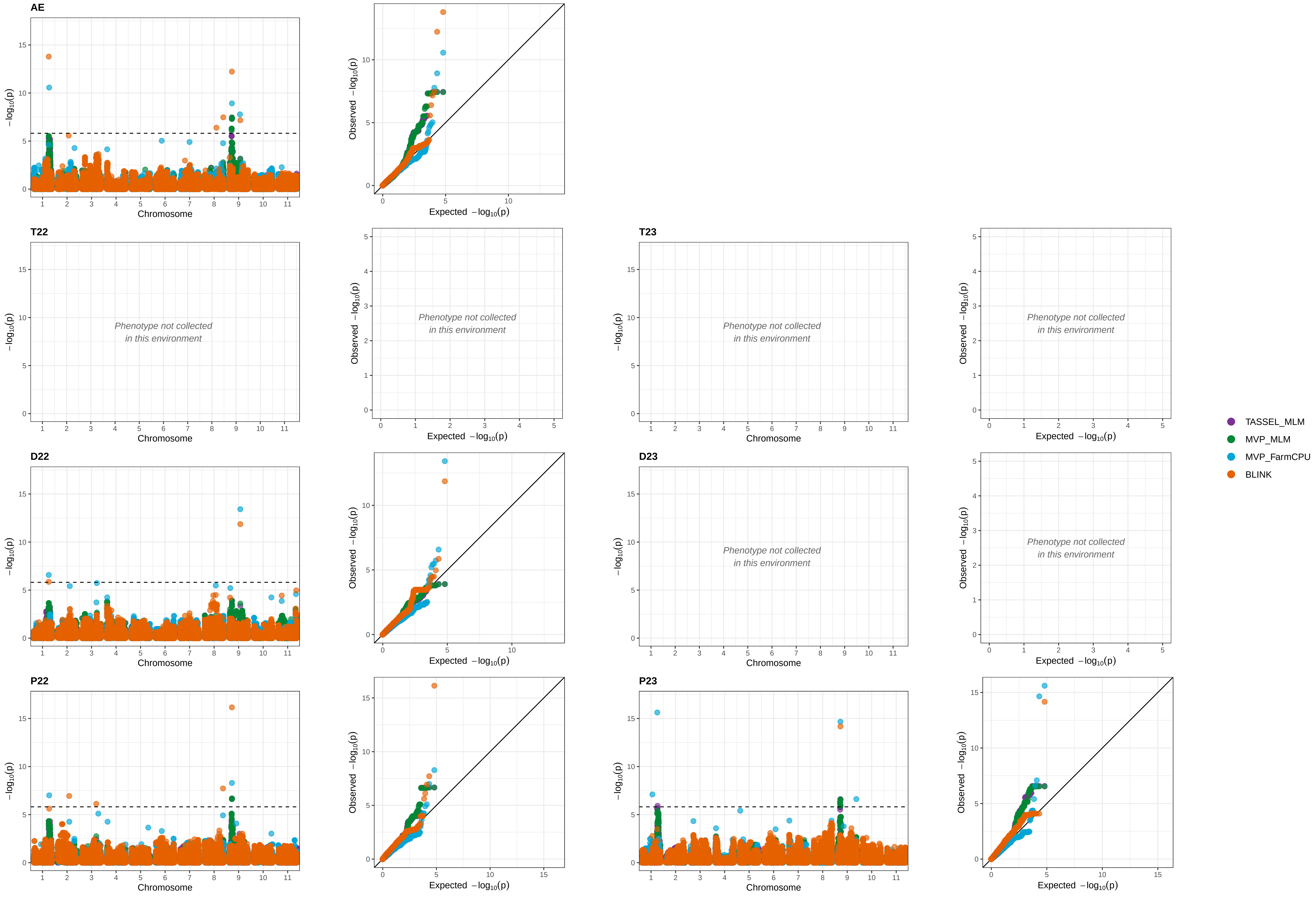

Trait: PtHt_T10  (Sensing)

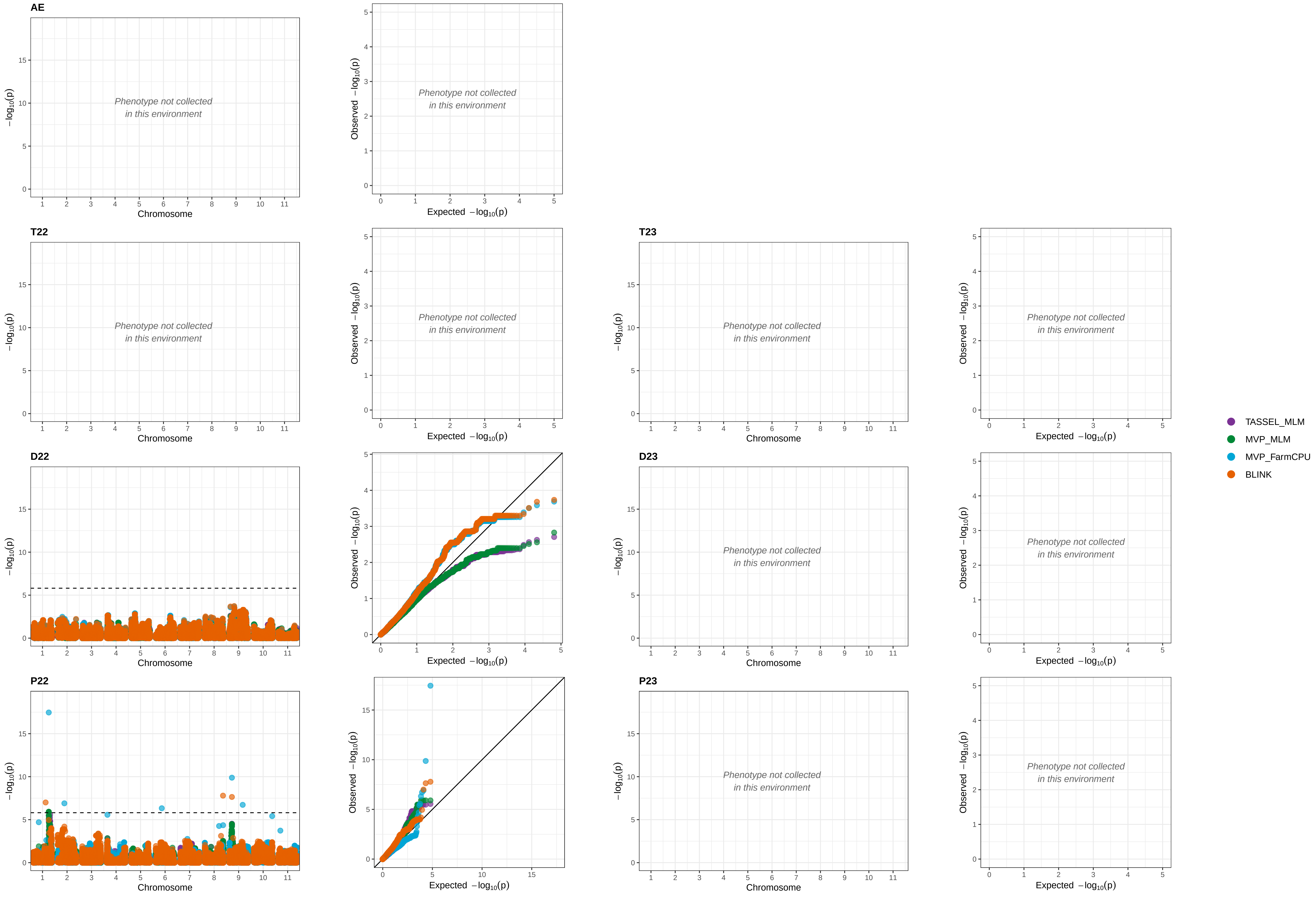

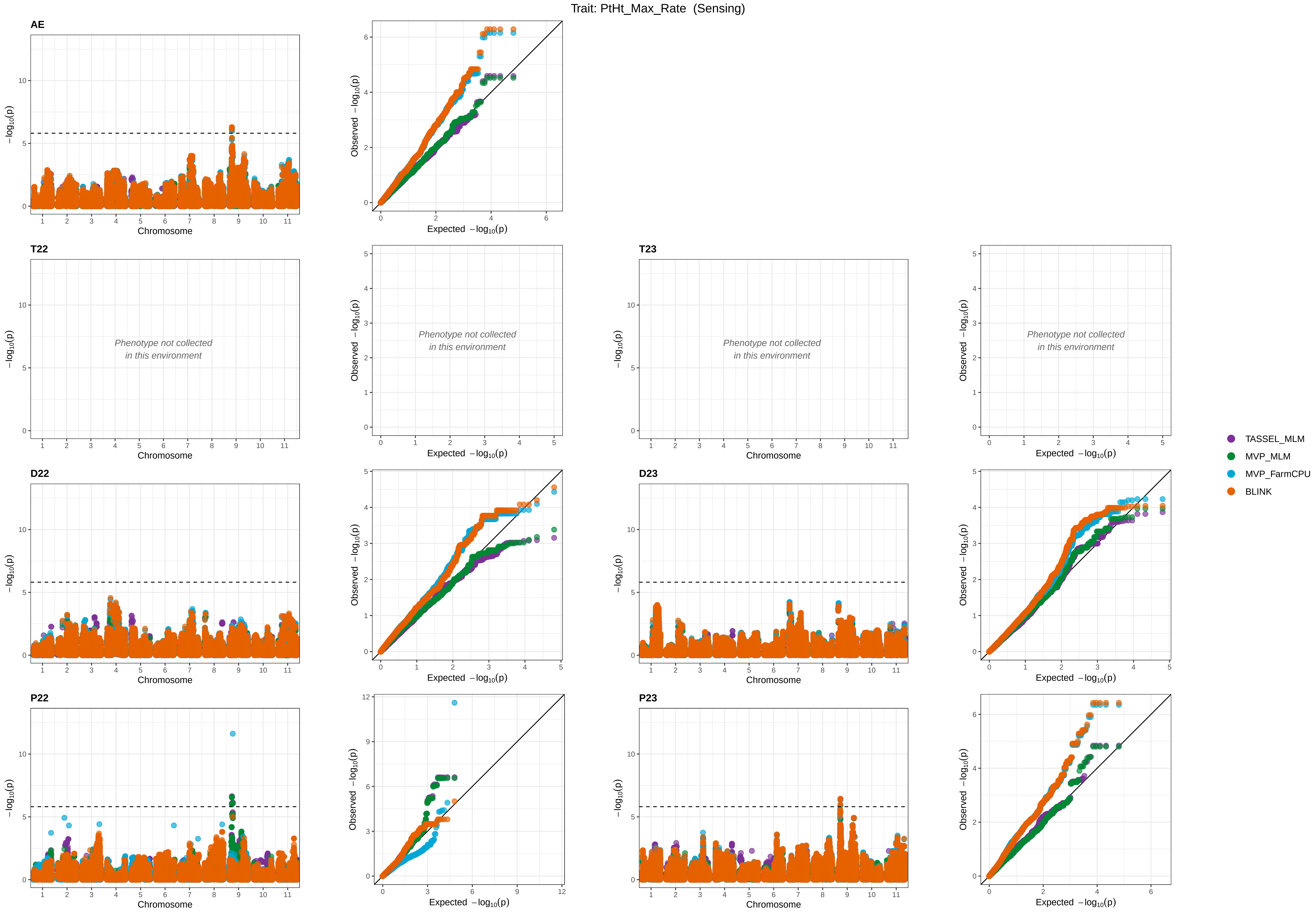

Trait: PtHt_Max_Rate (Sensing)
AE
T22
T23
D22
D23
P22
P23
−log10(p)
Observed −log10(p)
Expected −log10(p)
Chromosome
Phenotype not collected in this environment
TASSEL_MLM
MVP_MLM
MVP_FarmCPU
BLINK

Trait: PtHt_Max_Rate_DAP (Sensing)

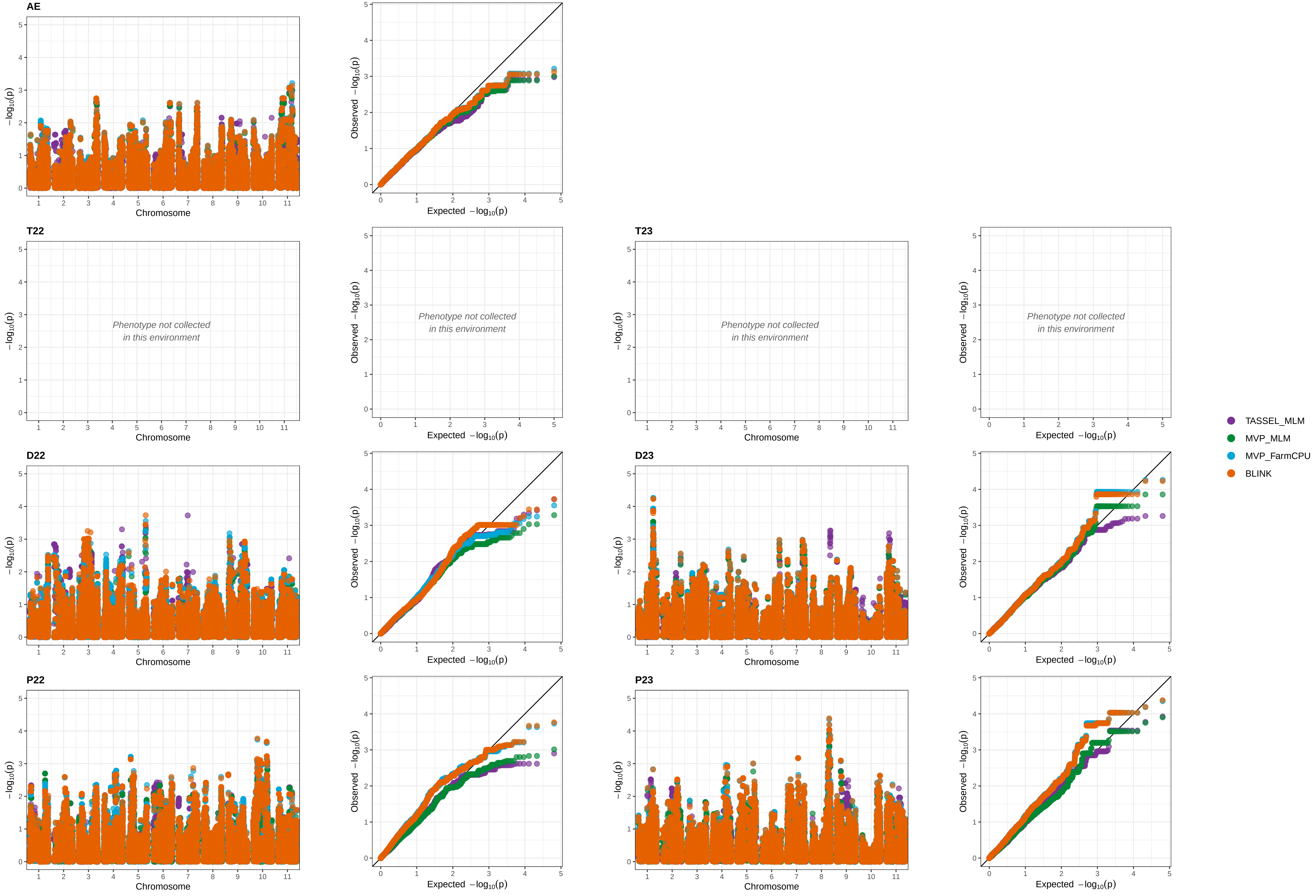

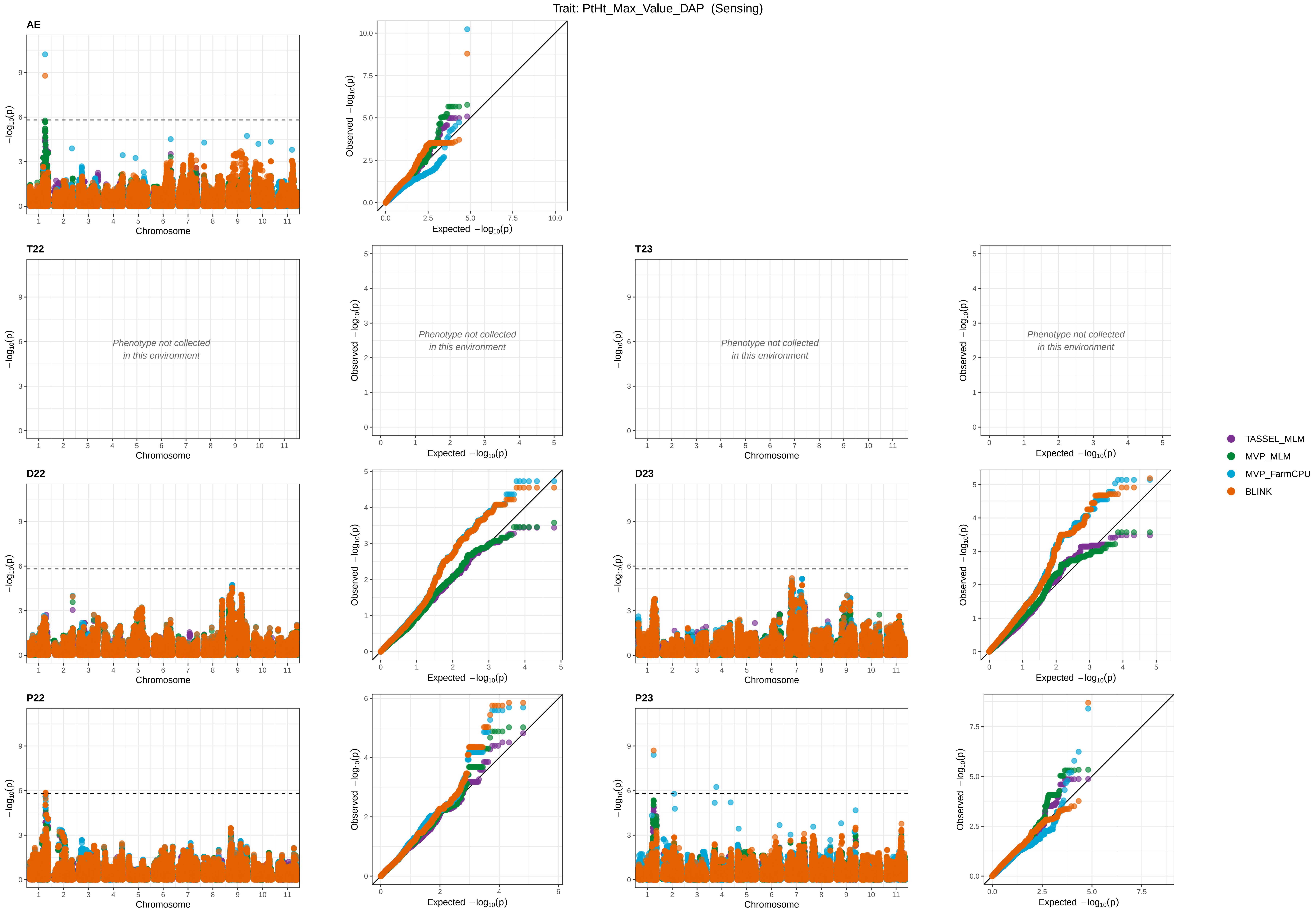

Trait: PtHt_Max_Value_DAP (Sensing)
AE
T22
T23
D22
D23
P22
P23
Phenotype not collected
in this environment
Chromosome
Expected −log10(p)
Observed −log10(p)
−log10(p)
TASSEL_MLM
MVP_MLM
MVP_FarmCPU
BLINK

Trait: PtHt_Min_Rate (Sensing)

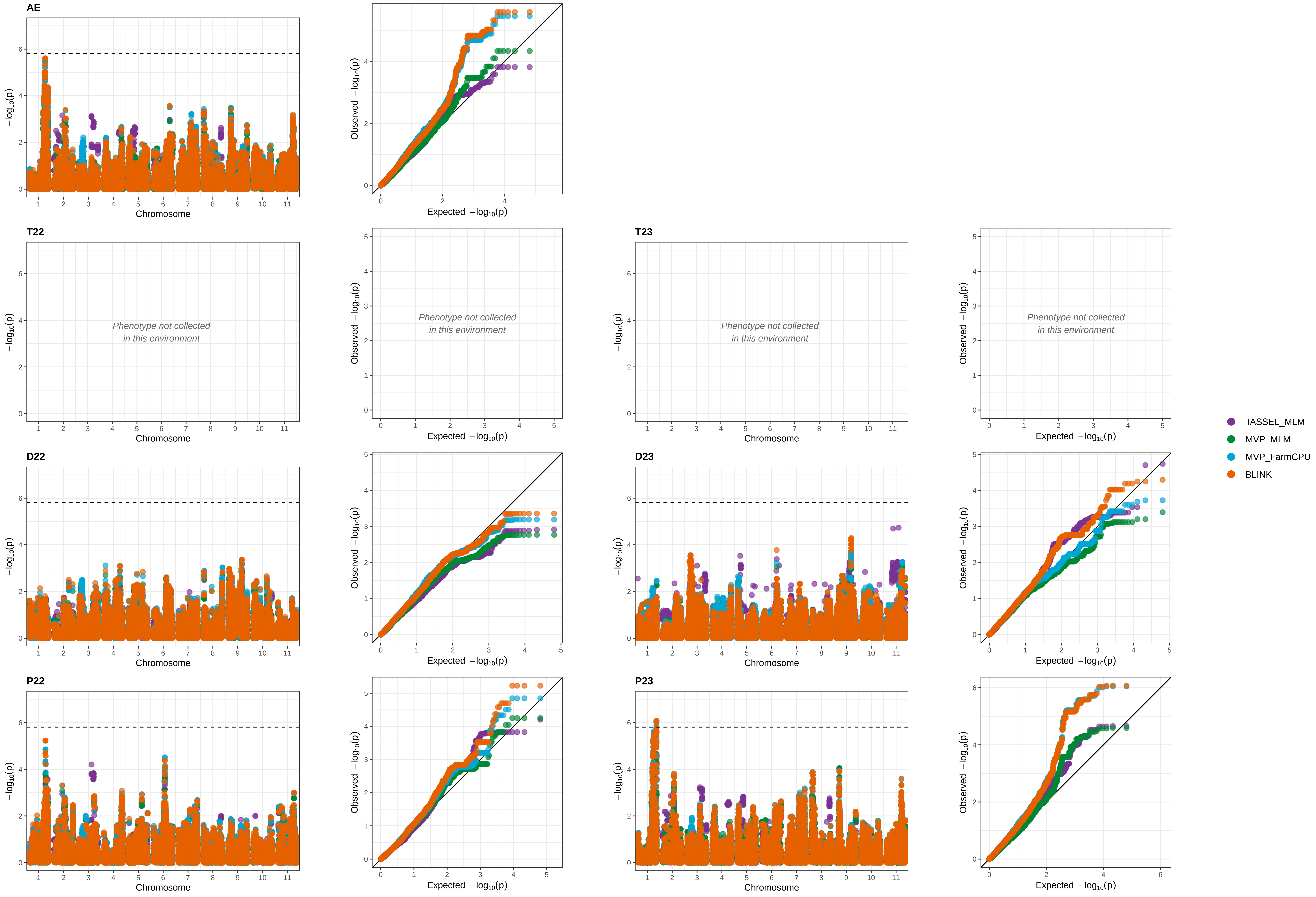

Trait: PtHt_Min_Rate_DAP (Sensing)

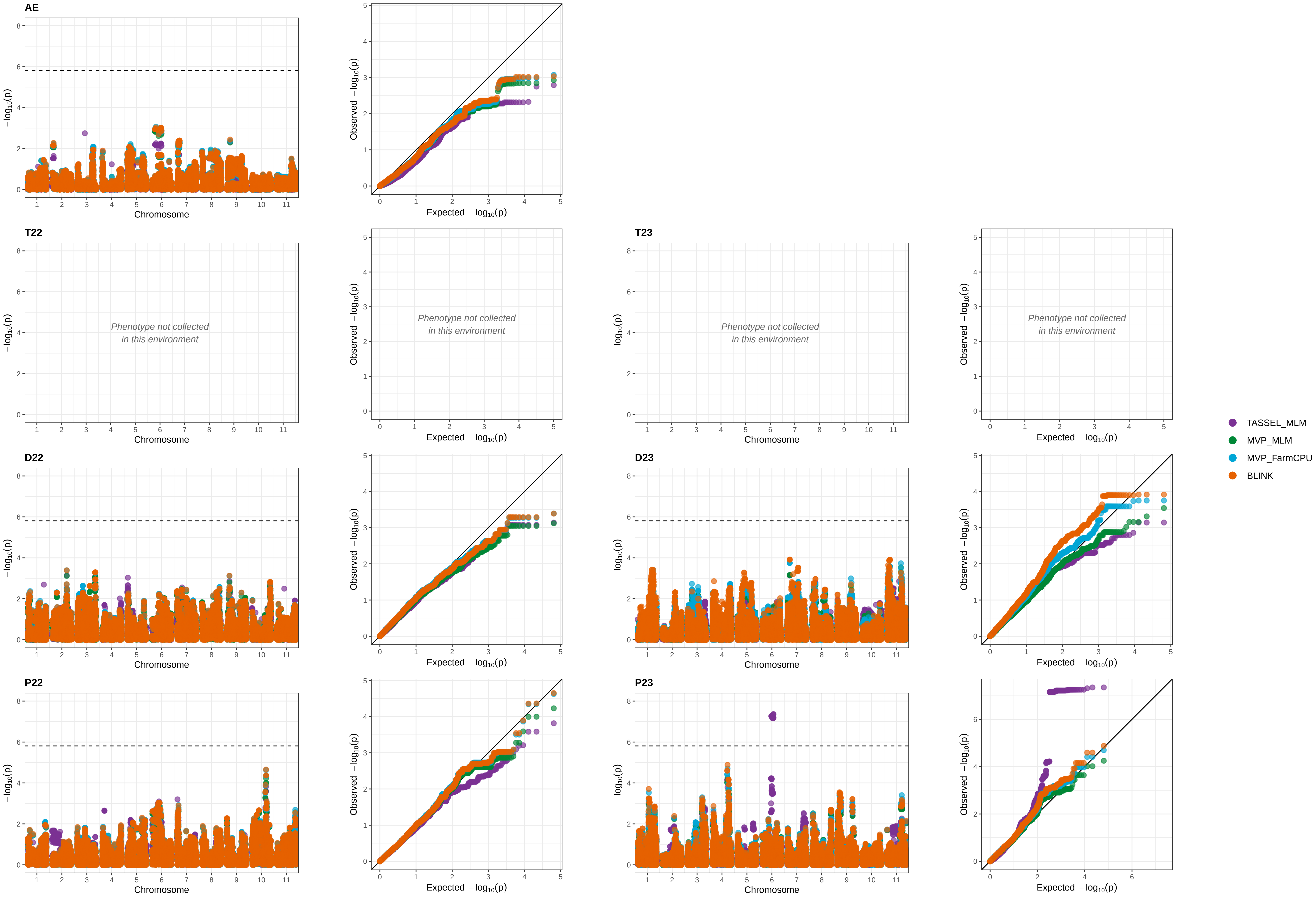

Trait: VgFr_T1 (Sensing)

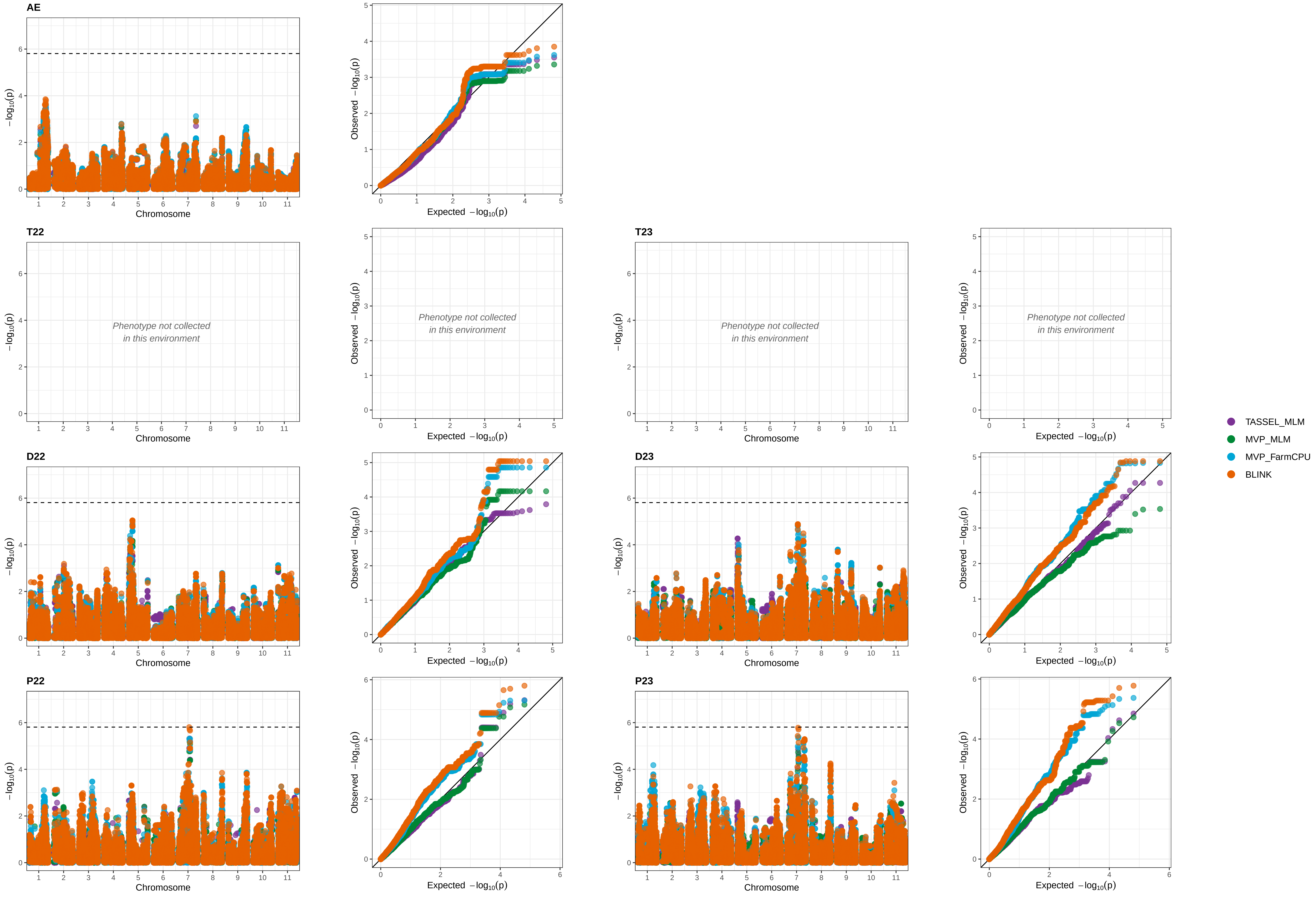

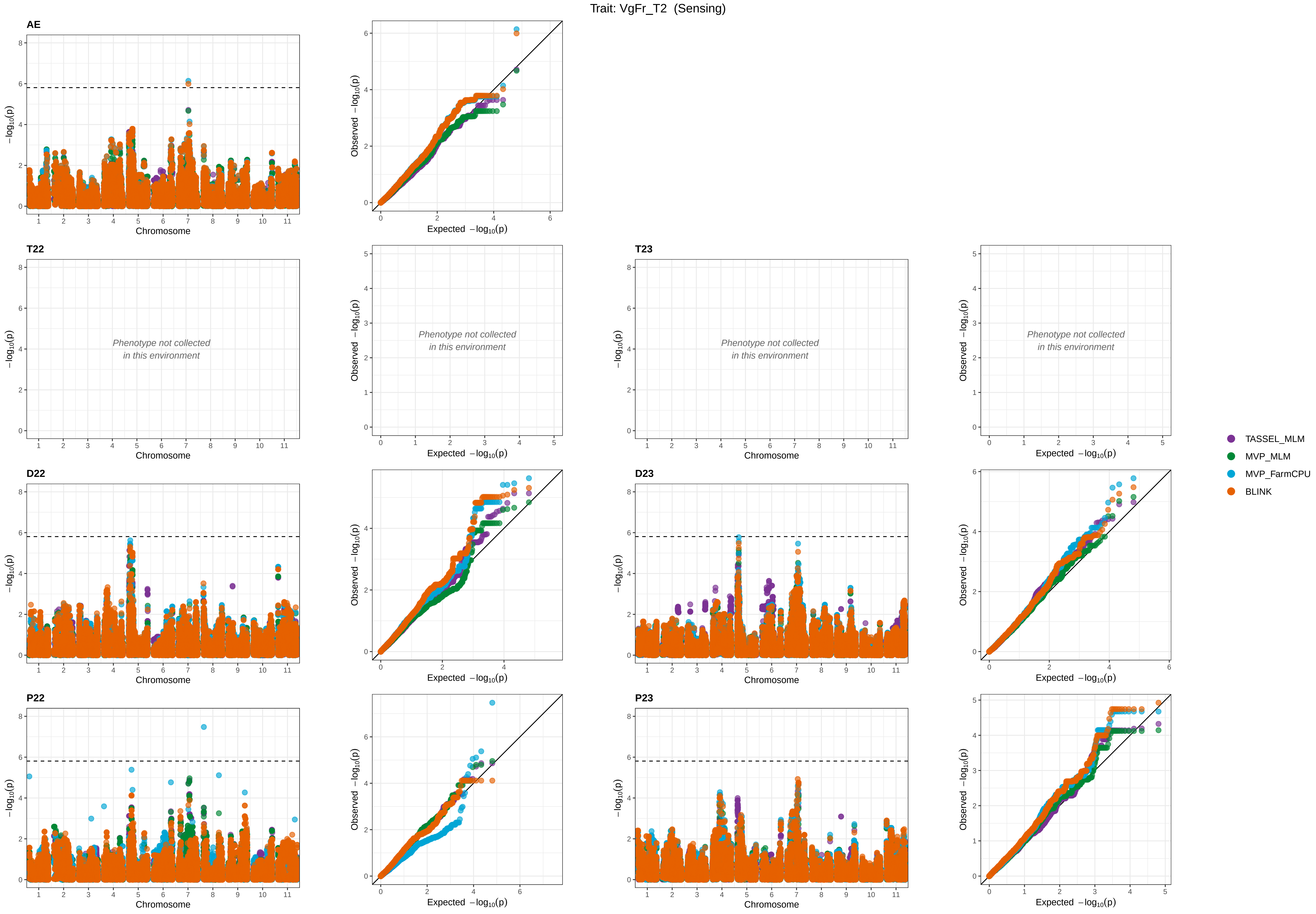

Trait: VgFr_T2 (Sensing)
AE
T22
T23
D22
D23
P22
P23
Phenotype not collected
in this environment
Chromosome
−log10(p)
Expected −log10(p)
Observed −log10(p)
TASSEL_MLM
MVP_MLM
MVP_FarmCPU
BLINK

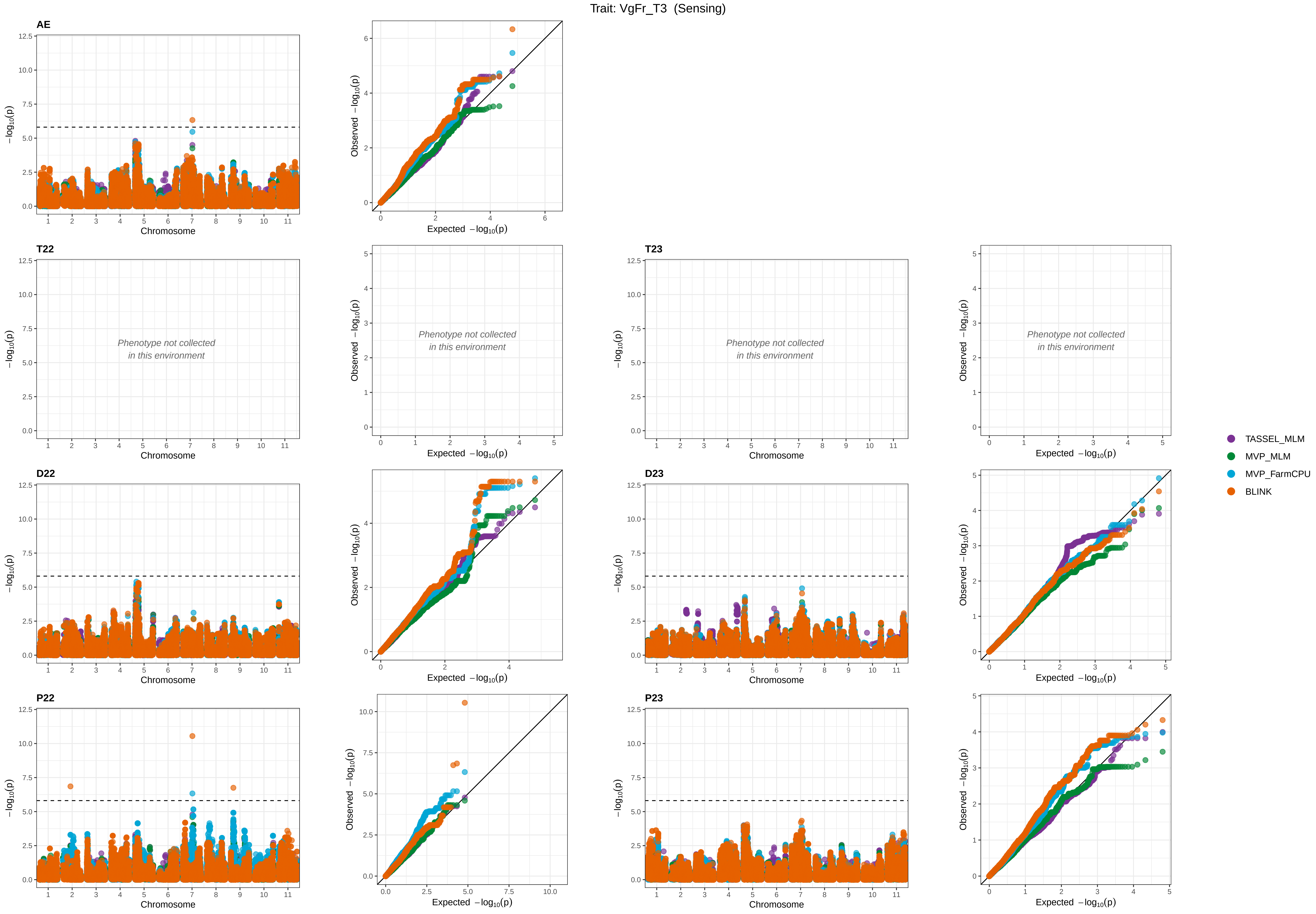

Trait: VgFr_T3 (Sensing)
AE
T22
T23
D22
D23
P22
P23
Phenotype not collected
in this environment
Chromosome
−log10(p)
Expected −log10(p)
Observed −log10(p)
TASSEL_MLM
MVP_MLM
MVP_FarmCPU
BLINK

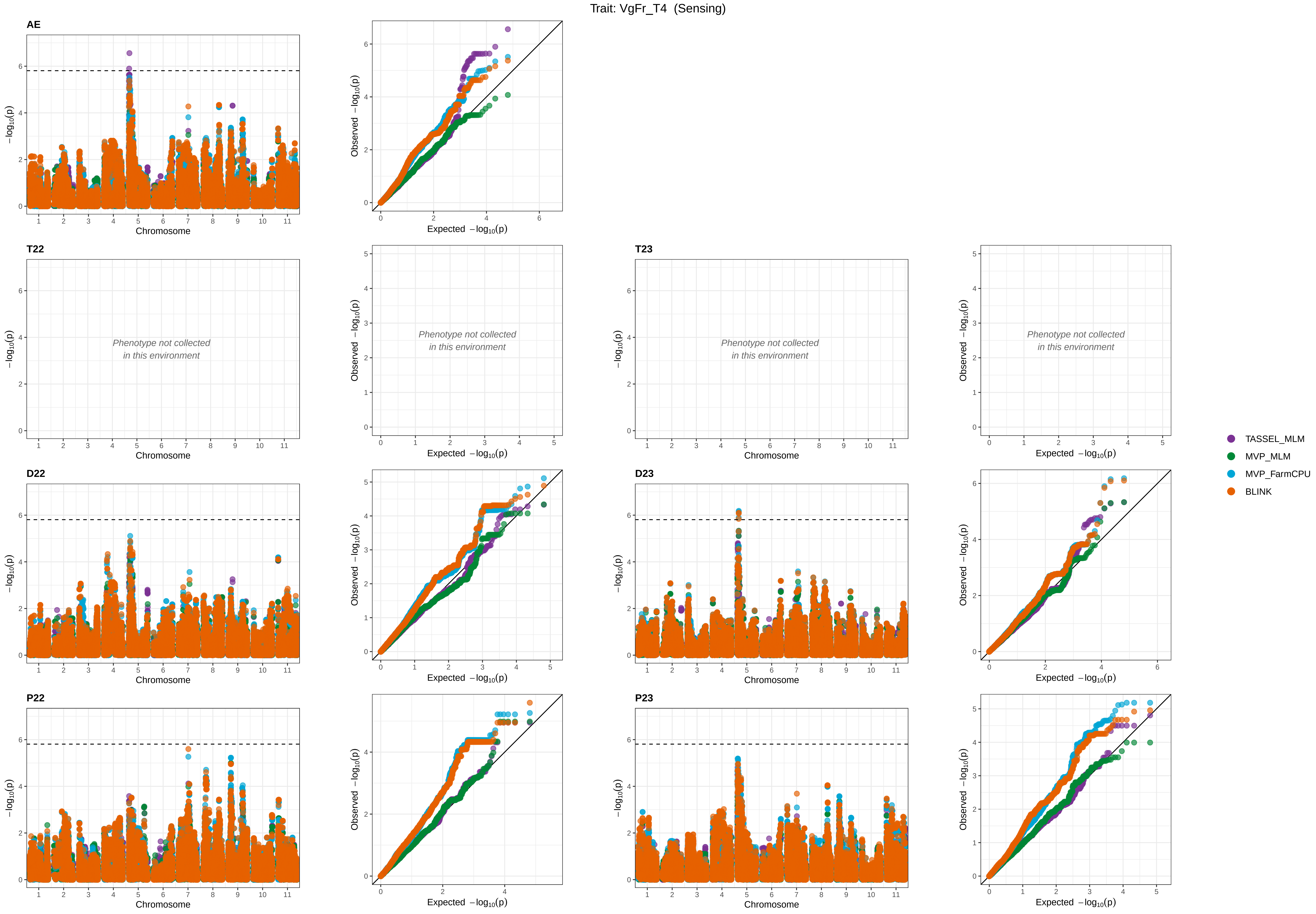

Trait: VgFr_T4 (Sensing)
AE
T22
T23
D22
D23
P22
P23
Phenotype not collected in this environment
Chromosome
−log10(p)
Expected −log10(p)
Observed −log10(p)
TASSEL_MLM
MVP_MLM
MVP_FarmCPU
BLINK

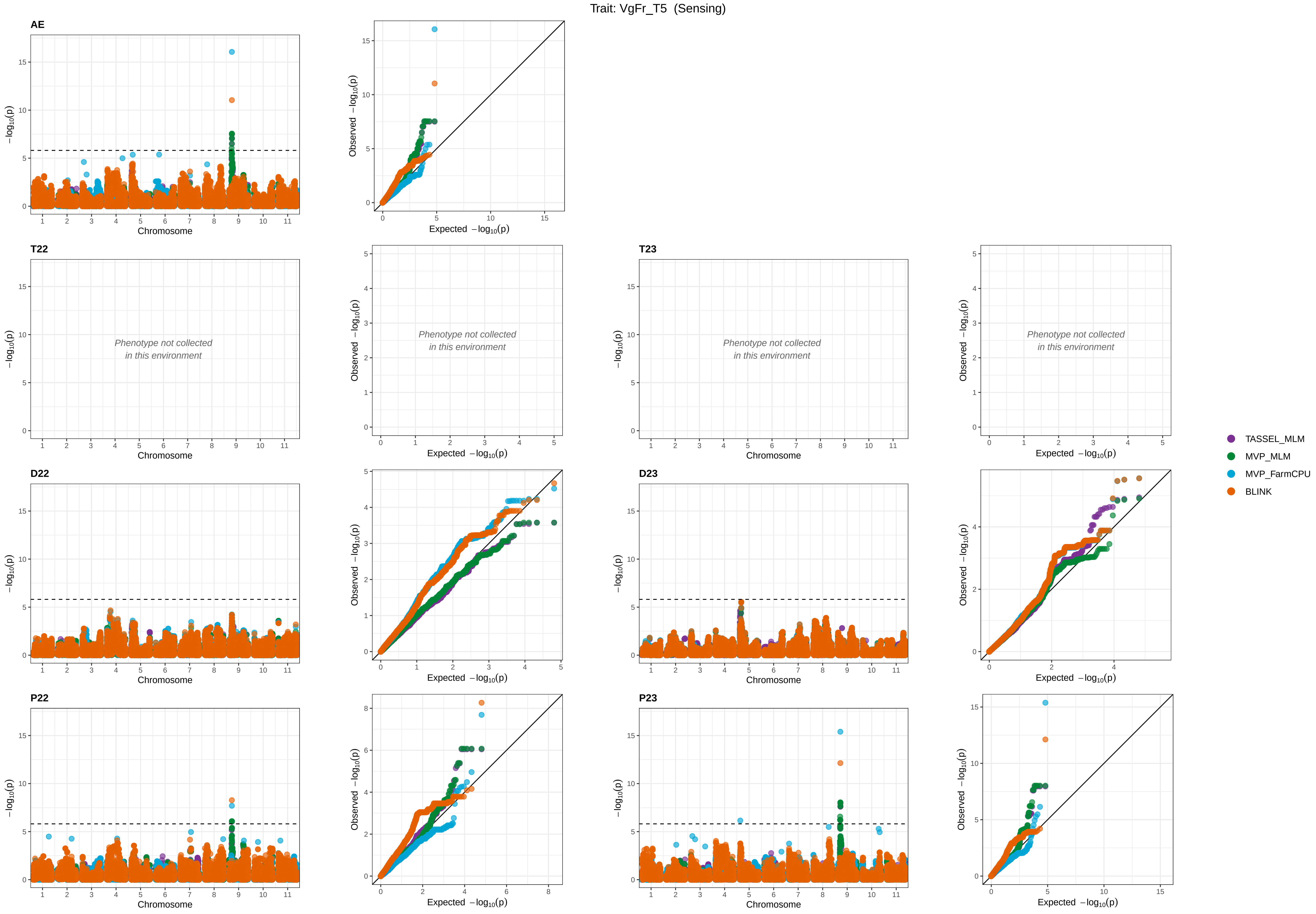

Trait: VgFr_T5 (Sensing)
AE
T22
T23
D22
D23
P22
P23
Phenotype not collected
in this environment
Chromosome
Expected −log10(p)
Observed −log10(p)
−log10(p)
TASSEL_MLM
MVP_MLM
MVP_FarmCPU
BLINK

Trait: VgFr_T6 (Sensing)

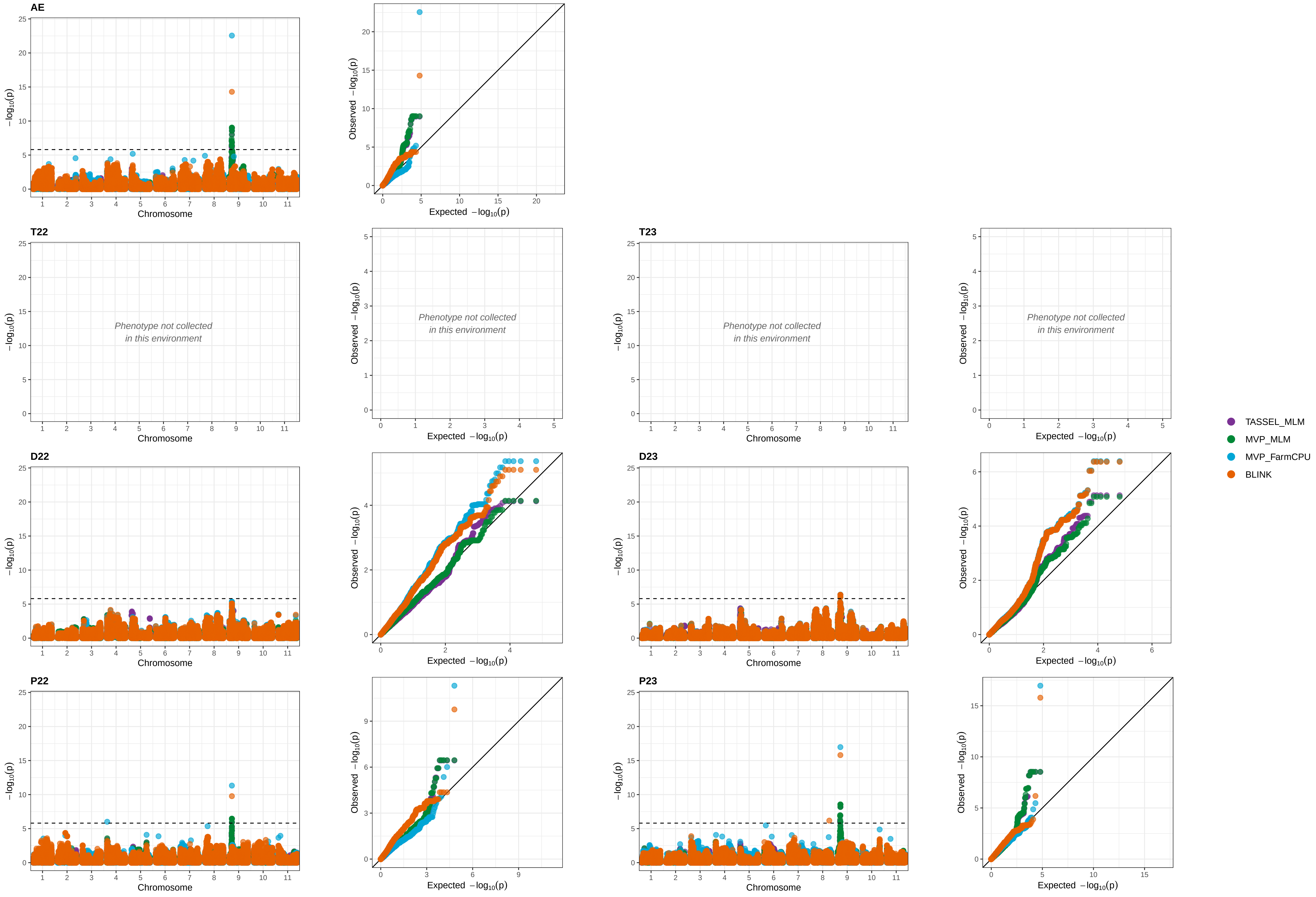

Trait: VgFr_T7 (Sensing)

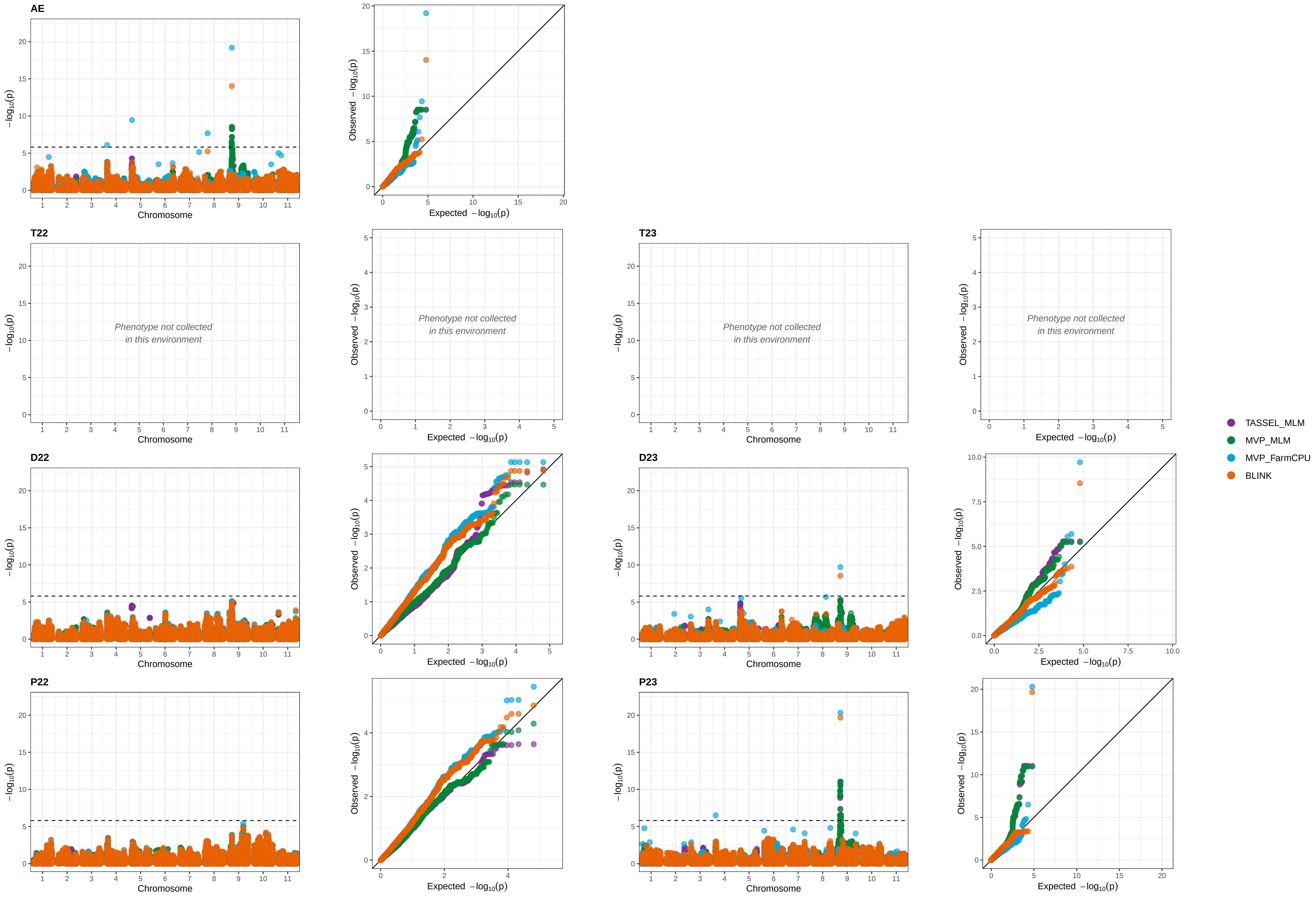

Trait: VgFr_T8 (Sensing)

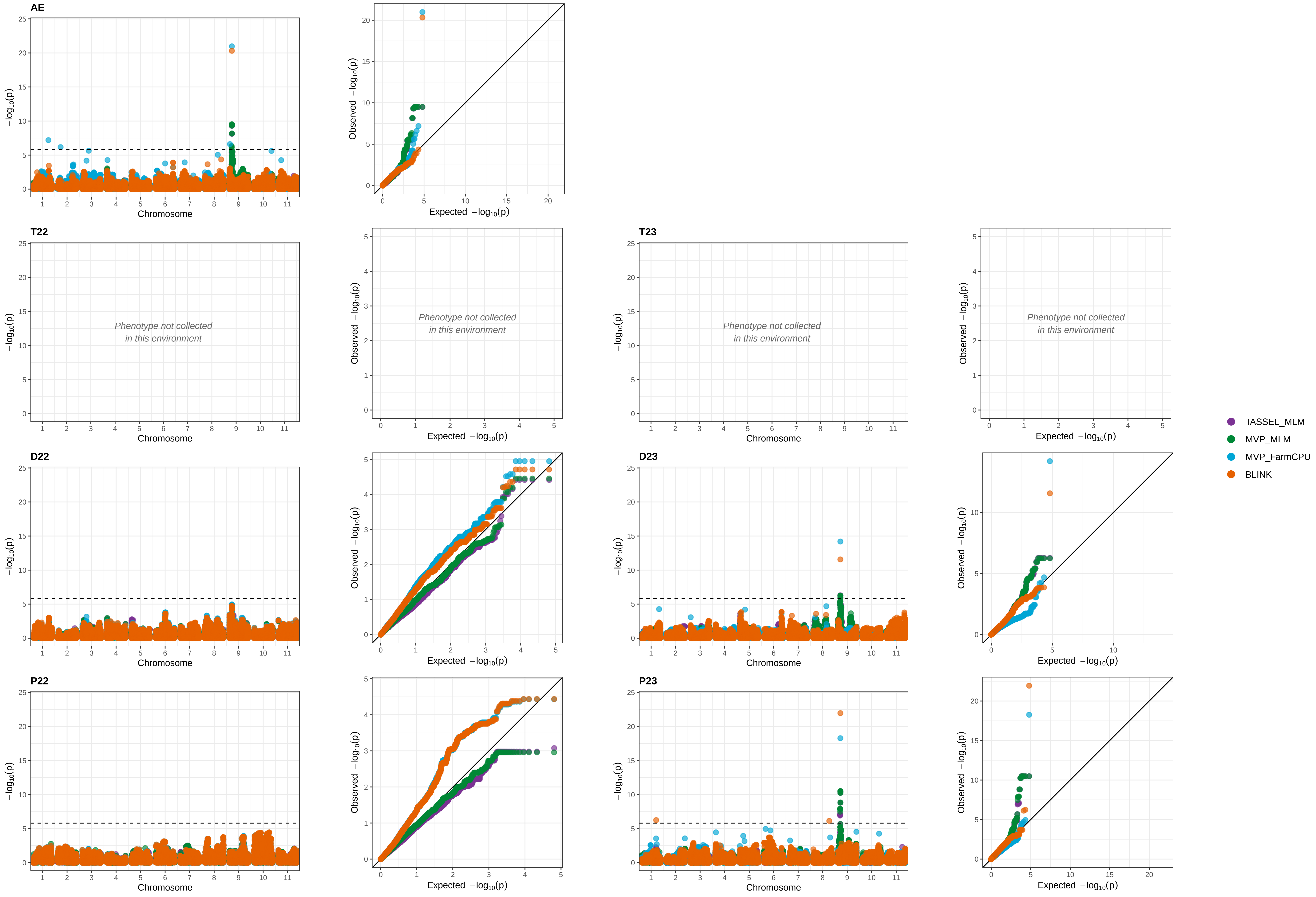

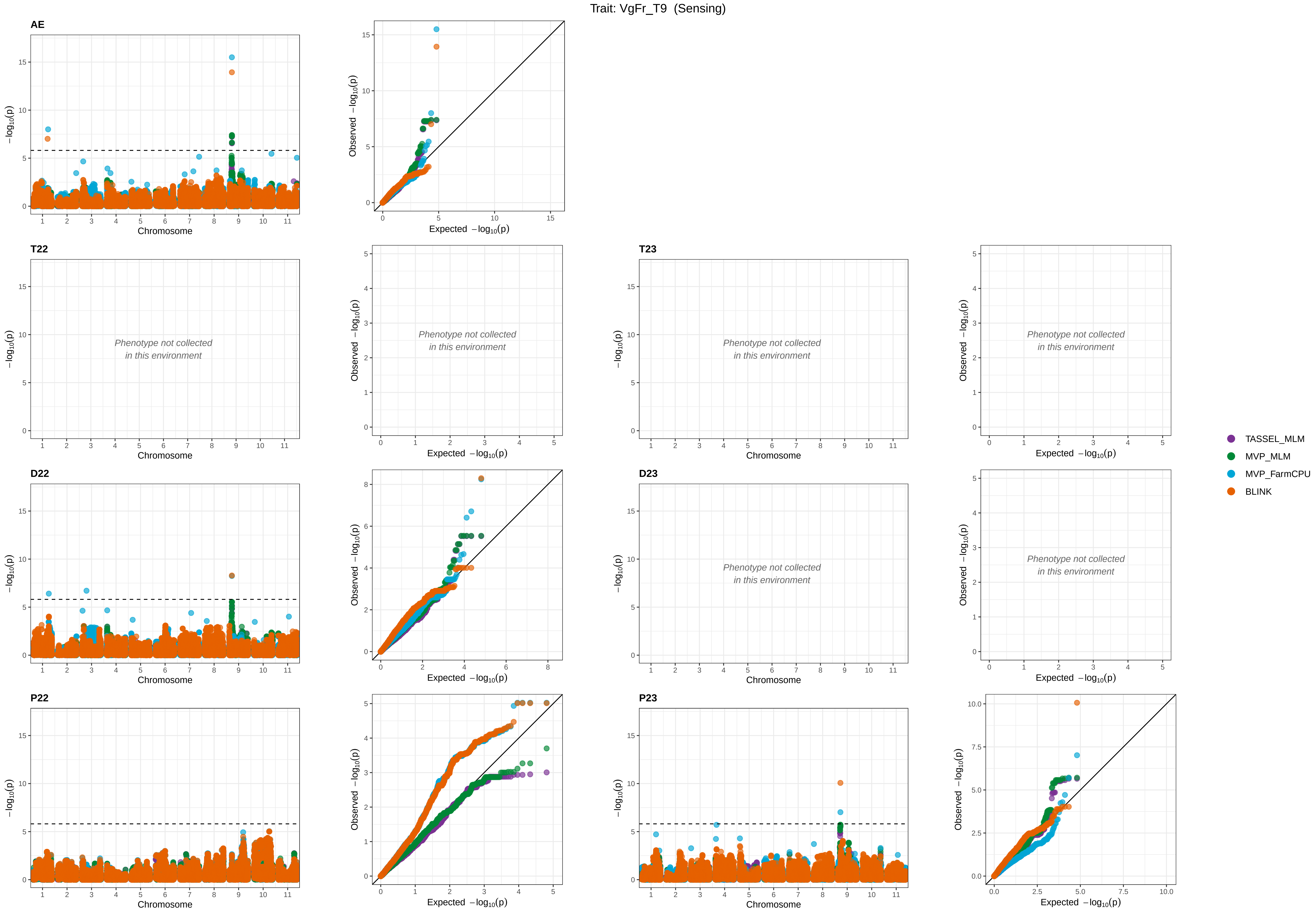
Trait: VgFr_T9 (Sensing)
AE
T22
T23
D22
D23
P22
P23
−log10(p)
Chromosome
Observed −log10(p)
Expected −log10(p)
Phenotype not collected
in this environment
TASSEL_MLM
MVP_MLM
MVP_FarmCPU
BLINK

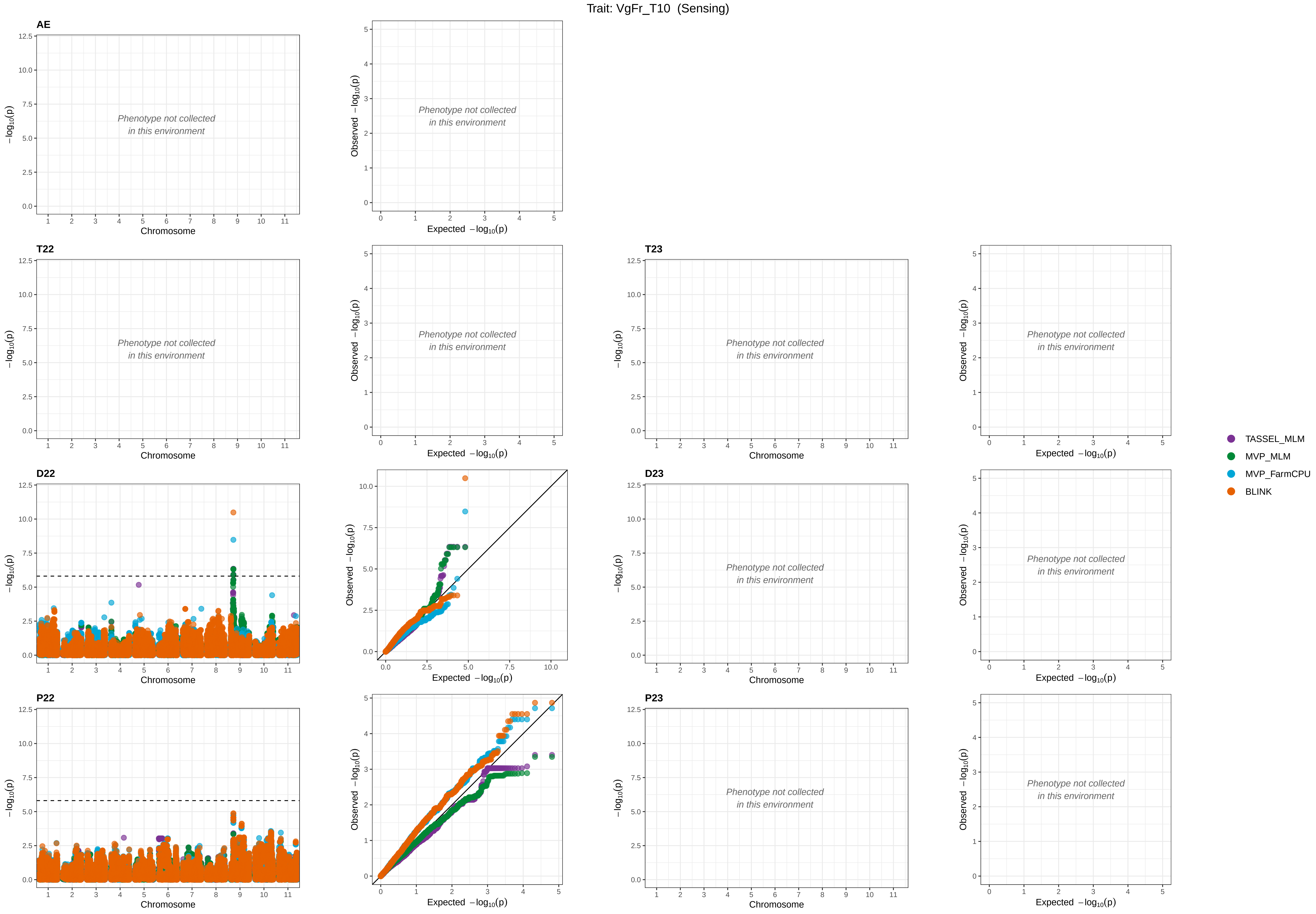

Trait: VgFr_T10 (Sensing)
AE
T22
T23
D22
D23
P22
P23
Phenotype not collected
in this environment
Chromosome
Expected −log10(p)
Observed −log10(p)
−log10(p)
TASSEL_MLM
MVP_MLM
MVP_FarmCPU
BLINK

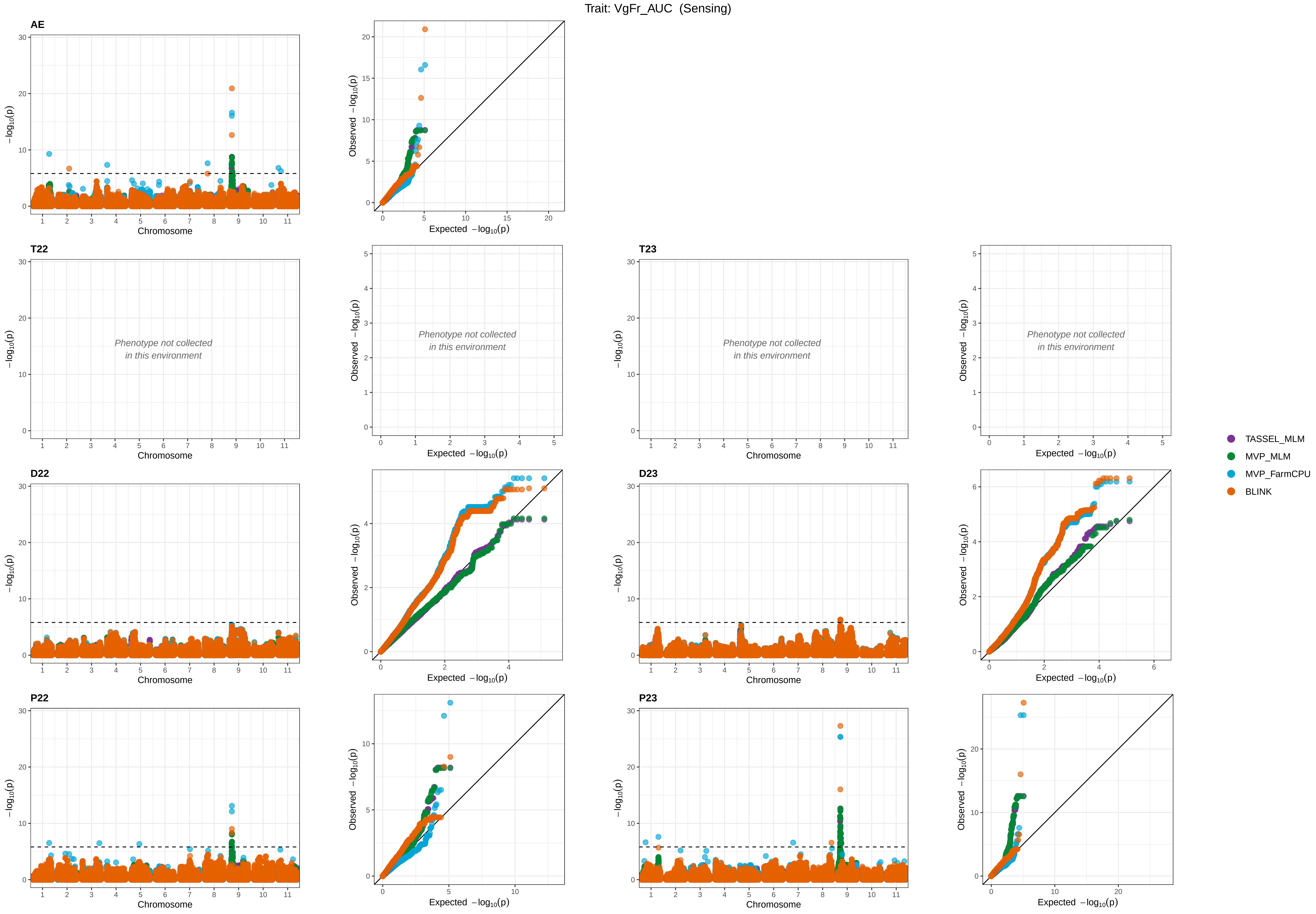

Trait: VgFr_AUC (Sensing)
AE
T22
T23
D22
D23
P22
P23
Phenotype not collected in this environment
Chromosome
Expected −log10(p)
Observed −log10(p)
−log10(p)
TASSEL_MLM
MVP_MLM
MVP_FarmCPU
BLINK

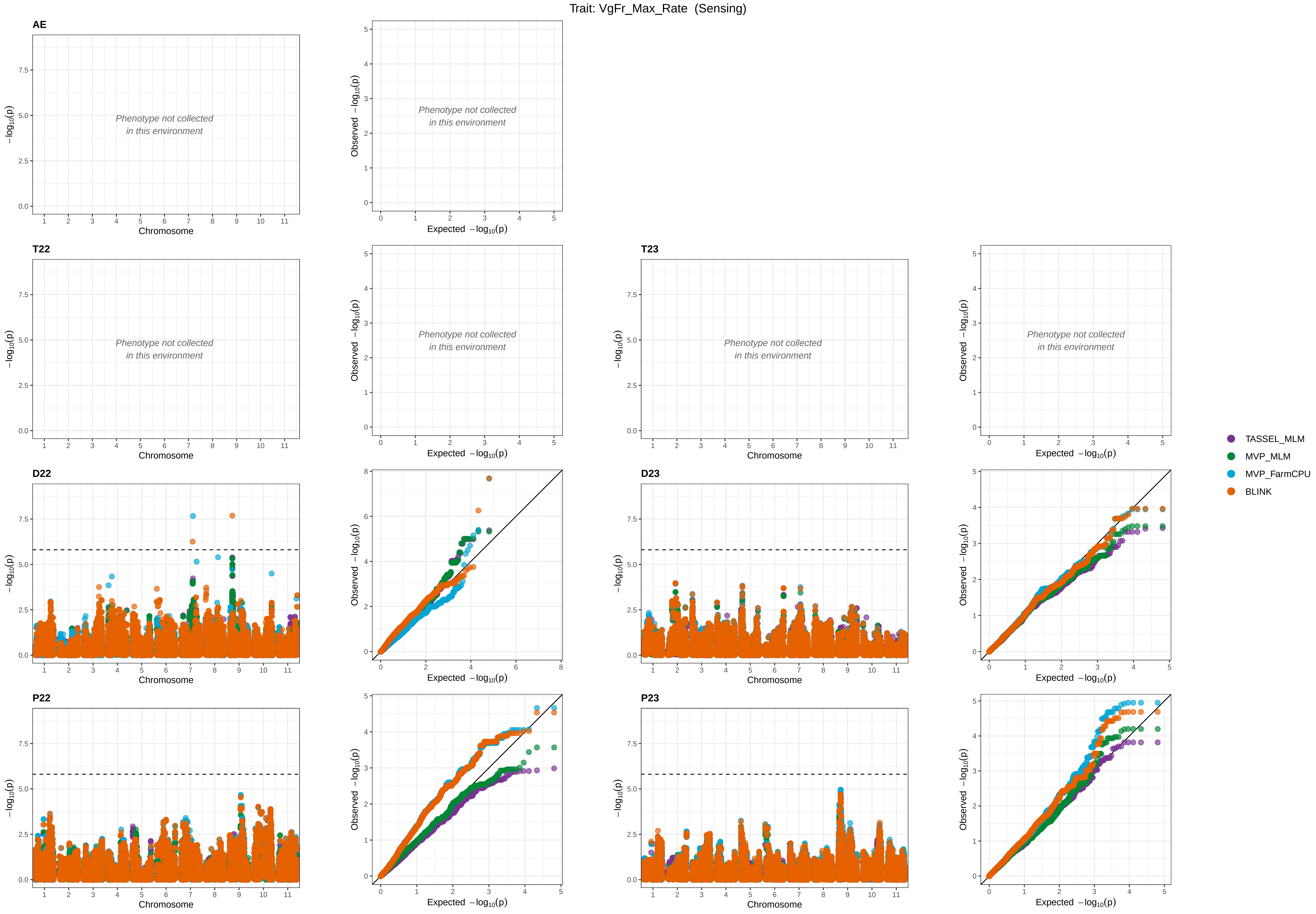

Trait: VgFr_Max_Rate (Sensing)
AE
T22
T23
D22
D23
P22
P23
Phenotype not collected
in this environment
Chromosome
Expected −log10(p)
Observed −log10(p)
−log10(p)
TASSEL_MLM
MVP_MLM
MVP_FarmCPU
BLINK

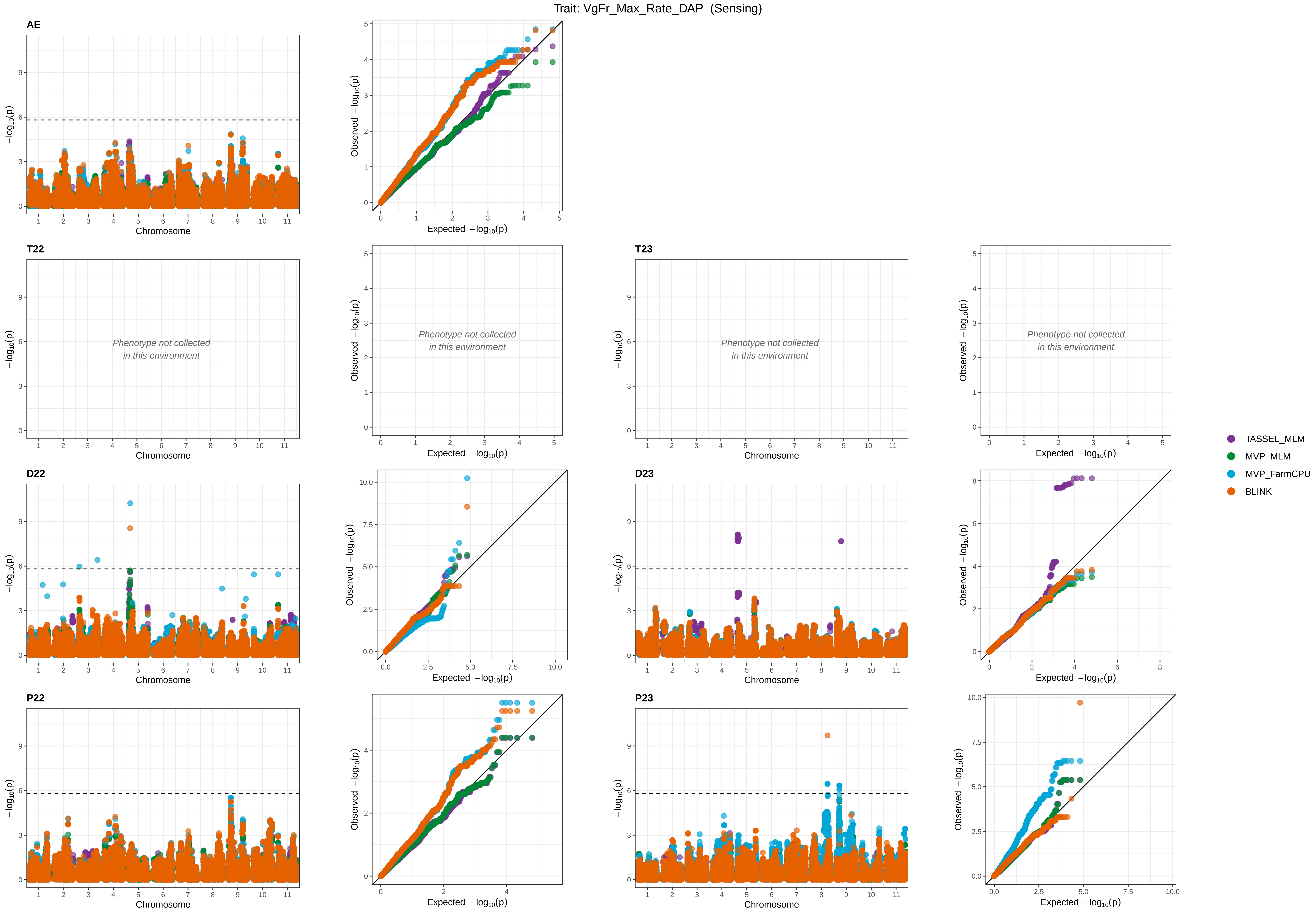

Trait: VgFr_Max_Rate_DAP (Sensing)
AE
T22
T23
D22
D23
P22
P23
Phenotype not collected in this environment
Chromosome
$-\log_{10}(p)$
Observed $-\log_{10}(p)$
Expected $-\log_{10}(p)$
TASSEL_MLM
MVP_MLM
MVP_FarmCPU
BLINK

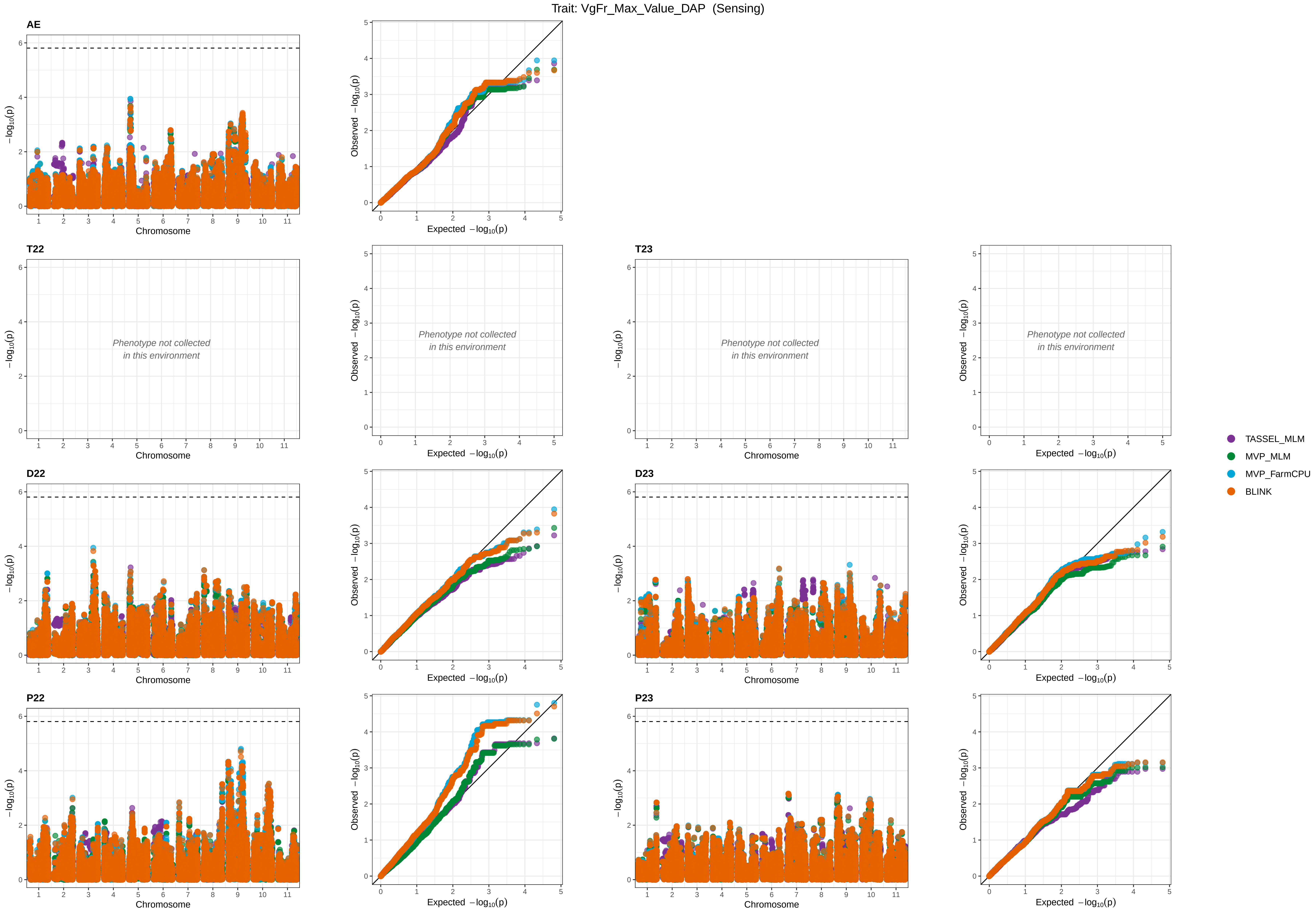

Trait: VgFr_Max_Value_DAP (Sensing)
AE
T22
T23
D22
D23
P22
P23
Phenotype not collected in this environment
Chromosome
Expected −log10(p)
Observed −log10(p)
−log10(p)
TASSEL_MLM
MVP_MLM
MVP_FarmCPU
BLINK

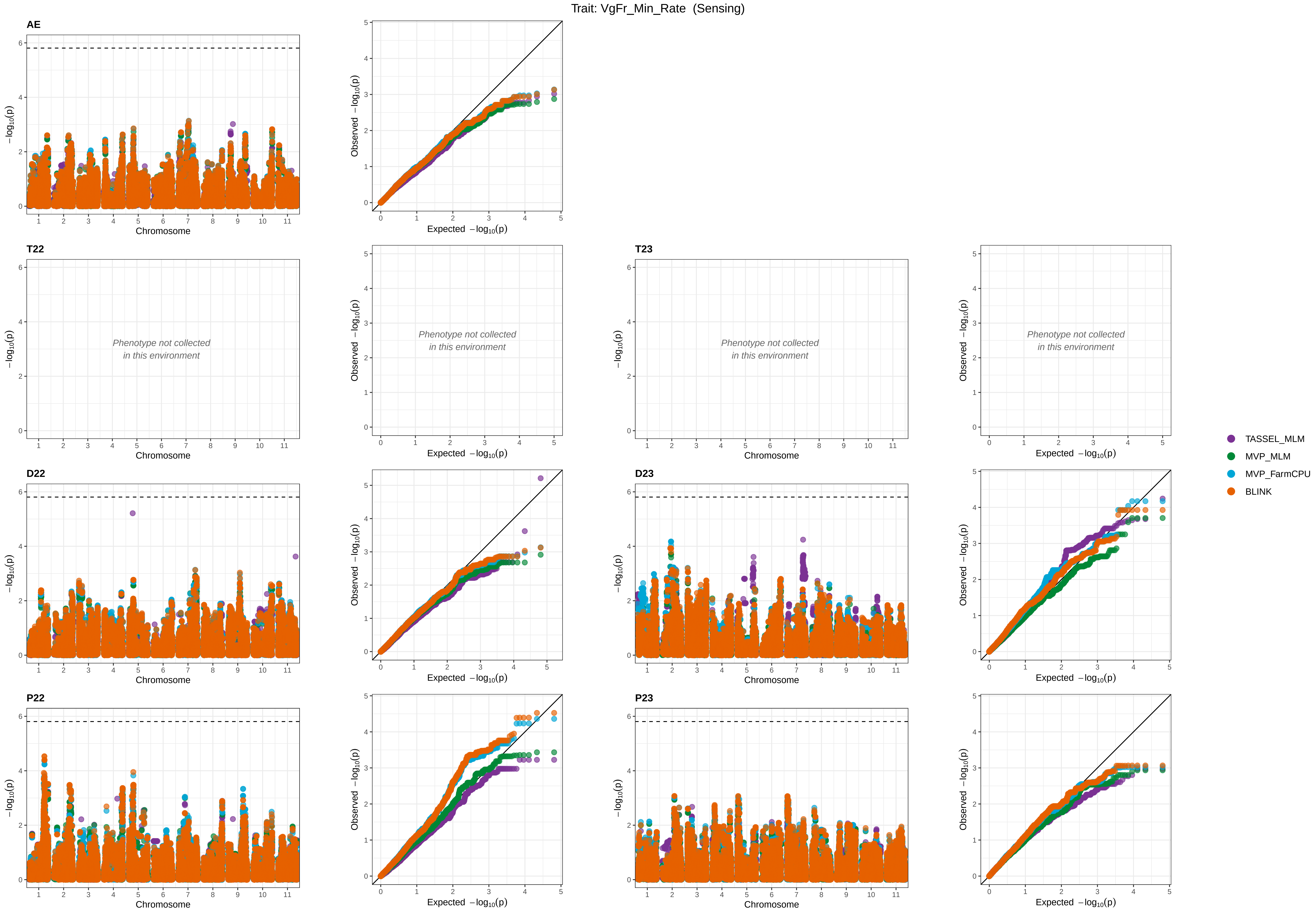

Trait: VgFr_Min_Rate (Sensing)
AE
T22
T23
D22
D23
P22
P23
Phenotype not collected
in this environment
−log10(p)
Chromosome
Observed −log10(p)
Expected −log10(p)
TASSEL_MLM
MVP_MLM
MVP_FarmCPU
BLINK

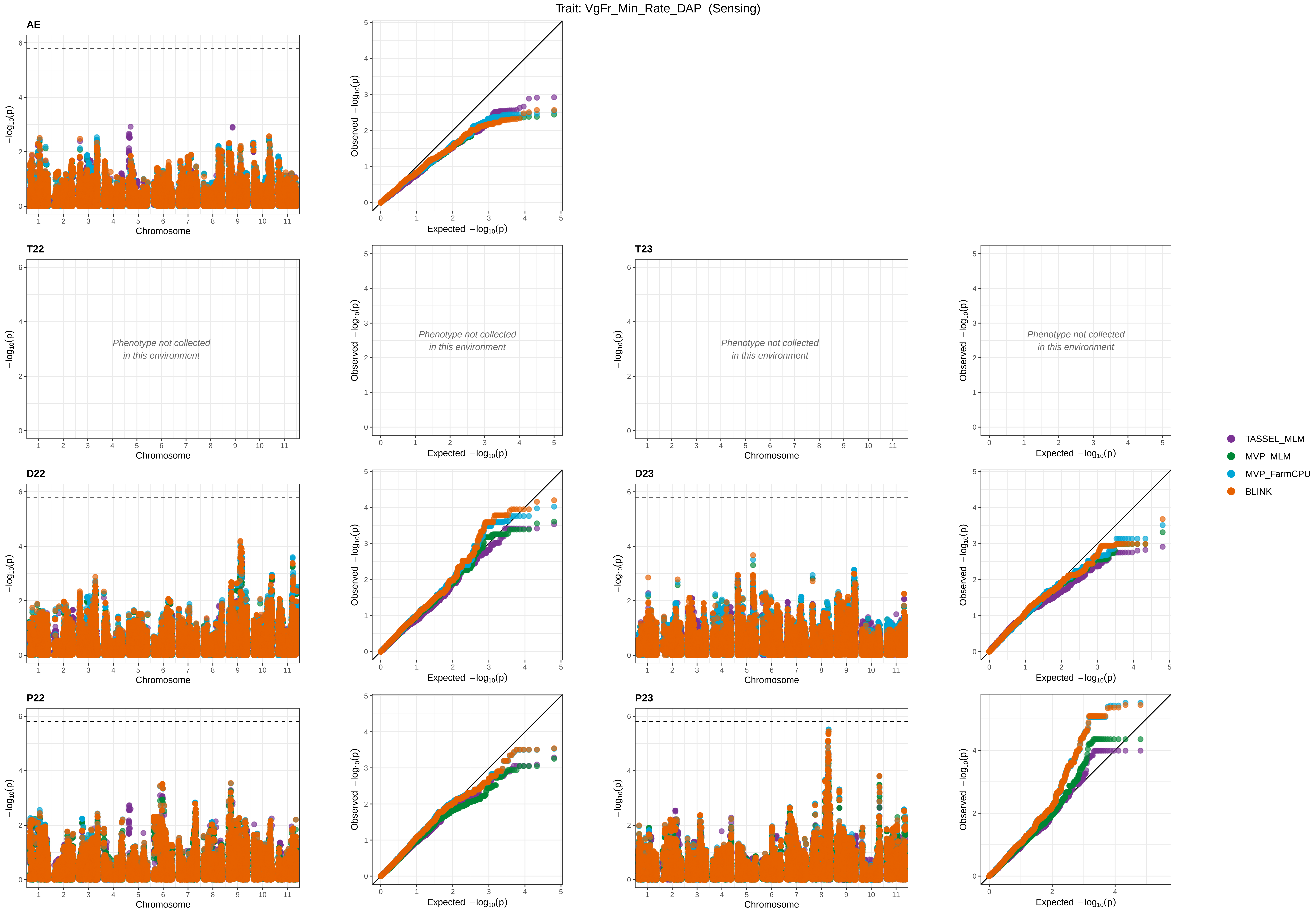

Trait: VgFr_Min_Rate_DAP (Sensing)
AE
T22
T23
D22
D23
P22
P23
Phenotype not collected in this environment
Chromosome
Expected −log10(p)
Observed −log10(p)
−log10(p)
TASSEL_MLM
MVP_MLM
MVP_FarmCPU
BLINK

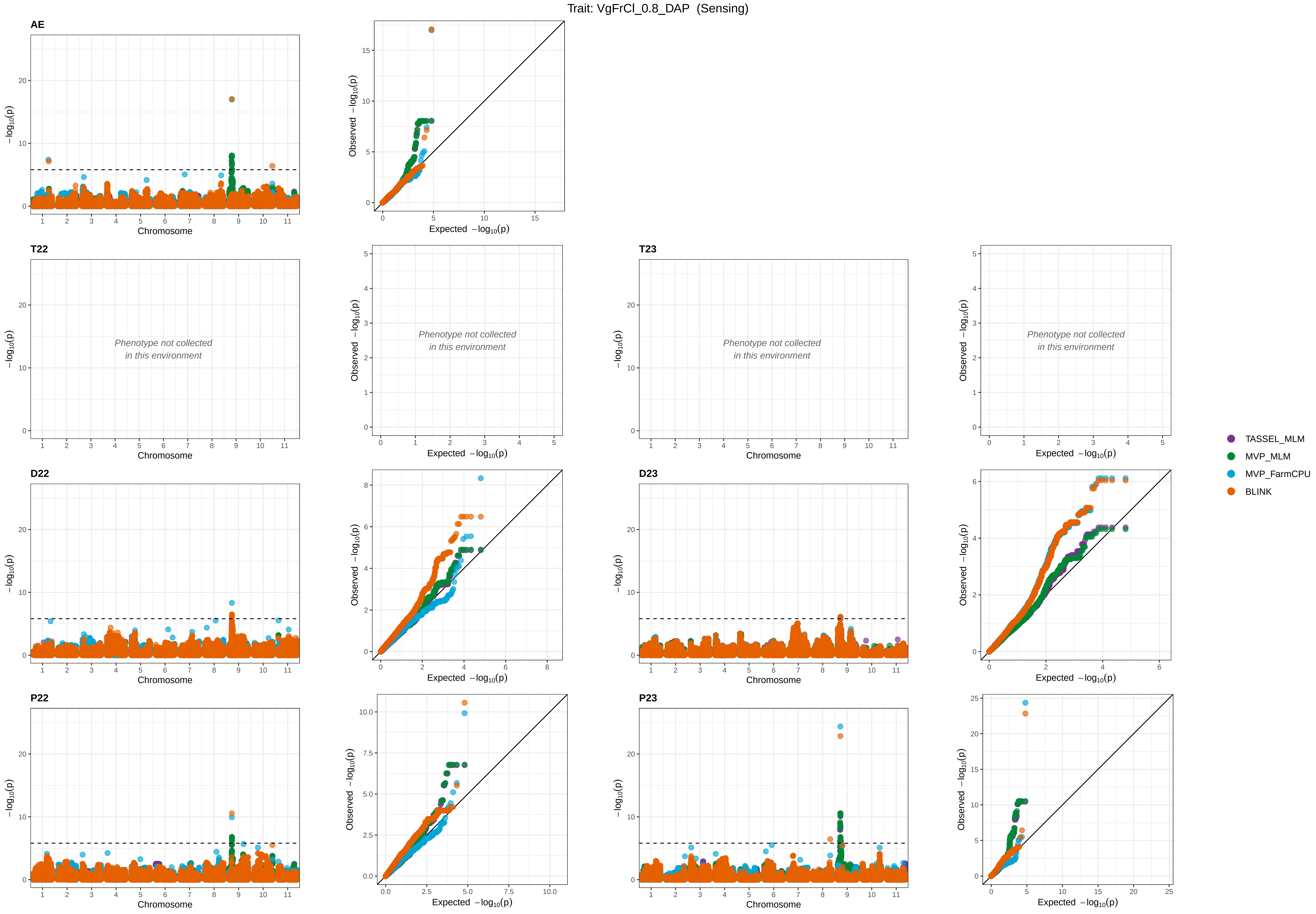

Trait: VgFrCl_0.8_DAP (Sensing)
AE
T22
T23
D22
D23
P22
P23
Phenotype not collected
in this environment
Chromosome
Expected −log10(p)
Observed −log10(p)
−log10(p)
TASSEL_MLM
MVP_MLM
MVP_FarmCPU
BLINK
