## Supplementary material for "Genetic mapping and genomic prediction for agronomic, grain compositional, and sensing-enabled traits in a cowpea MAGIC population along an environmental gradient": Large supplemental tables and figures, uploaded separately: SuppFig_10_AcrossEnv_QTLprofiles_FTconditional_Chr9.pdf

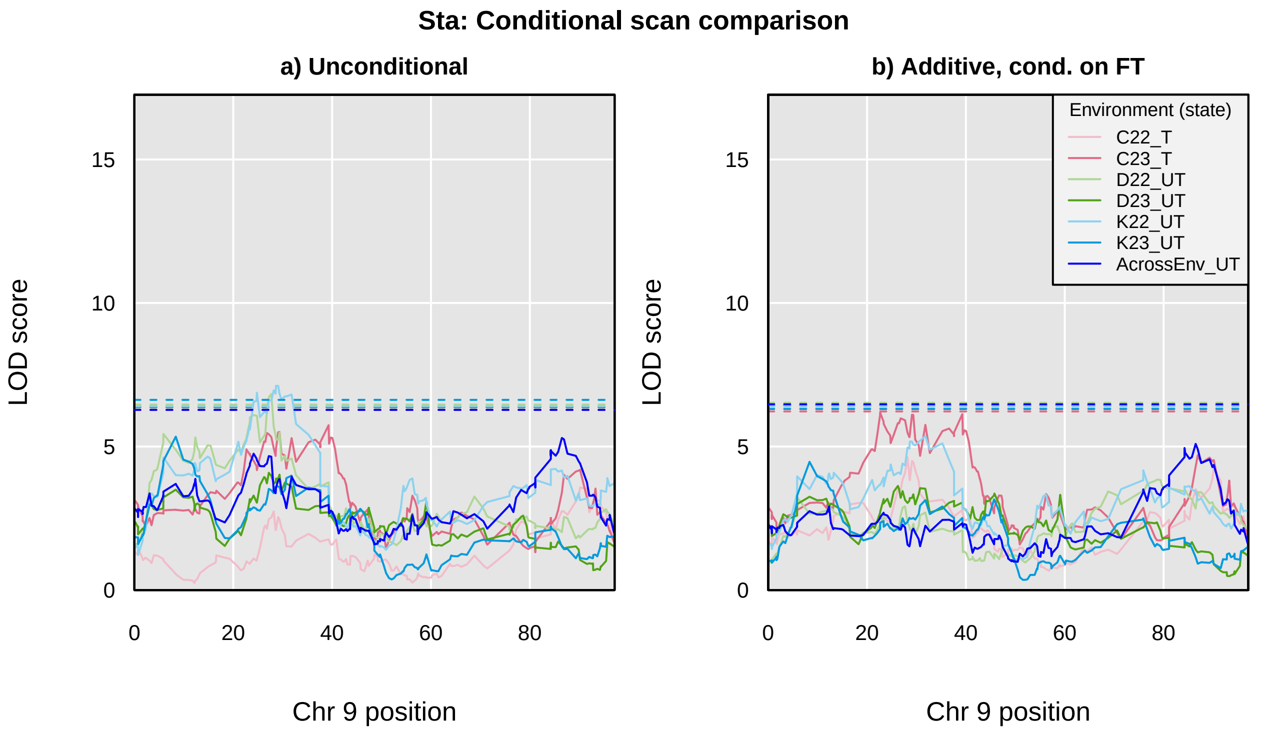
Sta: Conditional scan comparison
a) Unconditional
b) Additive, cond. on FT
LOD score
0
5
10
15
0
20
40
60
80
Chr 9 position
Environment (state)
C22_T
C23_T
D22_UT
D23_UT
K22_UT
K23_UT
AcrossEnv_UT

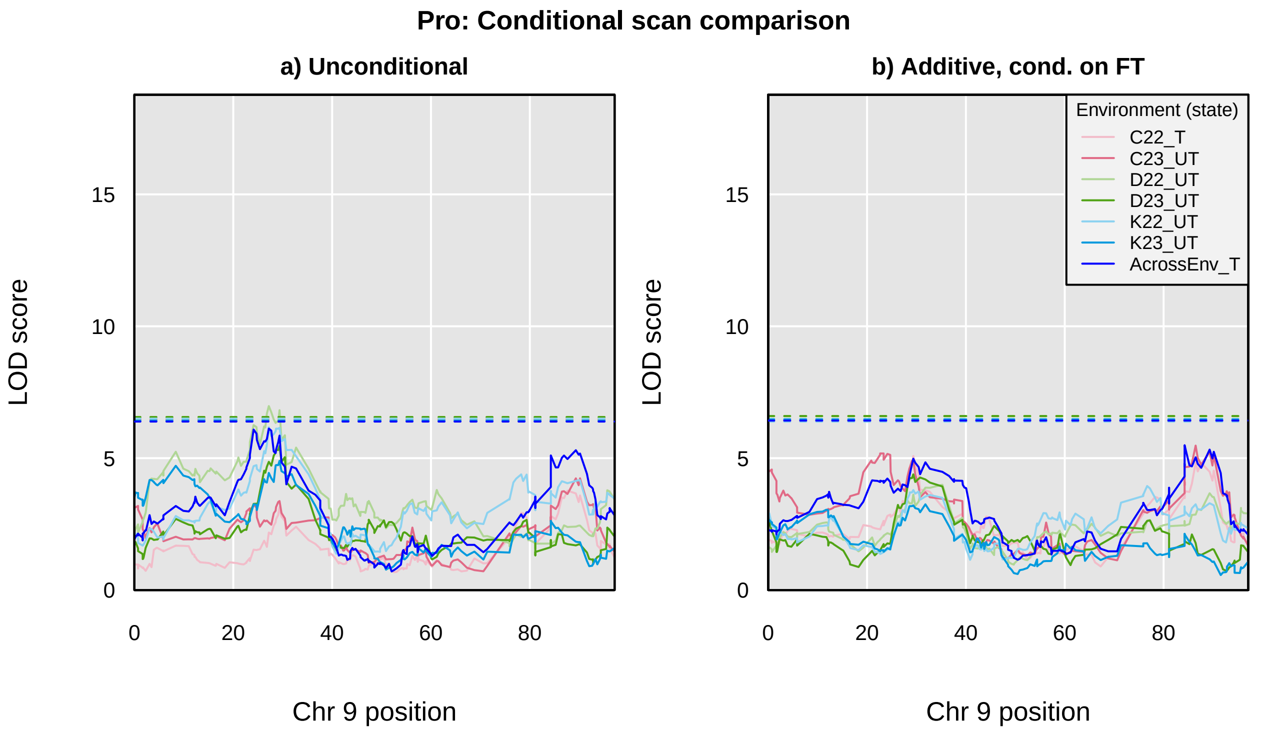

Pro: Conditional scan comparison
a) Unconditional
b) Additive, cond. on FT
LOD score
0
5
10
15
Chr 9 position
0
20
40
60
80
Environment (state)
C22_T
C23_UT
D22_UT
D23_UT
K22_UT
K23_UT
AcrossEnv_T

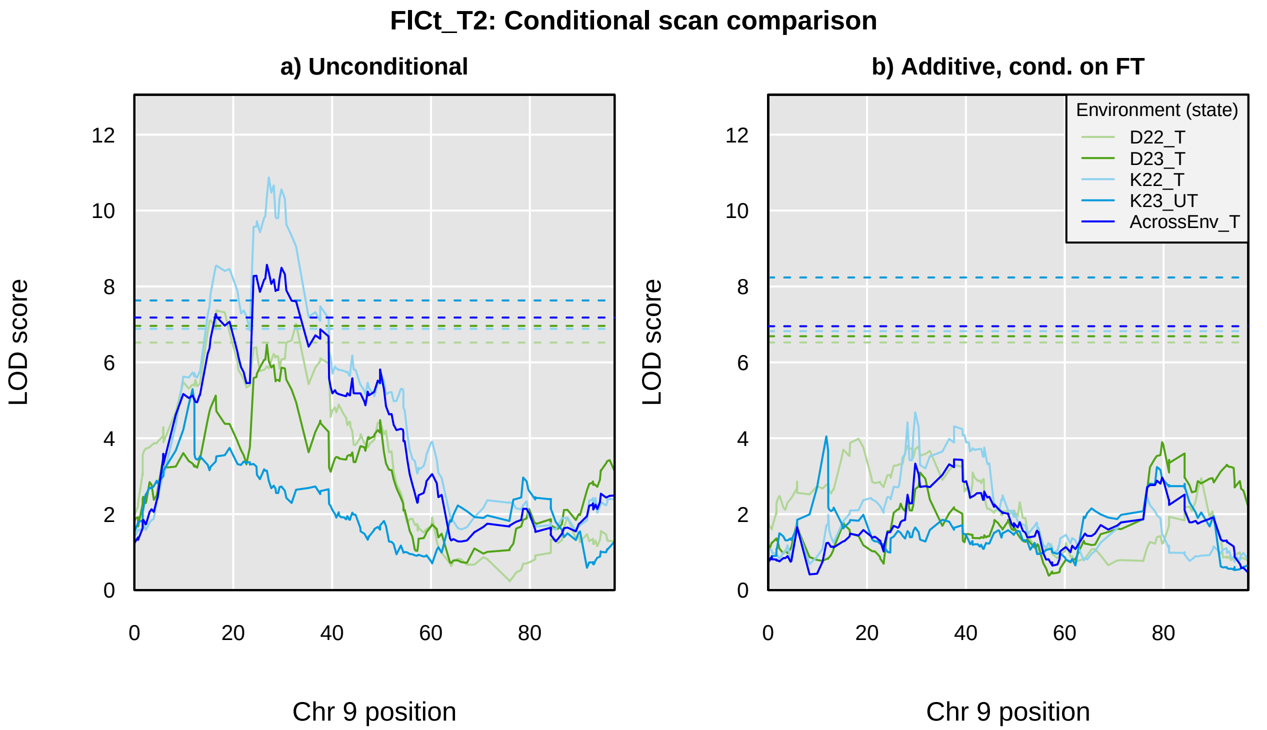
FlCt_T2: Conditional scan comparison
a) Unconditional
b) Additive, cond. on FT
LOD score
Chr 9 position
0
2
4
6
8
10
12
0
20
40
60
80
Environment (state)
D22_T
D23_T
K22_T
K23_UT
AcrossEnv_T

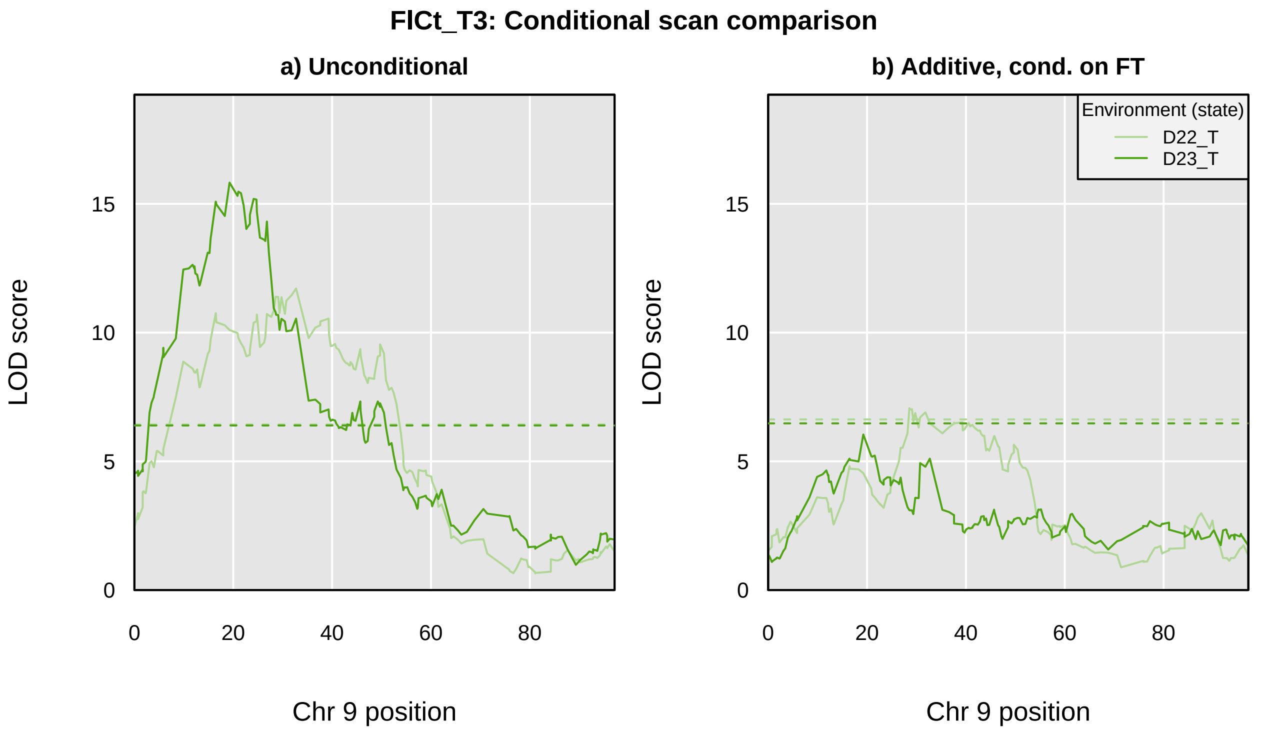
FlCt_T3: Conditional scan comparison
a) Unconditional
b) Additive, cond. on FT
LOD score
Chr 9 position
0
5
10
15
0
20
40
60
80
Environment (state)
D22_T
D23_T

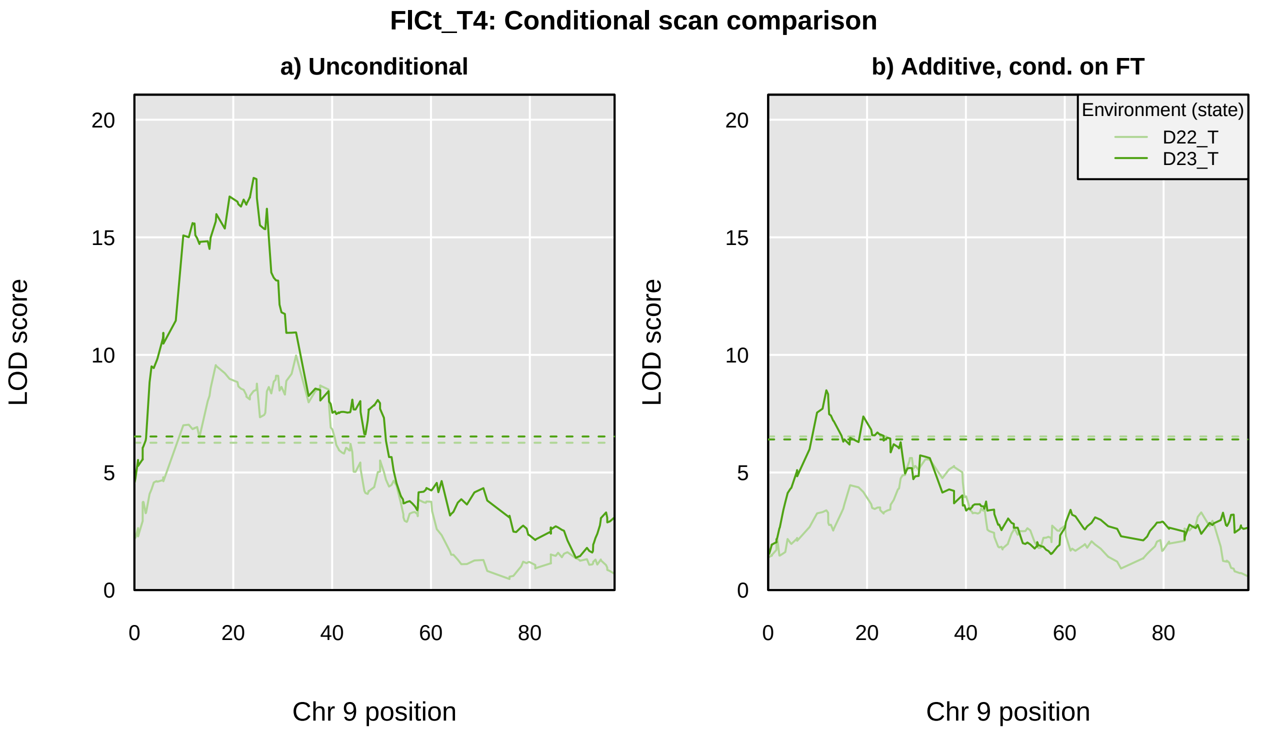

FlCt_T4: Conditional scan comparison
a) Unconditional
b) Additive, cond. on FT
LOD score
Chr 9 position
0
5
10
15
20
0
20
40
60
80
Environment (state)
D22_T
D23_T

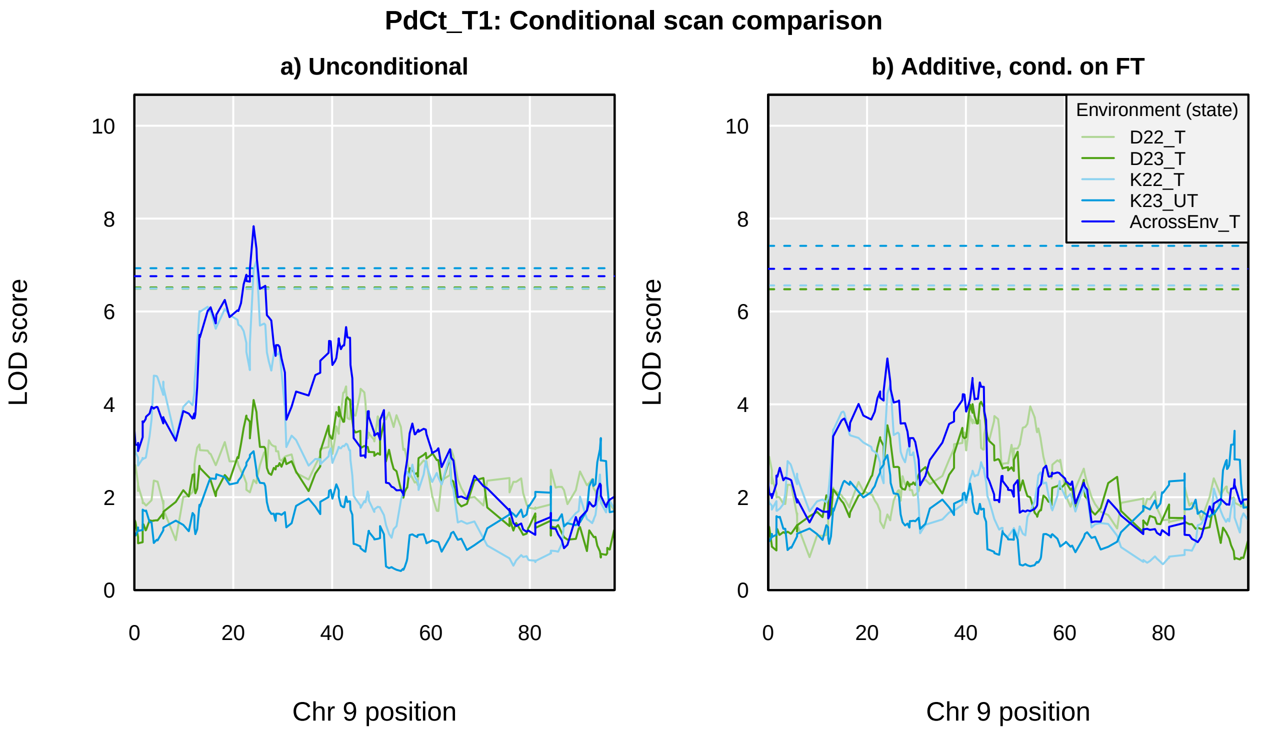
PdCt_T1: Conditional scan comparison
a) Unconditional
b) Additive, cond. on FT
LOD score
Chr 9 position
0
2
4
6
8
10
0
20
40
60
80
Environment (state)
D22_T
D23_T
K22_T
K23_UT
AcrossEnv_T

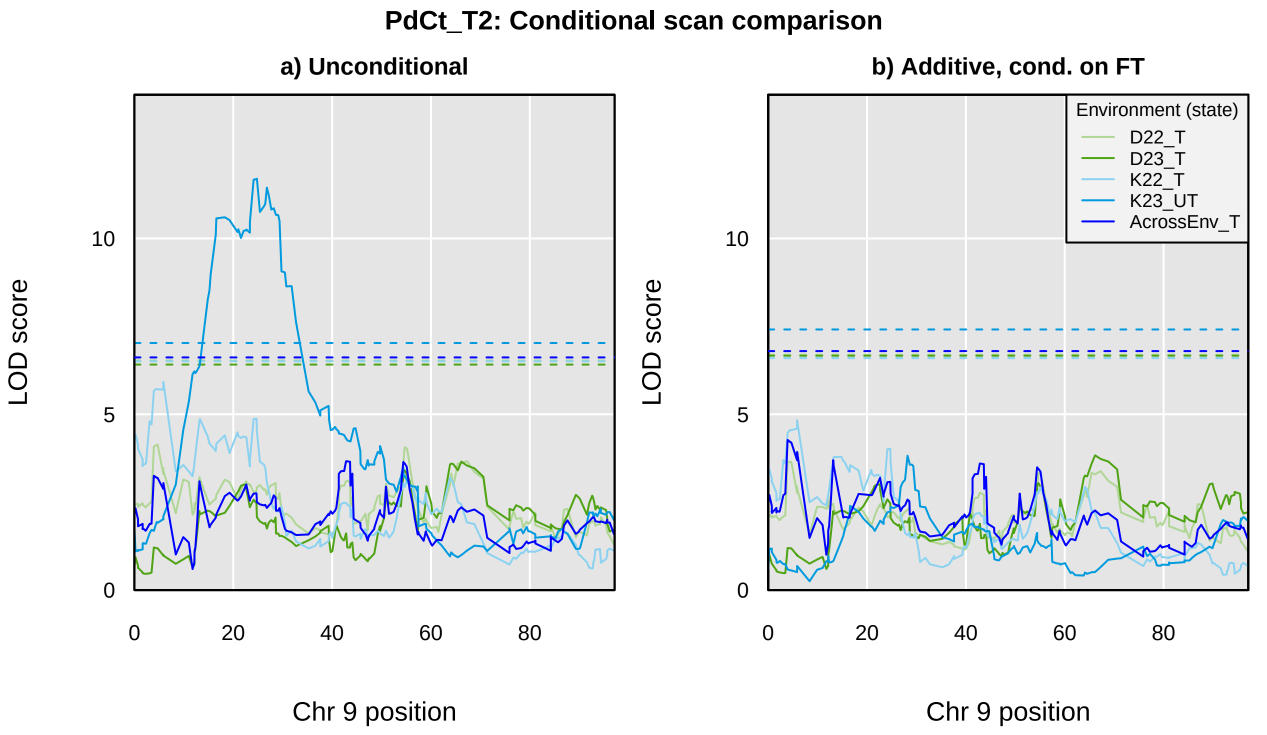

PdCt_T2: Conditional scan comparison
a) Unconditional
b) Additive, cond. on FT
LOD score
Chr 9 position
0
5
10
0
20
40
60
80
Environment (state)
D22_T
D23_T
K22_T
K23_UT
AcrossEnv_T

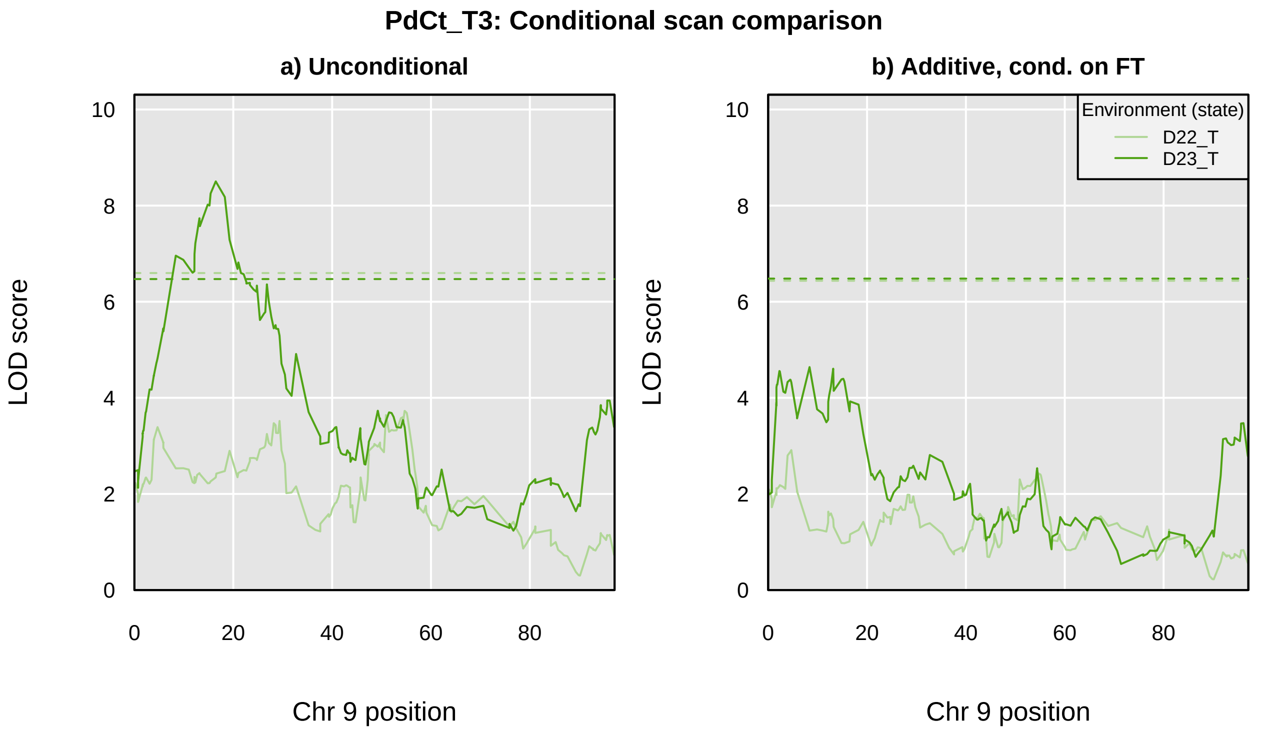
PdCt_T3: Conditional scan comparison
a) Unconditional
b) Additive, cond. on FT
LOD score
Chr 9 position
0
2
4
6
8
10
0
20
40
60
80
Environment (state)
D22_T
D23_T

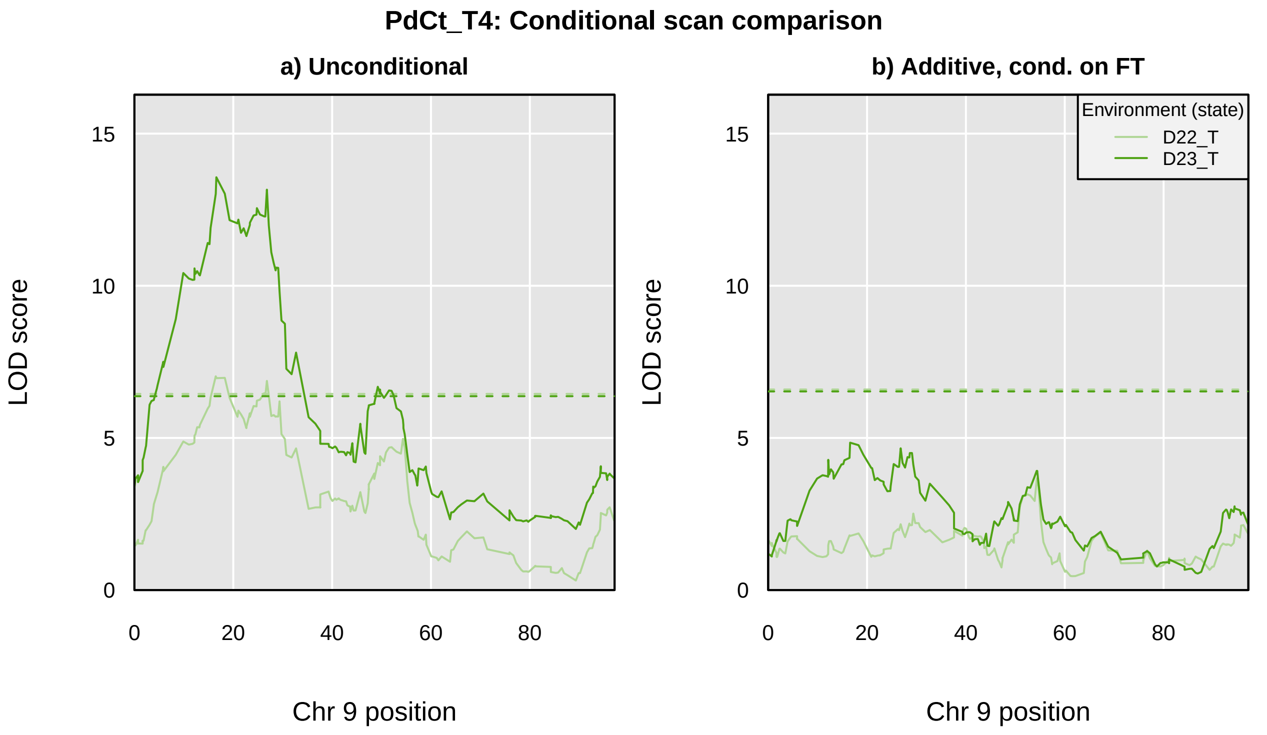
PdCt_T4: Conditional scan comparison
a) Unconditional
b) Additive, cond. on FT
Environment (state)
D22_T
D23_T
LOD score
0
5
10
15
0
20
40
60
80
Chr 9 position

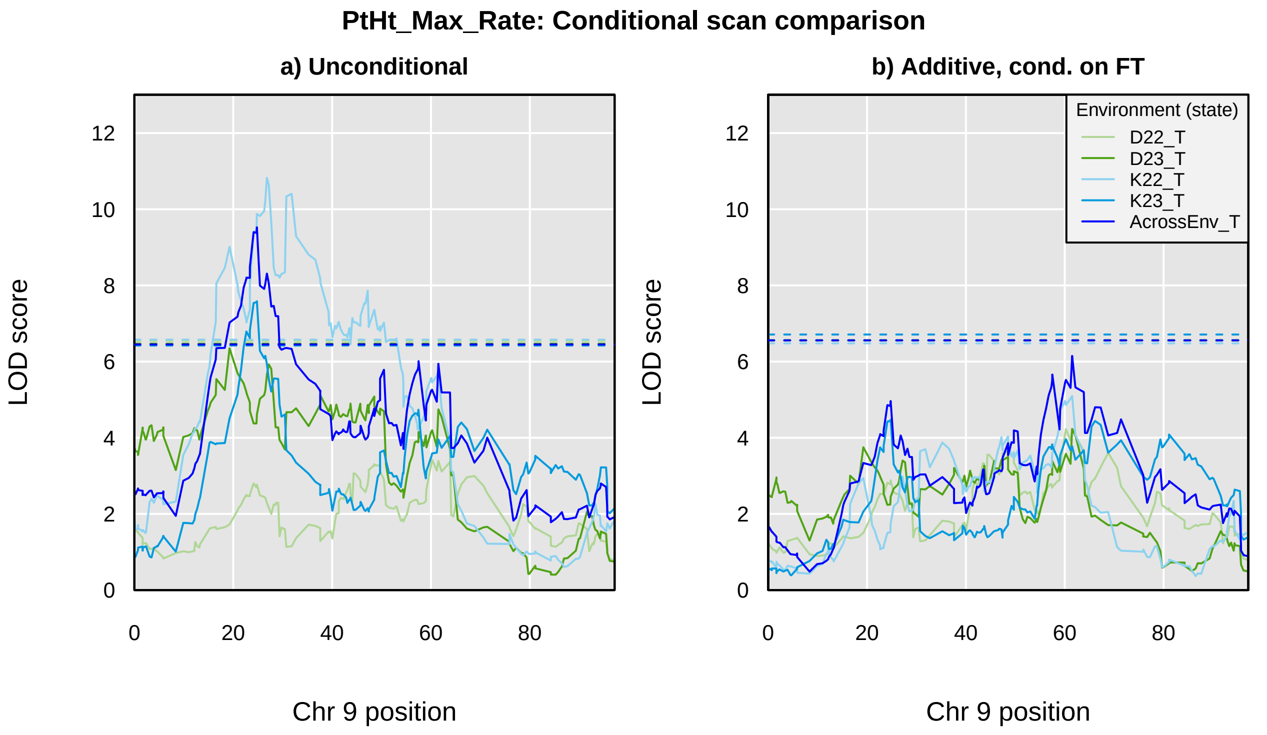
PtHt_Max_Rate: Conditional scan comparison
a) Unconditional
b) Additive, cond. on FT
LOD score
0
2
4
6
8
10
12
0
20
40
60
80
Chr 9 position
Environment (state)
D22_T
D23_T
K22_T
K23_T
AcrossEnv_T

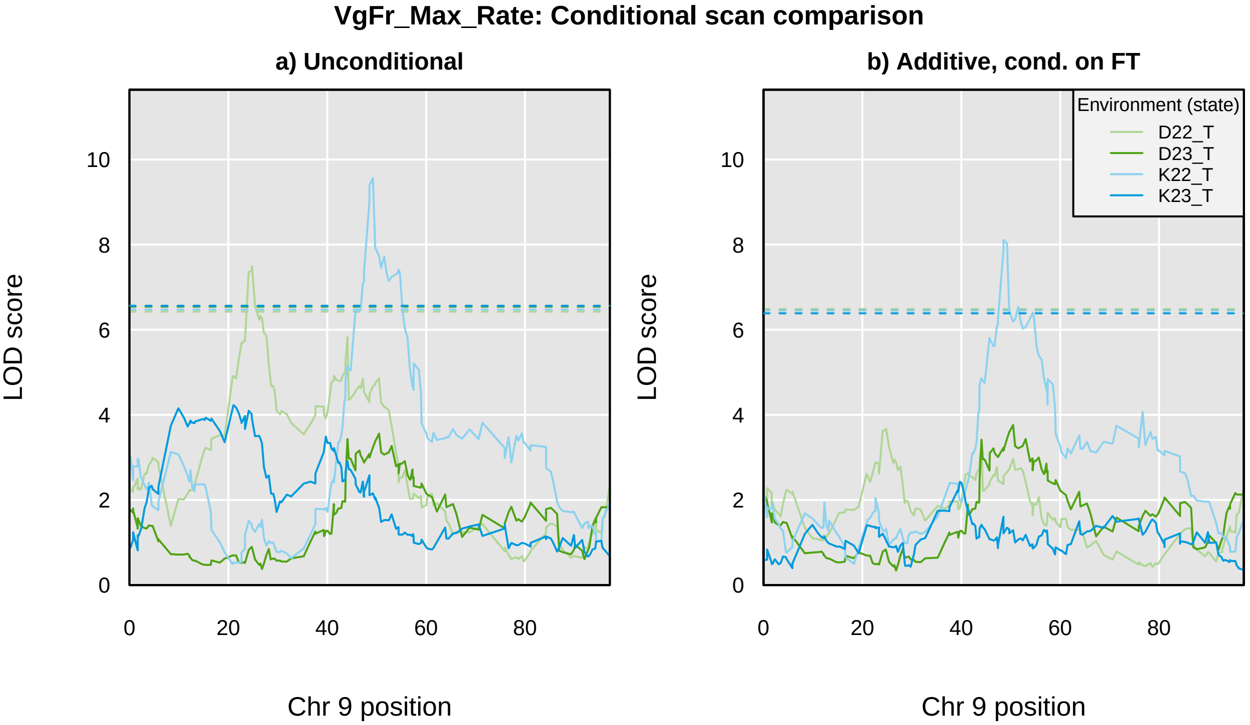

VgFr_Max_Rate: Conditional scan comparison
a) Unconditional
b) Additive, cond. on FT
LOD score
Chr 9 position
0
2
4
6
8
10
0
20
40
60
80
Environment (state)
D22_T
D23_T
K22_T
K23_T

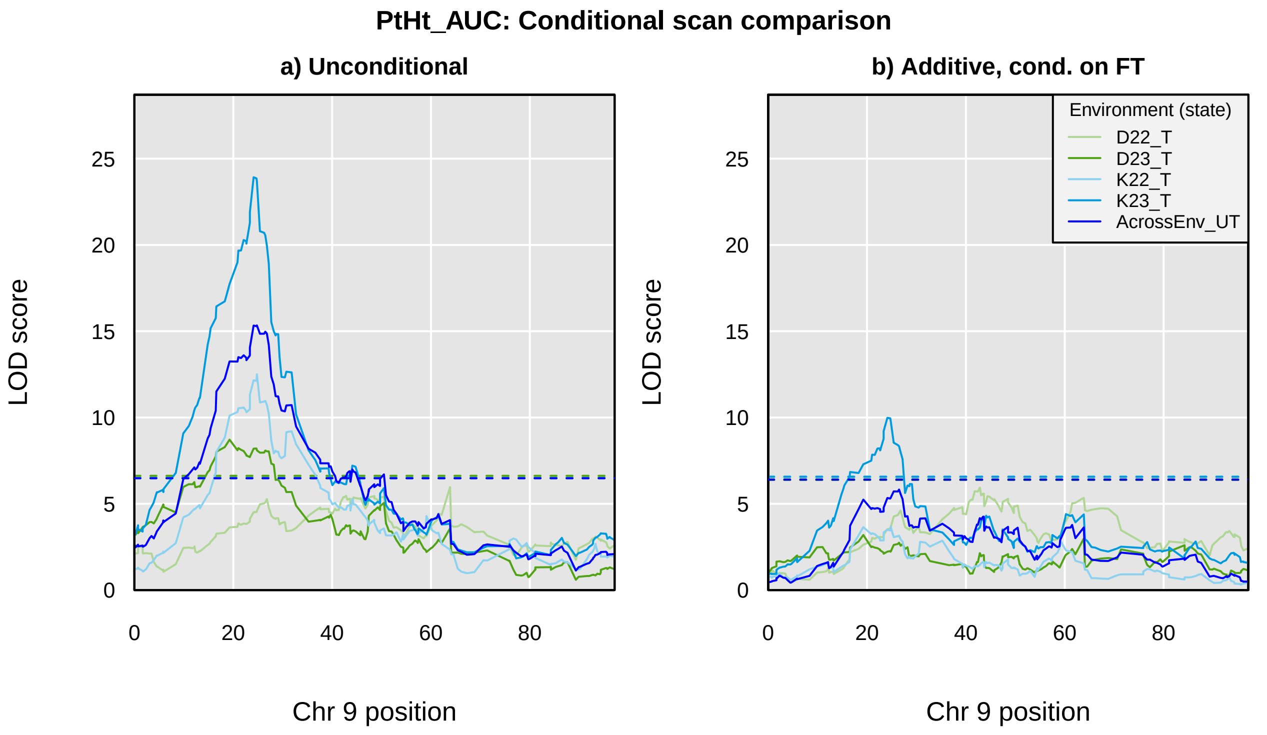
PtHt_AUC: Conditional scan comparison
a) Unconditional
b) Additive, cond. on FT
LOD score
Chr 9 position
0
5
10
15
20
25
0
20
40
60
80
Environment (state)
D22_T
D23_T
K22_T
K23_T
AcrossEnv_UT

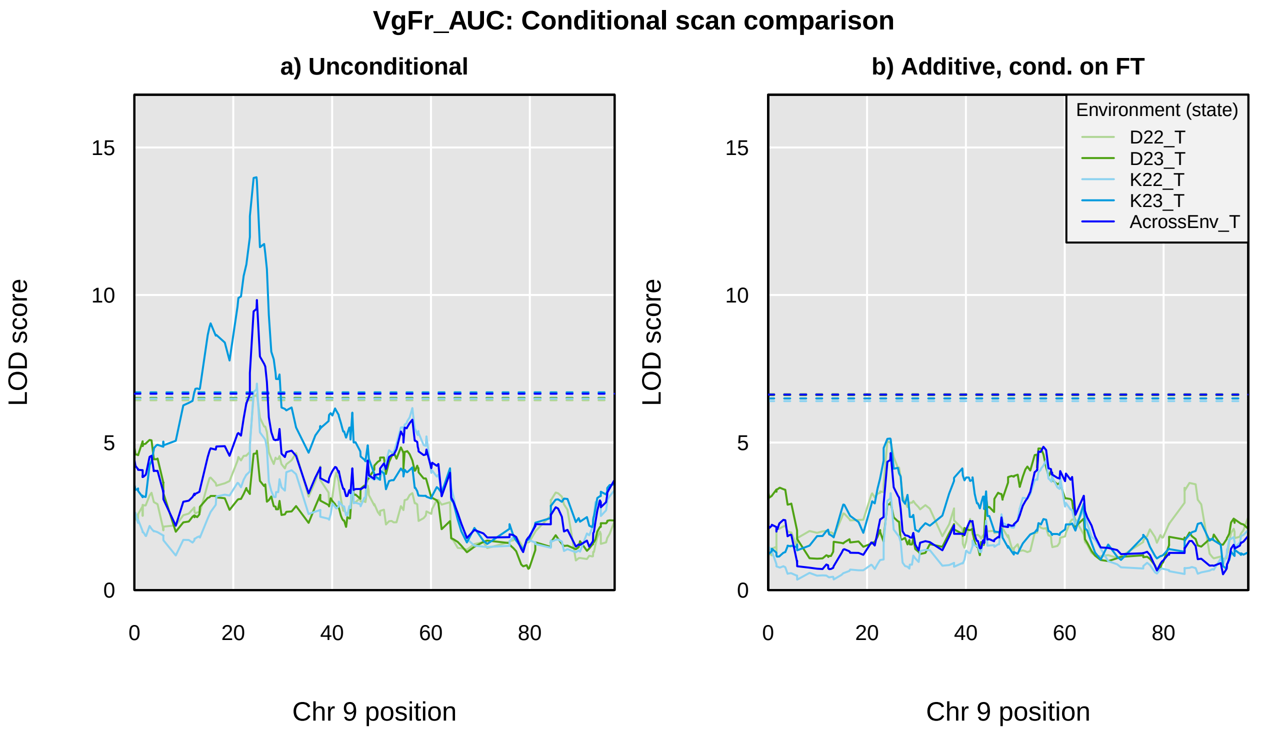
VgFr_AUC: Conditional scan comparison
a) Unconditional
b) Additive, cond. on FT
LOD score
Chr 9 position
0
5
10
15
0
20
40
60
80
Environment (state)
D22_T
D23_T
K22_T
K23_T
AcrossEnv_T

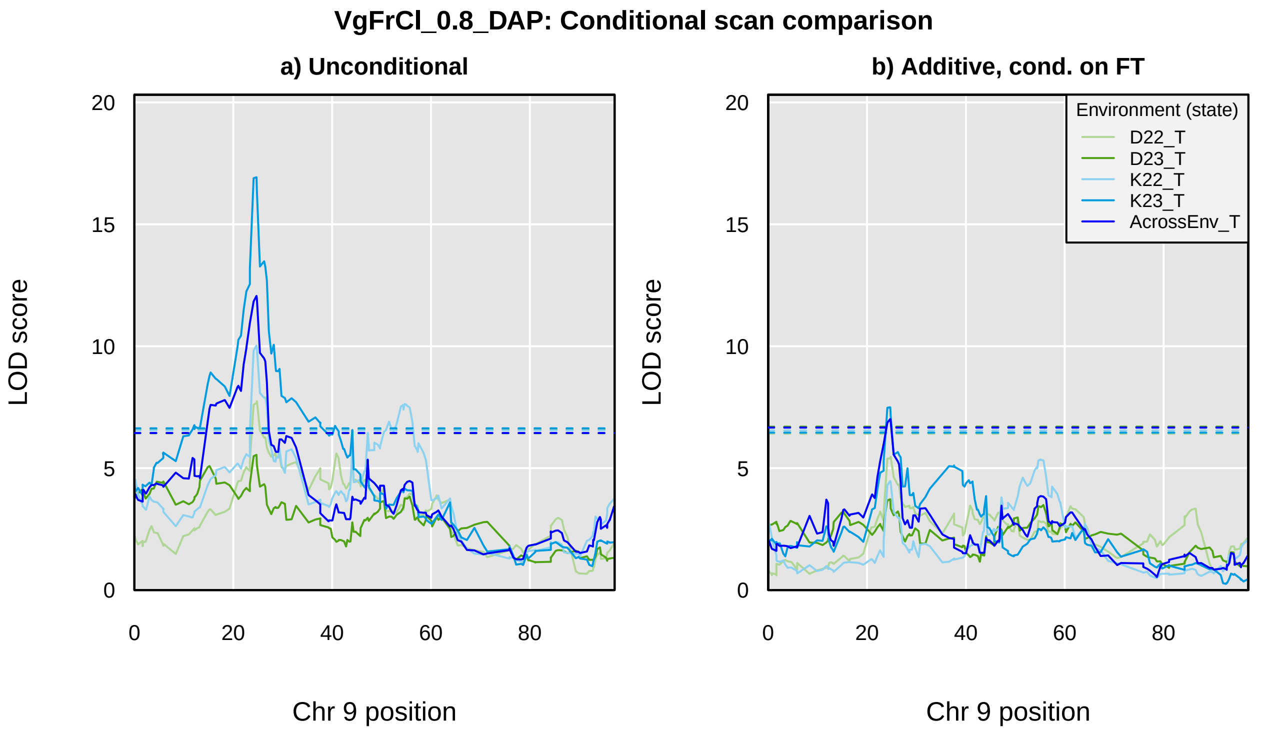

VgFrCl_0.8_DAP: Conditional scan comparison
a) Unconditional
b) Additive, cond. on FT
LOD score
Chr 9 position
0
5
10
15
20
0
20
40
60
80
Environment (state)
D22_T
D23_T
K22_T
K23_T
AcrossEnv_T

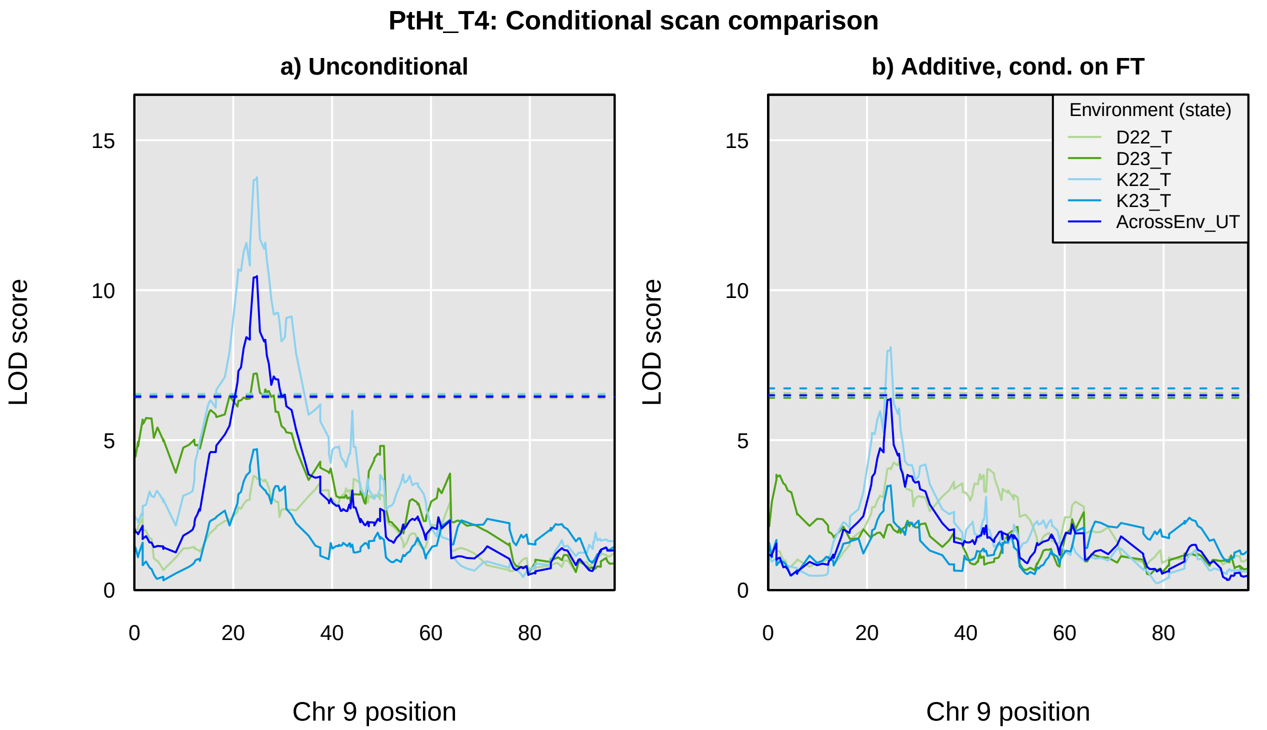
PtHt_T4: Conditional scan comparison
a) Unconditional
b) Additive, cond. on FT
LOD score
15
10
5
0
0
20
40
60
80
Chr 9 position
Environment (state)
D22_T
D23_T
K22_T
K23_T
AcrossEnv_UT

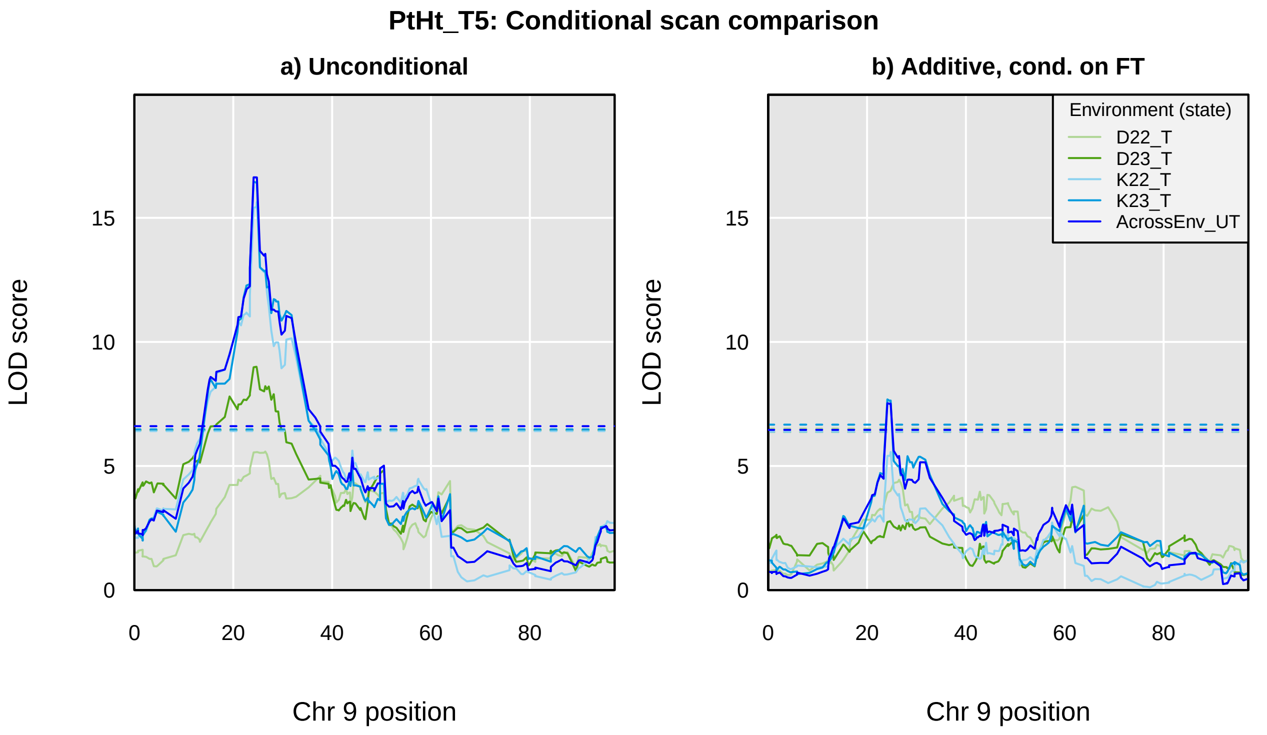
PtHt_T5: Conditional scan comparison
a) Unconditional
b) Additive, cond. on FT
LOD score
Chr 9 position
Environment (state)
D22_T
D23_T
K22_T
K23_T
AcrossEnv_UT
0
5
10
15
0
20
40
60
80

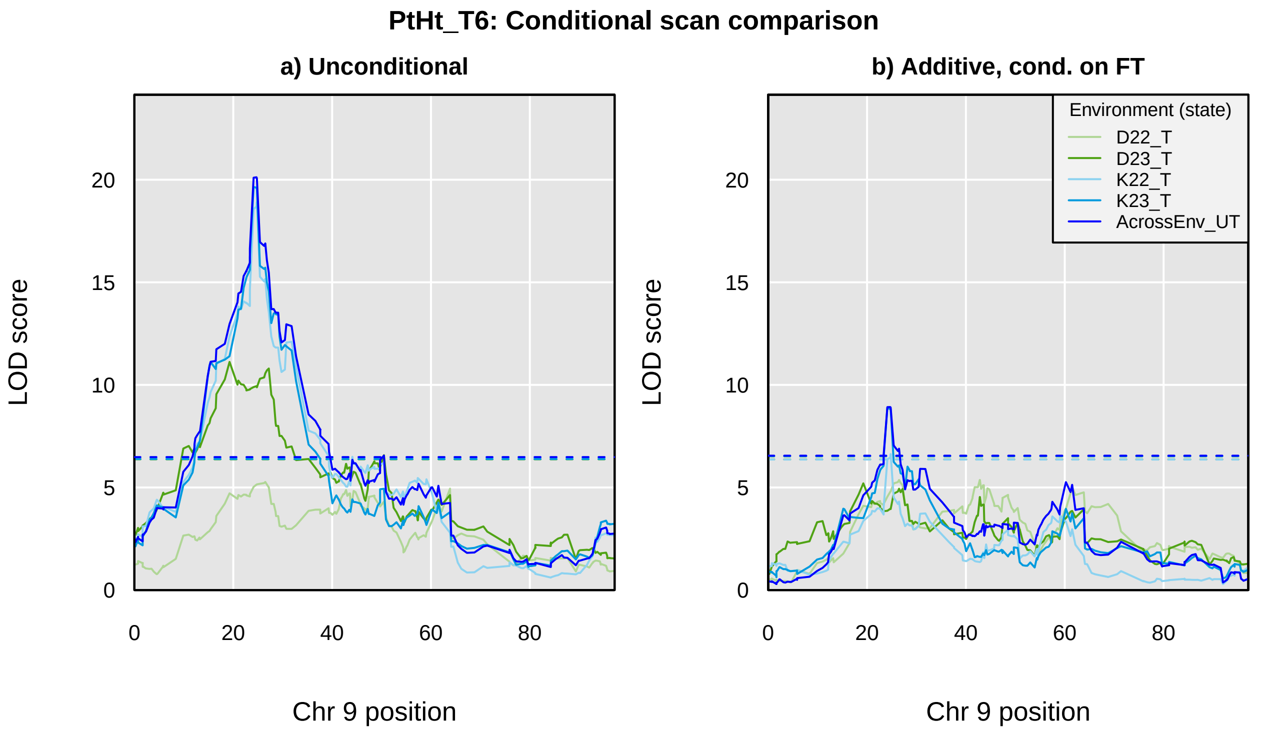
PtHt_T6: Conditional scan comparison
a) Unconditional
b) Additive, cond. on FT
LOD score
Chr 9 position
0
5
10
15
20
0
20
40
60
80
Environment (state)
D22_T
D23_T
K22_T
K23_T
AcrossEnv_UT

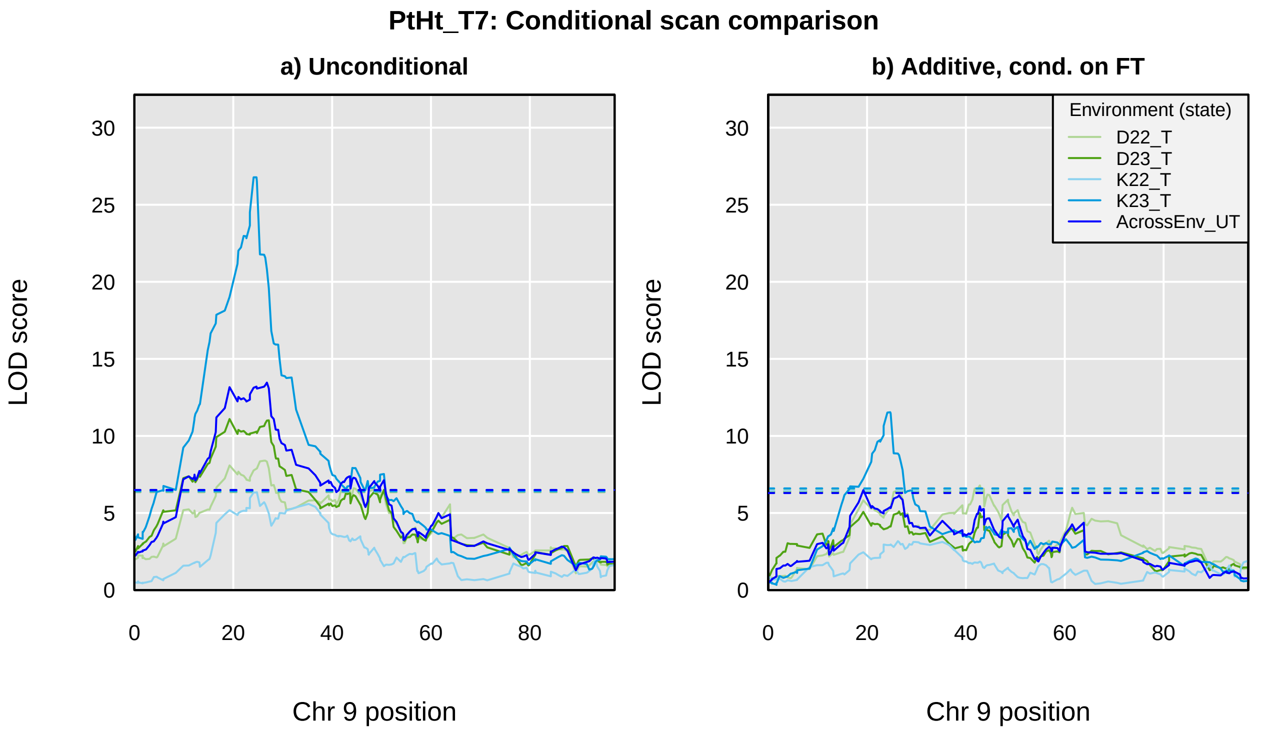
PtHt_T7: Conditional scan comparison
a) Unconditional
b) Additive, cond. on FT
LOD score
Chr 9 position
0
5
10
15
20
25
30
0
20
40
60
80
Environment (state)
D22_T
D23_T
K22_T
K23_T
AcrossEnv_UT

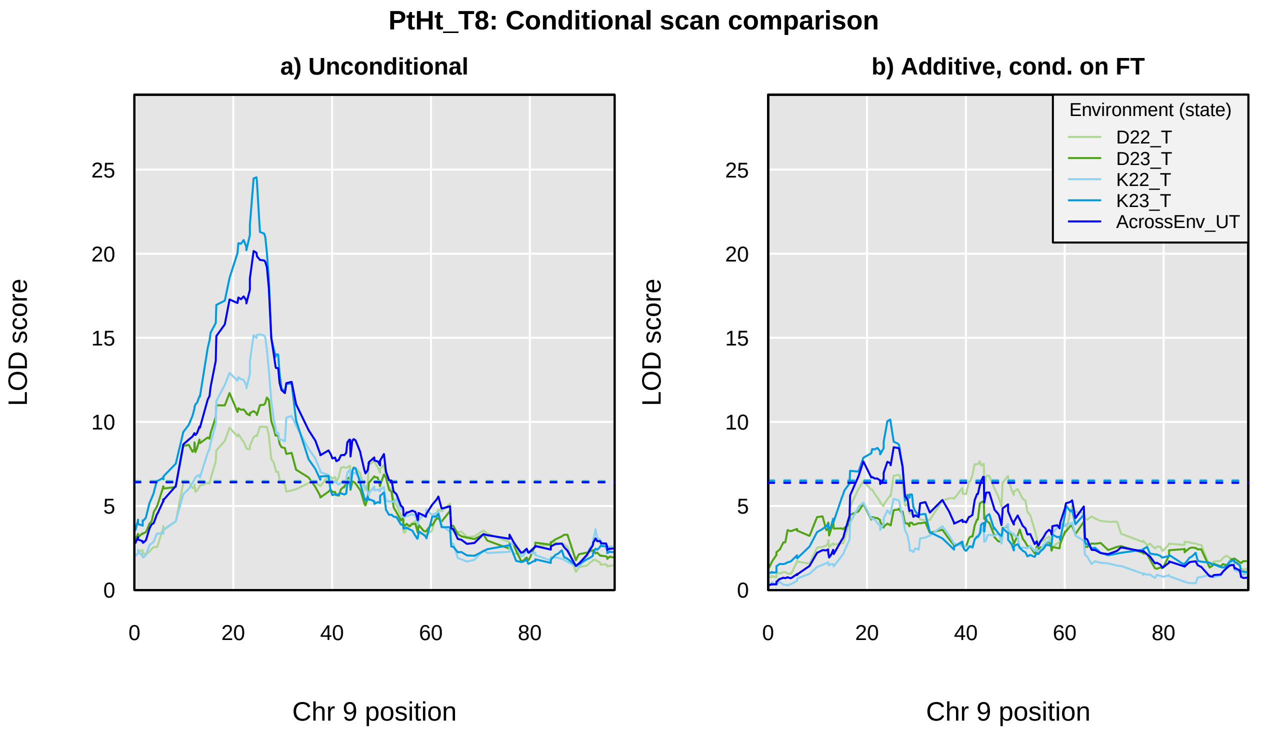

PtHt_T8: Conditional scan comparison
a) Unconditional
b) Additive, cond. on FT
LOD score
Chr 9 position
Environment (state)
D22_T
D23_T
K22_T
K23_T
AcrossEnv_UT
0
5
10
15
20
25
0
20
40
60
80

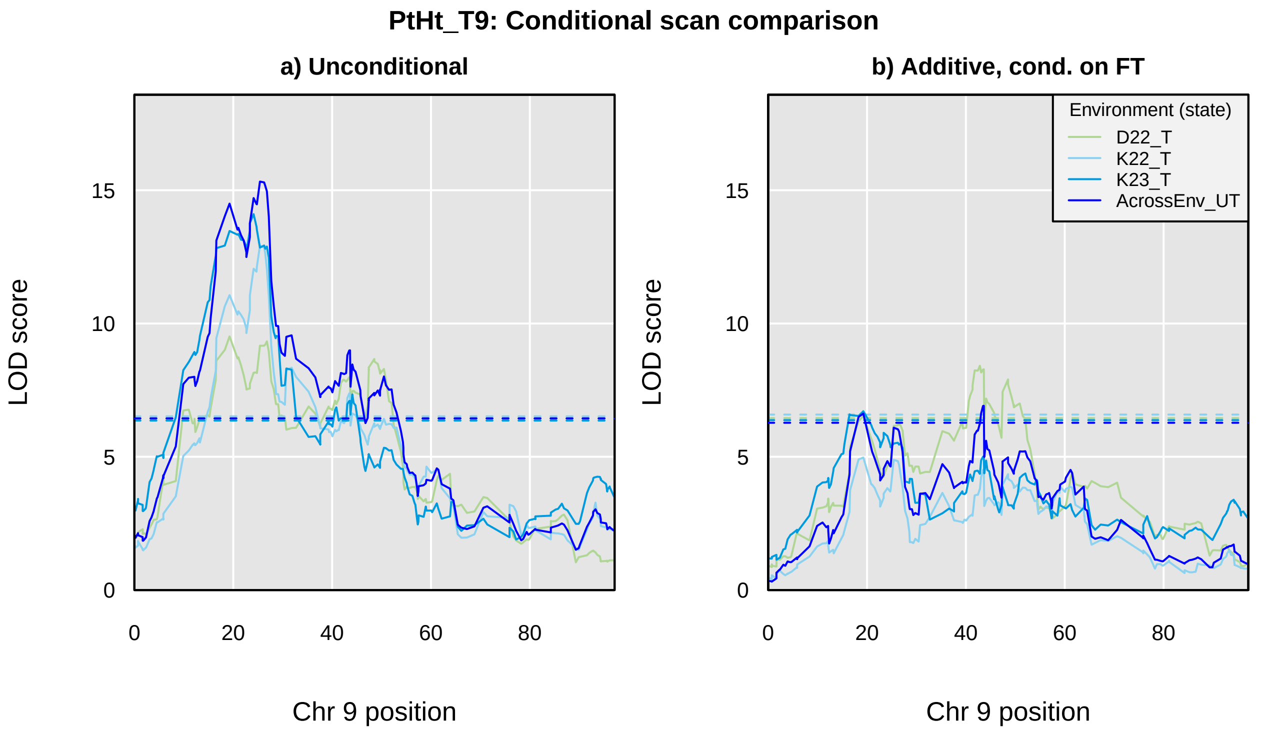
PtHt_T9: Conditional scan comparison
a) Unconditional
b) Additive, cond. on FT
LOD score
Chr 9 position
0
5
10
15
0
20
40
60
80
Environment (state)
D22_T
K22_T
K23_T
AcrossEnv_UT

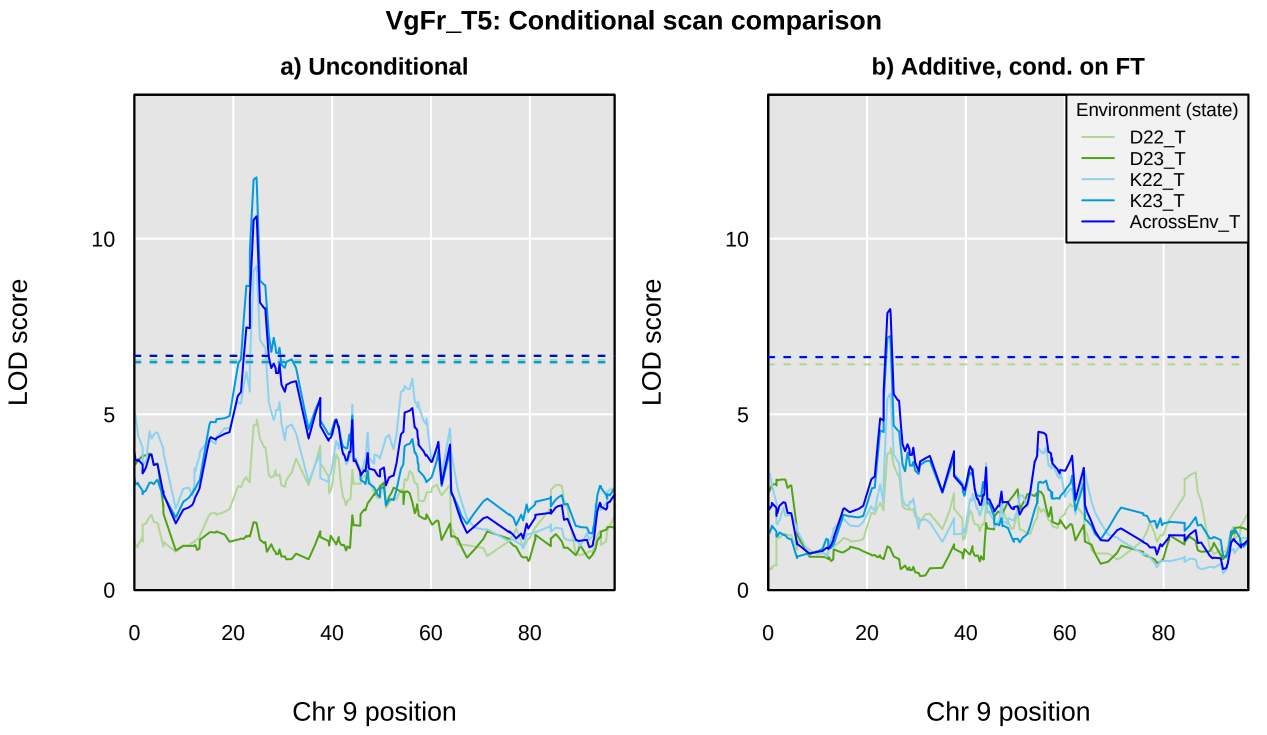
VgFr_T5: Conditional scan comparison
a) Unconditional
b) Additive, cond. on FT
LOD score
Chr 9 position
0
5
10
0
20
40
60
80
Environment (state)
D22_T
D23_T
K22_T
K23_T
AcrossEnv_T

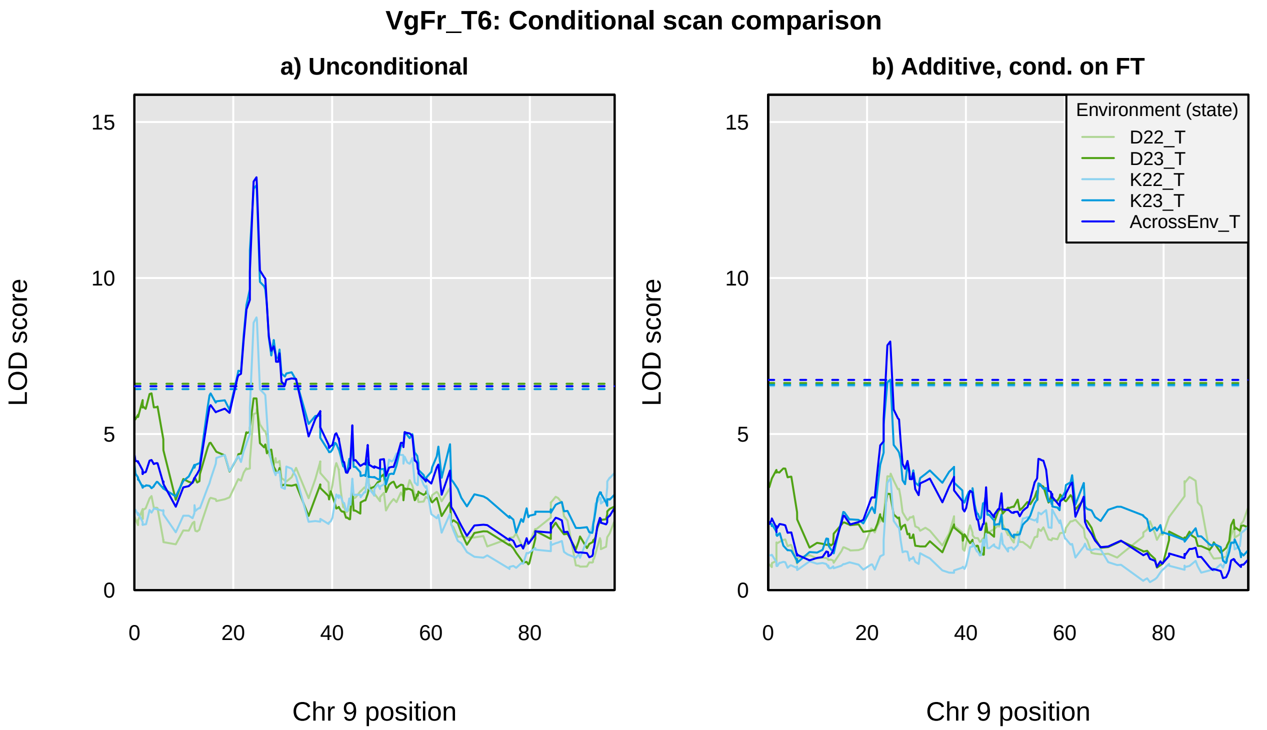
VgFr_T6: Conditional scan comparison
a) Unconditional
b) Additive, cond. on FT
LOD score
Chr 9 position
0
5
10
15
0
20
40
60
80
Environment (state)
D22_T
D23_T
K22_T
K23_T
AcrossEnv_T

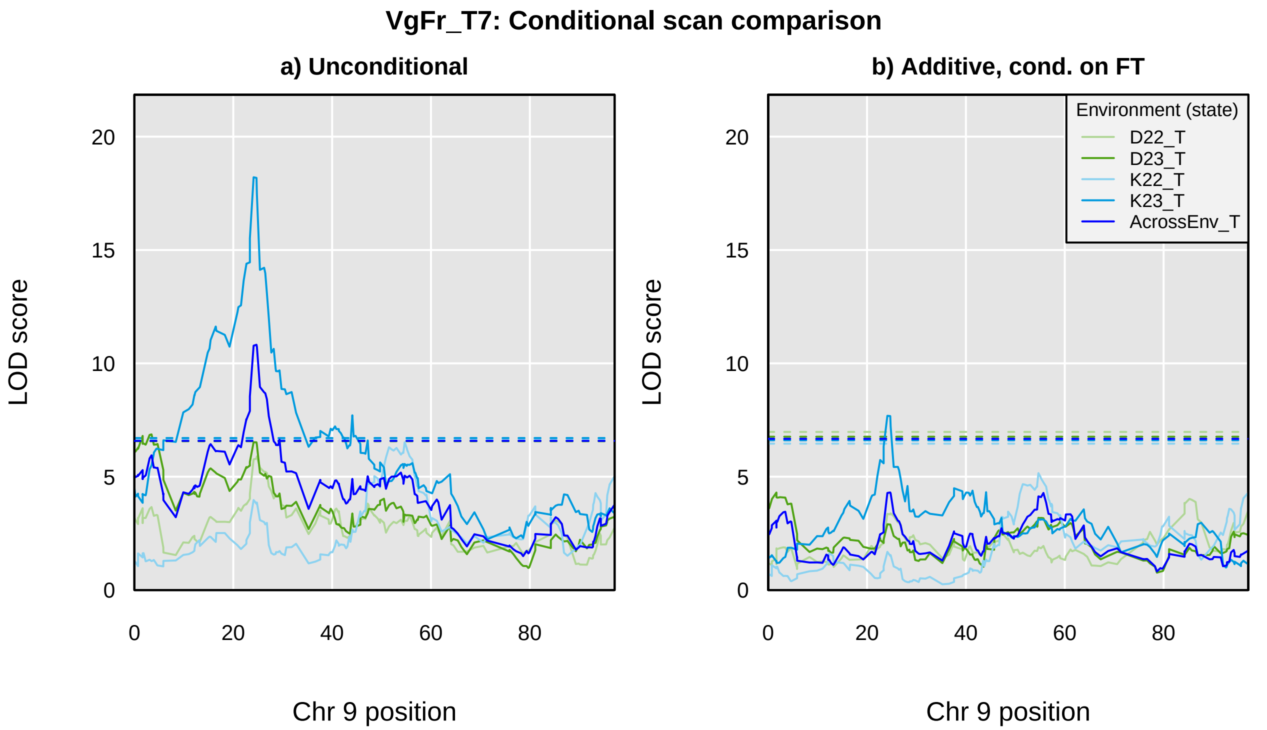
VgFr_T7: Conditional scan comparison
a) Unconditional
b) Additive, cond. on FT
LOD score
0
5
10
15
20
0
20
40
60
80
Chr 9 position
Environment (state)
D22_T
D23_T
K22_T
K23_T
AcrossEnv_T

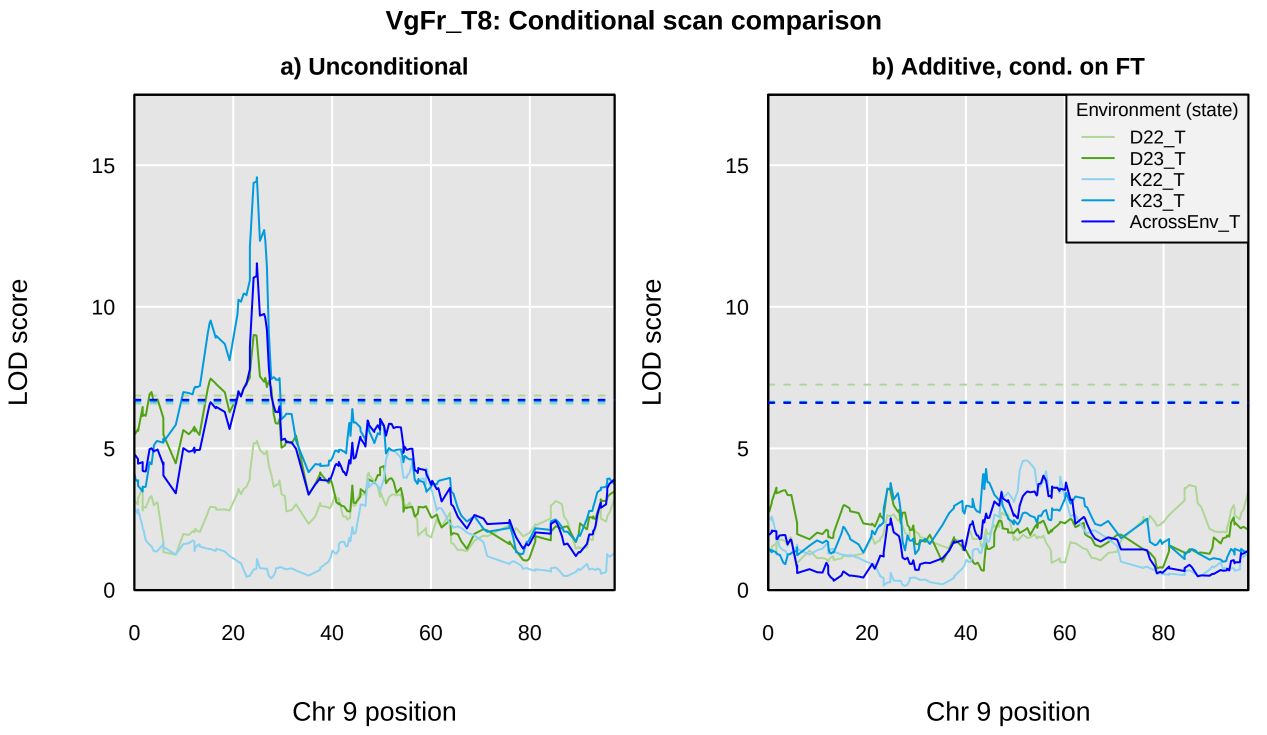
VgFr_T8: Conditional scan comparison
a) Unconditional
b) Additive, cond. on FT
LOD score
Chr 9 position
0
5
10
15
0
20
40
60
80
Environment (state)
D22_T
D23_T
K22_T
K23_T
AcrossEnv_T

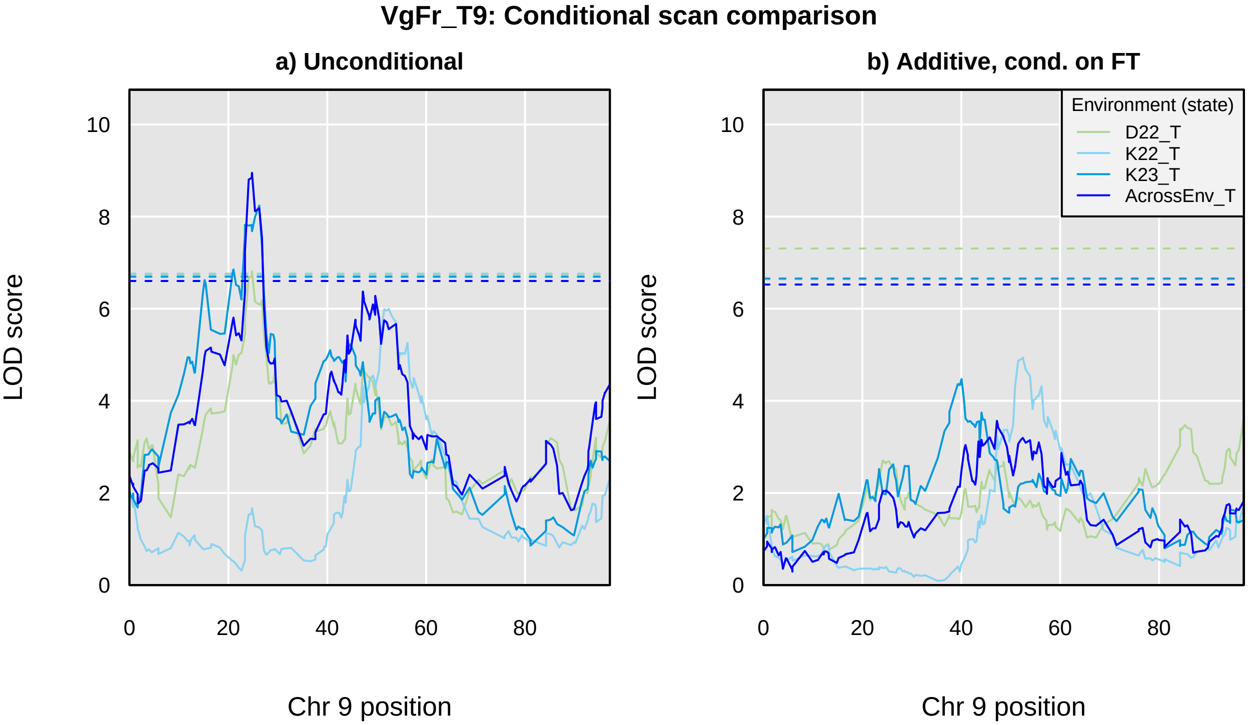
VgFr_T9: Conditional scan comparison
a) Unconditional
b) Additive, cond. on FT
Environment (state)
D22_T
K22_T
K23_T
AcrossEnv_T
LOD score
Chr 9 position
0
2
4
6
8
10
0
20
40
60
80
