## Supplementary material for "Genetic mapping and genomic prediction for agronomic, grain compositional, and sensing-enabled traits in a cowpea MAGIC population along an environmental gradient": Large supplemental tables and figures, uploaded separately: SuppFig_11_AcrossEnv_QTLprofiles_FTconditional.pdf

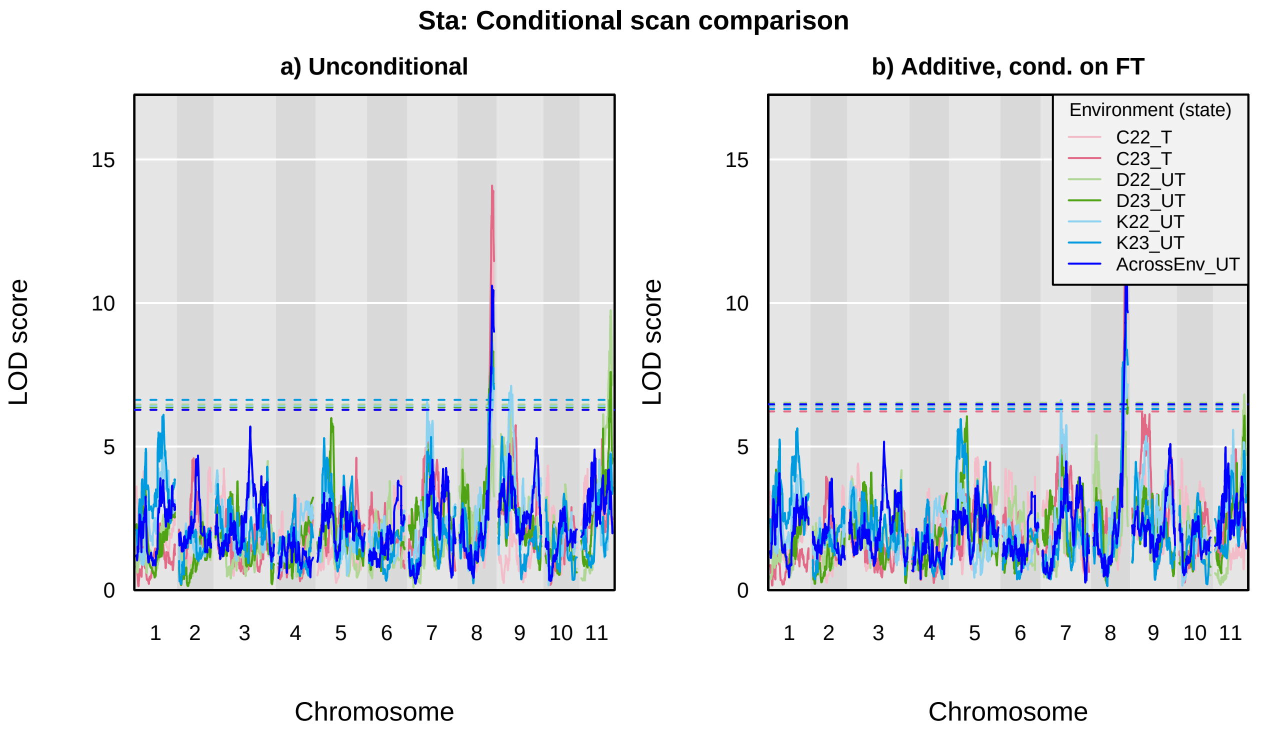

Sta: Conditional scan comparison
a) Unconditional
b) Additive, cond. on FT
LOD score
0
5
10
15
Chromosome
1
2
3
4
5
6
7
8
9
10
11
Environment (state)
C22_T
C23_T
D22_UT
D23_UT
K22_UT
K23_UT
AcrossEnv_UT

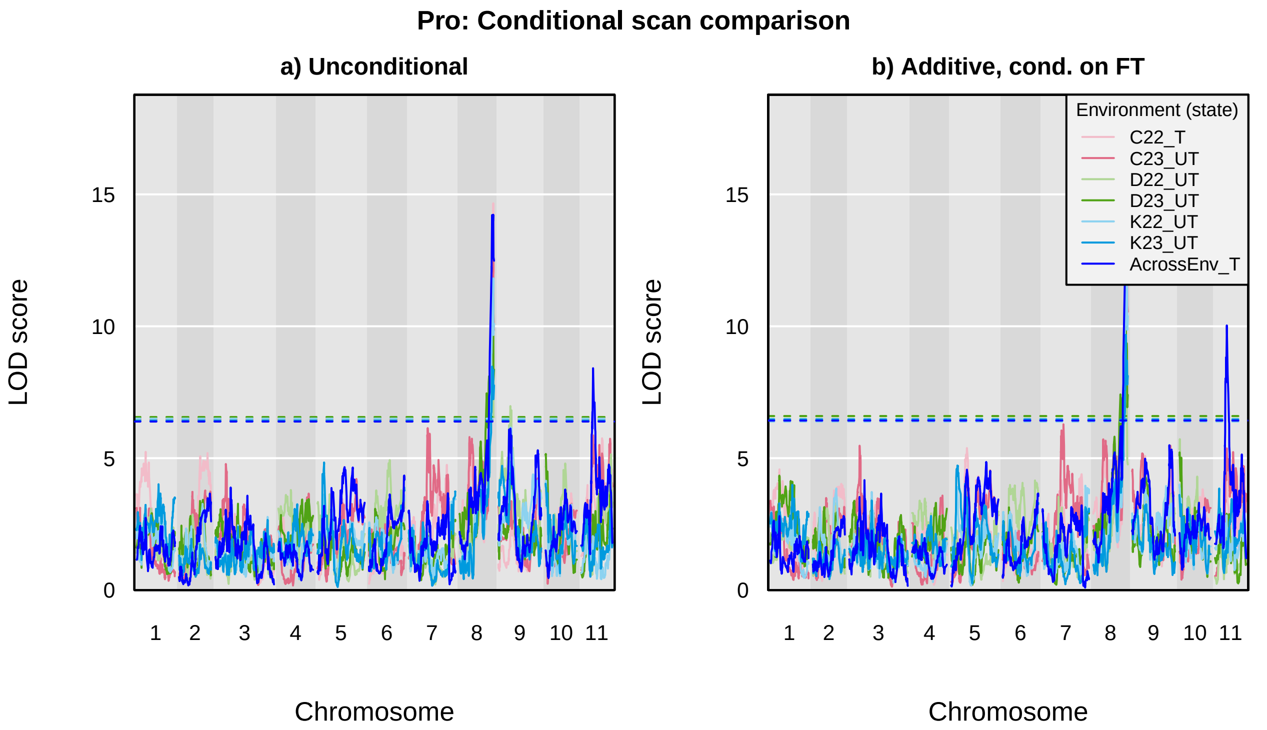

Pro: Conditional scan comparison
a) Unconditional
b) Additive, cond. on FT
LOD score
Chromosome
0
5
10
15
1
2
3
4
5
6
7
8
9
10
11
Environment (state)
C22_T
C23_UT
D22_UT
D23_UT
K22_UT
K23_UT
AcrossEnv_T

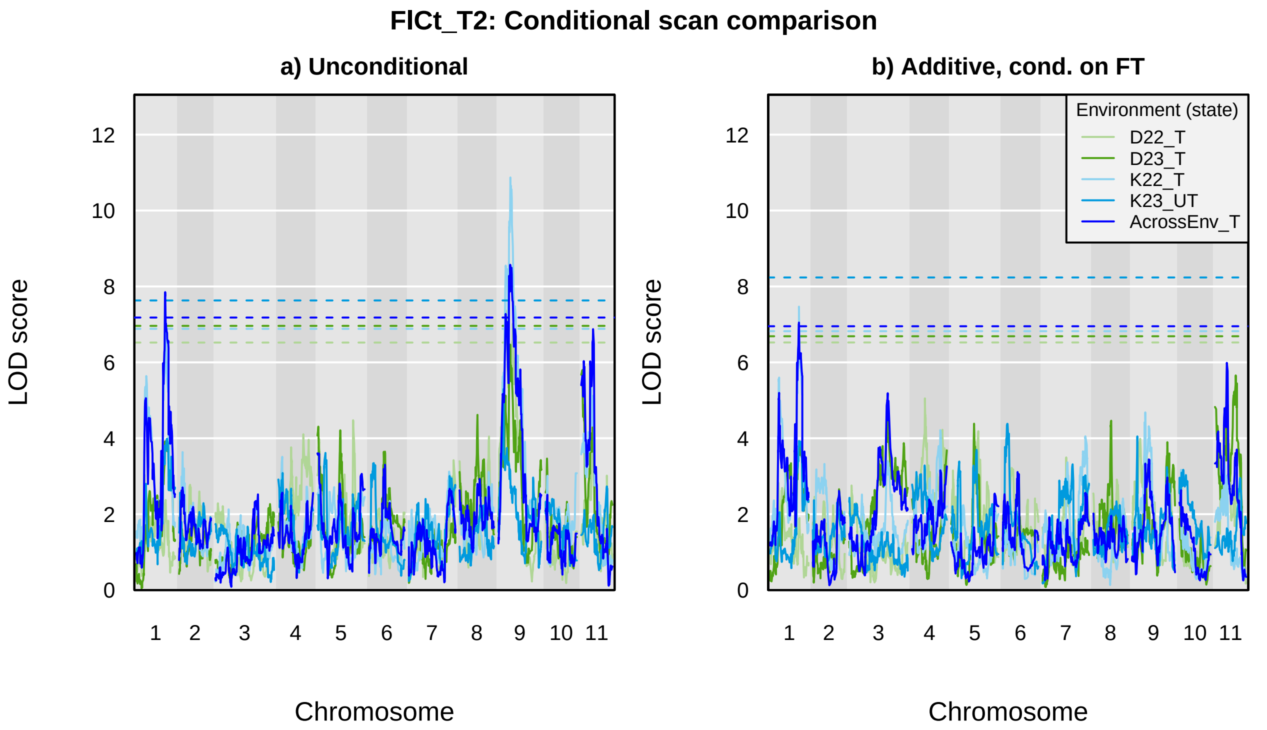
FlCt_T2: Conditional scan comparison
a) Unconditional
b) Additive, cond. on FT
LOD score
Chromosome
0
2
4
6
8
10
12
1
2
3
4
5
6
7
8
9
10
11
Environment (state)
D22_T
D23_T
K22_T
K23_UT
AcrossEnv_T

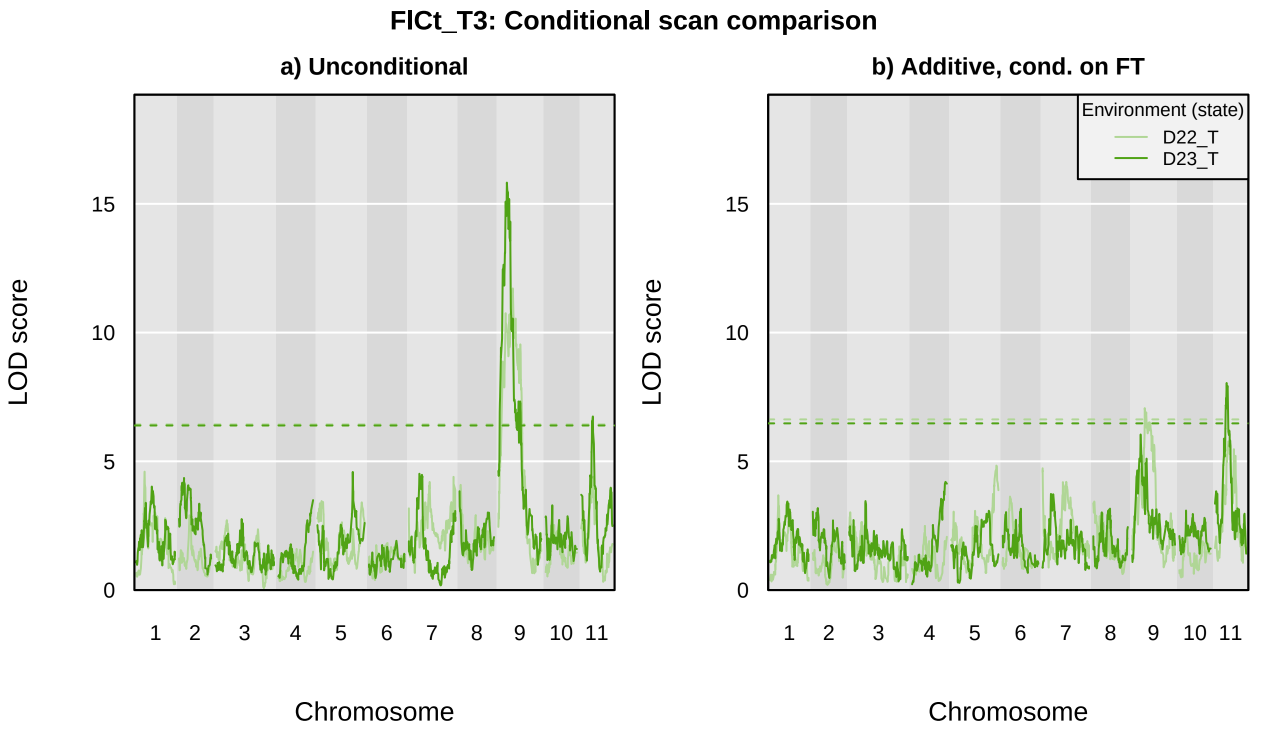
FlCt_T3: Conditional scan comparison
a) Unconditional
b) Additive, cond. on FT
Environment (state)
D22_T
D23_T
LOD score
0
5
10
15
Chromosome
1
2
3
4
5
6
7
8
9
10
11

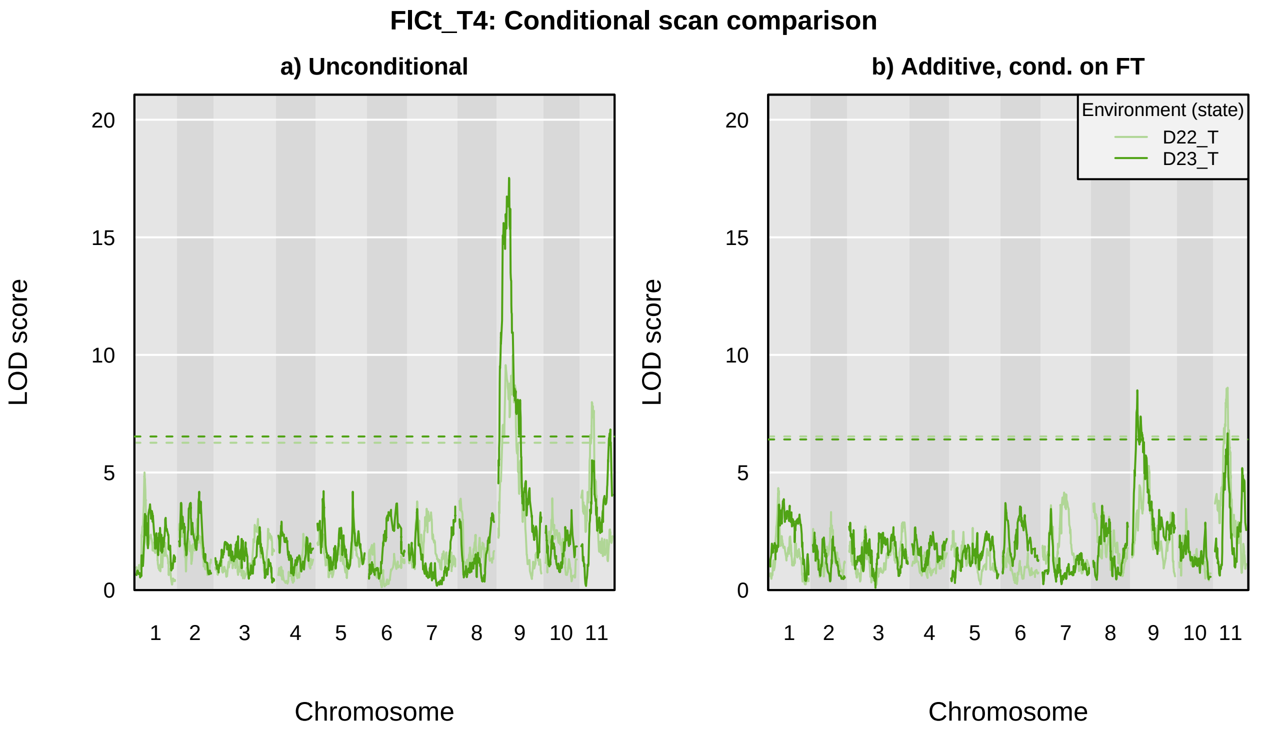

FlCt_T4: Conditional scan comparison
a) Unconditional
b) Additive, cond. on FT
Environment (state)
D22_T
D23_T
LOD score
Chromosome
0
5
10
15
20
1
2
3
4
5
6
7
8
9
10
11

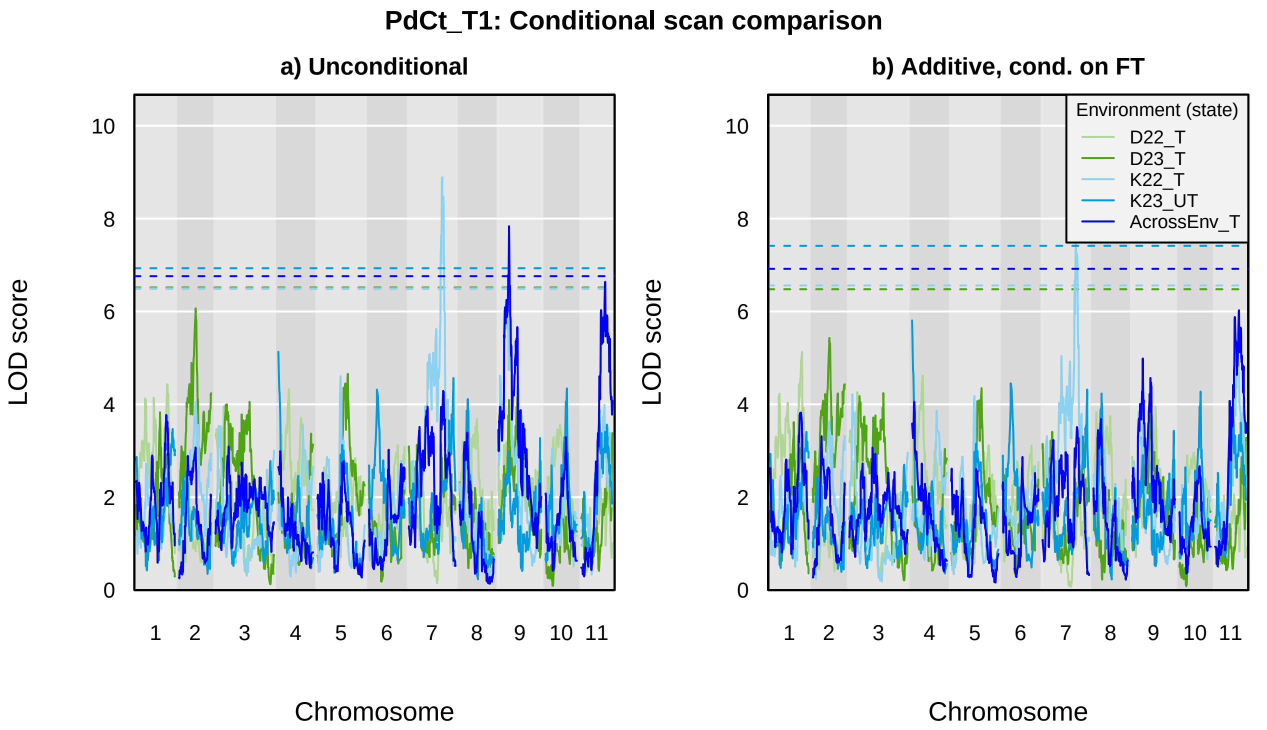
PdCt_T1: Conditional scan comparison
a) Unconditional
b) Additive, cond. on FT
LOD score
Chromosome
0
2
4
6
8
10
1
2
3
4
5
6
7
8
9
10
11
Environment (state)
D22_T
D23_T
K22_T
K23_UT
AcrossEnv_T

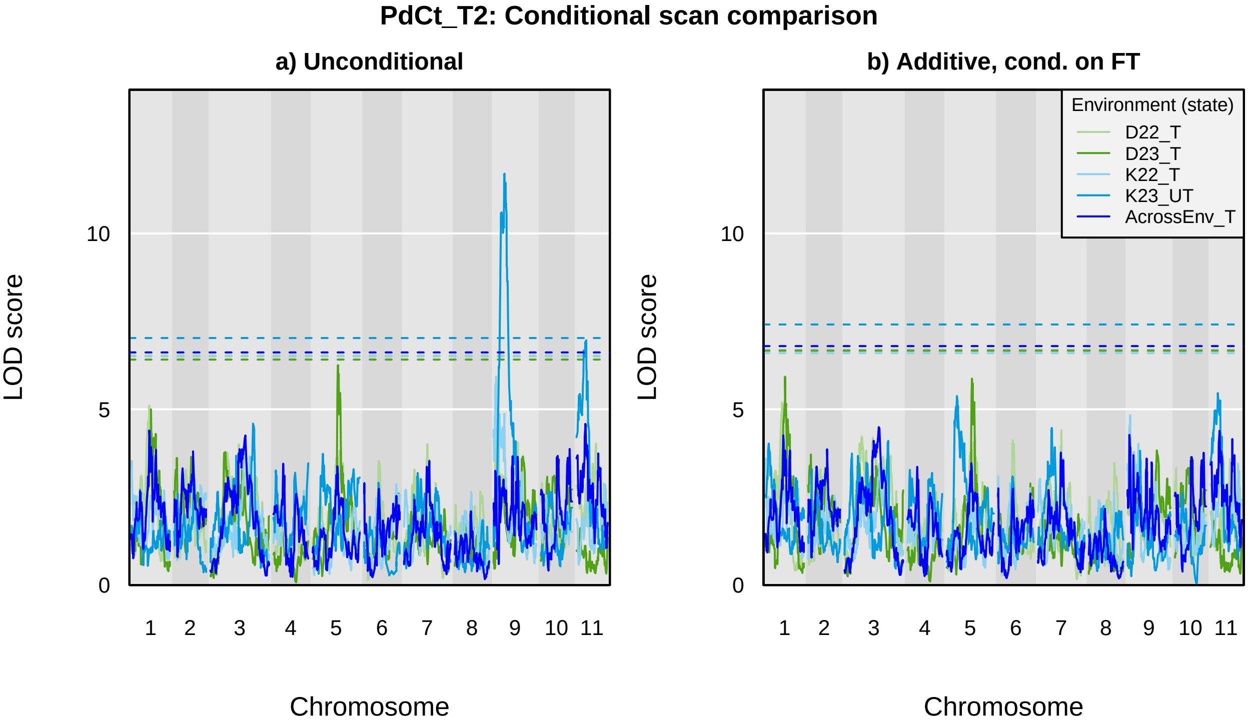
PdCt_T2: Conditional scan comparison
a) Unconditional
b) Additive, cond. on FT
Environment (state)
D22_T
D23_T
K22_T
K23_UT
AcrossEnv_T
LOD score
Chromosome
0
5
10
1
2
3
4
5
6
7
8
9
10
11

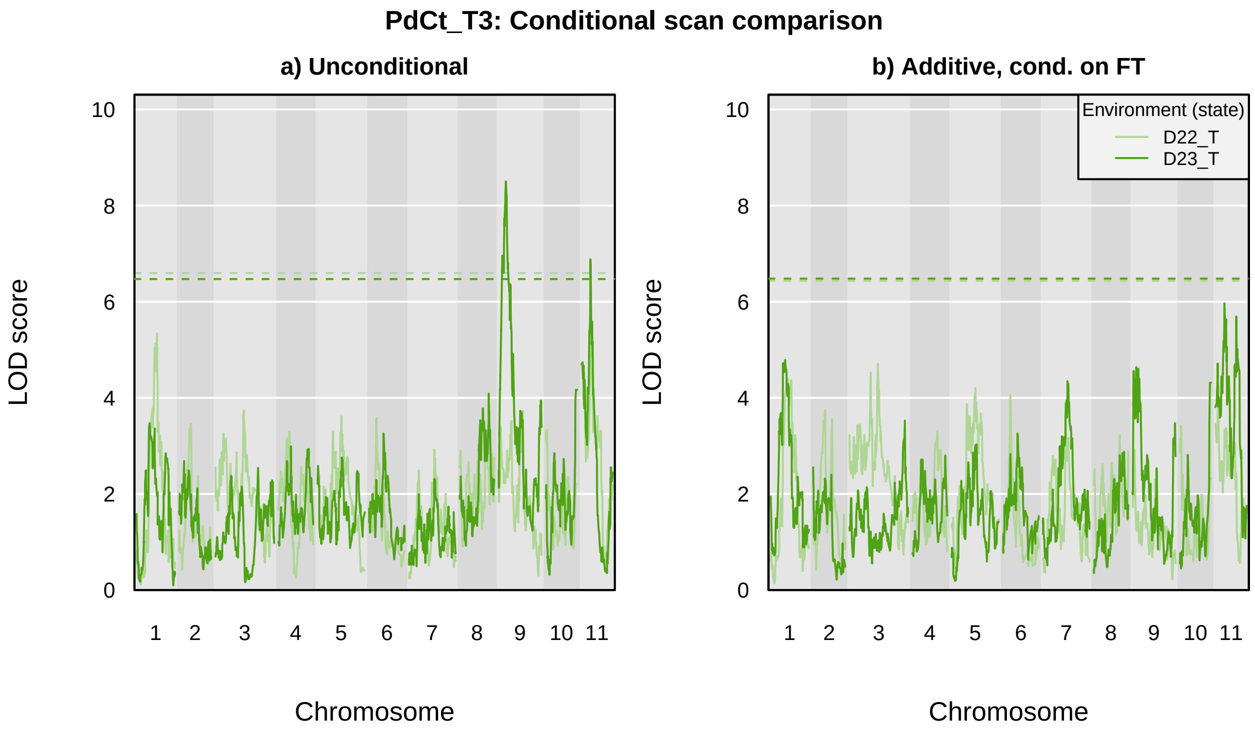
PdCt_T3: Conditional scan comparison
a) Unconditional
b) Additive, cond. on FT
LOD score
Chromosome
0
2
4
6
8
10
1
2
3
4
5
6
7
8
9
10
11
Environment (state)
D22_T
D23_T

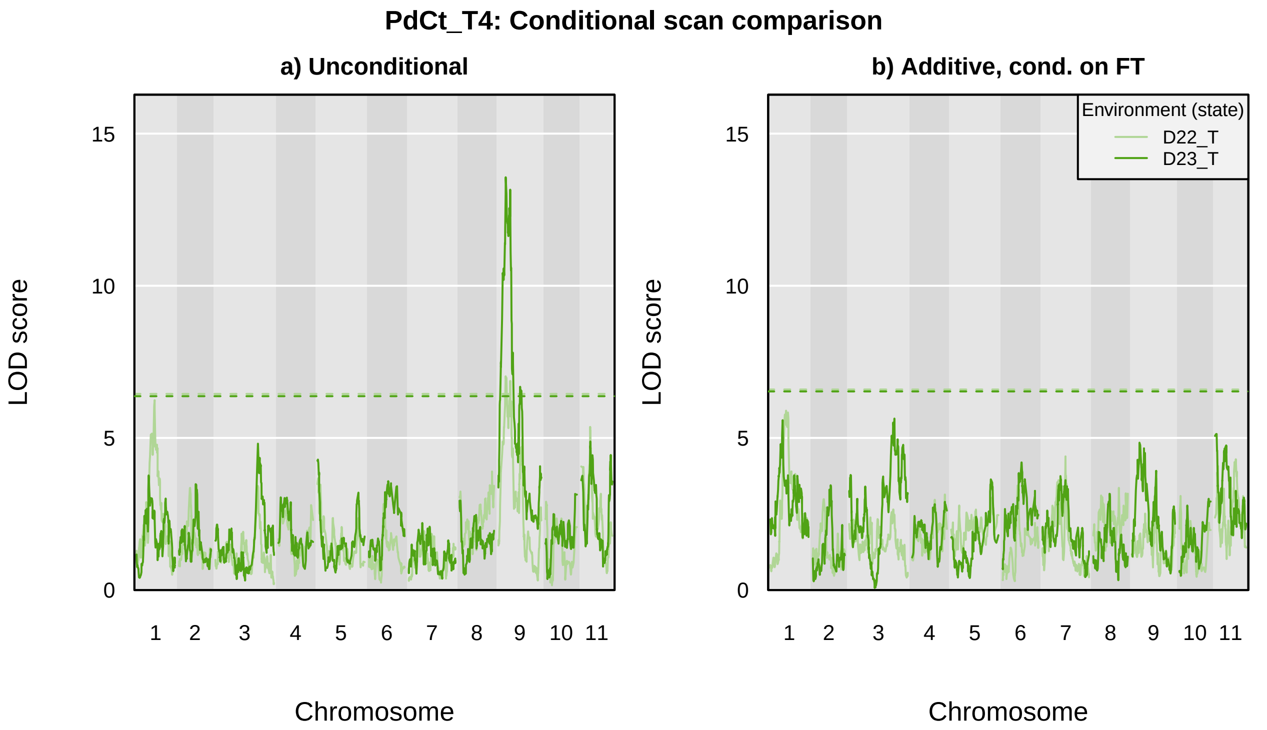

PdCt_T4: Conditional scan comparison
a) Unconditional
b) Additive, cond. on FT
LOD score
0
5
10
15
1
2
3
4
5
6
7
8
9
10
11
Chromosome
Environment (state)
D22_T
D23_T

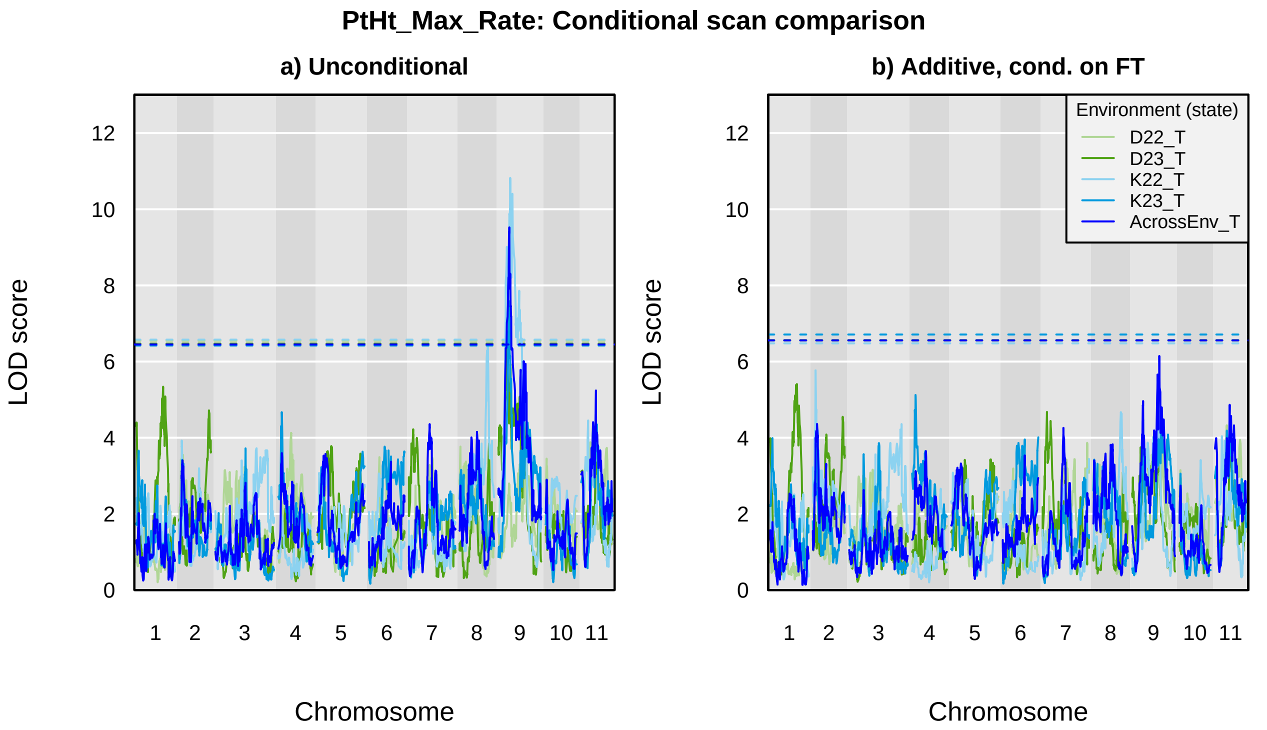
PtHt_Max_Rate: Conditional scan comparison
a) Unconditional
b) Additive, cond. on FT
LOD score
Chromosome
0
2
4
6
8
10
12
1
2
3
4
5
6
7
8
9
10
11
Environment (state)
D22_T
D23_T
K22_T
K23_T
AcrossEnv_T

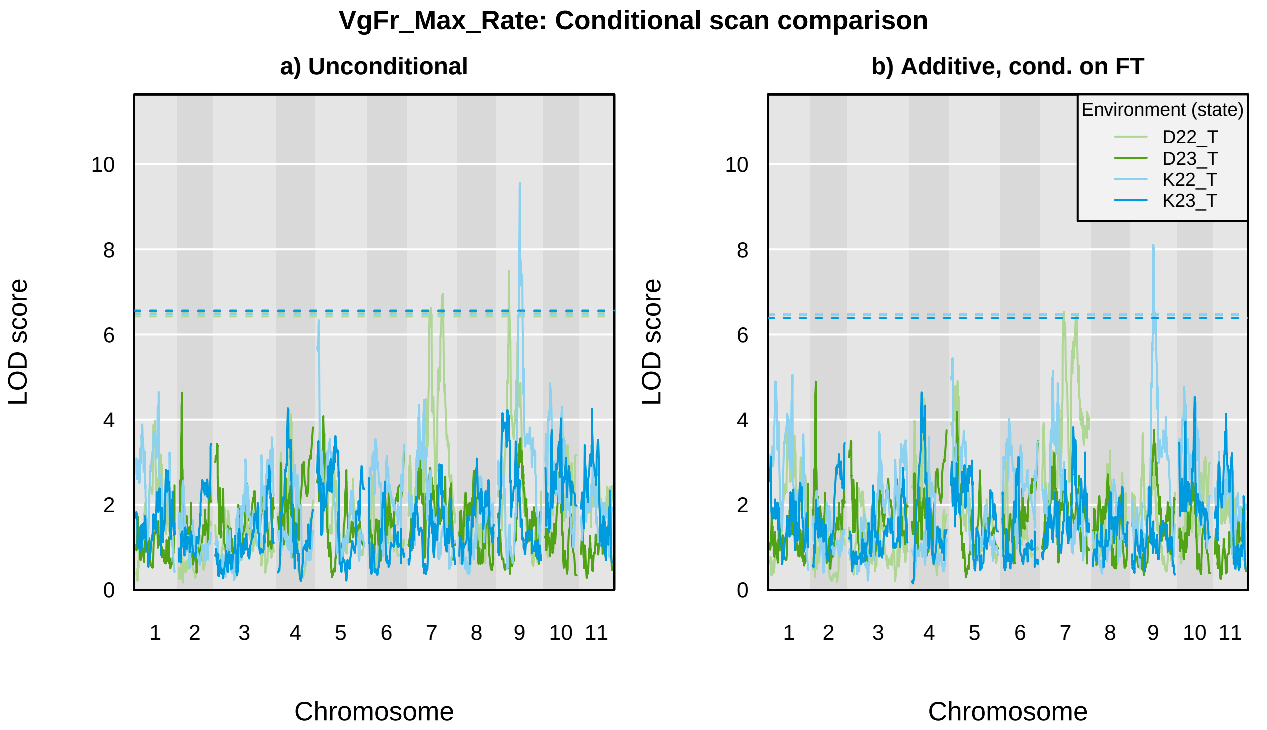
VgFr_Max_Rate: Conditional scan comparison
a) Unconditional
b) Additive, cond. on FT
Environment (state)
D22_T
D23_T
K22_T
K23_T
LOD score
0
2
4
6
8
10
Chromosome
1
2
3
4
5
6
7
8
9
10
11

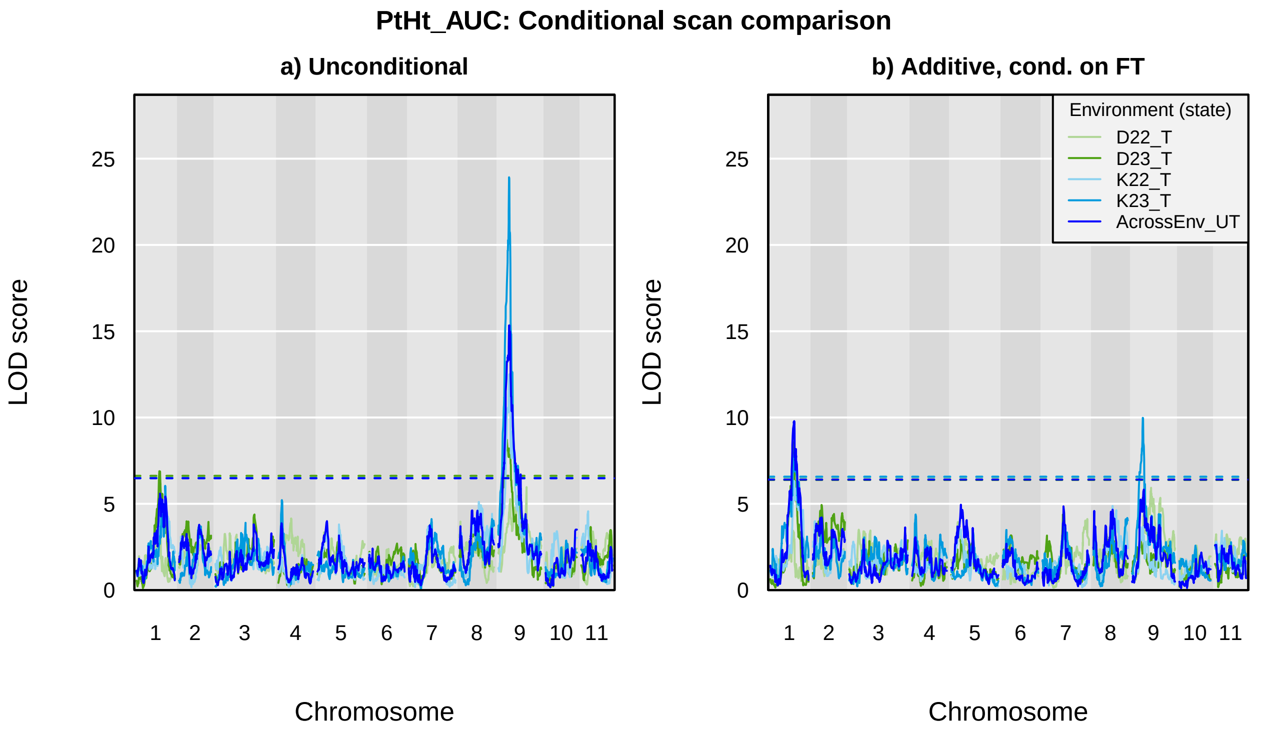

PtHt_AUC: Conditional scan comparison
a) Unconditional
b) Additive, cond. on FT
LOD score
Chromosome
Environment (state)
D22_T
D23_T
K22_T
K23_T
AcrossEnv_UT
0
5
10
15
20
25
1
2
3
4
5
6
7
8
9
10
11

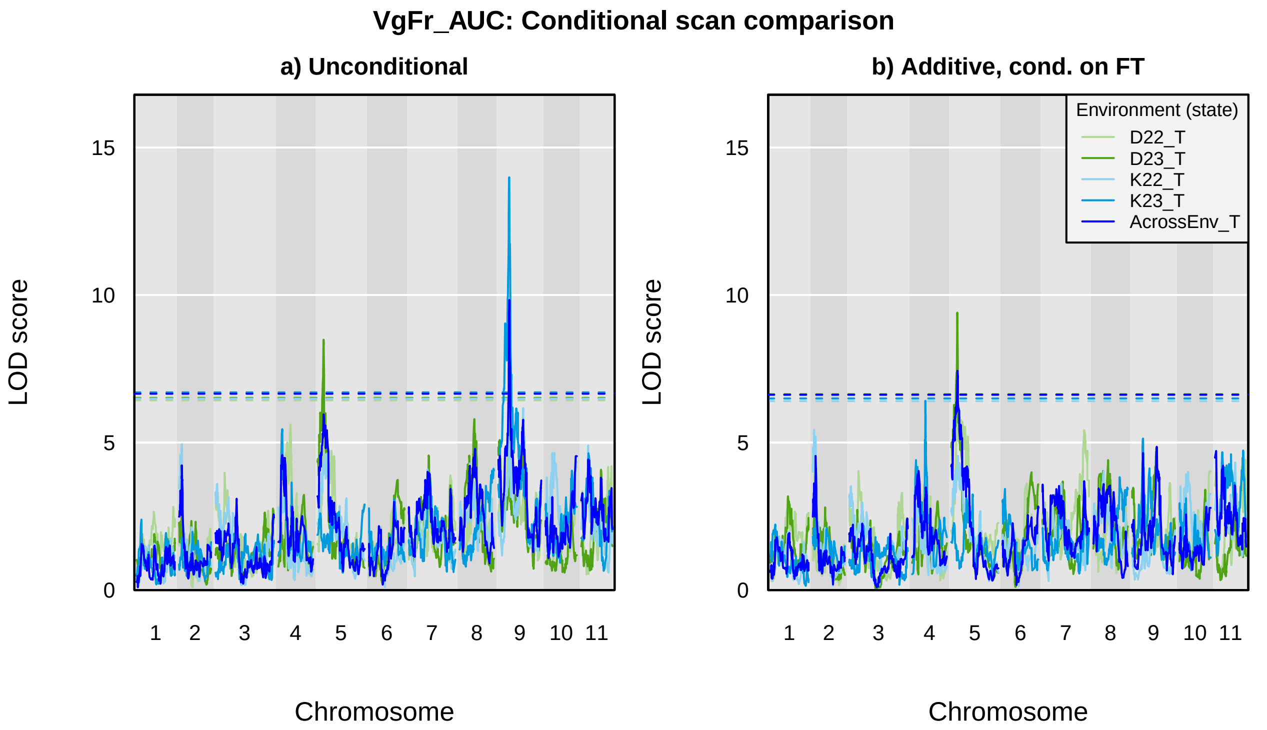

VgFr_AUC: Conditional scan comparison
a) Unconditional
b) Additive, cond. on FT
LOD score
0
5
10
15
Chromosome
1 2 3 4 5 6 7 8 9 10 11
Environment (state)
D22_T
D23_T
K22_T
K23_T
AcrossEnv_T

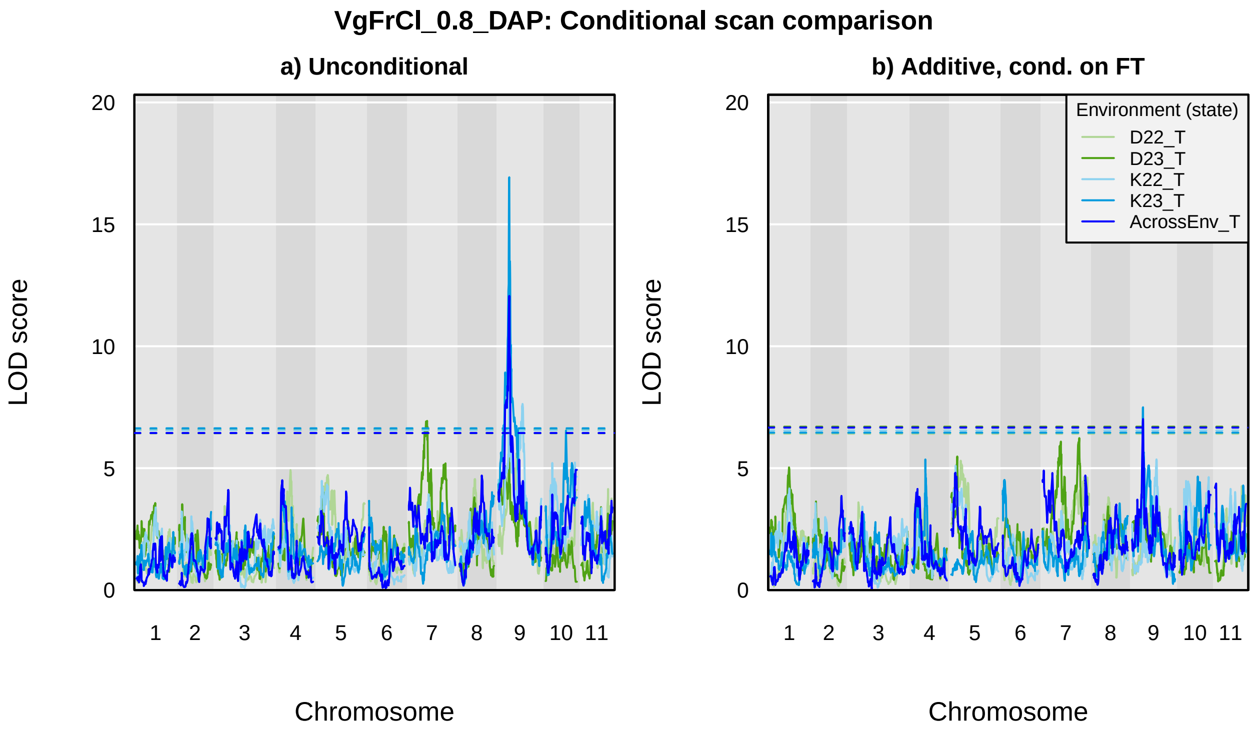
VgFrCl_0.8_DAP: Conditional scan comparison
a) Unconditional
b) Additive, cond. on FT
LOD score
0
5
10
15
20
Chromosome
1
2
3
4
5
6
7
8
9
10
11
Environment (state)
D22_T
D23_T
K22_T
K23_T
AcrossEnv_T

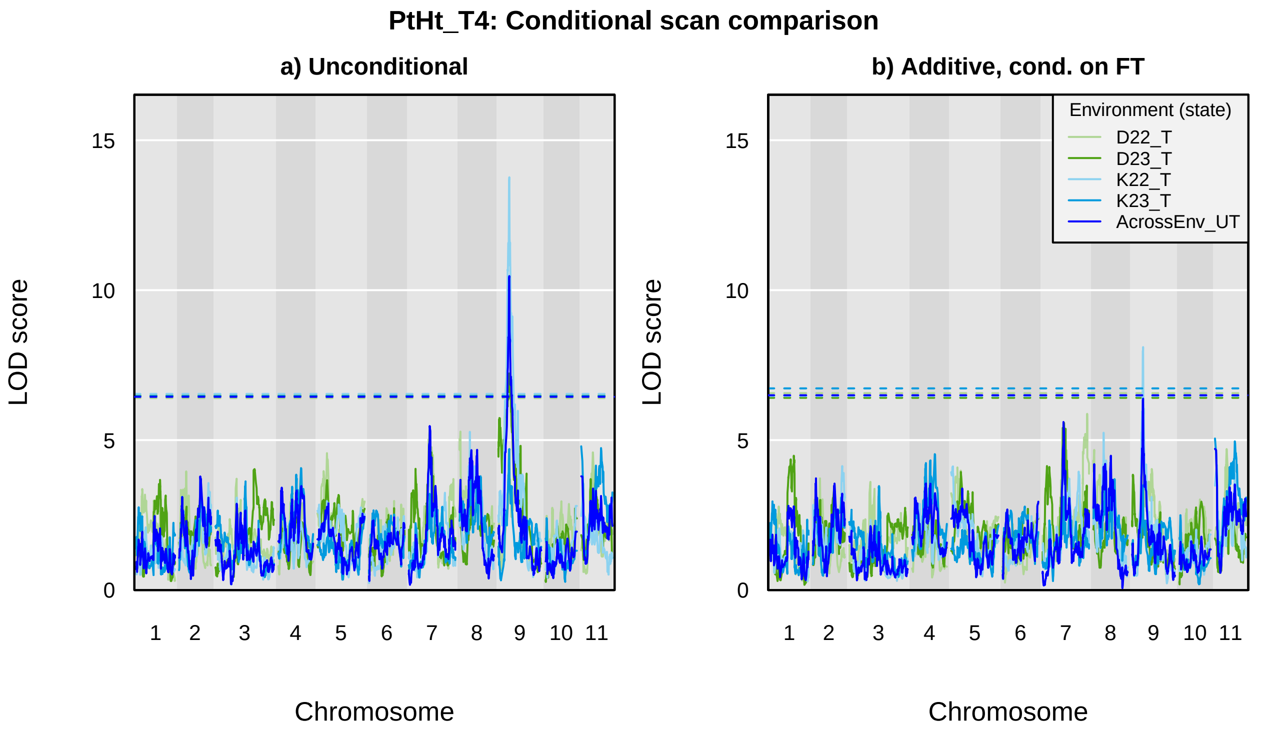
PtHt_T4: Conditional scan comparison
a) Unconditional
b) Additive, cond. on FT
LOD score
Chromosome
0
5
10
15
1
2
3
4
5
6
7
8
9
10
11
Environment (state)
D22_T
D23_T
K22_T
K23_T
AcrossEnv_UT

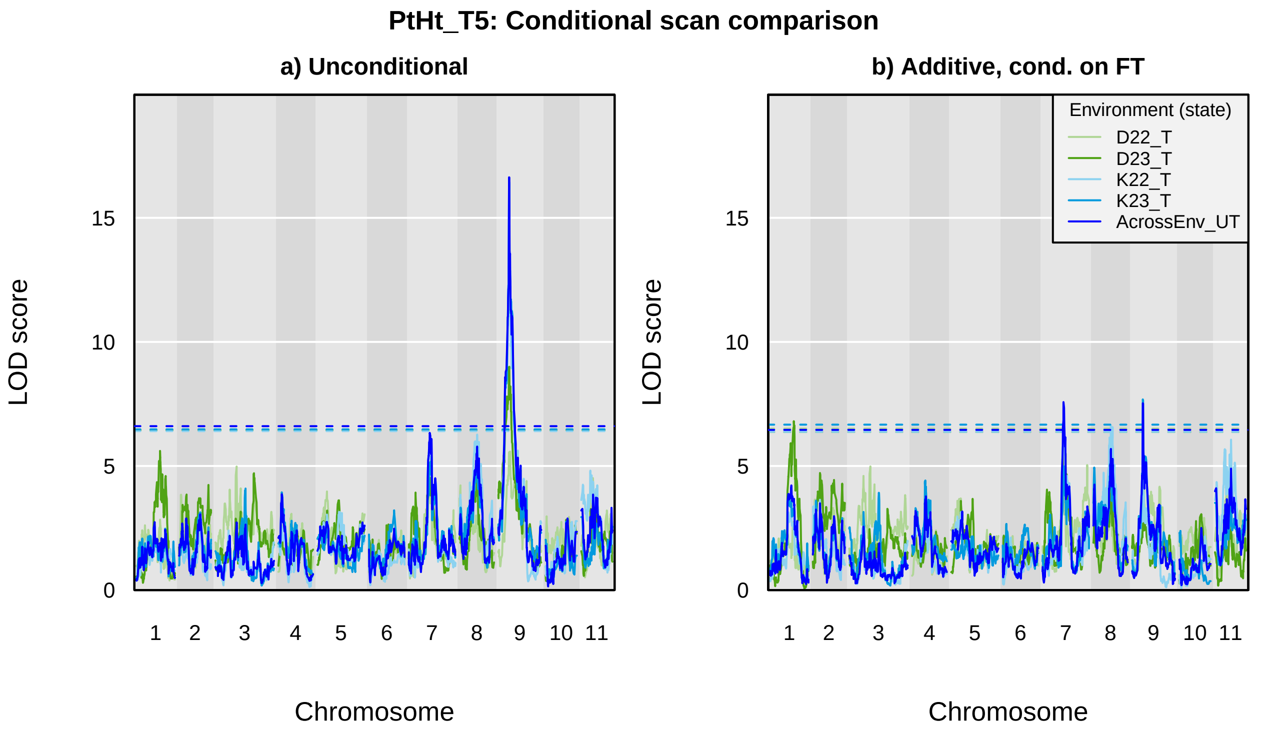

PtHt_T5: Conditional scan comparison
a) Unconditional
b) Additive, cond. on FT
LOD score
Chromosome
0
5
10
15
1
2
3
4
5
6
7
8
9
10
11
Environment (state)
D22_T
D23_T
K22_T
K23_T
AcrossEnv_UT

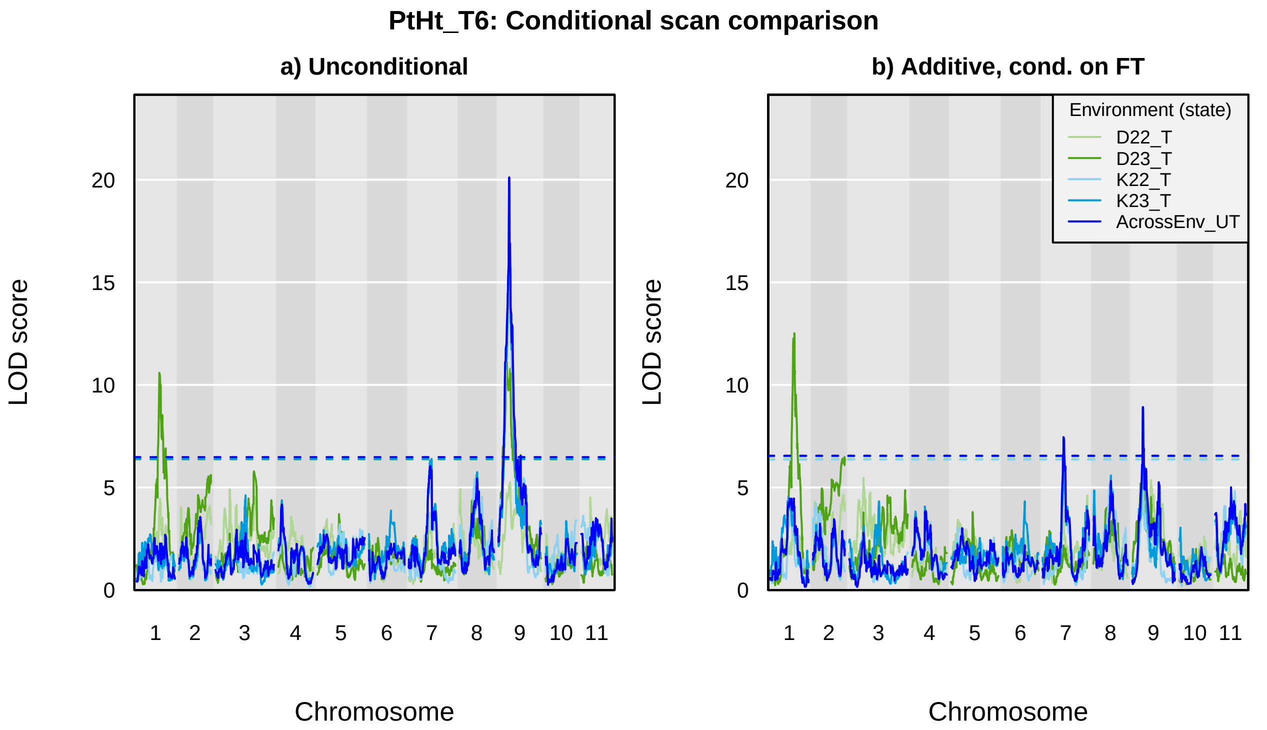
PtHt_T6: Conditional scan comparison
a) Unconditional
b) Additive, cond. on FT
Environment (state)
D22_T
D23_T
K22_T
K23_T
AcrossEnv_UT
LOD score
Chromosome
0
5
10
15
20
1 2 3 4 5 6 7 8 9 10 11

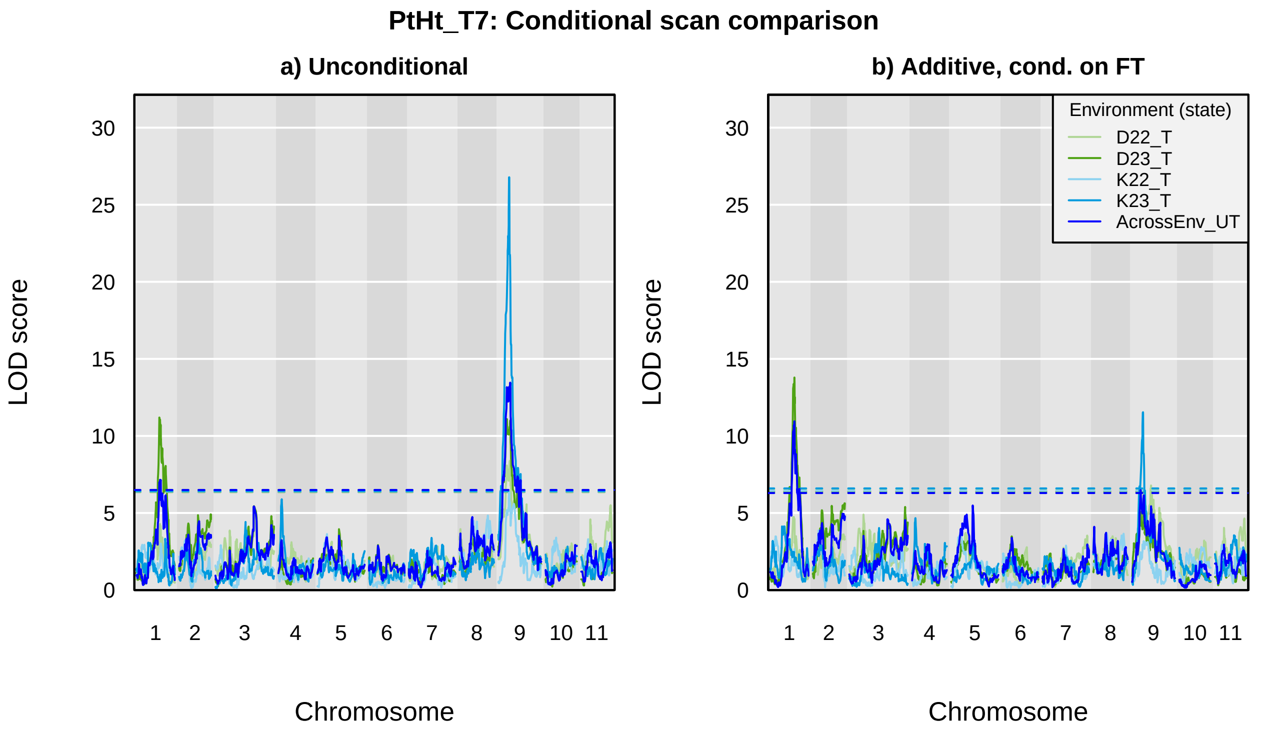

PtHt_T7: Conditional scan comparison
a) Unconditional
b) Additive, cond. on FT
LOD score
Chromosome
0
5
10
15
20
25
30
1
2
3
4
5
6
7
8
9
10
11
Environment (state)
D22_T
D23_T
K22_T
K23_T
AcrossEnv_UT

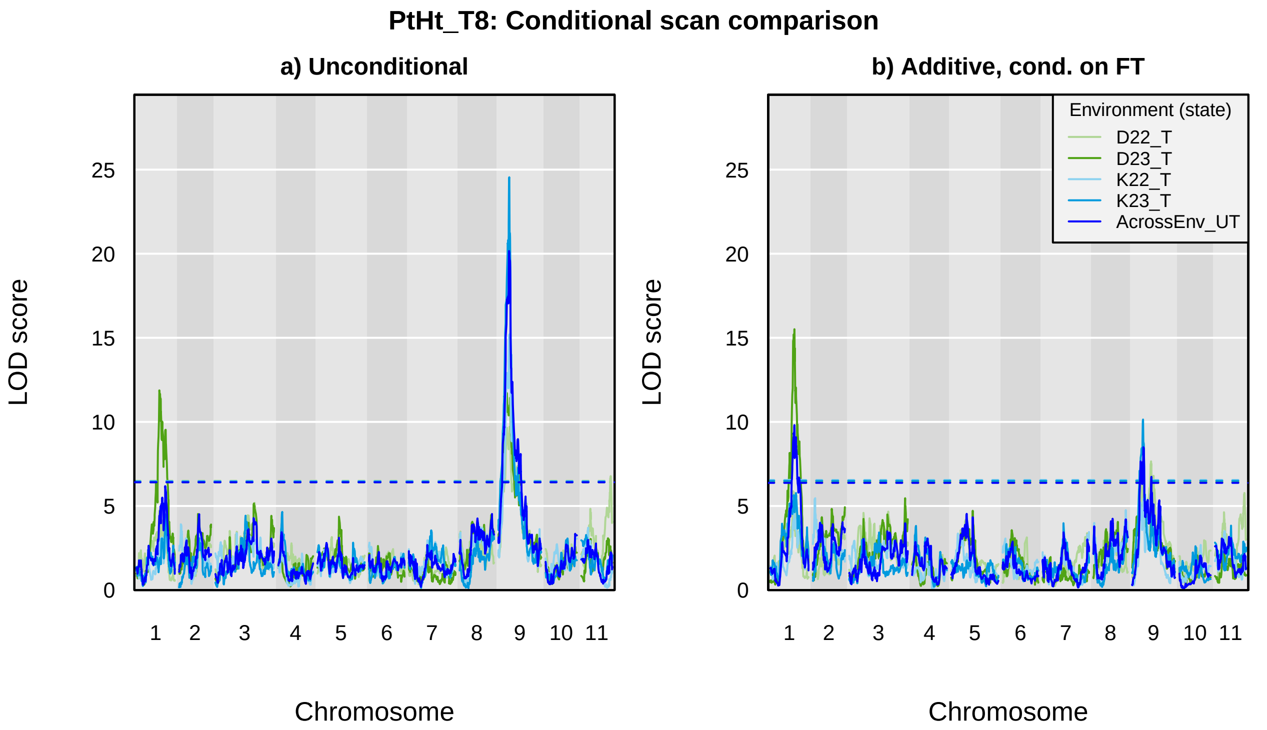
PtHt_T8: Conditional scan comparison
a) Unconditional
b) Additive, cond. on FT
LOD score
Chromosome
0
5
10
15
20
25
1
2
3
4
5
6
7
8
9
10
11
Environment (state)
D22_T
D23_T
K22_T
K23_T
AcrossEnv_UT

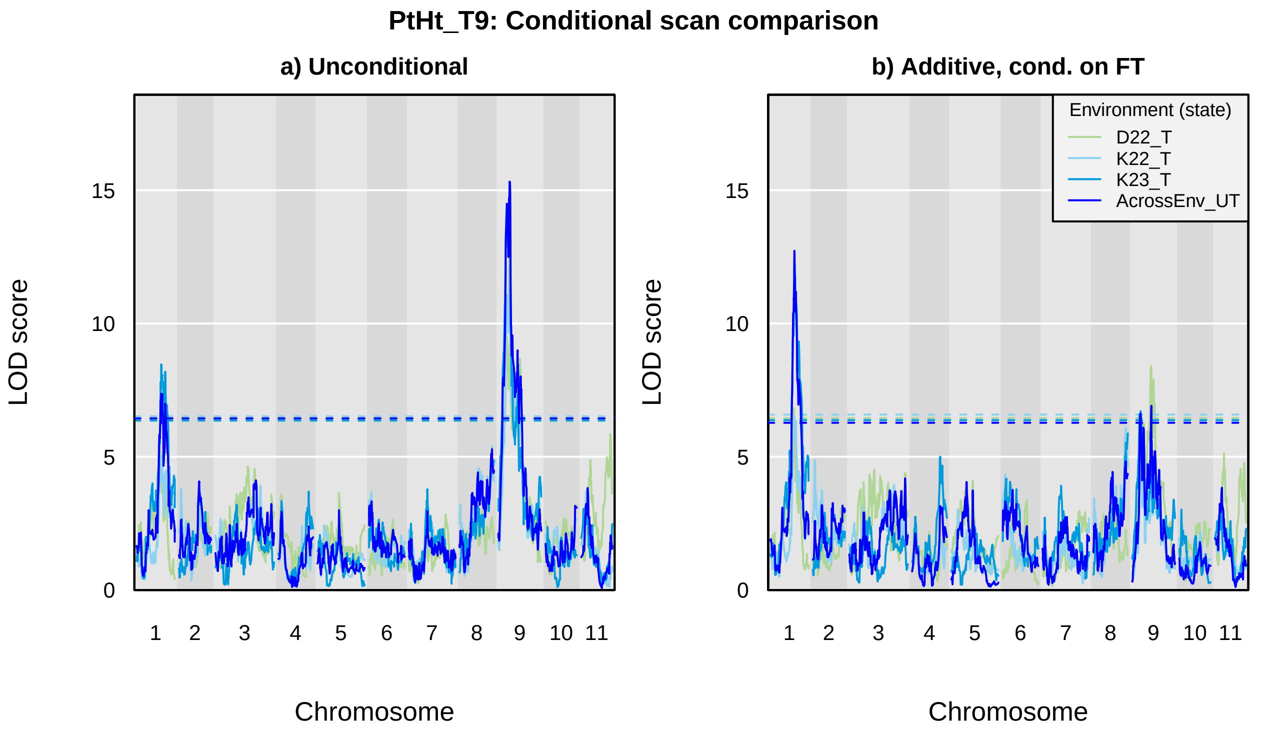

PtHt_T9: Conditional scan comparison
a) Unconditional
b) Additive, cond. on FT
LOD score
Chromosome
0
5
10
15
1
2
3
4
5
6
7
8
9
10
11
Environment (state)
D22_T
K22_T
K23_T
AcrossEnv_UT

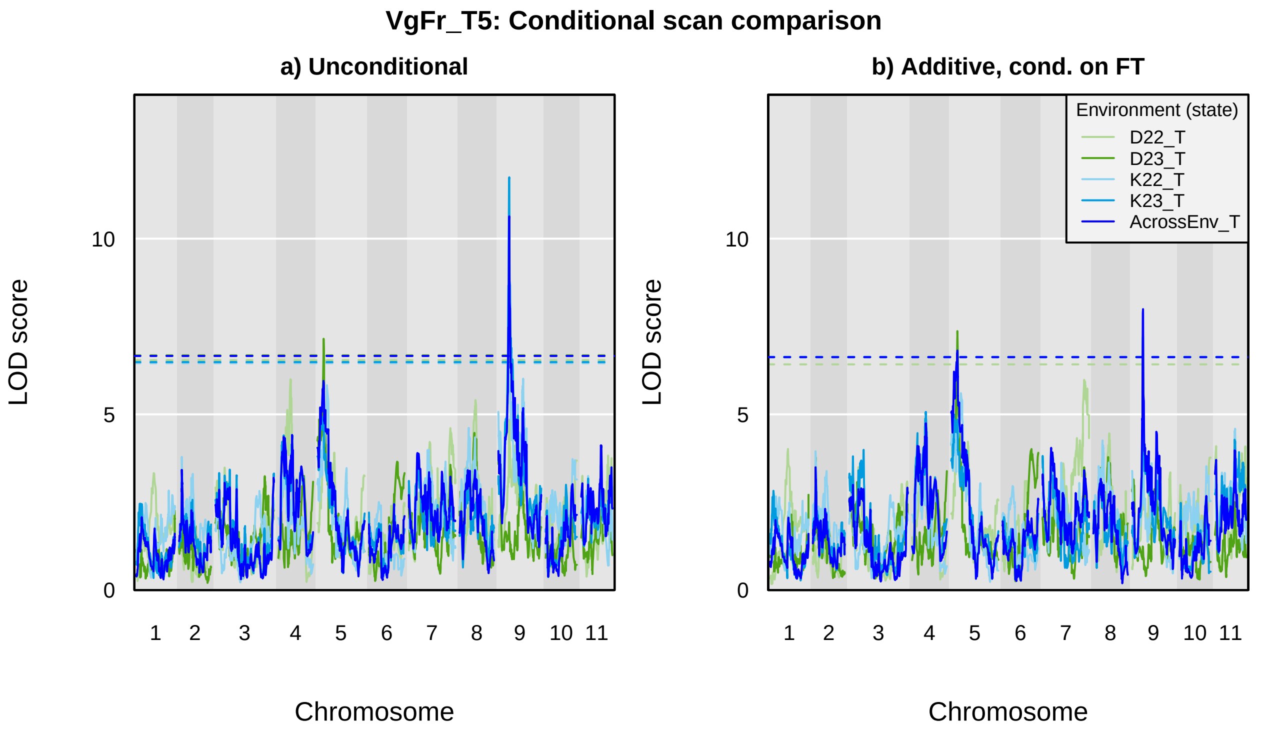

VgFr_T5: Conditional scan comparison
a) Unconditional
b) Additive, cond. on FT
LOD score
0
5
10
Chromosome
1
2
3
4
5
6
7
8
9
10
11
Environment (state)
D22_T
D23_T
K22_T
K23_T
AcrossEnv_T

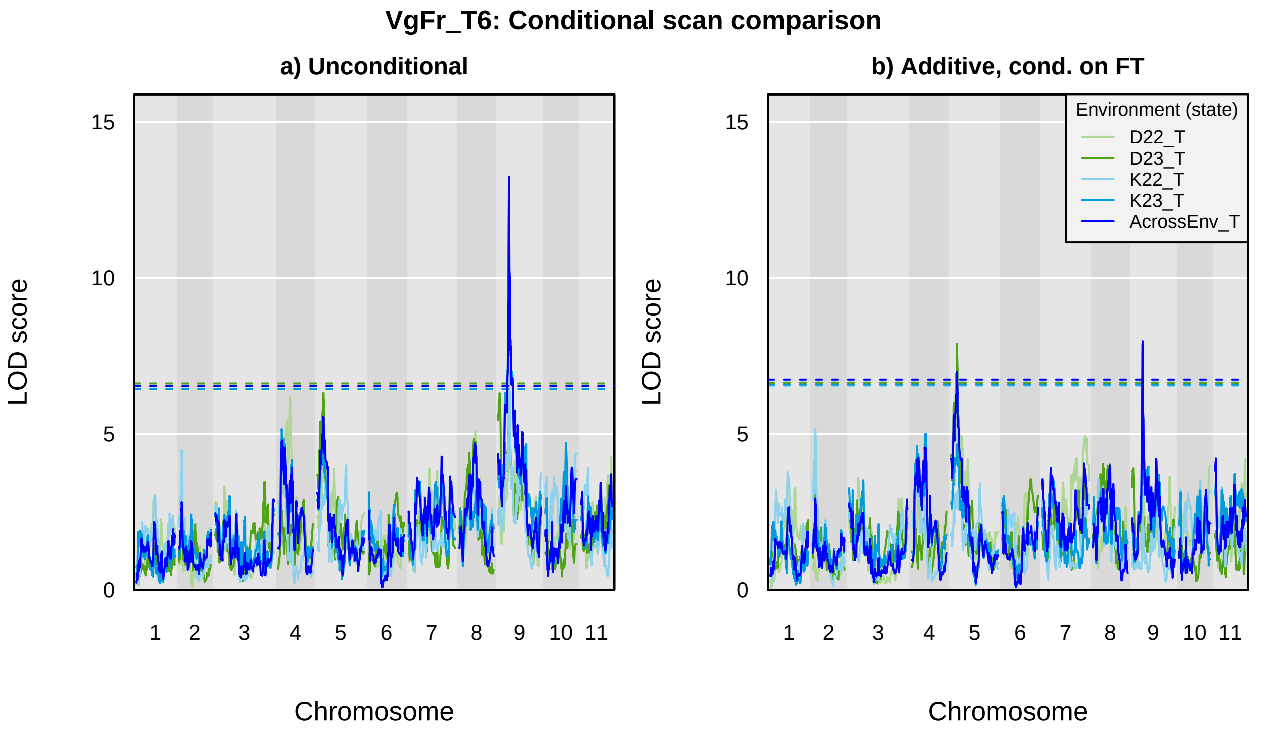
VgFr_T6: Conditional scan comparison
a) Unconditional
b) Additive, cond. on FT
LOD score
Chromosome
0
5
10
15
1
2
3
4
5
6
7
8
9
10
11
Environment (state)
D22_T
D23_T
K22_T
K23_T
AcrossEnv_T

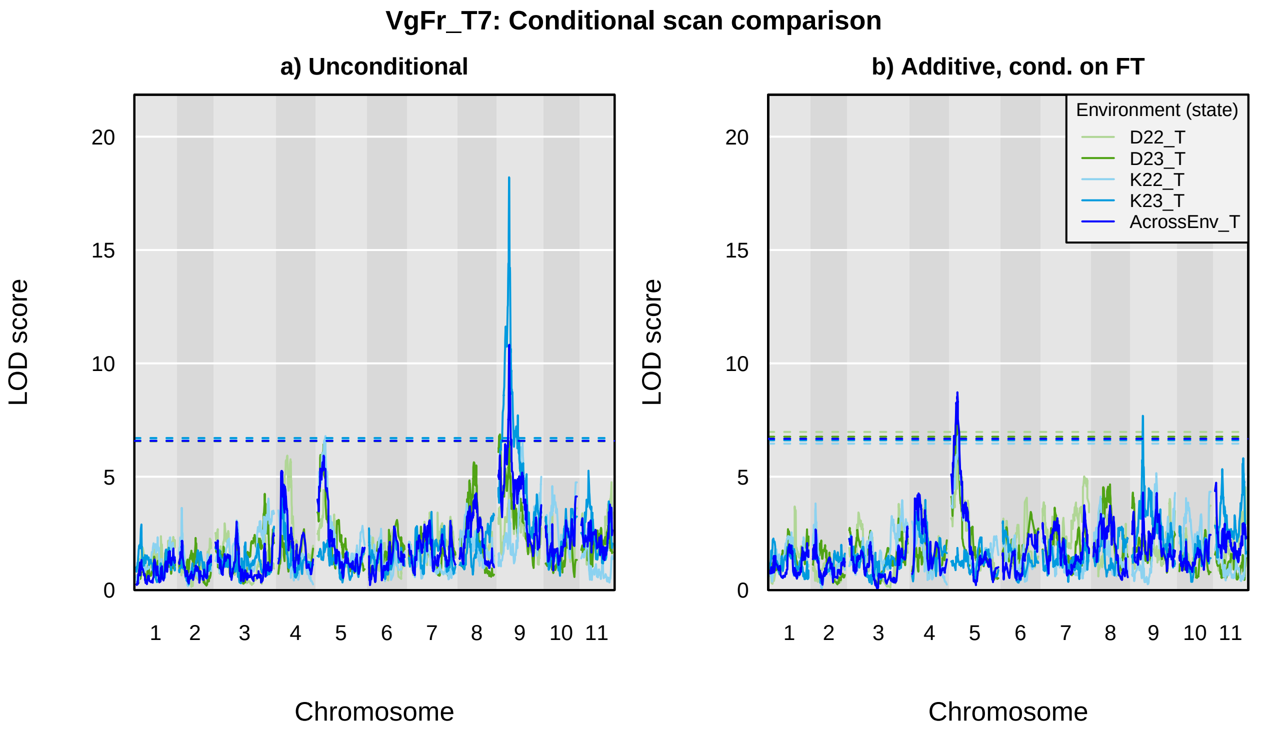
VgFr_T7: Conditional scan comparison
a) Unconditional
b) Additive, cond. on FT
LOD score
0
5
10
15
20
Chromosome
1
2
3
4
5
6
7
8
9
10
11
Environment (state)
D22_T
D23_T
K22_T
K23_T
AcrossEnv_T

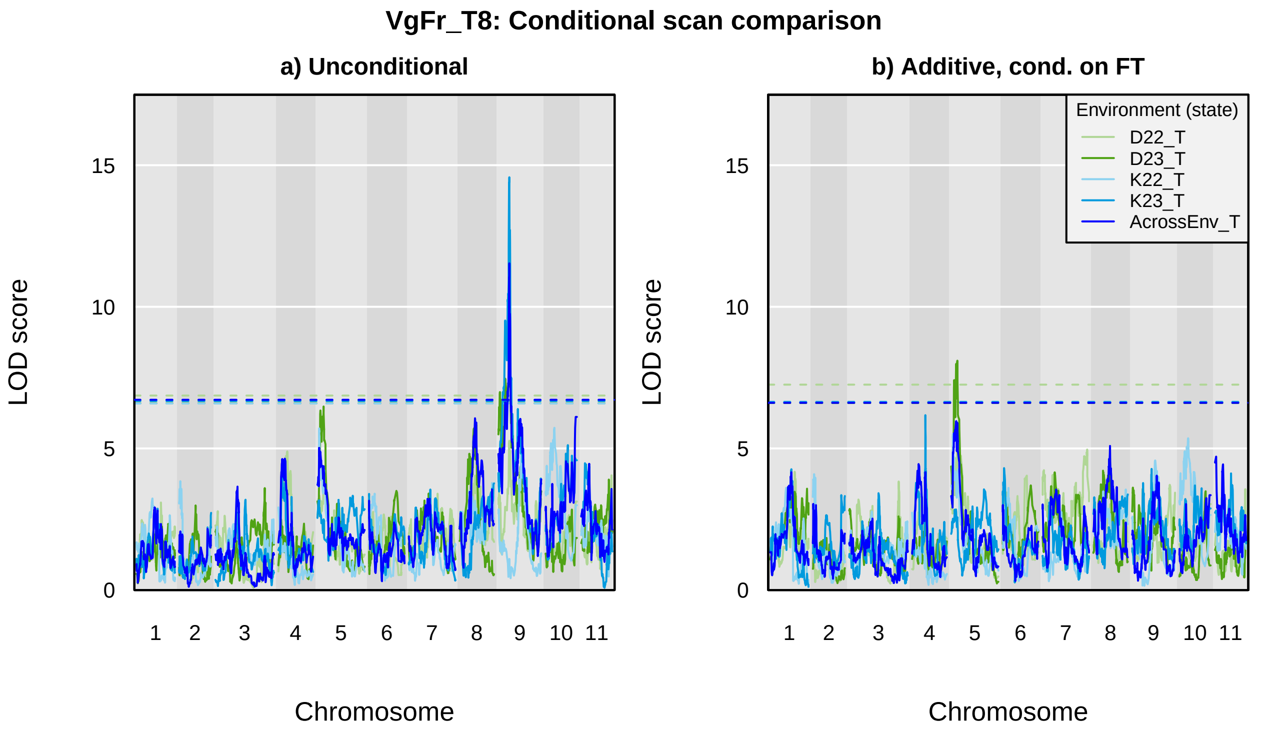
VgFr_T8: Conditional scan comparison
a) Unconditional
b) Additive, cond. on FT
LOD score
0
5
10
15
Chromosome
1
2
3
4
5
6
7
8
9
10
11
Environment (state)
D22_T
D23_T
K22_T
K23_T
AcrossEnv_T

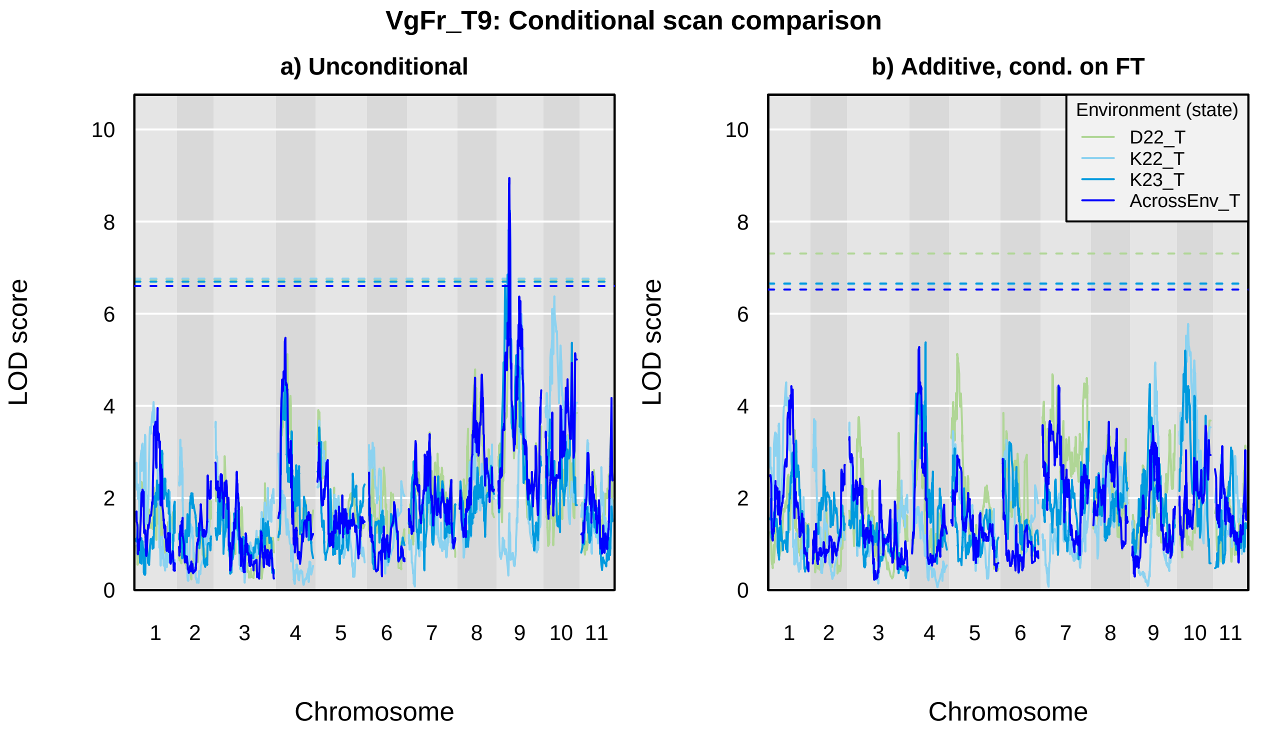
VgFr_T9: Conditional scan comparison
a) Unconditional
b) Additive, cond. on FT
LOD score
Chromosome
0
2
4
6
8
10
1
2
3
4
5
6
7
8
9
10
11
Environment (state)
D22_T
K22_T
K23_T
AcrossEnv_T
