## Supplementary material for "Genetic mapping and genomic prediction for agronomic, grain compositional, and sensing-enabled traits in a cowpea MAGIC population along an environmental gradient": Large supplemental tables and figures, uploaded separately: SuppFig_18_CowpeaMET_SMF_Diagnostics.pdf

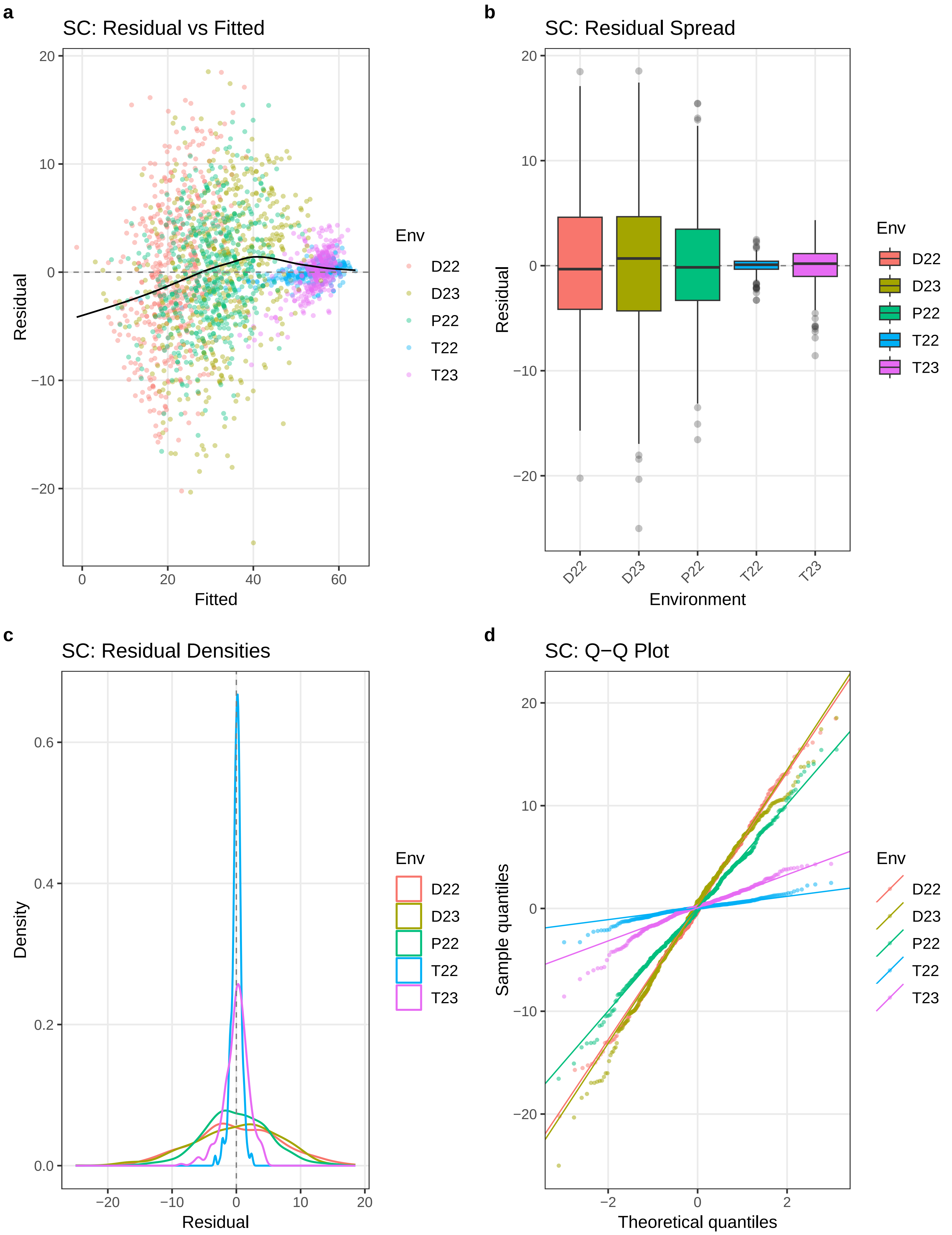

a
SC: Residual vs Fitted
Residual
Fitted
Env
D22
D23
P22
T22
T23
b
SC: Residual Spread
Residual
Environment
D22
D23
P22
T22
T23
c
SC: Residual Densities
Density
Residual
d
SC: Q−Q Plot
Sample quantiles
Theoretical quantiles

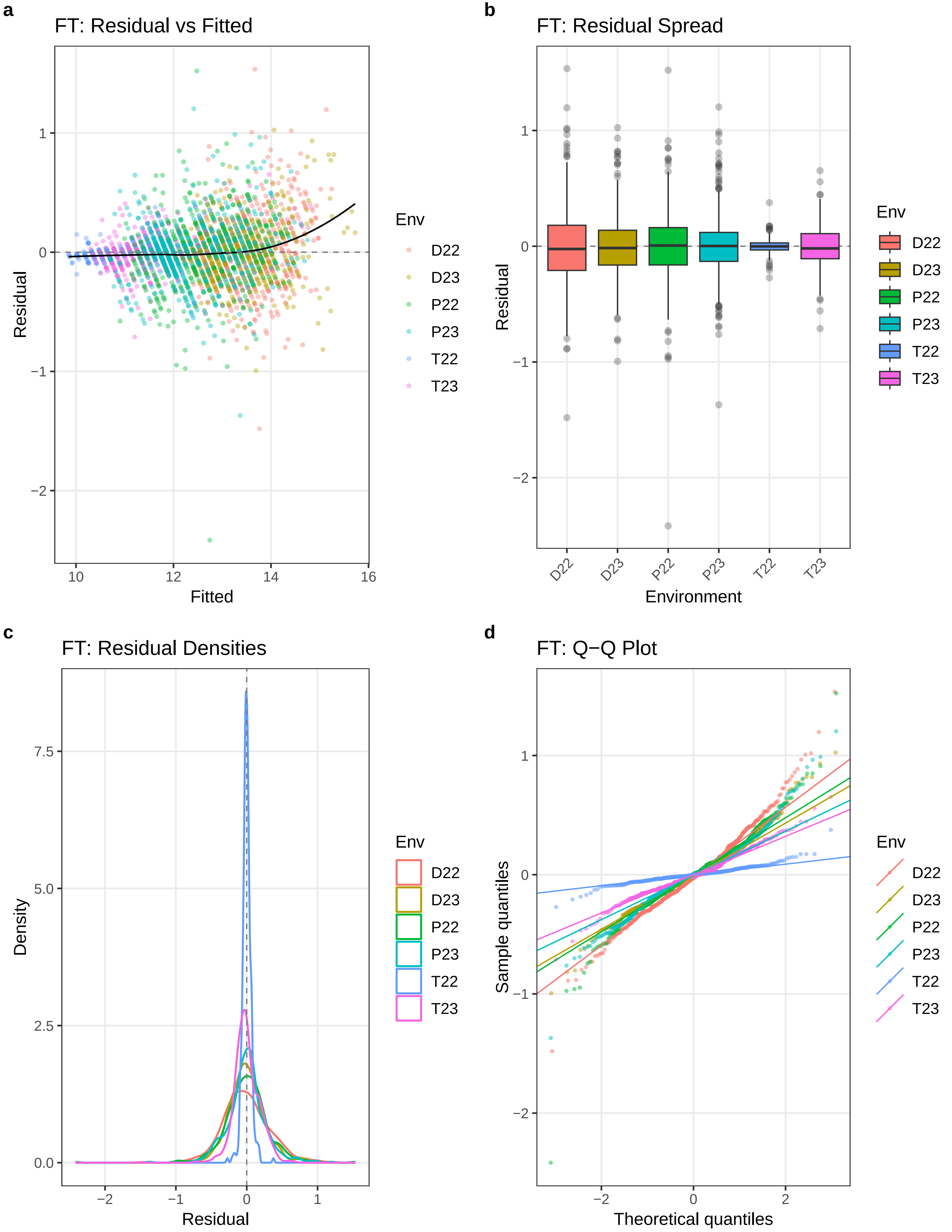
a
FT: Residual vs Fitted
Residual
Fitted
1
0
−1
−2
10
12
14
16
Env
D22
D23
P22
P23
T22
T23
b
FT: Residual Spread
Residual
Environment
D22
D23
P22
P23
T22
T23
c
FT: Residual Densities
Density
Residual
7.5
5.0
2.5
0.0
−2
−1
0
1
d
FT: Q−Q Plot
Sample quantiles
Theoretical quantiles
−2
0
2

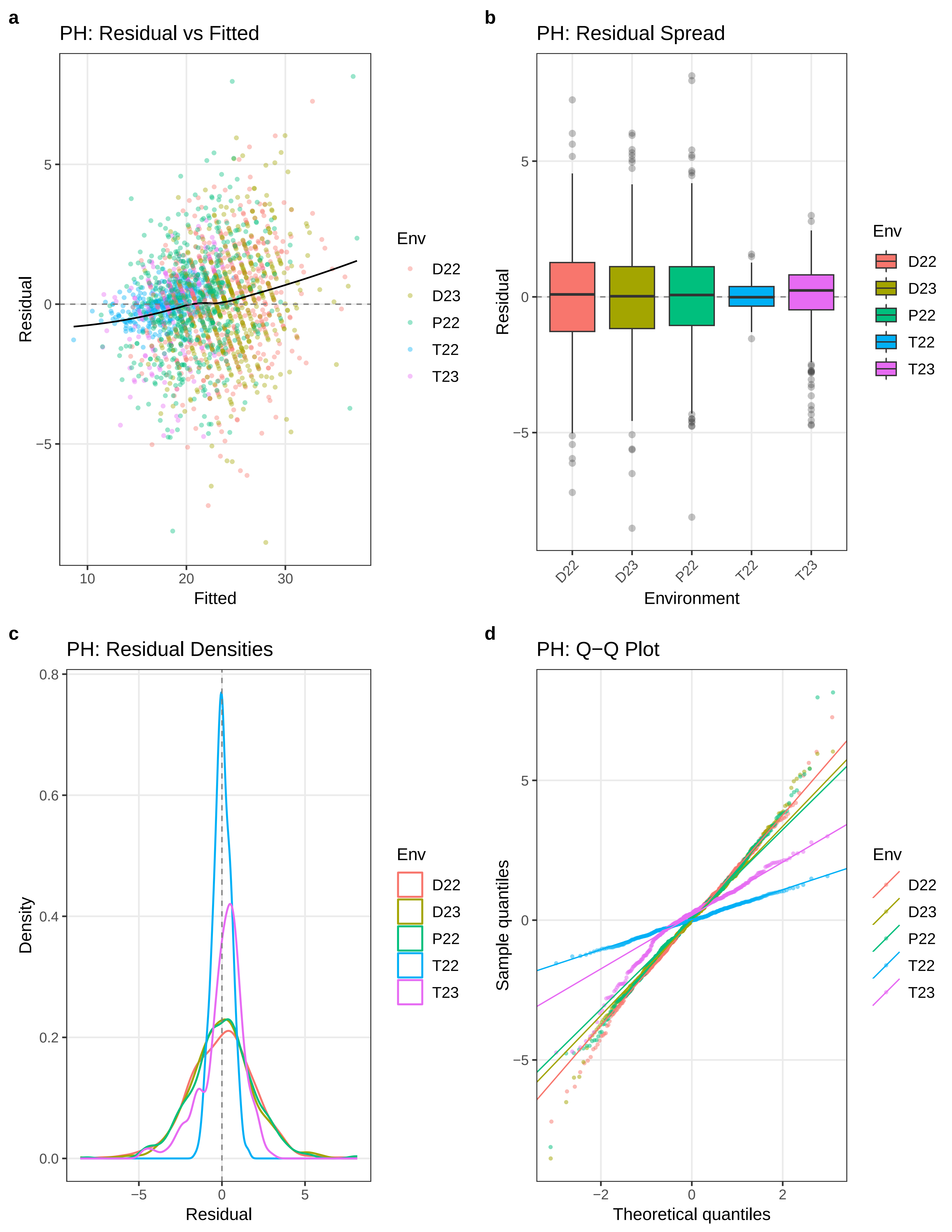

a
PH: Residual vs Fitted
Residual
Fitted
5
0
−5
10
20
30
Env
D22
D23
P22
T22
T23
b
PH: Residual Spread
Residual
Environment
5
0
−5
D22
D23
P22
T22
T23
Env
D22
D23
P22
T22
T23
c
PH: Residual Densities
Density
Residual
0.8
0.6
0.4
0.2
0.0
−5
0
5
Env
D22
D23
P22
T22
T23
d
PH: Q−Q Plot
Sample quantiles
Theoretical quantiles
5
0
−5
−2
0
2
Env
D22
D23
P22
T22
T23

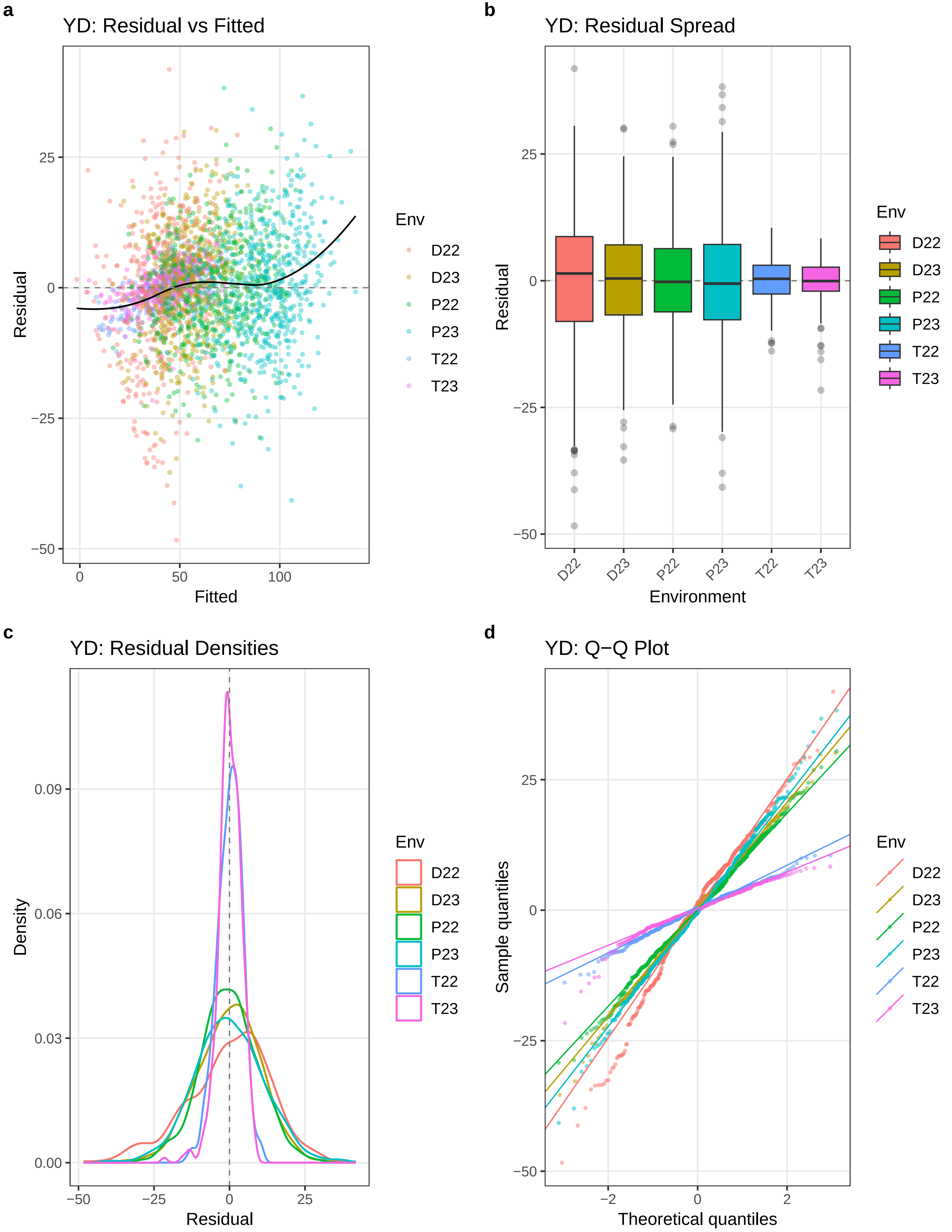
a
YD: Residual vs Fitted
Residual
Fitted
Env
D22
D23
P22
P23
T22
T23
b
YD: Residual Spread
Residual
Environment
c
YD: Residual Densities
Density
Residual
d
YD: Q−Q Plot
Sample quantiles
Theoretical quantiles

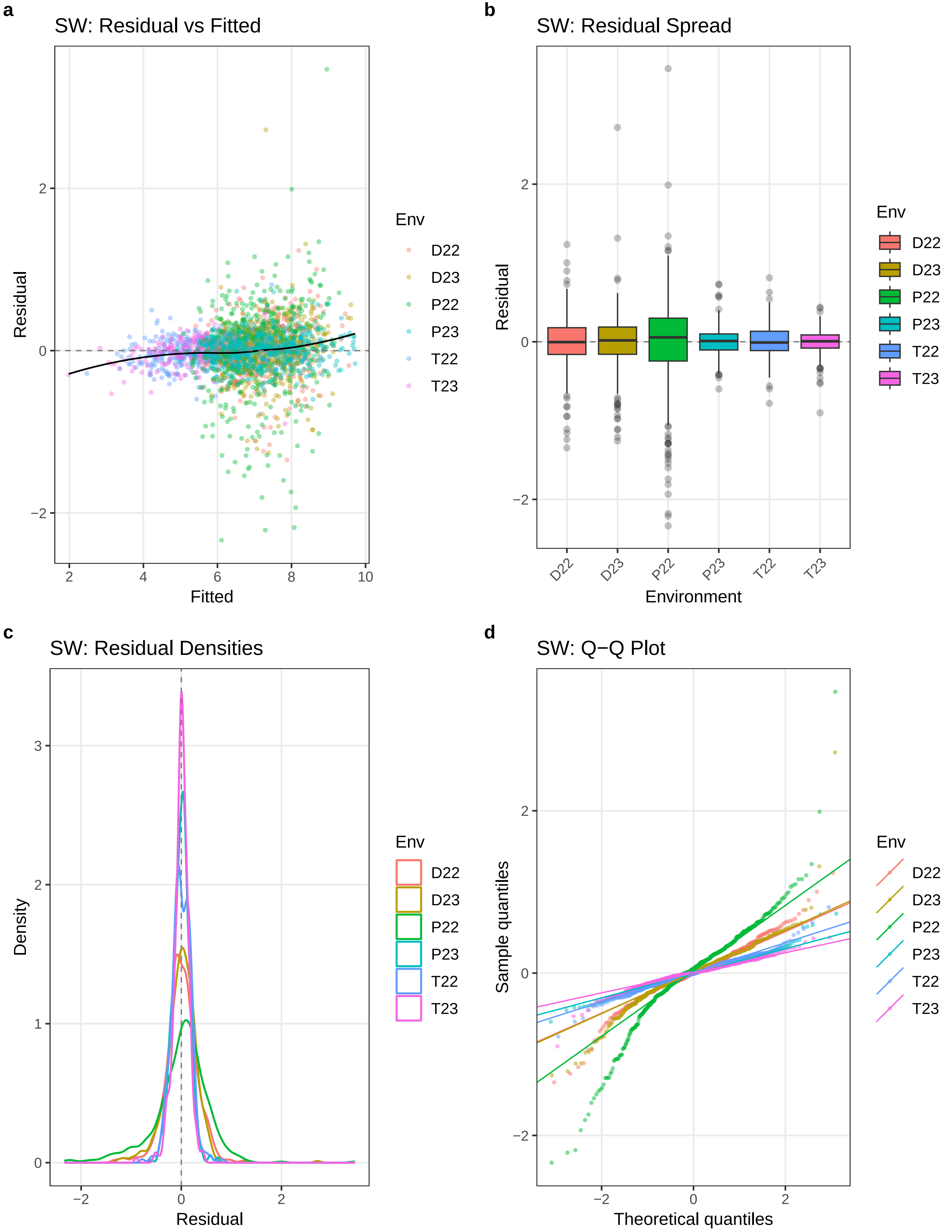
a
SW: Residual vs Fitted
Residual
Fitted
Env
D22
D23
P22
P23
T22
T23
b
SW: Residual Spread
Residual
Environment
c
SW: Residual Densities
Density
Residual
d
SW: Q−Q Plot
Sample quantiles
Theoretical quantiles

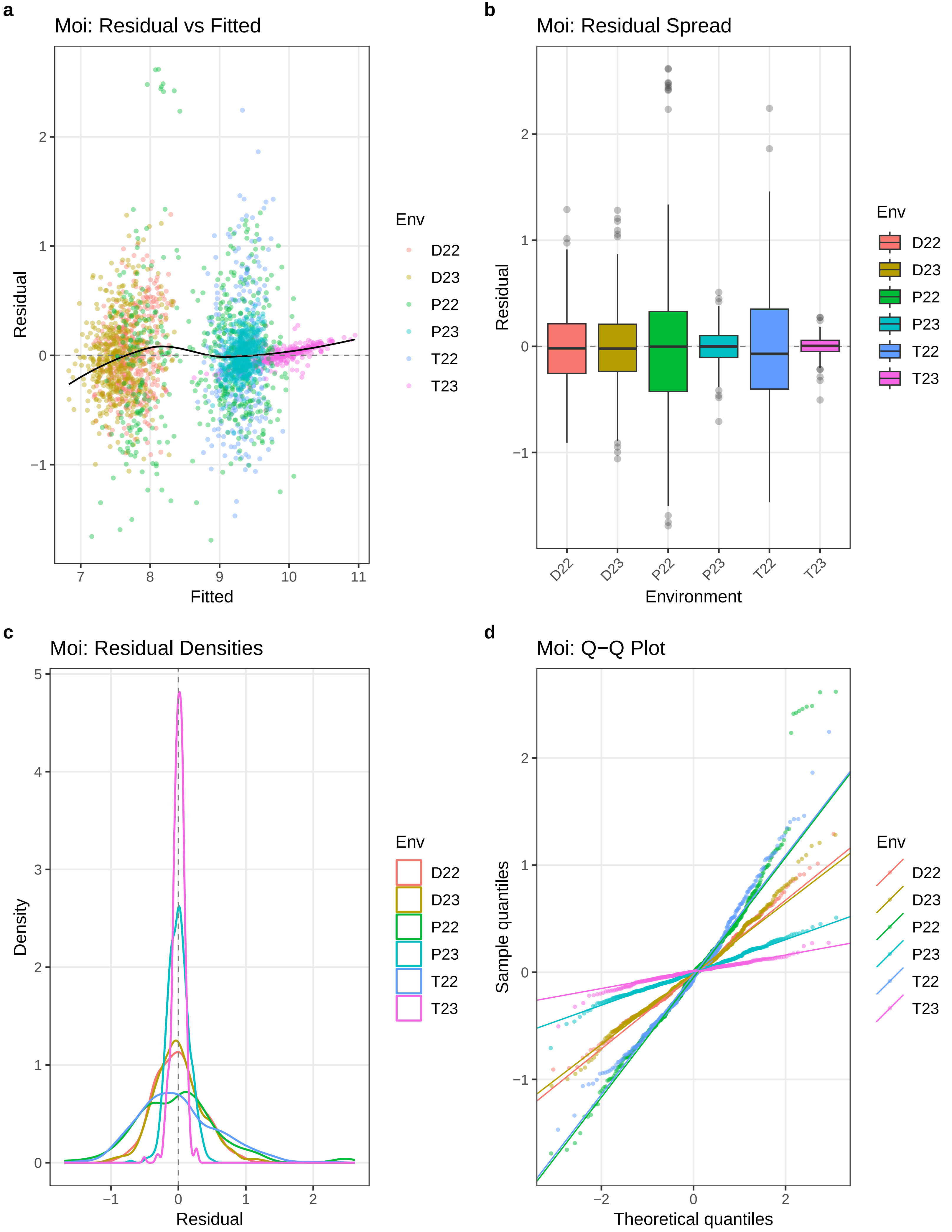

a
Moi: Residual vs Fitted
Residual
Fitted
Env
D22
D23
P22
P23
T22
T23
b
Moi: Residual Spread
Residual
Environment
D22
D23
P22
P23
T22
T23
Env
D22
D23
P22
P23
T22
T23
c
Moi: Residual Densities
Density
Residual
Env
D22
D23
P22
P23
T22
T23
d
Moi: Q−Q Plot
Sample quantiles
Theoretical quantiles
Env
D22
D23
P22
P23
T22
T23

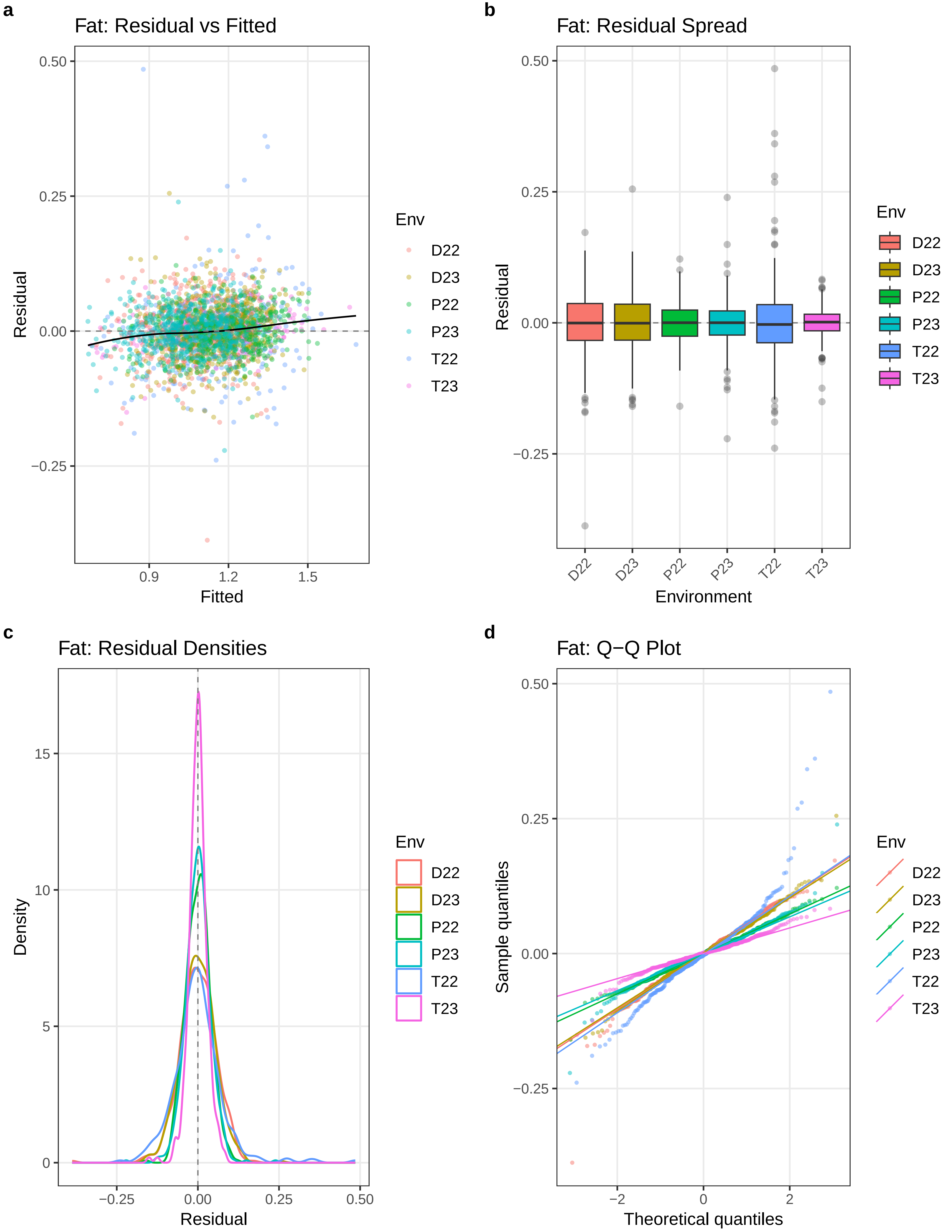
a
Fat: Residual vs Fitted
Residual
Fitted
0.50
0.25
0.00
−0.25
0.9
1.2
1.5
Env
D22
D23
P22
P23
T22
T23
b
Fat: Residual Spread
Residual
Environment
D22
D23
P22
P23
T22
T23
c
Fat: Residual Densities
Density
Residual
15
10
5
0
−0.25
0.00
0.25
0.50
d
Fat: Q−Q Plot
Sample quantiles
Theoretical quantiles
−2
0
2

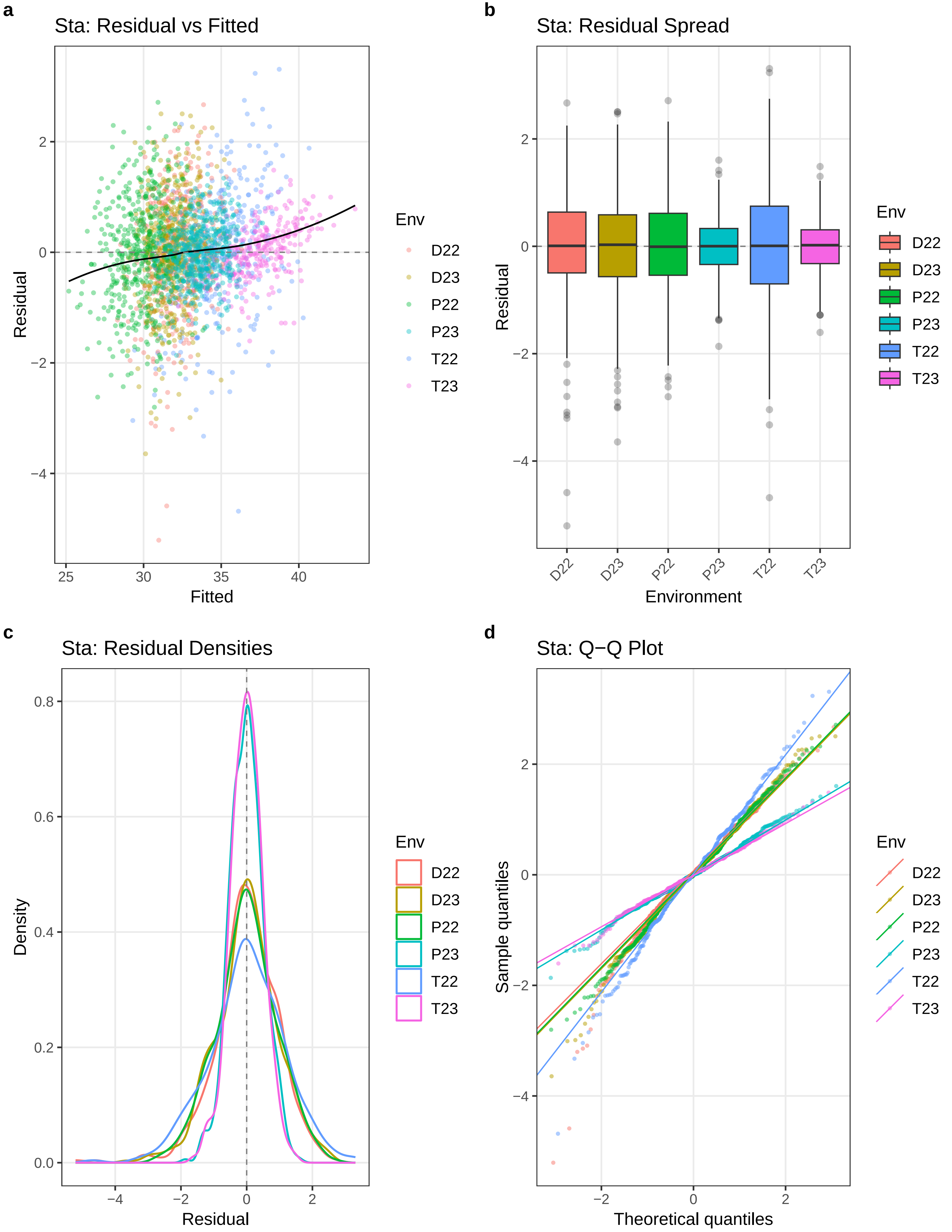
a
Sta: Residual vs Fitted
Residual
Fitted
Env
D22
D23
P22
P23
T22
T23
b
Sta: Residual Spread
Residual
Environment
c
Sta: Residual Densities
Density
Residual
d
Sta: Q−Q Plot
Sample quantiles
Theoretical quantiles

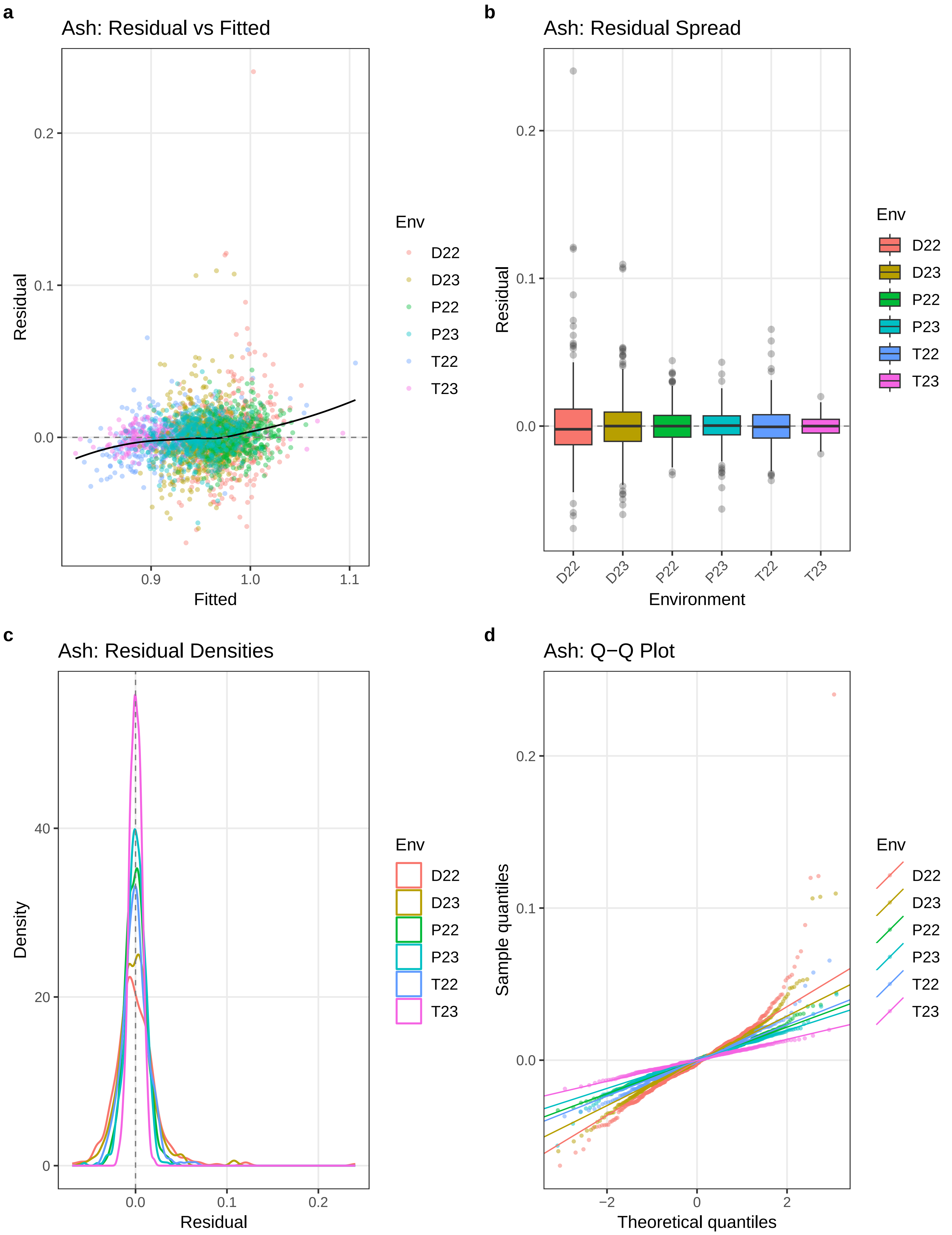
a
Ash: Residual vs Fitted
Residual
Fitted
0.2
0.1
0.0
0.9
1.0
1.1
Env
D22
D23
P22
P23
T22
T23
b
Ash: Residual Spread
Residual
Environment
D22
D23
P22
P23
T22
T23
c
Ash: Residual Densities
Density
Residual
40
20
0
0.0
0.1
0.2
d
Ash: Q−Q Plot
Sample quantiles
Theoretical quantiles
−2
0
2

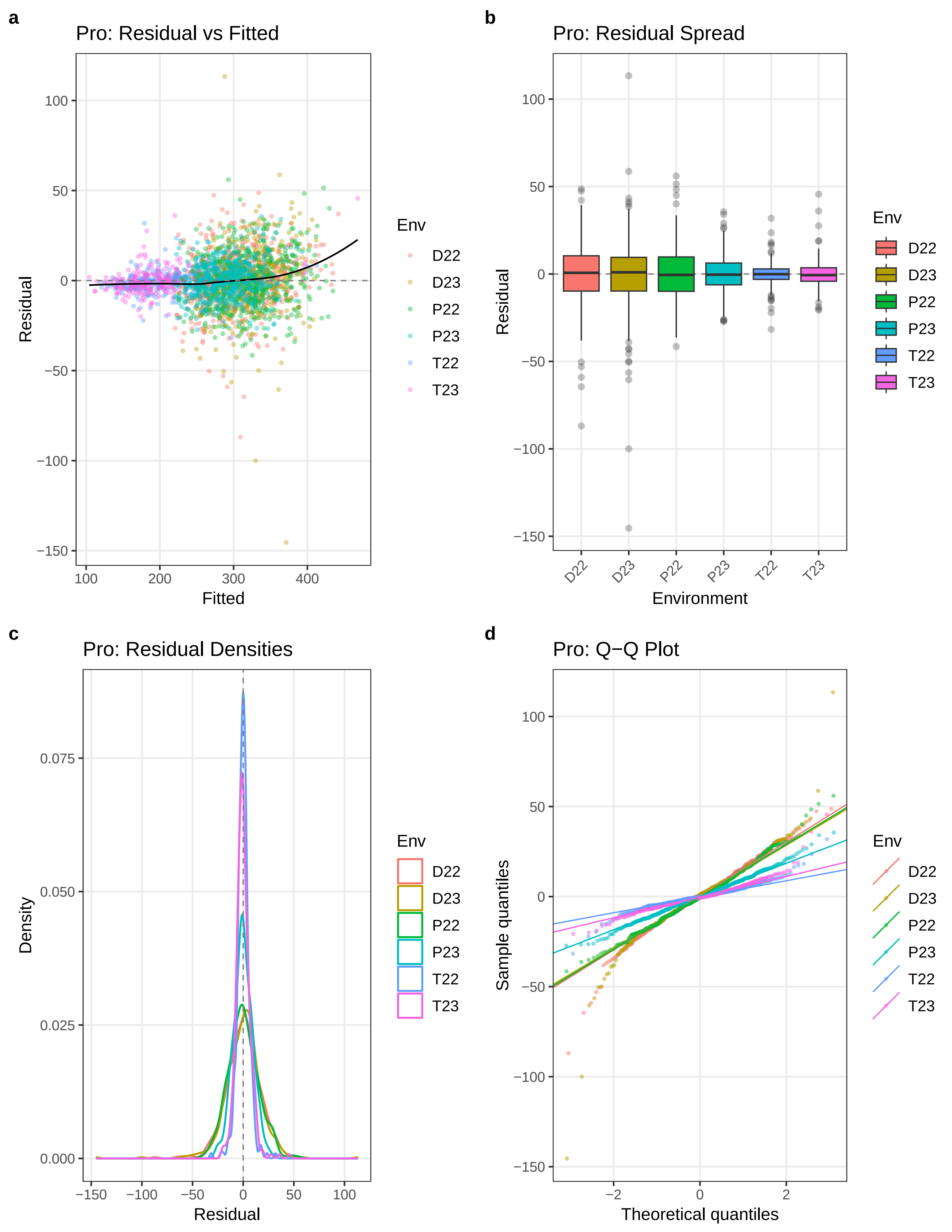

a
Pro: Residual vs Fitted
Residual
Fitted
b
Pro: Residual Spread
Residual
Environment
c
Pro: Residual Densities
Density
Residual
d
Pro: Q−Q Plot
Sample quantiles
Theoretical quantiles
Env
D22
D23
P22
P23
T22
T23

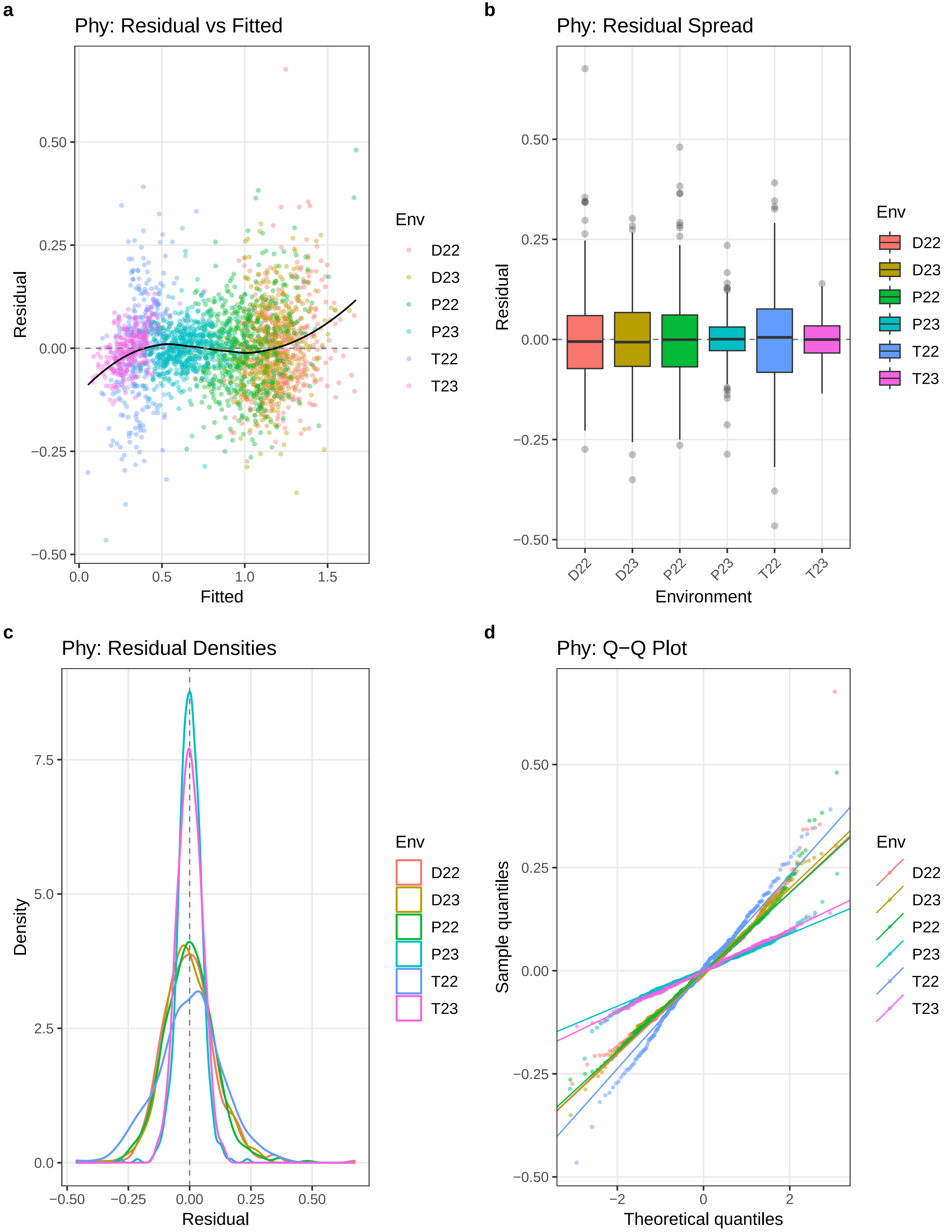

a
Phy: Residual vs Fitted
Residual
Fitted
Env
D22
D23
P22
P23
T22
T23
b
Phy: Residual Spread
Residual
Environment
c
Phy: Residual Densities
Density
Residual
d
Phy: Q−Q Plot
Sample quantiles
Theoretical quantiles

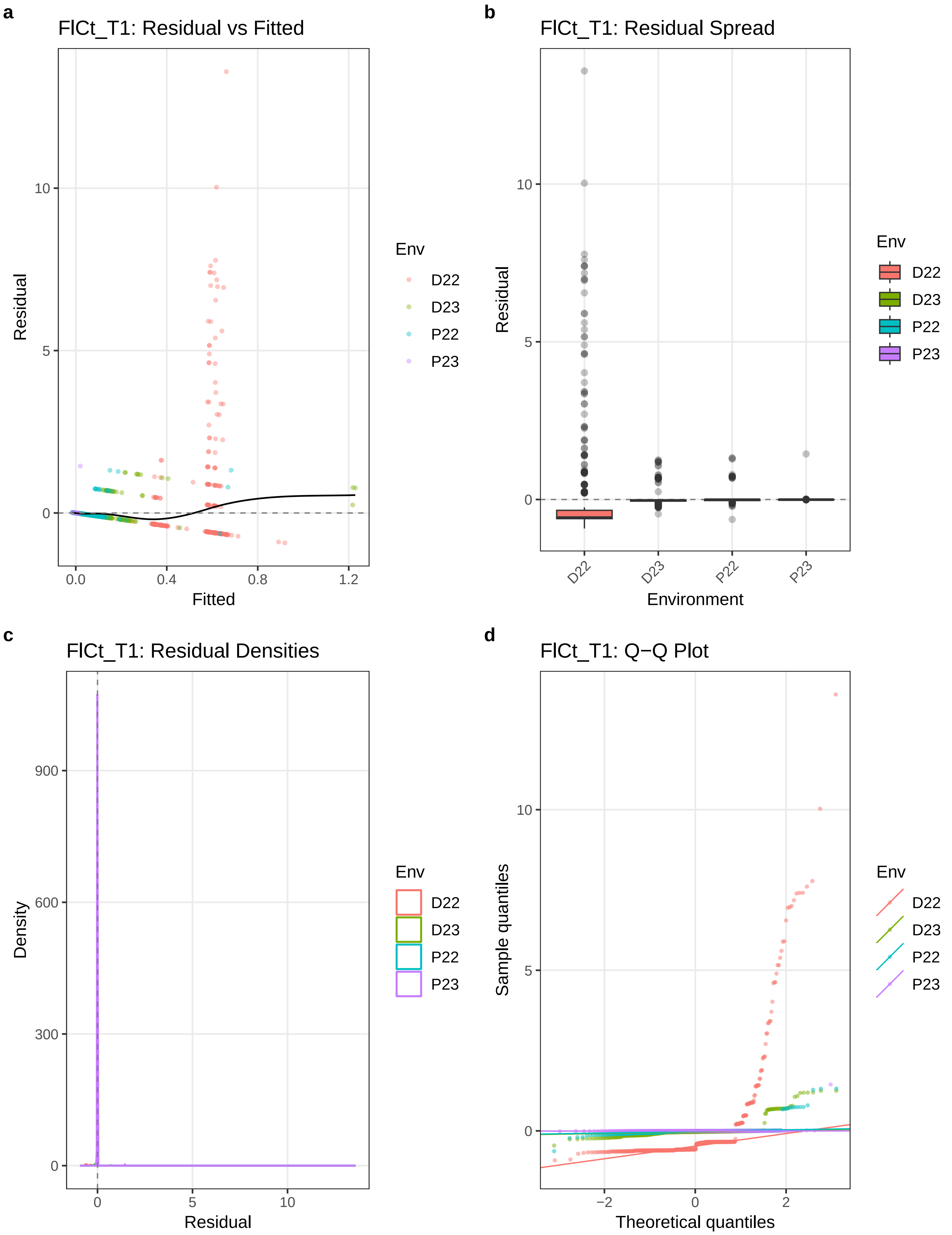
a
FlCt_T1: Residual vs Fitted
Residual
Fitted
0
5
10
0.0
0.4
0.8
1.2
Env
D22
D23
P22
P23
b
FlCt_T1: Residual Spread
Residual
Environment
0
5
10
D22
D23
P22
P23
Env
D22
D23
P22
P23
c
FlCt_T1: Residual Densities
Density
Residual
0
300
600
900
0
5
10
Env
D22
D23
P22
P23
d
FlCt_T1: Q−Q Plot
Sample quantiles
Theoretical quantiles
0
5
10
−2
0
2
Env
D22
D23
P22
P23

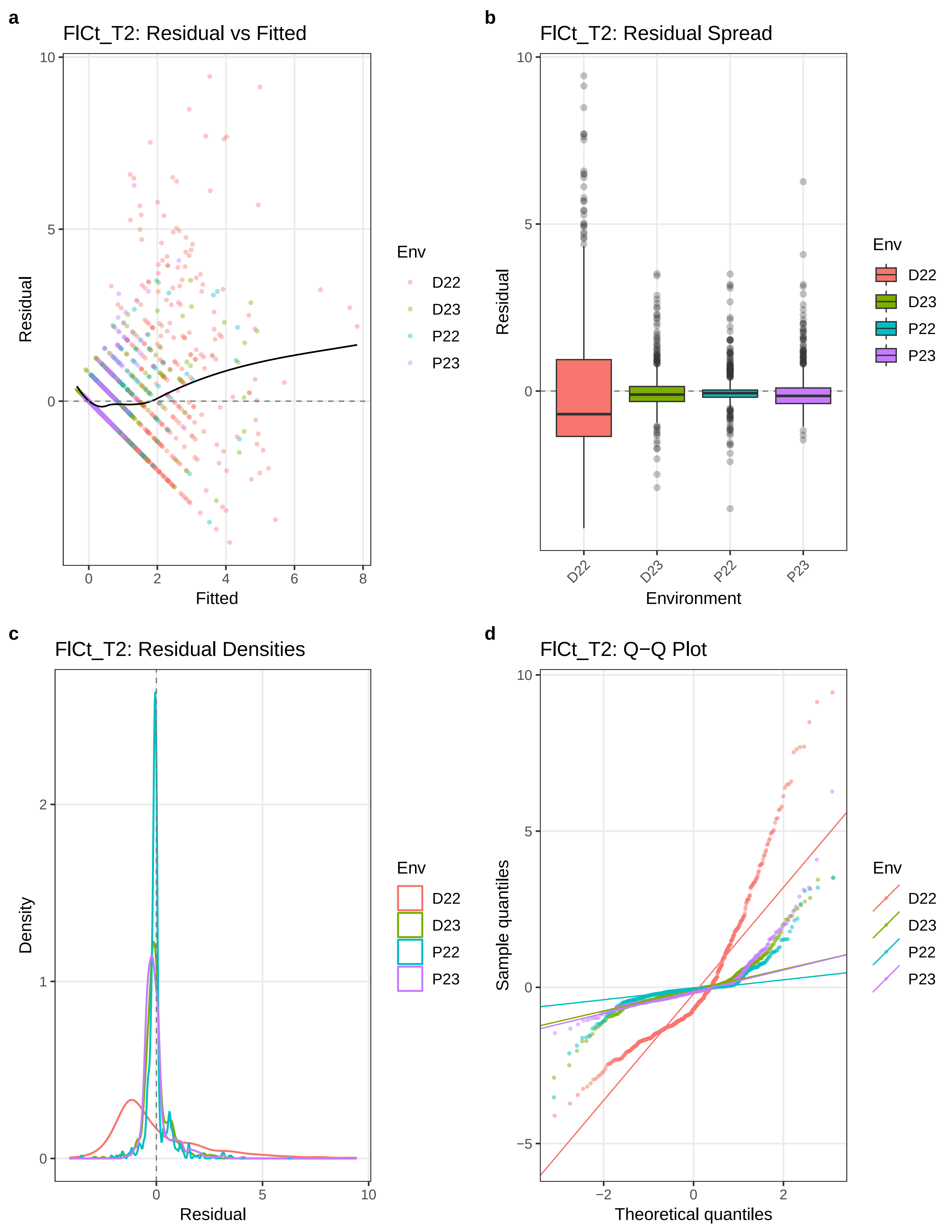
a
FlCt_T2: Residual vs Fitted
Residual
Fitted
10
5
0
0
2
4
6
8
Env
D22
D23
P22
P23
b
FlCt_T2: Residual Spread
Residual
Environment
D22
D23
P22
P23
c
FlCt_T2: Residual Densities
Density
Residual
2
1
0
0
5
10
d
FlCt_T2: Q−Q Plot
Sample quantiles
Theoretical quantiles
−5
−2
0
2

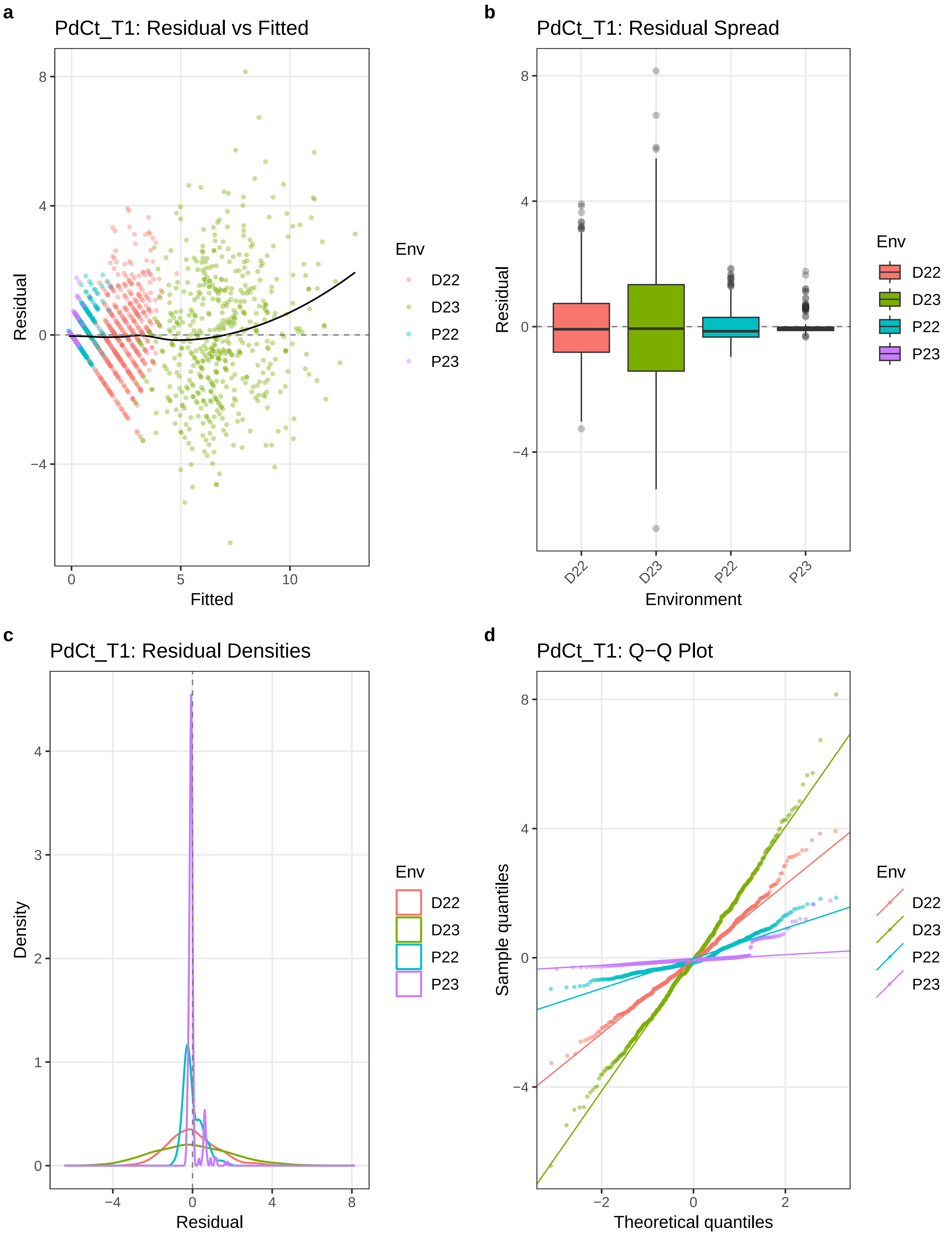
a
PdCt_T1: Residual vs Fitted
Residual
Fitted
Env
D22
D23
P22
P23
b
PdCt_T1: Residual Spread
Residual
Environment
c
PdCt_T1: Residual Densities
Density
Residual
d
PdCt_T1: Q−Q Plot
Sample quantiles
Theoretical quantiles

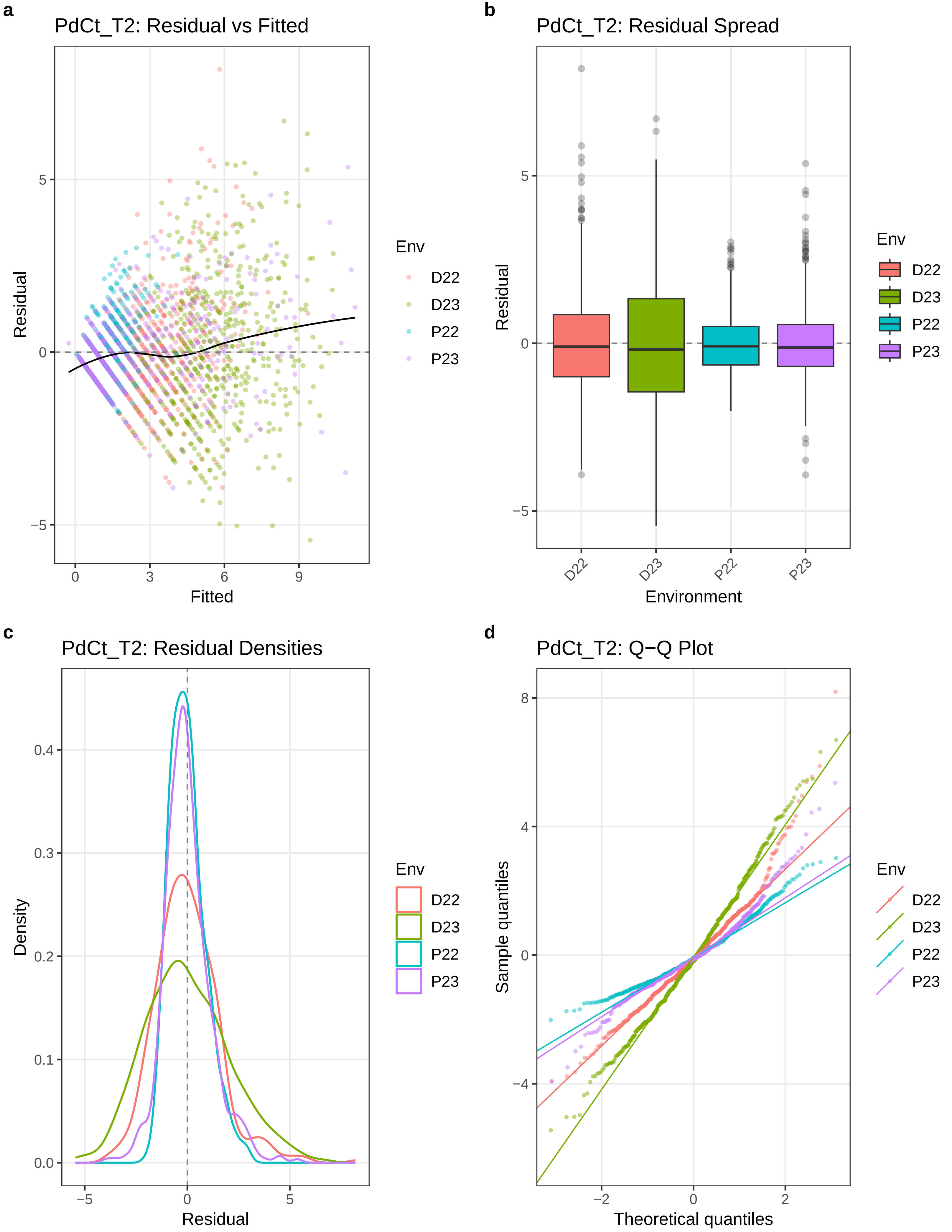
a
PdCt_T2: Residual vs Fitted
Residual
Fitted
Env
D22
D23
P22
P23
b
PdCt_T2: Residual Spread
Residual
Environment
c
PdCt_T2: Residual Densities
Density
Residual
d
PdCt_T2: Q−Q Plot
Sample quantiles
Theoretical quantiles

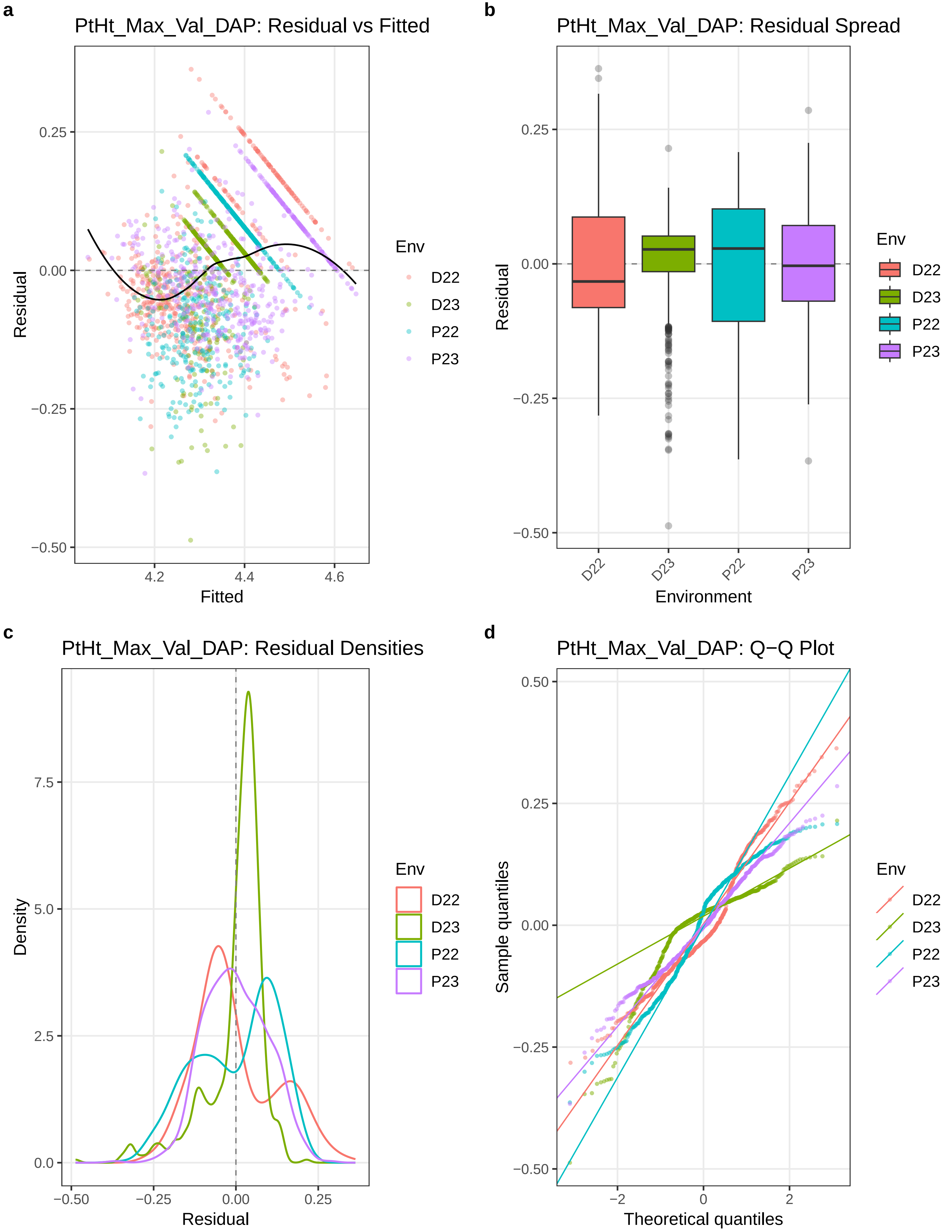

a
PtHt_Max_Val_DAP: Residual vs Fitted
Residual
Fitted
0.25
0.00
−0.25
−0.50
4.2
4.4
4.6
Env
D22
D23
P22
P23
b
PtHt_Max_Val_DAP: Residual Spread
Residual
Environment
D22
D23
P22
P23
c
PtHt_Max_Val_DAP: Residual Densities
Density
Residual
7.5
5.0
2.5
0.0
−0.50
−0.25
0.00
0.25
d
PtHt_Max_Val_DAP: Q−Q Plot
Sample quantiles
Theoretical quantiles
0.50
−2
0
2

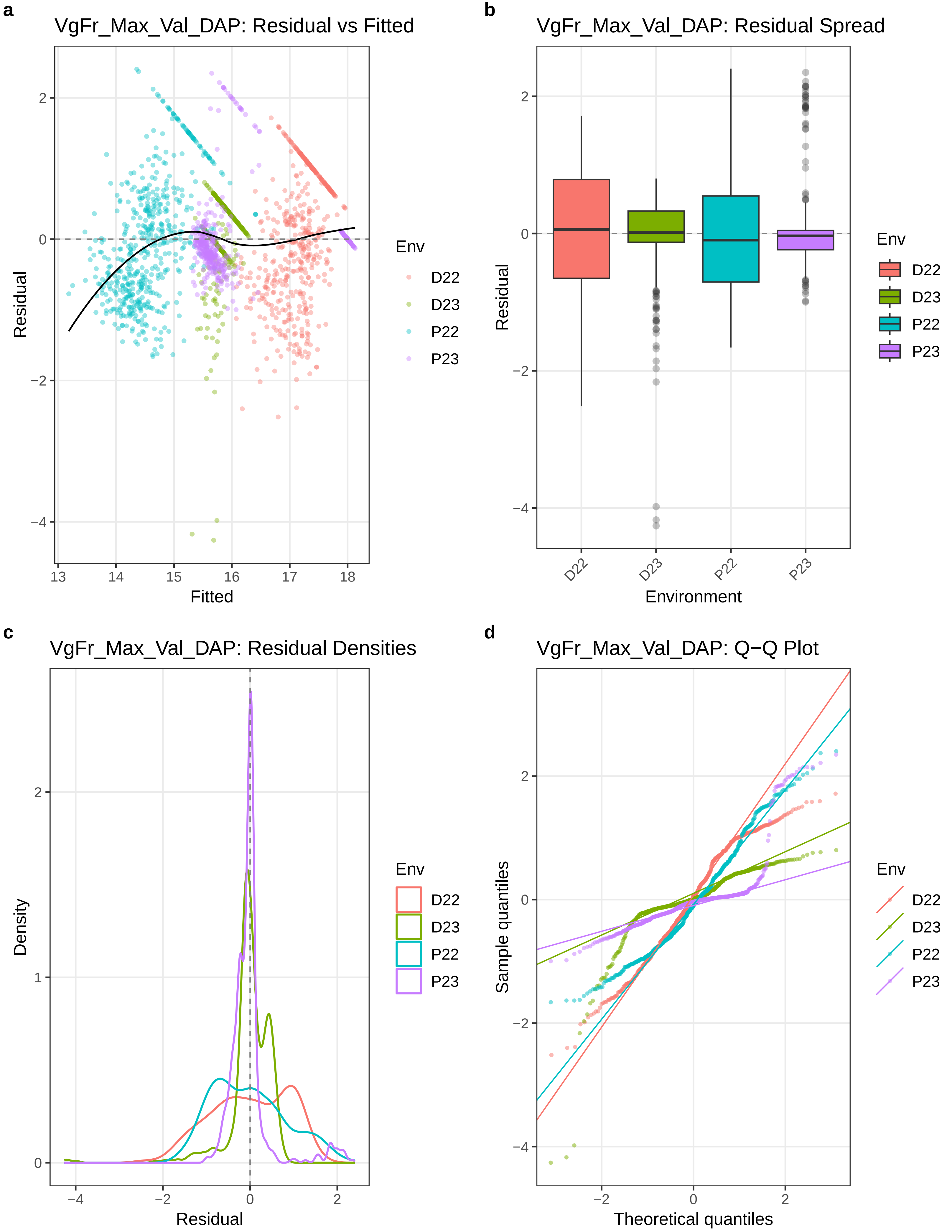
a
VgFr_Max_Val_DAP: Residual vs Fitted
Residual
Fitted
2
0
−2
−4
13
14
15
16
17
18
Env
D22
D23
P22
P23
b
VgFr_Max_Val_DAP: Residual Spread
Residual
Environment
D22
D23
P22
P23
Env
D22
D23
P22
P23
c
VgFr_Max_Val_DAP: Residual Densities
Density
Residual
2
1
0
−4
−2
0
2
Env
D22
D23
P22
P23
d
VgFr_Max_Val_DAP: Q−Q Plot
Sample quantiles
Theoretical quantiles
2
0
−2
−4
Env
D22
D23
P22
P23

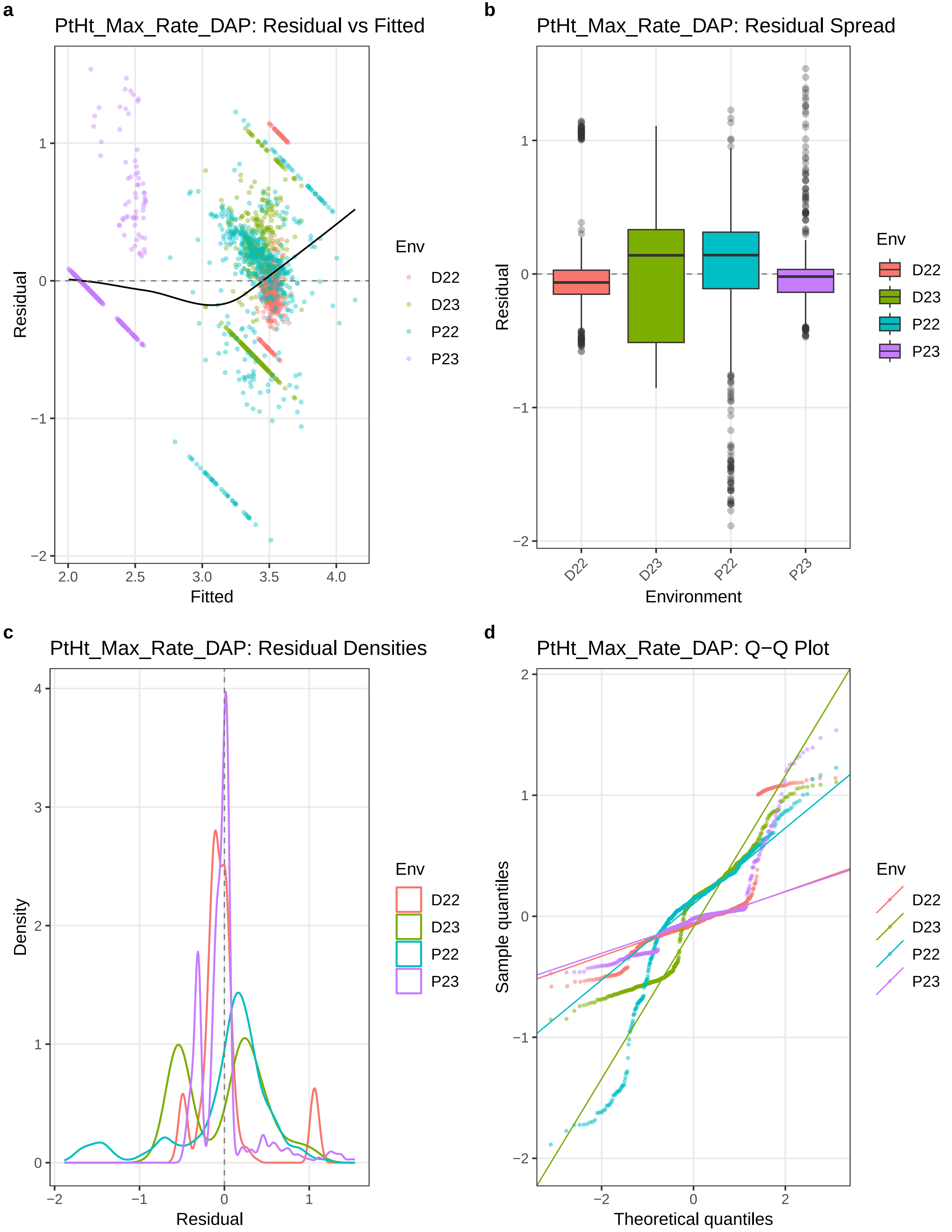
a
PtHt_Max_Rate_DAP: Residual vs Fitted
Residual
Fitted
Env
D22
D23
P22
P23
b
PtHt_Max_Rate_DAP: Residual Spread
Residual
Environment
D22
D23
P22
P23
c
PtHt_Max_Rate_DAP: Residual Densities
Density
Residual
d
PtHt_Max_Rate_DAP: Q−Q Plot
Sample quantiles
Theoretical quantiles

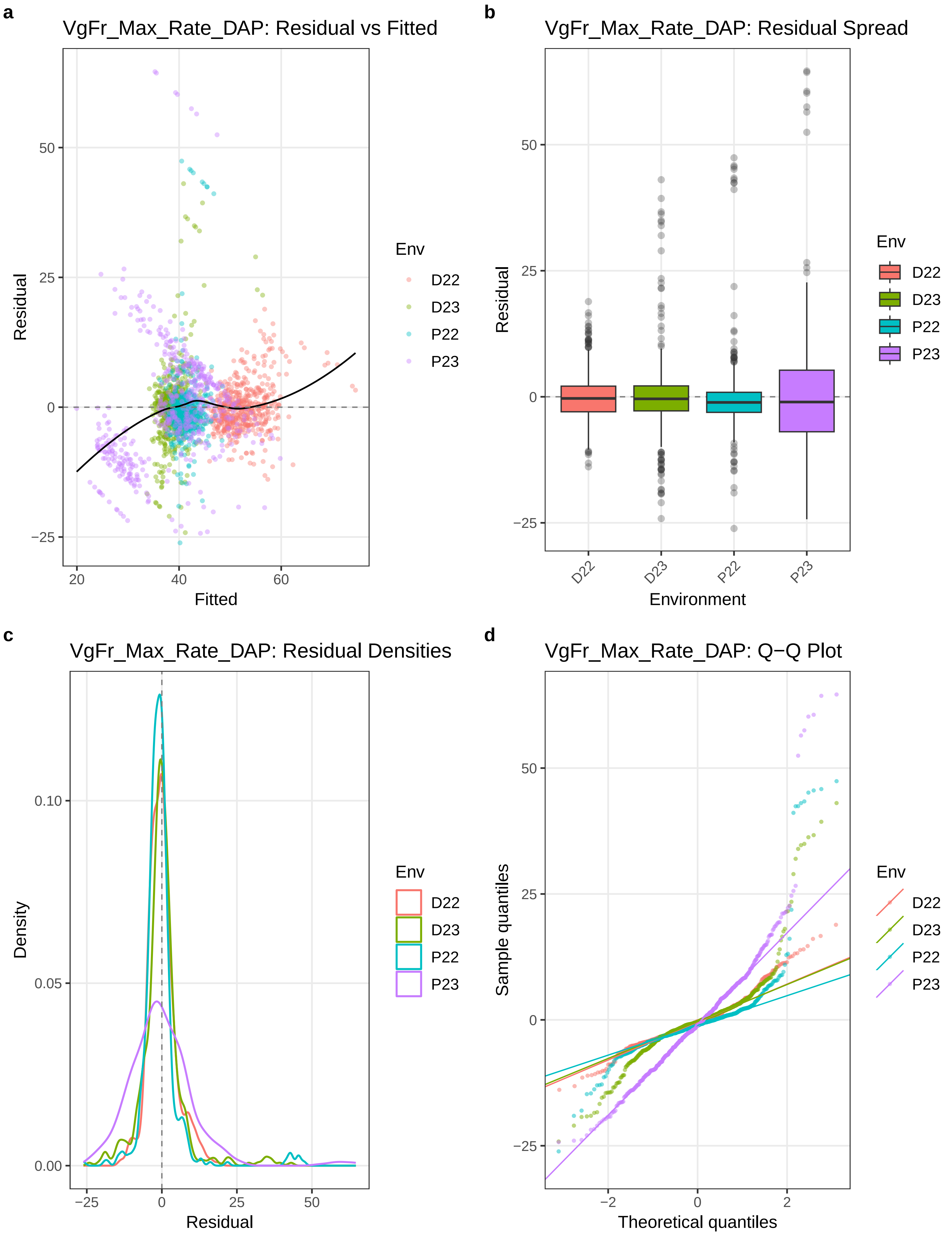

a
VgFr_Max_Rate_DAP: Residual vs Fitted
Residual
Fitted
Env
D22
D23
P22
P23
b
VgFr_Max_Rate_DAP: Residual Spread
Residual
Environment
c
VgFr_Max_Rate_DAP: Residual Densities
Density
Residual
d
VgFr_Max_Rate_DAP: Q−Q Plot
Sample quantiles
Theoretical quantiles

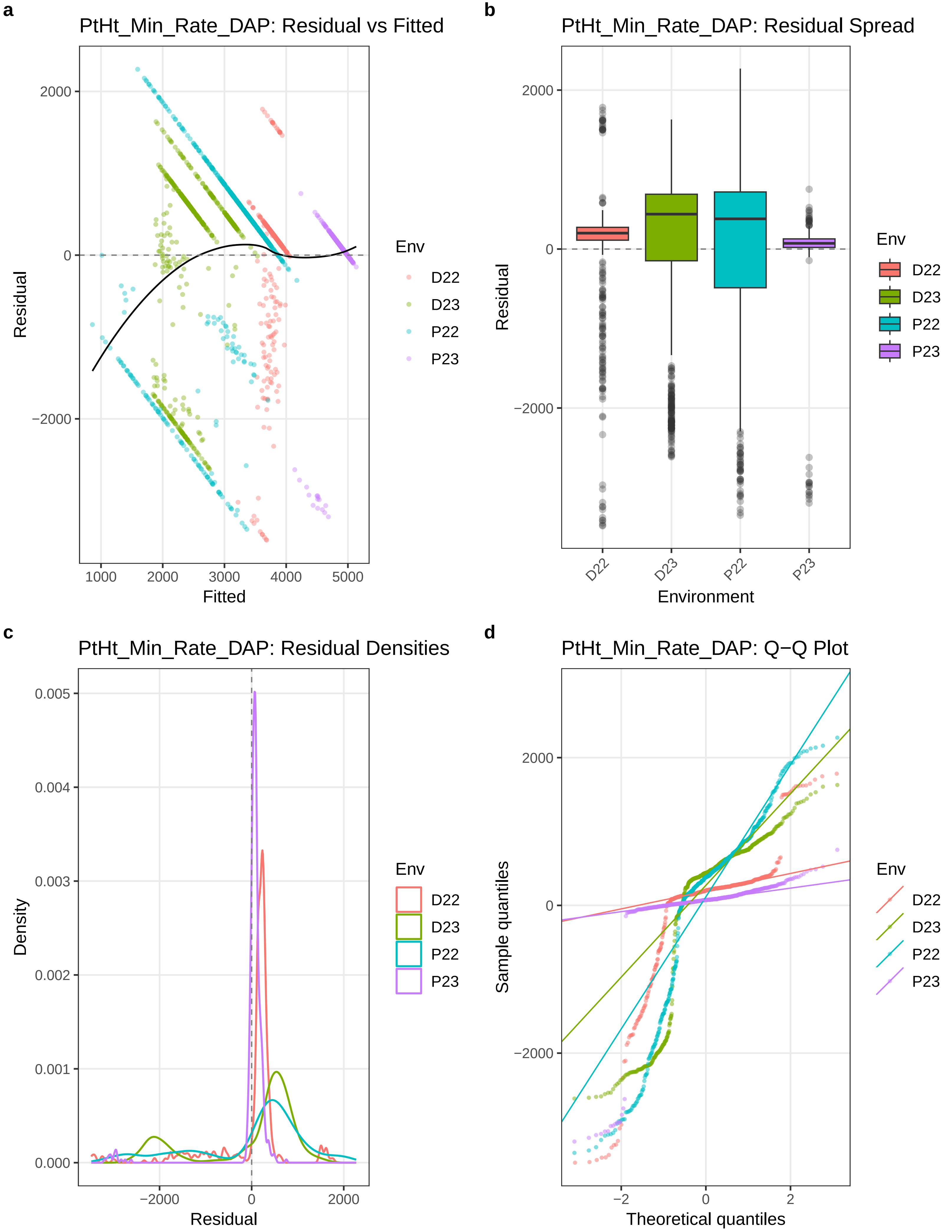

a
PtHt_Min_Rate_DAP: Residual vs Fitted
Residual
Fitted
2000
0
−2000
1000
2000
3000
4000
5000
Env
D22
D23
P22
P23
b
PtHt_Min_Rate_DAP: Residual Spread
Residual
Environment
D22
D23
P22
P23
c
PtHt_Min_Rate_DAP: Residual Densities
Density
Residual
0.005
0.004
0.003
0.002
0.001
0.000
−2000
0
2000
d
PtHt_Min_Rate_DAP: Q−Q Plot
Sample quantiles
Theoretical quantiles
−2
0
2

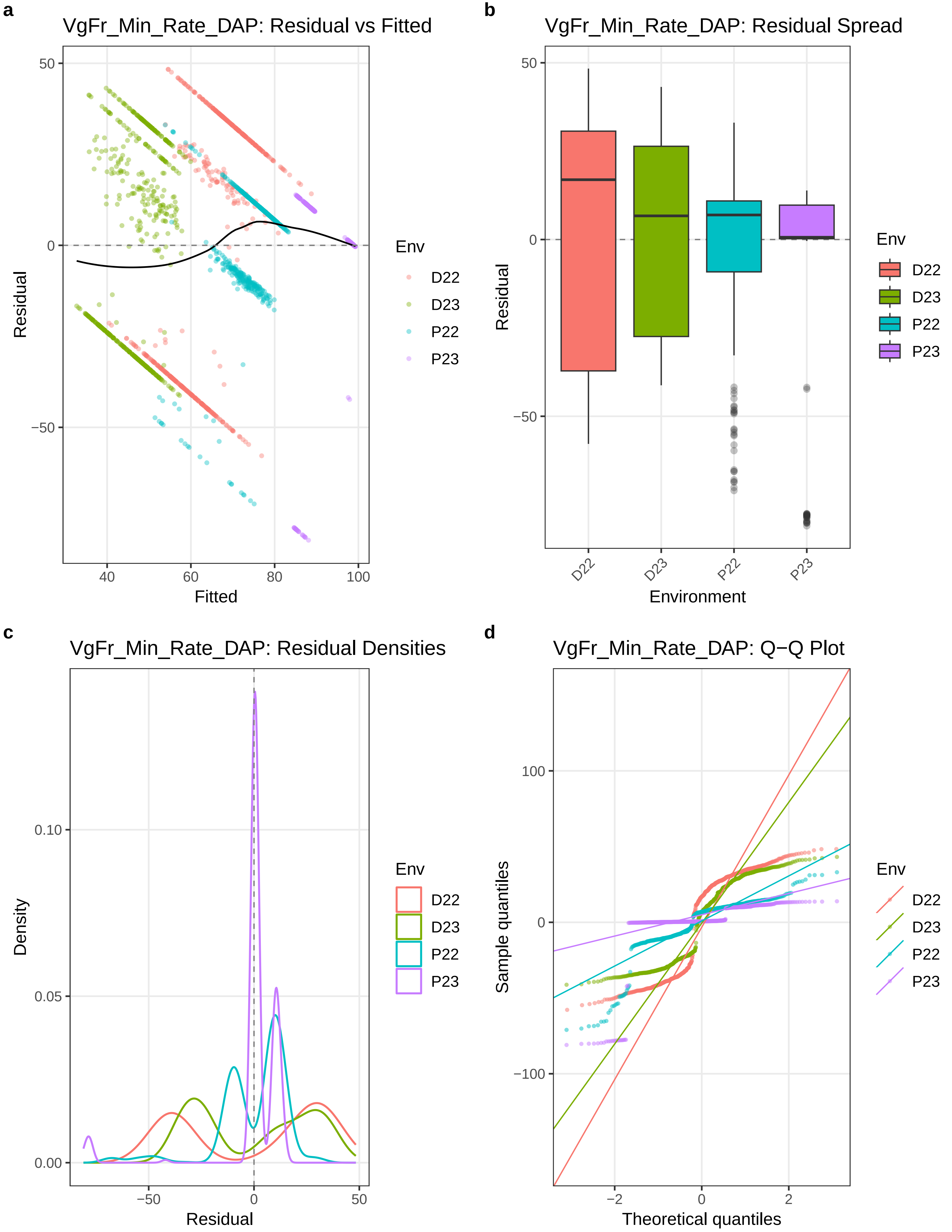

a
VgFr_Min_Rate_DAP: Residual vs Fitted
Residual
Fitted
Env
D22
D23
P22
P23
b
VgFr_Min_Rate_DAP: Residual Spread
Residual
Environment
c
VgFr_Min_Rate_DAP: Residual Densities
Density
Residual
d
VgFr_Min_Rate_DAP: Q−Q Plot
Sample quantiles
Theoretical quantiles

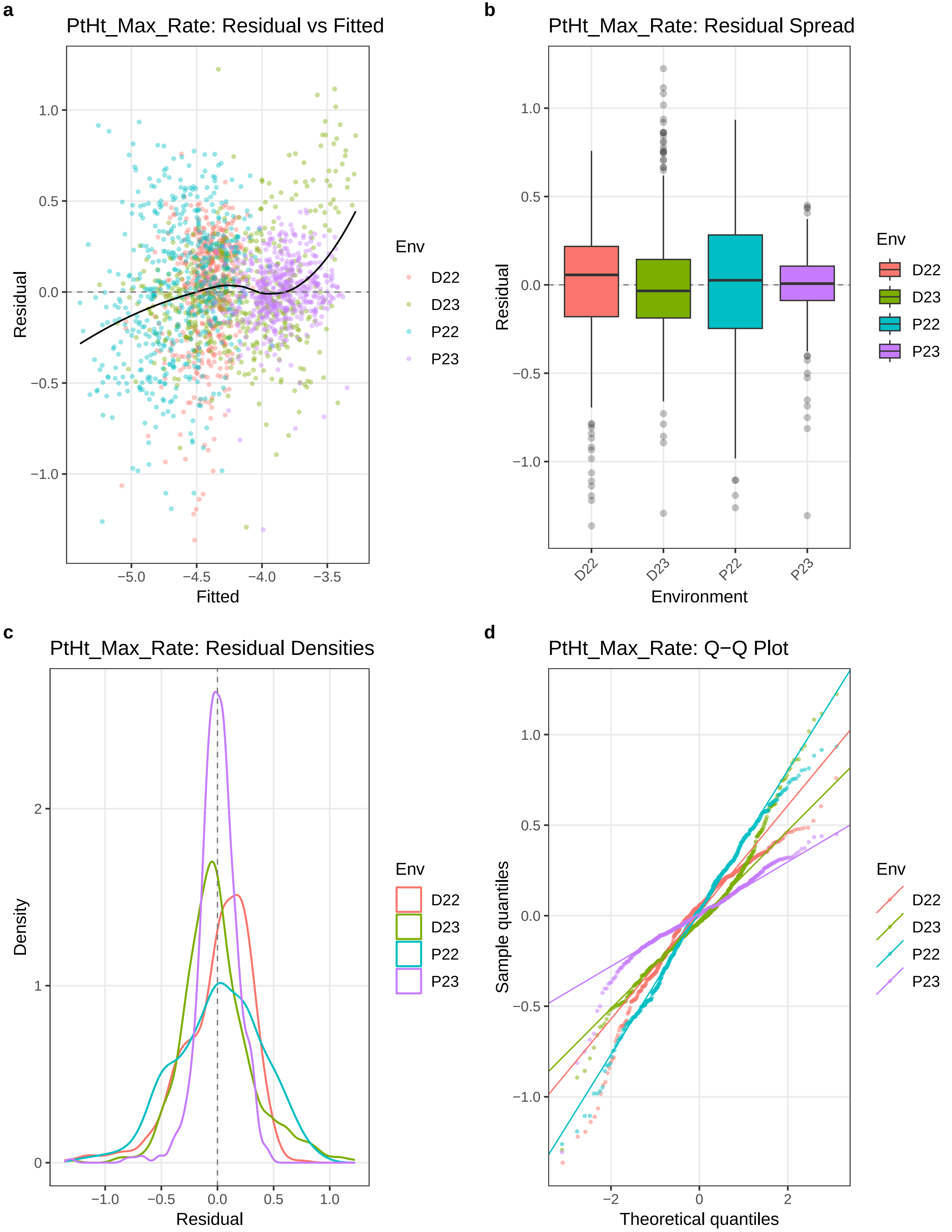

a
PtHt_Max_Rate: Residual vs Fitted
Residual
Fitted
Env
D22
D23
P22
P23
b
PtHt_Max_Rate: Residual Spread
Residual
Environment
c
PtHt_Max_Rate: Residual Densities
Density
Residual
d
PtHt_Max_Rate: Q−Q Plot
Sample quantiles
Theoretical quantiles

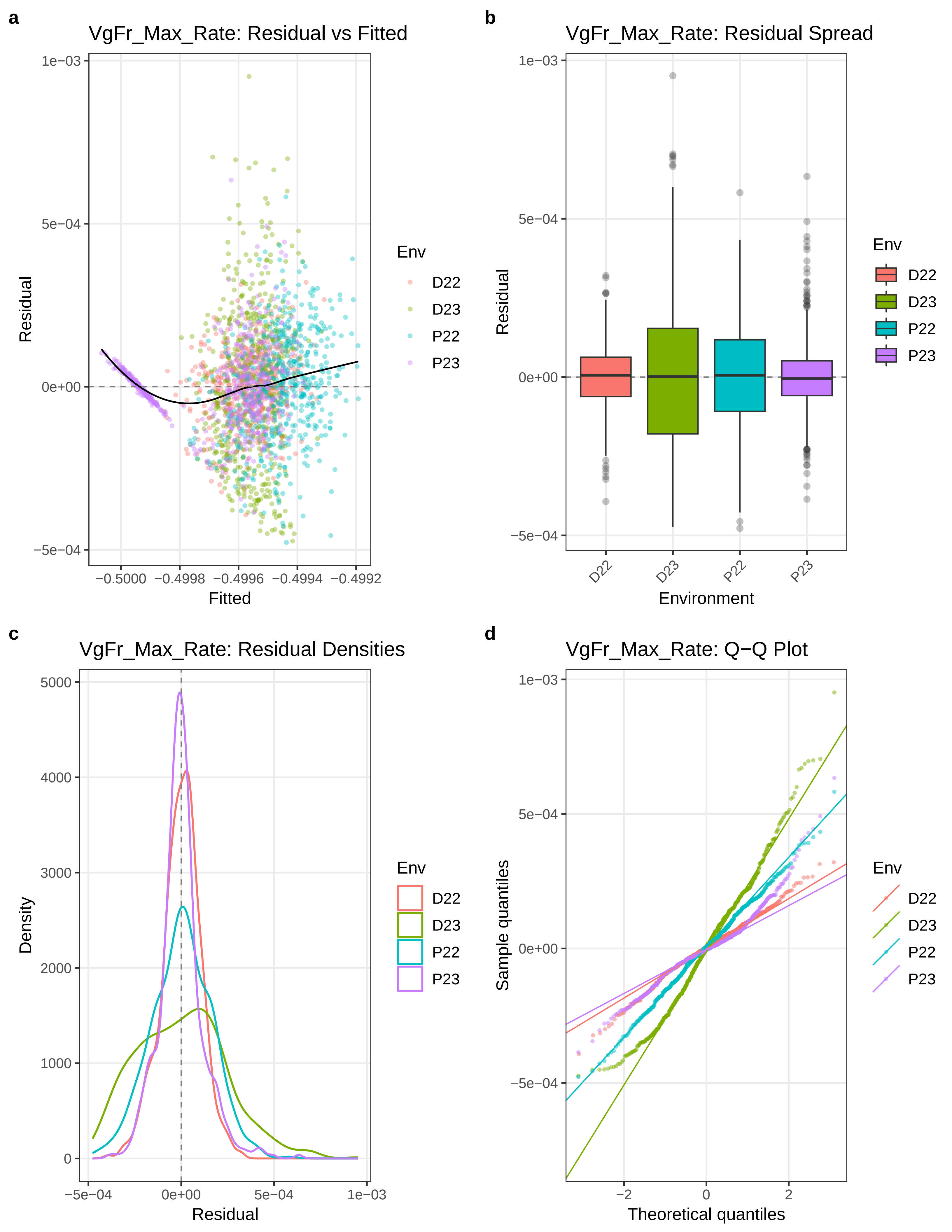
a
VgFr_Max_Rate: Residual vs Fitted
Residual
Fitted
Env
D22
D23
P22
P23
b
VgFr_Max_Rate: Residual Spread
Residual
Environment
c
VgFr_Max_Rate: Residual Densities
Density
Residual
d
VgFr_Max_Rate: Q−Q Plot
Sample quantiles
Theoretical quantiles

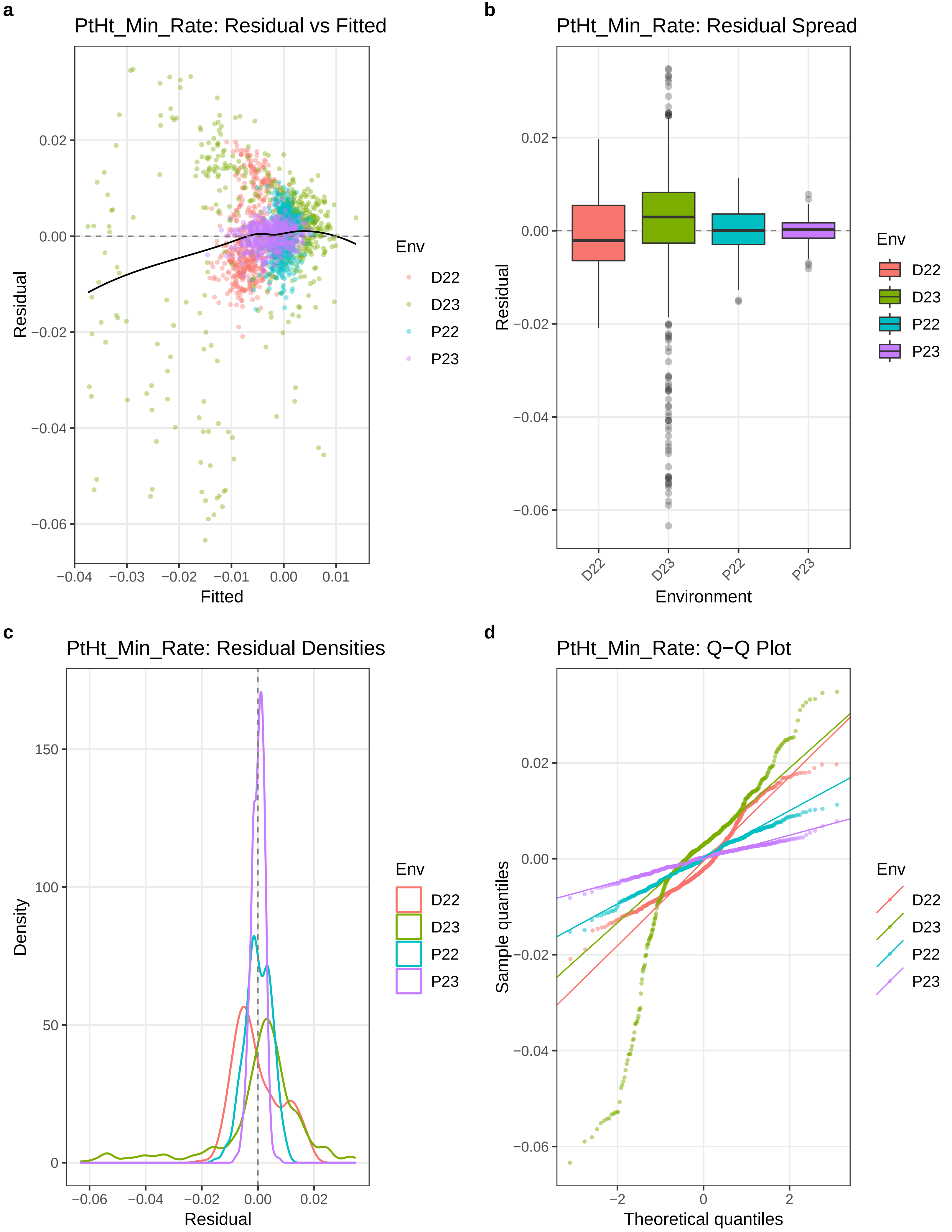
a
PtHt_Min_Rate: Residual vs Fitted
Residual
Fitted
0.02
0.00
−0.02
−0.04
−0.06
−0.04
−0.03
−0.02
−0.01
0.00
0.01
Env
D22
D23
P22
P23
b
PtHt_Min_Rate: Residual Spread
Residual
Environment
D22
D23
P22
P23
c
PtHt_Min_Rate: Residual Densities
Density
Residual
150
100
50
0
−0.06
−0.04
−0.02
0.00
0.02
d
PtHt_Min_Rate: Q−Q Plot
Sample quantiles
Theoretical quantiles
−2
0
2

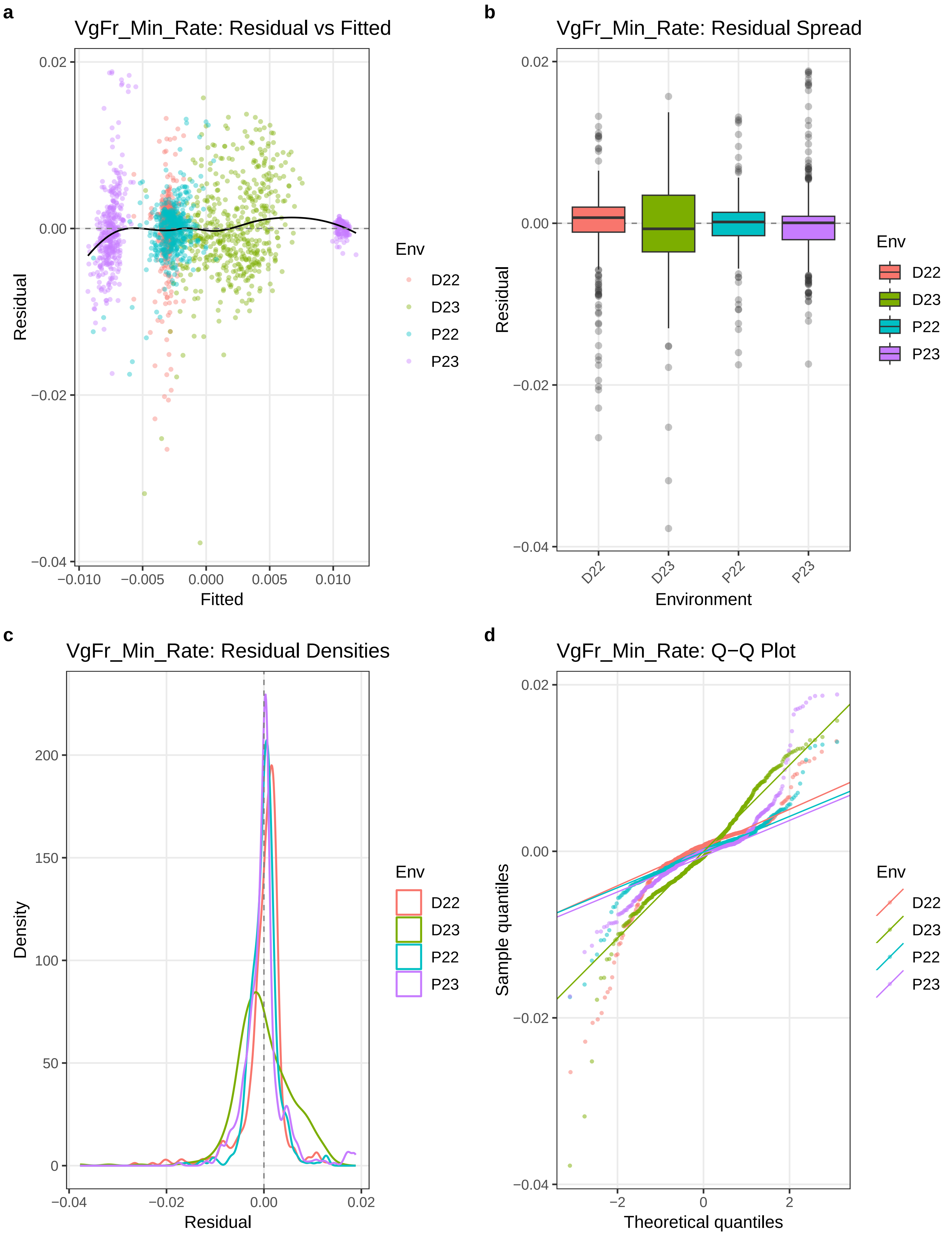
a
VgFr_Min_Rate: Residual vs Fitted
Residual
Fitted
0.02
0.00
−0.02
−0.04
−0.010
−0.005
0.000
0.005
0.010
Env
D22
D23
P22
P23
b
VgFr_Min_Rate: Residual Spread
Residual
Environment
0.02
0.00
−0.02
−0.04
D22
D23
P22
P23
Env
D22
D23
P22
P23
c
VgFr_Min_Rate: Residual Densities
Density
Residual
200
150
100
50
0
−0.04
−0.02
0.00
0.02
Env
D22
D23
P22
P23
d
VgFr_Min_Rate: Q−Q Plot
Sample quantiles
Theoretical quantiles
0.02
0.00
−0.02
−0.04
−2
0
2
Env
D22
D23
P22
P23

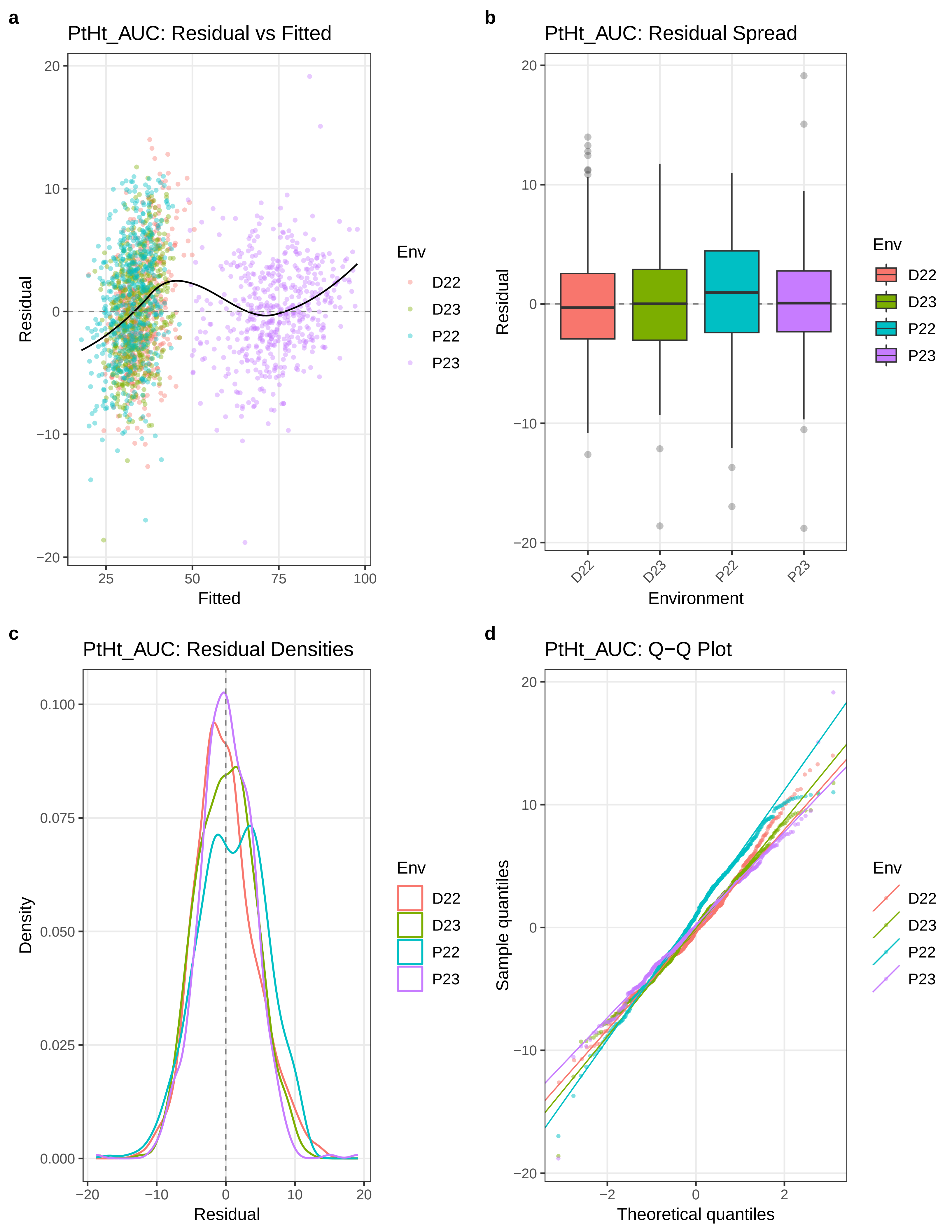

a
PtHt_AUC: Residual vs Fitted
Residual
Fitted
Env
D22
D23
P22
P23
b
PtHt_AUC: Residual Spread
Residual
Environment
c
PtHt_AUC: Residual Densities
Density
Residual
d
PtHt_AUC: Q−Q Plot
Sample quantiles
Theoretical quantiles

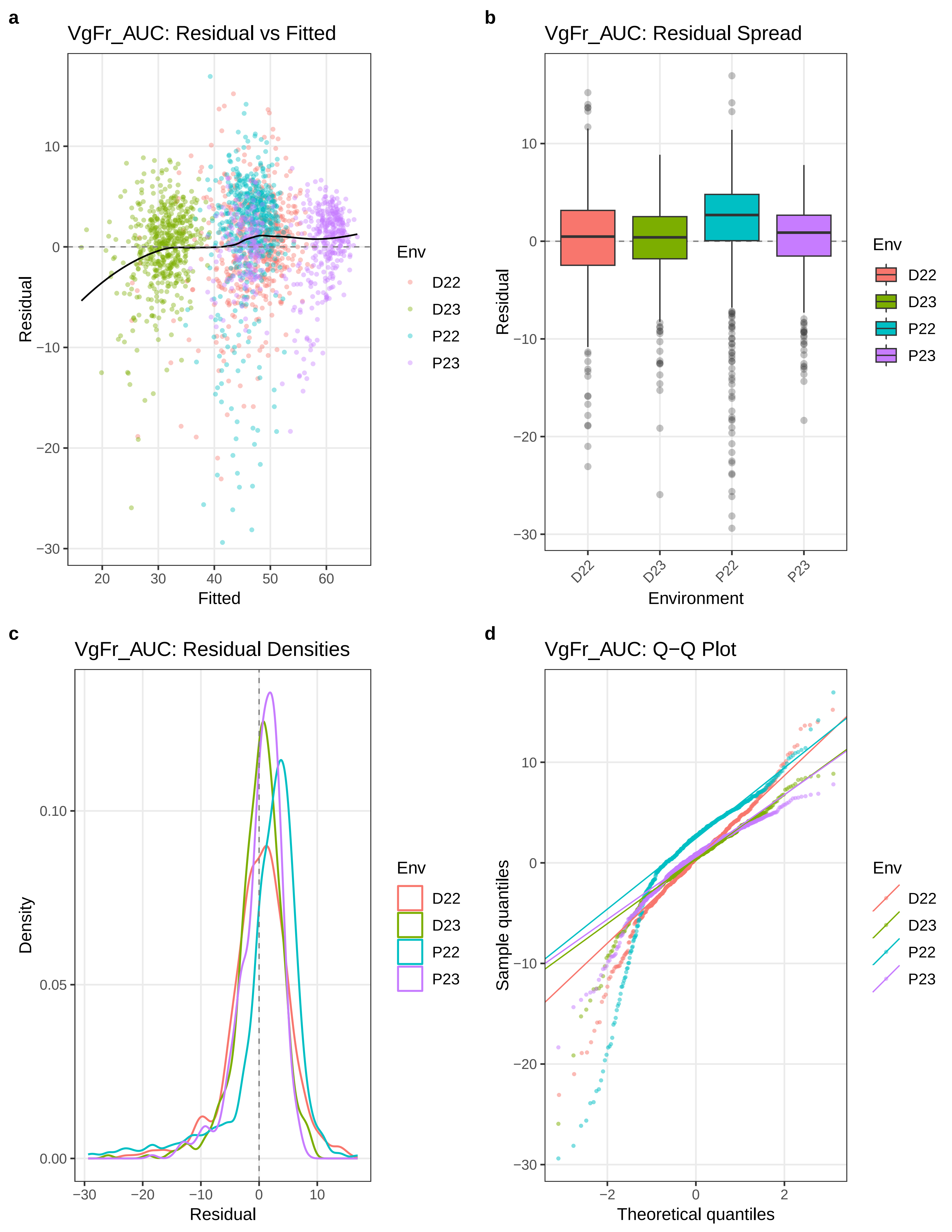

a
VgFr_AUC: Residual vs Fitted
Residual
Fitted
Env
D22
D23
P22
P23
b
VgFr_AUC: Residual Spread
Residual
Environment
D22
D23
P22
P23
Env
D22
D23
P22
P23
c
VgFr_AUC: Residual Densities
Density
Residual
Env
D22
D23
P22
P23
d
VgFr_AUC: Q−Q Plot
Sample quantiles
Theoretical quantiles
Env
D22
D23
P22
P23

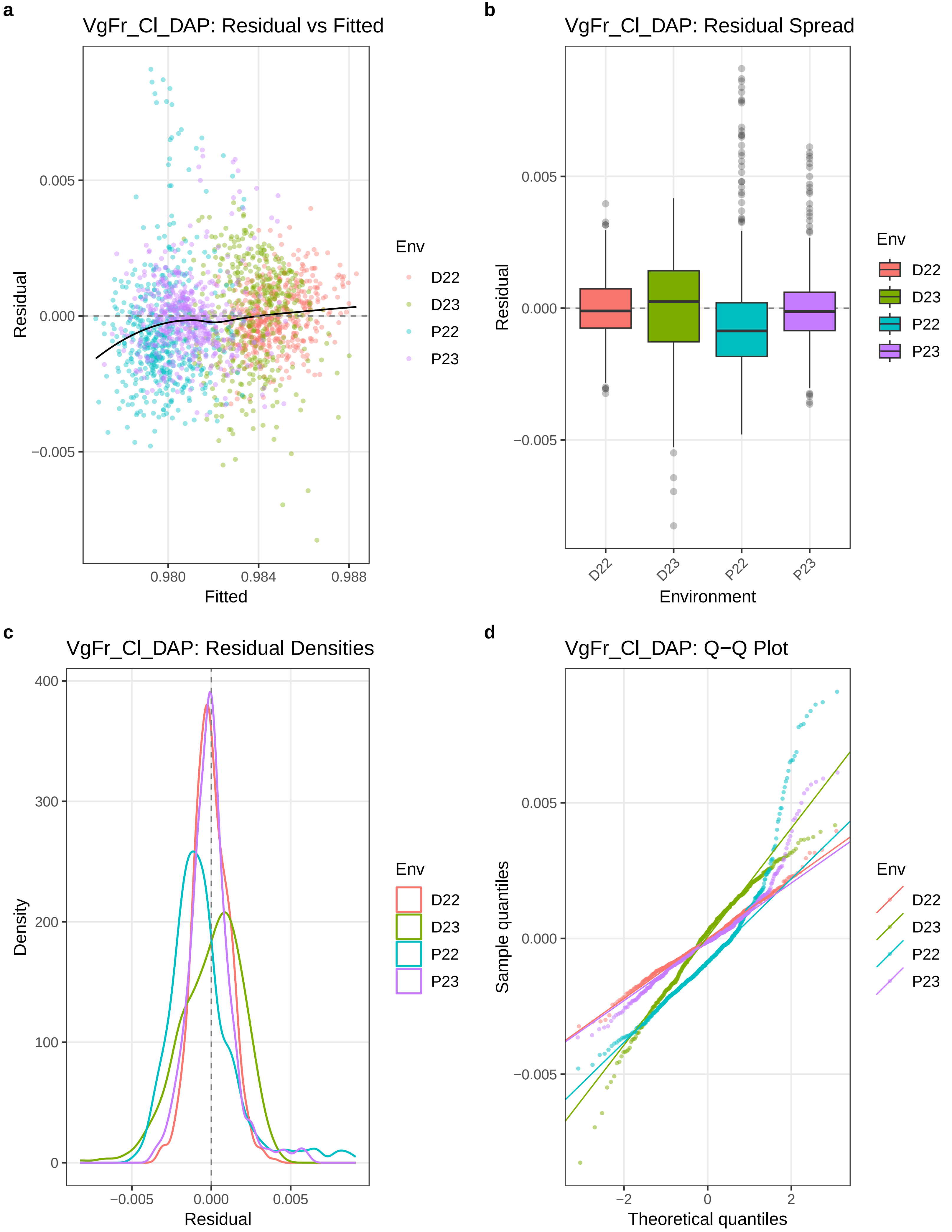
a
VgFr_Cl_DAP: Residual vs Fitted
Residual
Fitted
0.005
0.000
−0.005
0.980
0.984
0.988
Env
D22
D23
P22
P23
b
VgFr_Cl_DAP: Residual Spread
Residual
Environment
D22
D23
P22
P23
c
VgFr_Cl_DAP: Residual Densities
Density
Residual
400
300
200
100
0
−0.005
0.000
0.005
d
VgFr_Cl_DAP: Q−Q Plot
Sample quantiles
Theoretical quantiles
−2
0
2

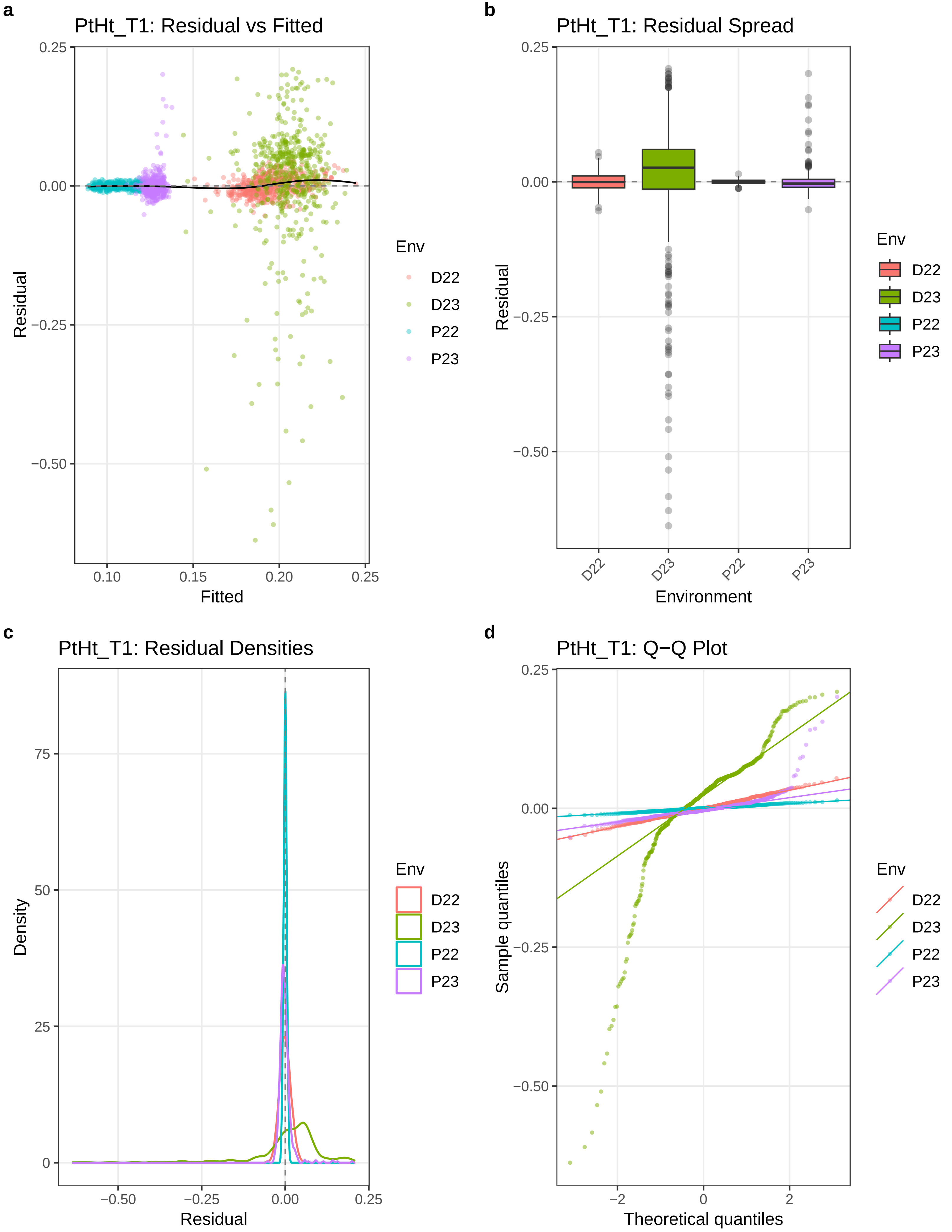
a
PtHt_T1: Residual vs Fitted
Residual
Fitted
0.25
0.00
−0.25
−0.50
0.10
0.15
0.20
0.25
Env
D22
D23
P22
P23
b
PtHt_T1: Residual Spread
Residual
Environment
D22
D23
P22
P23
c
PtHt_T1: Residual Densities
Density
Residual
75
50
25
0
−0.50
−0.25
0.00
0.25
d
PtHt_T1: Q−Q Plot
Sample quantiles
Theoretical quantiles
−2
0
2

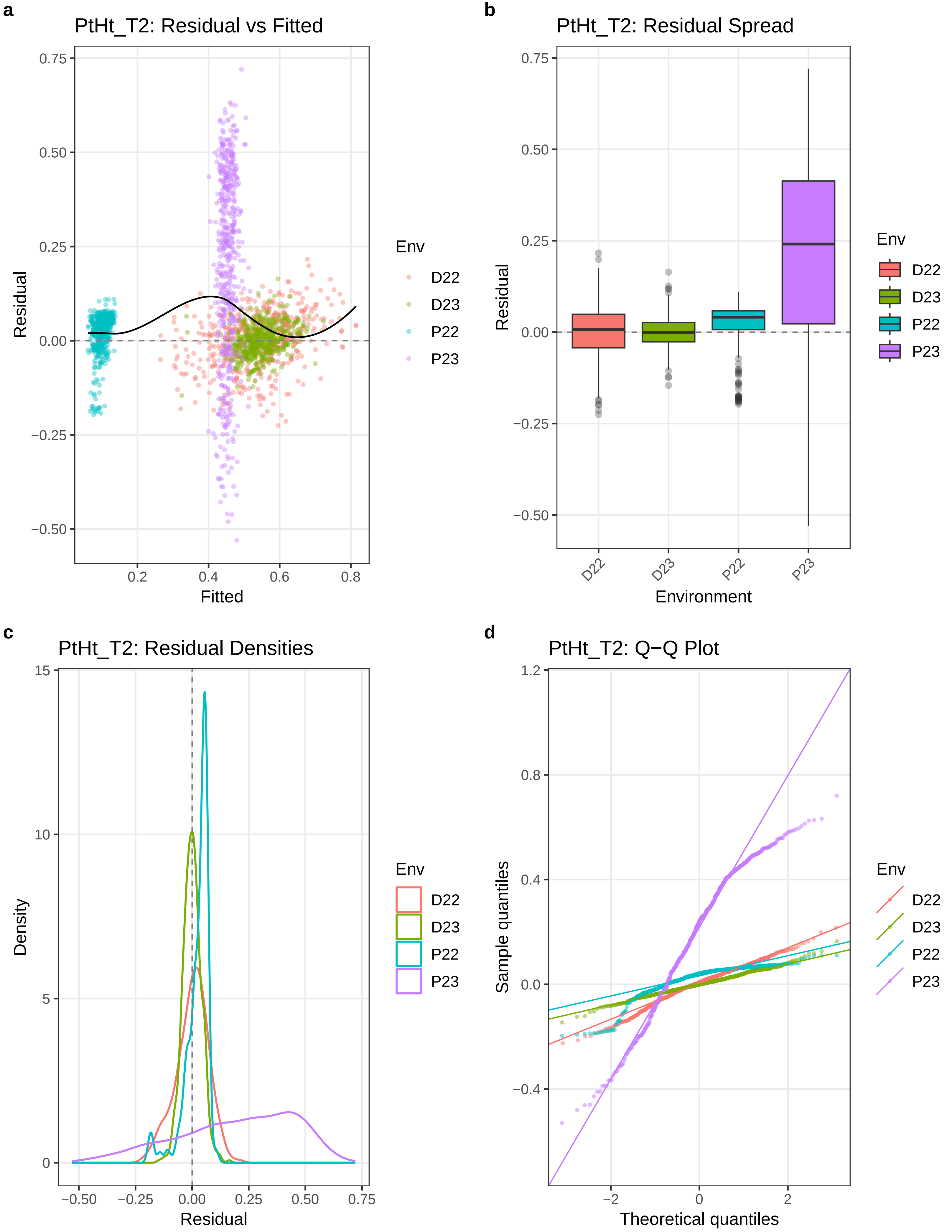

a
PtHt_T2: Residual vs Fitted
Residual
Fitted
0.75
0.50
0.25
0.00
−0.25
−0.50
0.2
0.4
0.6
0.8
Env
D22
D23
P22
P23
b
PtHt_T2: Residual Spread
Residual
Environment
D22
D23
P22
P23
c
PtHt_T2: Residual Densities
Density
Residual
15
10
5
0
−0.50
−0.25
0.00
0.25
0.50
0.75
d
PtHt_T2: Q−Q Plot
Sample quantiles
Theoretical quantiles
1.2
0.8
0.4
0.0
−0.4
−2
0
2

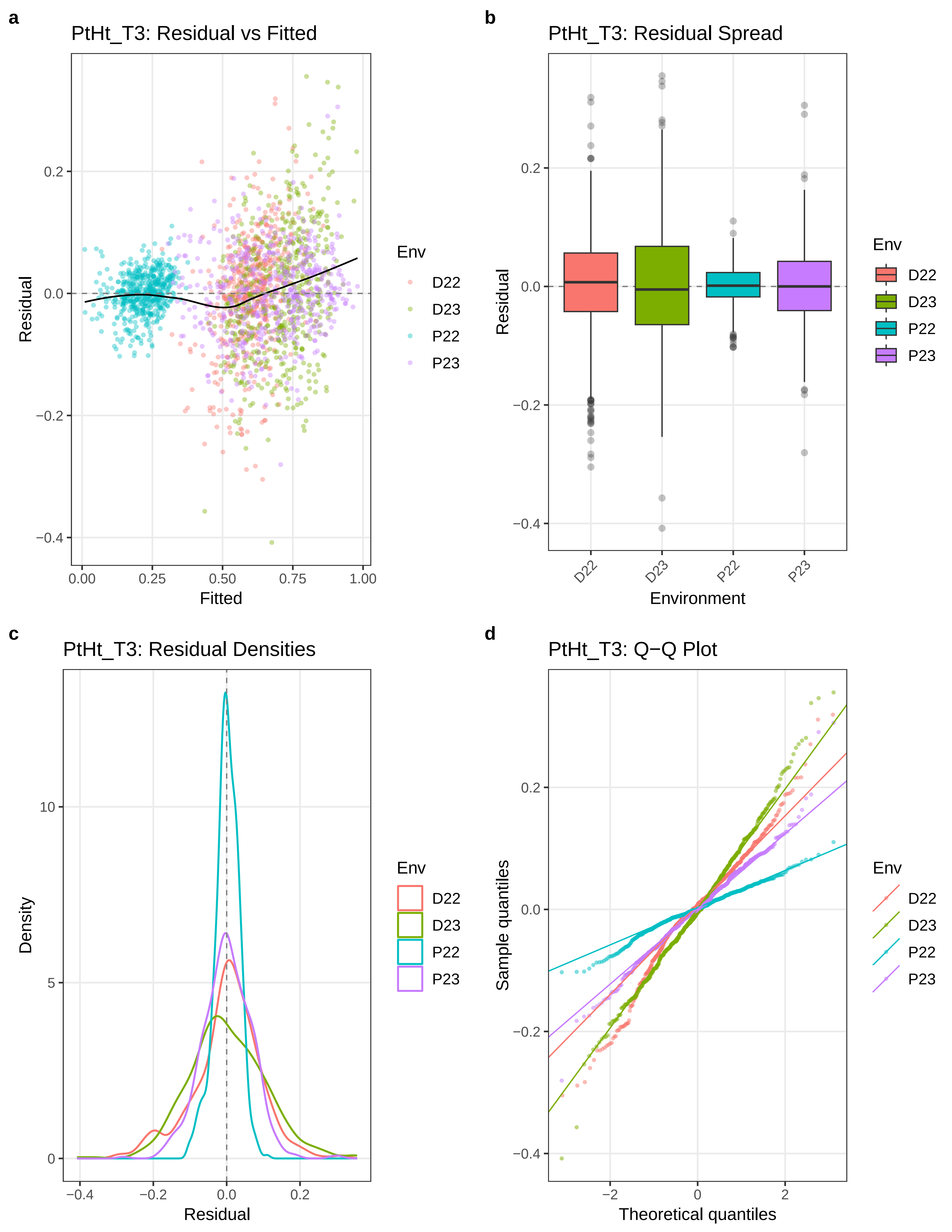
a
PtHt_T3: Residual vs Fitted
Residual
Fitted
0.2
0.0
−0.2
−0.4
0.00
0.25
0.50
0.75
1.00
Env
D22
D23
P22
P23
b
PtHt_T3: Residual Spread
Residual
Environment
D22
D23
P22
P23
c
PtHt_T3: Residual Densities
Density
Residual
10
5
0
−0.4
−0.2
0.0
0.2
d
PtHt_T3: Q−Q Plot
Sample quantiles
Theoretical quantiles
−2
0
2

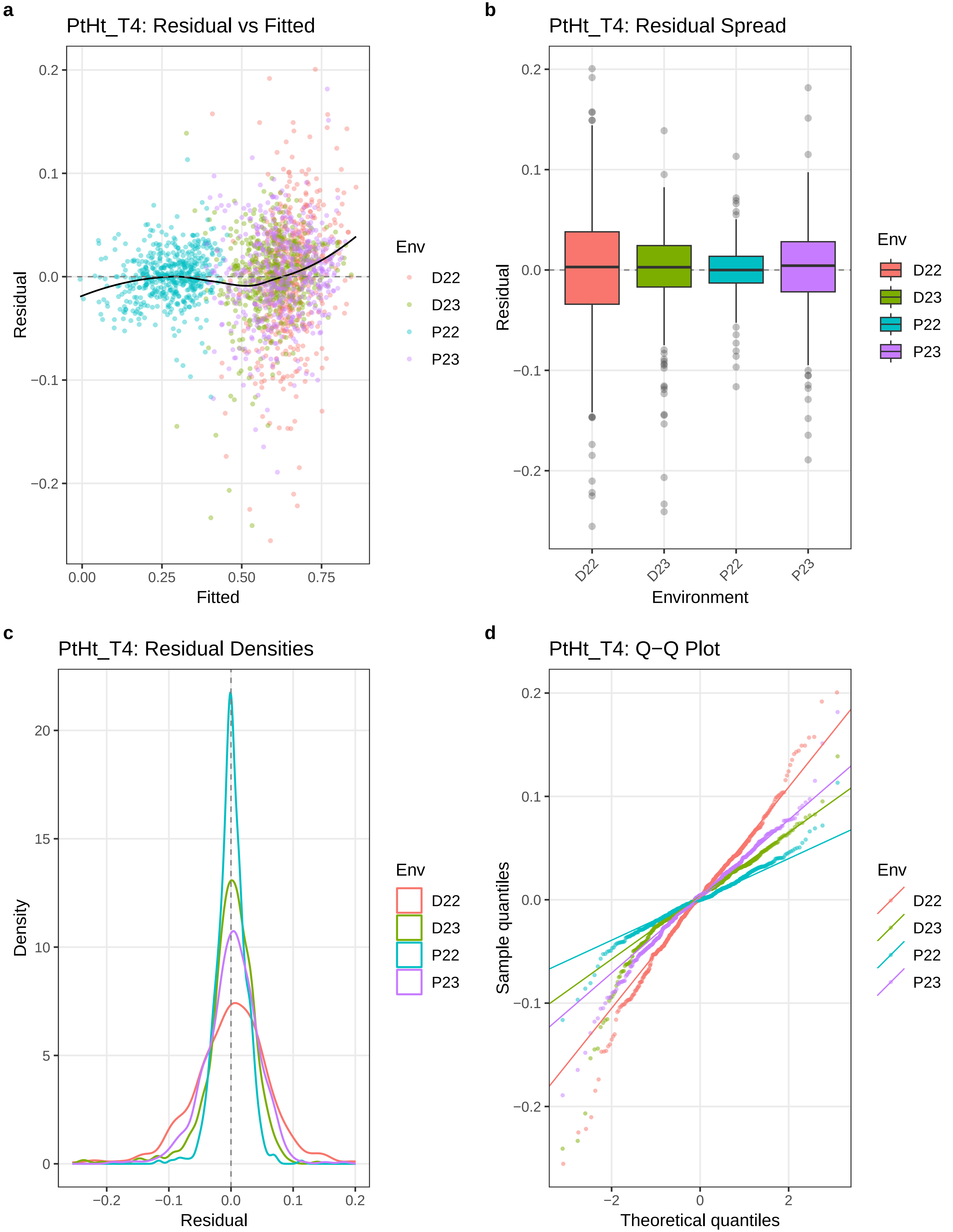
a
PtHt_T4: Residual vs Fitted
Residual
Fitted
0.2
0.1
0.0
−0.1
−0.2
0.00
0.25
0.50
0.75
Env
D22
D23
P22
P23
b
PtHt_T4: Residual Spread
Residual
Environment
D22
D23
P22
P23
c
PtHt_T4: Residual Densities
Density
Residual
20
15
10
5
0
−0.2
−0.1
0.0
0.1
0.2
d
PtHt_T4: Q−Q Plot
Sample quantiles
Theoretical quantiles
−2
0
2

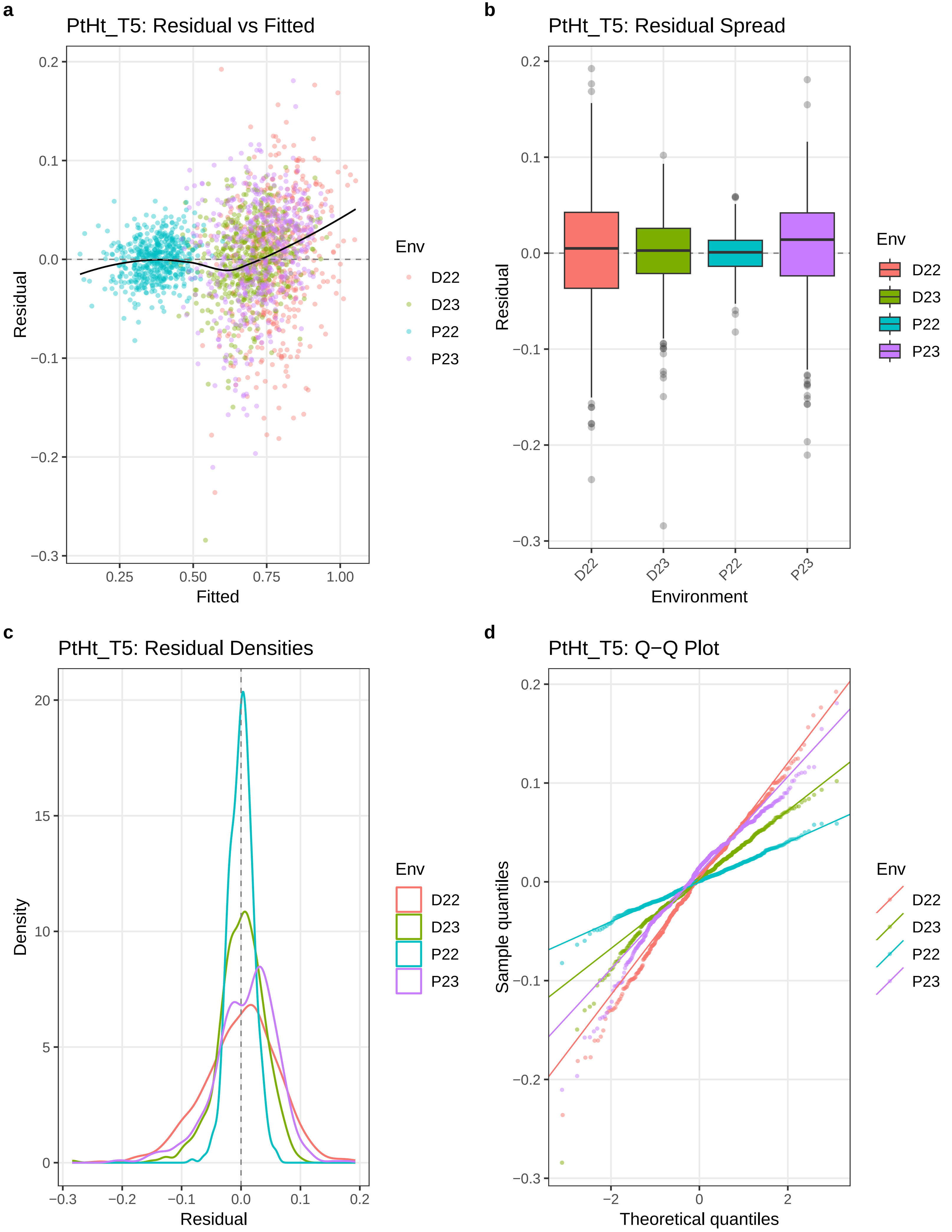

a
PtHt_T5: Residual vs Fitted
Residual
Fitted
0.2
0.1
0.0
−0.1
−0.2
−0.3
0.25
0.50
0.75
1.00
Env
D22
D23
P22
P23
b
PtHt_T5: Residual Spread
Residual
Environment
D22
D23
P22
P23
c
PtHt_T5: Residual Densities
Density
Residual
20
15
10
5
0
−0.3
−0.2
−0.1
0.0
0.1
0.2
d
PtHt_T5: Q−Q Plot
Sample quantiles
Theoretical quantiles
−2
0
2

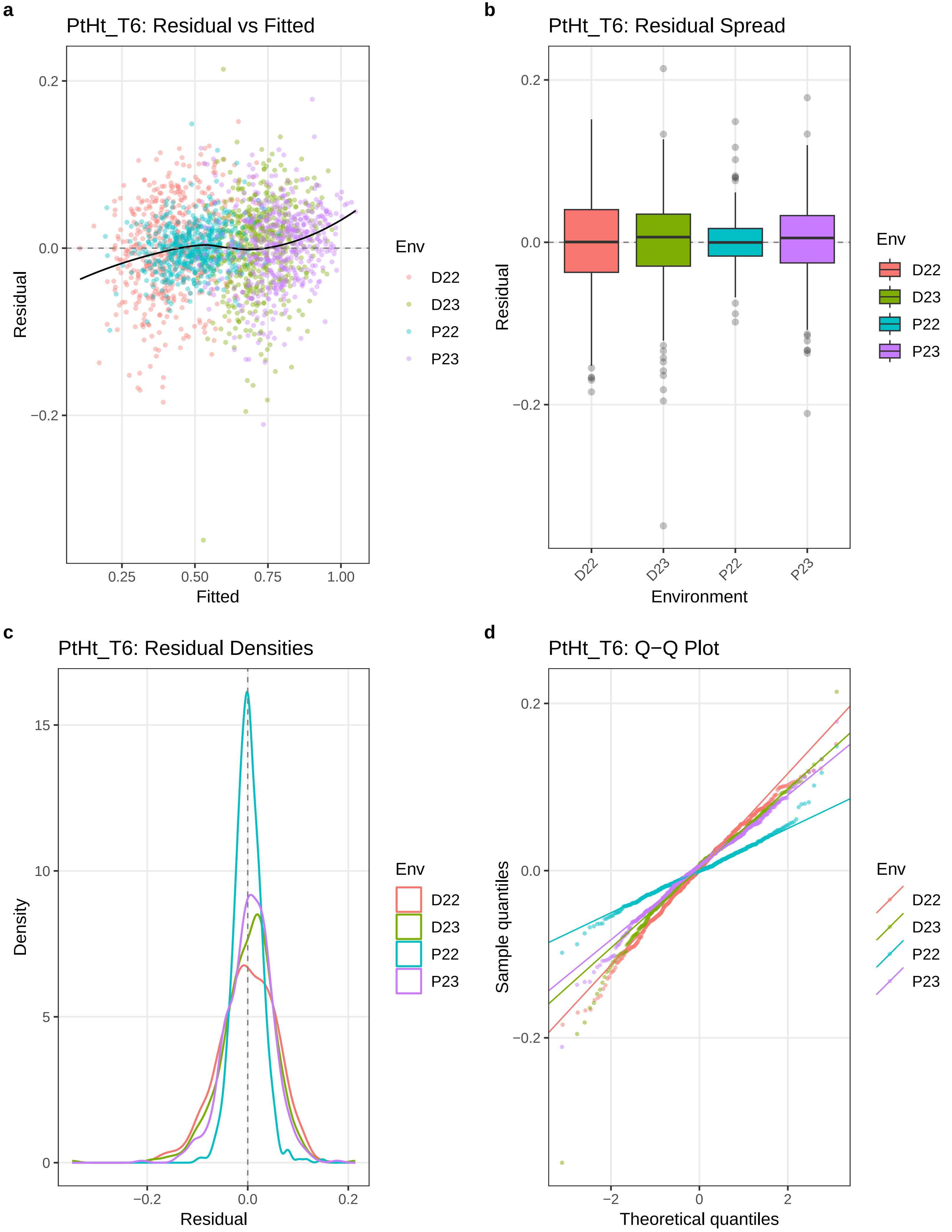

a
PtHt_T6: Residual vs Fitted
Residual
Fitted
0.2
0.0
−0.2
0.25
0.50
0.75
1.00
Env
D22
D23
P22
P23
b
PtHt_T6: Residual Spread
Residual
Environment
0.2
0.0
−0.2
D22
D23
P22
P23
Env
D22
D23
P22
P23
c
PtHt_T6: Residual Densities
Density
Residual
15
10
5
0
−0.2
0.0
0.2
Env
D22
D23
P22
P23
d
PtHt_T6: Q−Q Plot
Sample quantiles
Theoretical quantiles
0.2
0.0
−0.2
−2
0
2
Env
D22
D23
P22
P23

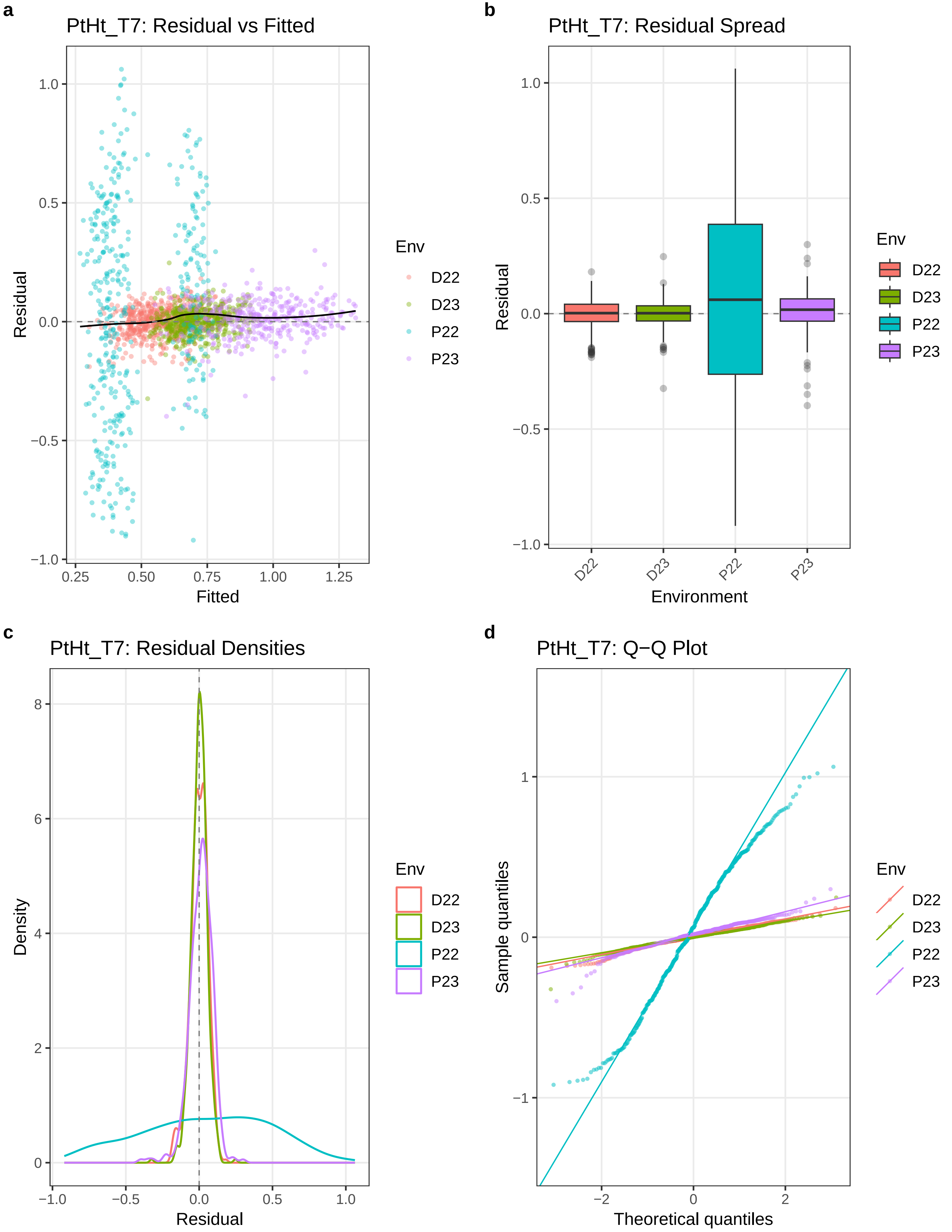
a
PtHt_T7: Residual vs Fitted
Residual
Fitted
Env
D22
D23
P22
P23
b
PtHt_T7: Residual Spread
Residual
Environment
Env
D22
D23
P22
P23
c
PtHt_T7: Residual Densities
Density
Residual
Env
D22
D23
P22
P23
d
PtHt_T7: Q−Q Plot
Sample quantiles
Theoretical quantiles
Env
D22
D23
P22
P23

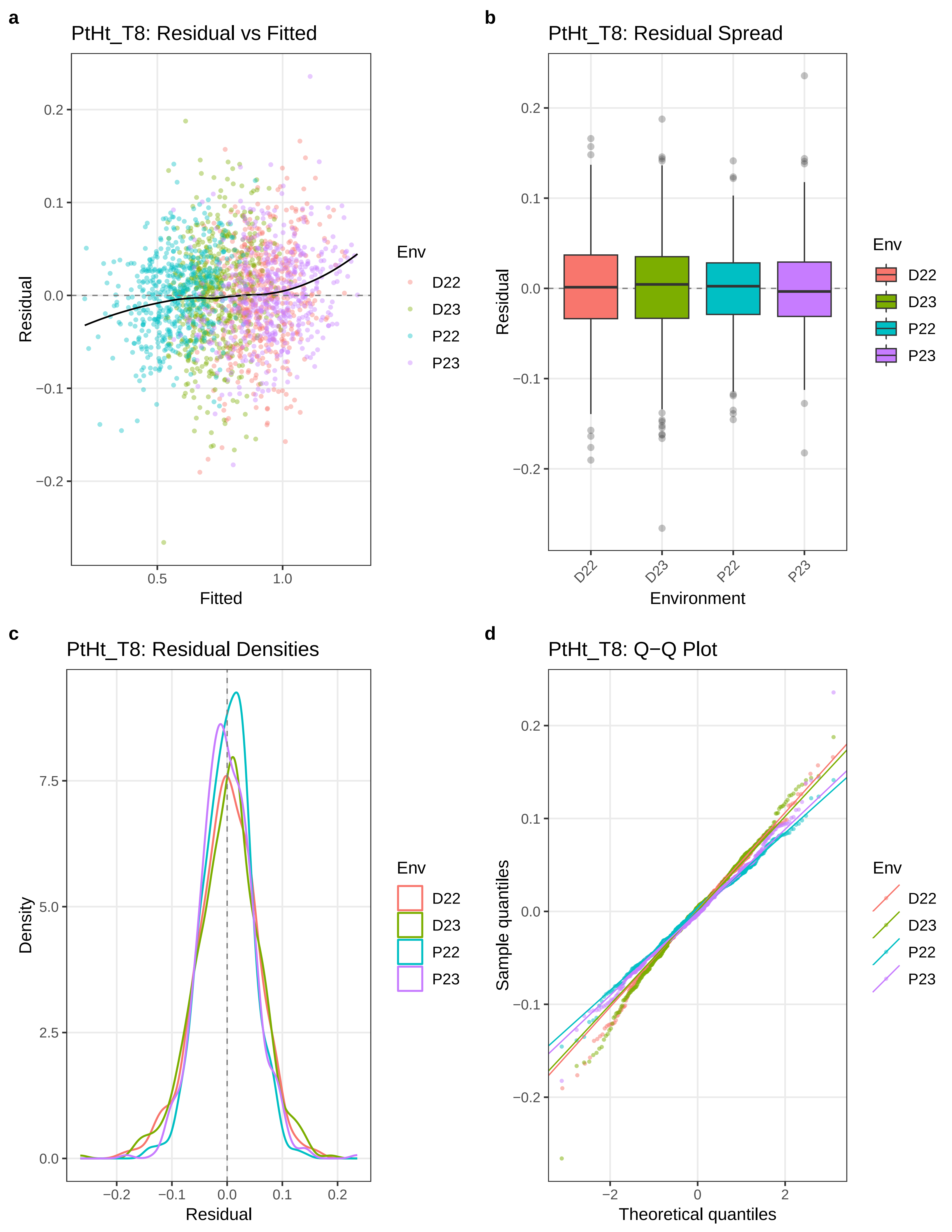

a
PtHt_T8: Residual vs Fitted
Residual
Fitted
Env
D22
D23
P22
P23
b
PtHt_T8: Residual Spread
Residual
Environment
c
PtHt_T8: Residual Densities
Density
Residual
d
PtHt_T8: Q−Q Plot
Sample quantiles
Theoretical quantiles

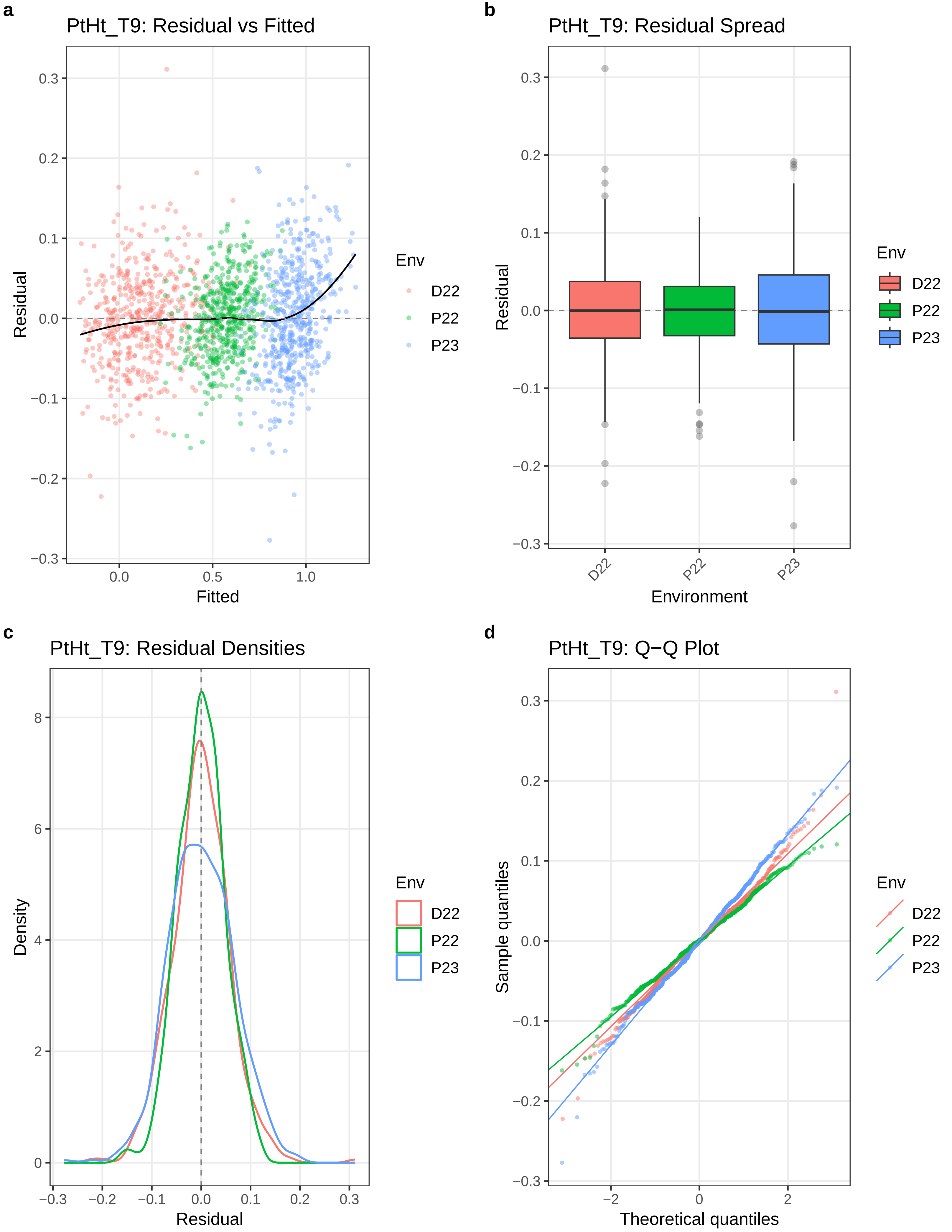

a
PtHt_T9: Residual vs Fitted
Residual
Fitted
0.3
0.2
0.1
0.0
−0.1
−0.2
−0.3
0.0
0.5
1.0
Env
D22
P22
P23
b
PtHt_T9: Residual Spread
Residual
Environment
D22
P22
P23
Env
D22
P22
P23
c
PtHt_T9: Residual Densities
Density
Residual
8
6
4
2
0
−0.3
−0.2
−0.1
0.0
0.1
0.2
0.3
Env
D22
P22
P23
d
PtHt_T9: Q−Q Plot
Sample quantiles
Theoretical quantiles
−2
0
2
Env
D22
P22
P23

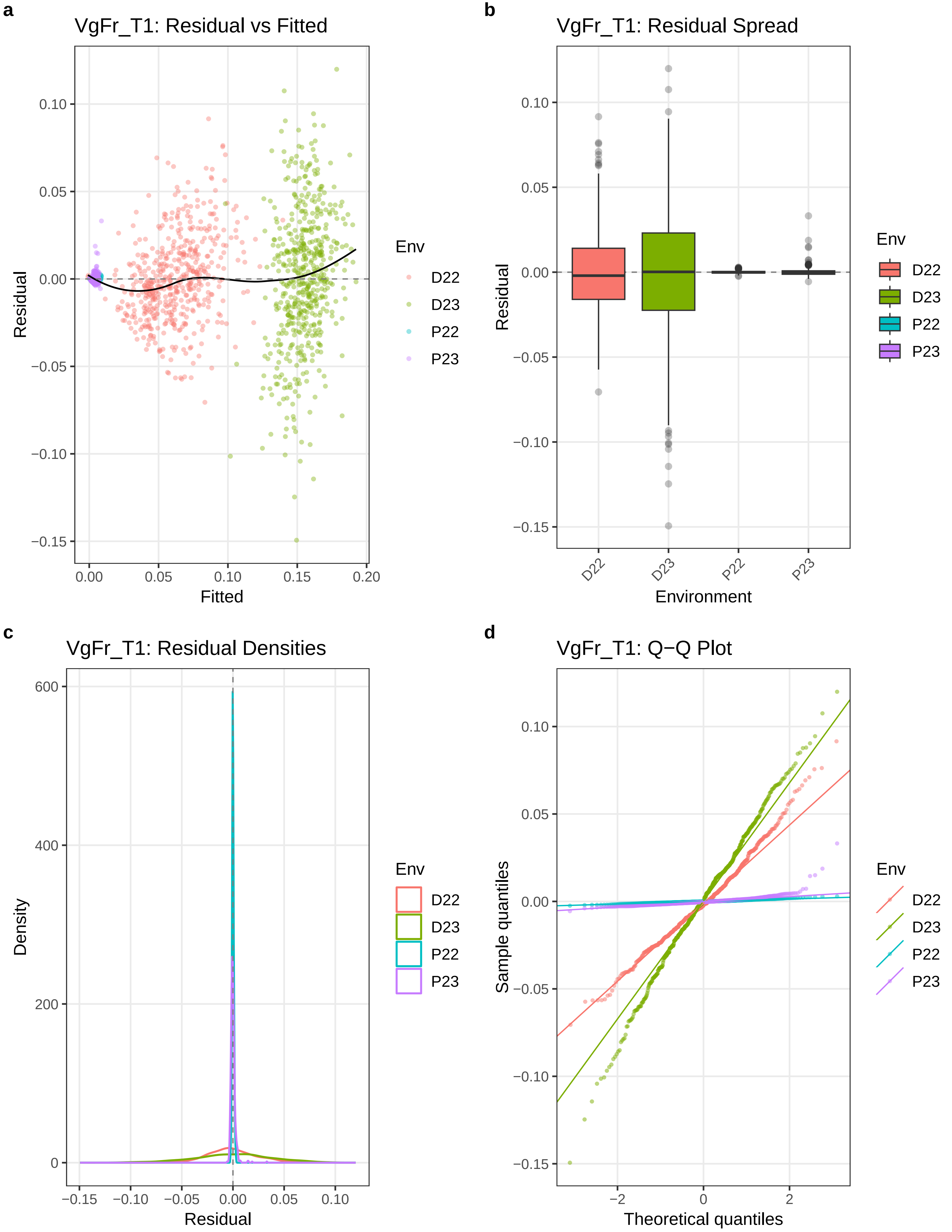
a
VgFr_T1: Residual vs Fitted
Residual
Fitted
Env
D22
D23
P22
P23
b
VgFr_T1: Residual Spread
Residual
Environment
D22
D23
P22
P23
c
VgFr_T1: Residual Densities
Density
Residual
d
VgFr_T1: Q−Q Plot
Sample quantiles
Theoretical quantiles

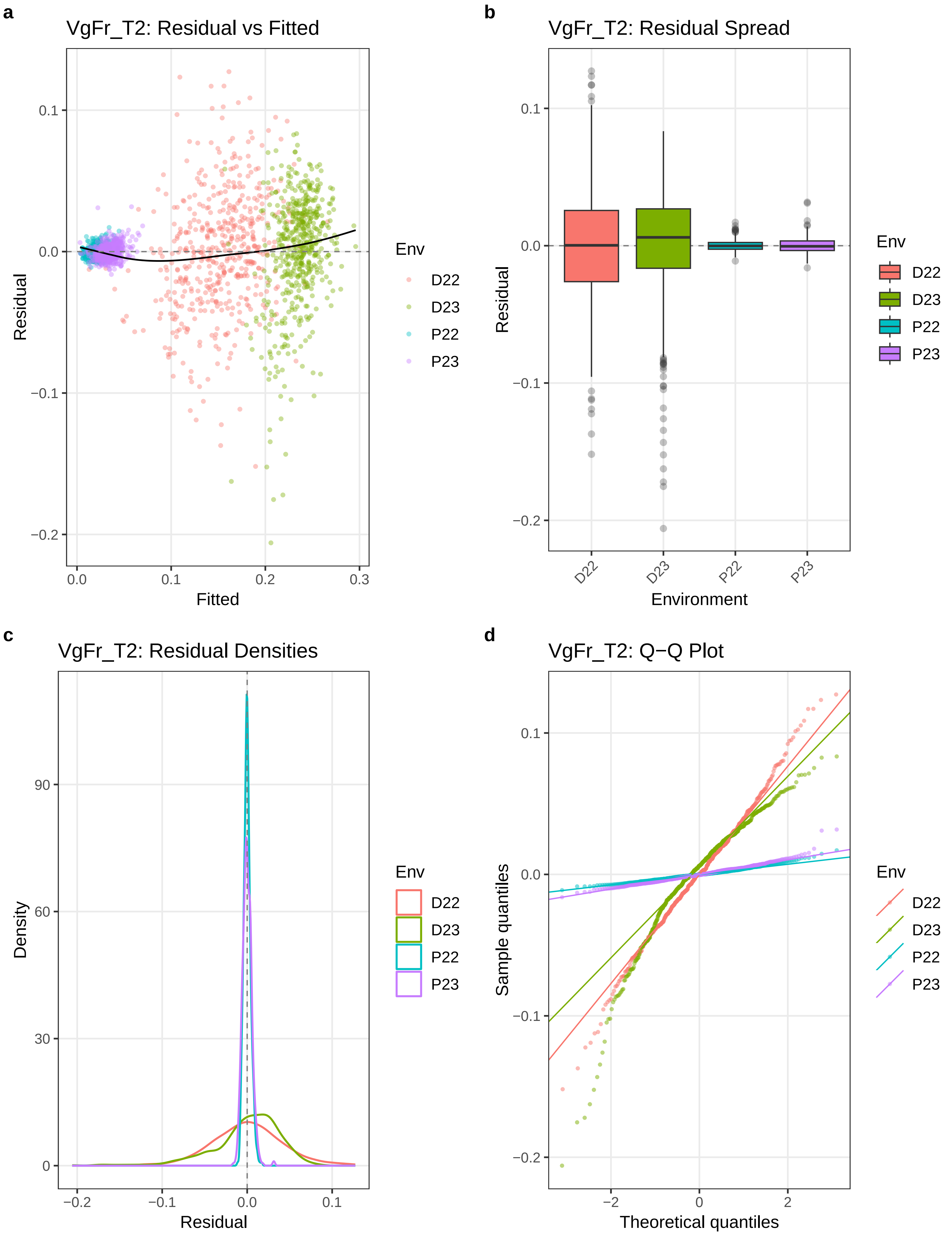
a
VgFr_T2: Residual vs Fitted
Residual
Fitted
Env
D22
D23
P22
P23
b
VgFr_T2: Residual Spread
Residual
Environment
c
VgFr_T2: Residual Densities
Density
Residual
d
VgFr_T2: Q−Q Plot
Sample quantiles
Theoretical quantiles

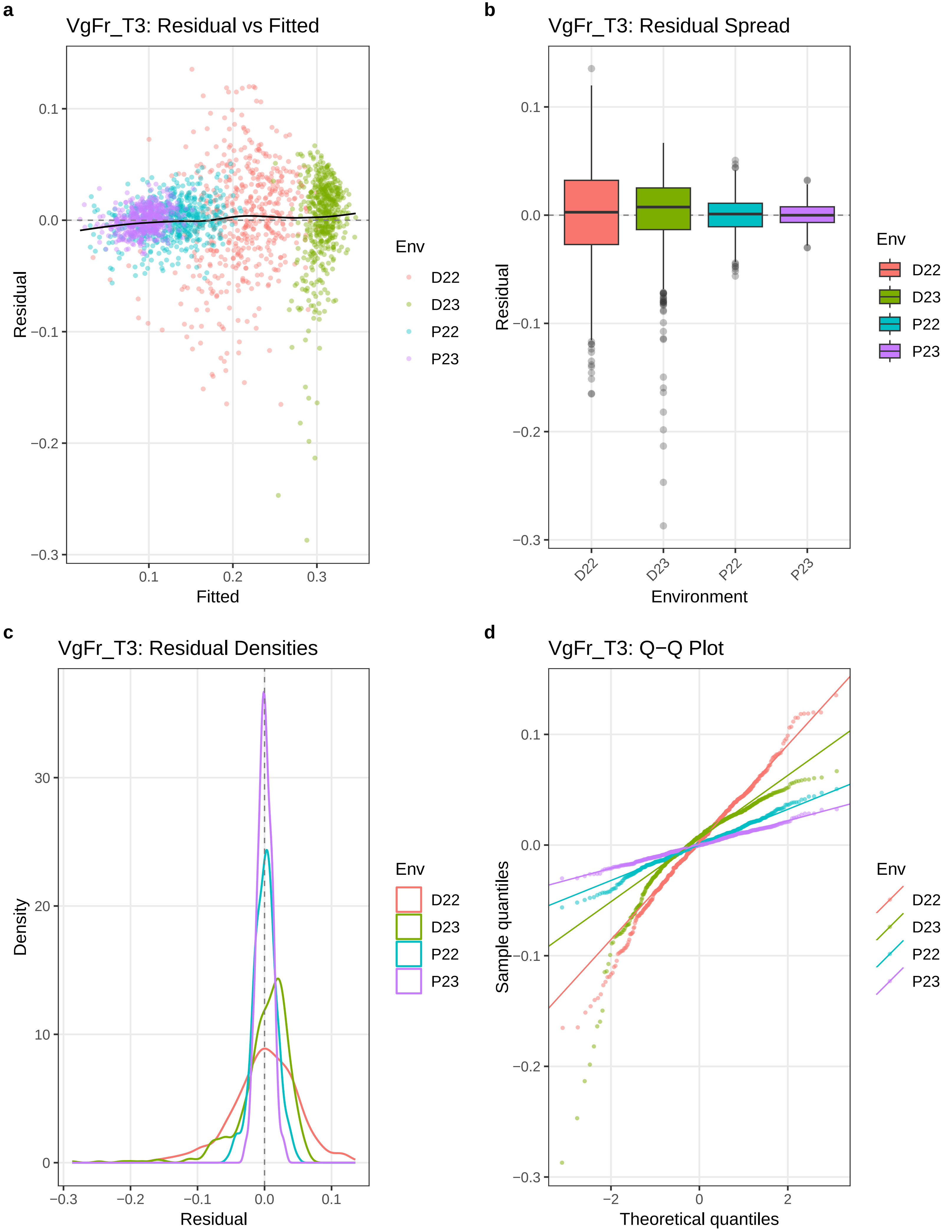

a
VgFr_T3: Residual vs Fitted
Residual
Fitted
0.1
0.0
−0.1
−0.2
−0.3
0.1
0.2
0.3
Env
D22
D23
P22
P23
b
VgFr_T3: Residual Spread
Residual
Environment
D22
D23
P22
P23
c
VgFr_T3: Residual Densities
Density
Residual
30
20
10
0
−0.3
−0.2
−0.1
0.0
0.1
d
VgFr_T3: Q−Q Plot
Sample quantiles
Theoretical quantiles
−2
0
2

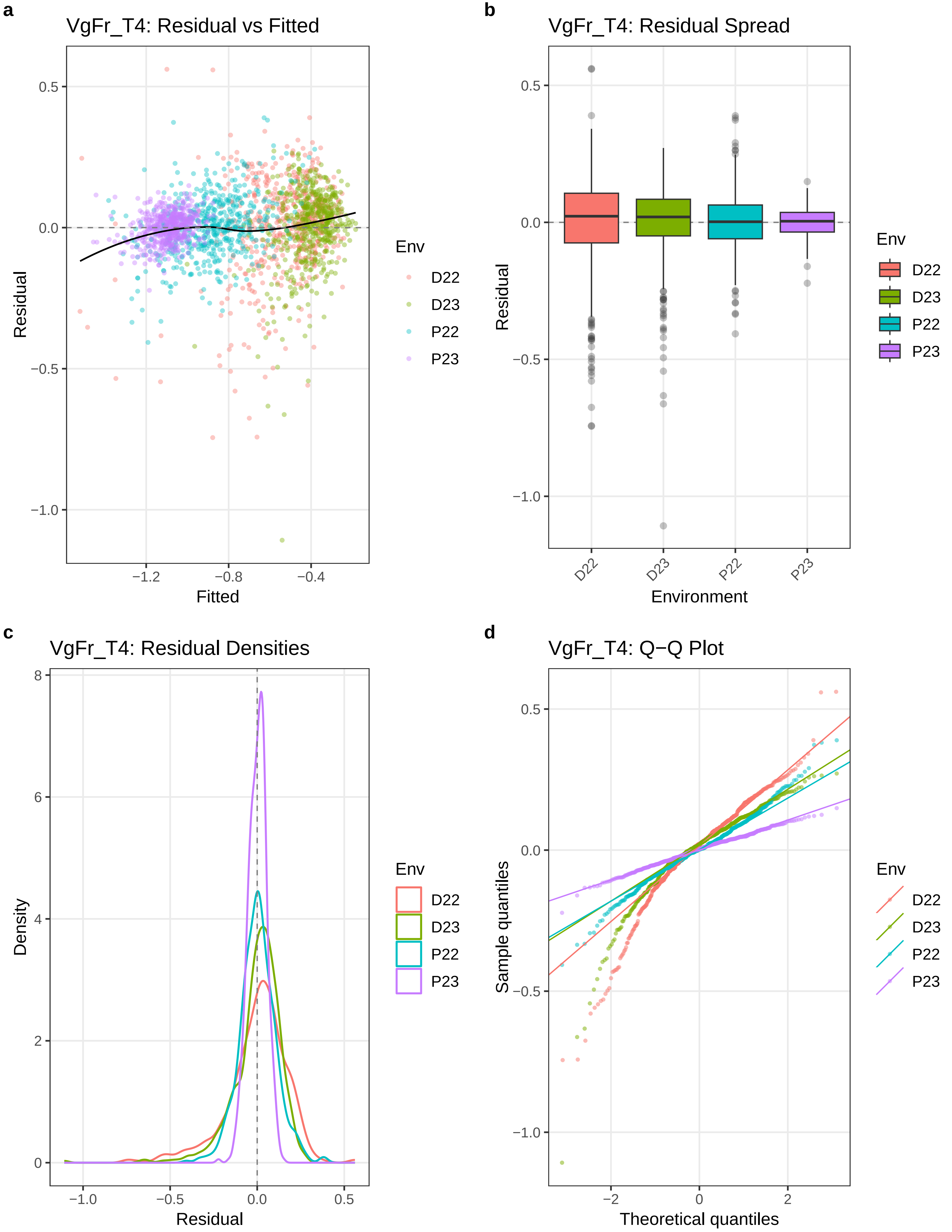

a
VgFr_T4: Residual vs Fitted
Residual
Fitted
0.5
0.0
−0.5
−1.0
−1.2
−0.8
−0.4
Env
D22
D23
P22
P23
b
VgFr_T4: Residual Spread
Residual
Environment
D22
D23
P22
P23
c
VgFr_T4: Residual Densities
Density
Residual
8
6
4
2
0
−1.0
−0.5
0.0
0.5
d
VgFr_T4: Q−Q Plot
Sample quantiles
Theoretical quantiles
−2
0
2

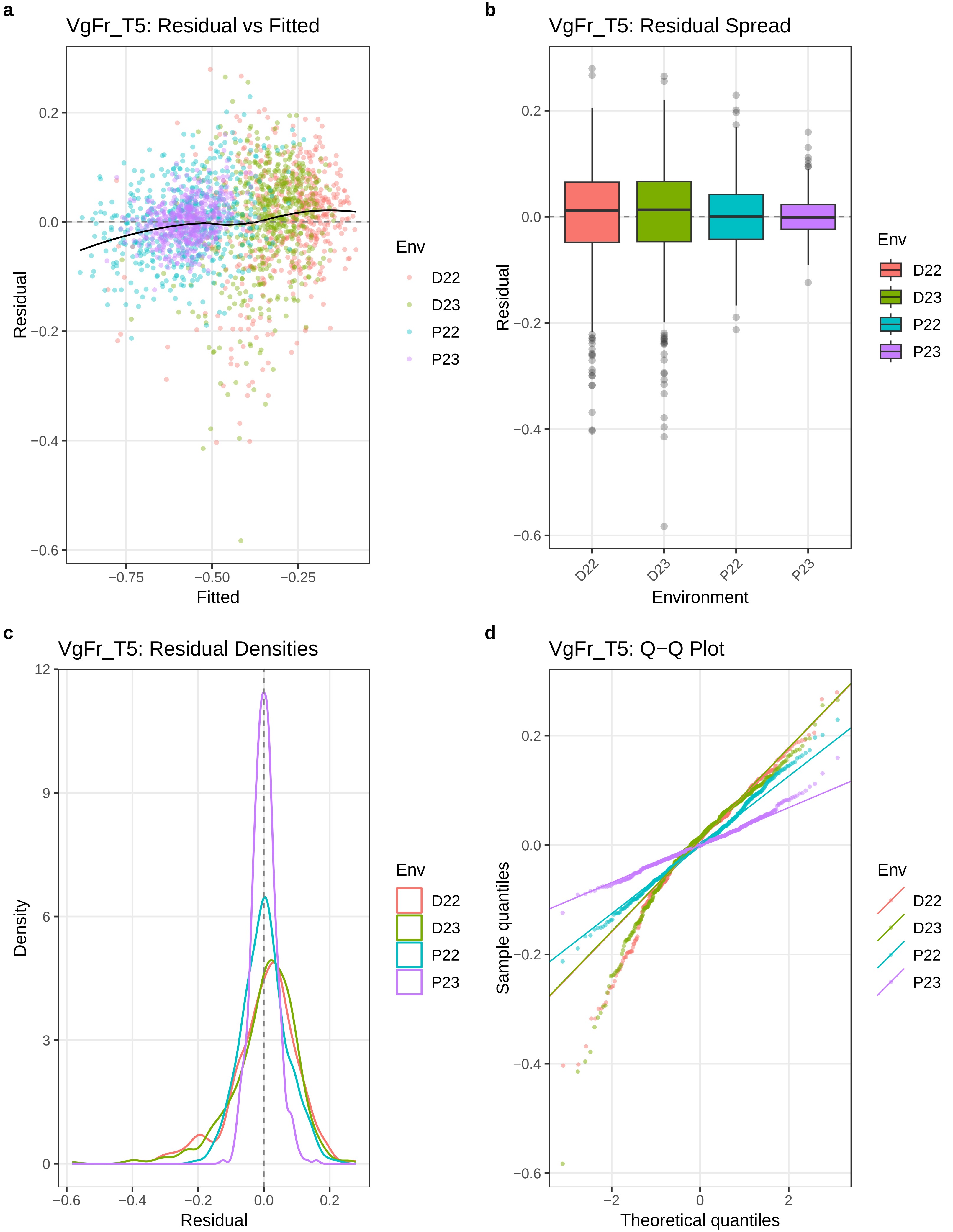
a
VgFr_T5: Residual vs Fitted
Residual
Fitted
0.2
0.0
−0.2
−0.4
−0.6
−0.75
−0.50
−0.25
Env
D22
D23
P22
P23
b
VgFr_T5: Residual Spread
Residual
Environment
D22
D23
P22
P23
c
VgFr_T5: Residual Densities
Density
Residual
12
9
6
3
0
−0.6
−0.4
−0.2
0.0
0.2
d
VgFr_T5: Q−Q Plot
Sample quantiles
Theoretical quantiles
−2
0
2

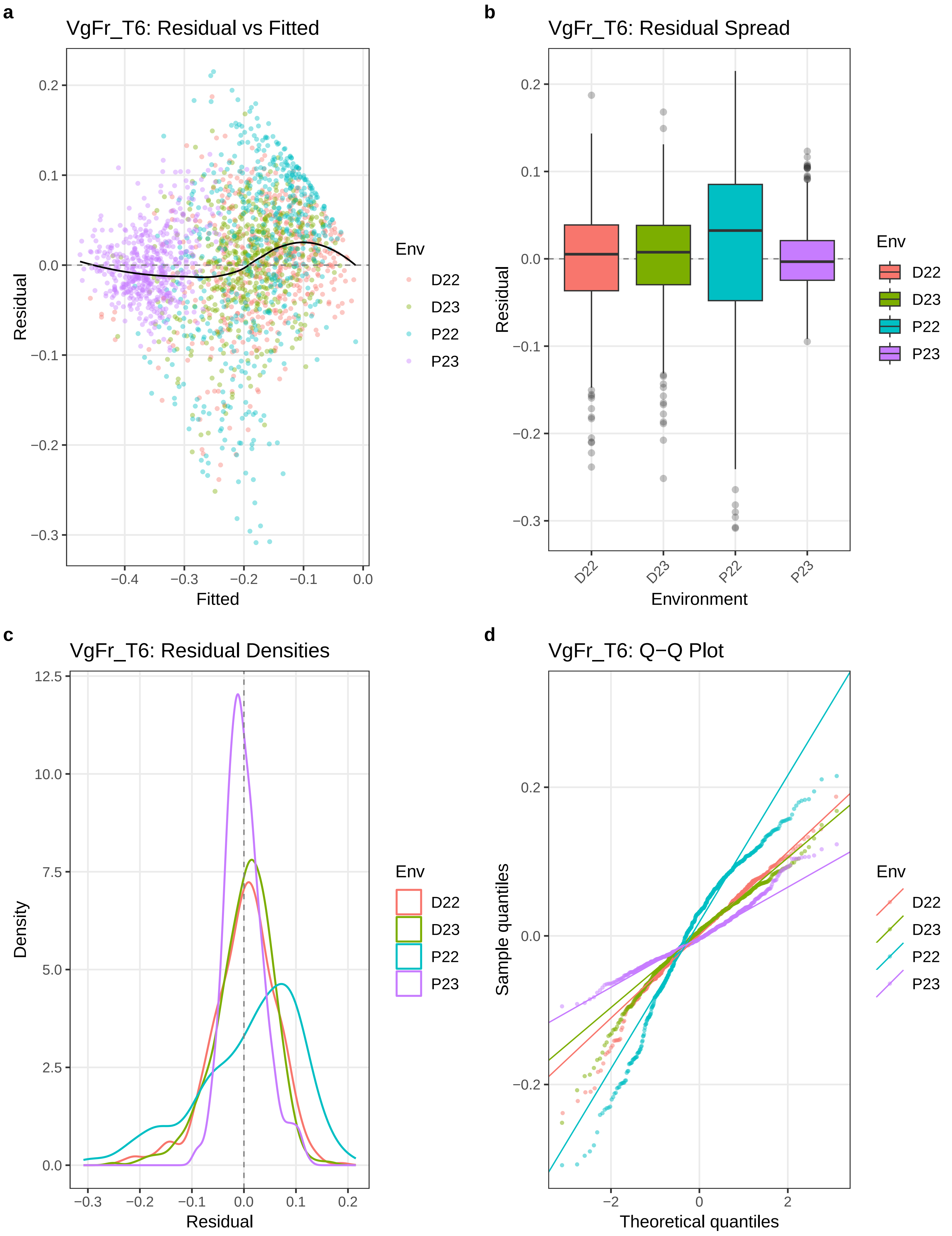
a
VgFr_T6: Residual vs Fitted
Residual
0.2
0.1
0.0
−0.1
−0.2
−0.3
Fitted
−0.4
−0.3
−0.2
−0.1
0.0
Env
D22
D23
P22
P23
b
VgFr_T6: Residual Spread
Residual
0.2
0.1
0.0
−0.1
−0.2
−0.3
Environment
D22
D23
P22
P23
Env
D22
D23
P22
P23
c
VgFr_T6: Residual Densities
Density
12.5
10.0
7.5
5.0
2.5
0.0
Residual
−0.3
−0.2
−0.1
0.0
0.1
0.2
Env
D22
D23
P22
P23
d
VgFr_T6: Q−Q Plot
Sample quantiles
0.2
0.0
−0.2
Theoretical quantiles
−2
0
2
Env
D22
D23
P22
P23

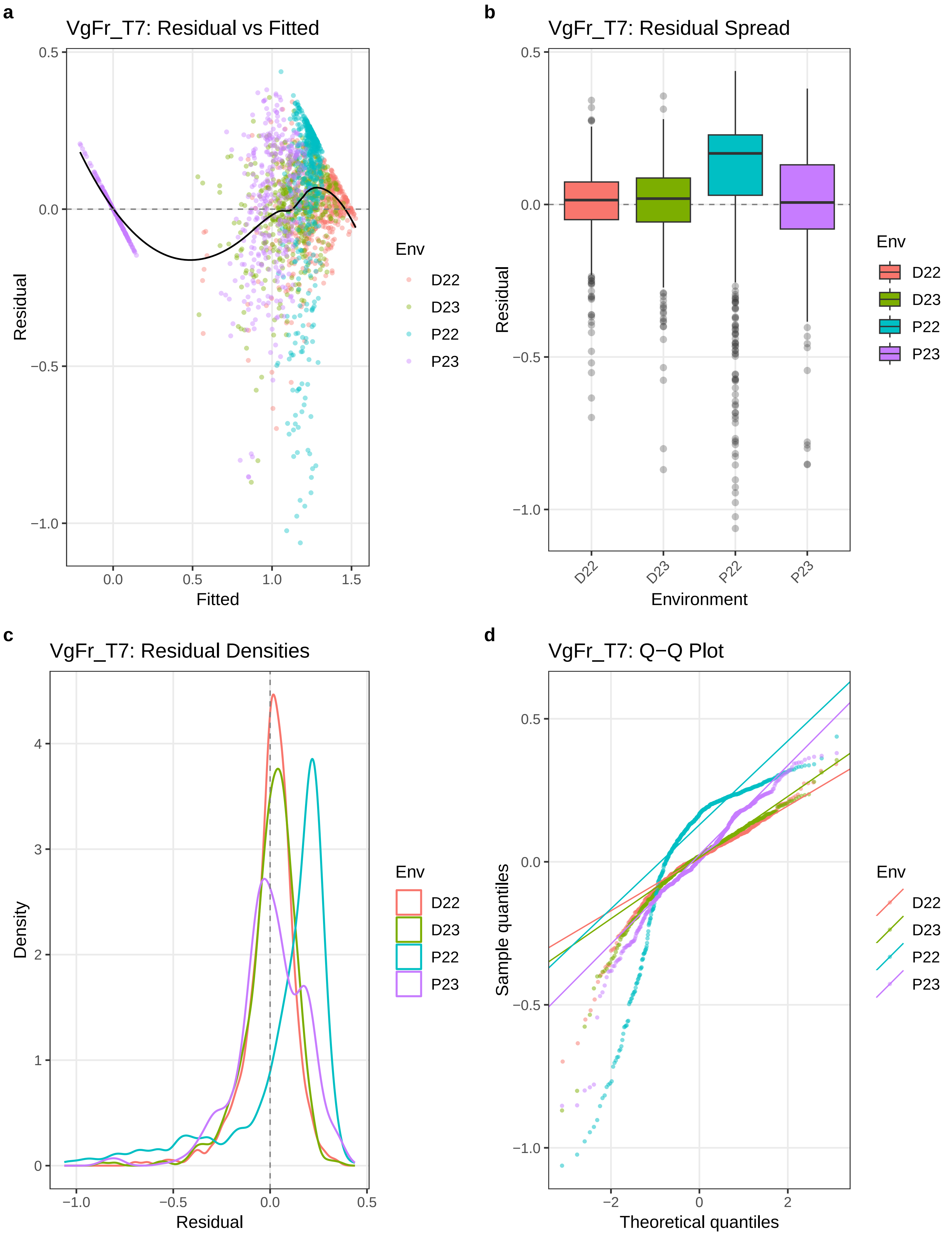
a
VgFr_T7: Residual vs Fitted
Residual
Fitted
Env
D22
D23
P22
P23
b
VgFr_T7: Residual Spread
Residual
Environment
c
VgFr_T7: Residual Densities
Density
Residual
d
VgFr_T7: Q−Q Plot
Sample quantiles
Theoretical quantiles

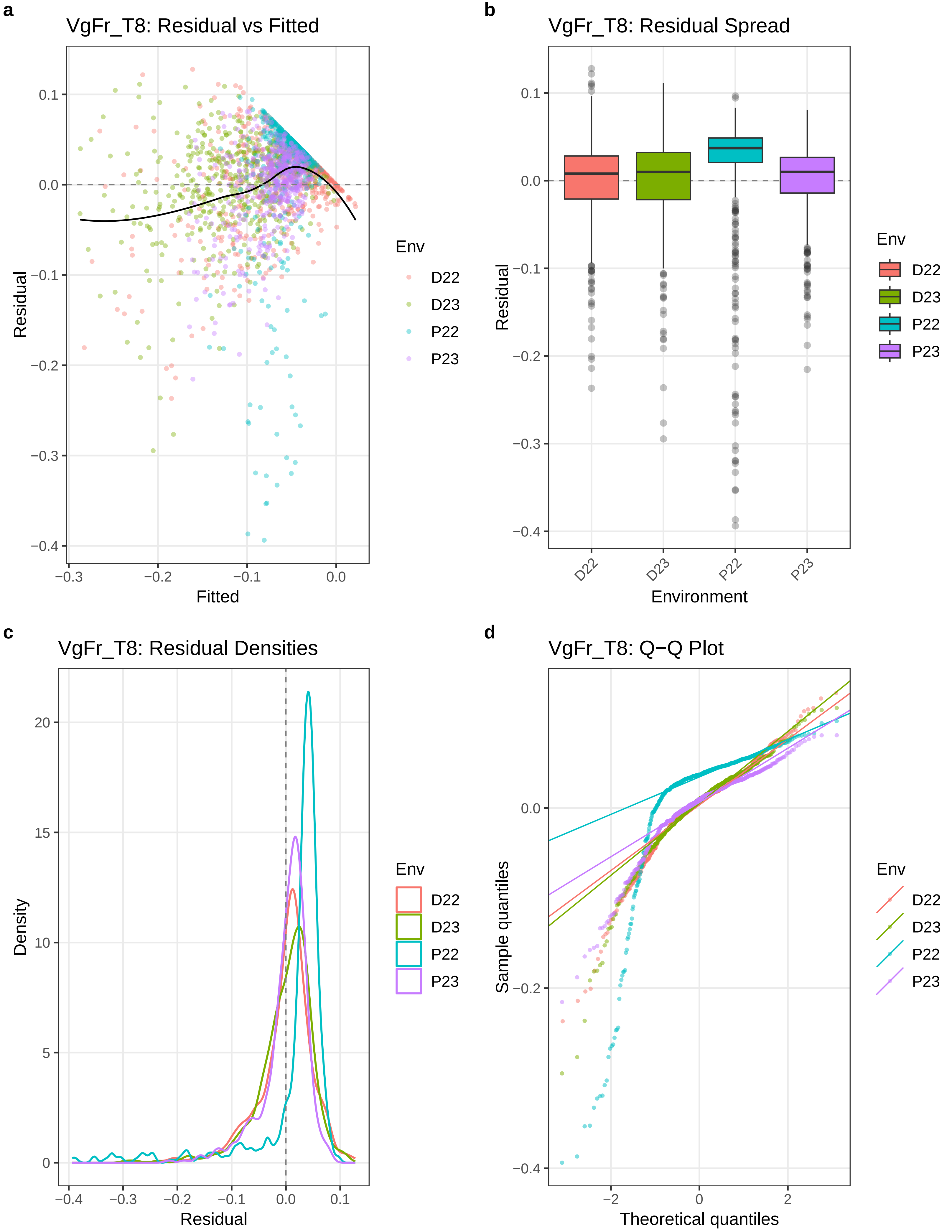
a
VgFr_T8: Residual vs Fitted
Residual
Fitted
0.1
0.0
−0.1
−0.2
−0.3
−0.4
−0.3
−0.2
−0.1
0.0
Env
D22
D23
P22
P23
b
VgFr_T8: Residual Spread
Residual
Environment
0.1
0.0
−0.1
−0.2
−0.3
−0.4
D22
D23
P22
P23
Env
D22
D23
P22
P23
c
VgFr_T8: Residual Densities
Density
Residual
20
15
10
5
0
−0.4
−0.3
−0.2
−0.1
0.0
0.1
Env
D22
D23
P22
P23
d
VgFr_T8: Q−Q Plot
Sample quantiles
Theoretical quantiles
0.0
−0.2
−0.4
−2
0
2
Env
D22
D23
P22
P23

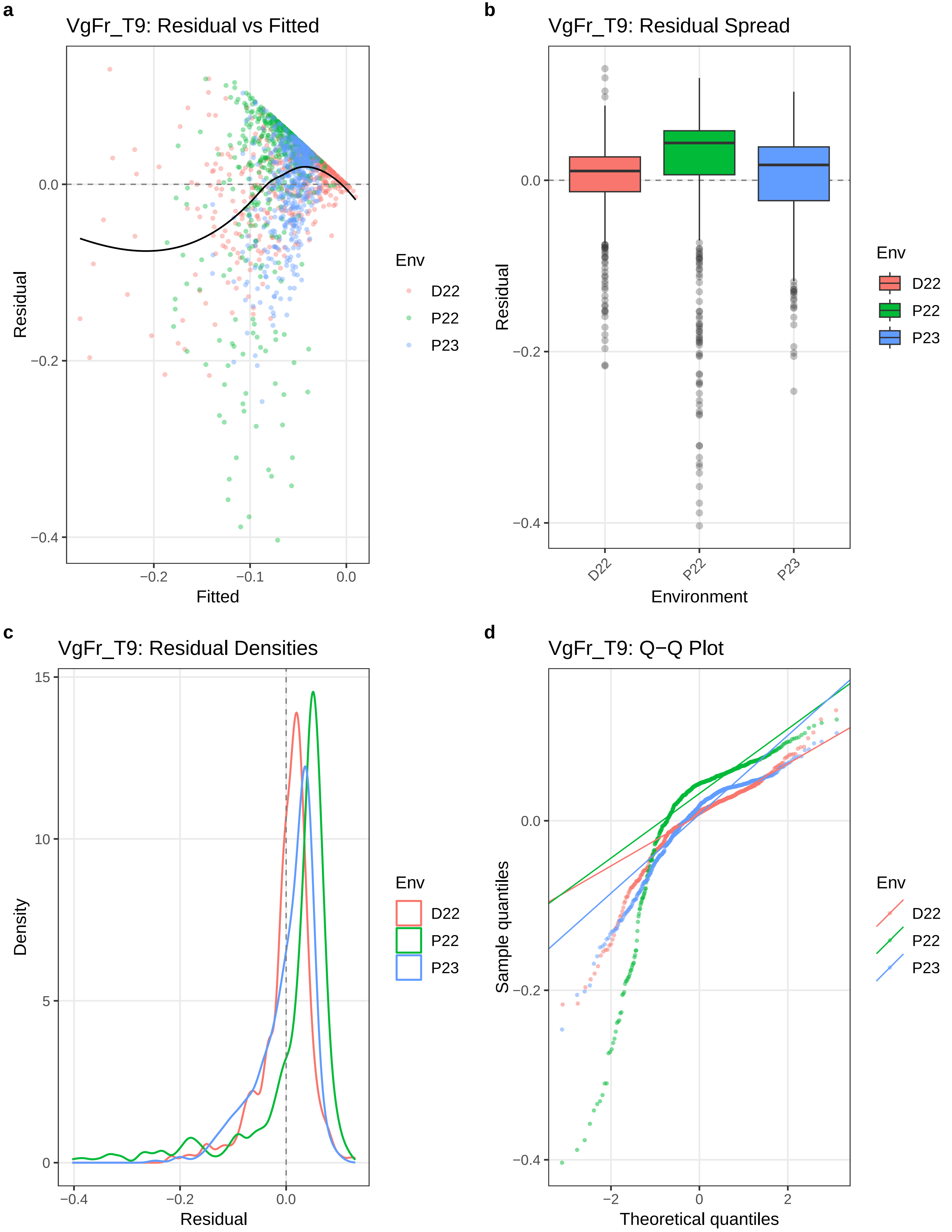
a
VgFr_T9: Residual vs Fitted
Residual
Fitted
0.0
−0.2
−0.4
−0.2
−0.1
0.0
Env
D22
P22
P23
b
VgFr_T9: Residual Spread
Residual
Environment
0.0
−0.2
−0.4
D22
P22
P23
Env
D22
P22
P23
c
VgFr_T9: Residual Densities
Density
Residual
15
10
5
0
−0.4
−0.2
0.0
Env
D22
P22
P23
d
VgFr_T9: Q−Q Plot
Sample quantiles
Theoretical quantiles
0.0
−0.2
−0.4
−2
0
2
Env
D22
P22
P23
