## Supplementary material for "Genetic mapping and genomic prediction for agronomic, grain compositional, and sensing-enabled traits in a cowpea MAGIC population along an environmental gradient": Large supplemental tables and figures, uploaded separately: SuppFig_19_QTLprofiles_wPseudo_HK_LMM_LOCO.pdf

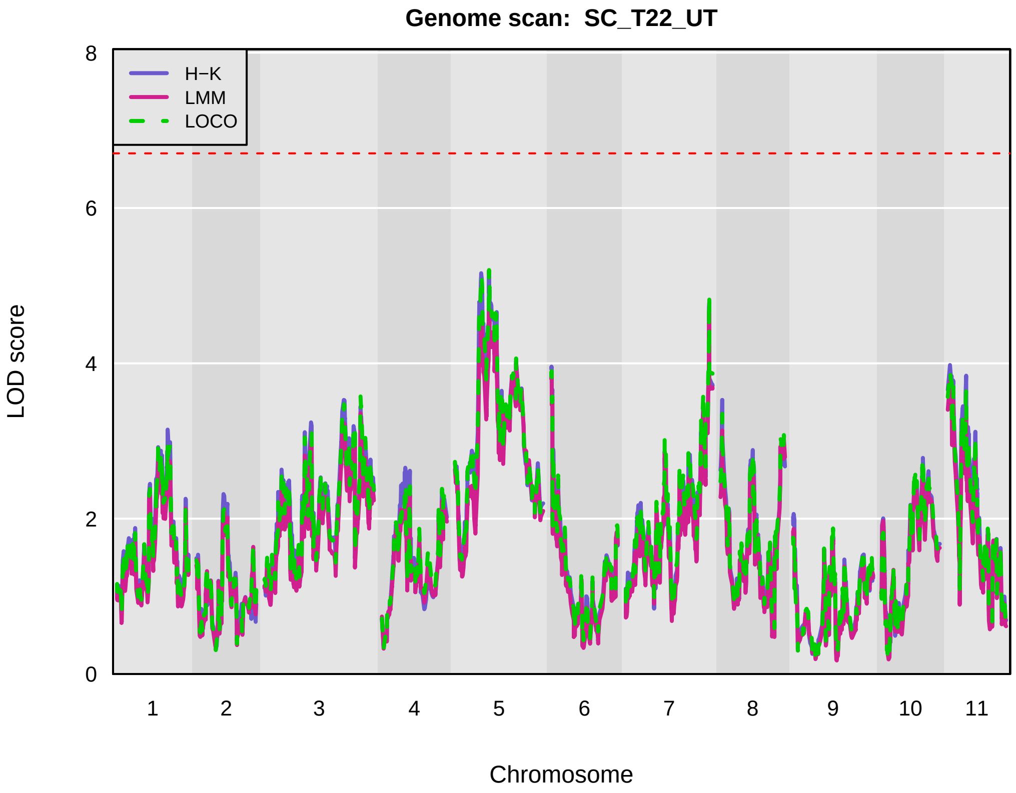
Genome scan: SC_T22_UT
H−K
LMM
LOCO
LOD score
0
2
4
6
8
Chromosome
1 2 3 4 5 6 7 8 9 10 11

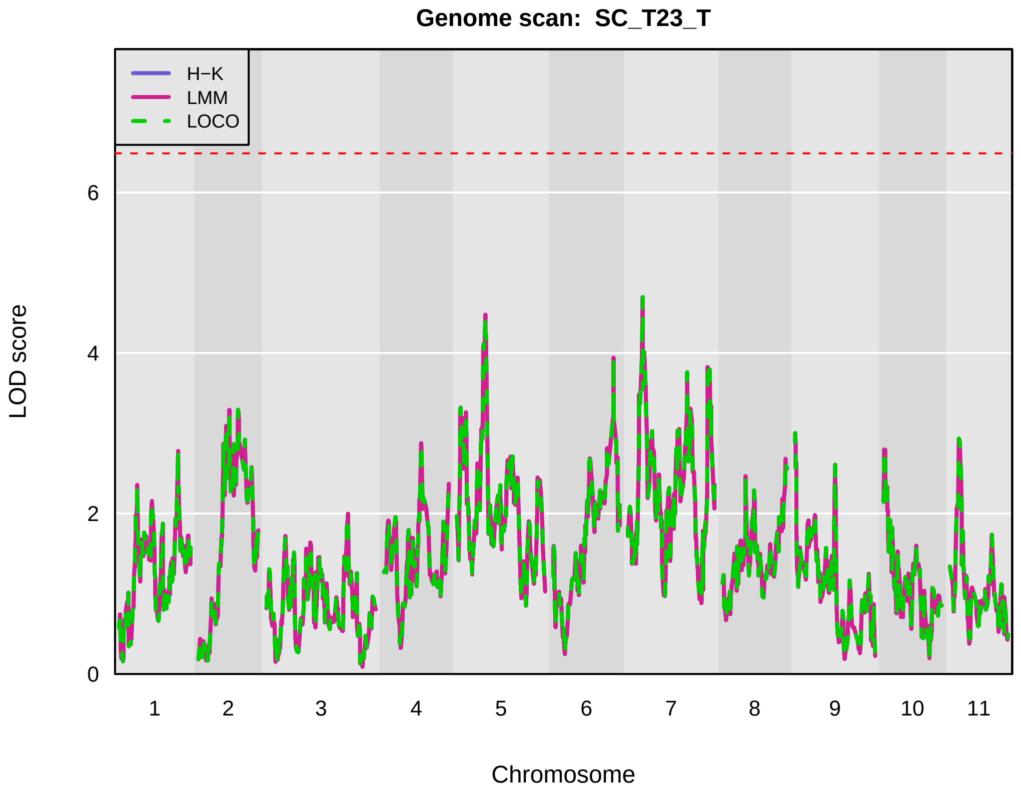
Genome scan: SC_T23_T
H−K
LMM
LOCO
LOD score
0
2
4
6
Chromosome
1 2 3 4 5 6 7 8 9 10 11

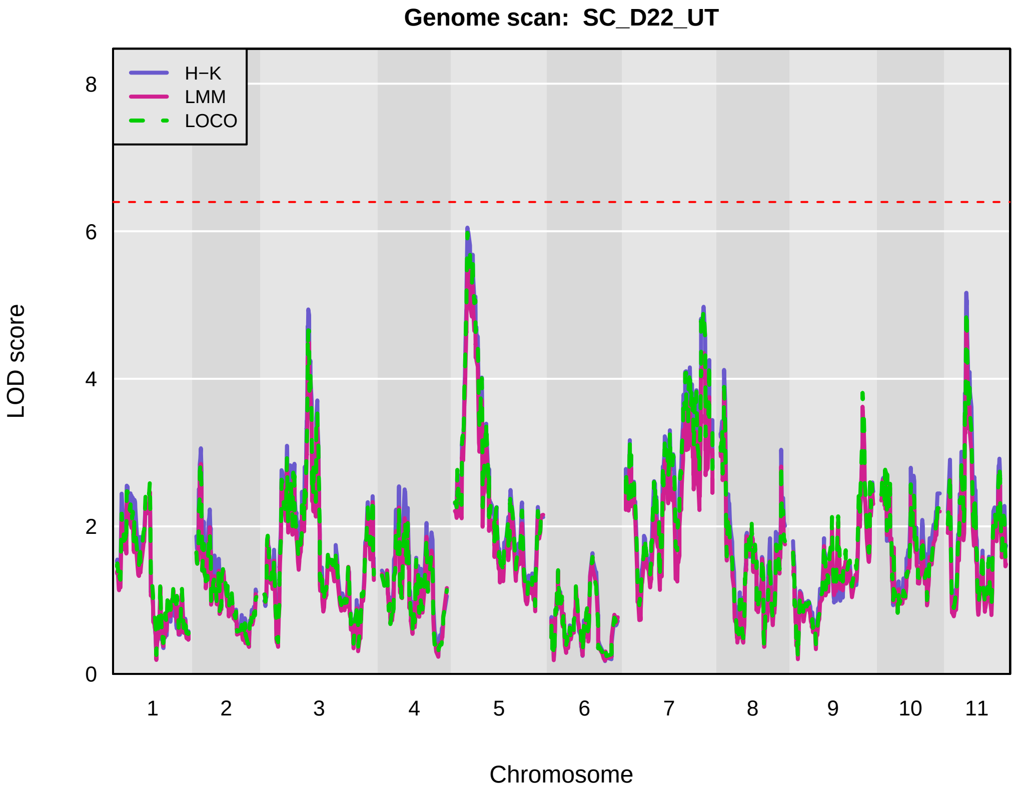
Genome scan: SC_D22_UT
H−K
LMM
LOCO
LOD score
0
2
4
6
8
Chromosome
1 2 3 4 5 6 7 8 9 10 11

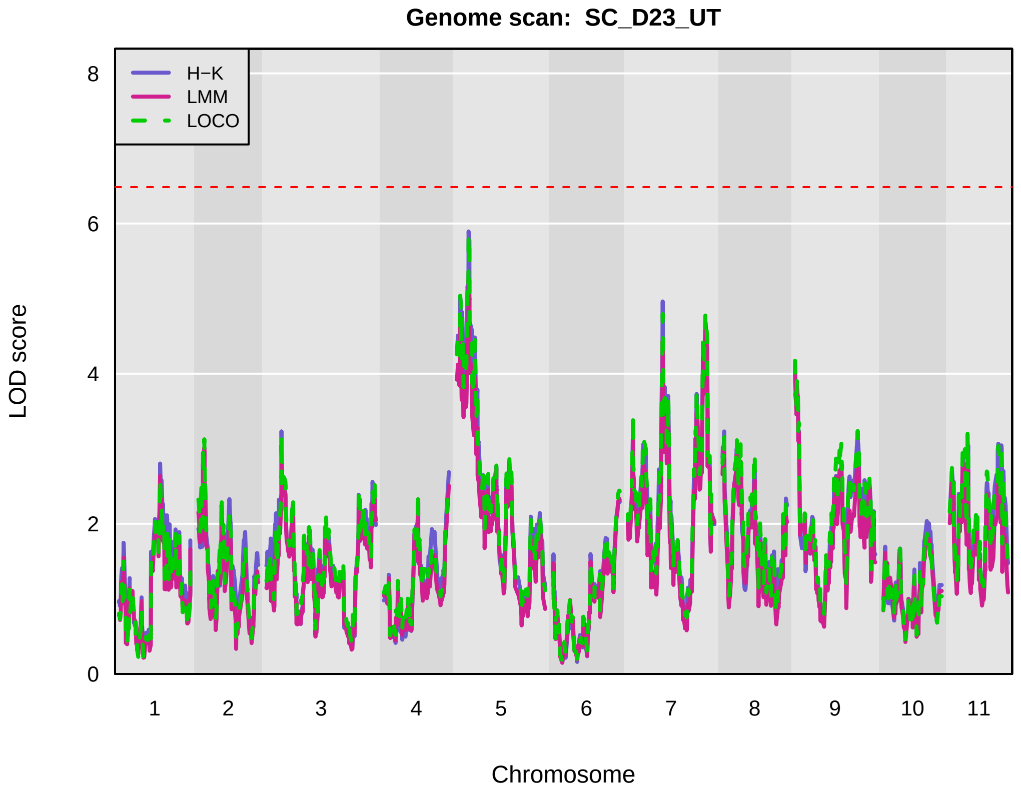
Genome scan: SC_D23_UT
H−K
LMM
LOCO
LOD score
0
2
4
6
8
Chromosome
1 2 3 4 5 6 7 8 9 10 11

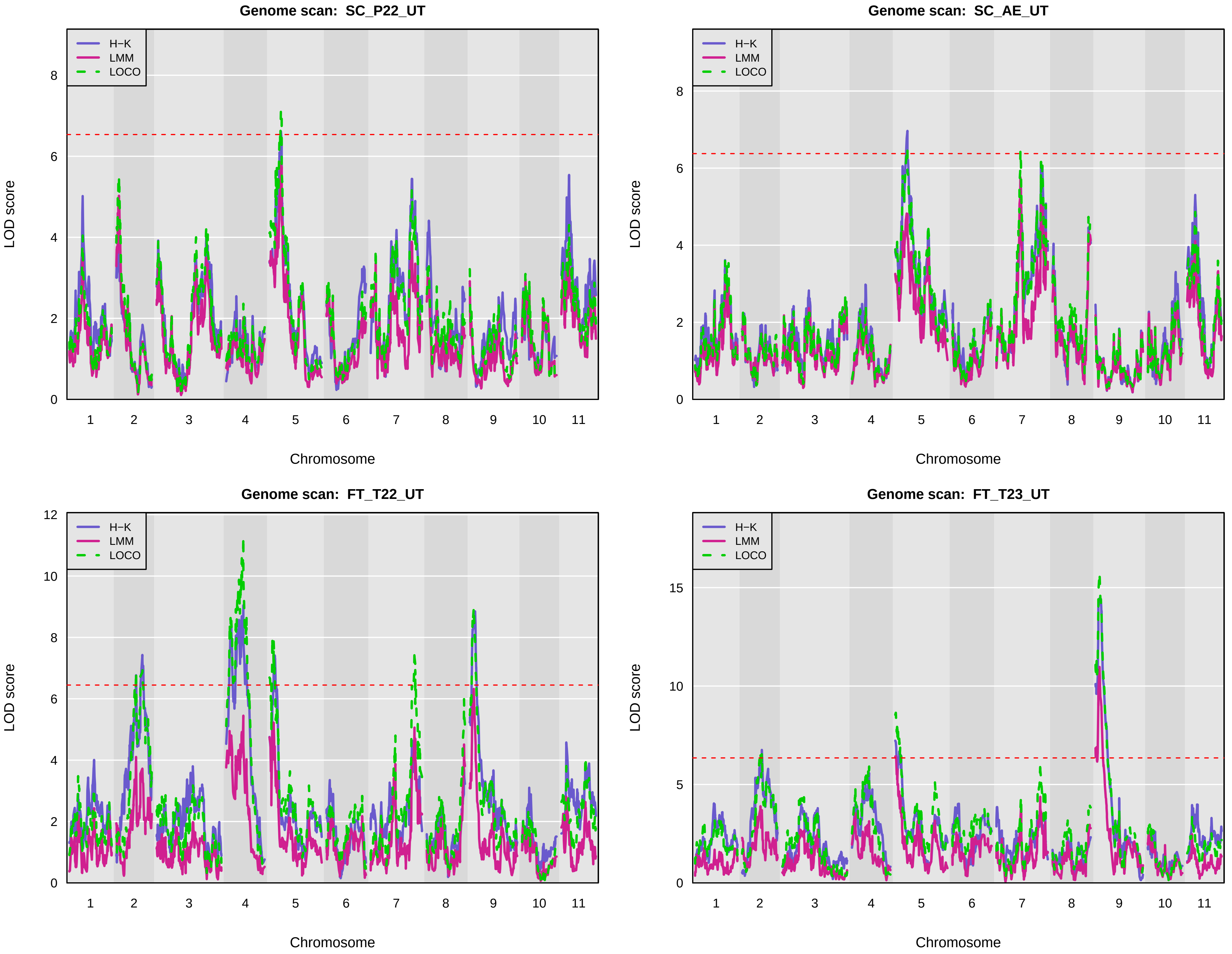
Genome scan: SC_P22_UT
Genome scan: SC_AE_UT
Genome scan: FT_T22_UT
Genome scan: FT_T23_UT
H−K
LMM
LOCO
LOD score
Chromosome
1
2
3
4
5
6
7
8
9
10
11
0
2
4
6
8
10
12
15

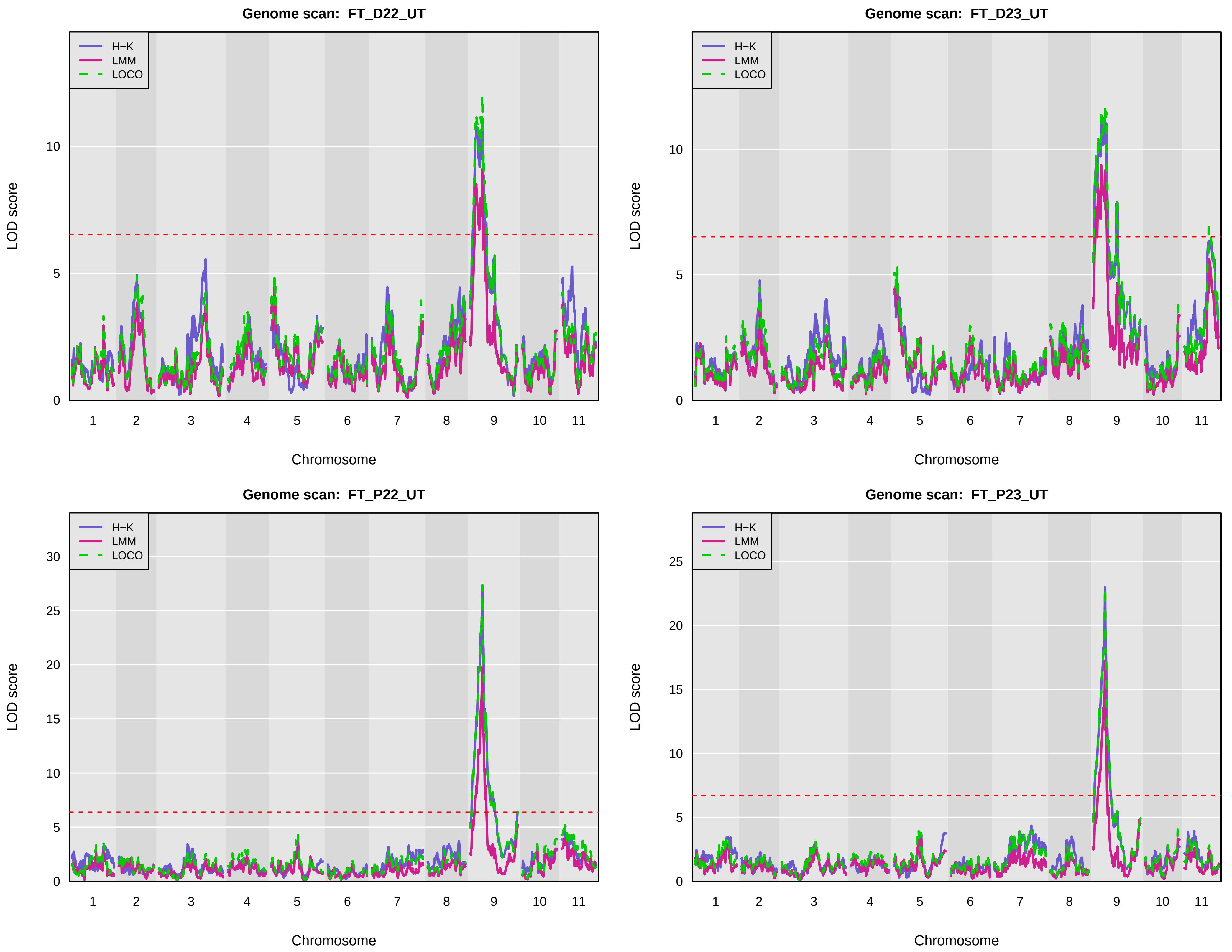
Genome scan: FT_D22_UT
Genome scan: FT_D23_UT
Genome scan: FT_P22_UT
Genome scan: FT_P23_UT
H−K
LMM
LOCO
LOD score
Chromosome
1 2 3 4 5 6 7 8 9 10 11
0 5 10
0 5 10 15 20 25 30
0 5 10 15 20 25

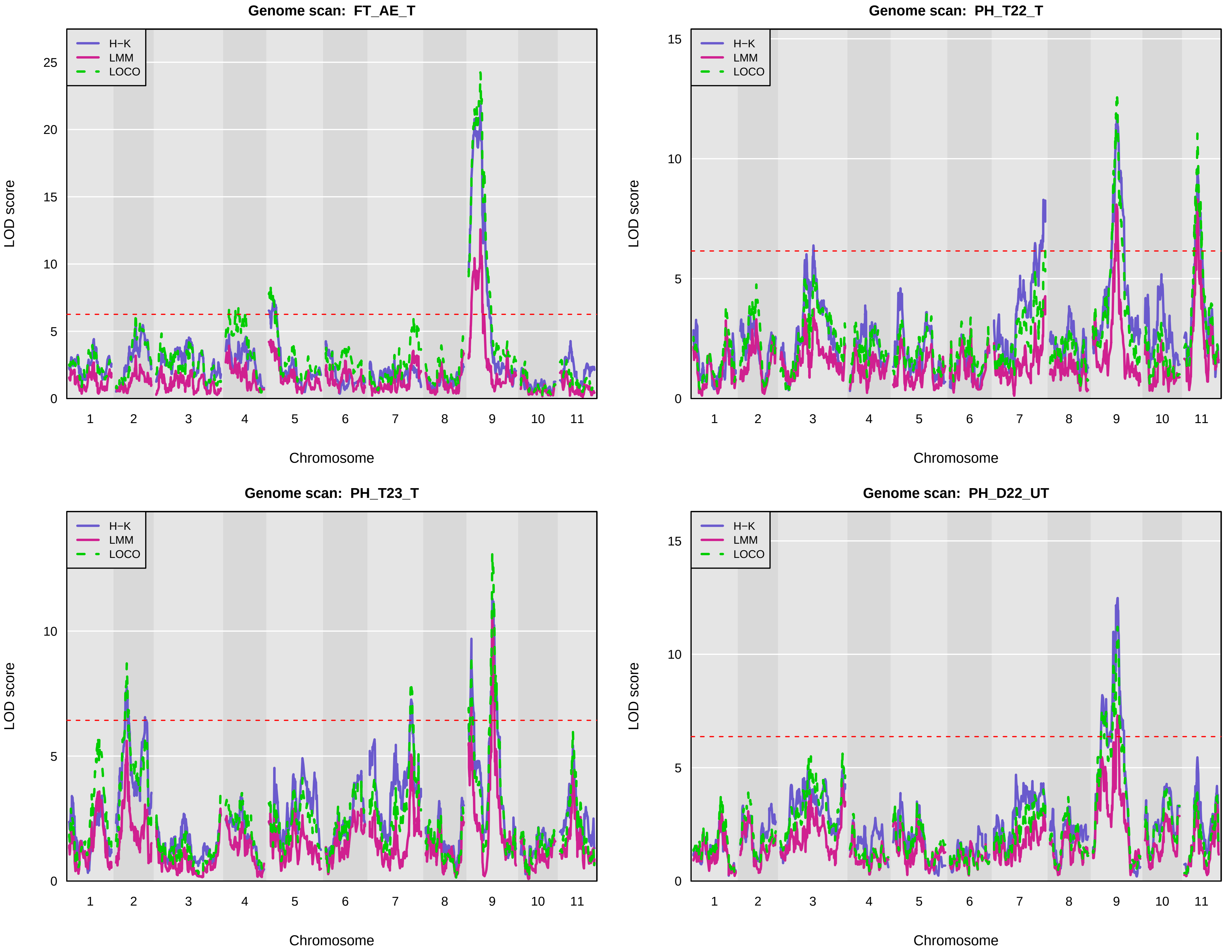
Genome scan: FT_AE_T
Genome scan: PH_T22_T
Genome scan: PH_T23_T
Genome scan: PH_D22_UT
H−K
LMM
LOCO
LOD score
Chromosome
1 2 3 4 5 6 7 8 9 10 11

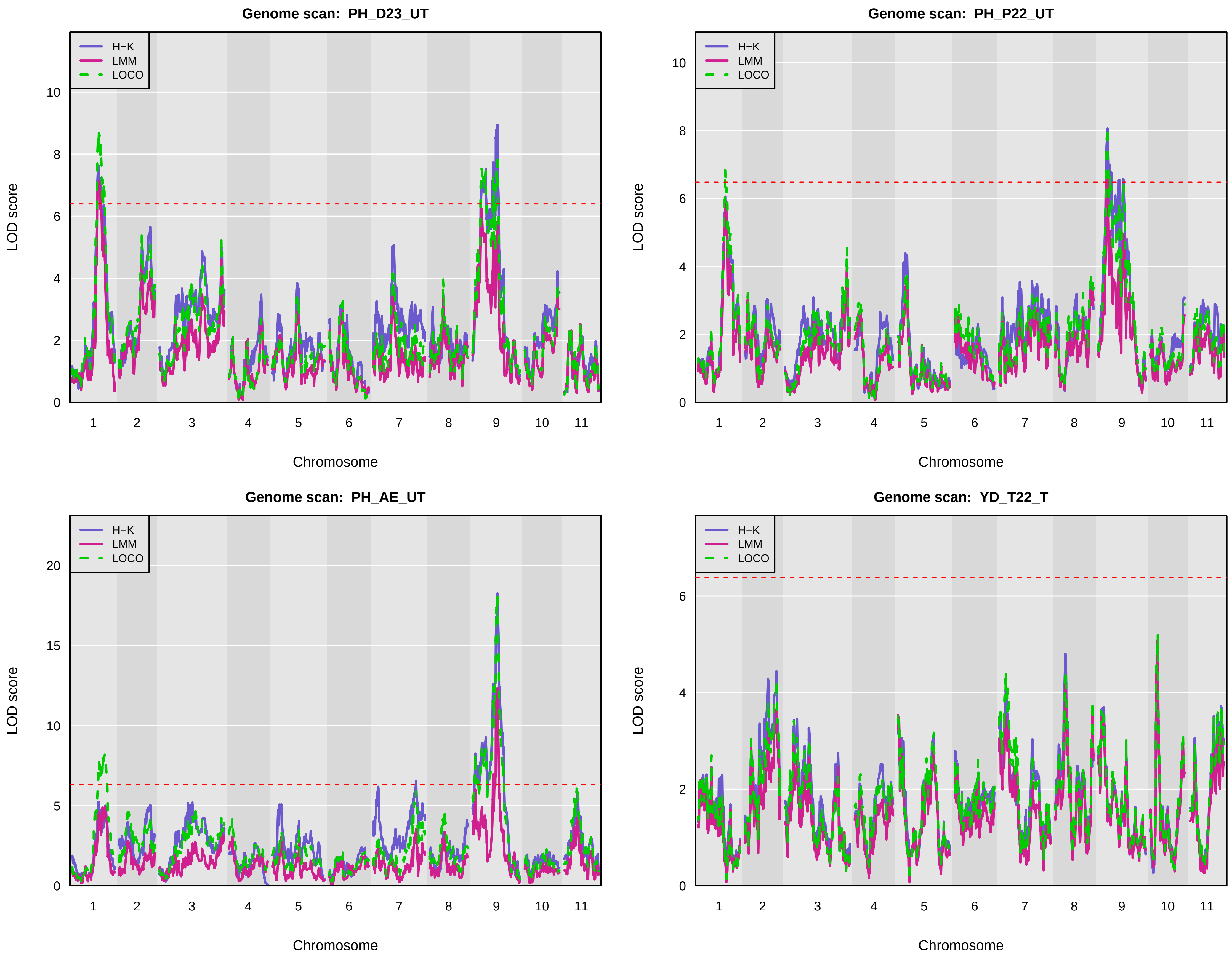
Genome scan: PH_D23_UT
Genome scan: PH_P22_UT
Genome scan: PH_AE_UT
Genome scan: YD_T22_T
H−K
LMM
LOCO
LOD score
Chromosome
1
2
3
4
5
6
7
8
9
10
11

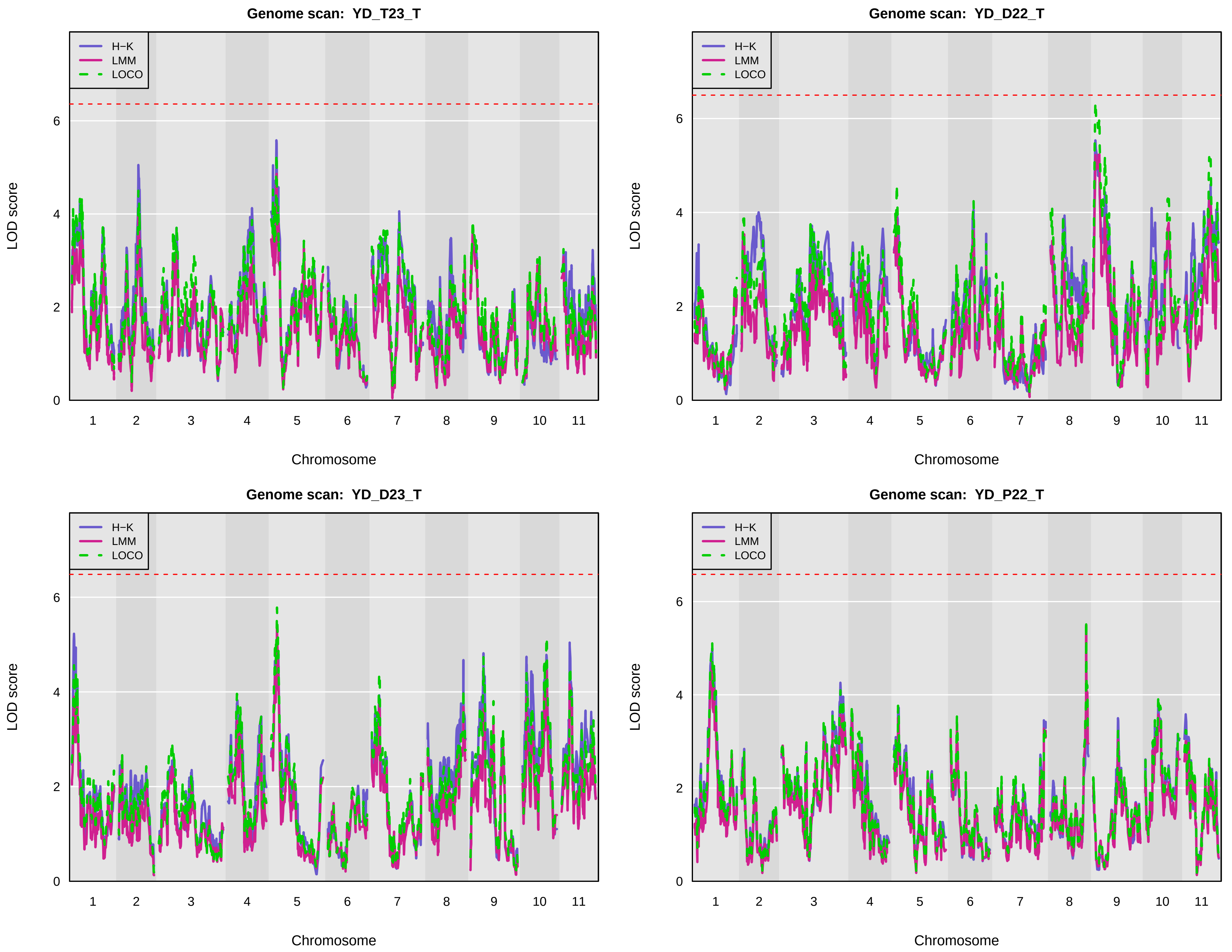

Genome scan: YD_T23_T
Genome scan: YD_D22_T
Genome scan: YD_D23_T
Genome scan: YD_P22_T
H−K
LMM
LOCO
LOD score
Chromosome
0
2
4
6
1
2
3
4
5
6
7
8
9
10
11

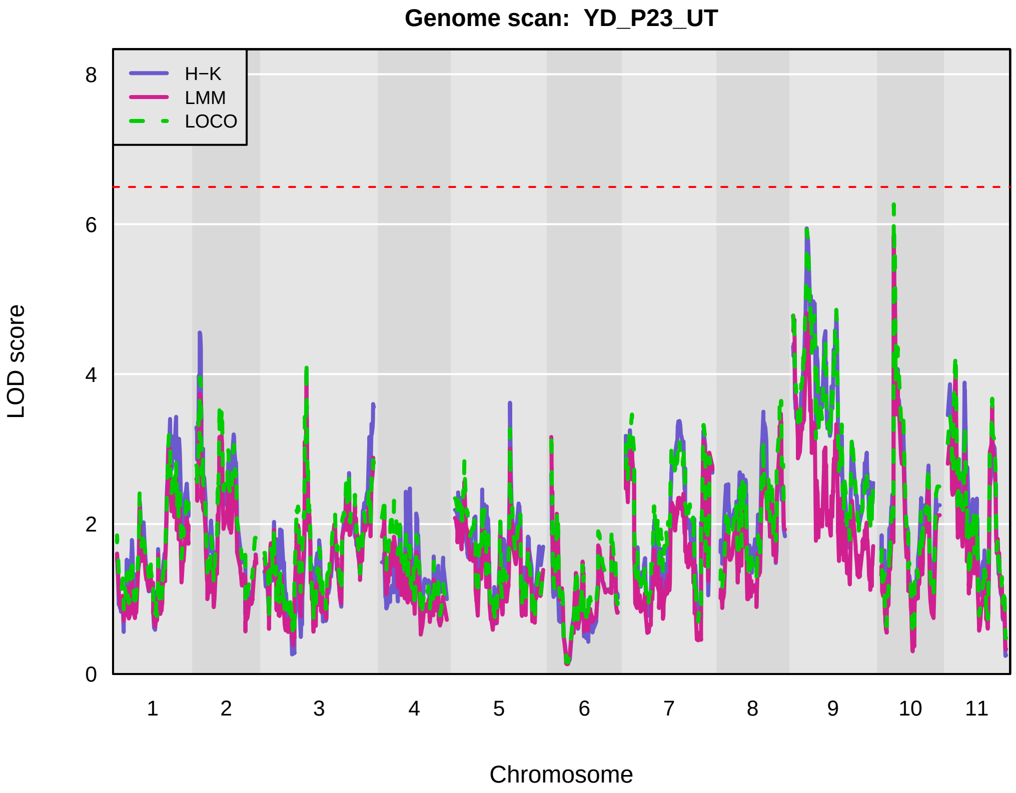
Genome scan: YD_P23_UT
H−K
LMM
LOCO
LOD score
0
2
4
6
8
1 2 3 4 5 6 7 8 9 10 11
Chromosome

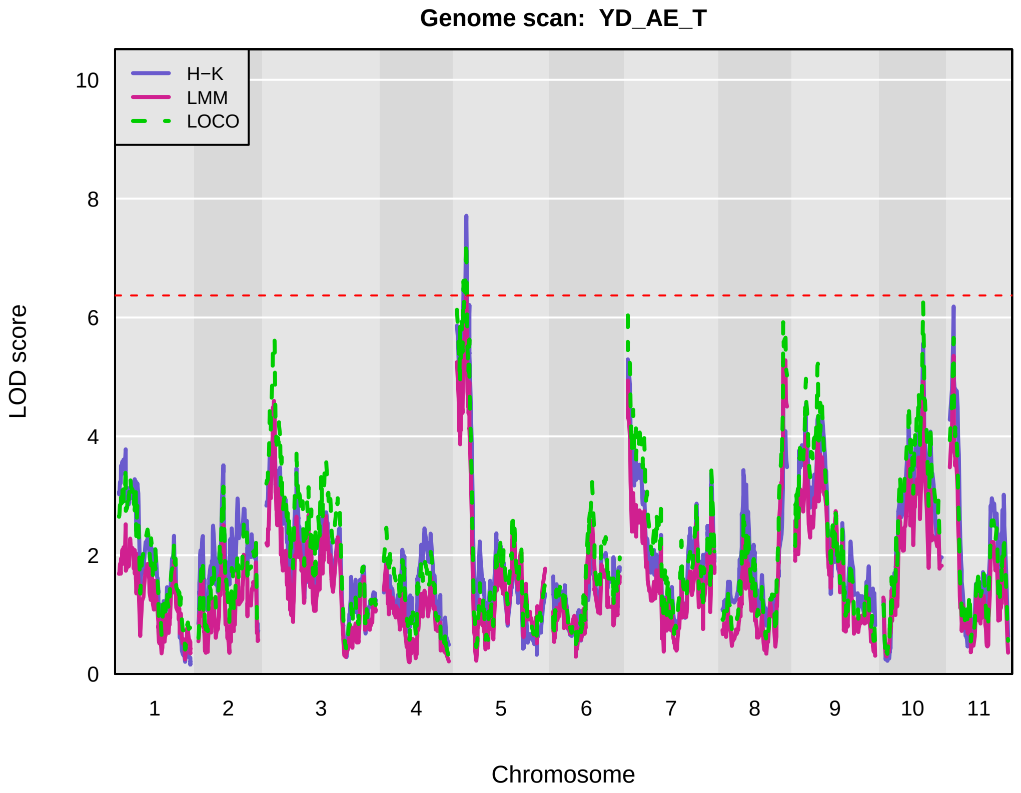
Genome scan: YD_AE_T
H−K
LMM
LOCO
LOD score
0
2
4
6
8
10
1 2 3 4 5 6 7 8 9 10 11
Chromosome

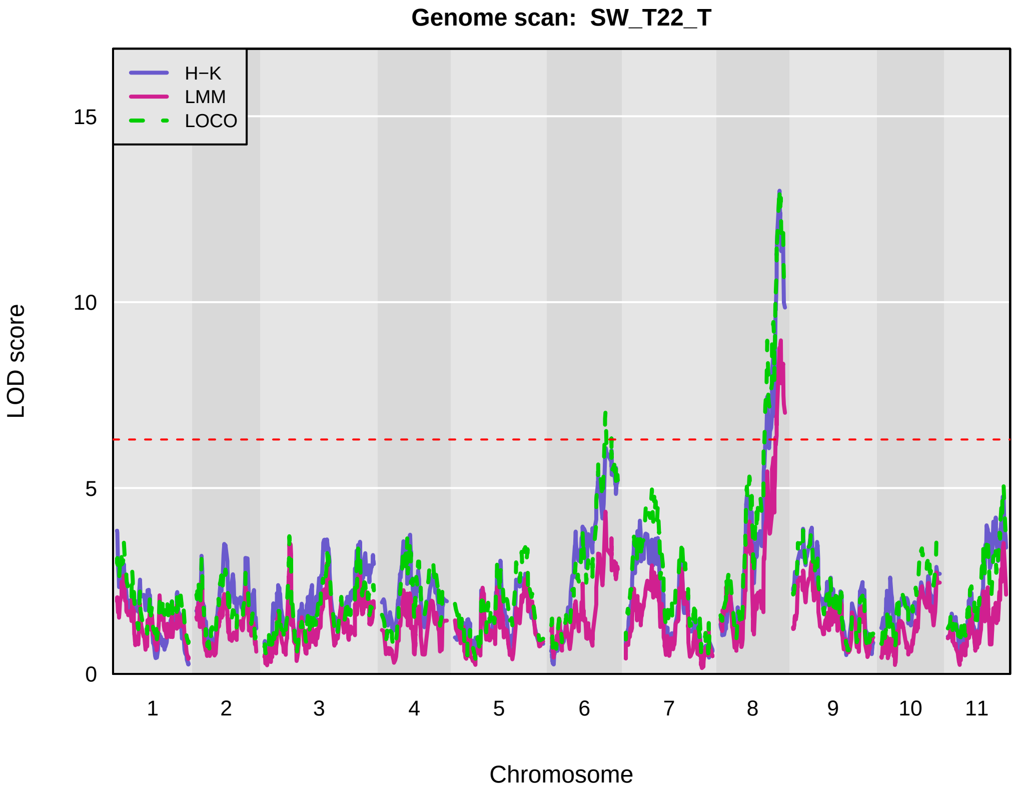
Genome scan: SW_T22_T
H−K
LMM
LOCO
LOD score
0
5
10
15
1 2 3 4 5 6 7 8 9 10 11
Chromosome

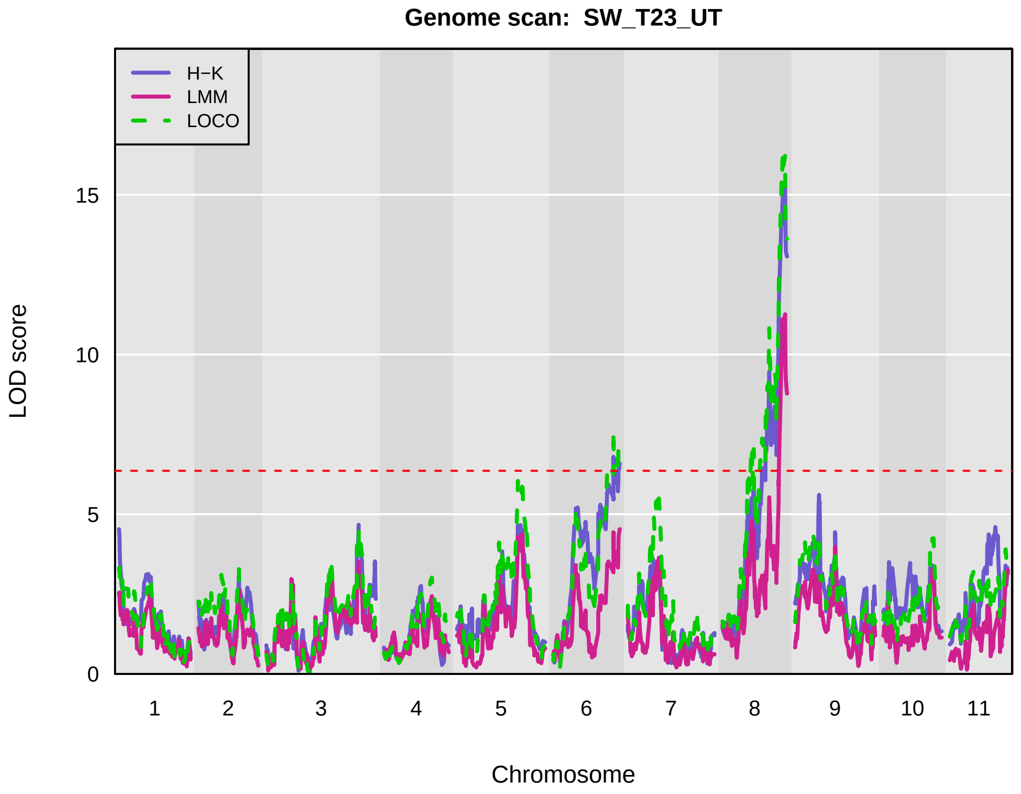
Genome scan: SW_T23_UT
H−K
LMM
LOCO
LOD score
0
5
10
15
1 2 3 4 5 6 7 8 9 10 11
Chromosome

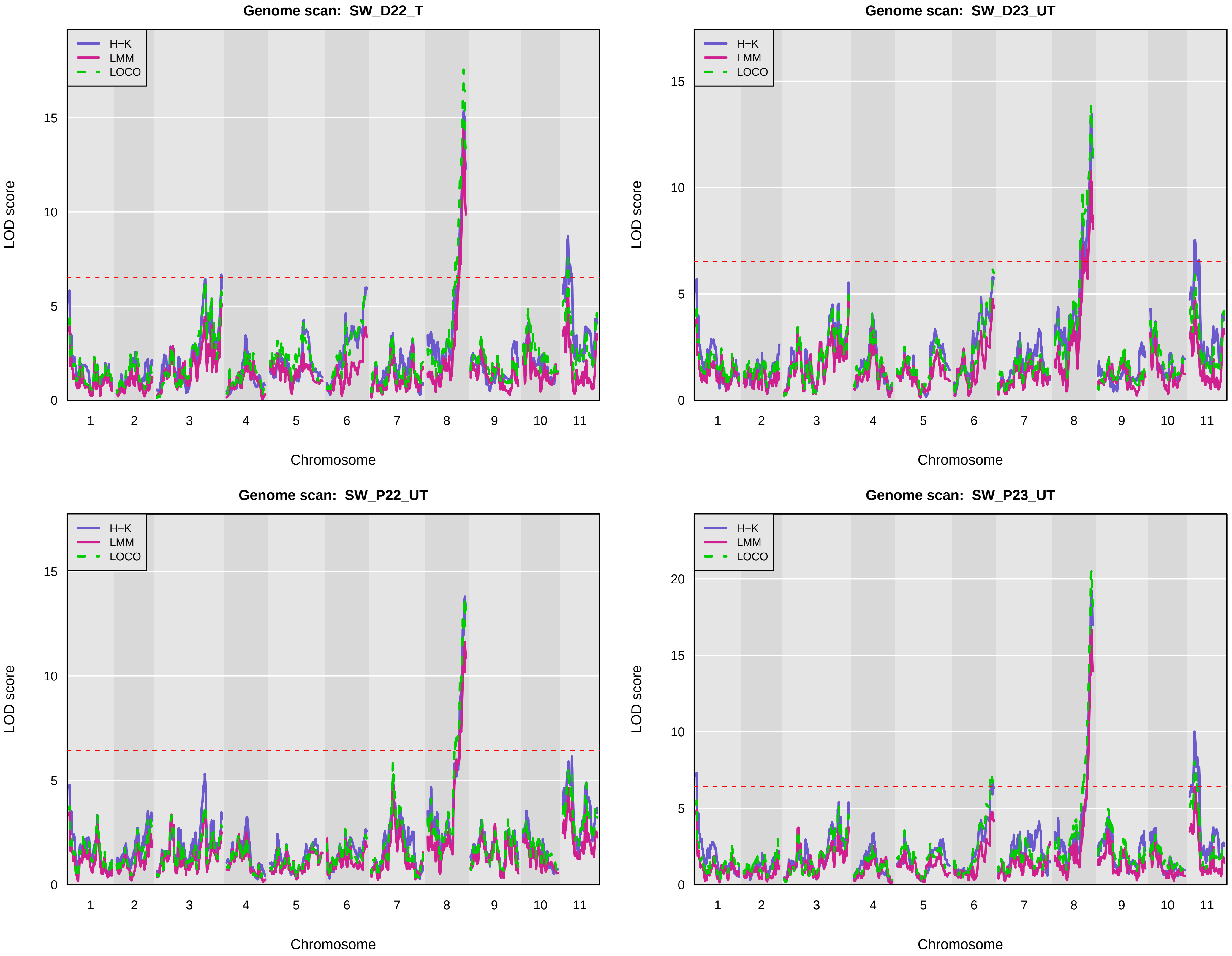
Genome scan: SW_D22_T
Genome scan: SW_D23_UT
Genome scan: SW_P22_UT
Genome scan: SW_P23_UT
H−K
LMM
LOCO
LOD score
Chromosome
1
2
3
4
5
6
7
8
9
10
11
0
5
10
15
20

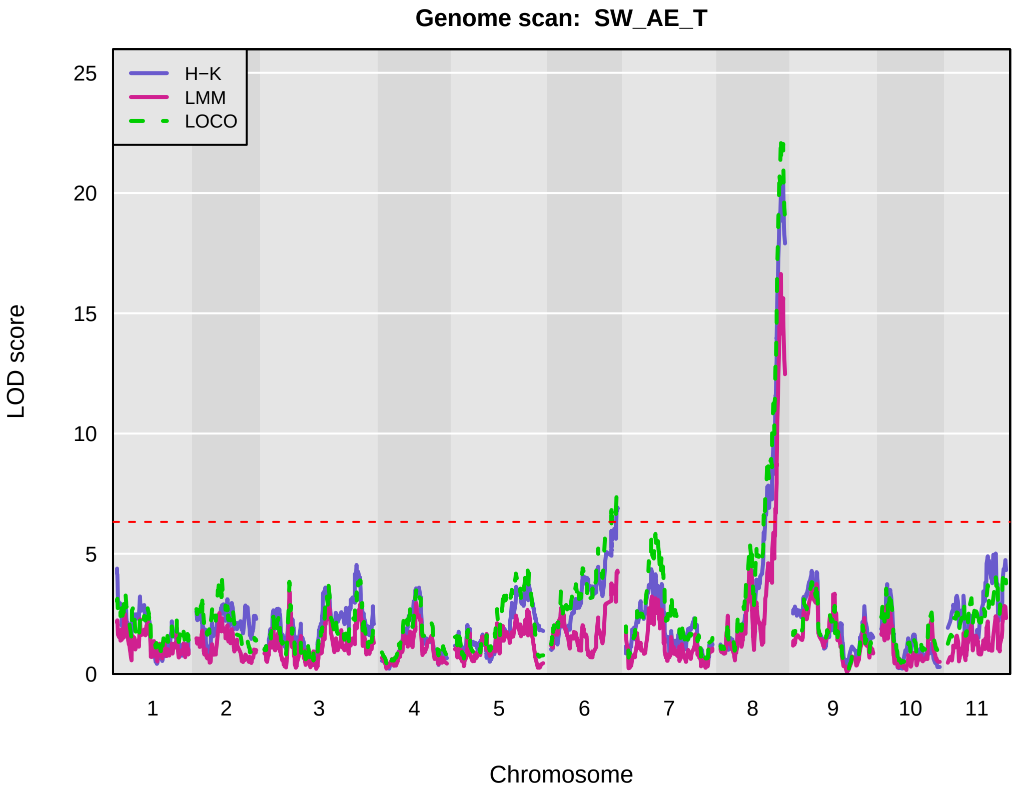
Genome scan: SW_AE_T
H−K
LMM
LOCO
LOD score
0
5
10
15
20
25
1 2 3 4 5 6 7 8 9 10 11
Chromosome

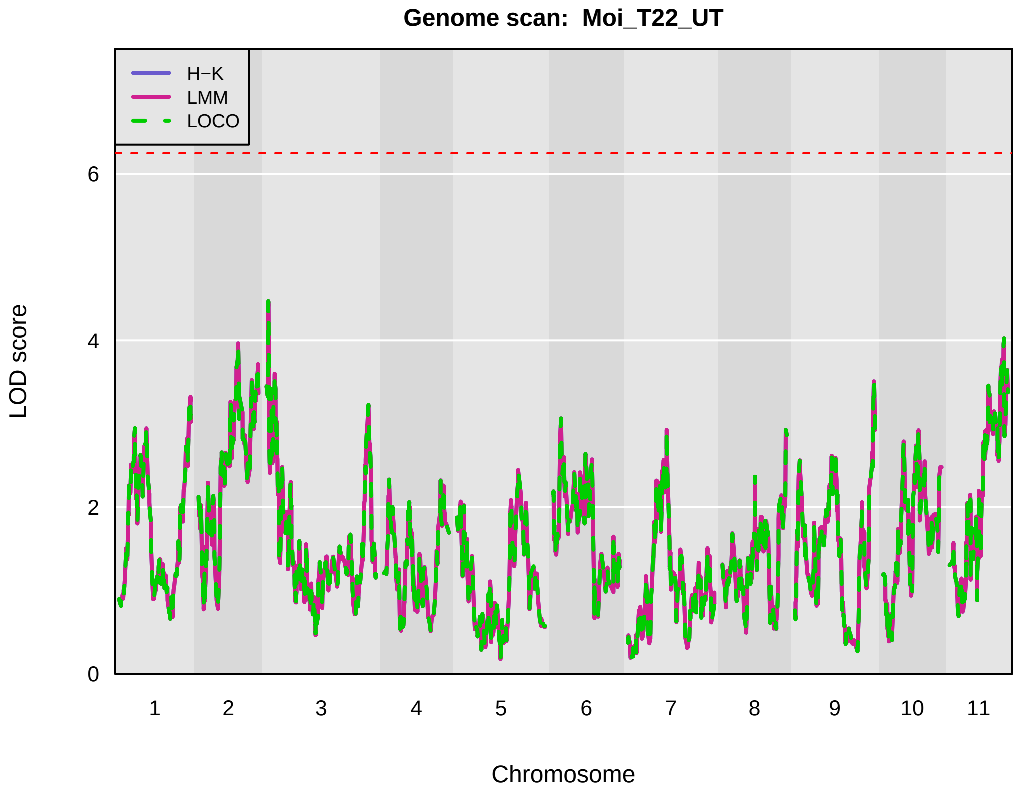
Genome scan: Moi_T22_UT
H−K
LMM
LOCO
LOD score
0
2
4
6
1 2 3 4 5 6 7 8 9 10 11
Chromosome

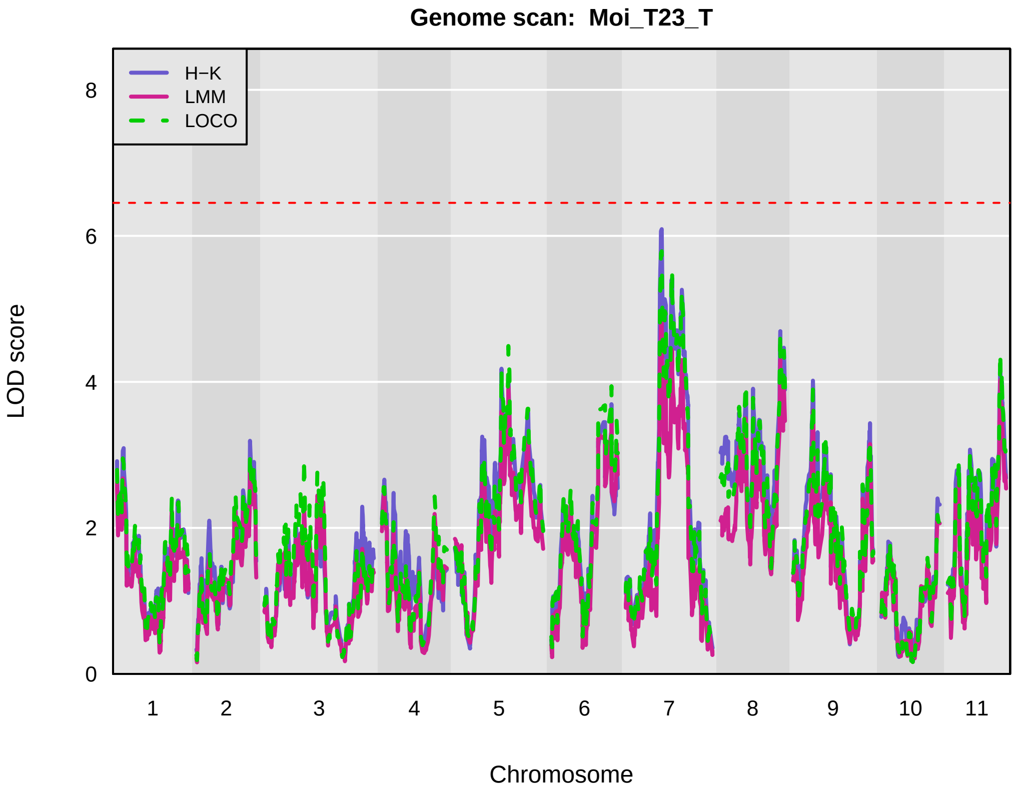
Genome scan: Moi_T23_T
H−K
LMM
LOCO
LOD score
0
2
4
6
8
1 2 3 4 5 6 7 8 9 10 11
Chromosome

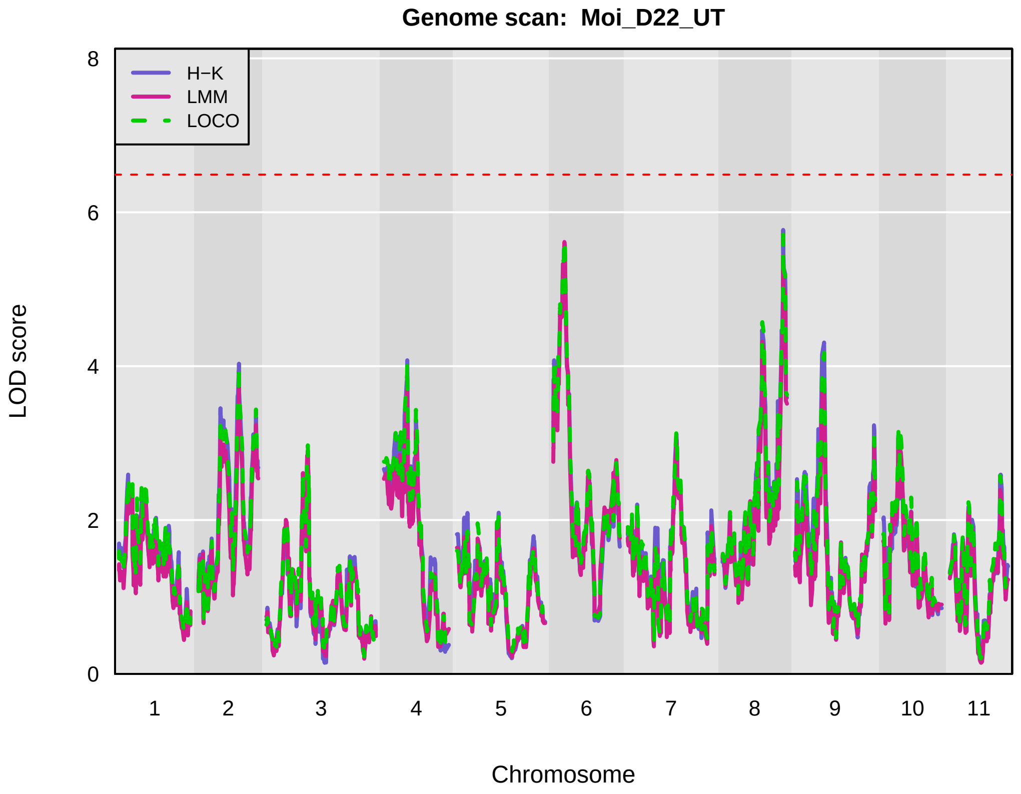
Genome scan: Moi_D22_UT
H−K
LMM
LOCO
LOD score
0
2
4
6
8
1 2 3 4 5 6 7 8 9 10 11
Chromosome

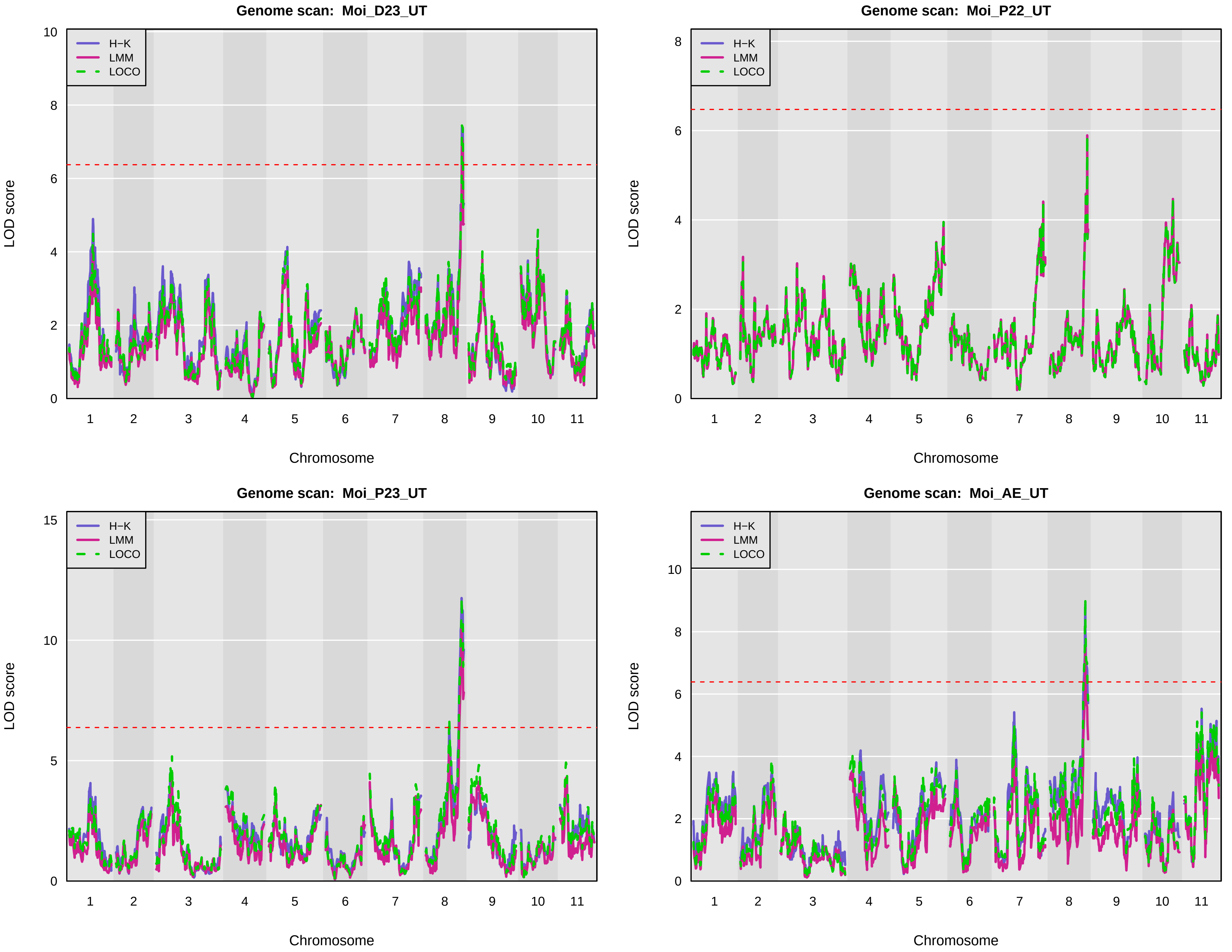

Genome scan: Moi_D23_UT
Genome scan: Moi_P22_UT
Genome scan: Moi_P23_UT
Genome scan: Moi_AE_UT
H−K
LMM
LOCO
LOD score
Chromosome
1 2 3 4 5 6 7 8 9 10 11

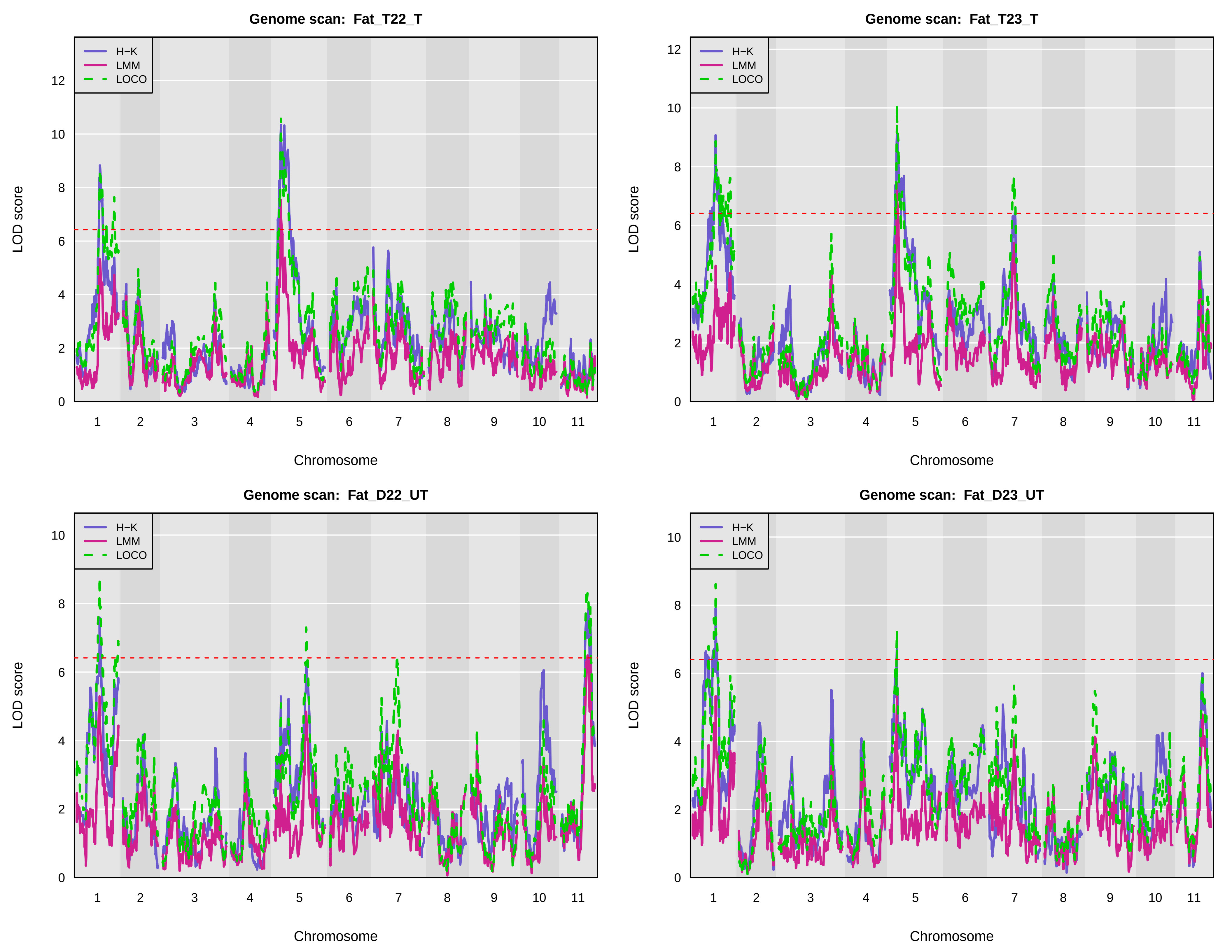
Genome scan: Fat_T22_T
Genome scan: Fat_T23_T
Genome scan: Fat_D22_UT
Genome scan: Fat_D23_UT
H−K
LMM
LOCO
LOD score
Chromosome
1 2 3 4 5 6 7 8 9 10 11
0 2 4 6 8 10 12

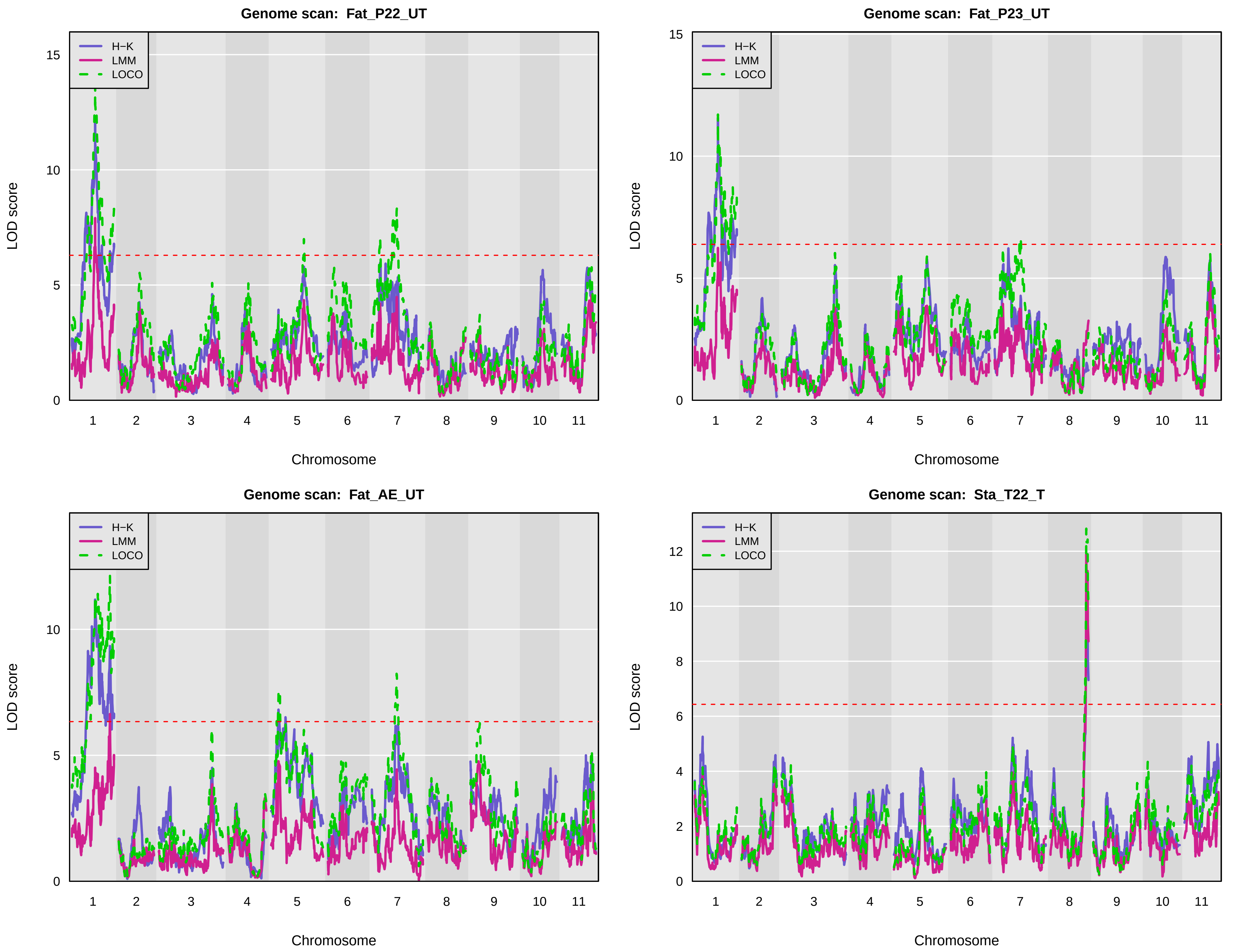
Genome scan: Fat_P22_UT
Genome scan: Fat_P23_UT
Genome scan: Fat_AE_UT
Genome scan: Sta_T22_T
H−K
LMM
LOCO
LOD score
Chromosome
1 2 3 4 5 6 7 8 9 10 11
0
5
10
15
0
2
4
6
8
10
12

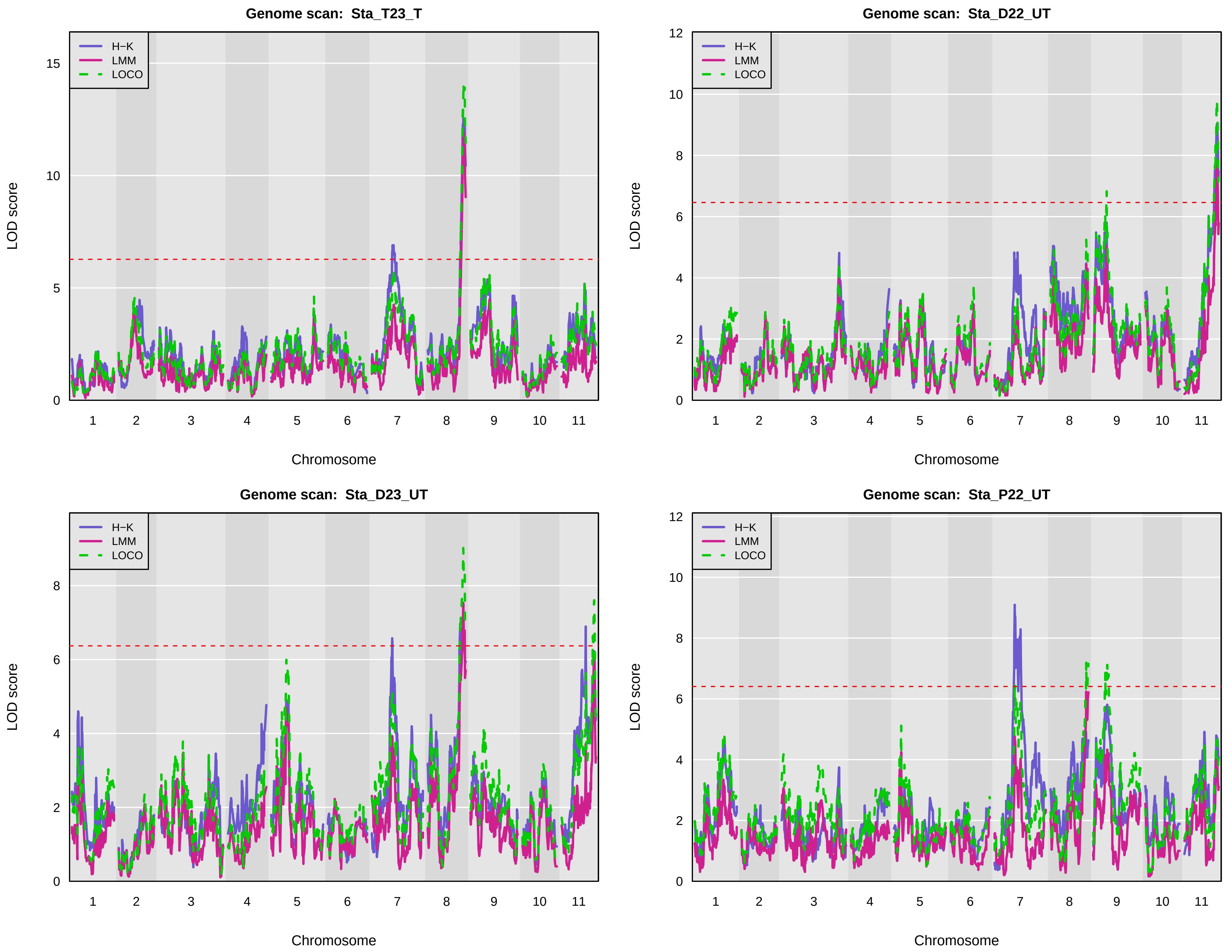
Genome scan: Sta_T23_T
H−K
LMM
LOCO
LOD score
Chromosome
1 2 3 4 5 6 7 8 9 10 11
0 5 10 15
Genome scan: Sta_D22_UT
H−K
LMM
LOCO
LOD score
Chromosome
1 2 3 4 5 6 7 8 9 10 11
0 2 4 6 8 10 12
Genome scan: Sta_D23_UT
H−K
LMM
LOCO
LOD score
Chromosome
1 2 3 4 5 6 7 8 9 10 11
0 2 4 6 8
Genome scan: Sta_P22_UT
H−K
LMM
LOCO
LOD score
Chromosome
1 2 3 4 5 6 7 8 9 10 11
0 2 4 6 8 10 12

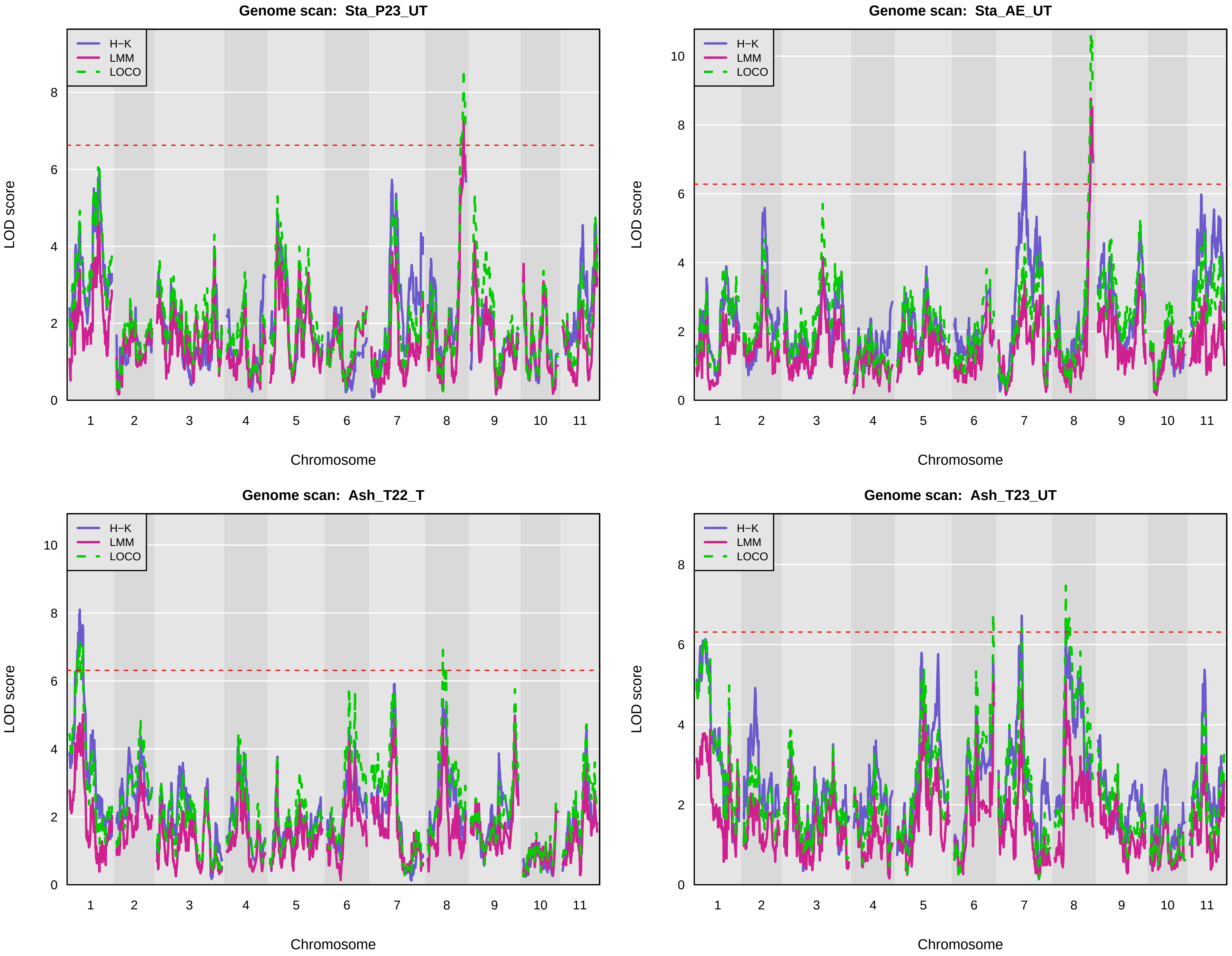

Genome scan: Sta_P23_UT
Genome scan: Sta_AE_UT
Genome scan: Ash_T22_T
Genome scan: Ash_T23_UT
H−K
LMM
LOCO
LOD score
Chromosome
1 2 3 4 5 6 7 8 9 10 11

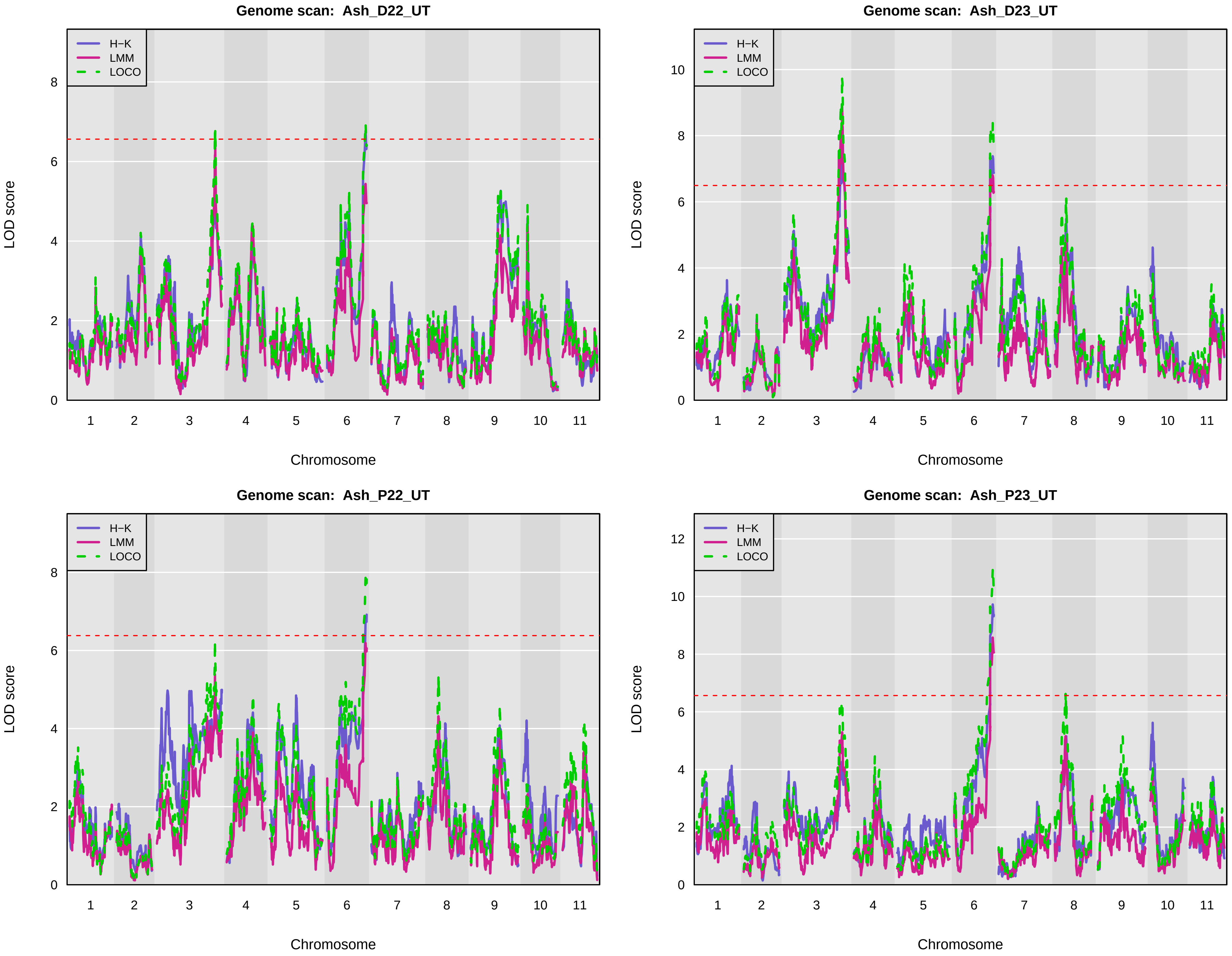
Genome scan: Ash_D22_UT
Genome scan: Ash_D23_UT
Genome scan: Ash_P22_UT
Genome scan: Ash_P23_UT
H−K
LMM
LOCO
LOD score
Chromosome
1 2 3 4 5 6 7 8 9 10 11
0 2 4 6 8 10 12

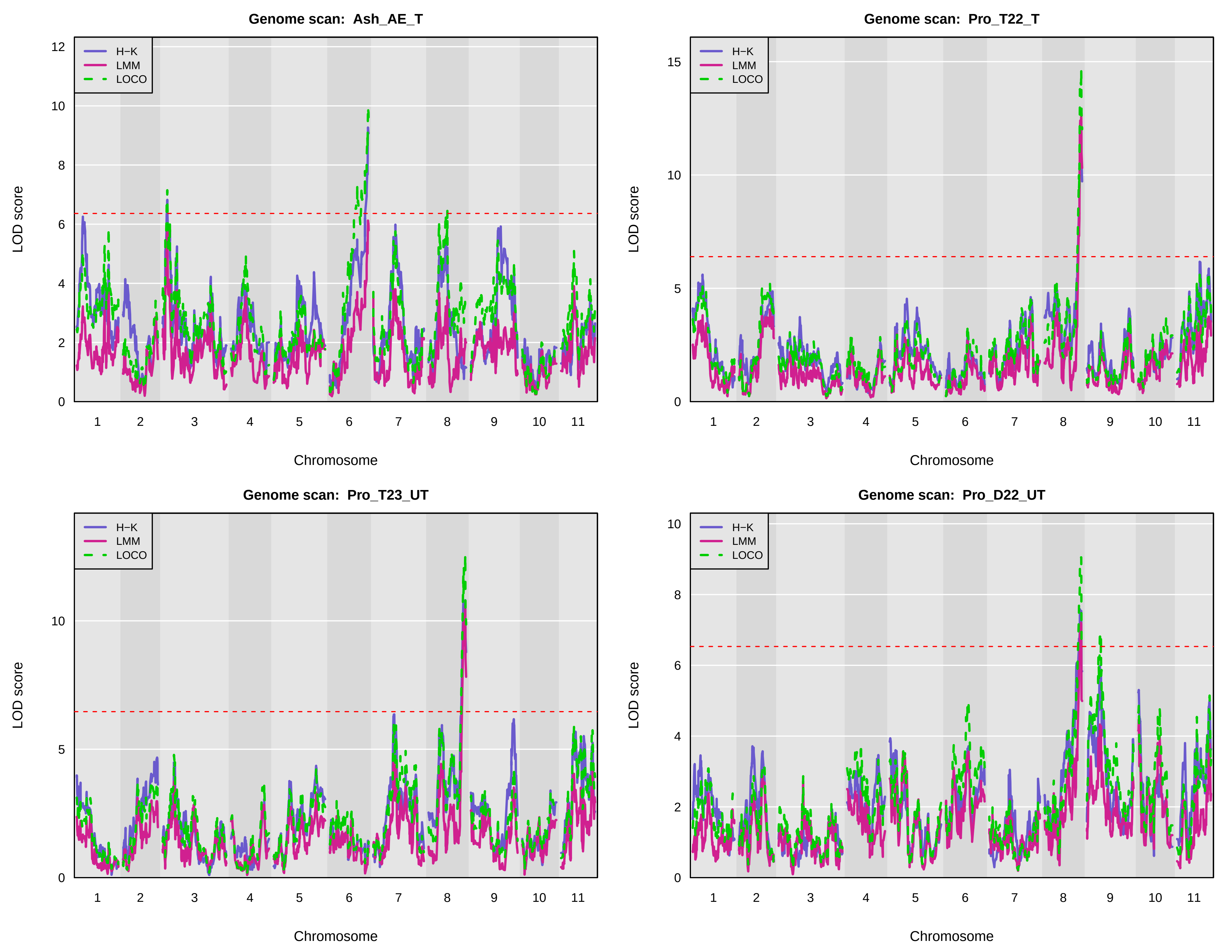
Genome scan: Ash_AE_T
Genome scan: Pro_T22_T
Genome scan: Pro_T23_UT
Genome scan: Pro_D22_UT
H−K
LMM
LOCO
LOD score
Chromosome
1 2 3 4 5 6 7 8 9 10 11
0 2 4 6 8 10 12
0 5 10 15
0 5 10
0 2 4 6 8 10

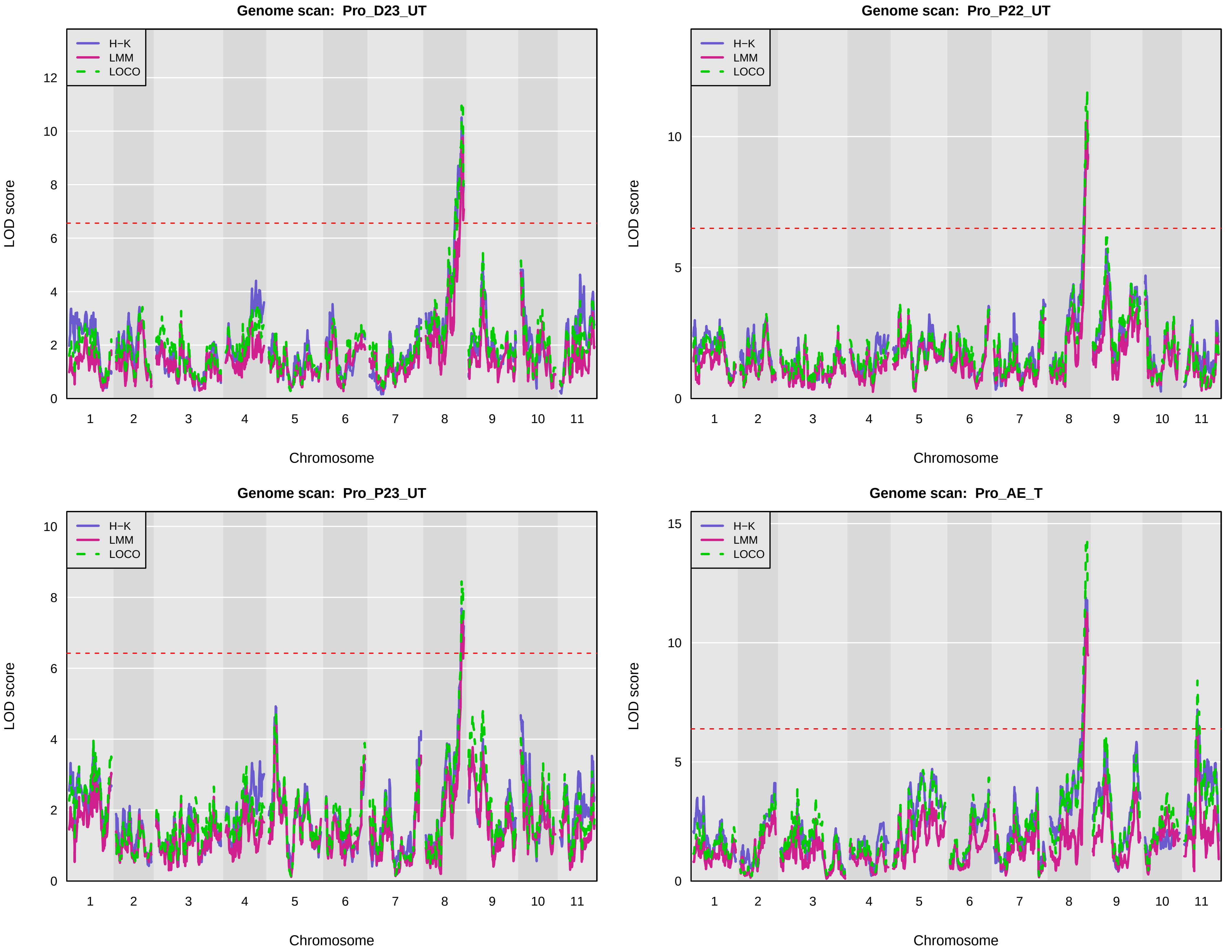
Genome scan: Pro_D23_UT
H−K
LMM
LOCO
LOD score
0 2 4 6 8 10 12
1 2 3 4 5 6 7 8 9 10 11
Chromosome
Genome scan: Pro_P22_UT
H−K
LMM
LOCO
LOD score
0 5 10
1 2 3 4 5 6 7 8 9 10 11
Chromosome
Genome scan: Pro_P23_UT
H−K
LMM
LOCO
LOD score
0 2 4 6 8 10
1 2 3 4 5 6 7 8 9 10 11
Chromosome
Genome scan: Pro_AE_T
H−K
LMM
LOCO
LOD score
0 5 10 15
1 2 3 4 5 6 7 8 9 10 11
Chromosome

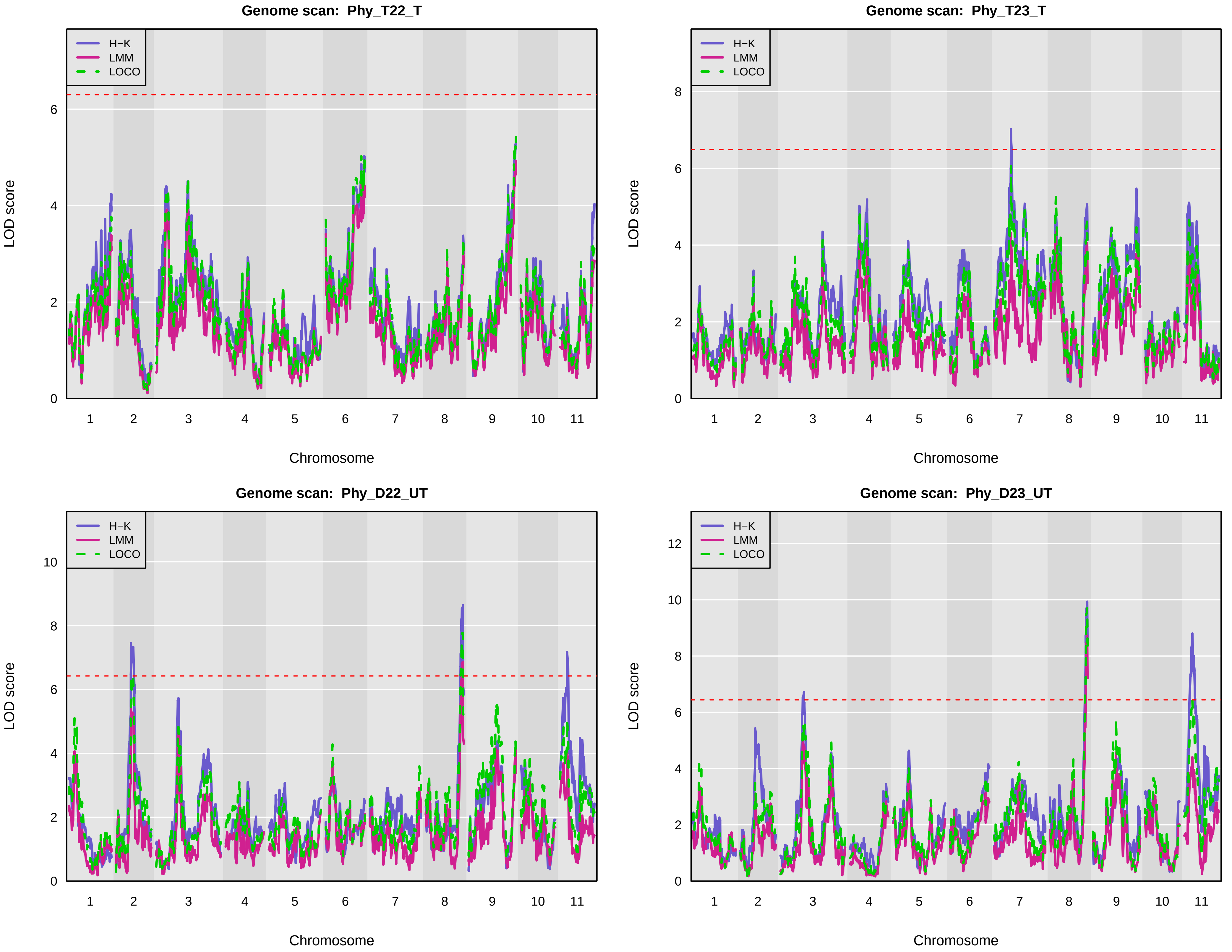

Genome scan: Phy_T22_T
Genome scan: Phy_T23_T
Genome scan: Phy_D22_UT
Genome scan: Phy_D23_UT
H−K
LMM
LOCO
LOD score
Chromosome
1 2 3 4 5 6 7 8 9 10 11

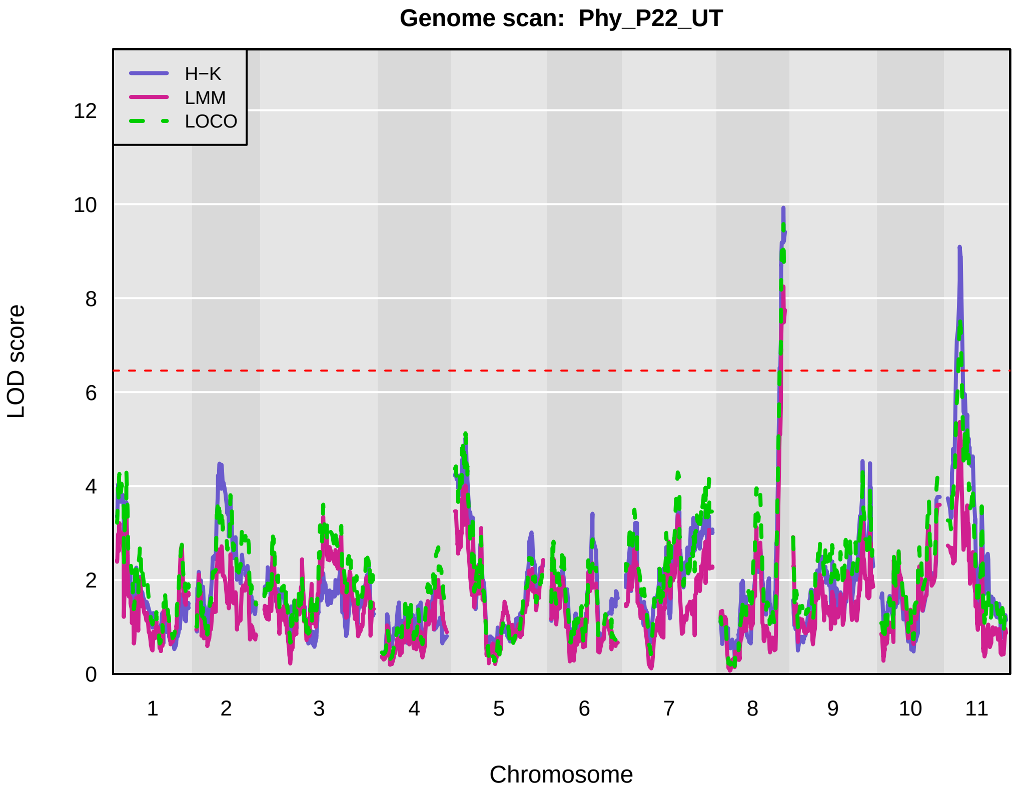
Genome scan: Phy_P22_UT
H−K
LMM
LOCO
LOD score
0
2
4
6
8
10
12
1 2 3 4 5 6 7 8 9 10 11
Chromosome

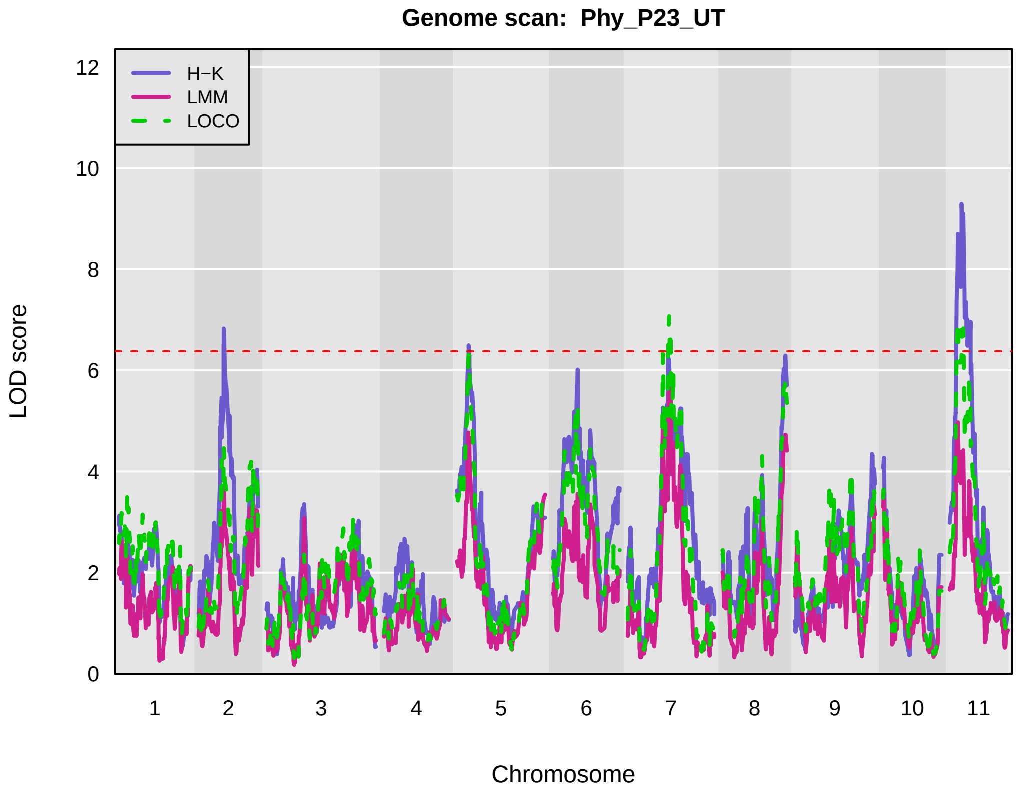
Genome scan: Phy_P23_UT
H−K
LMM
LOCO
LOD score
0
2
4
6
8
10
12
1 2 3 4 5 6 7 8 9 10 11
Chromosome

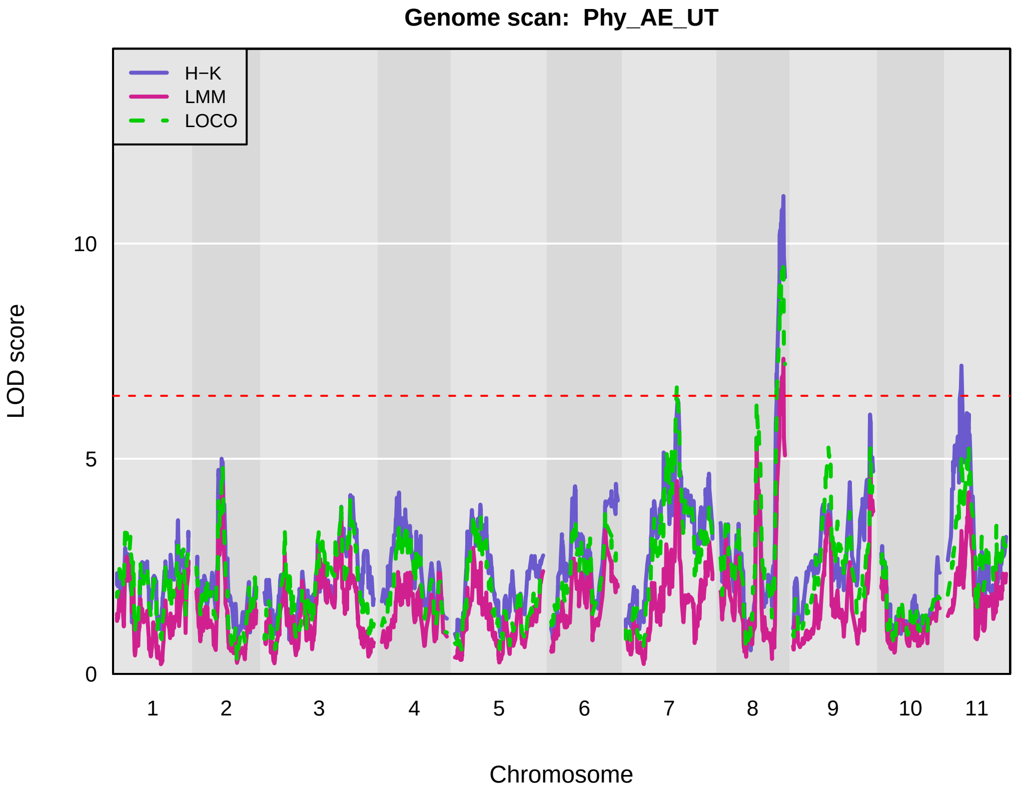
Genome scan: Phy_AE_UT
H−K
LMM
LOCO
LOD score
0
5
10
1 2 3 4 5 6 7 8 9 10 11
Chromosome

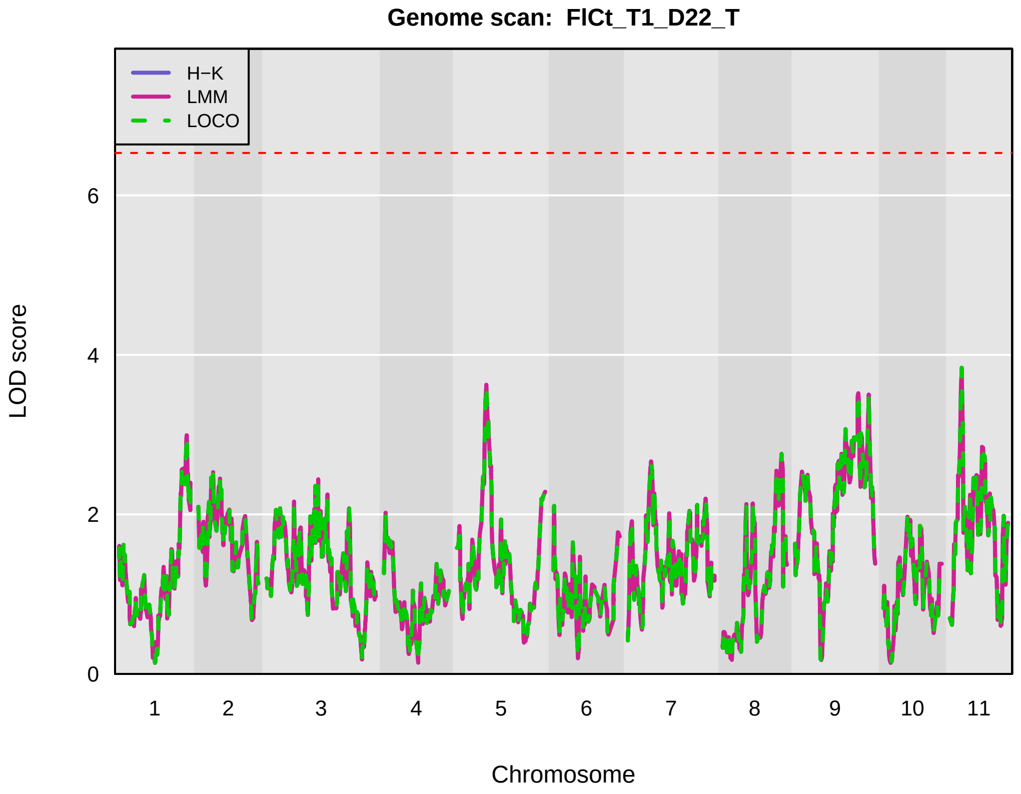
Genome scan: FlCt_T1_D22_T
H−K
LMM
LOCO
LOD score
0
2
4
6
1 2 3 4 5 6 7 8 9 10 11
Chromosome

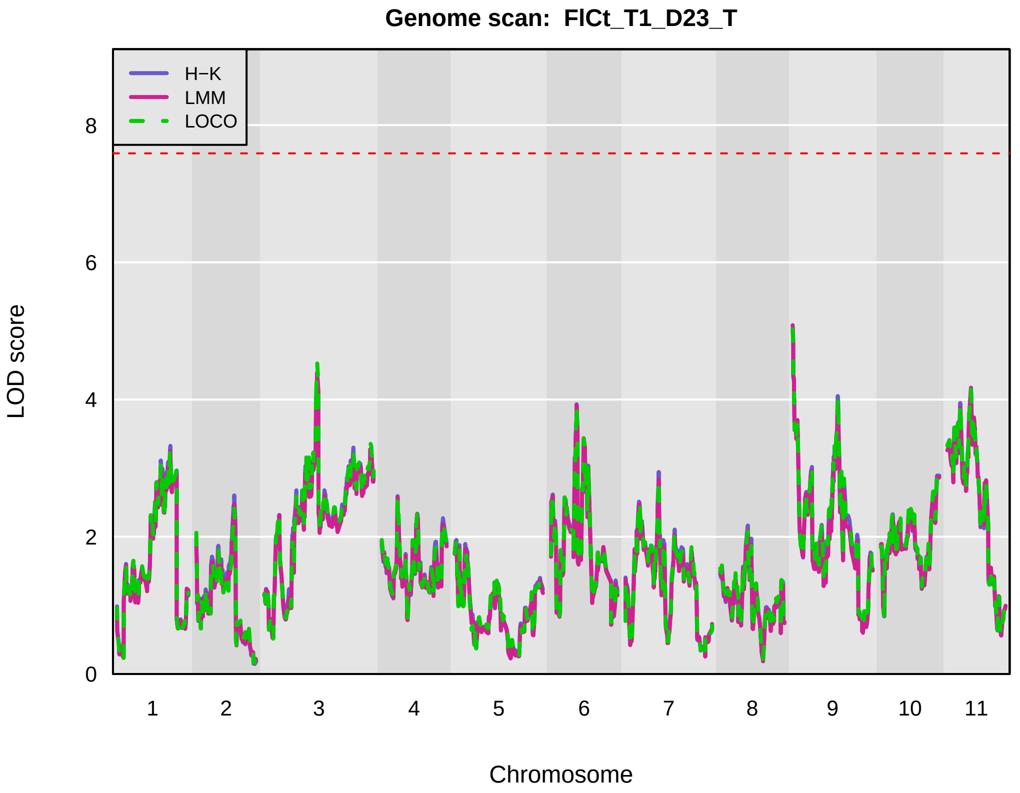

Genome scan: FlCt_T1_D23_T
H−K
LMM
LOCO
LOD score
0
2
4
6
8
1 2 3 4 5 6 7 8 9 10 11
Chromosome


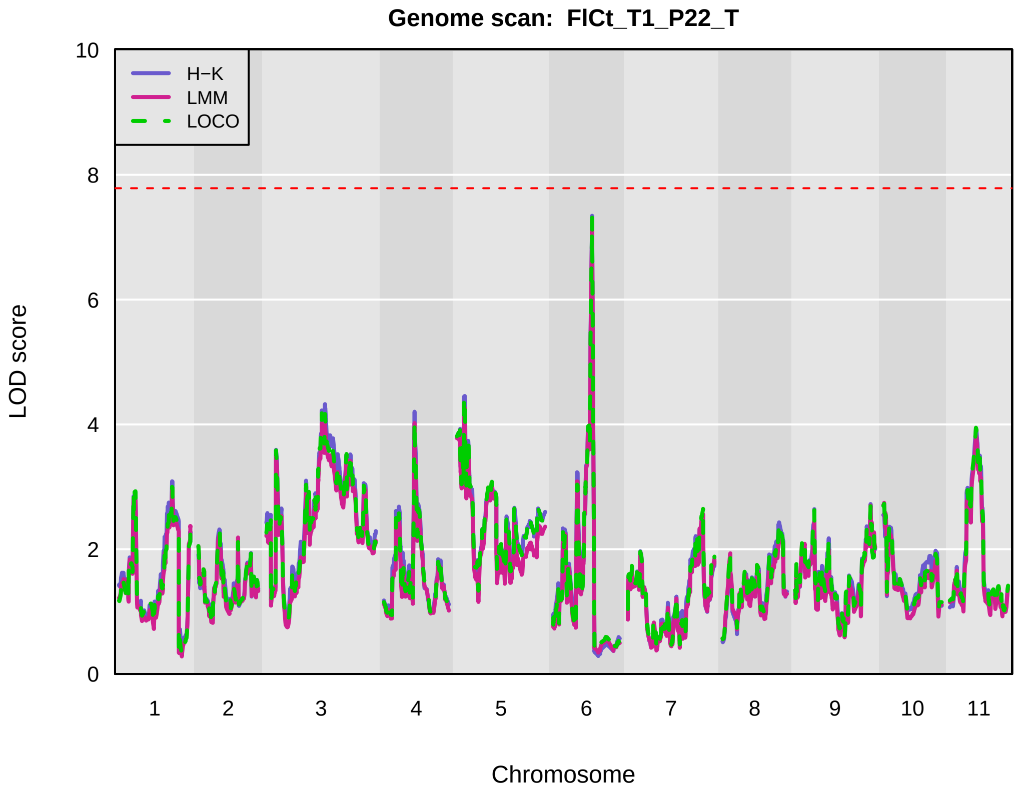

Genome scan: FlCt_T1_P22_T
H−K
LMM
LOCO
LOD score
0
2
4
6
8
10
1 2 3 4 5 6 7 8 9 10 11
Chromosome


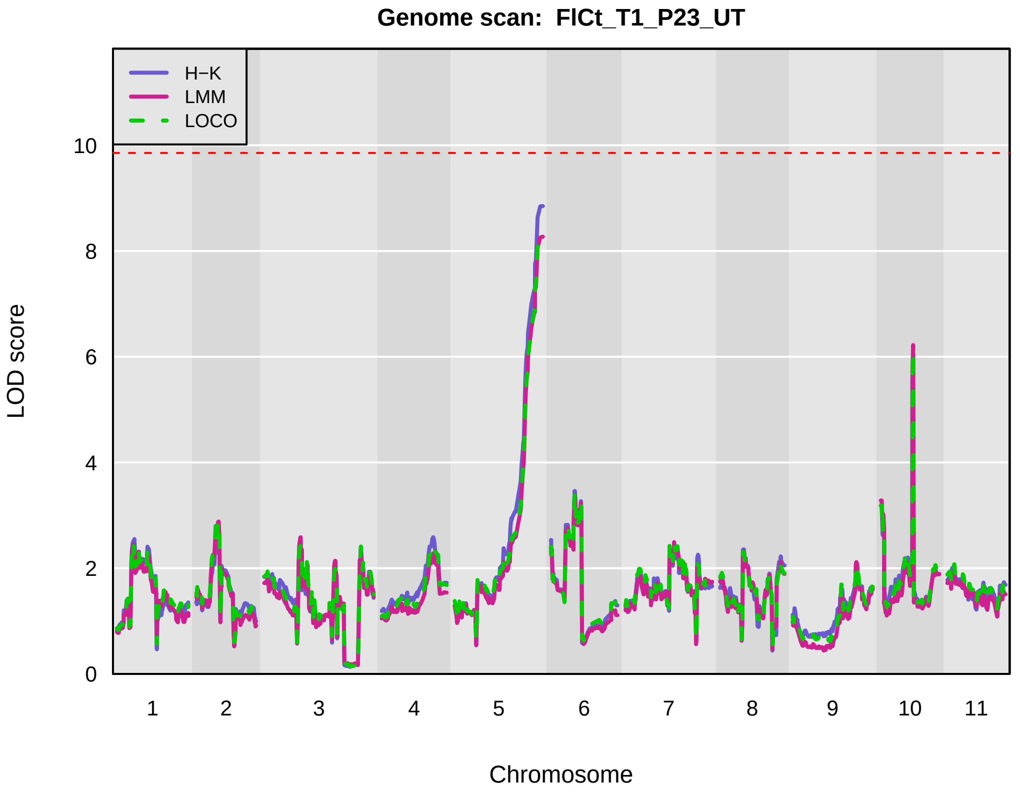

Genome scan: FlCt_T1_P23_UT
H−K
LMM
LOCO
LOD score
0
2
4
6
8
10
1 2 3 4 5 6 7 8 9 10 11
Chromosome


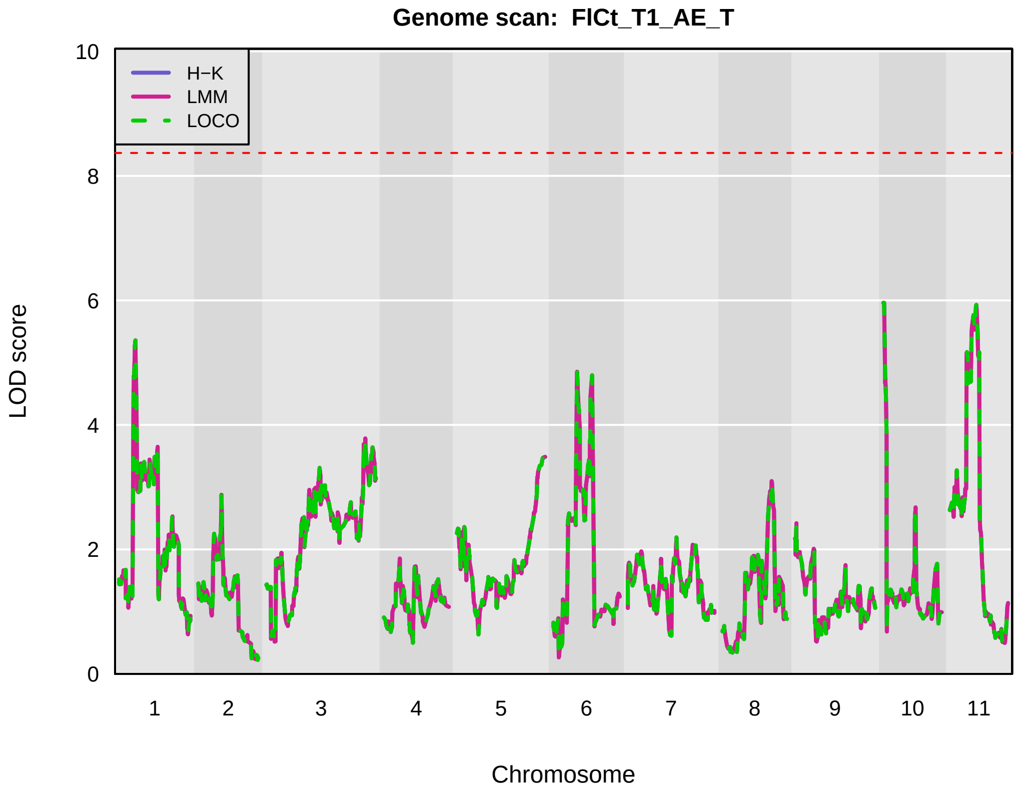

Genome scan: FlCt_T1_AE_T
H−K
LMM
LOCO
LOD score
0
2
4
6
8
10
1 2 3 4 5 6 7 8 9 10 11
Chromosome

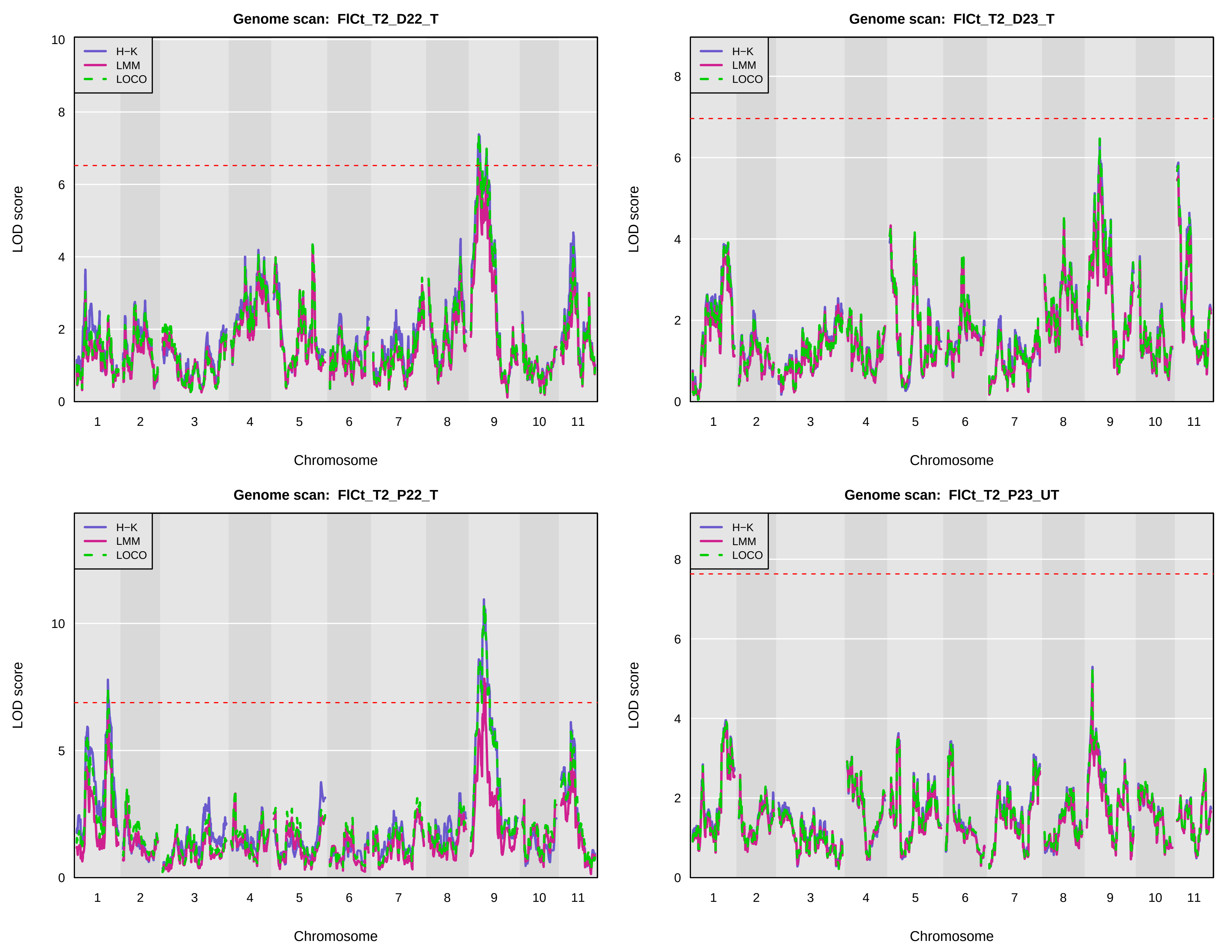
Genome scan: FlCt_T2_D22_T
Genome scan: FlCt_T2_D23_T
Genome scan: FlCt_T2_P22_T
Genome scan: FlCt_T2_P23_UT
H−K
LMM
LOCO
LOD score
Chromosome
1 2 3 4 5 6 7 8 9 10 11

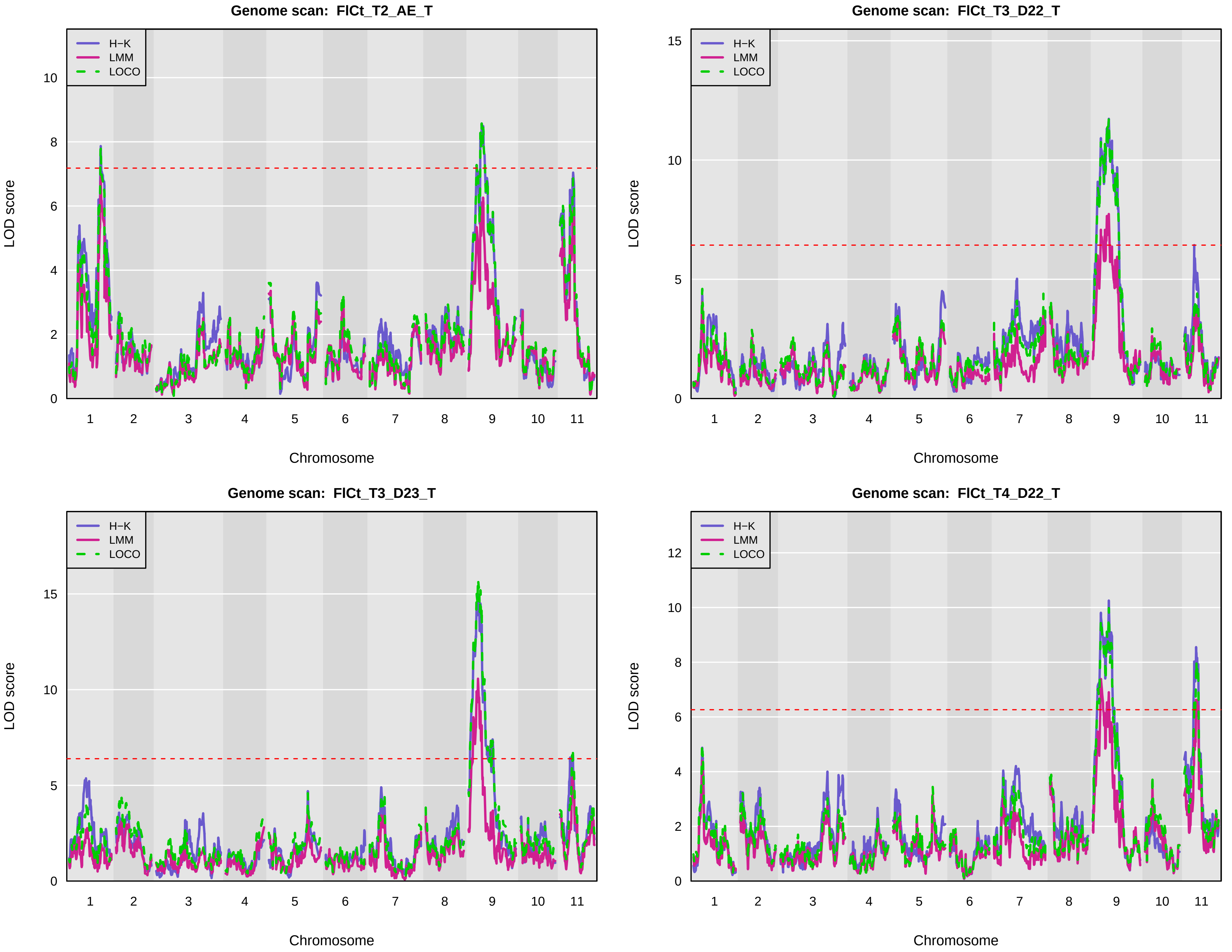
Genome scan: FlCt_T2_AE_T
Genome scan: FlCt_T3_D22_T
Genome scan: FlCt_T3_D23_T
Genome scan: FlCt_T4_D22_T
H−K
LMM
LOCO
LOD score
Chromosome
1 2 3 4 5 6 7 8 9 10 11

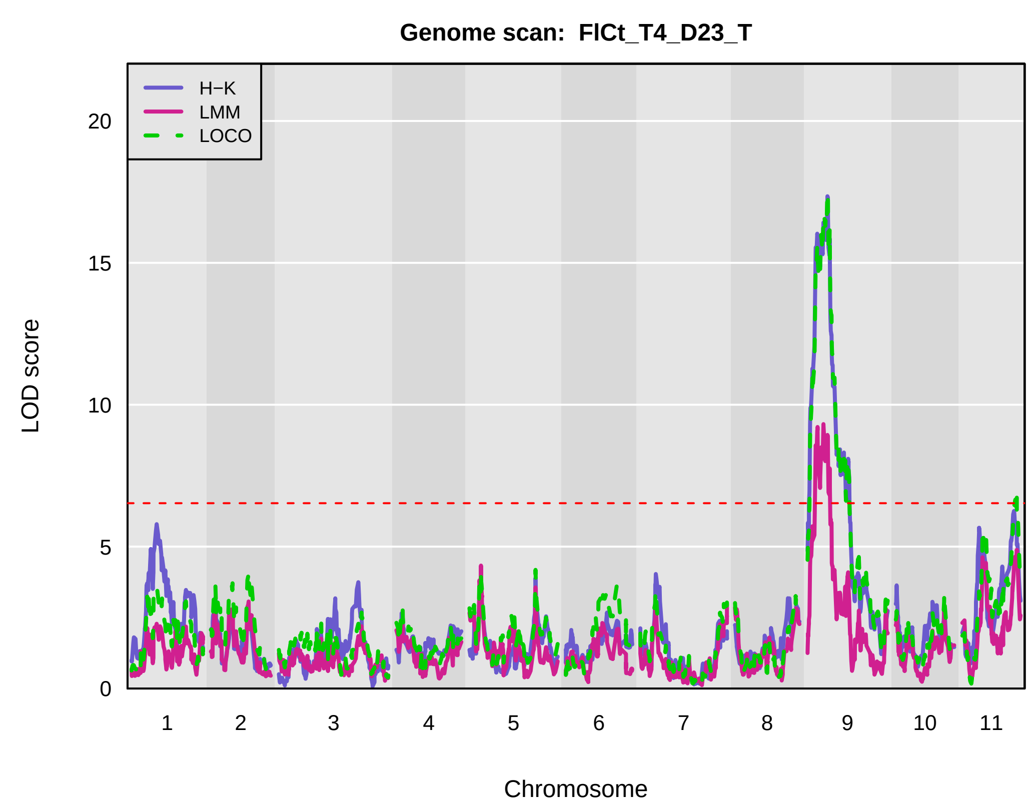
Genome scan: FlCt_T4_D23_T
H−K
LMM
LOCO
LOD score
0 5 10 15 20
1 2 3 4 5 6 7 8 9 10 11
Chromosome

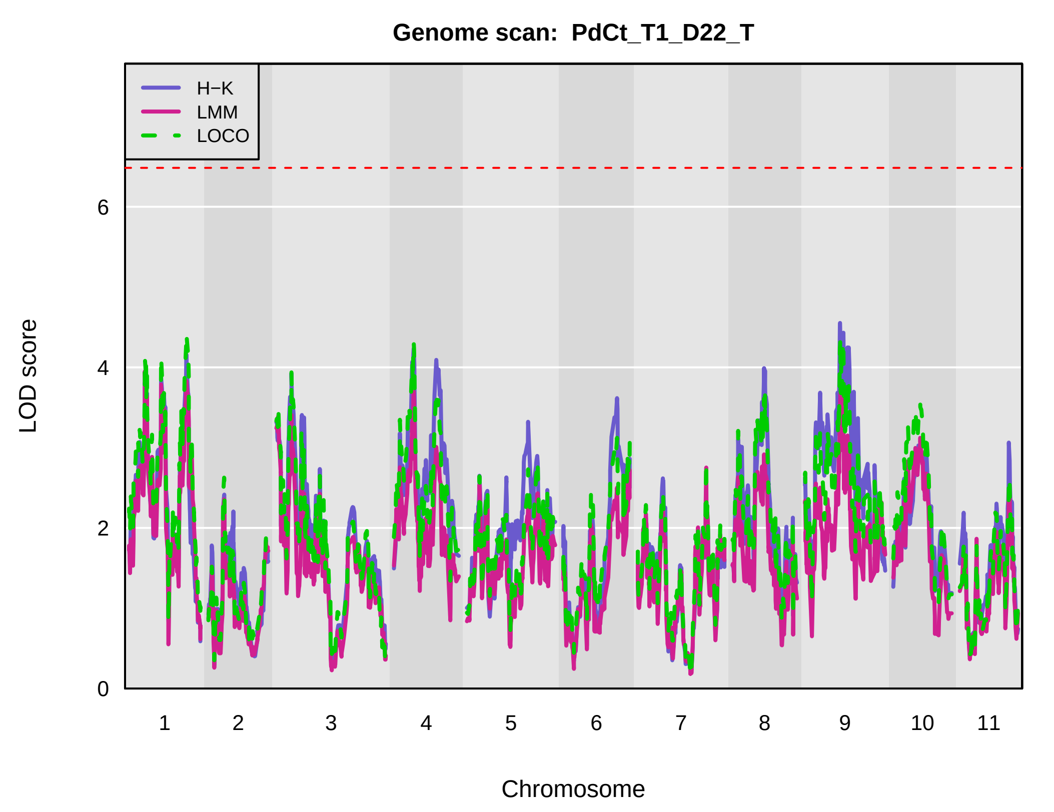
Genome scan: PdCt_T1_D22_T
H−K
LMM
LOCO
LOD score
0 2 4 6
1 2 3 4 5 6 7 8 9 10 11
Chromosome

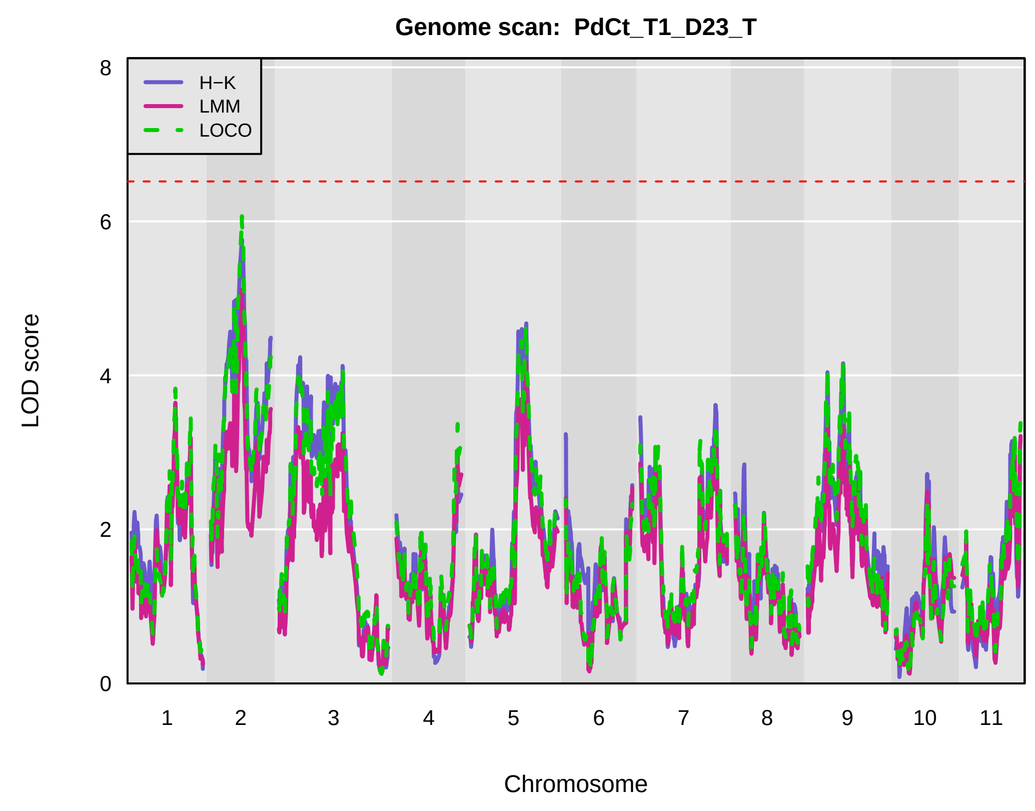
Genome scan: PdCt_T1_D23_T
H−K
LMM
LOCO
LOD score
0 2 4 6 8
1 2 3 4 5 6 7 8 9 10 11
Chromosome

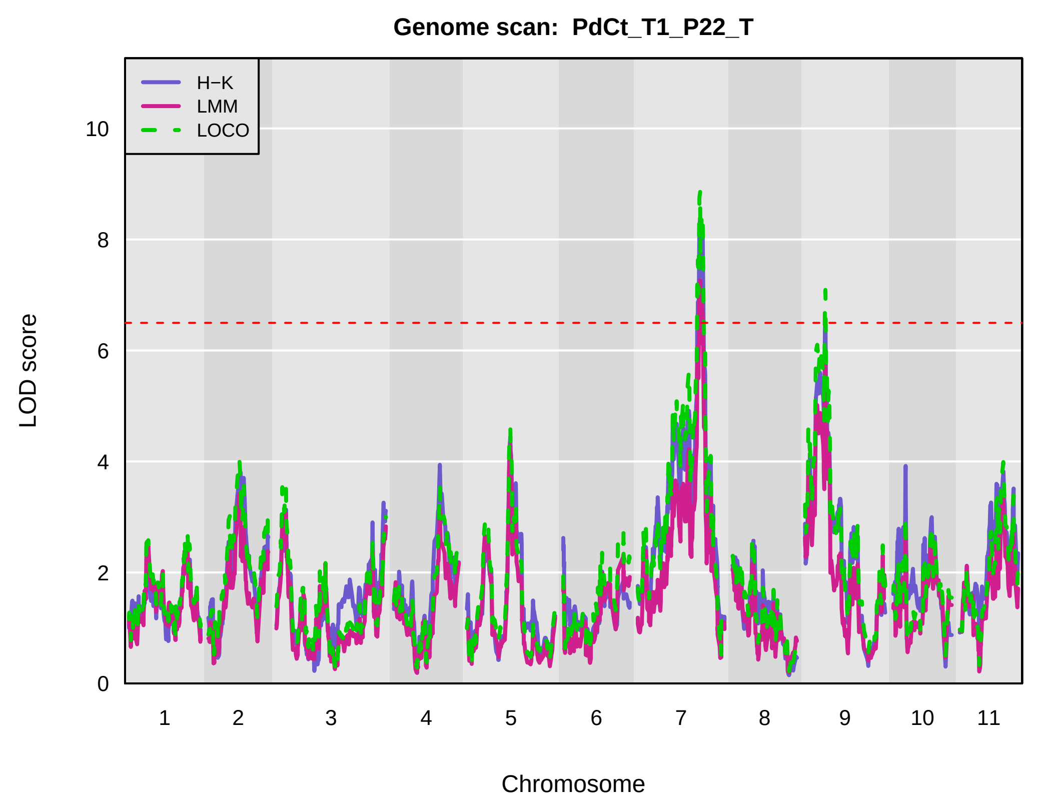
Genome scan: PdCt_T1_P22_T
H−K
LMM
LOCO
LOD score
0 2 4 6 8 10
1 2 3 4 5 6 7 8 9 10 11
Chromosome

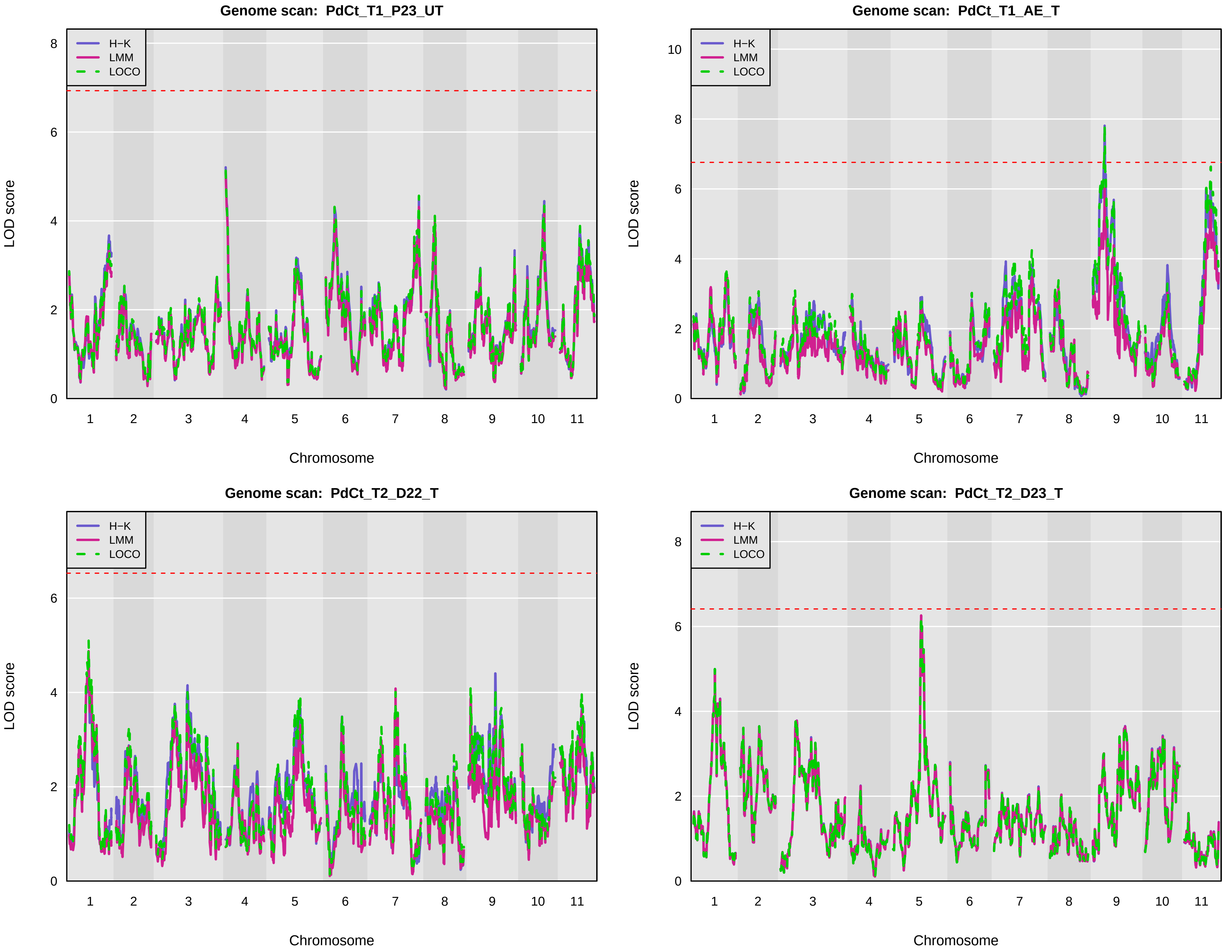

Genome scan: PdCt_T1_P23_UT
Genome scan: PdCt_T1_AE_T
Genome scan: PdCt_T2_D22_T
Genome scan: PdCt_T2_D23_T
H−K
LMM
LOCO
LOD score
Chromosome
1 2 3 4 5 6 7 8 9 10 11

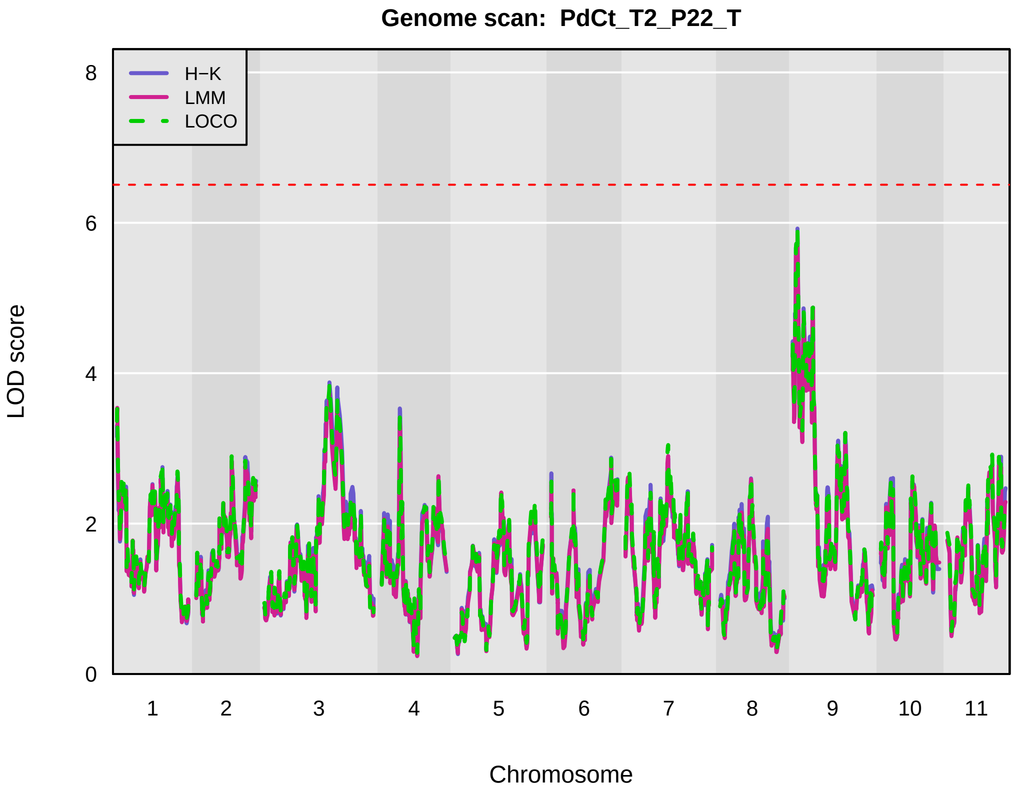
Genome scan: PdCt_T2_P22_T
H−K
LMM
LOCO
LOD score
Chromosome
0
2
4
6
8
1
2
3
4
5
6
7
8
9
10
11

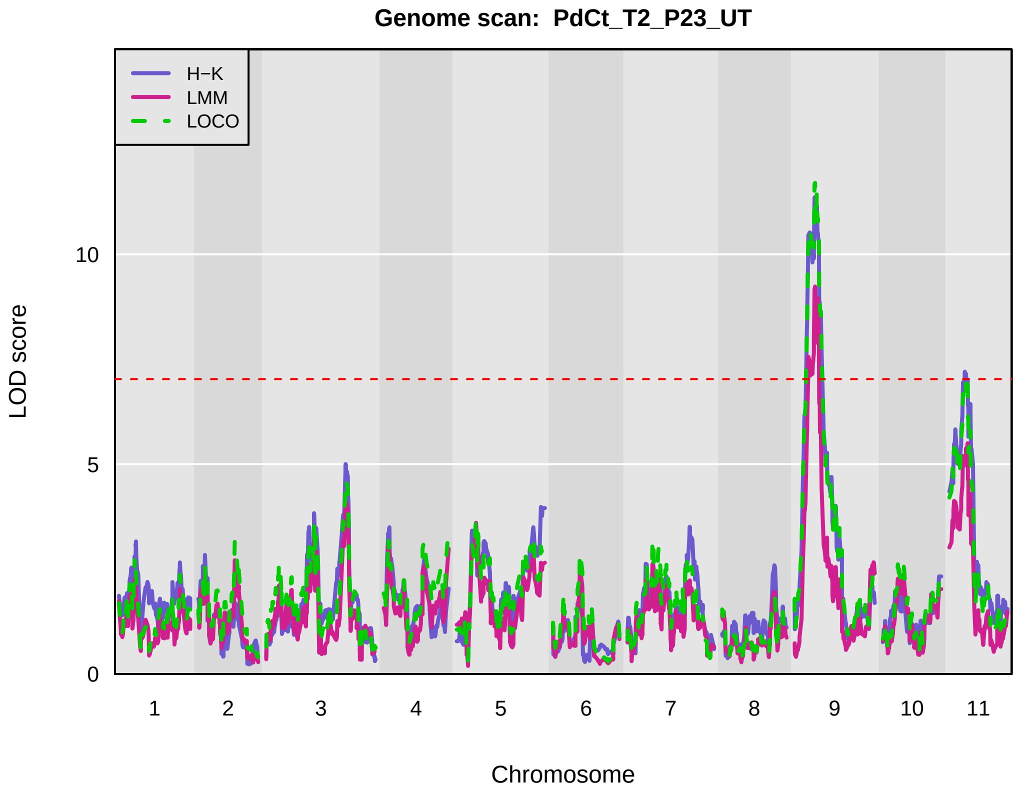
Genome scan: PdCt_T2_P23_UT
H−K
LMM
LOCO
LOD score
Chromosome
0
5
10
1
2
3
4
5
6
7
8
9
10
11

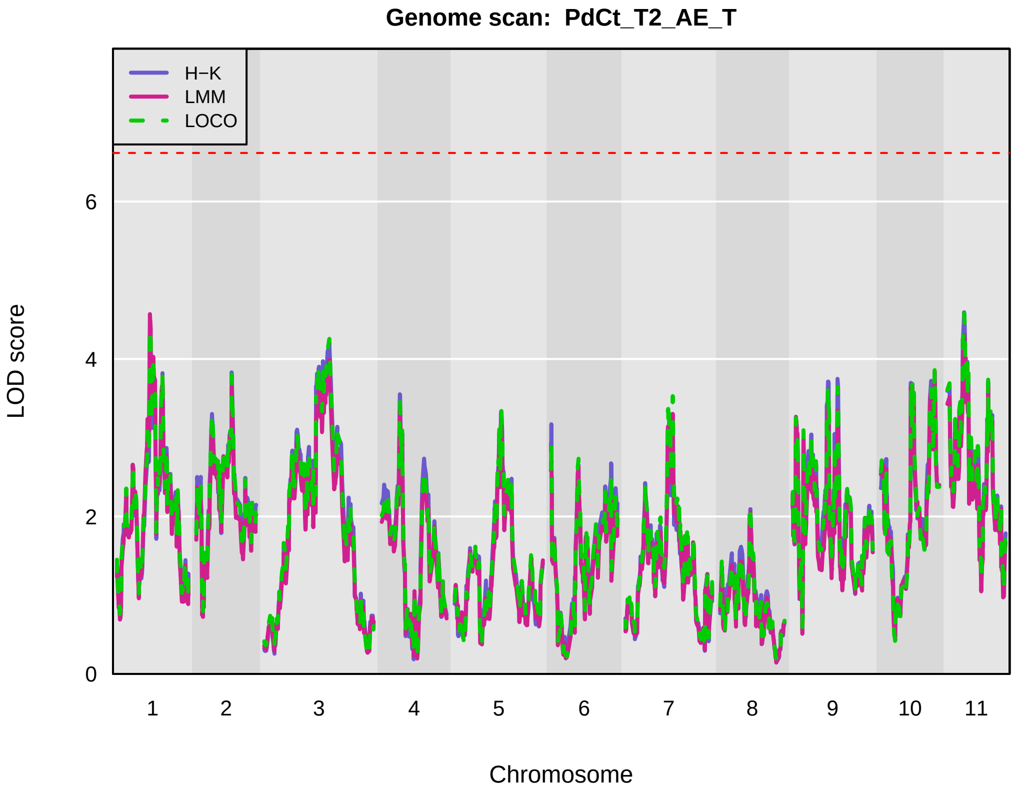
Genome scan: PdCt_T2_AE_T
H−K
LMM
LOCO
LOD score
Chromosome
0
2
4
6
1
2
3
4
5
6
7
8
9
10
11

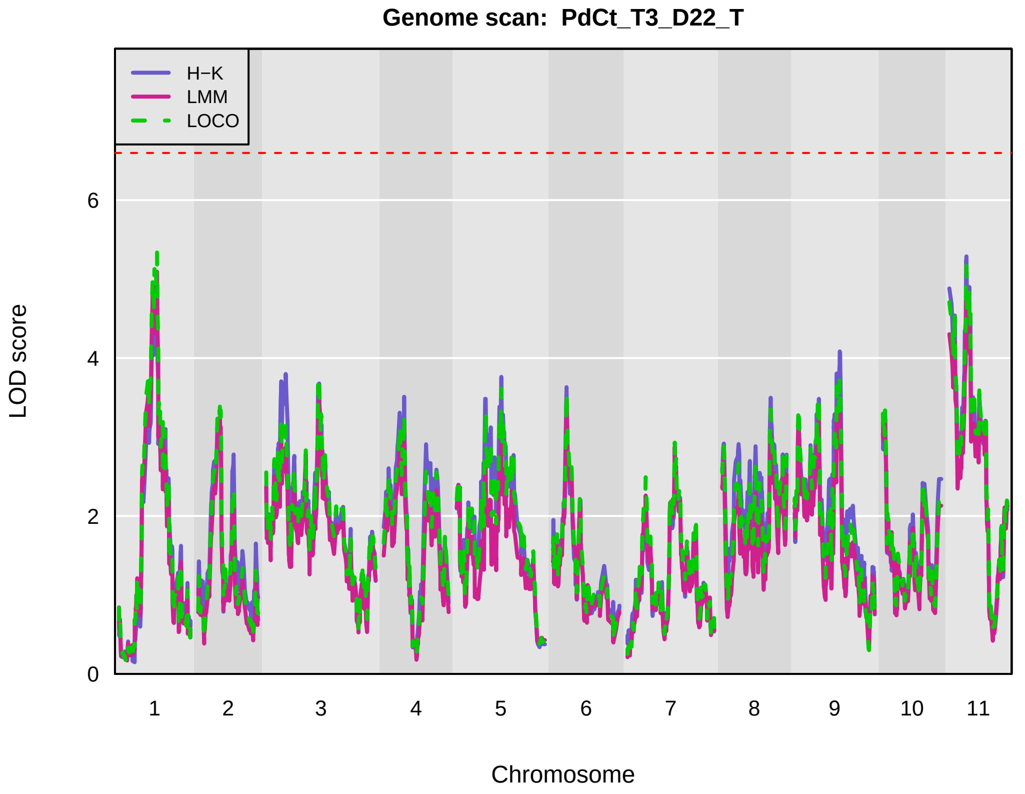
Genome scan: PdCt_T3_D22_T
H−K
LMM
LOCO
LOD score
Chromosome
0
2
4
6
1
2
3
4
5
6
7
8
9
10
11

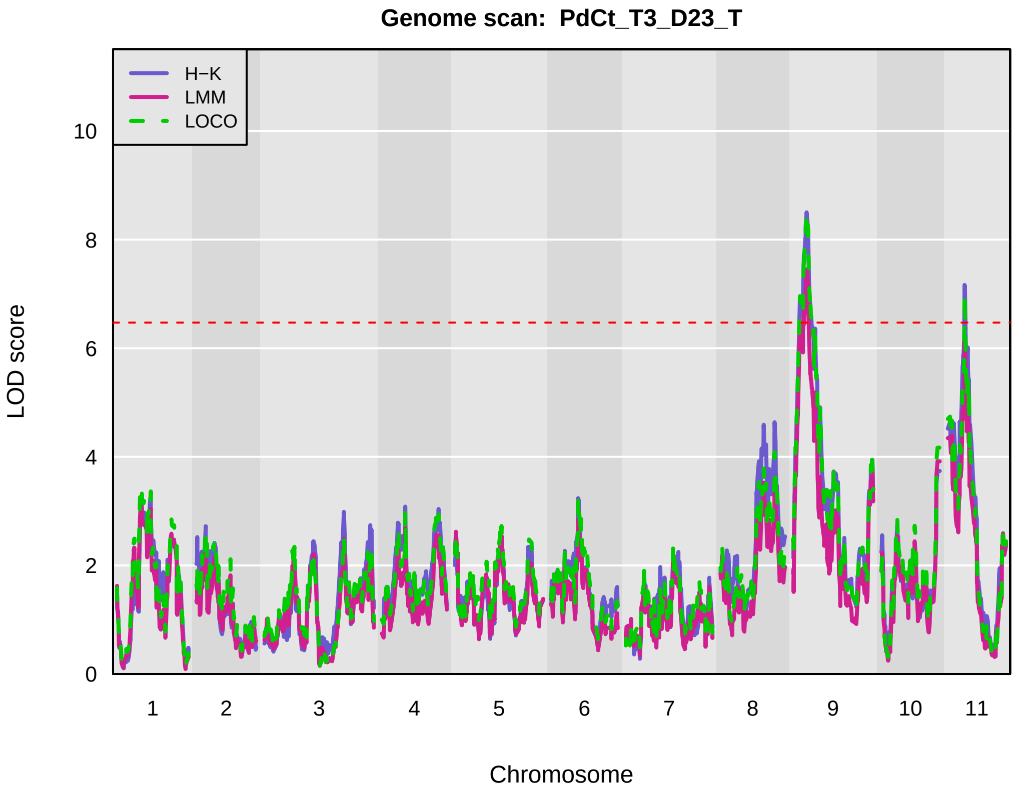
Genome scan: PdCt_T3_D23_T
H−K
LMM
LOCO
LOD score
0
2
4
6
8
10
1 2 3 4 5 6 7 8 9 10 11
Chromosome

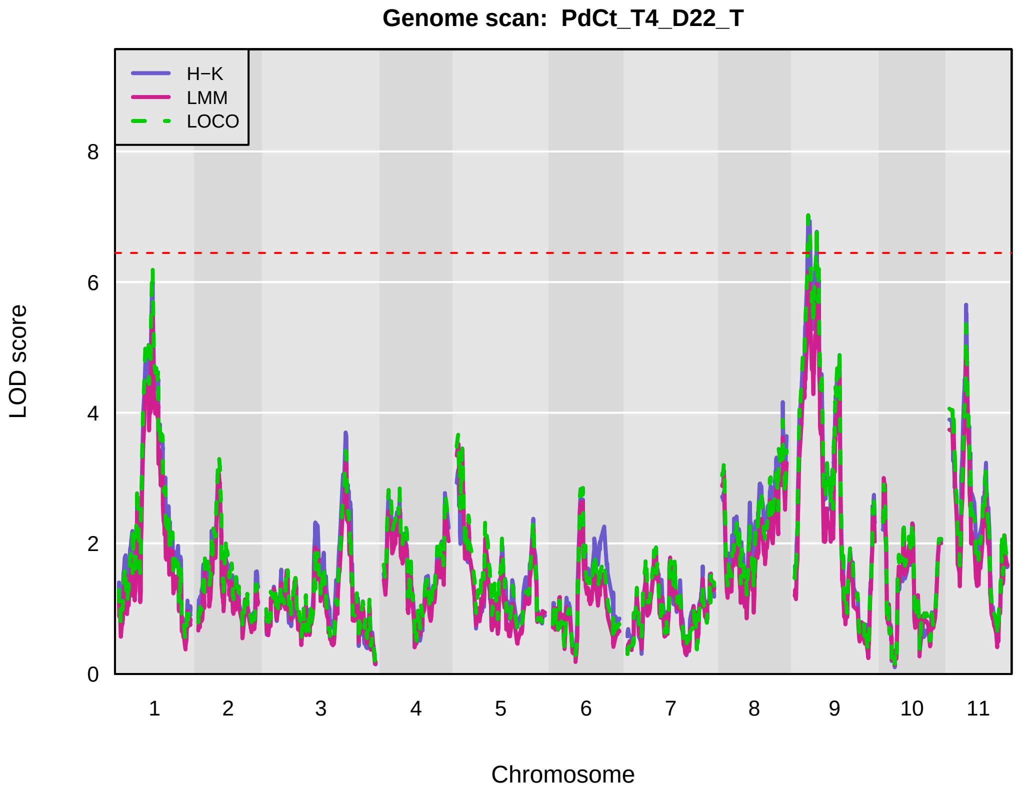
Genome scan: PdCt_T4_D22_T
H−K
LMM
LOCO
LOD score
0
2
4
6
8
1 2 3 4 5 6 7 8 9 10 11
Chromosome

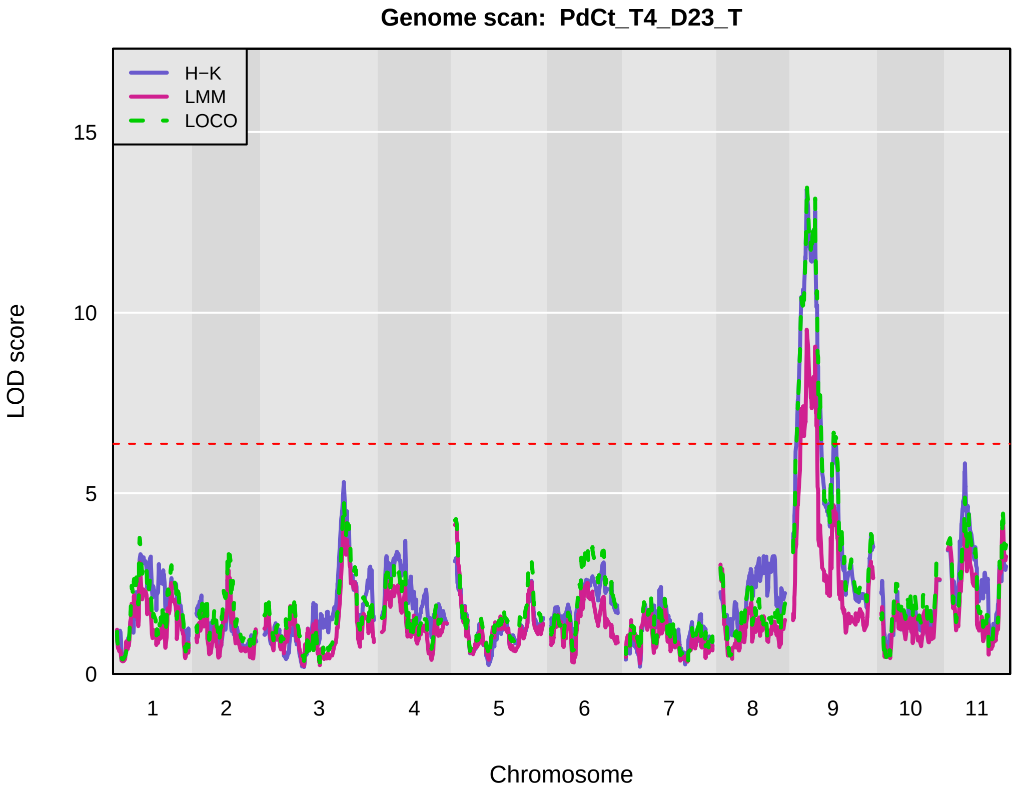
Genome scan: PdCt_T4_D23_T
H−K
LMM
LOCO
LOD score
0
5
10
15
1 2 3 4 5 6 7 8 9 10 11
Chromosome

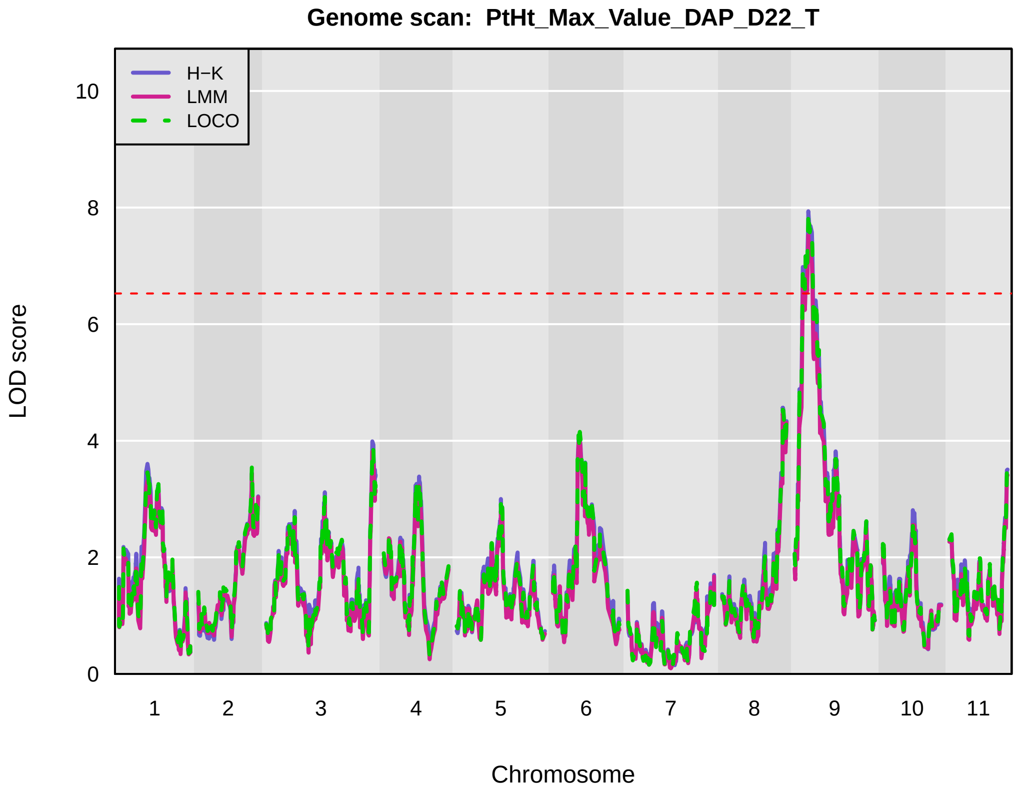
Genome scan: PtHt_Max_Value_DAP_D22_T
H−K
LMM
LOCO
LOD score
0
2
4
6
8
10
1 2 3 4 5 6 7 8 9 10 11
Chromosome

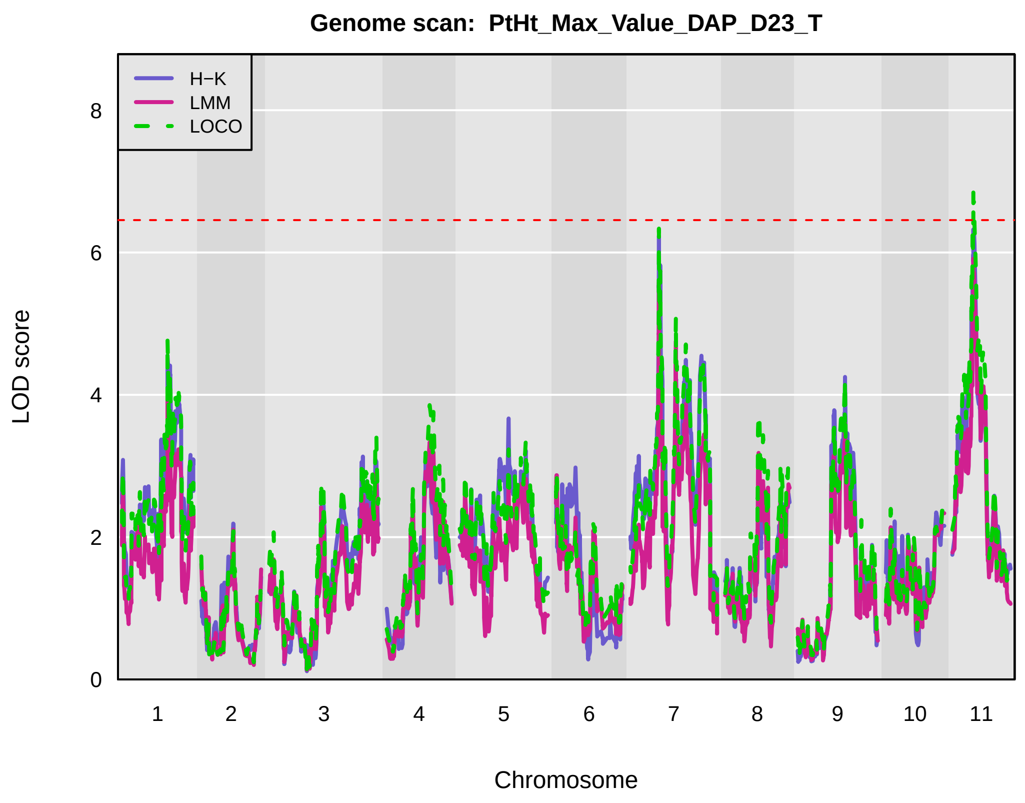
Genome scan: PtHt_Max_Value_DAP_D23_T
H−K
LMM
LOCO
LOD score
0
2
4
6
8
Chromosome
1 2 3 4 5 6 7 8 9 10 11

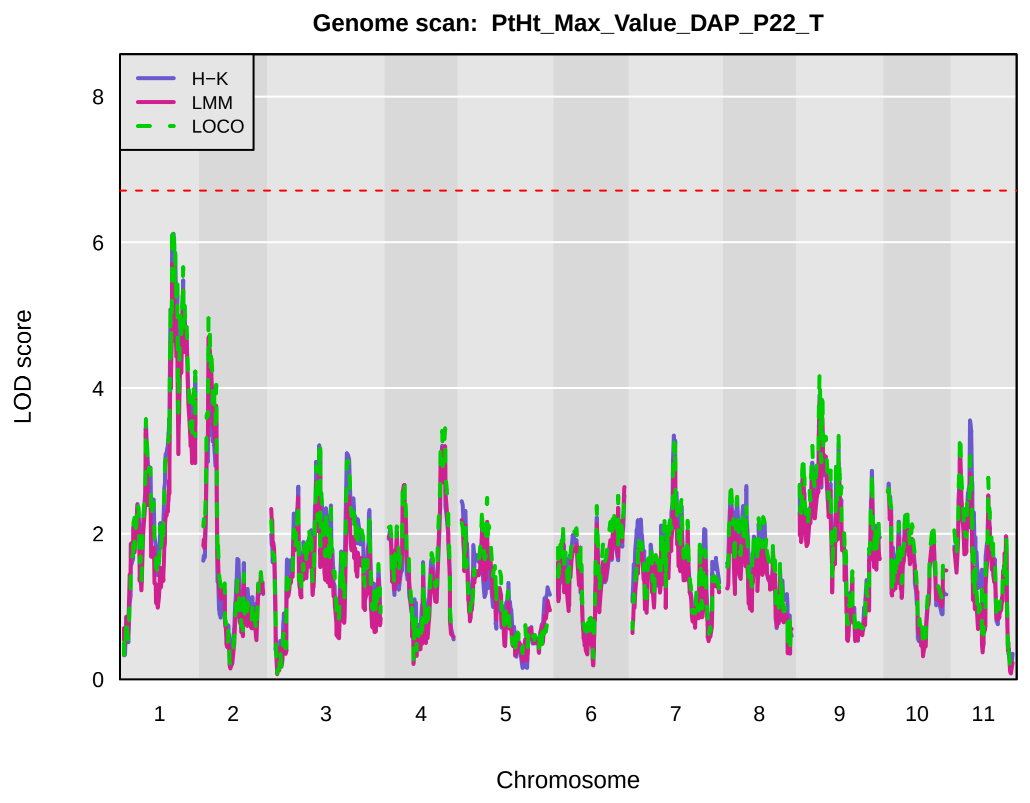
Genome scan: PtHt_Max_Value_DAP_P22_T
H−K
LMM
LOCO
LOD score
0
2
4
6
8
Chromosome
1 2 3 4 5 6 7 8 9 10 11

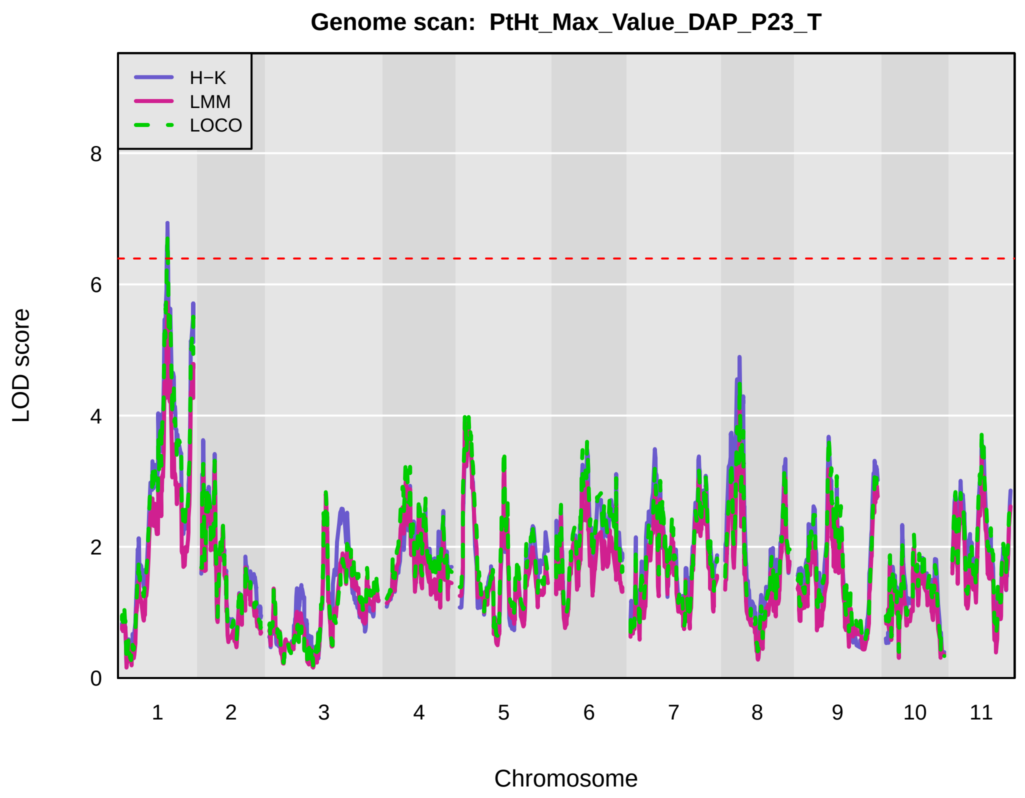
Genome scan: PtHt_Max_Value_DAP_P23_T
H−K
LMM
LOCO
LOD score
0
2
4
6
8
Chromosome
1 2 3 4 5 6 7 8 9 10 11

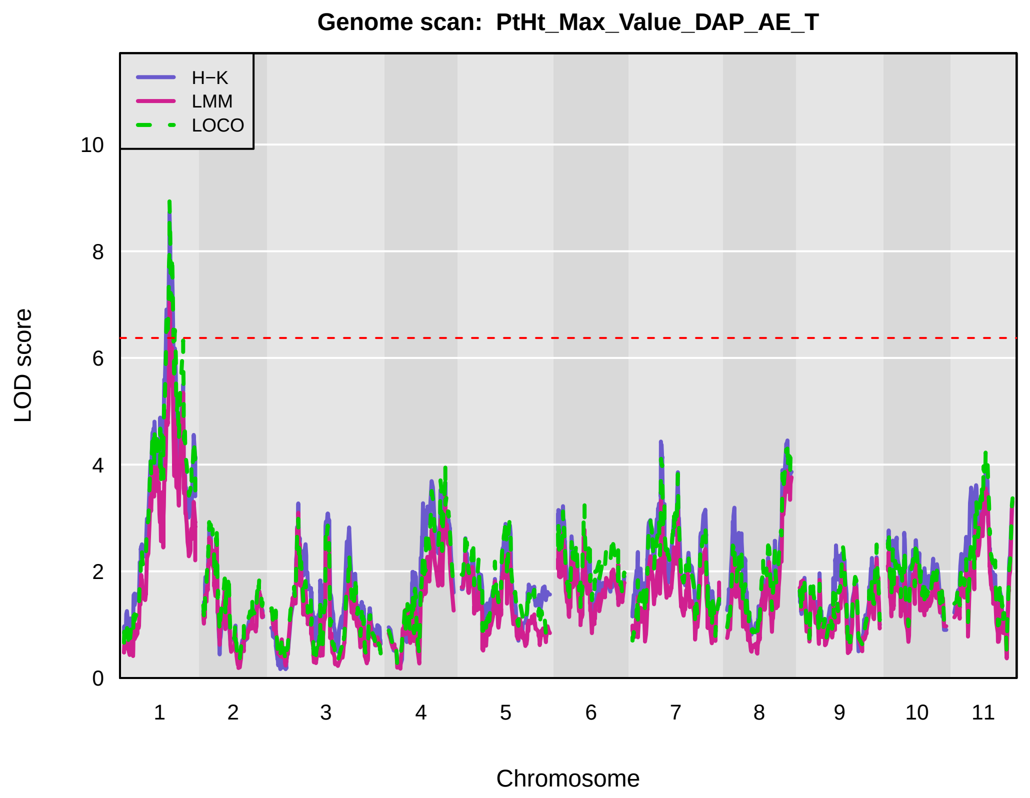
Genome scan: PtHt_Max_Value_DAP_AE_T
H−K
LMM
LOCO
LOD score
0
2
4
6
8
10
Chromosome
1 2 3 4 5 6 7 8 9 10 11

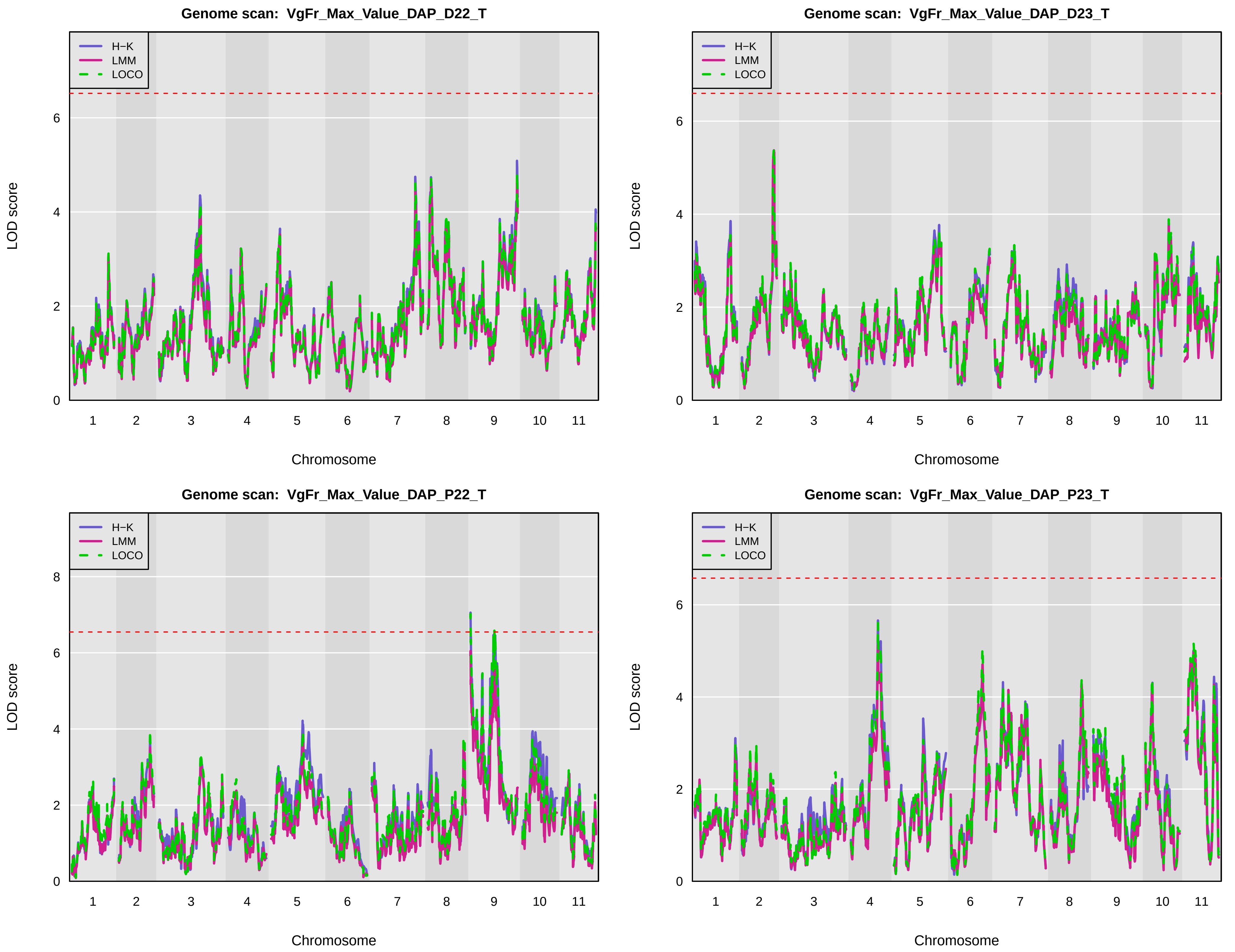
Genome scan: VgFr_Max_Value_DAP_D22_T
Genome scan: VgFr_Max_Value_DAP_D23_T
Genome scan: VgFr_Max_Value_DAP_P22_T
Genome scan: VgFr_Max_Value_DAP_P23_T
H−K
LMM
LOCO
LOD score
Chromosome
1
2
3
4
5
6
7
8
9
10
11
0
2
4
6
8

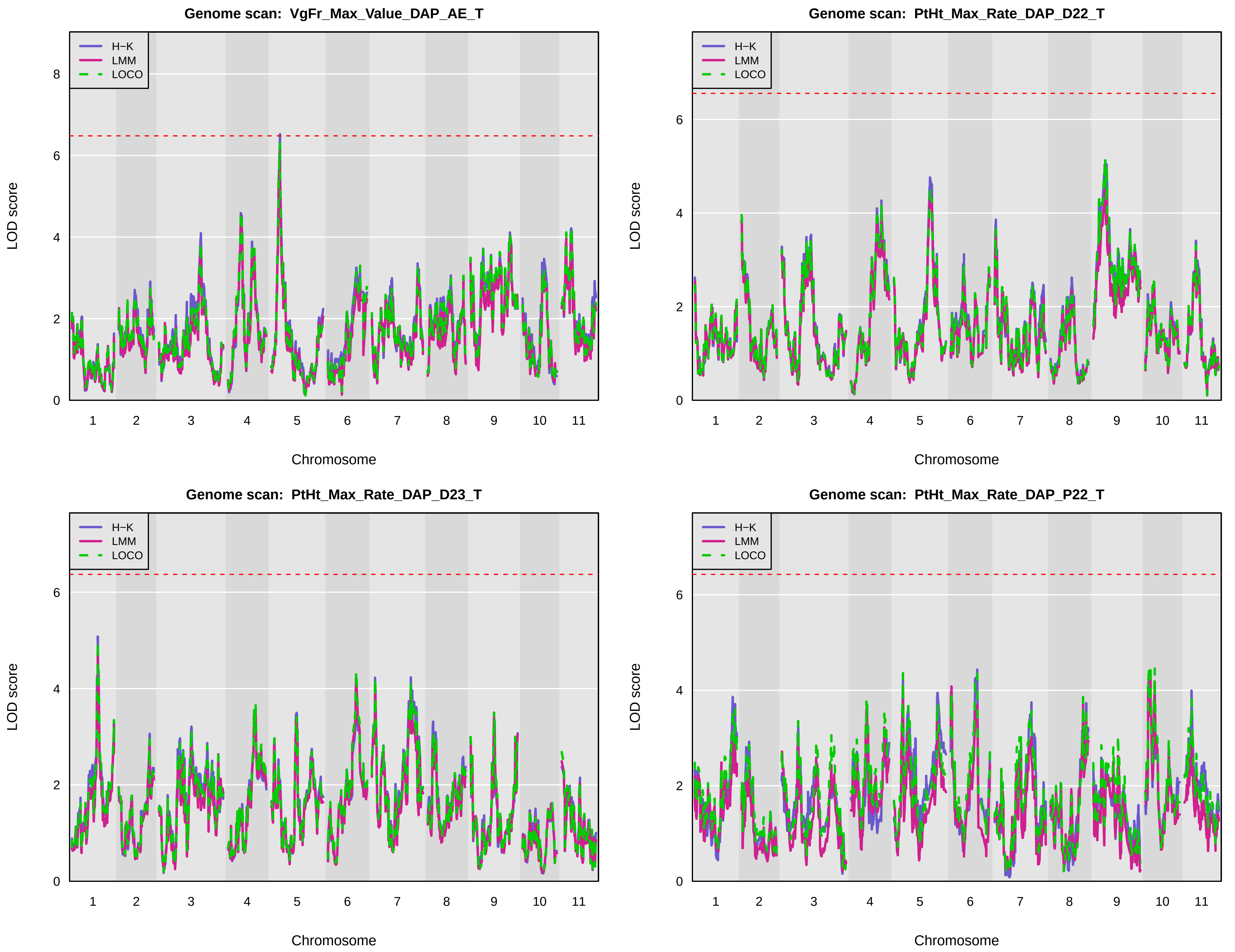
Genome scan: VgFr_Max_Value_DAP_AE_T
Genome scan: PtHt_Max_Rate_DAP_D22_T
Genome scan: PtHt_Max_Rate_DAP_D23_T
Genome scan: PtHt_Max_Rate_DAP_P22_T
H−K
LMM
LOCO
LOD score
Chromosome
1 2 3 4 5 6 7 8 9 10 11

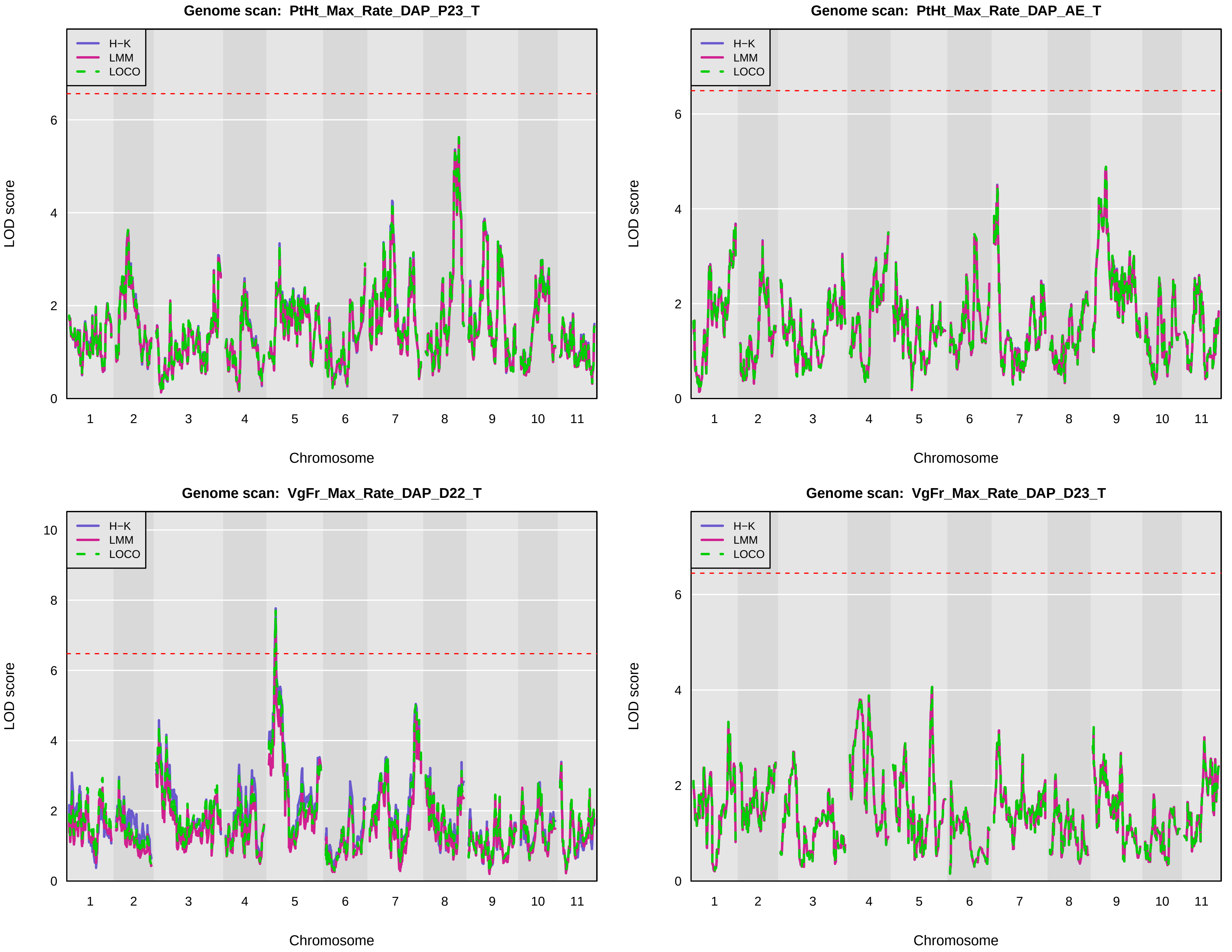
Genome scan: PtHt_Max_Rate_DAP_P23_T
Genome scan: PtHt_Max_Rate_DAP_AE_T
Genome scan: VgFr_Max_Rate_DAP_D22_T
Genome scan: VgFr_Max_Rate_DAP_D23_T
H−K
LMM
LOCO
LOD score
Chromosome
1 2 3 4 5 6 7 8 9 10 11
0 2 4 6 8 10

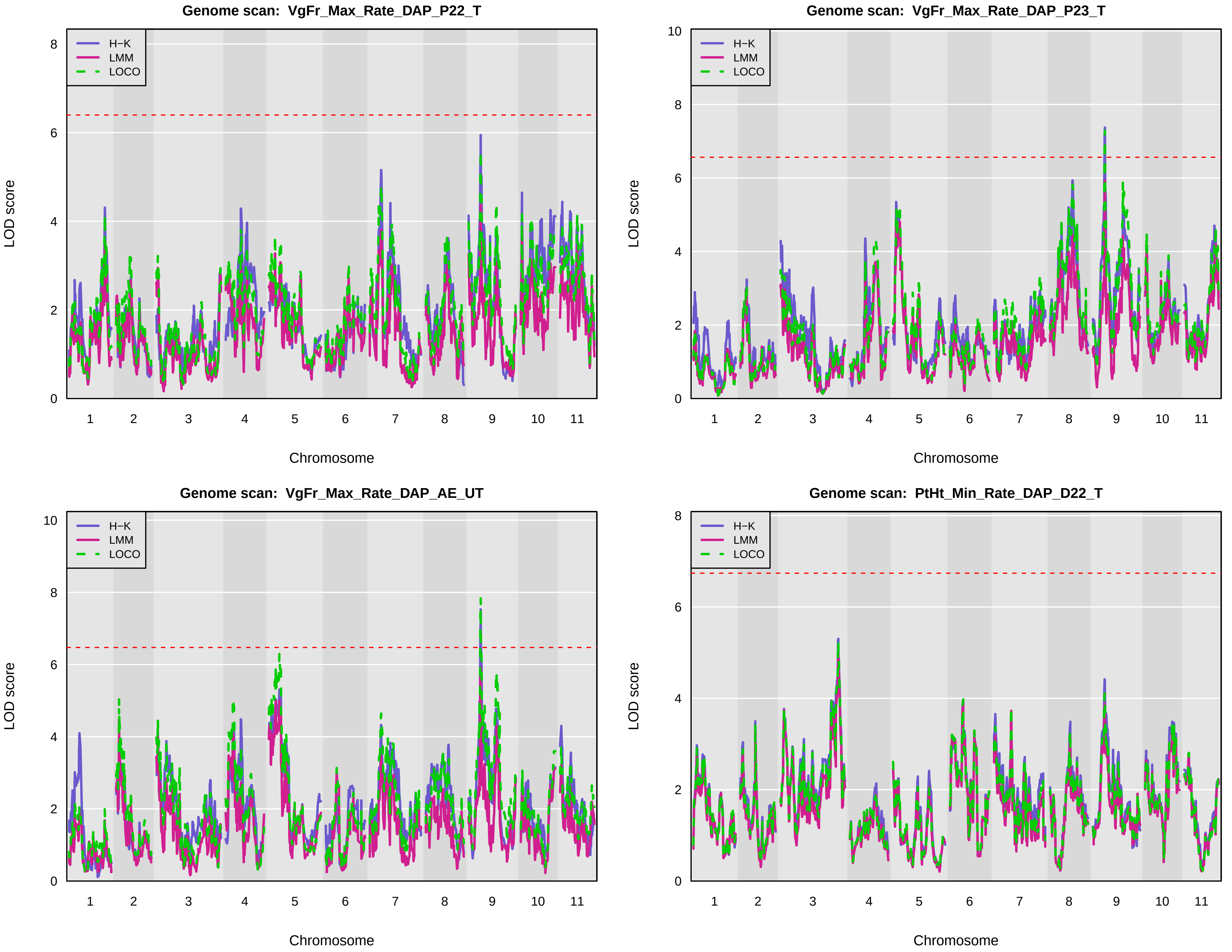
Genome scan: VgFr_Max_Rate_DAP_P22_T
Genome scan: VgFr_Max_Rate_DAP_P23_T
Genome scan: VgFr_Max_Rate_DAP_AE_UT
Genome scan: PtHt_Min_Rate_DAP_D22_T
H−K
LMM
LOCO
LOD score
Chromosome
1 2 3 4 5 6 7 8 9 10 11
0 2 4 6 8 10

Genome scan:  PtHt_Min_Rate_DAP_D23_T

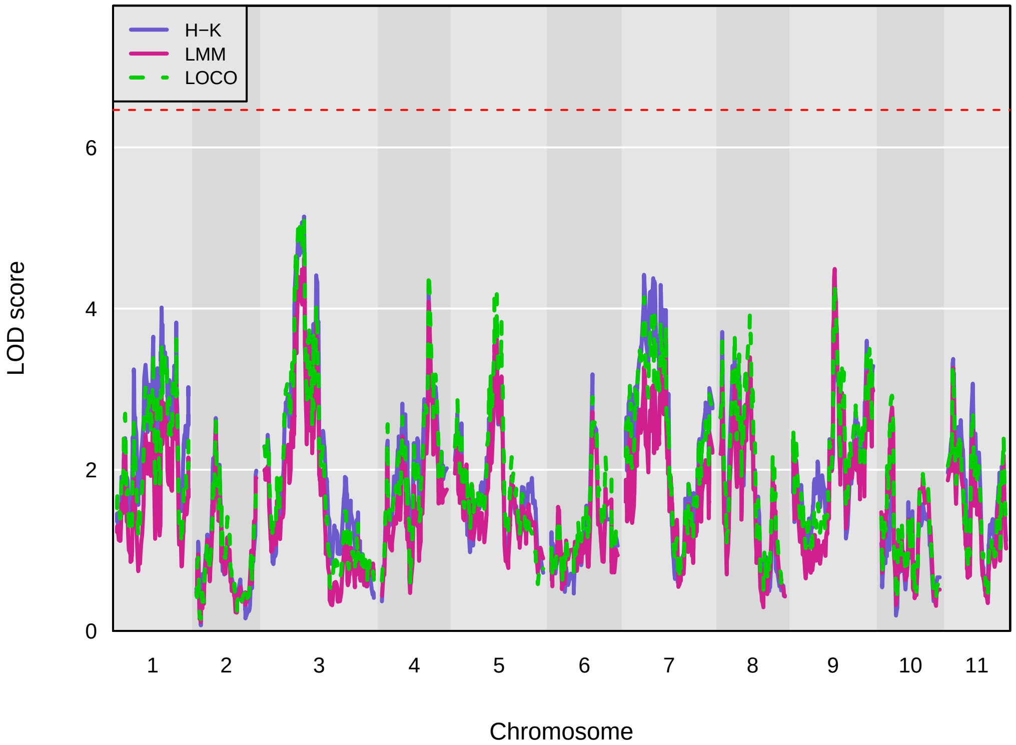


Genome scan:  PtHt_Min_Rate_DAP_P22_T

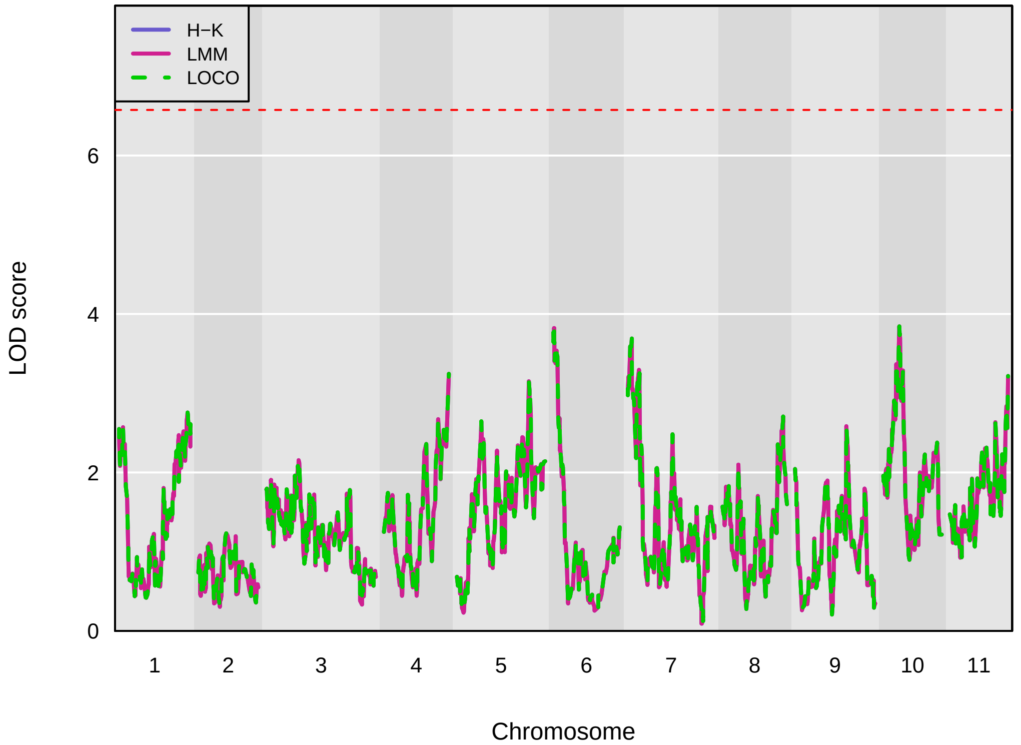


Genome scan:  PtHt_Min_Rate_DAP_P23_T

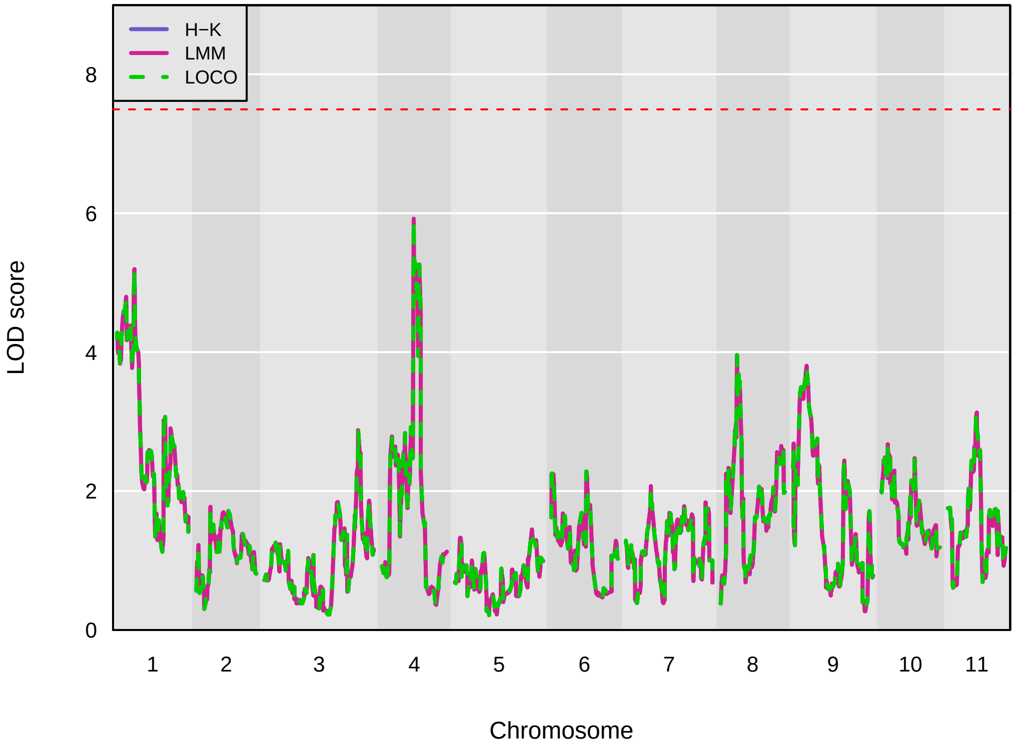


Genome scan:  PtHt_Min_Rate_DAP_AE_T

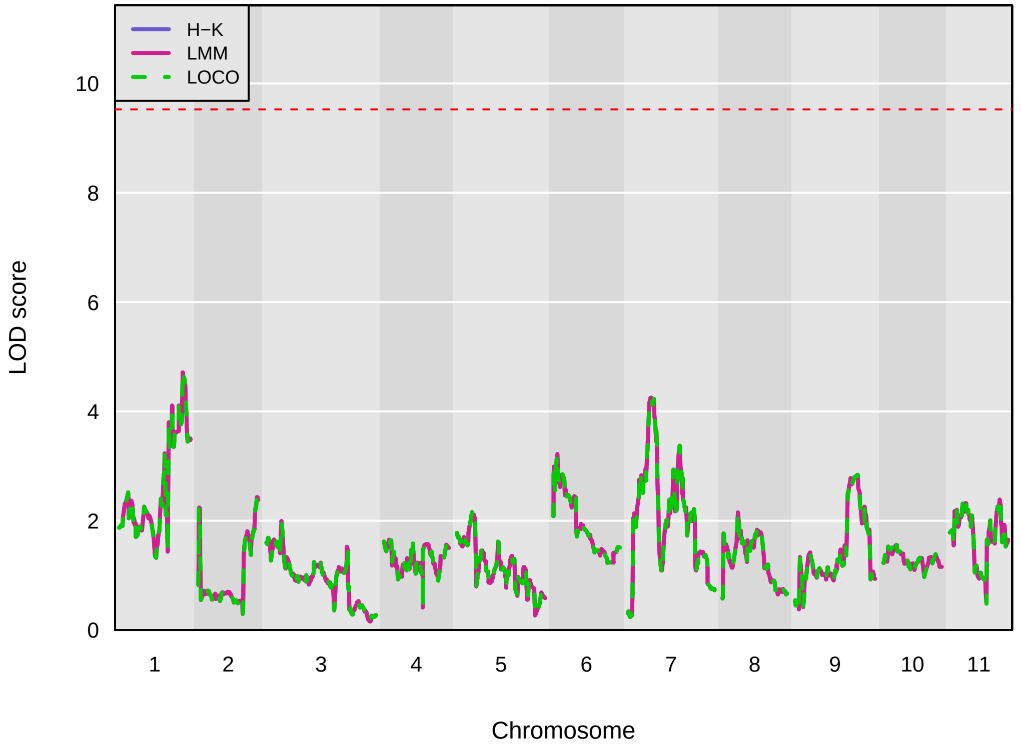

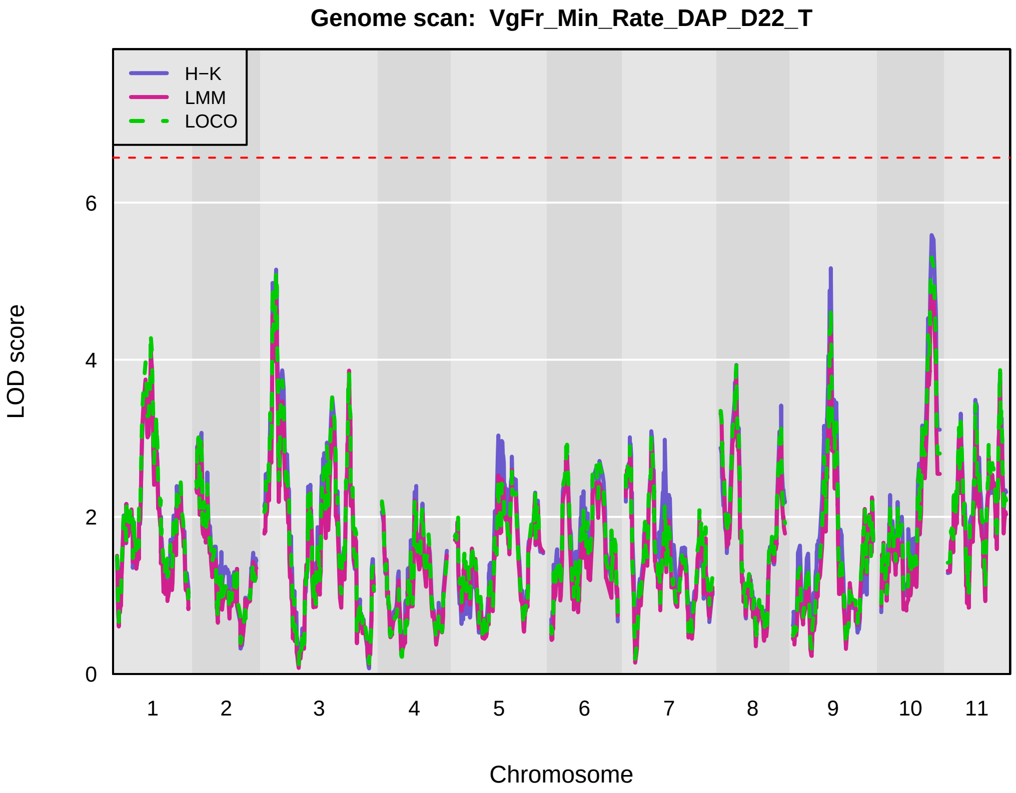
Genome scan: VgFr_Min_Rate_DAP_D22_T
H−K
LMM
LOCO
LOD score
0
2
4
6
1 2 3 4 5 6 7 8 9 10 11
Chromosome

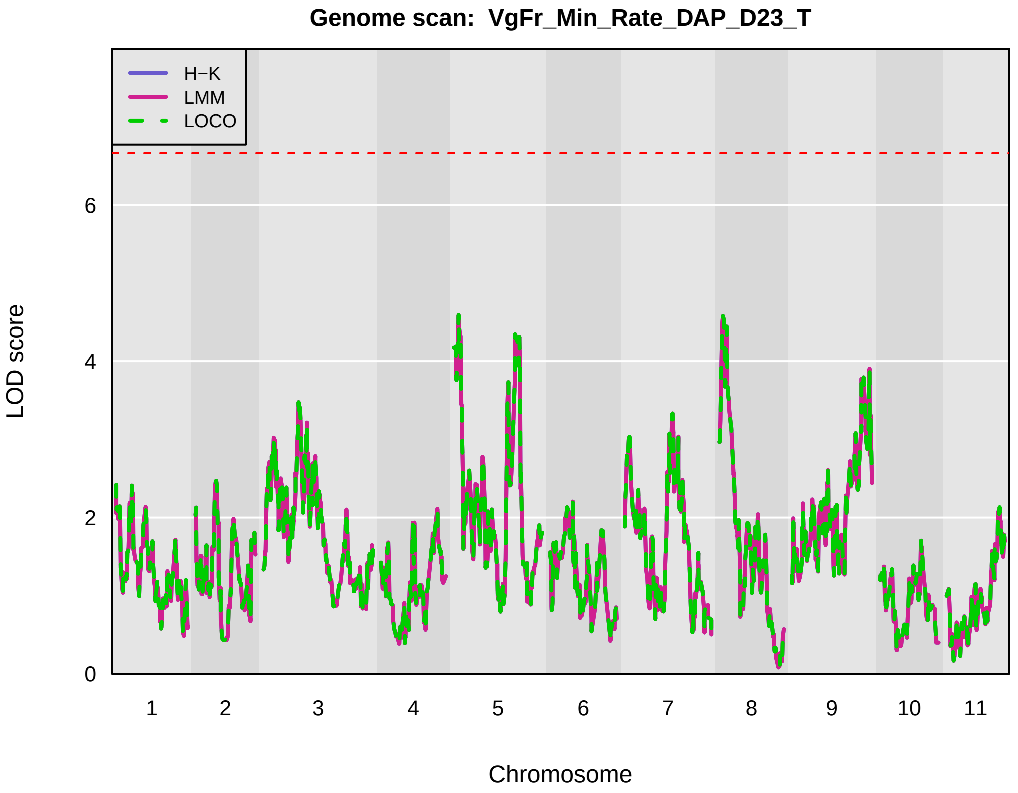
Genome scan: VgFr_Min_Rate_DAP_D23_T
H−K
LMM
LOCO
LOD score
0
2
4
6
1 2 3 4 5 6 7 8 9 10 11
Chromosome

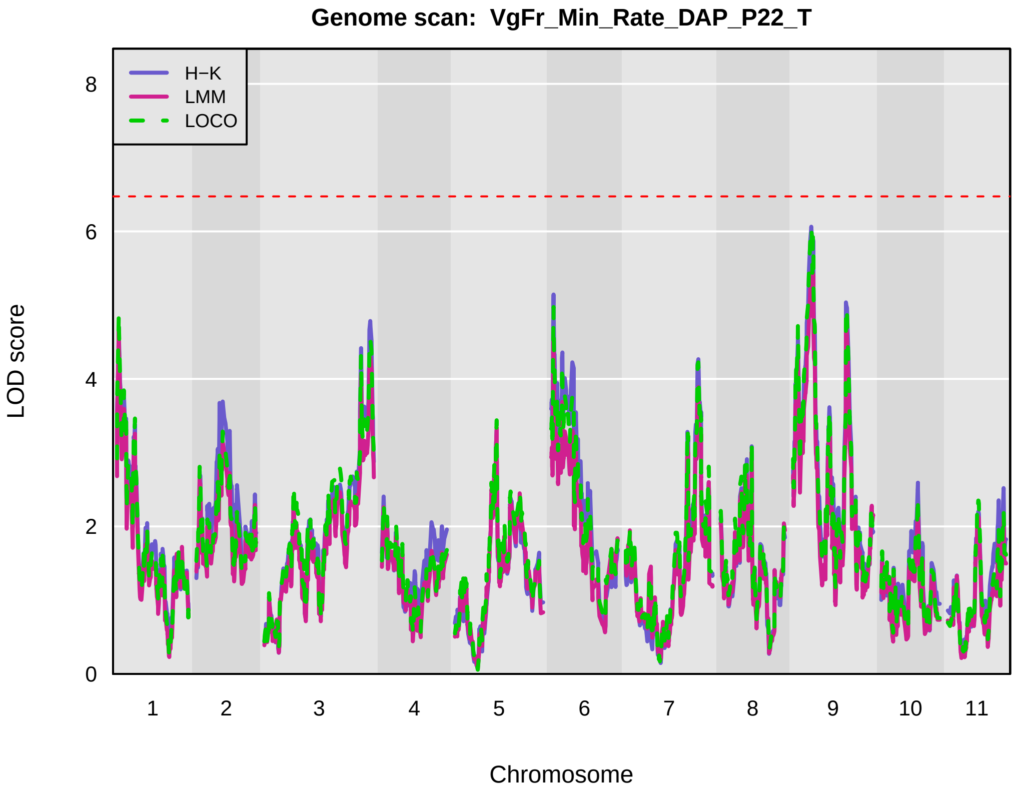
Genome scan: VgFr_Min_Rate_DAP_P22_T
H−K
LMM
LOCO
LOD score
0
2
4
6
8
1 2 3 4 5 6 7 8 9 10 11
Chromosome

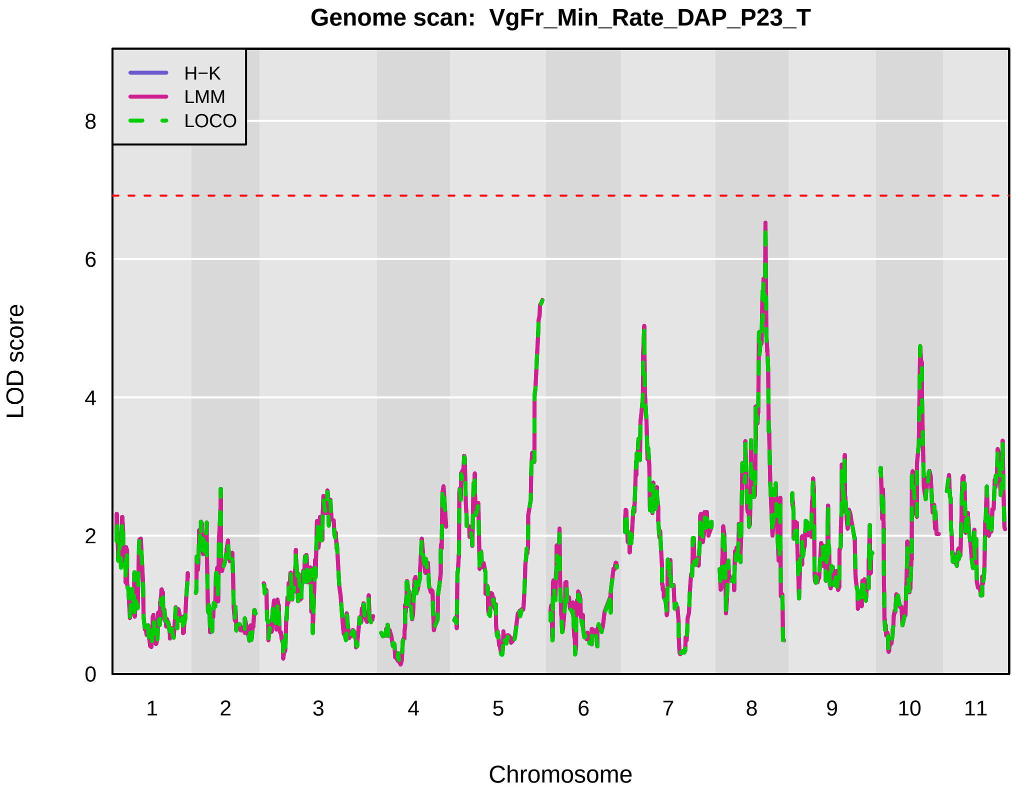
Genome scan: VgFr_Min_Rate_DAP_P23_T
H−K
LMM
LOCO
LOD score
0
2
4
6
8
1 2 3 4 5 6 7 8 9 10 11
Chromosome

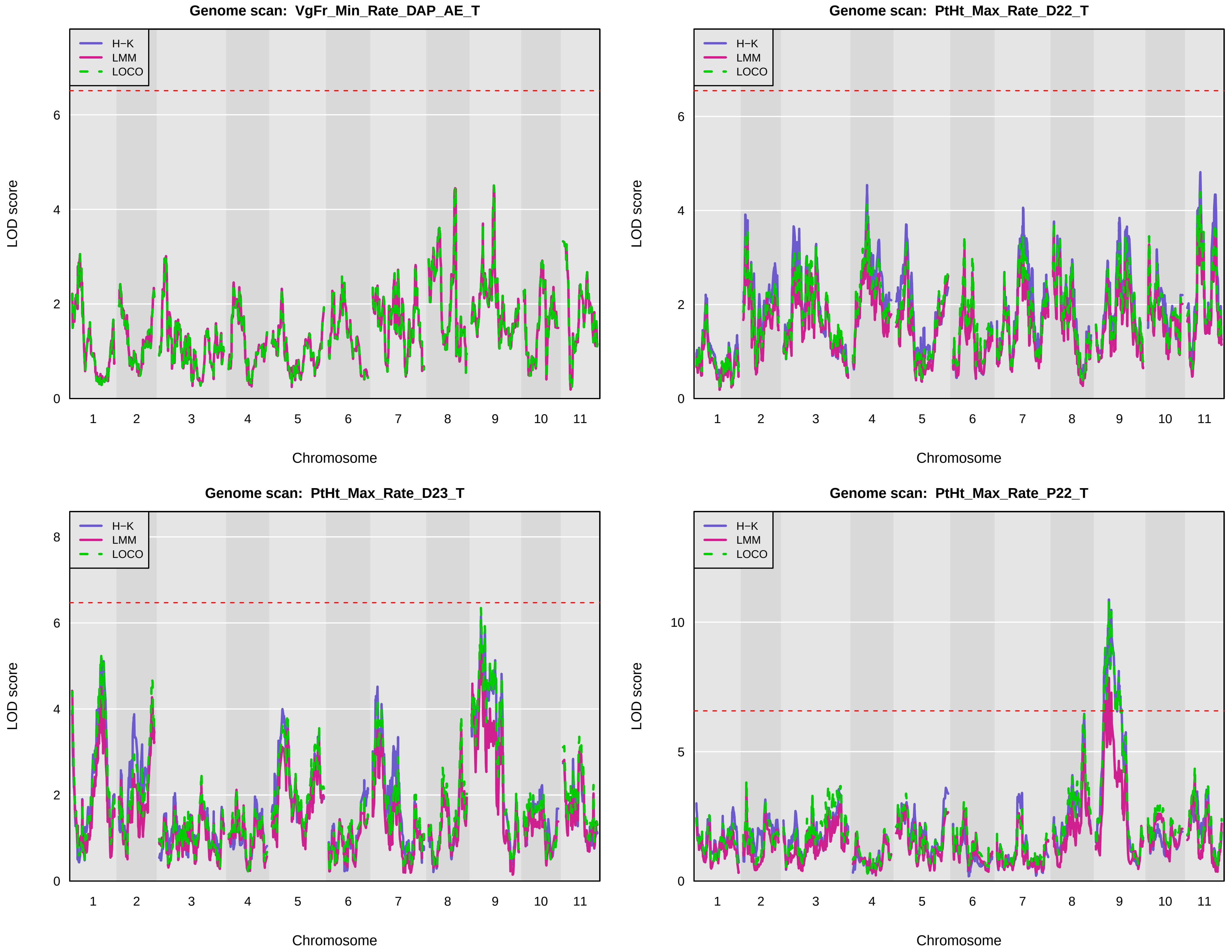
Genome scan: VgFr_Min_Rate_DAP_AE_T
Genome scan: PtHt_Max_Rate_D22_T
Genome scan: PtHt_Max_Rate_D23_T
Genome scan: PtHt_Max_Rate_P22_T
H−K
LMM
LOCO
LOD score
Chromosome
1 2 3 4 5 6 7 8 9 10 11

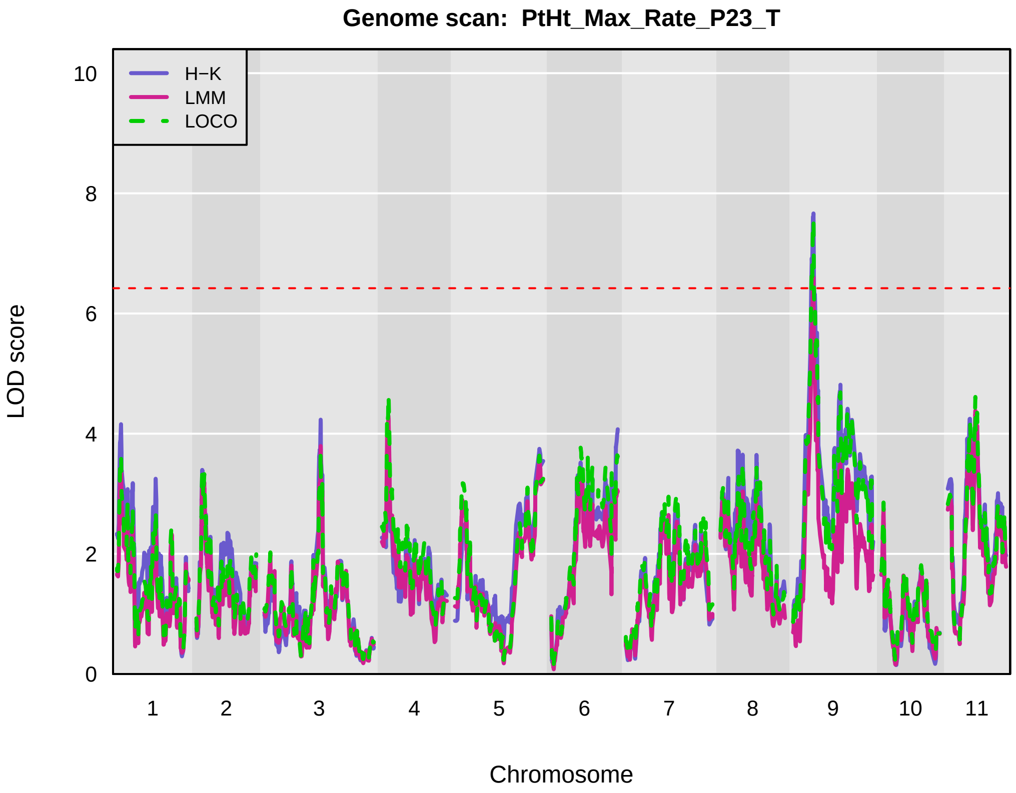
Genome scan: PtHt_Max_Rate_P23_T
H−K
LMM
LOCO
LOD score
Chromosome

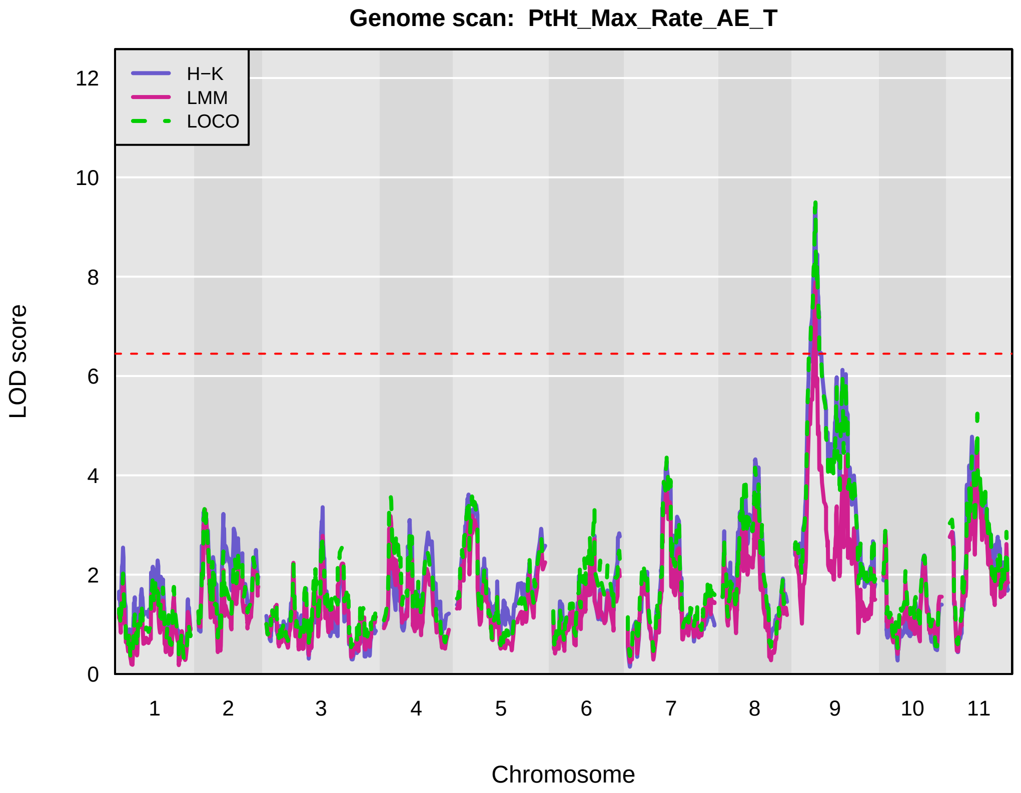
Genome scan: PtHt_Max_Rate_AE_T
H−K
LMM
LOCO
LOD score
Chromosome

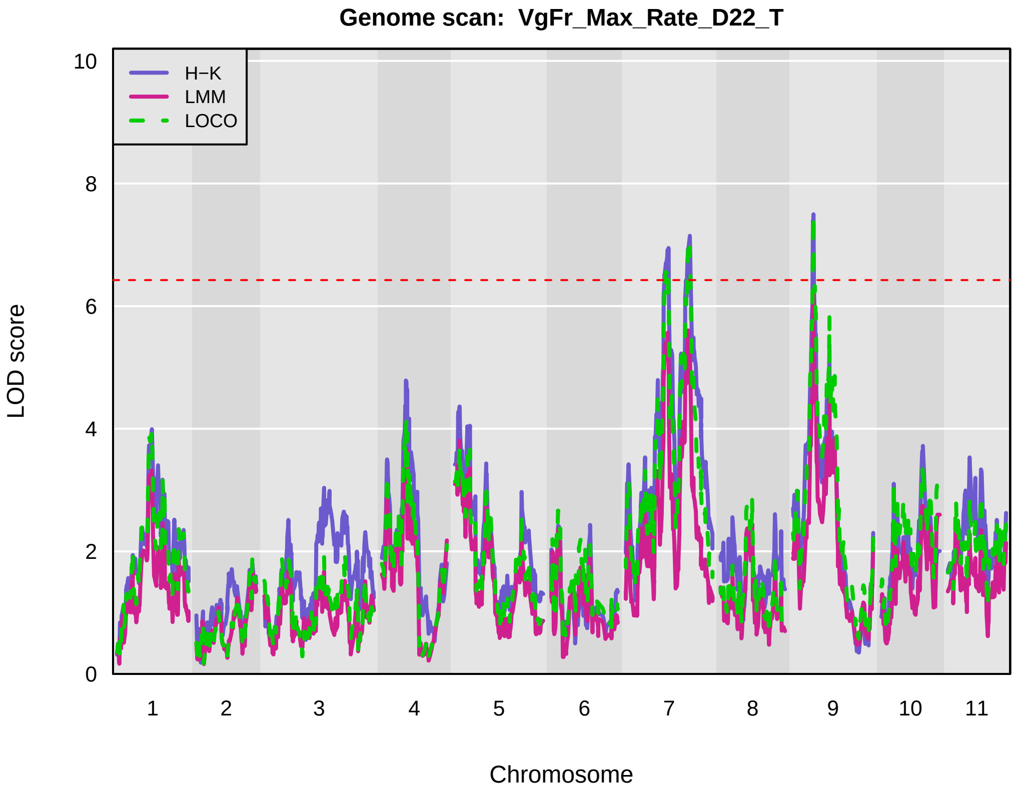
Genome scan: VgFr_Max_Rate_D22_T
H−K
LMM
LOCO
LOD score
Chromosome

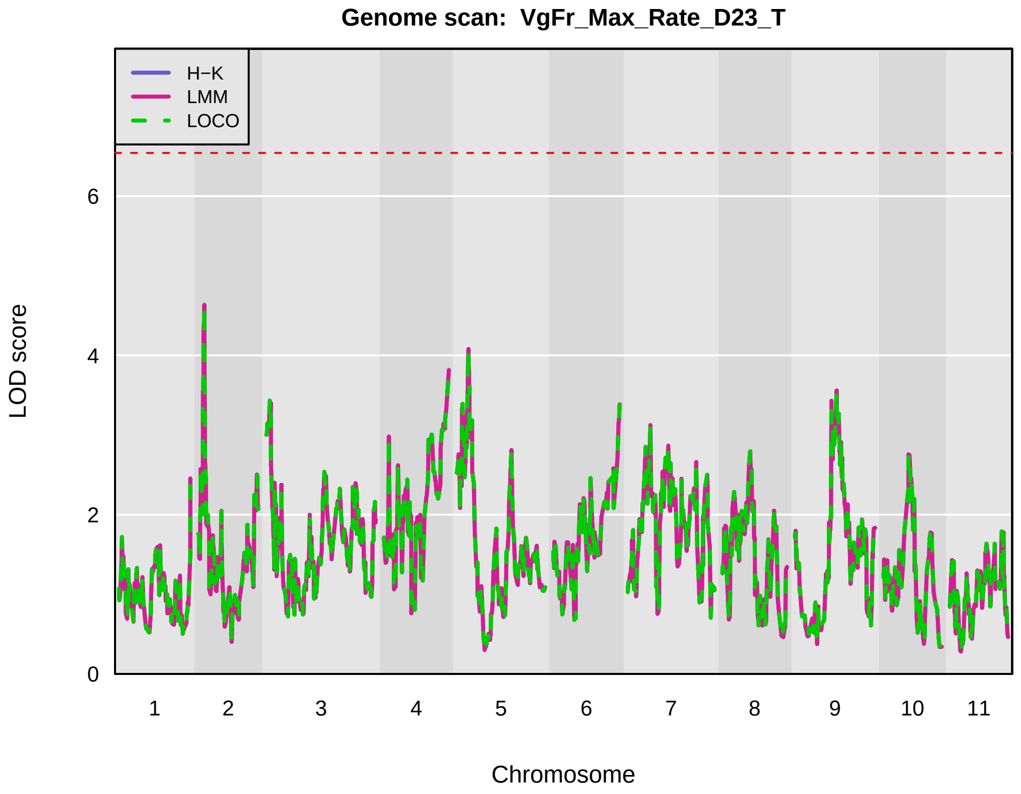
Genome scan: VgFr_Max_Rate_D23_T
H−K
LMM
LOCO
LOD score
Chromosome

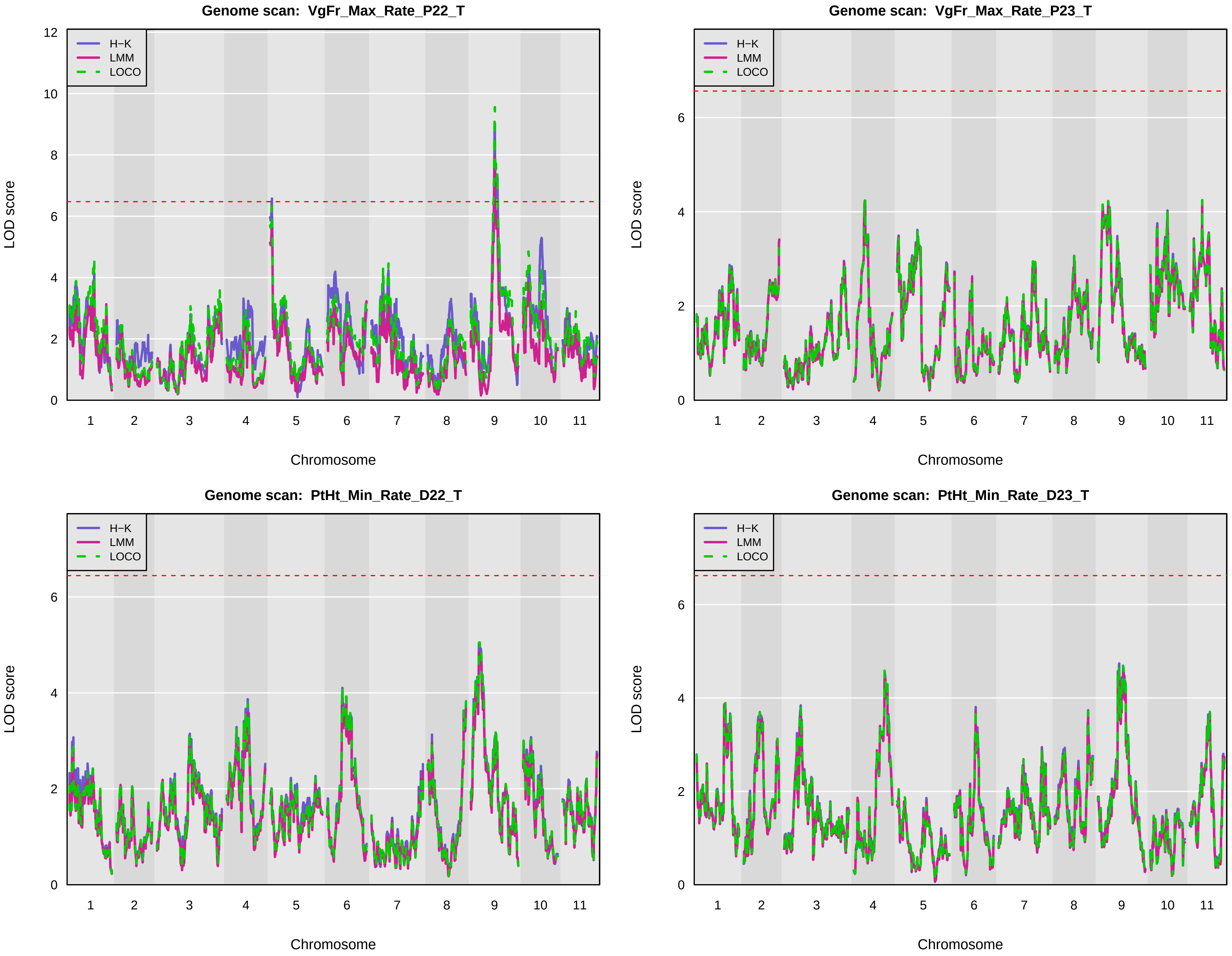
Genome scan: VgFr_Max_Rate_P22_T
Genome scan: VgFr_Max_Rate_P23_T
Genome scan: PtHt_Min_Rate_D22_T
Genome scan: PtHt_Min_Rate_D23_T
H−K
LMM
LOCO
LOD score
Chromosome
0
2
4
6
8
10
12
1
2
3
4
5
6
7
8
9
10
11

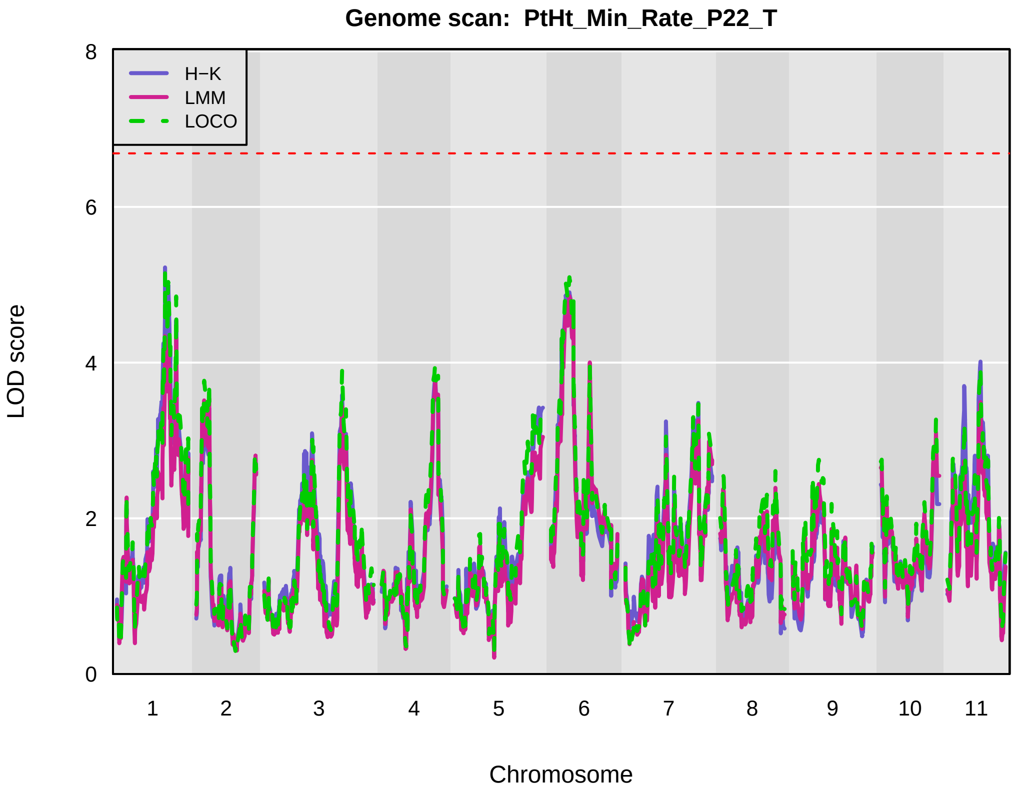
Genome scan: PtHt_Min_Rate_P22_T
H−K
LMM
LOCO
LOD score
Chromosome
1 2 3 4 5 6 7 8 9 10 11
0 2 4 6 8

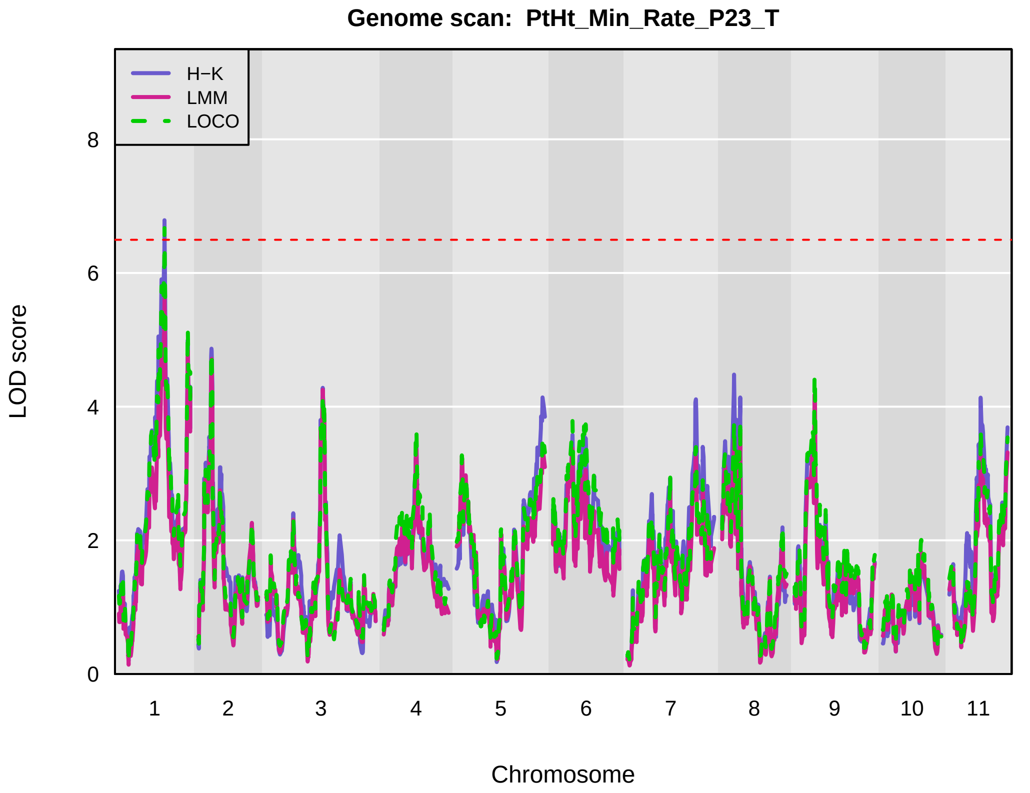
Genome scan: PtHt_Min_Rate_P23_T
H−K
LMM
LOCO
LOD score
Chromosome
1 2 3 4 5 6 7 8 9 10 11
0 2 4 6 8

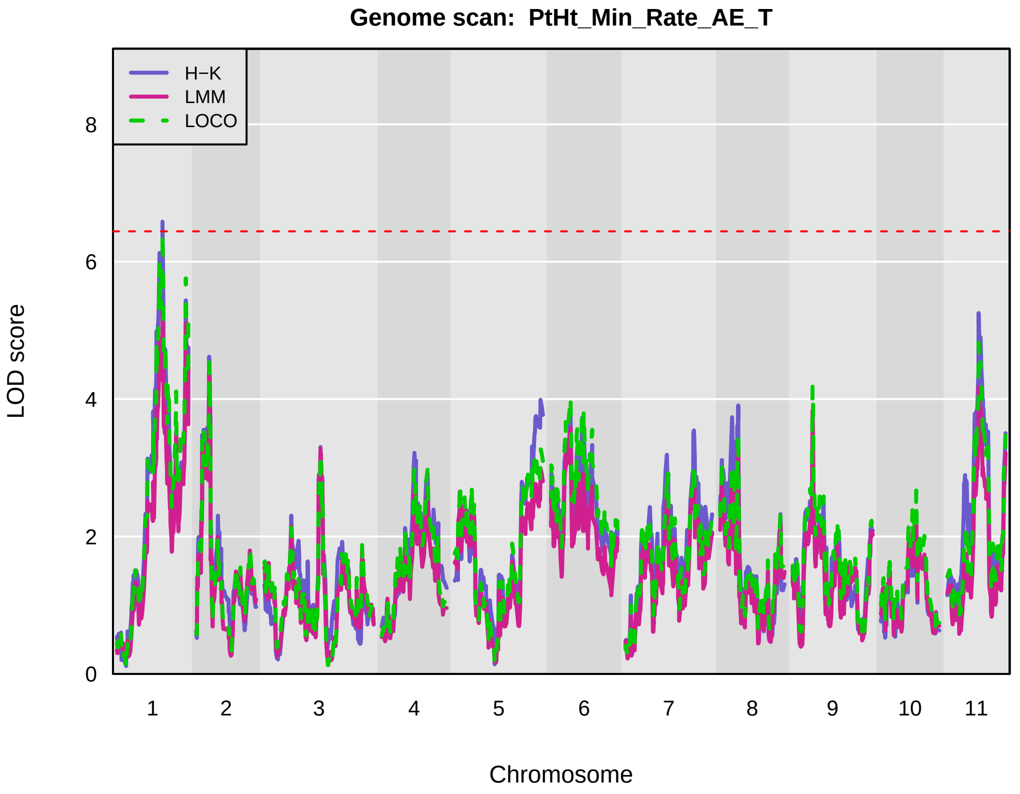
Genome scan: PtHt_Min_Rate_AE_T
H−K
LMM
LOCO
LOD score
Chromosome
1 2 3 4 5 6 7 8 9 10 11
0 2 4 6 8

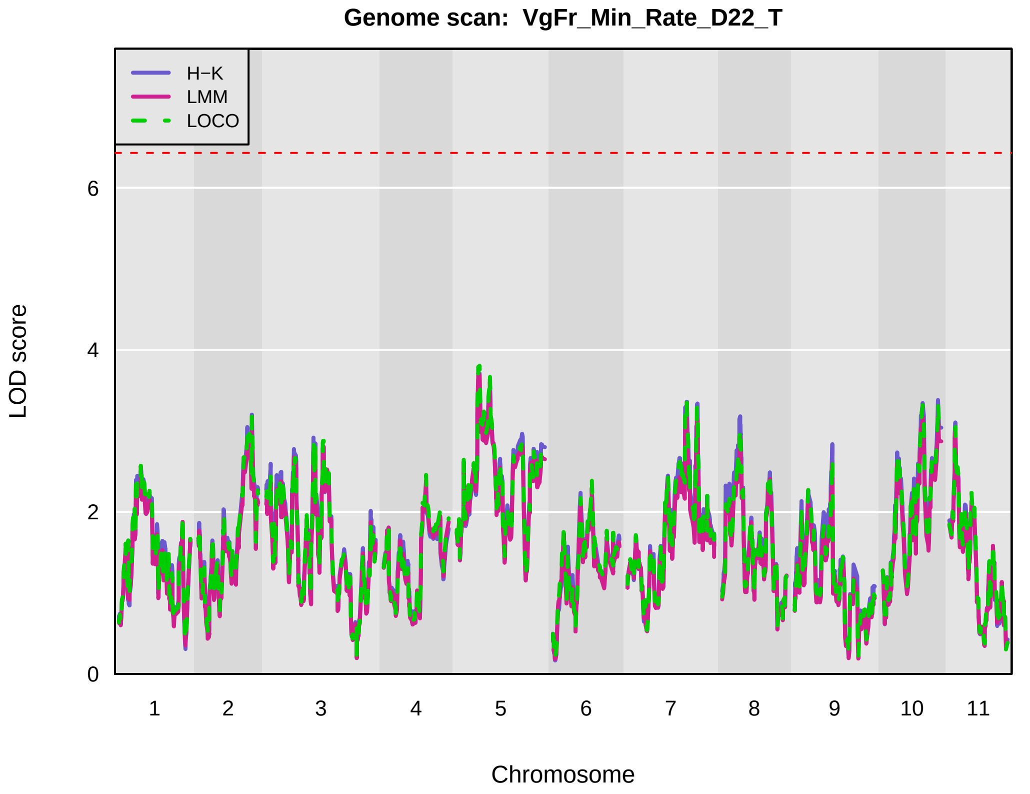
Genome scan: VgFr_Min_Rate_D22_T
H−K
LMM
LOCO
LOD score
Chromosome
1 2 3 4 5 6 7 8 9 10 11
0 2 4 6

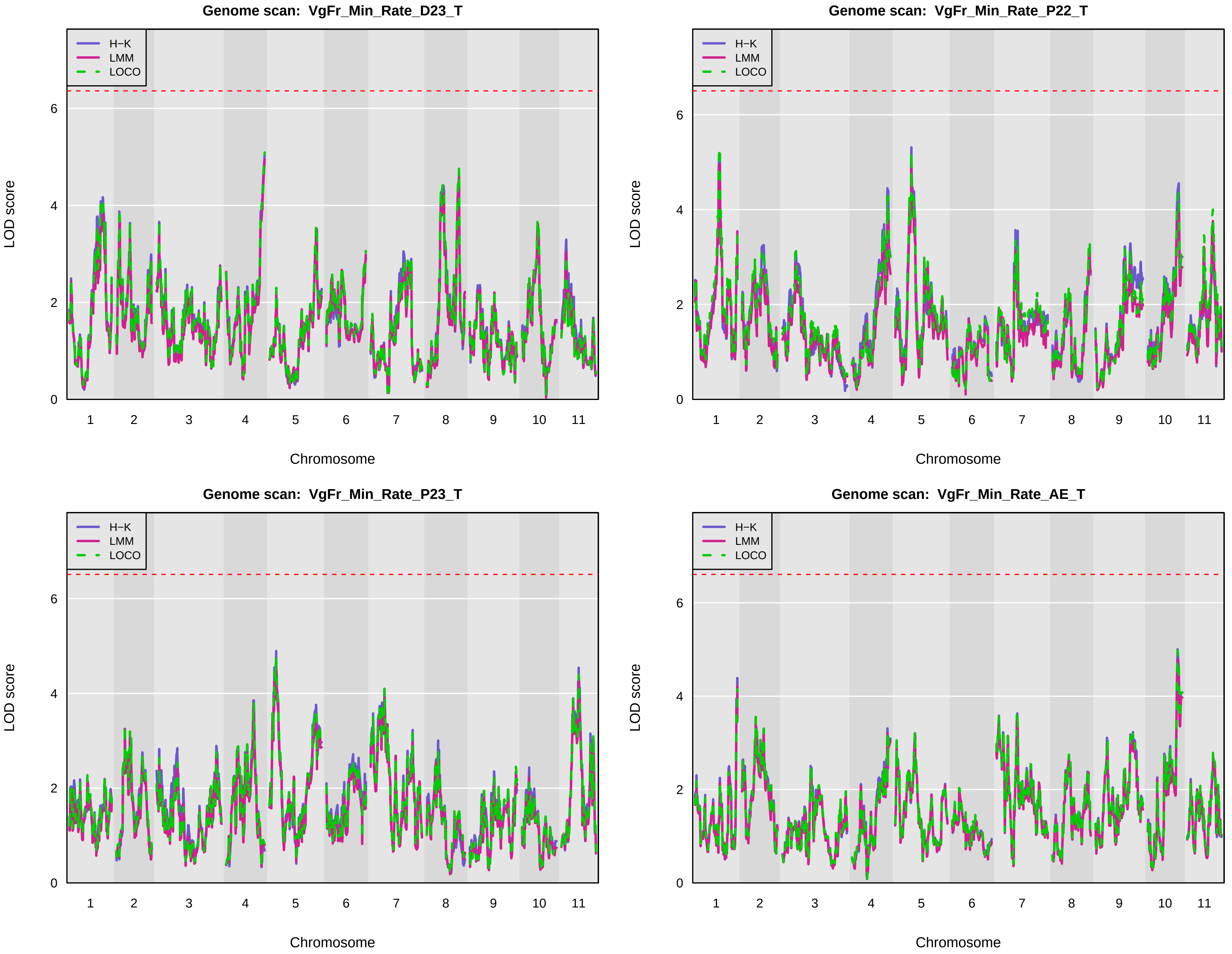
Genome scan: VgFr_Min_Rate_D23_T
Genome scan: VgFr_Min_Rate_P22_T
Genome scan: VgFr_Min_Rate_P23_T
Genome scan: VgFr_Min_Rate_AE_T
H−K
LMM
LOCO
LOD score
Chromosome
0
2
4
6
1
2
3
4
5
6
7
8
9
10
11

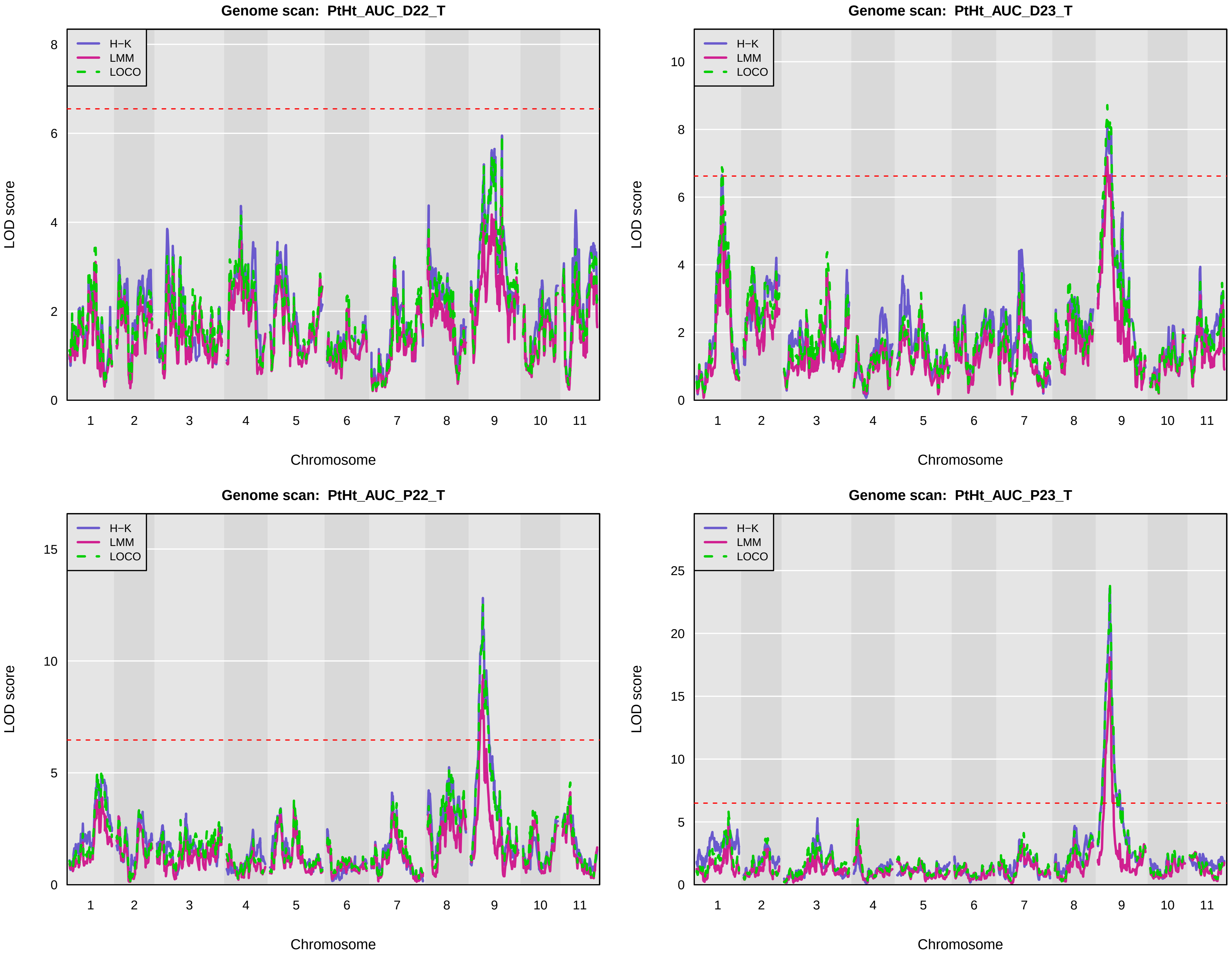

Genome scan: PtHt_AUC_D22_T
Genome scan: PtHt_AUC_D23_T
Genome scan: PtHt_AUC_P22_T
Genome scan: PtHt_AUC_P23_T
H−K
LMM
LOCO
LOD score
Chromosome
1 2 3 4 5 6 7 8 9 10 11

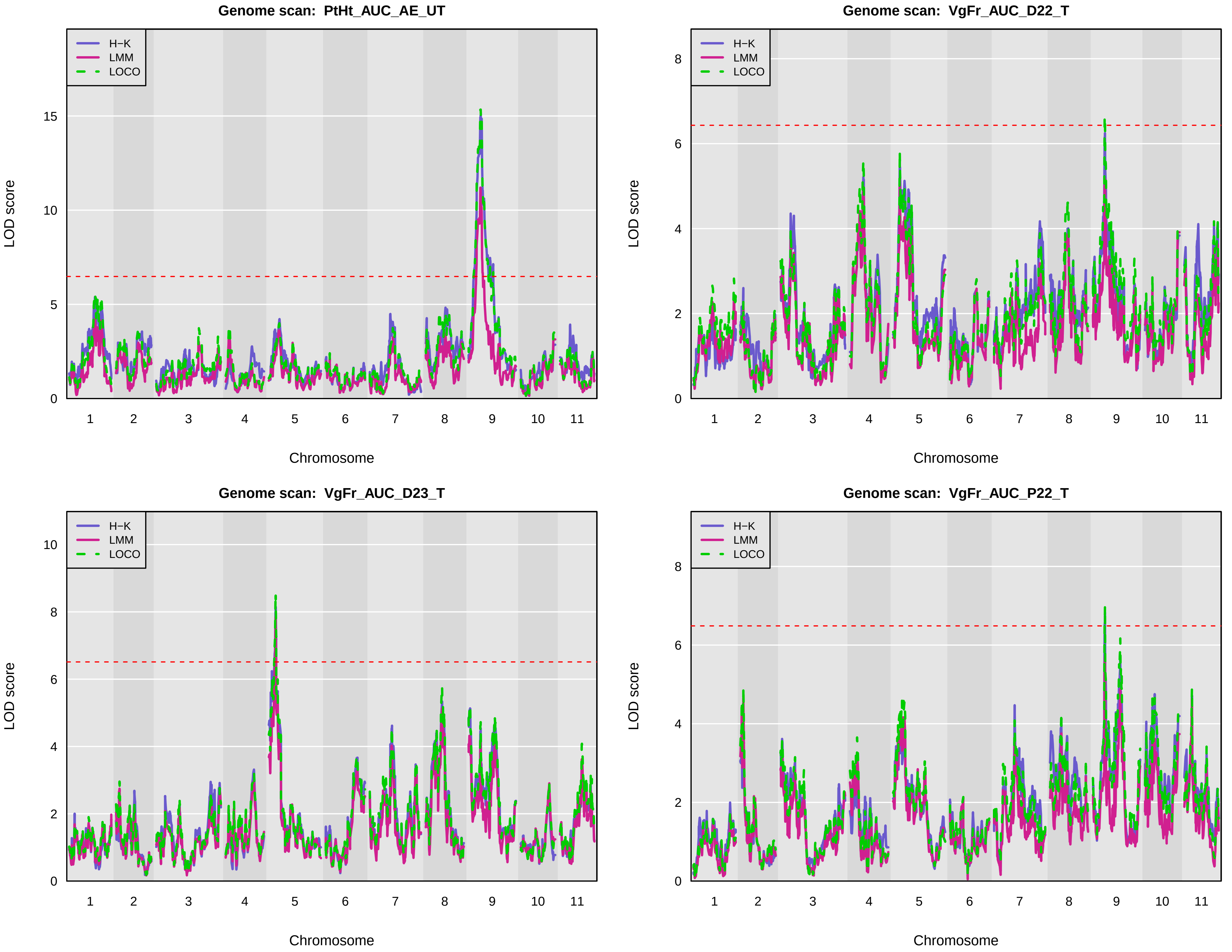

Genome scan: PtHt_AUC_AE_UT
H−K
LMM
LOCO
LOD score
0
5
10
15
1
2
3
4
5
6
7
8
9
10
11
Chromosome
Genome scan: VgFr_AUC_D22_T
H−K
LMM
LOCO
LOD score
0
2
4
6
8
1
2
3
4
5
6
7
8
9
10
11
Chromosome
Genome scan: VgFr_AUC_D23_T
H−K
LMM
LOCO
LOD score
0
2
4
6
8
10
1
2
3
4
5
6
7
8
9
10
11
Chromosome
Genome scan: VgFr_AUC_P22_T
H−K
LMM
LOCO
LOD score
0
2
4
6
8
1
2
3
4
5
6
7
8
9
10
11
Chromosome

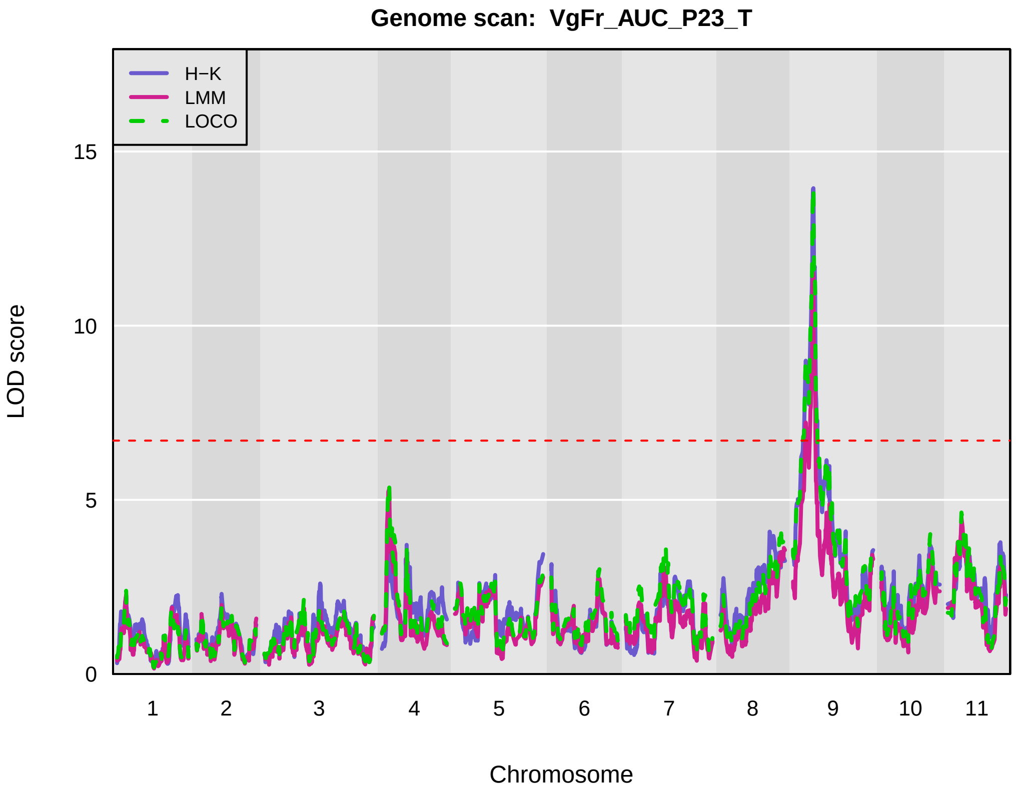
Genome scan: VgFr_AUC_P23_T
H−K
LMM
LOCO
LOD score
0
5
10
15
1
2
3
4
5
6
7
8
9
10
11
Chromosome

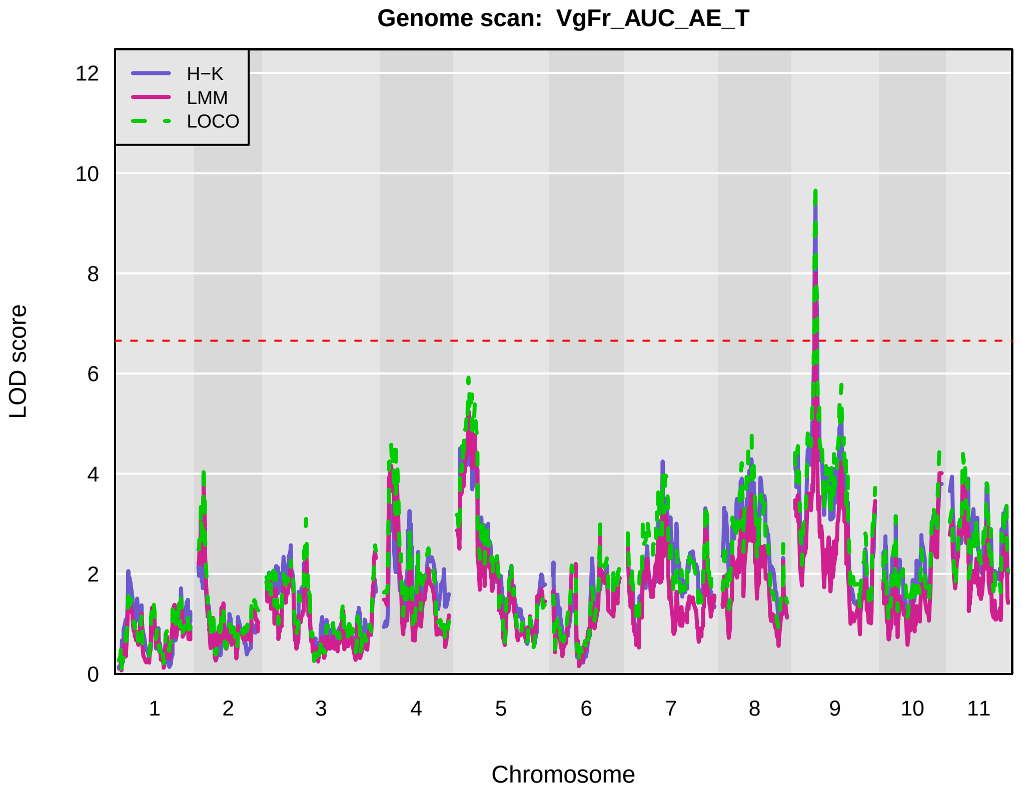
Genome scan: VgFr_AUC_AE_T
H−K
LMM
LOCO
LOD score
0
2
4
6
8
10
12
1
2
3
4
5
6
7
8
9
10
11
Chromosome

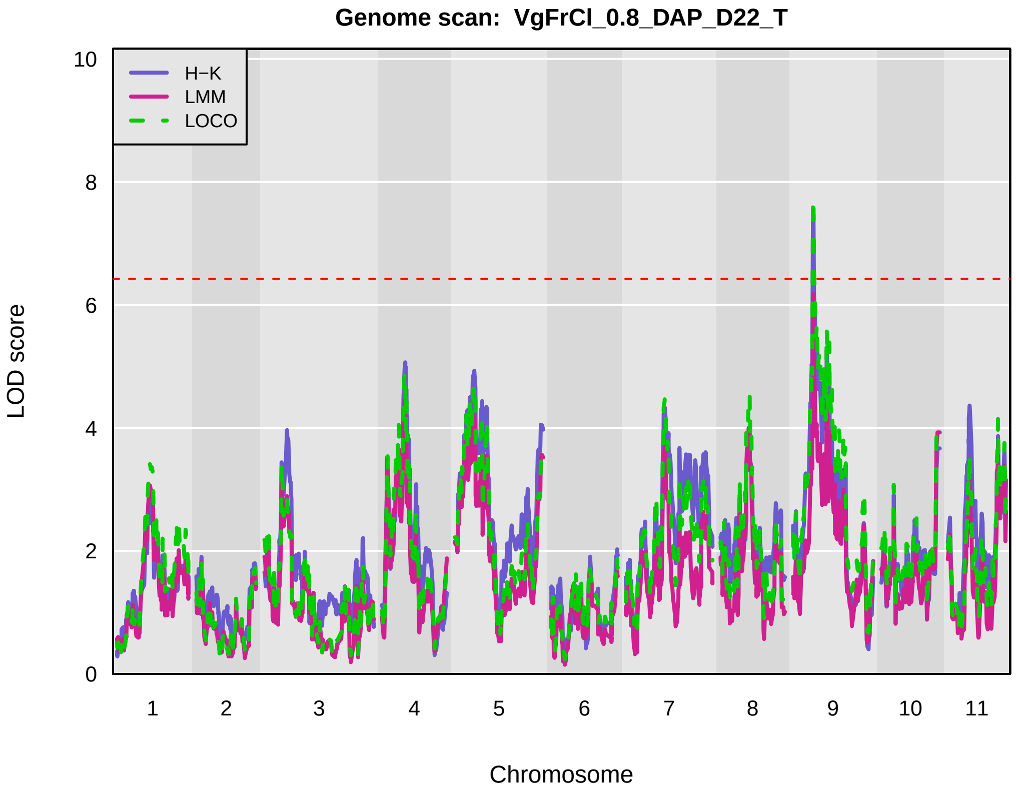
Genome scan: VgFrCl_0.8_DAP_D22_T
H−K
LMM
LOCO
LOD score
0
2
4
6
8
10
1
2
3
4
5
6
7
8
9
10
11
Chromosome

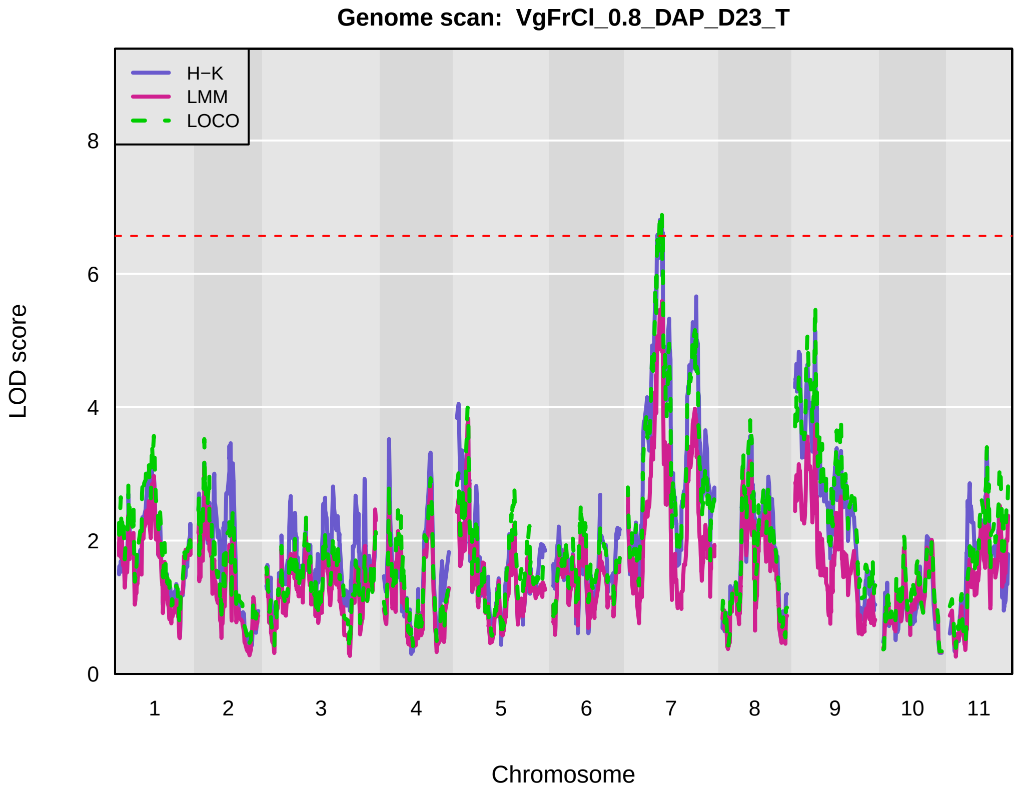
Genome scan: VgFrCl_0.8_DAP_D23_T
H−K
LMM
LOCO
LOD score
0
2
4
6
8
1
2
3
4
5
6
7
8
9
10
11
Chromosome

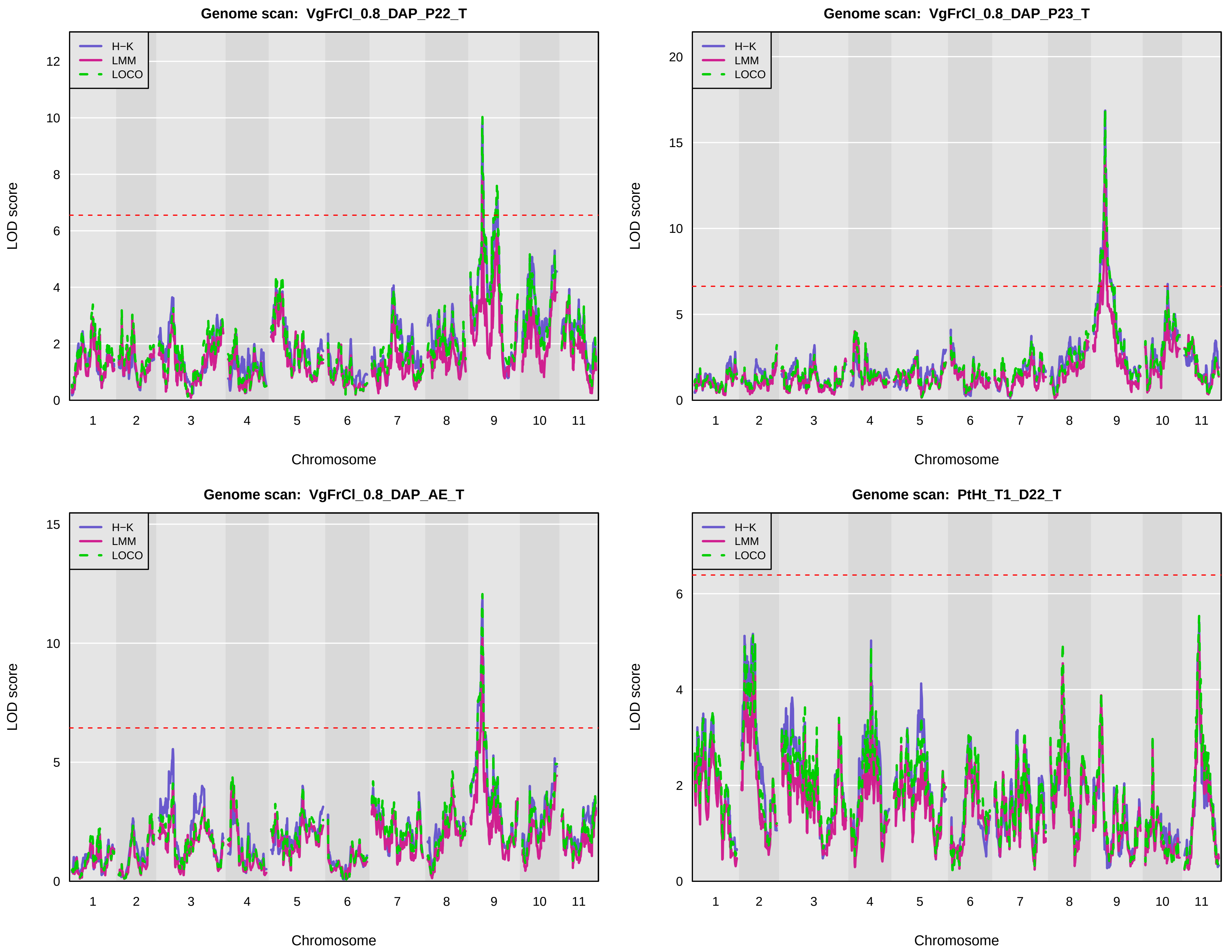

Genome scan: VgFrCl_0.8_DAP_P22_T
Genome scan: VgFrCl_0.8_DAP_P23_T
Genome scan: VgFrCl_0.8_DAP_AE_T
Genome scan: PtHt_T1_D22_T
H−K
LMM
LOCO
LOD score
Chromosome
0
2
4
6
8
10
12
5
15
20
1
3
7
9
11

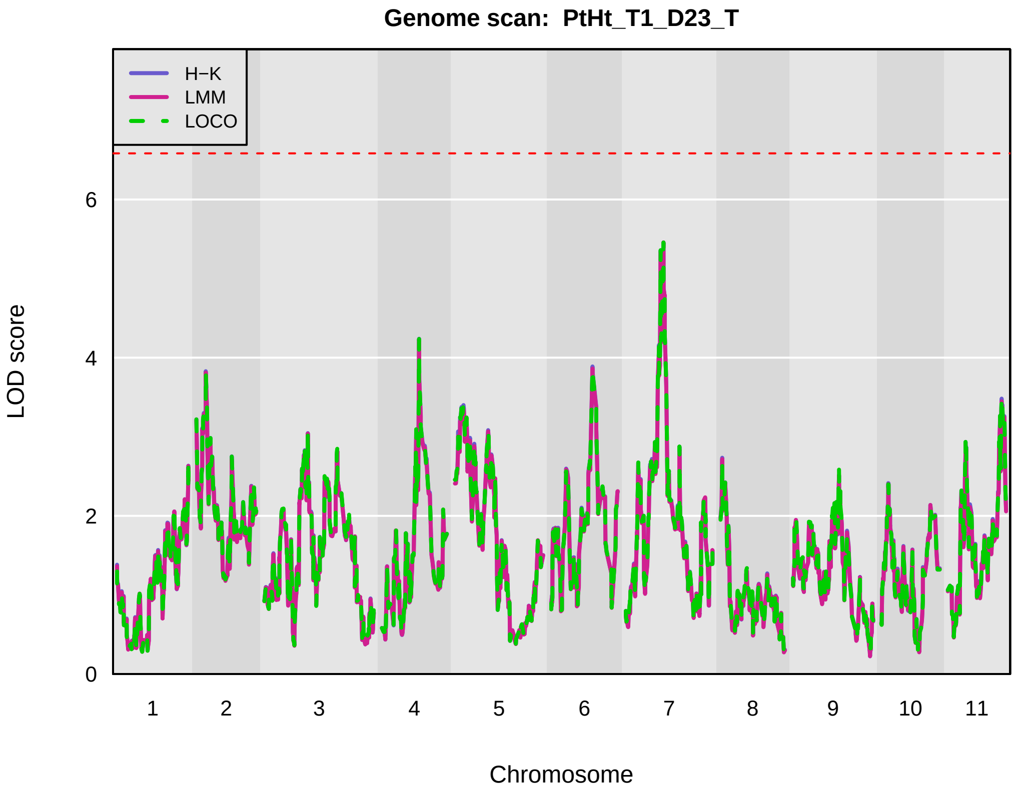
Genome scan: PtHt_T1_D23_T
H−K
LMM
LOCO
LOD score
0
2
4
6
1
2
3
4
5
6
7
8
9
10
11
Chromosome

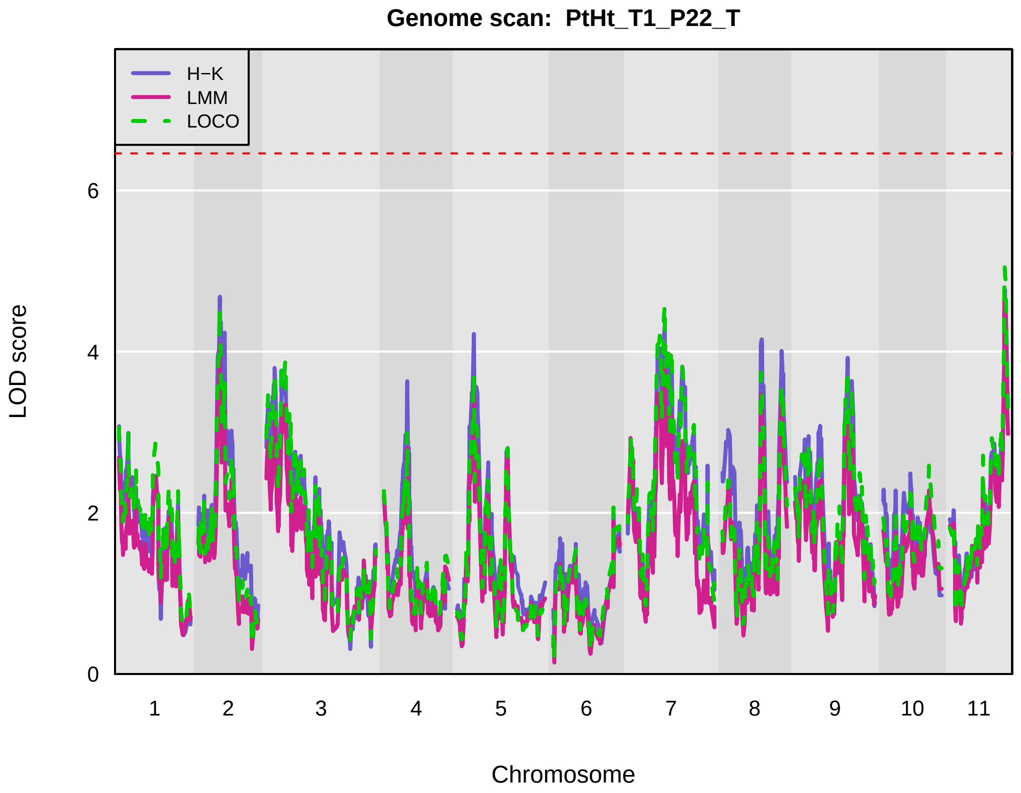
Genome scan: PtHt_T1_P22_T
H−K
LMM
LOCO
LOD score
0
2
4
6
1
2
3
4
5
6
7
8
9
10
11
Chromosome

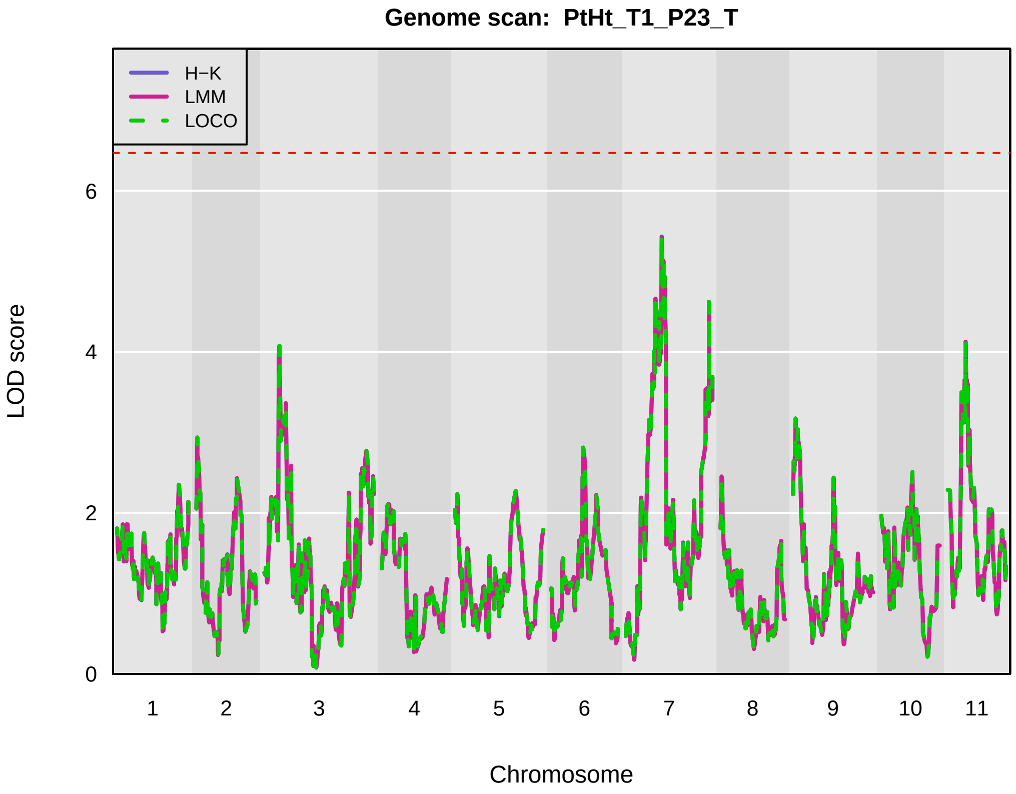
Genome scan: PtHt_T1_P23_T
H−K
LMM
LOCO
LOD score
0
2
4
6
1
2
3
4
5
6
7
8
9
10
11
Chromosome

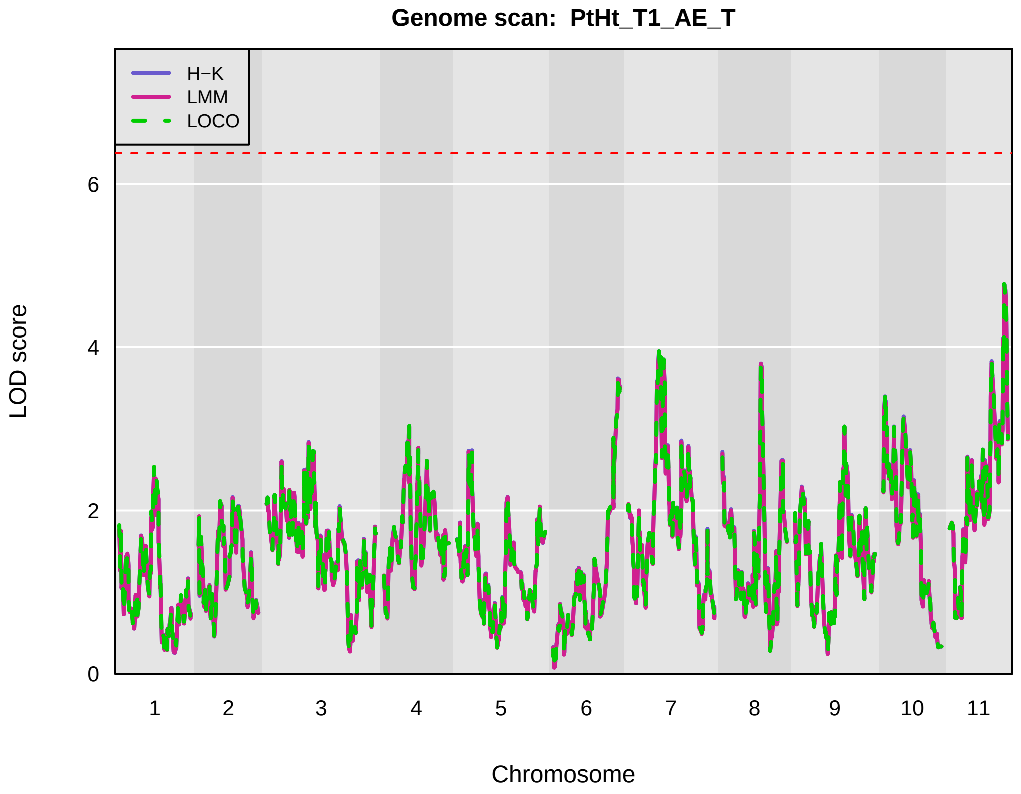
Genome scan: PtHt_T1_AE_T
H−K
LMM
LOCO
LOD score
0
2
4
6
1
2
3
4
5
6
7
8
9
10
11
Chromosome

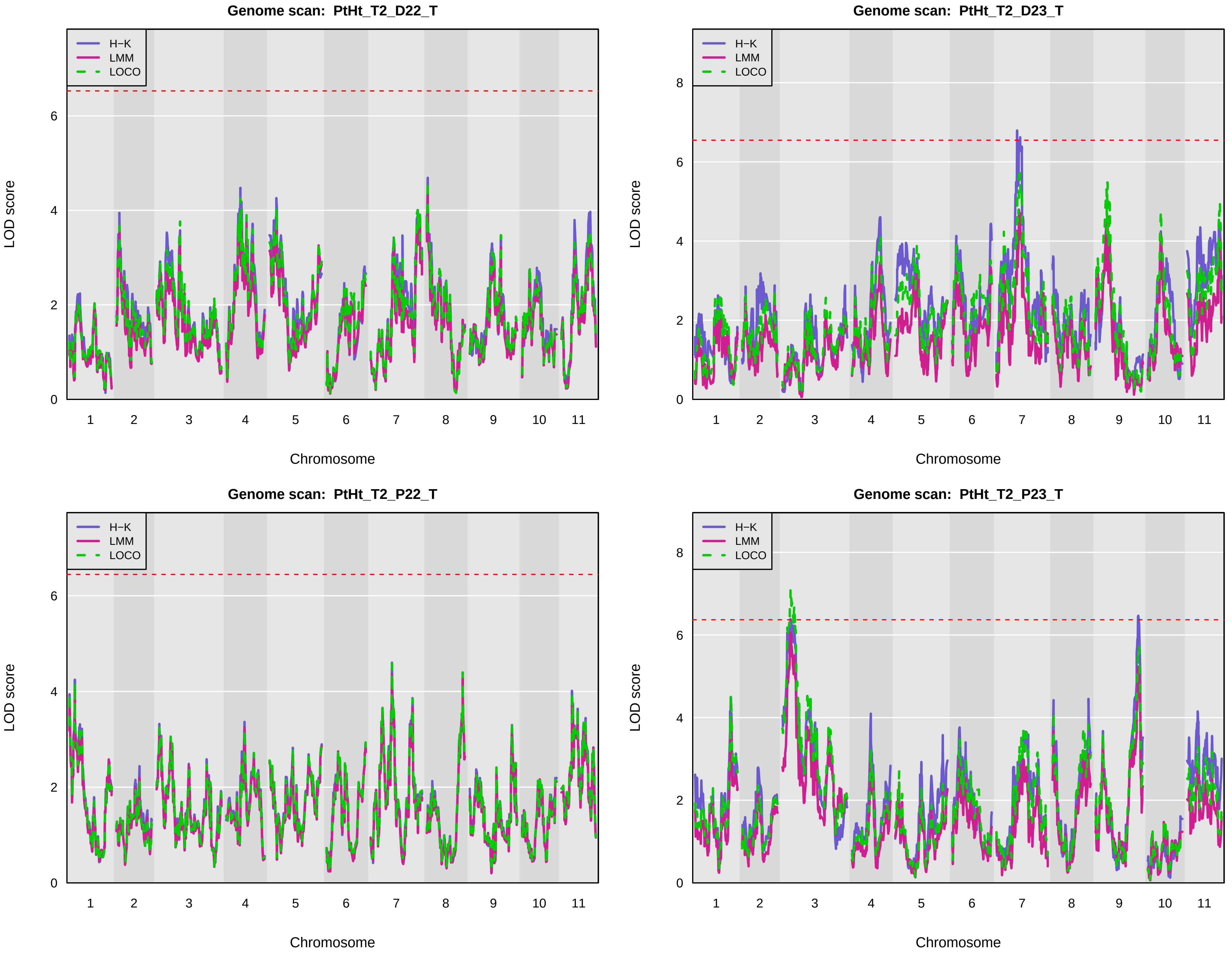
Genome scan: PtHt_T2_D22_T
Genome scan: PtHt_T2_D23_T
Genome scan: PtHt_T2_P22_T
Genome scan: PtHt_T2_P23_T
H−K
LMM
LOCO
LOD score
Chromosome
0
2
4
6
8
1
2
3
4
5
6
7
8
9
10
11

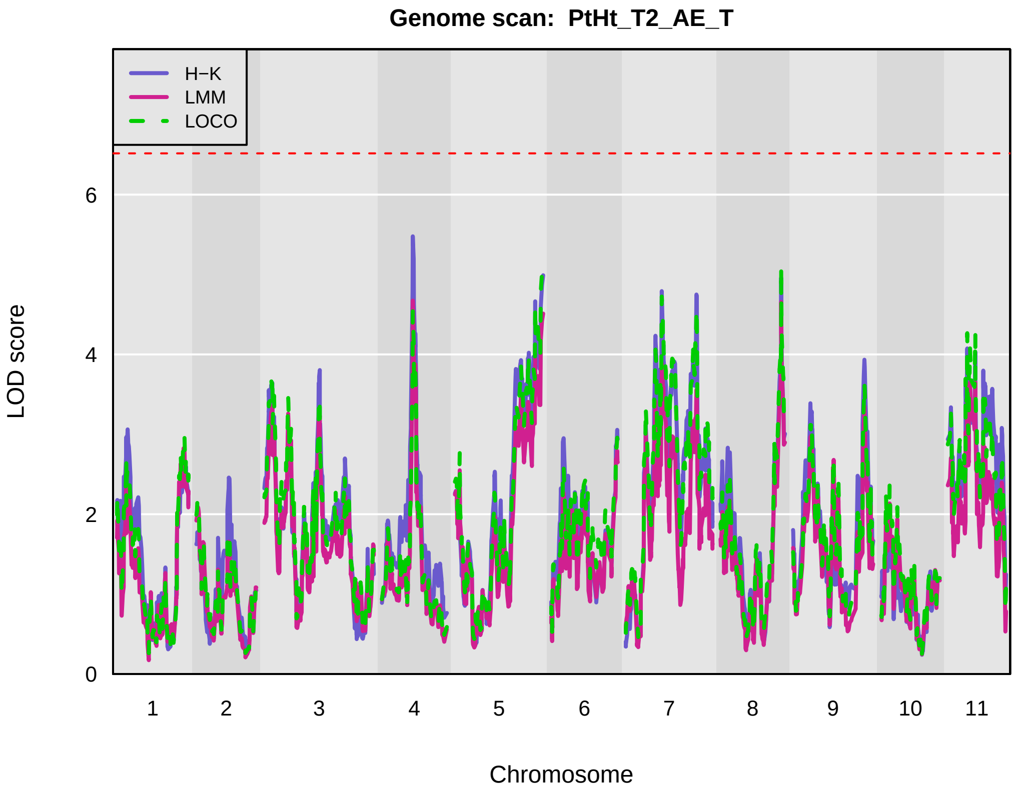
Genome scan: PtHt_T2_AE_T
H−K
LMM
LOCO
LOD score
0
2
4
6
1
2
3
4
5
6
7
8
9
10
11
Chromosome

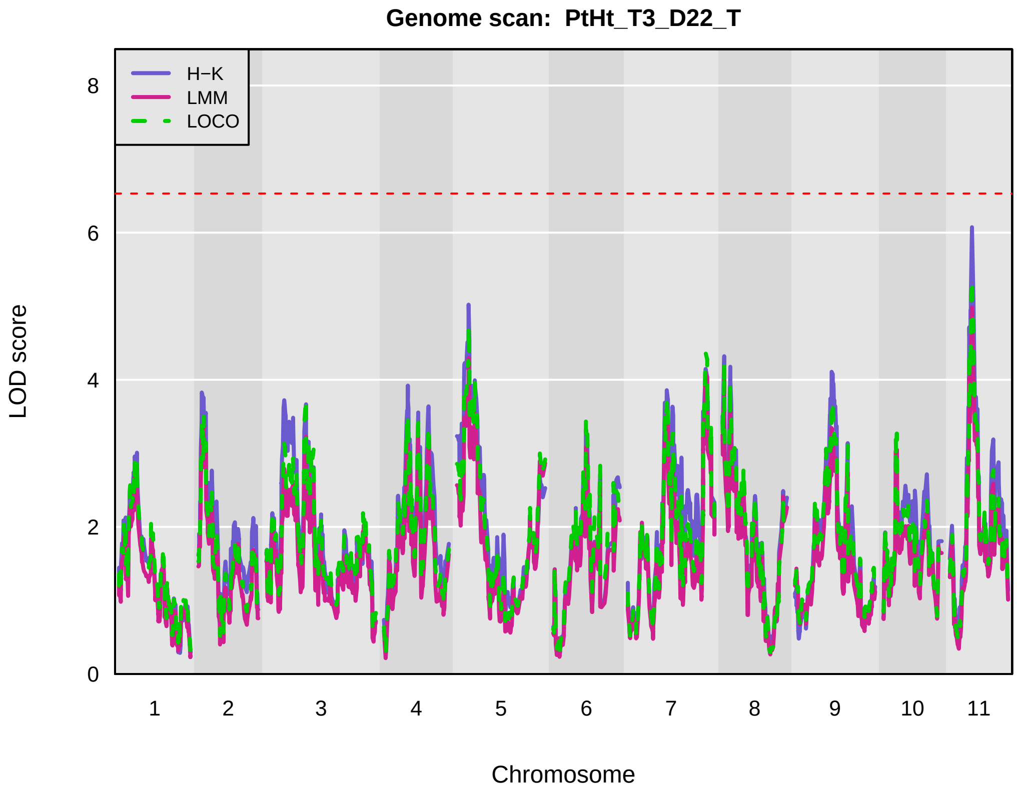
Genome scan: PtHt_T3_D22_T
H−K
LMM
LOCO
LOD score
0
2
4
6
8
1
2
3
4
5
6
7
8
9
10
11
Chromosome

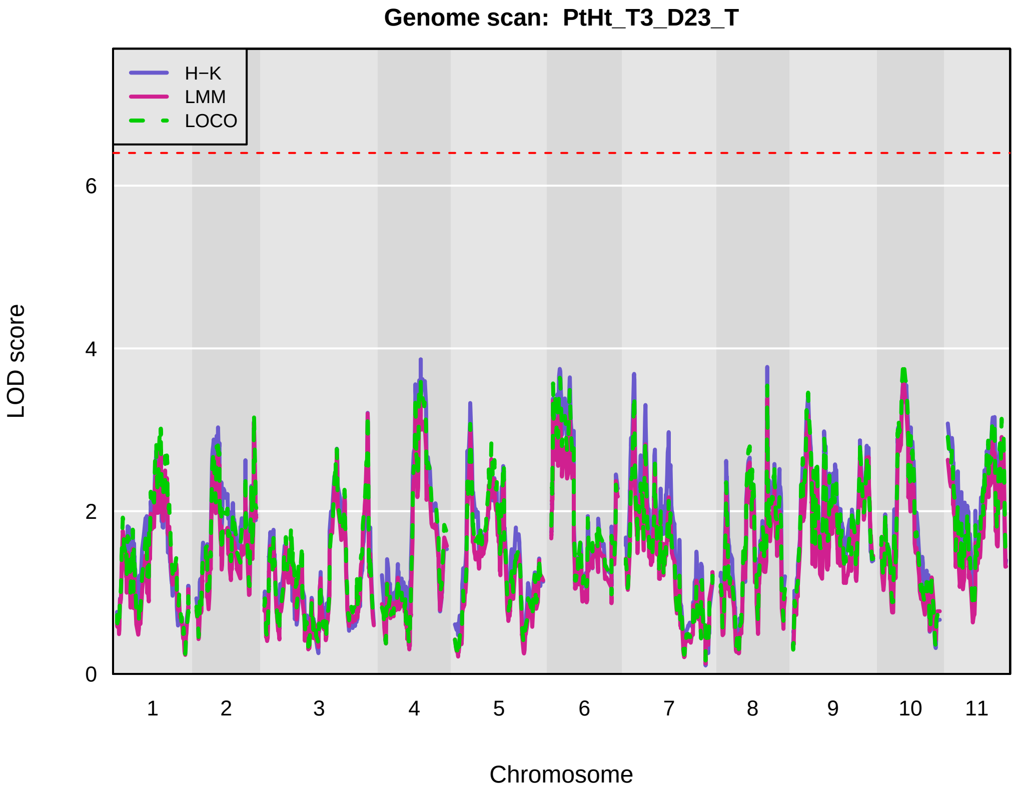
Genome scan: PtHt_T3_D23_T
H−K
LMM
LOCO
LOD score
0
2
4
6
1
2
3
4
5
6
7
8
9
10
11
Chromosome

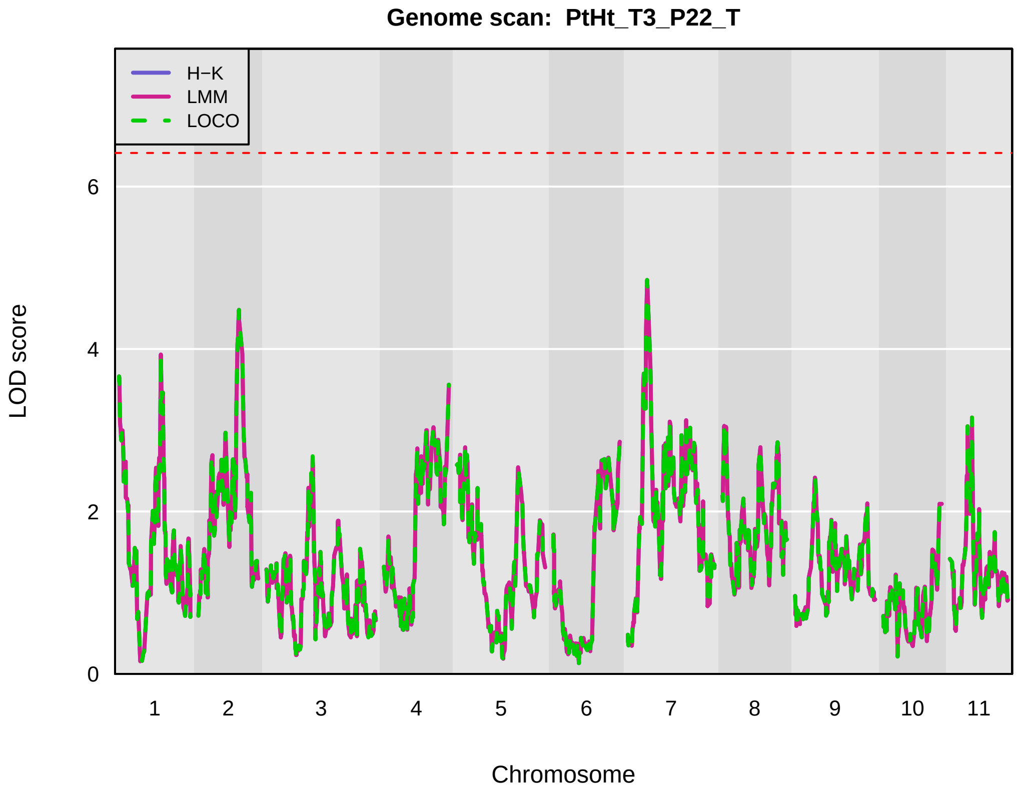
Genome scan: PtHt_T3_P22_T
H−K
LMM
LOCO
LOD score
0
2
4
6
1
2
3
4
5
6
7
8
9
10
11
Chromosome

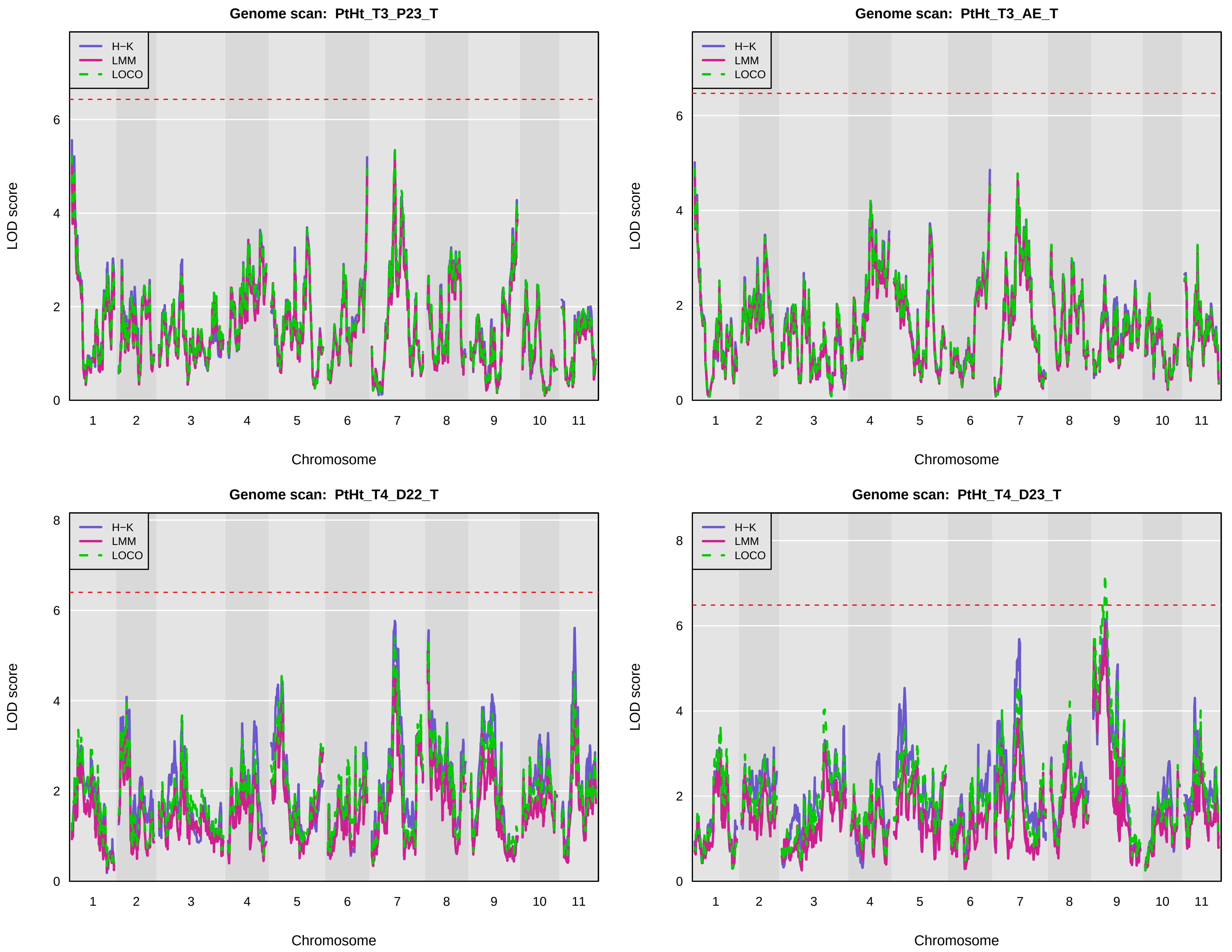
Genome scan: PtHt_T3_P23_T
Genome scan: PtHt_T3_AE_T
Genome scan: PtHt_T4_D22_T
Genome scan: PtHt_T4_D23_T
H−K
LMM
LOCO
LOD score
Chromosome
0
2
4
6
8
1
2
3
4
5
6
7
8
9
10
11

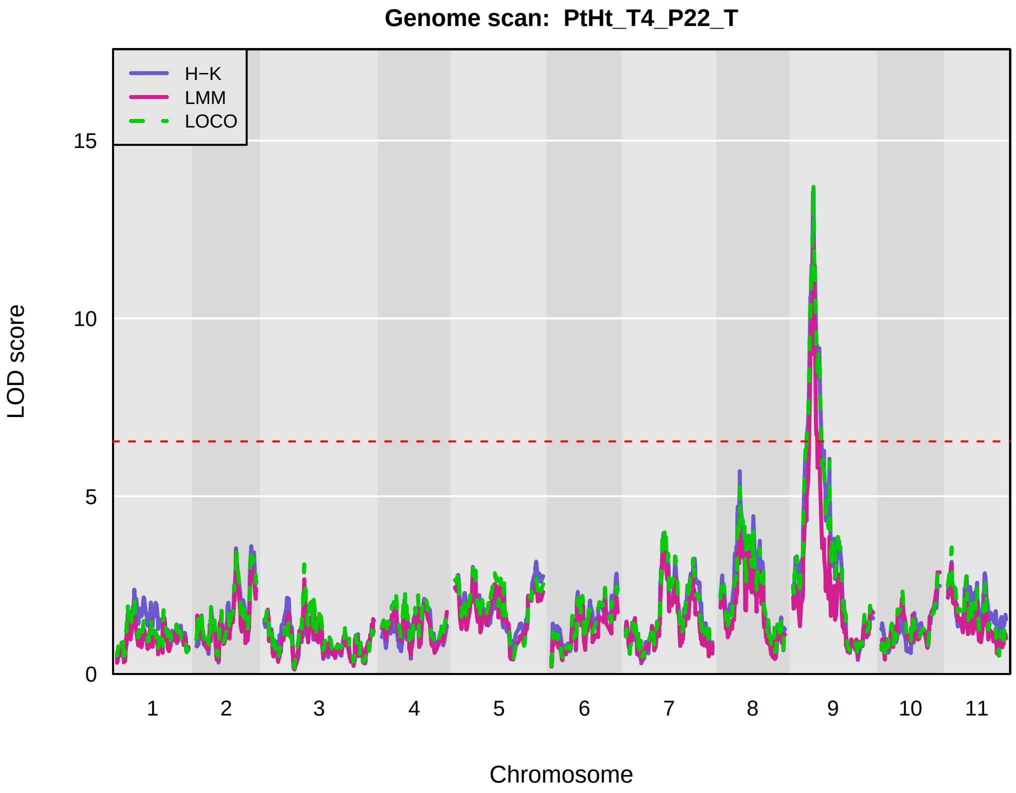
Genome scan: PtHt_T4_P22_T
H−K
LMM
LOCO
LOD score
0
5
10
15
1
2
3
4
5
6
7
8
9
10
11
Chromosome

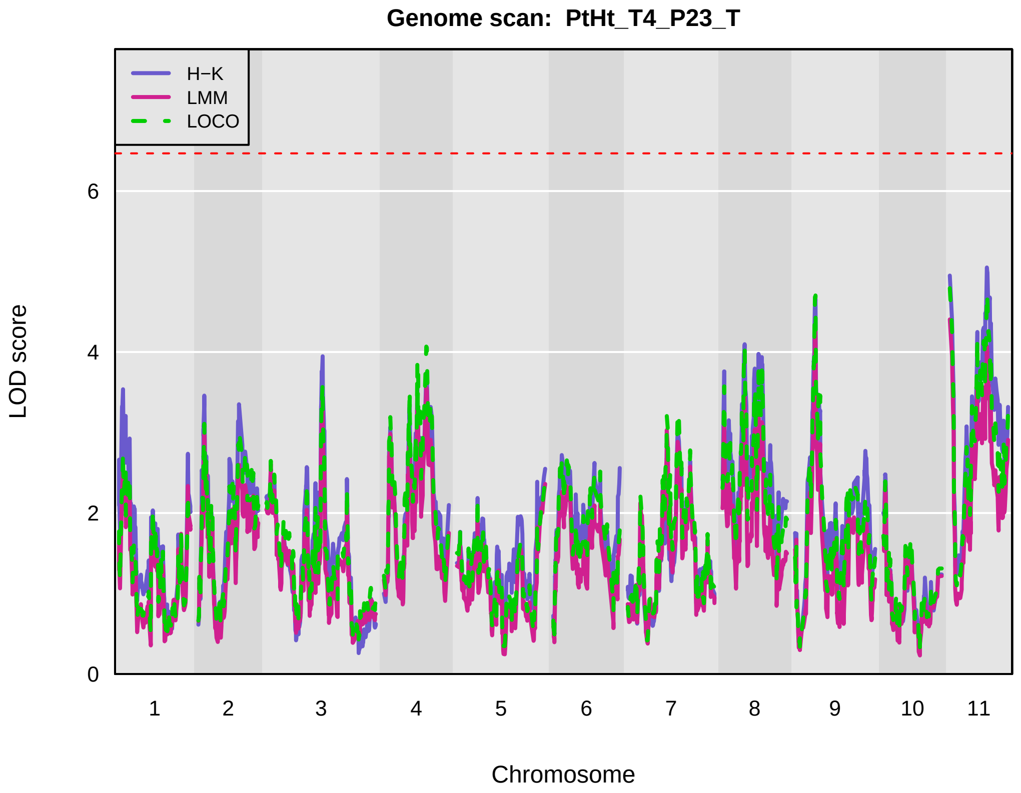
Genome scan: PtHt_T4_P23_T
H−K
LMM
LOCO
LOD score
0
2
4
6
1
2
3
4
5
6
7
8
9
10
11
Chromosome

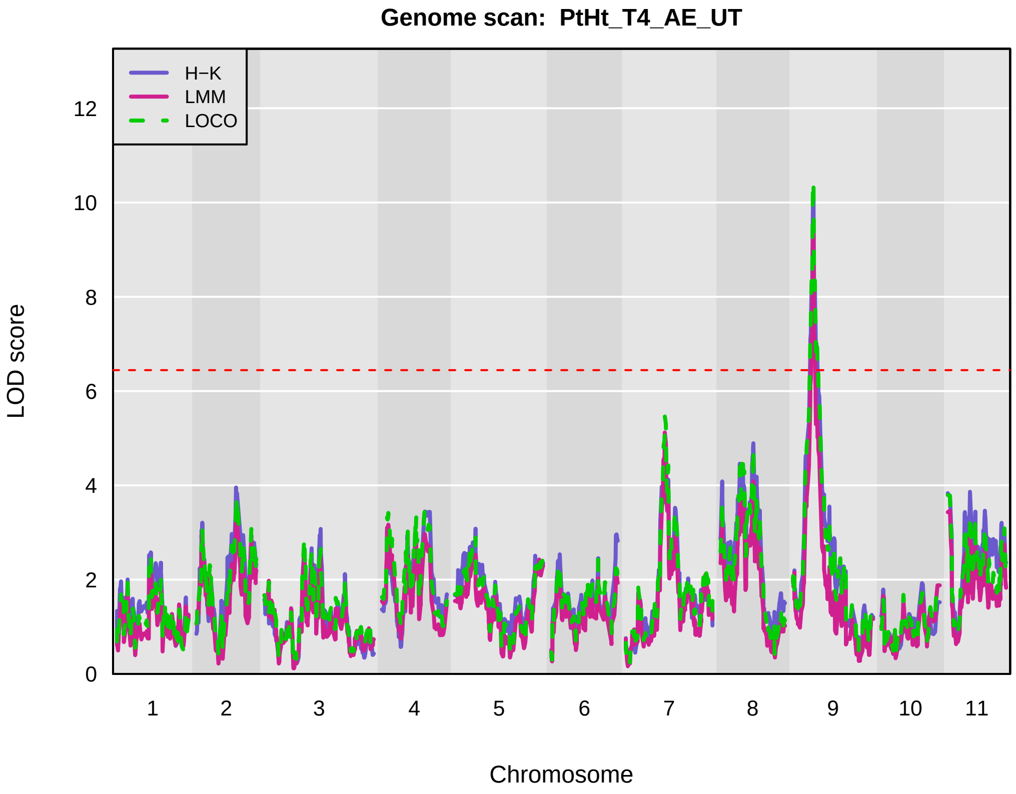
Genome scan: PtHt_T4_AE_UT
H−K
LMM
LOCO
LOD score
0
2
4
6
8
10
12
1
2
3
4
5
6
7
8
9
10
11
Chromosome

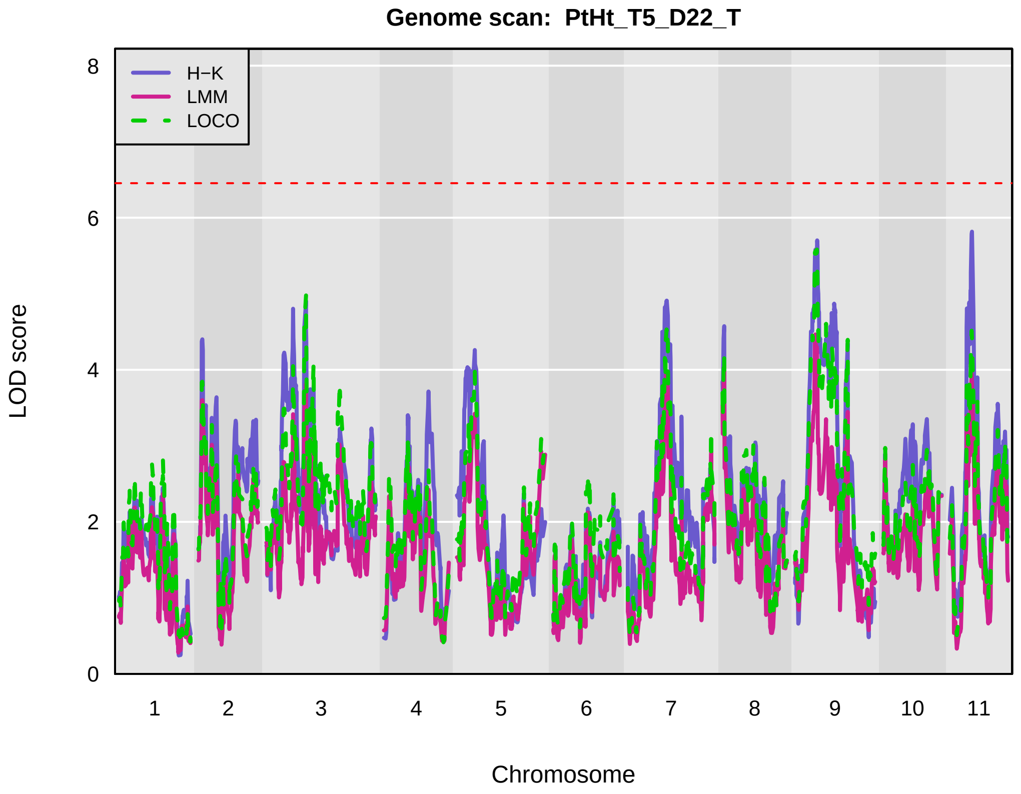
Genome scan: PtHt_T5_D22_T
H−K
LMM
LOCO
LOD score
0
2
4
6
8
1
2
3
4
5
6
7
8
9
10
11
Chromosome

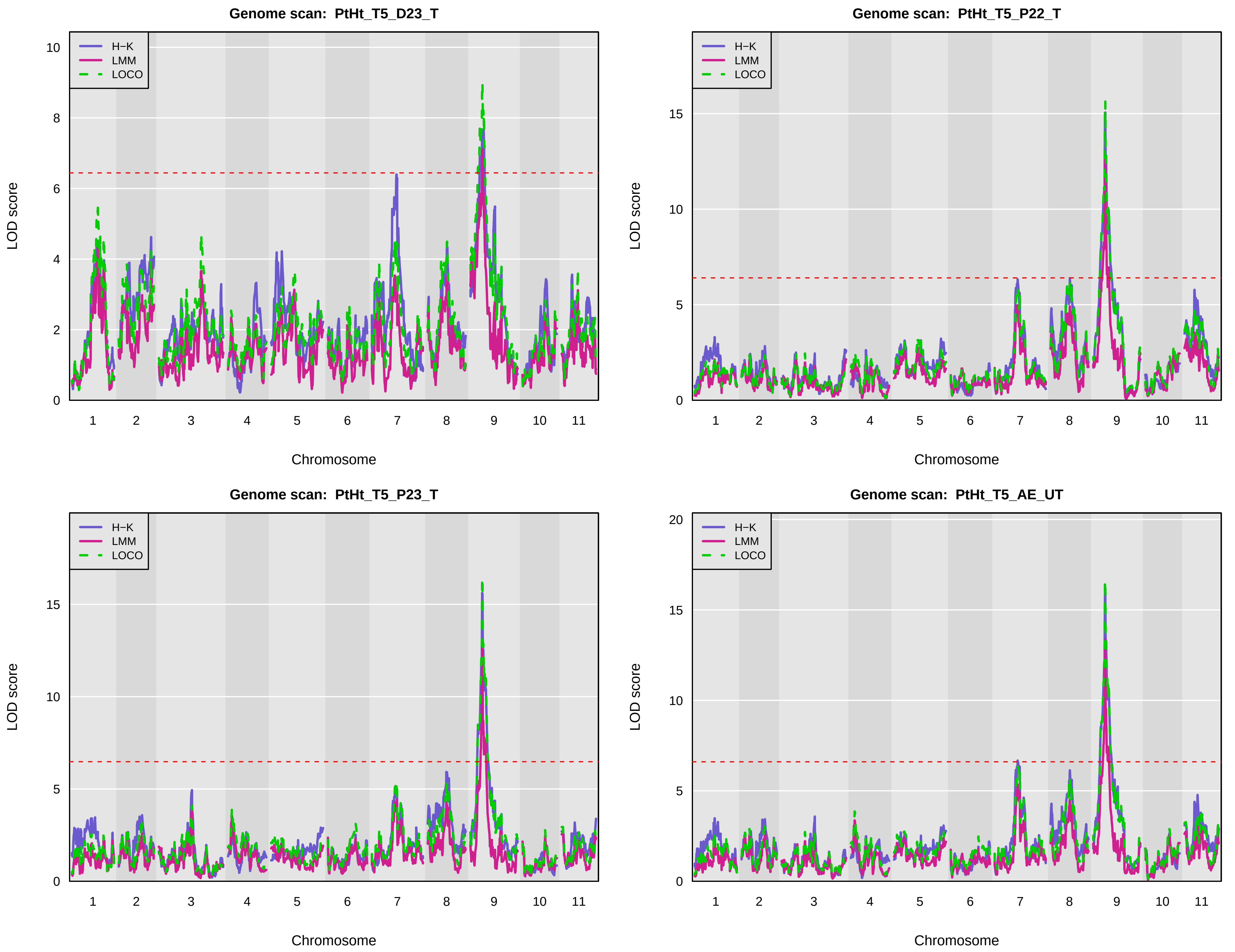
Genome scan: PtHt_T5_D23_T
Genome scan: PtHt_T5_P22_T
Genome scan: PtHt_T5_P23_T
Genome scan: PtHt_T5_AE_UT
H−K
LMM
LOCO
LOD score
Chromosome
1 2 3 4 5 6 7 8 9 10 11

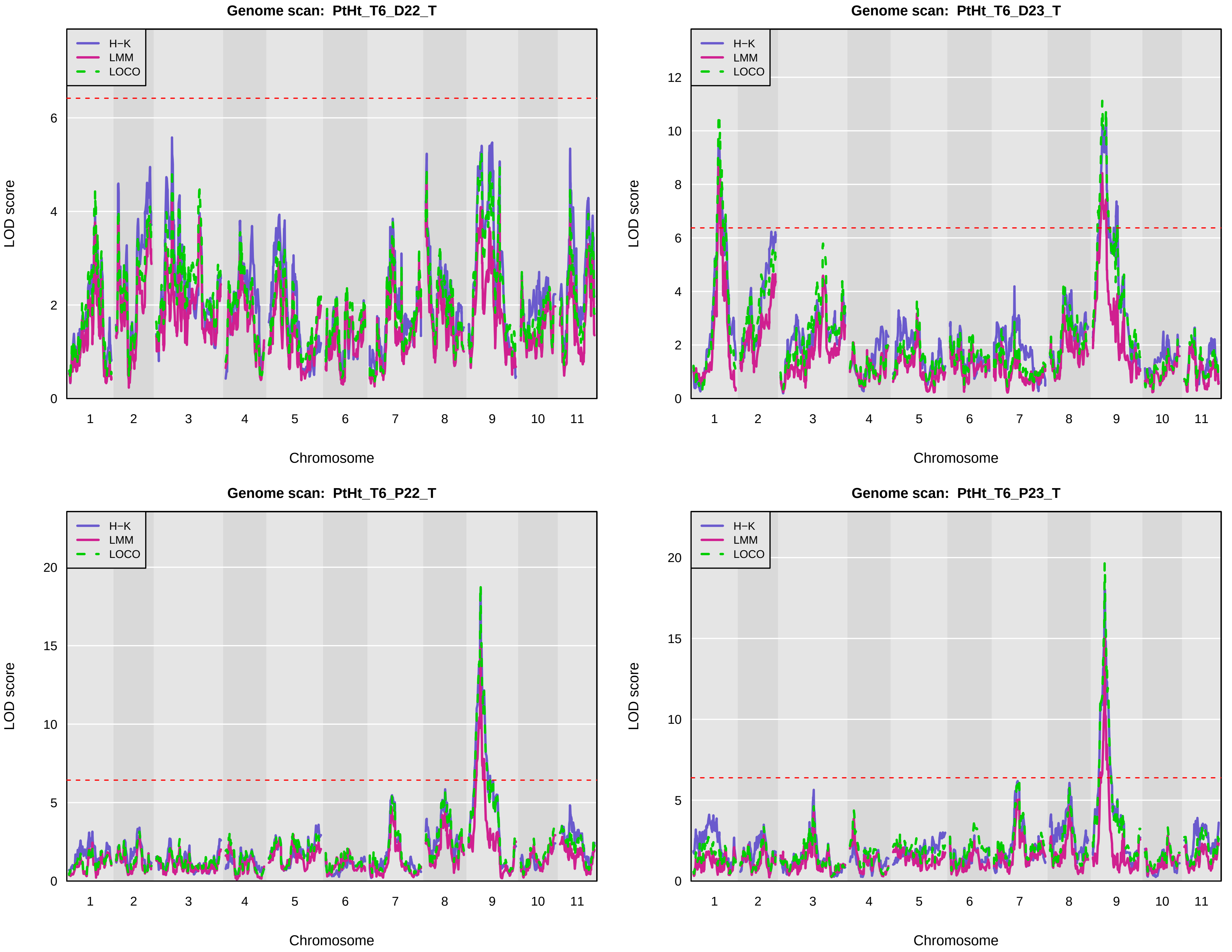
Genome scan: PtHt_T6_D22_T
Genome scan: PtHt_T6_D23_T
Genome scan: PtHt_T6_P22_T
Genome scan: PtHt_T6_P23_T
H−K
LMM
LOCO
LOD score
Chromosome
1
2
3
4
5
6
7
8
9
10
11

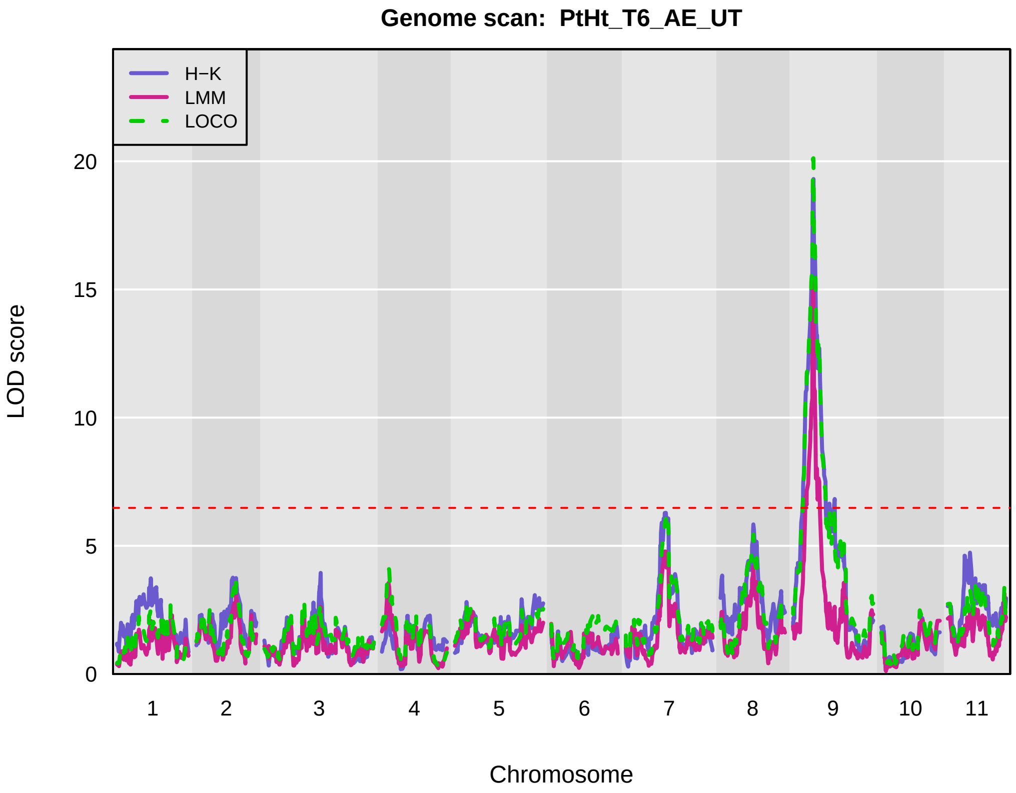
Genome scan: PtHt_T6_AE_UT
H−K
LMM
LOCO
LOD score
0
5
10
15
20
1 2 3 4 5 6 7 8 9 10 11
Chromosome

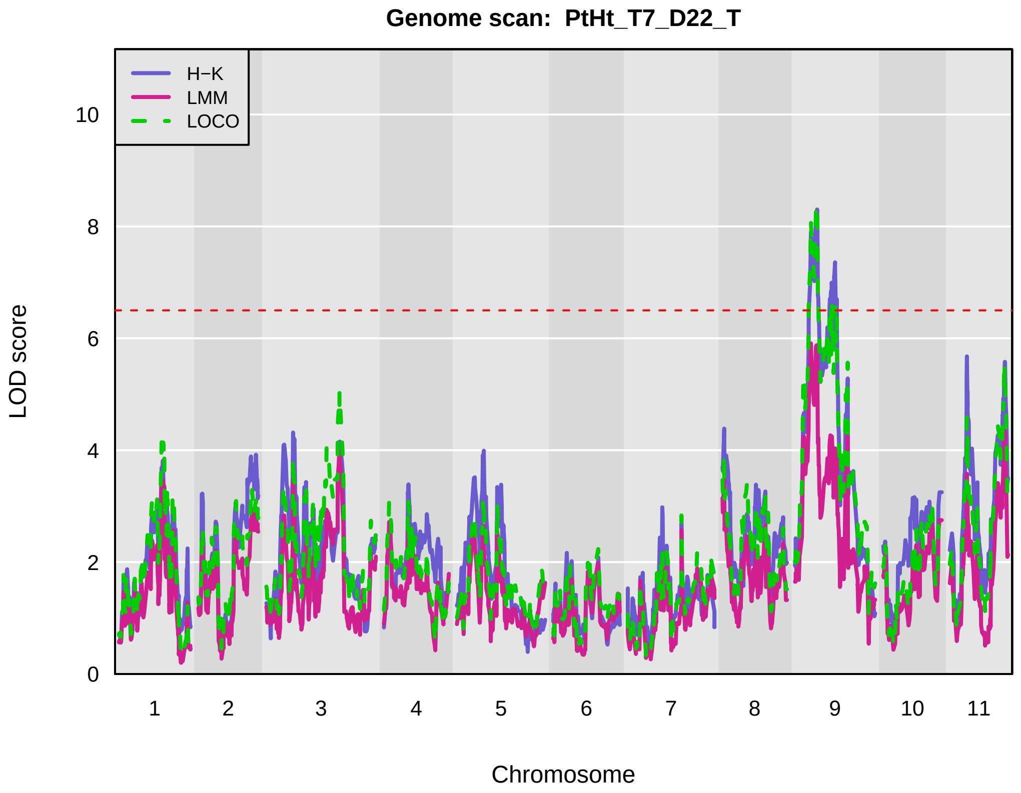
Genome scan: PtHt_T7_D22_T
H−K
LMM
LOCO
LOD score
0
2
4
6
8
10
1 2 3 4 5 6 7 8 9 10 11
Chromosome

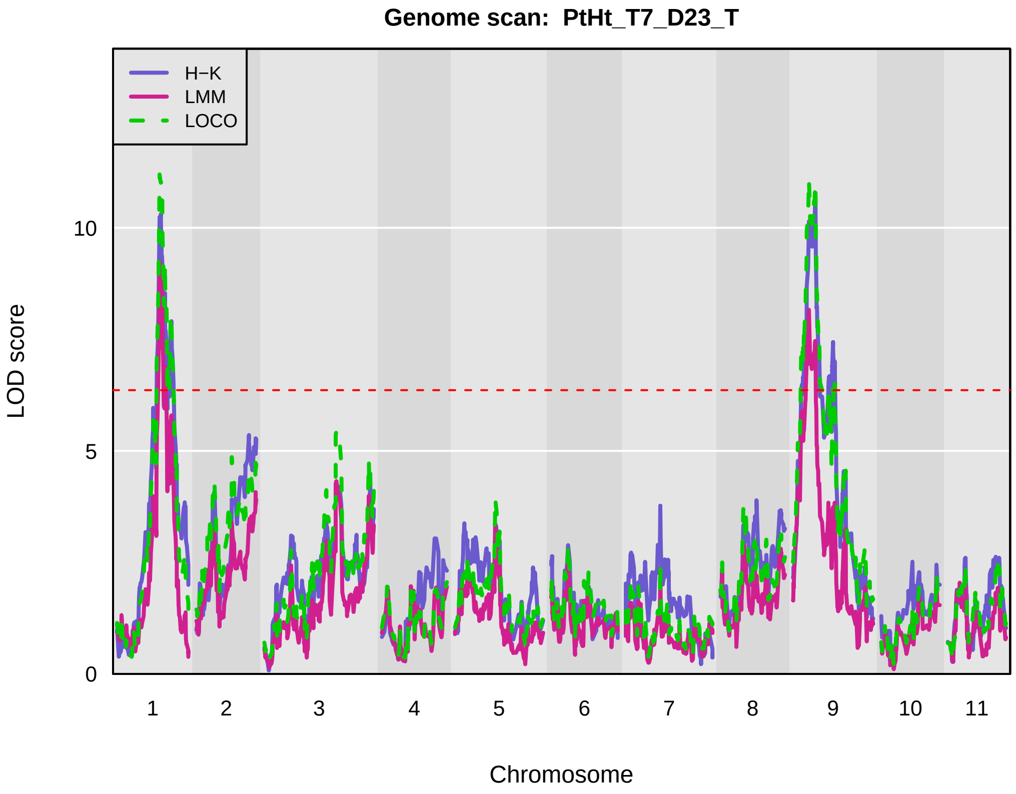
Genome scan: PtHt_T7_D23_T
H−K
LMM
LOCO
LOD score
0
5
10
1 2 3 4 5 6 7 8 9 10 11
Chromosome

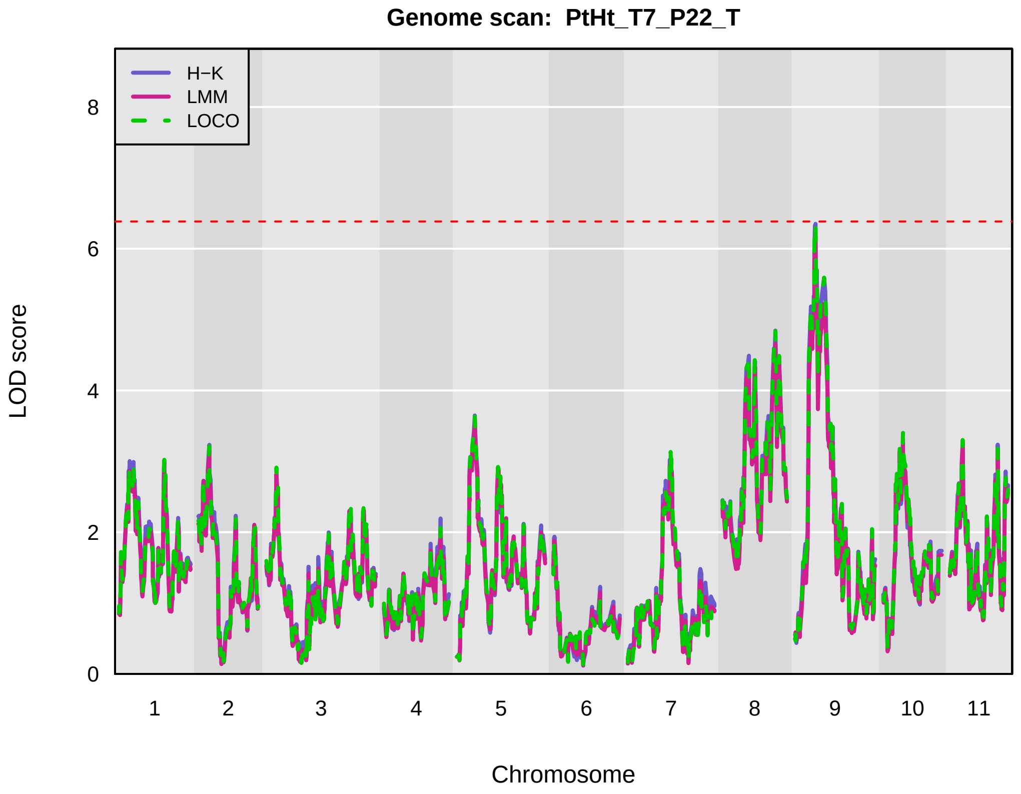
Genome scan: PtHt_T7_P22_T
H−K
LMM
LOCO
LOD score
0
2
4
6
8
1 2 3 4 5 6 7 8 9 10 11
Chromosome

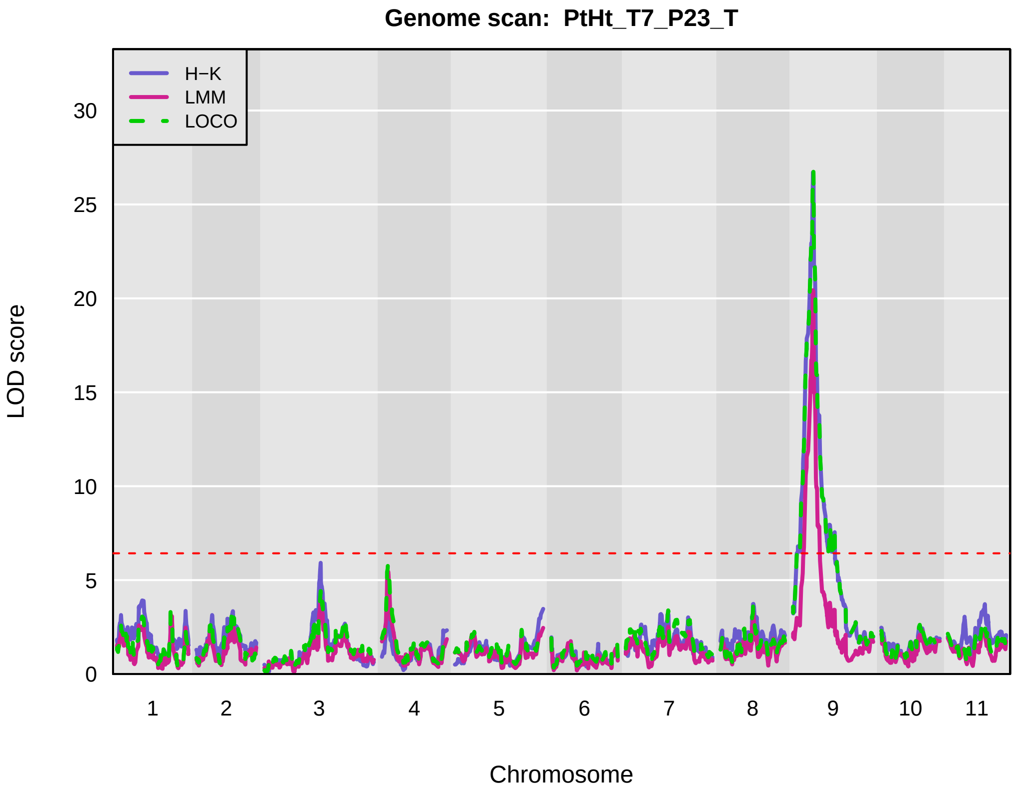
Genome scan: PtHt_T7_P23_T
H−K
LMM
LOCO
LOD score
0
5
10
15
20
25
30
1 2 3 4 5 6 7 8 9 10 11
Chromosome

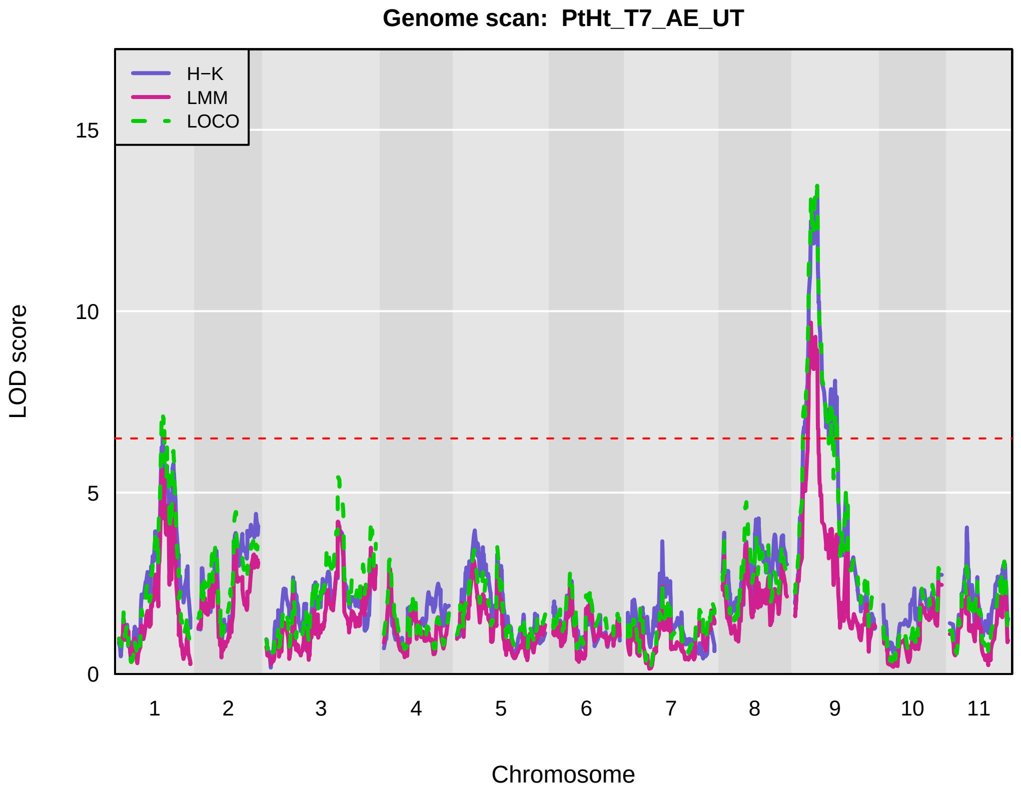
Genome scan: PtHt_T7_AE_UT
H−K
LMM
LOCO
LOD score
0
5
10
15
1 2 3 4 5 6 7 8 9 10 11
Chromosome

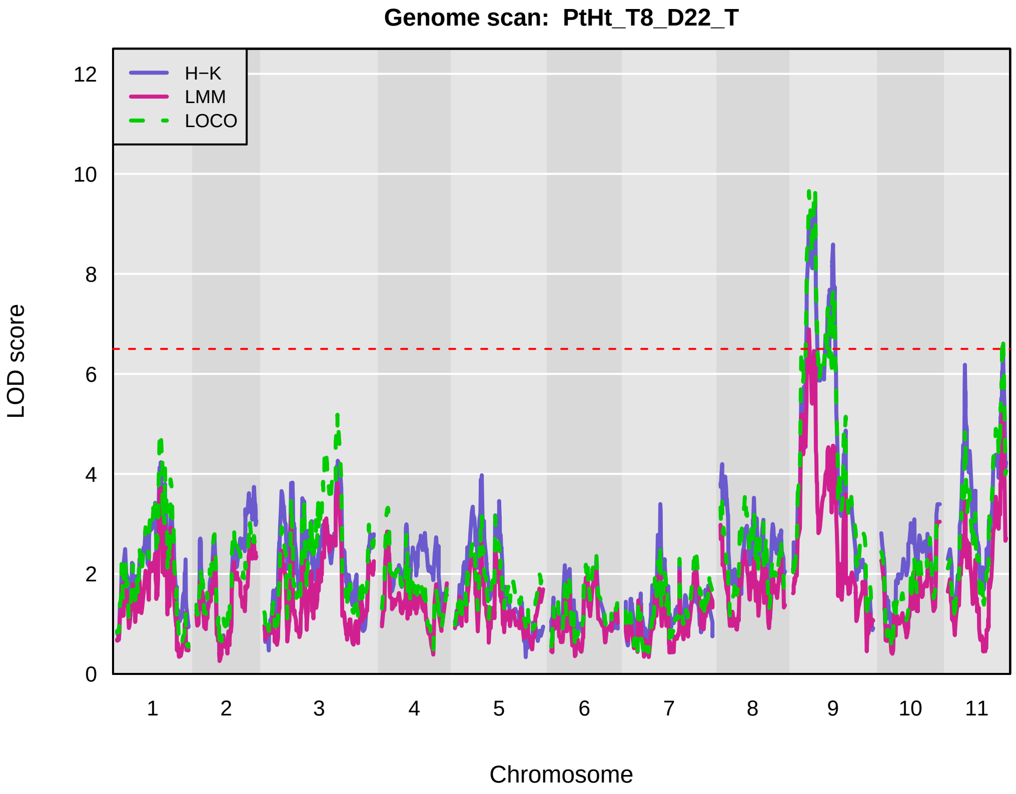
Genome scan: PtHt_T8_D22_T
H−K
LMM
LOCO
LOD score
0
2
4
6
8
10
12
1 2 3 4 5 6 7 8 9 10 11
Chromosome

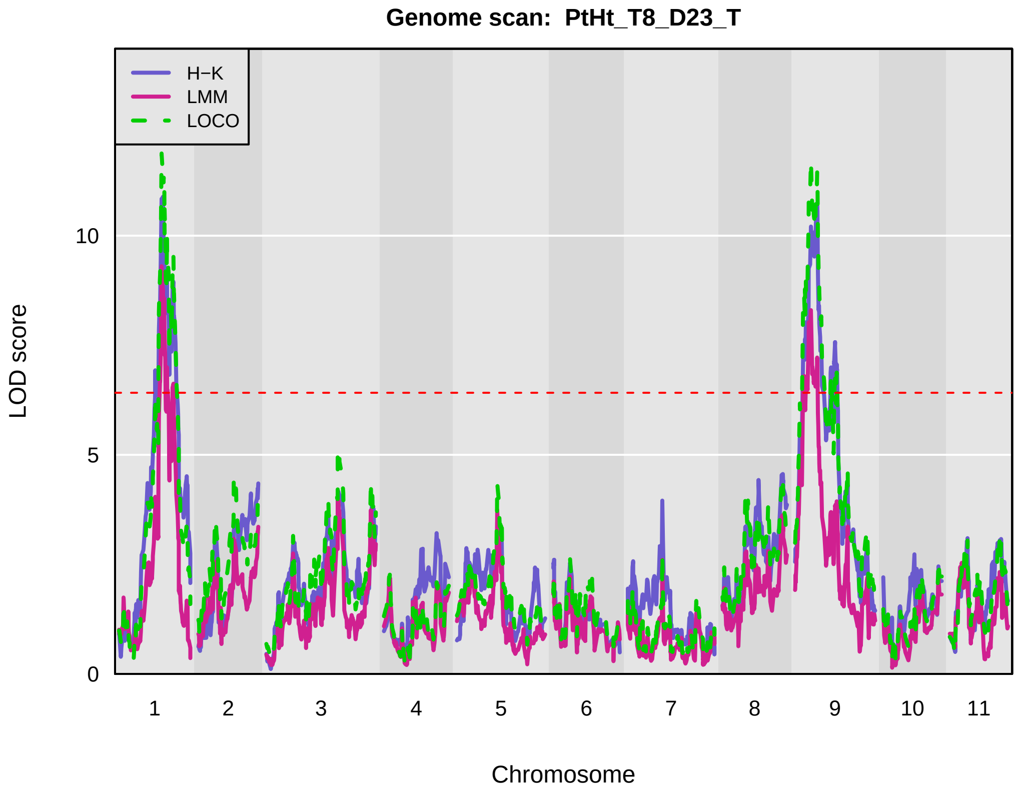
Genome scan: PtHt_T8_D23_T
H−K
LMM
LOCO
LOD score
0
5
10
1 2 3 4 5 6 7 8 9 10 11
Chromosome

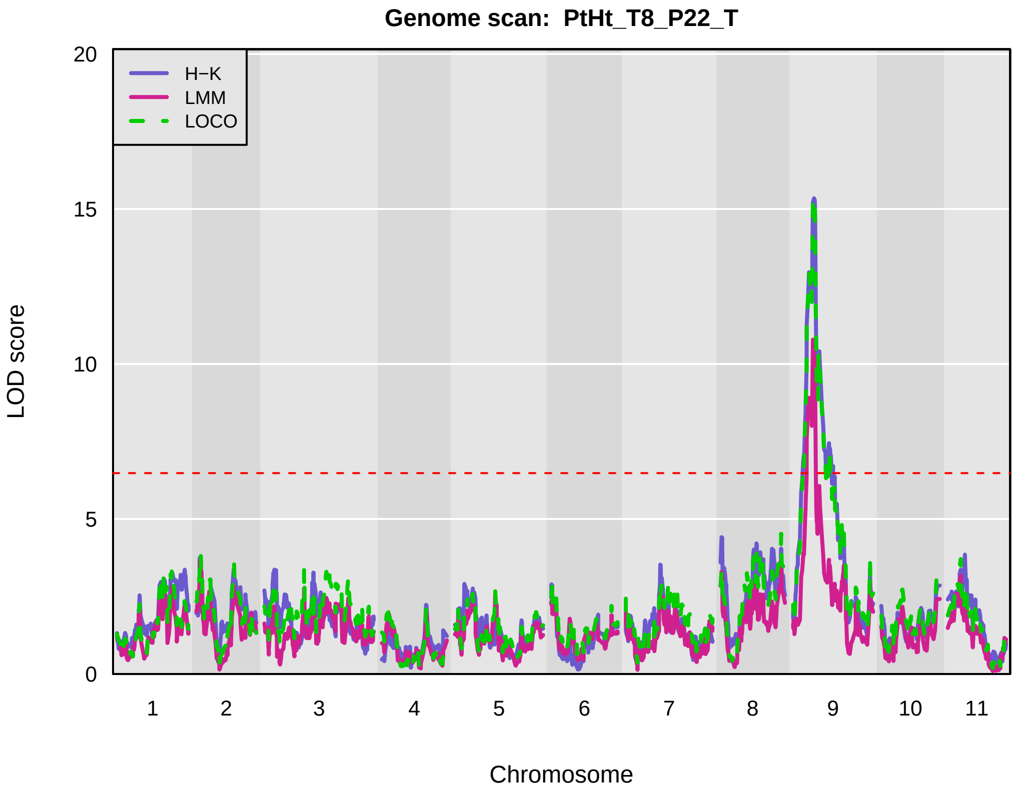
Genome scan: PtHt_T8_P22_T
H−K
LMM
LOCO
LOD score
0
5
10
15
20
1 2 3 4 5 6 7 8 9 10 11
Chromosome

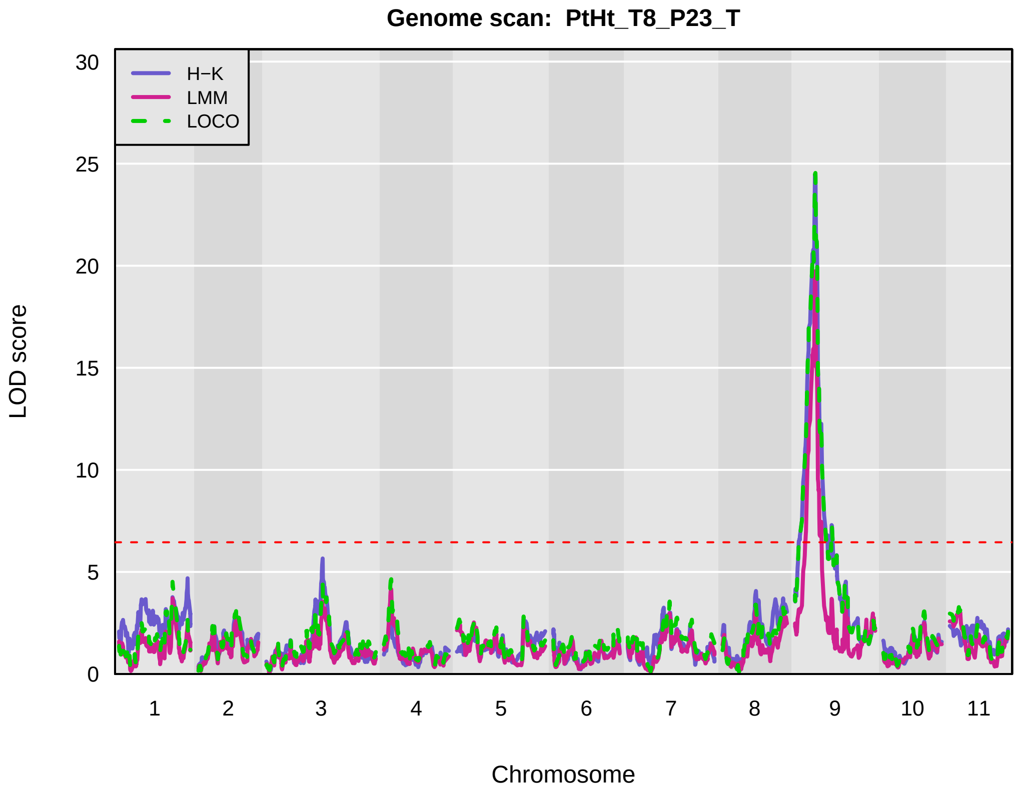
Genome scan: PtHt_T8_P23_T
H−K
LMM
LOCO
LOD score
0
5
10
15
20
25
30
1 2 3 4 5 6 7 8 9 10 11
Chromosome

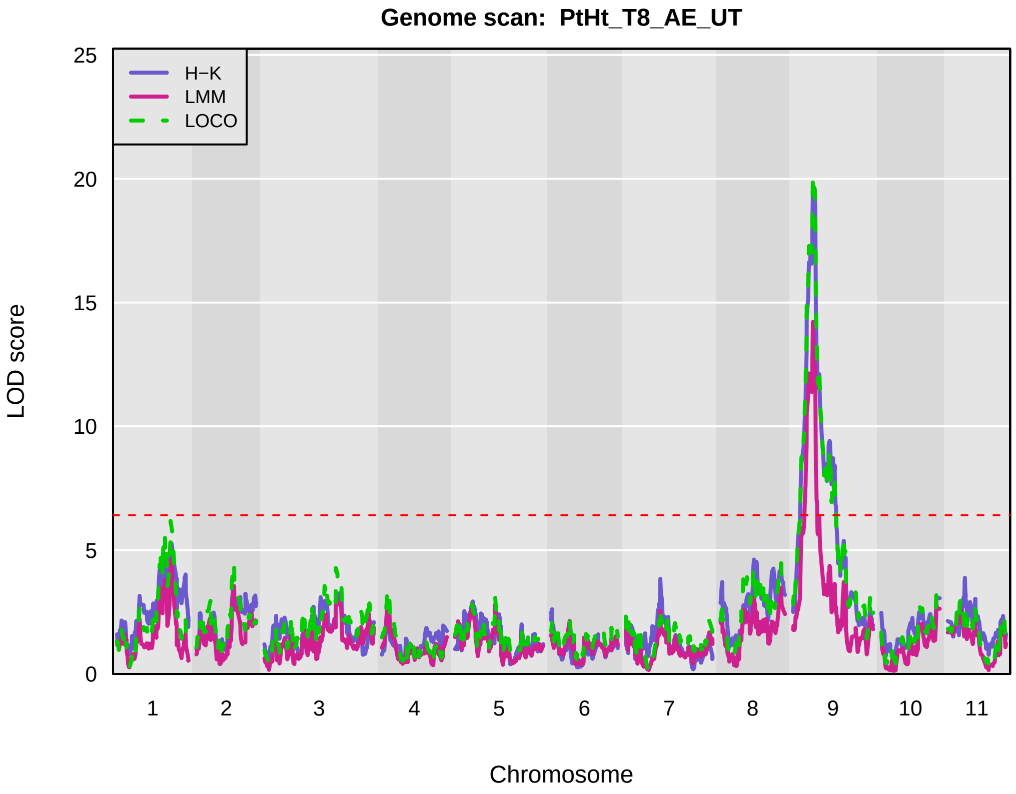
Genome scan: PtHt_T8_AE_UT
H−K
LMM
LOCO
LOD score
0
5
10
15
20
25
1 2 3 4 5 6 7 8 9 10 11
Chromosome

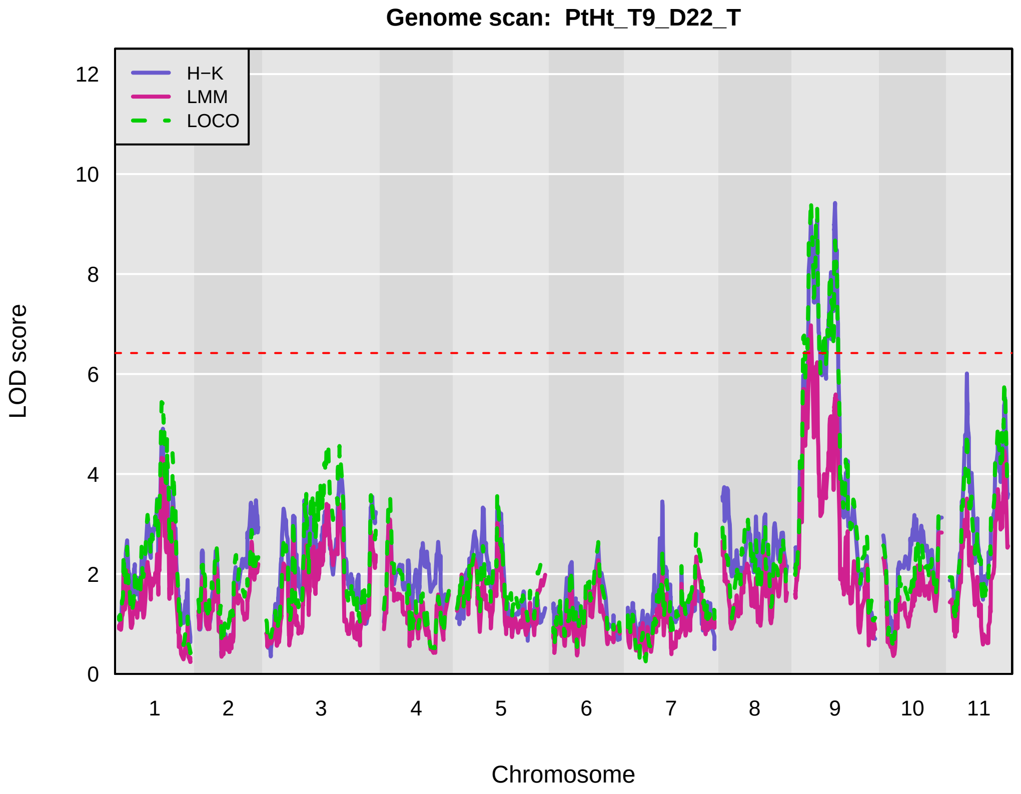
Genome scan: PtHt_T9_D22_T
H−K
LMM
LOCO
LOD score
0
2
4
6
8
10
12
1 2 3 4 5 6 7 8 9 10 11
Chromosome

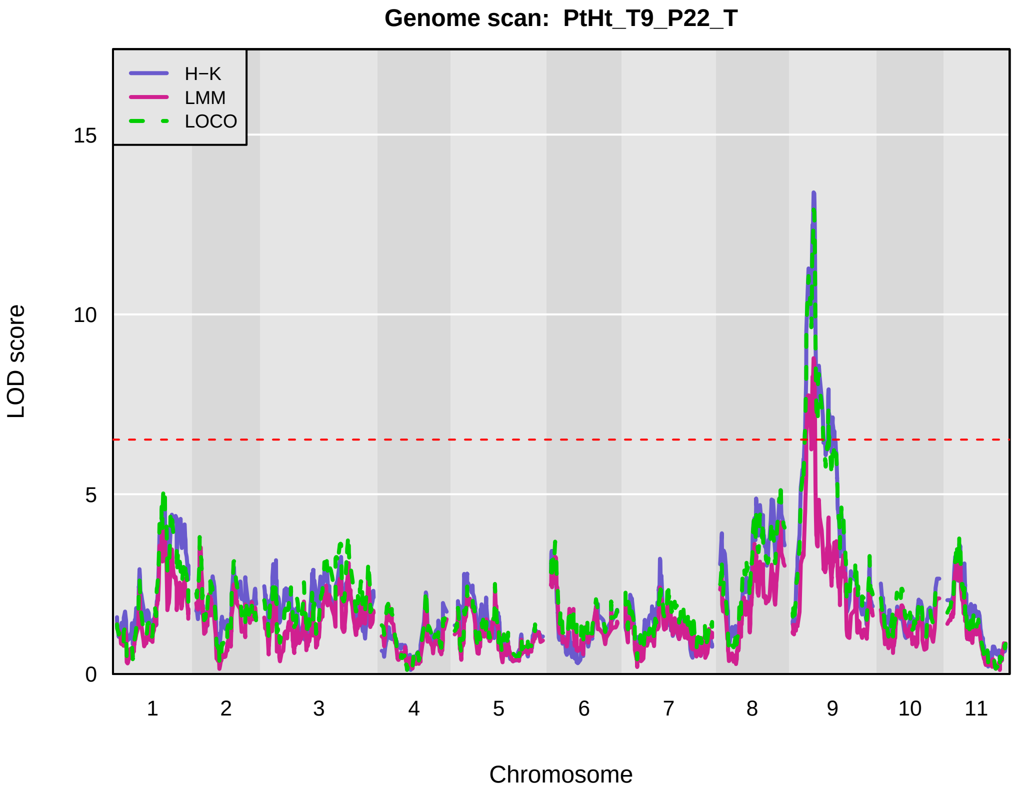
Genome scan: PtHt_T9_P22_T
H−K
LMM
LOCO
LOD score
0
5
10
15
1
2
3
4
5
6
7
8
9
10
11
Chromosome

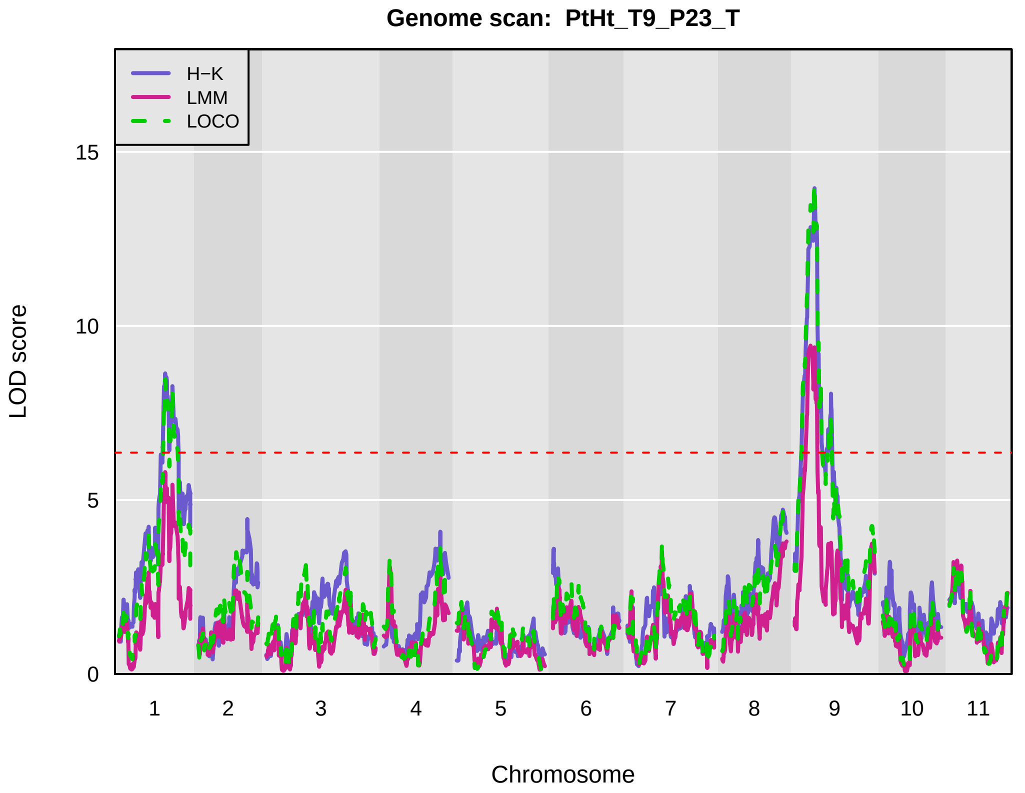
Genome scan: PtHt_T9_P23_T
H−K
LMM
LOCO
LOD score
0
5
10
15
1
2
3
4
5
6
7
8
9
10
11
Chromosome

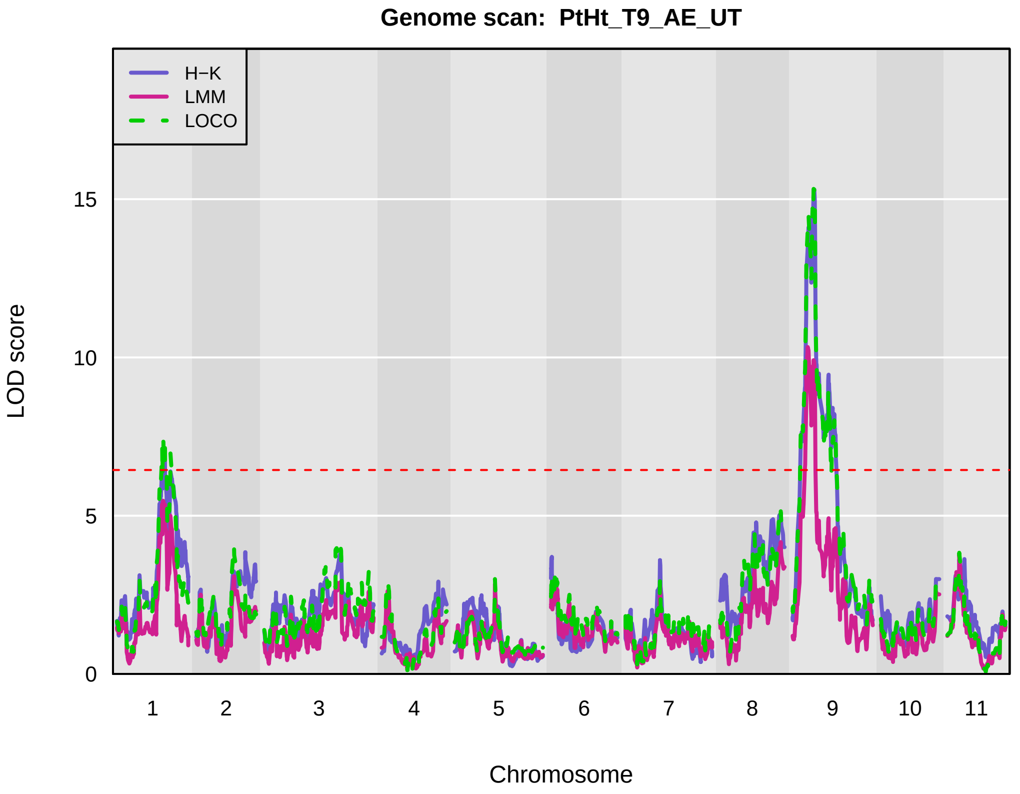
Genome scan: PtHt_T9_AE_UT
H−K
LMM
LOCO
LOD score
0
5
10
15
1
2
3
4
5
6
7
8
9
10
11
Chromosome

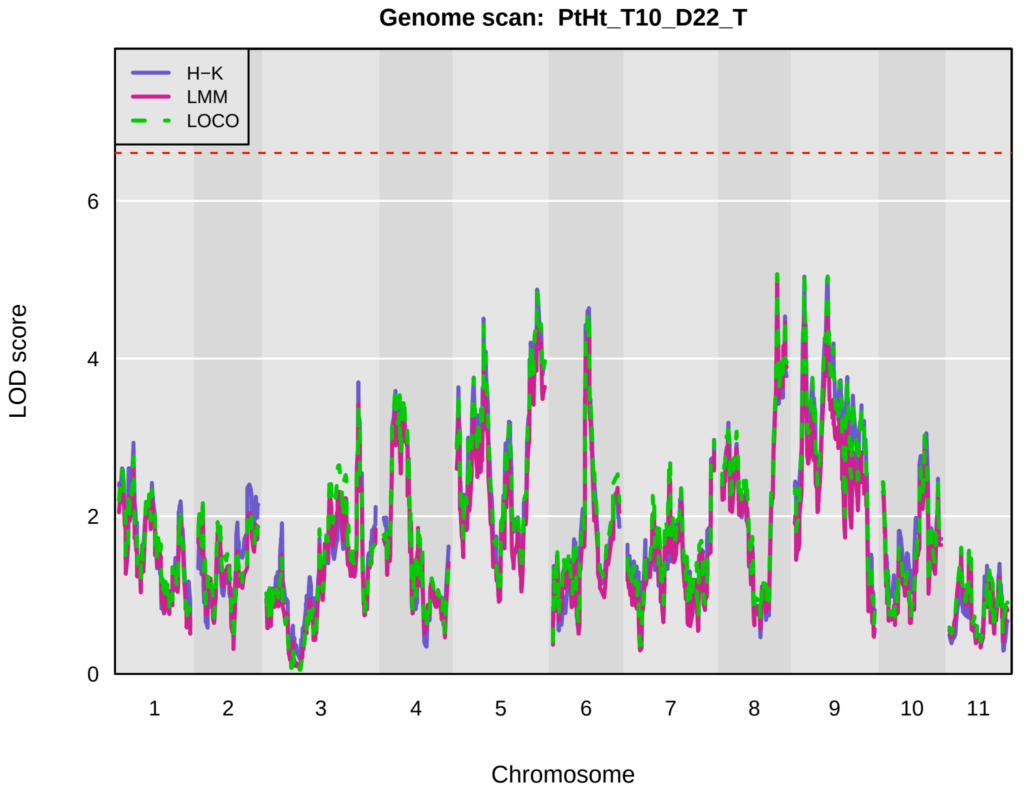
Genome scan: PtHt_T10_D22_T
H−K
LMM
LOCO
LOD score
0
2
4
6
1
2
3
4
5
6
7
8
9
10
11
Chromosome

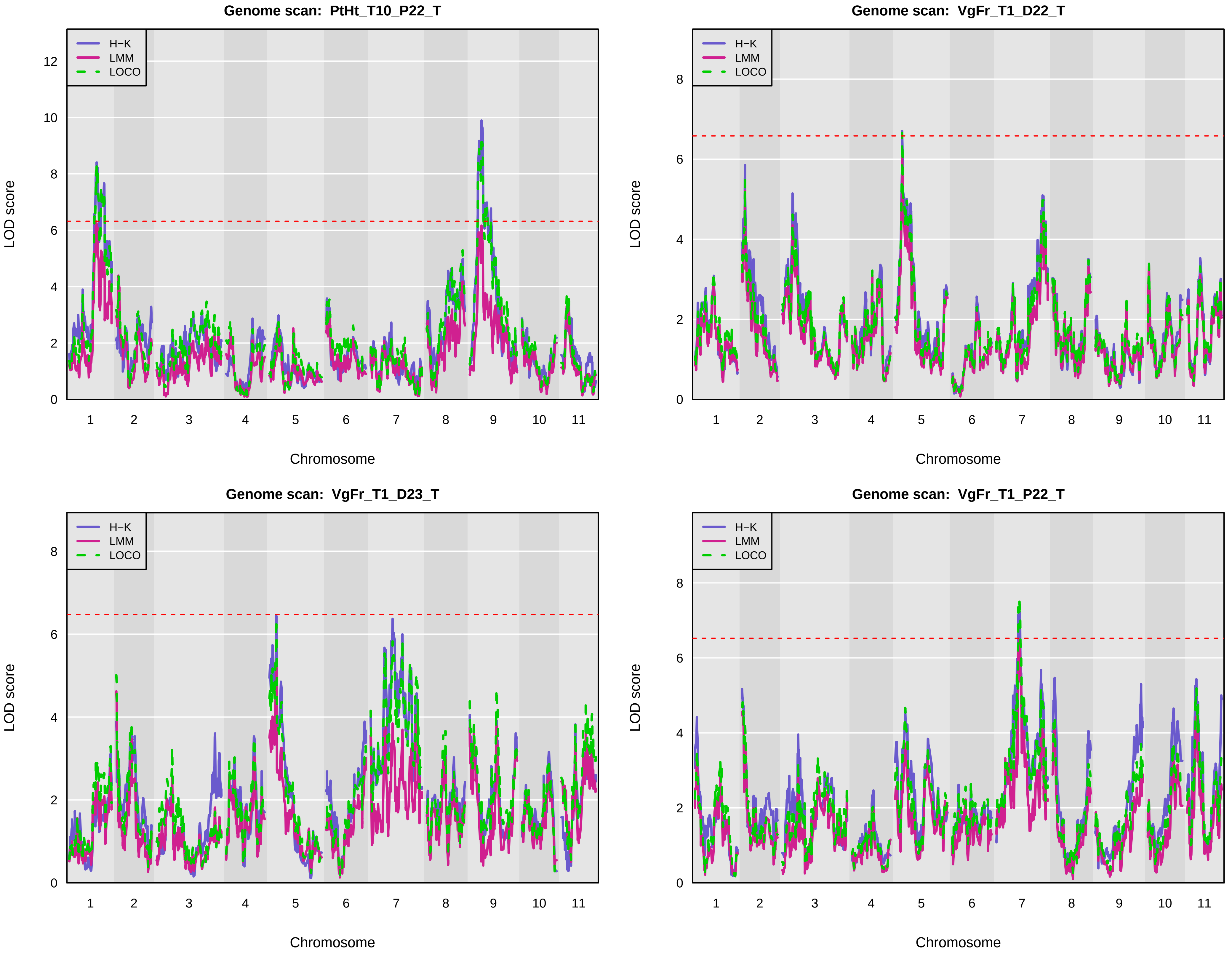

Genome scan: PtHt_T10_P22_T
Genome scan: VgFr_T1_D22_T
Genome scan: VgFr_T1_D23_T
Genome scan: VgFr_T1_P22_T
H−K
LMM
LOCO
LOD score
Chromosome
1
2
3
4
5
6
7
8
9
10
11

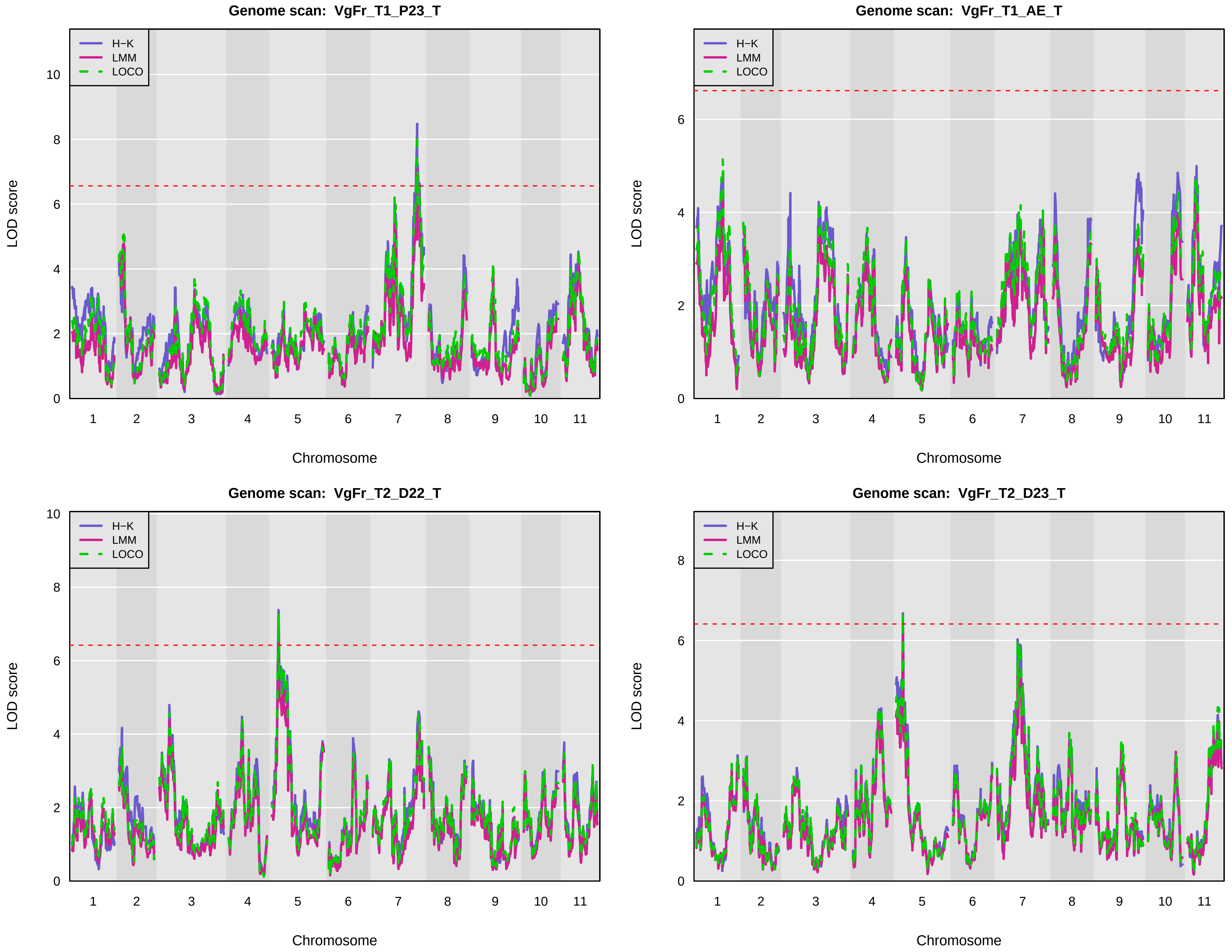

Genome scan: VgFr_T1_P23_T
Genome scan: VgFr_T1_AE_T
Genome scan: VgFr_T2_D22_T
Genome scan: VgFr_T2_D23_T
H−K
LMM
LOCO
LOD score
Chromosome
1 2 3 4 5 6 7 8 9 10 11

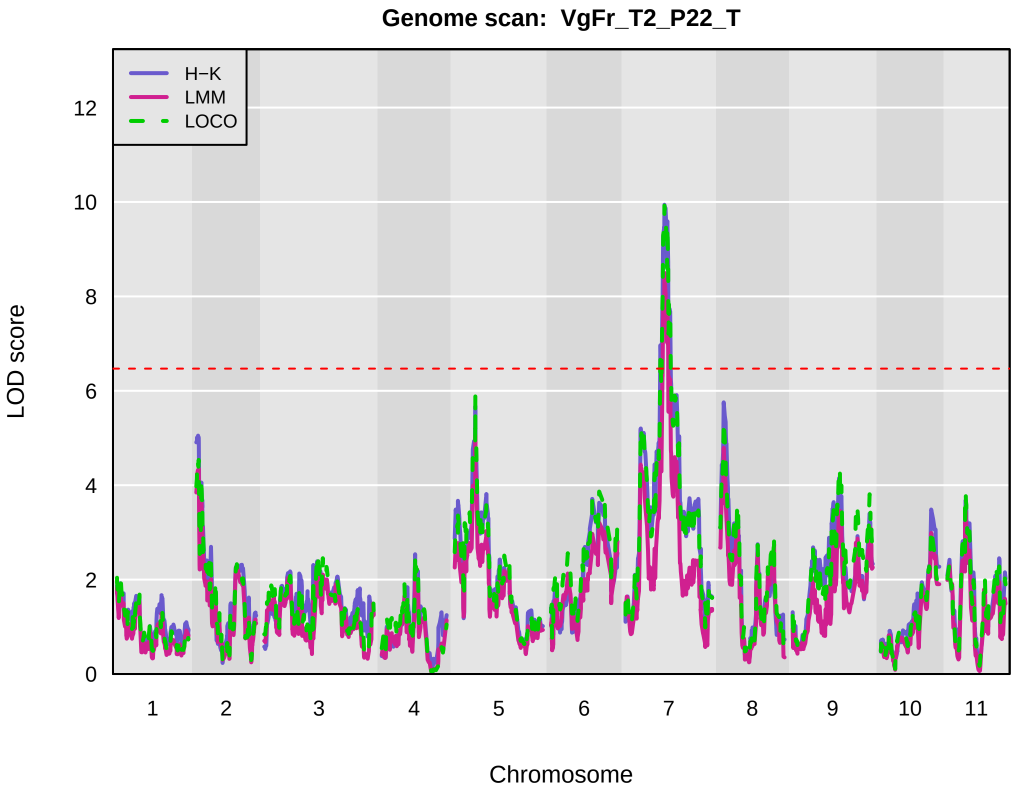
Genome scan: VgFr_T2_P22_T
H−K
LMM
LOCO
LOD score
0
2
4
6
8
10
12
Chromosome
1 2 3 4 5 6 7 8 9 10 11

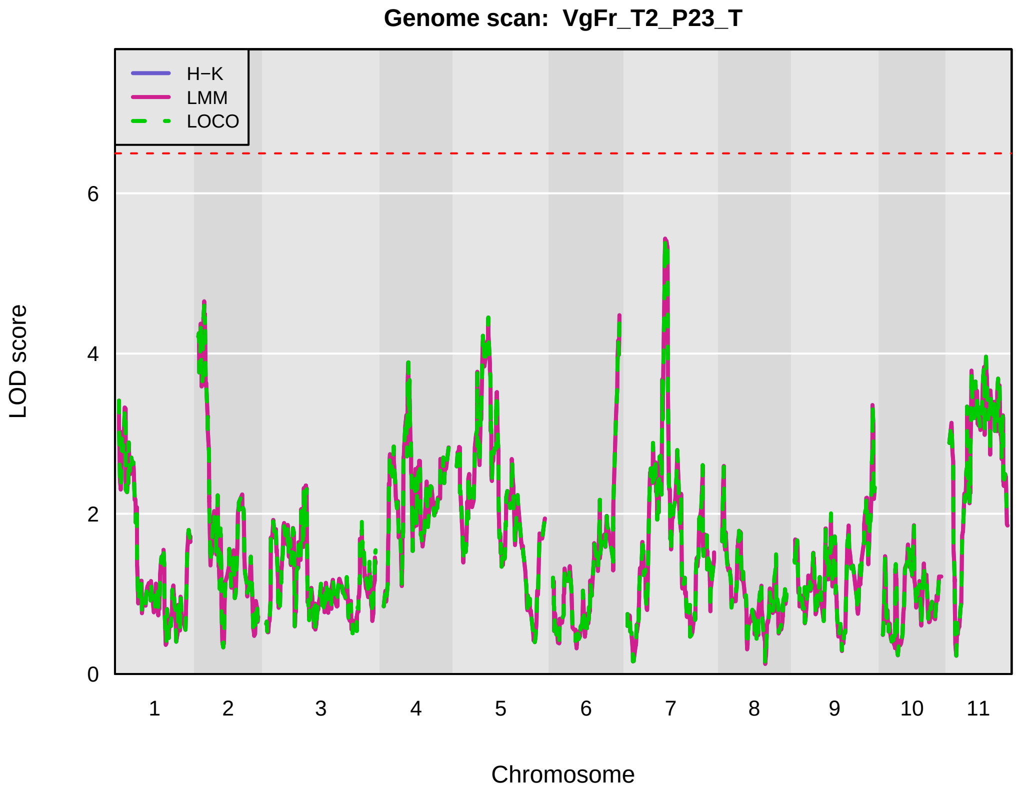
Genome scan: VgFr_T2_P23_T
H−K
LMM
LOCO
LOD score
0
2
4
6
Chromosome
1 2 3 4 5 6 7 8 9 10 11

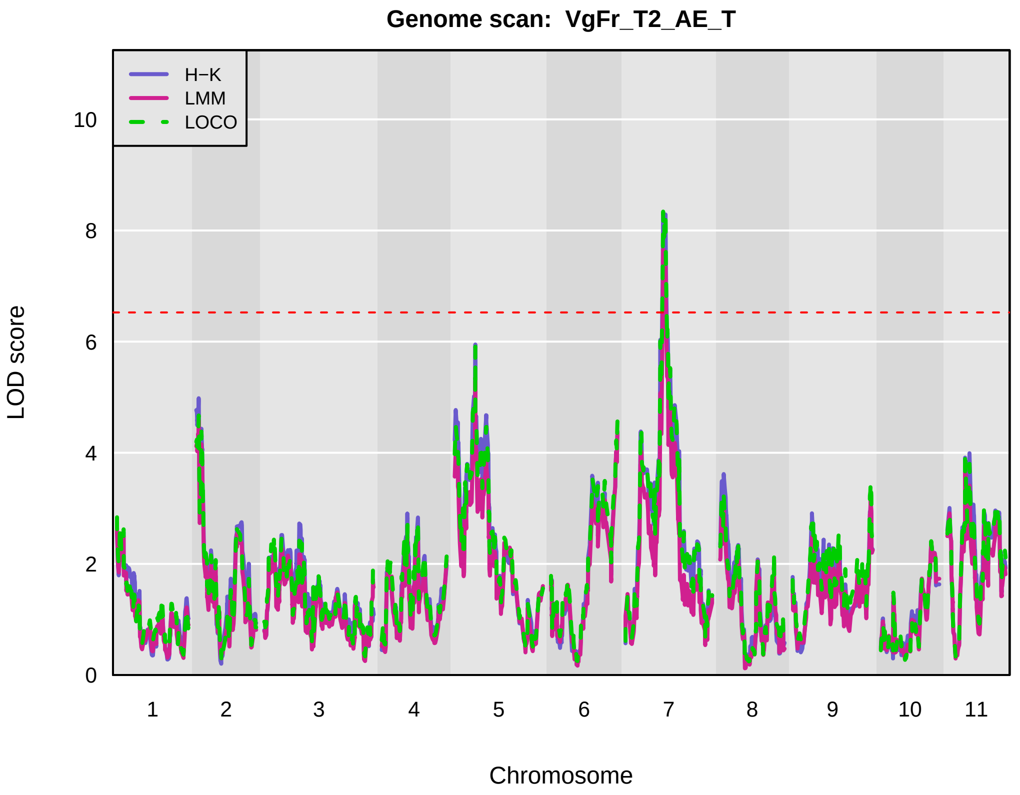
Genome scan: VgFr_T2_AE_T
H−K
LMM
LOCO
LOD score
0
2
4
6
8
10
Chromosome
1 2 3 4 5 6 7 8 9 10 11

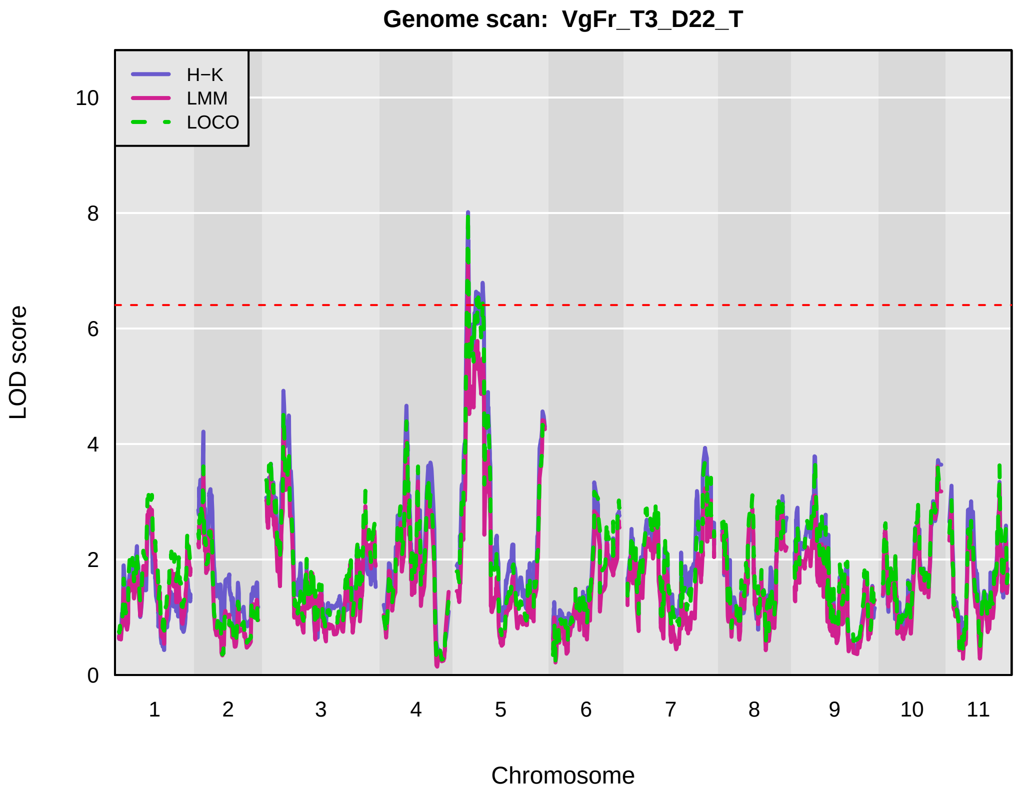
Genome scan: VgFr_T3_D22_T
H−K
LMM
LOCO
LOD score
0
2
4
6
8
10
Chromosome
1 2 3 4 5 6 7 8 9 10 11

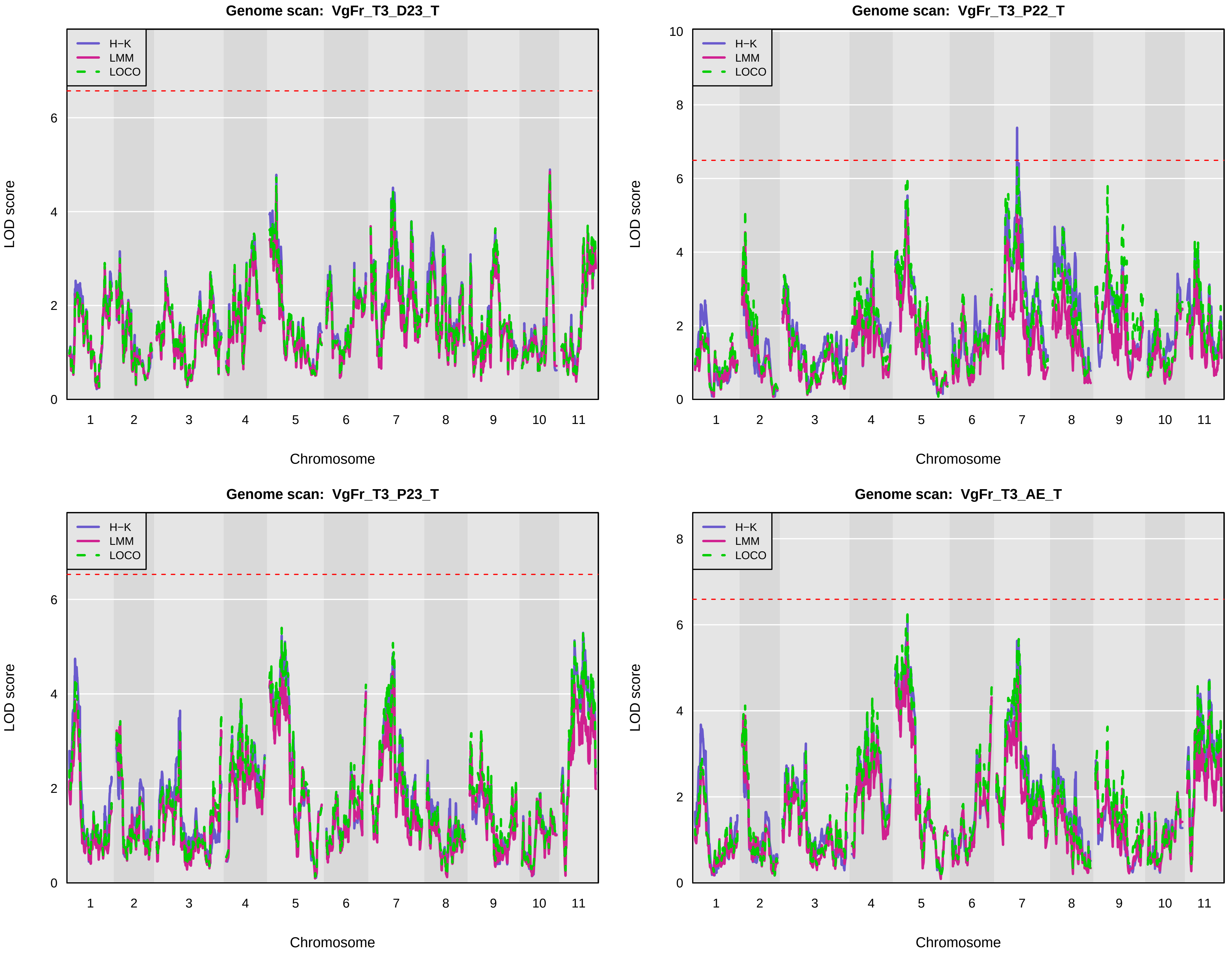
Genome scan: VgFr_T3_D23_T
H−K
LMM
LOCO
LOD score
0
2
4
6
1
2
3
4
5
6
7
8
9
10
11
Chromosome
Genome scan: VgFr_T3_P22_T
H−K
LMM
LOCO
LOD score
0
2
4
6
8
10
1
2
3
4
5
6
7
8
9
10
11
Chromosome
Genome scan: VgFr_T3_P23_T
H−K
LMM
LOCO
LOD score
0
2
4
6
1
2
3
4
5
6
7
8
9
10
11
Chromosome
Genome scan: VgFr_T3_AE_T
H−K
LMM
LOCO
LOD score
0
2
4
6
8
1
2
3
4
5
6
7
8
9
10
11
Chromosome

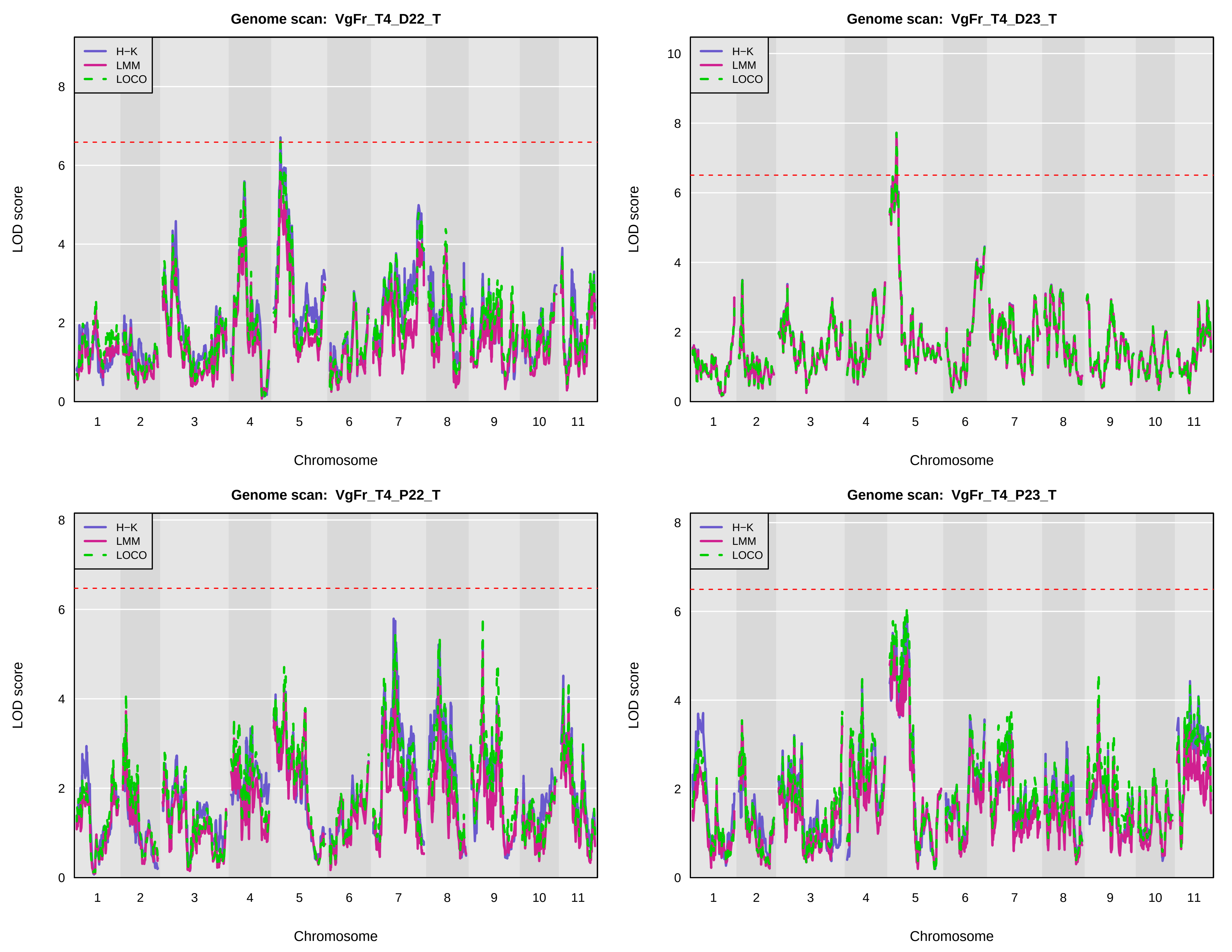

Genome scan: VgFr_T4_D22_T
H−K
LMM
LOCO
LOD score
Chromosome
1 2 3 4 5 6 7 8 9 10 11
Genome scan: VgFr_T4_D23_T
H−K
LMM
LOCO
LOD score
Chromosome
1 2 3 4 5 6 7 8 9 10 11
Genome scan: VgFr_T4_P22_T
H−K
LMM
LOCO
LOD score
Chromosome
1 2 3 4 5 6 7 8 9 10 11
Genome scan: VgFr_T4_P23_T
H−K
LMM
LOCO
LOD score
Chromosome
1 2 3 4 5 6 7 8 9 10 11

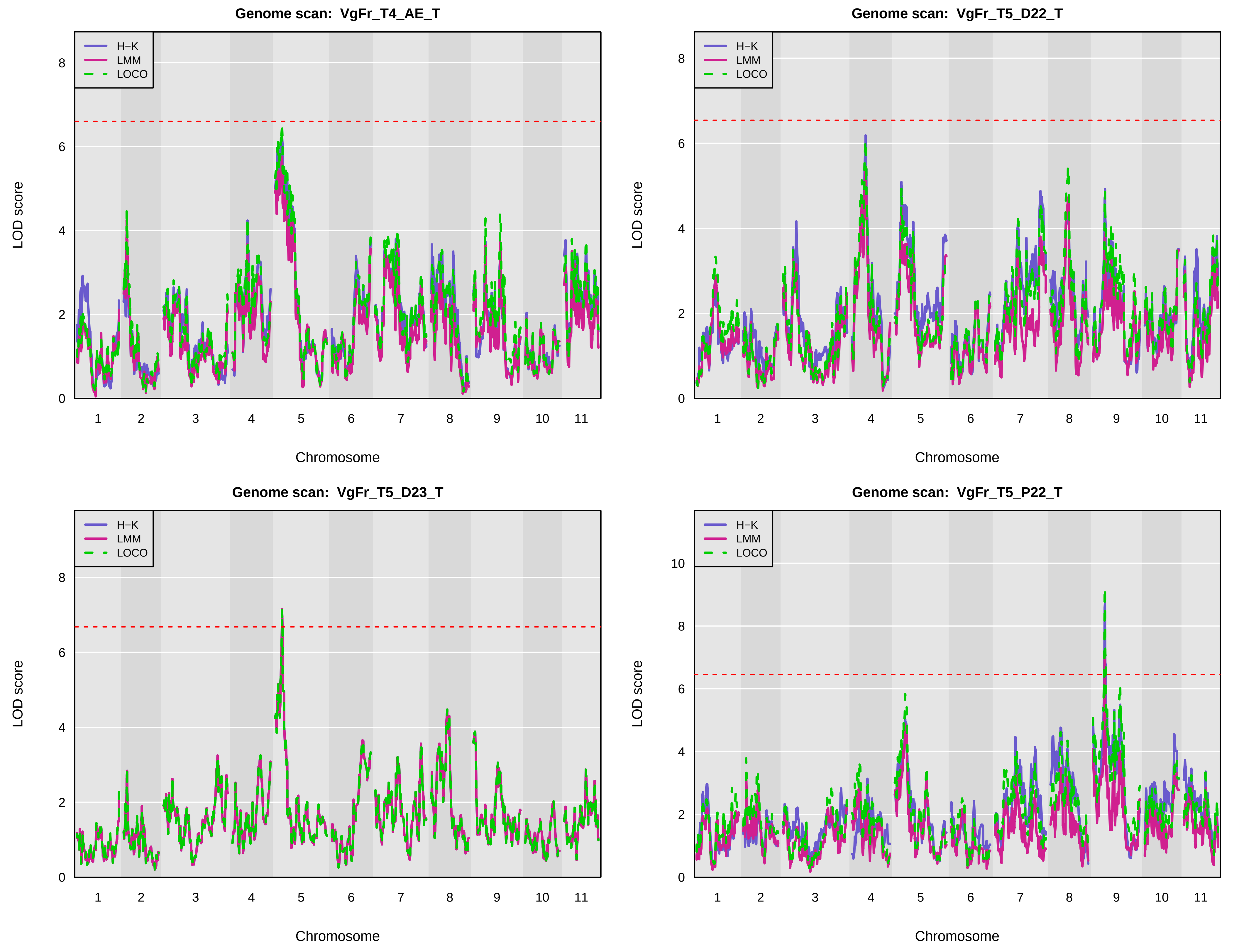
Genome scan: VgFr_T4_AE_T
Genome scan: VgFr_T5_D22_T
Genome scan: VgFr_T5_D23_T
Genome scan: VgFr_T5_P22_T
H−K
LMM
LOCO
LOD score
Chromosome
0
2
4
6
8
10
1
2
3
4
5
6
7
8
9
10
11

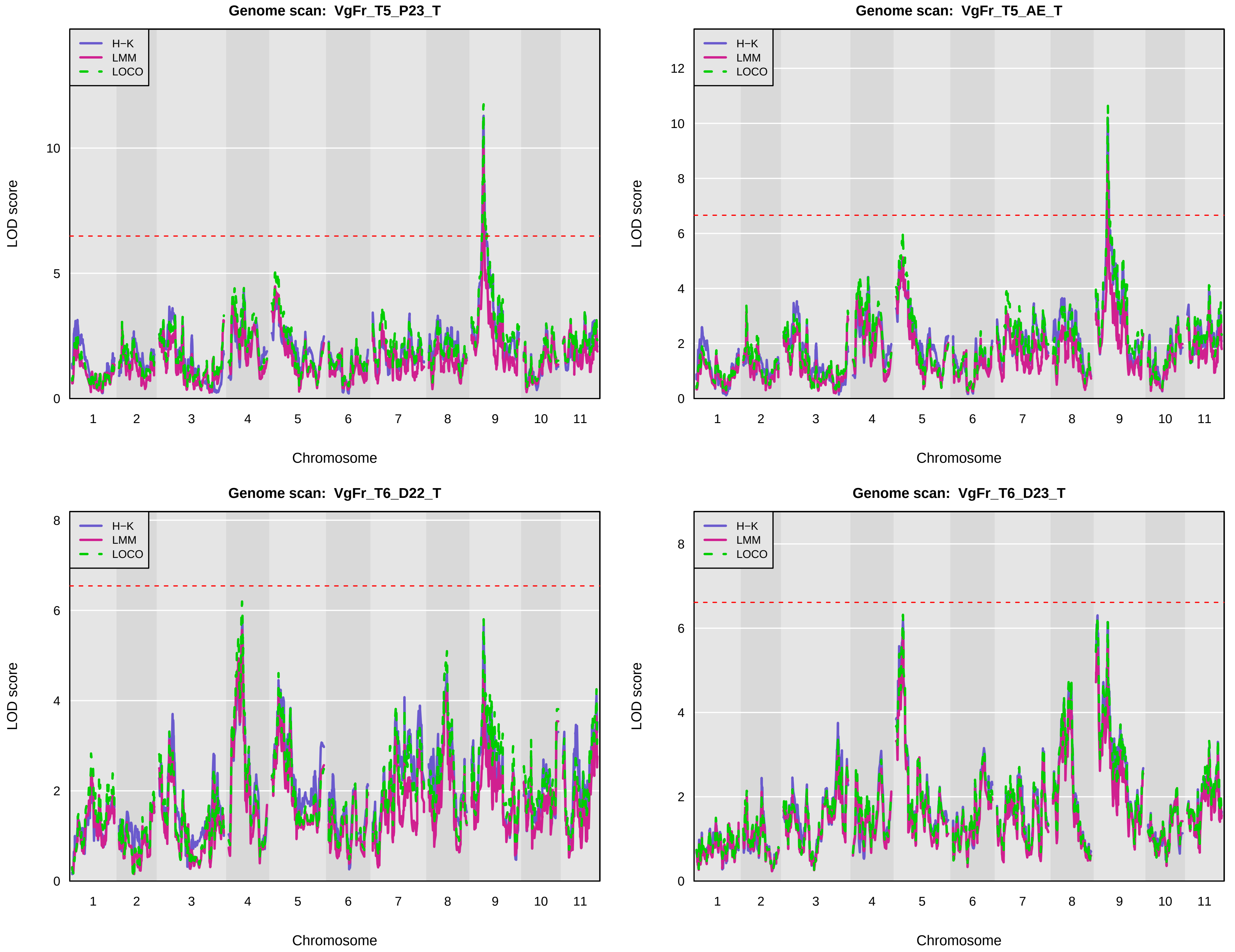

Genome scan: VgFr_T5_P23_T
H−K
LMM
LOCO
LOD score
0
5
10
1
2
3
4
5
6
7
8
9
10
11
Chromosome
Genome scan: VgFr_T5_AE_T
H−K
LMM
LOCO
LOD score
0
2
4
6
8
10
12
1
2
3
4
5
6
7
8
9
10
11
Chromosome
Genome scan: VgFr_T6_D22_T
H−K
LMM
LOCO
LOD score
0
2
4
6
8
1
2
3
4
5
6
7
8
9
10
11
Chromosome
Genome scan: VgFr_T6_D23_T
H−K
LMM
LOCO
LOD score
0
2
4
6
8
1
2
3
4
5
6
7
8
9
10
11
Chromosome

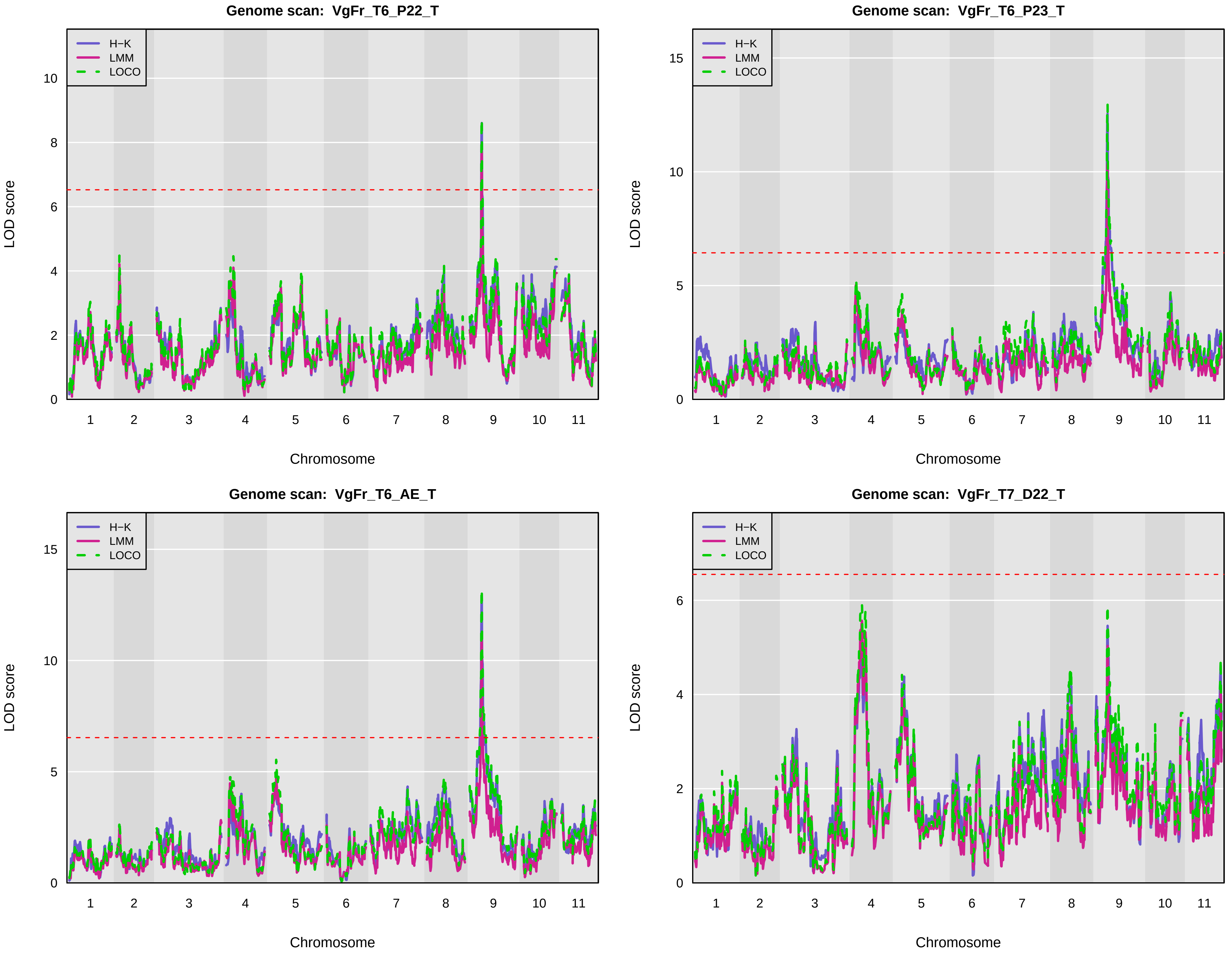
Genome scan: VgFr_T6_P22_T
Genome scan: VgFr_T6_P23_T
Genome scan: VgFr_T6_AE_T
Genome scan: VgFr_T7_D22_T
H−K
LMM
LOCO
LOD score
Chromosome
1 2 3 4 5 6 7 8 9 10 11

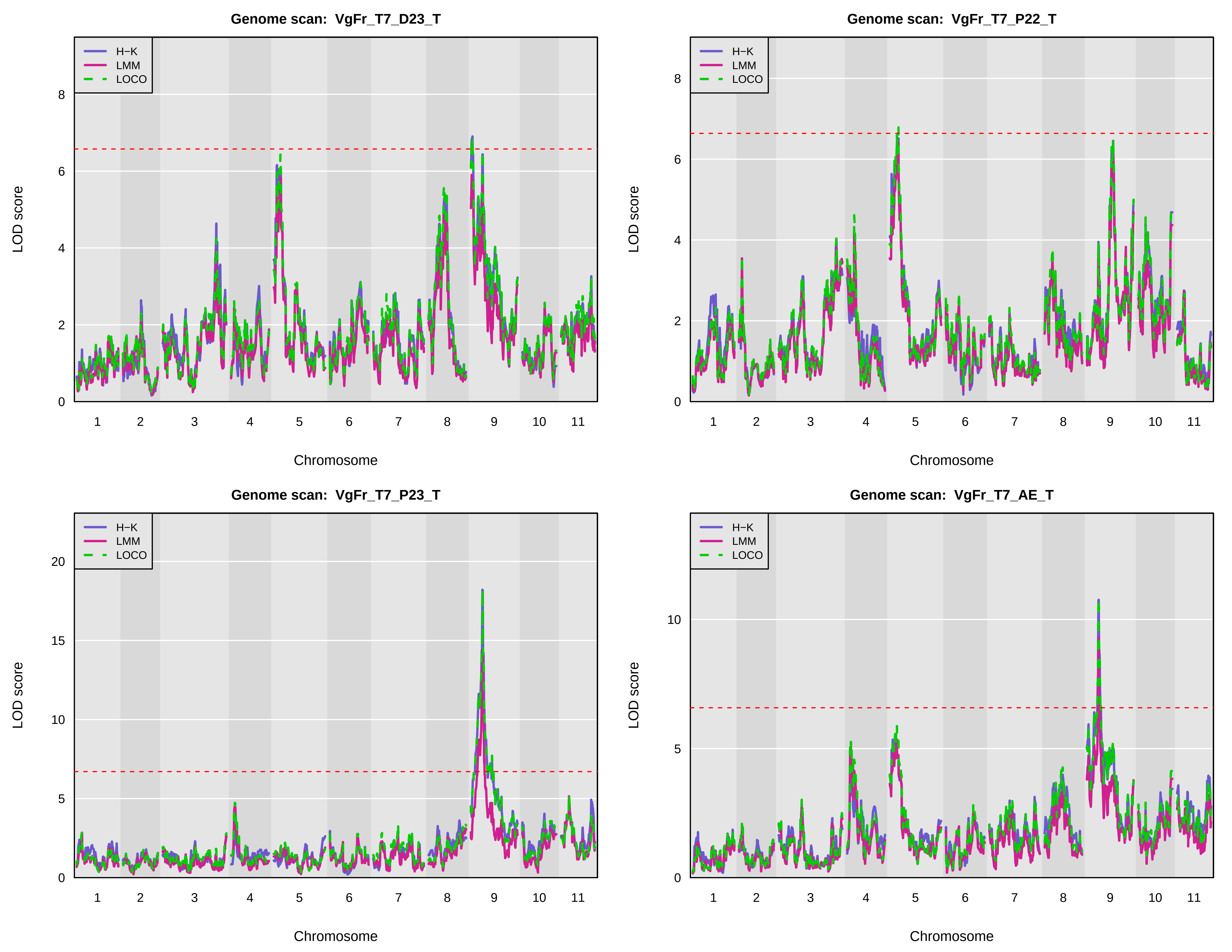
Genome scan: VgFr_T7_D23_T
Genome scan: VgFr_T7_P22_T
Genome scan: VgFr_T7_P23_T
Genome scan: VgFr_T7_AE_T
H−K
LMM
LOCO
LOD score
Chromosome
1 2 3 4 5 6 7 8 9 10 11

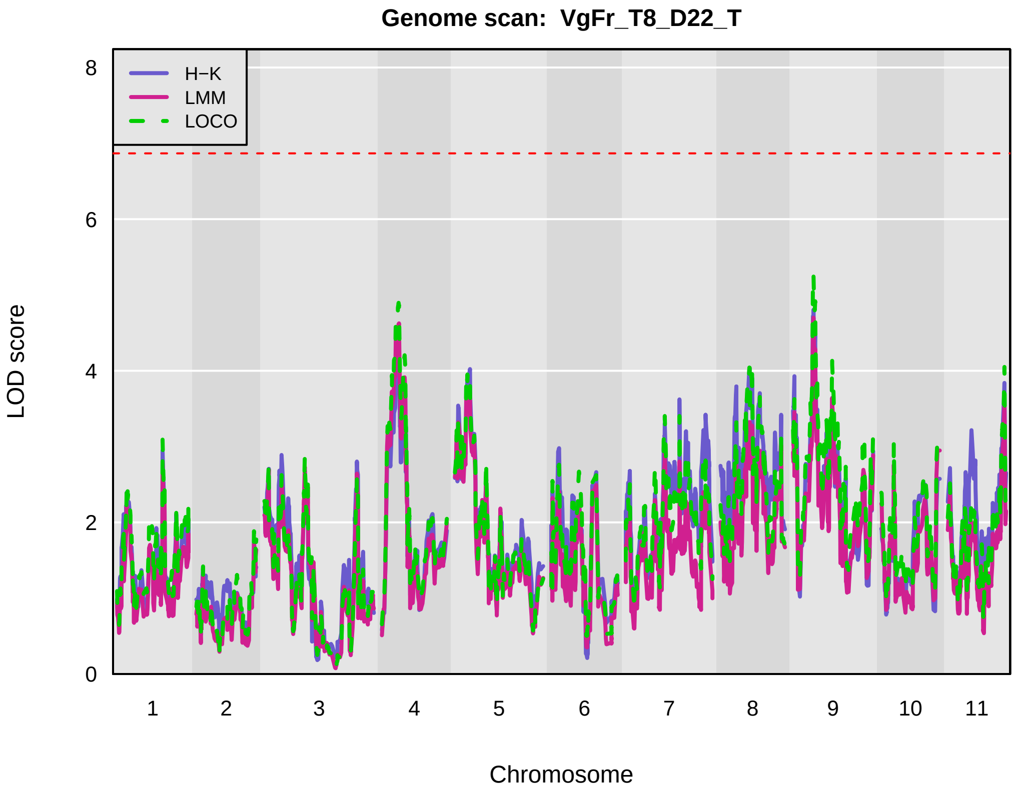
Genome scan: VgFr_T8_D22_T
H−K
LMM
LOCO
LOD score
Chromosome
0
2
4
6
8
1
2
3
4
5
6
7
8
9
10
11

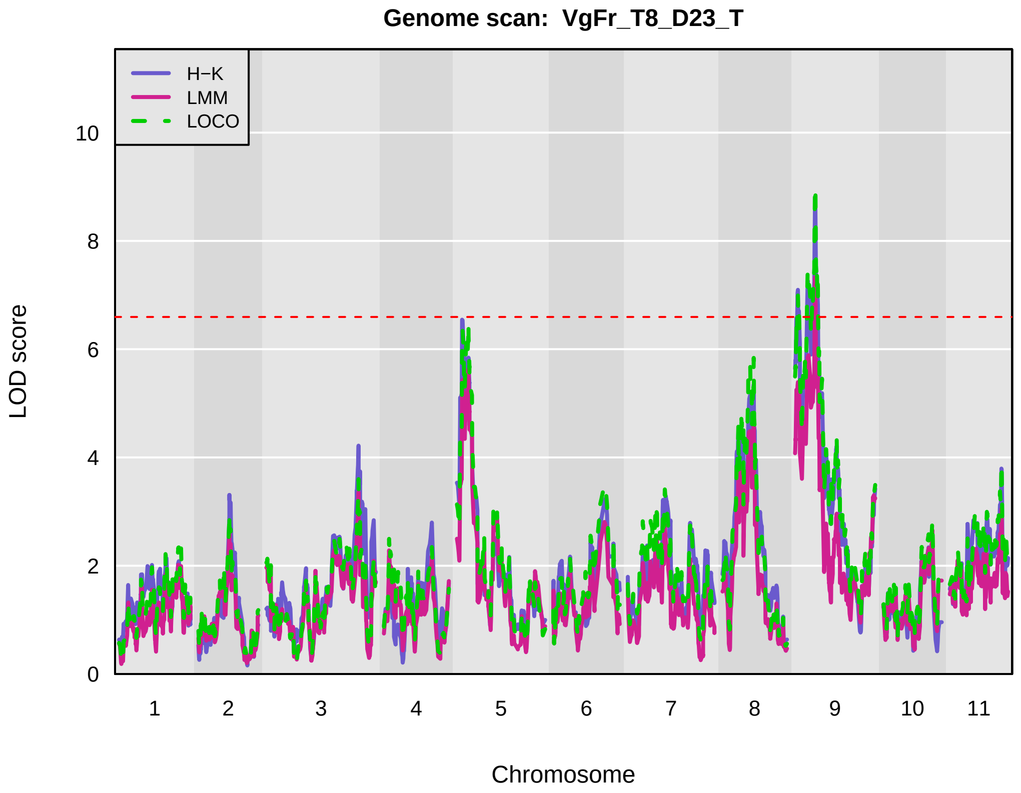
Genome scan: VgFr_T8_D23_T
H−K
LMM
LOCO
LOD score
Chromosome
0
2
4
6
8
10
1
2
3
4
5
6
7
8
9
10
11

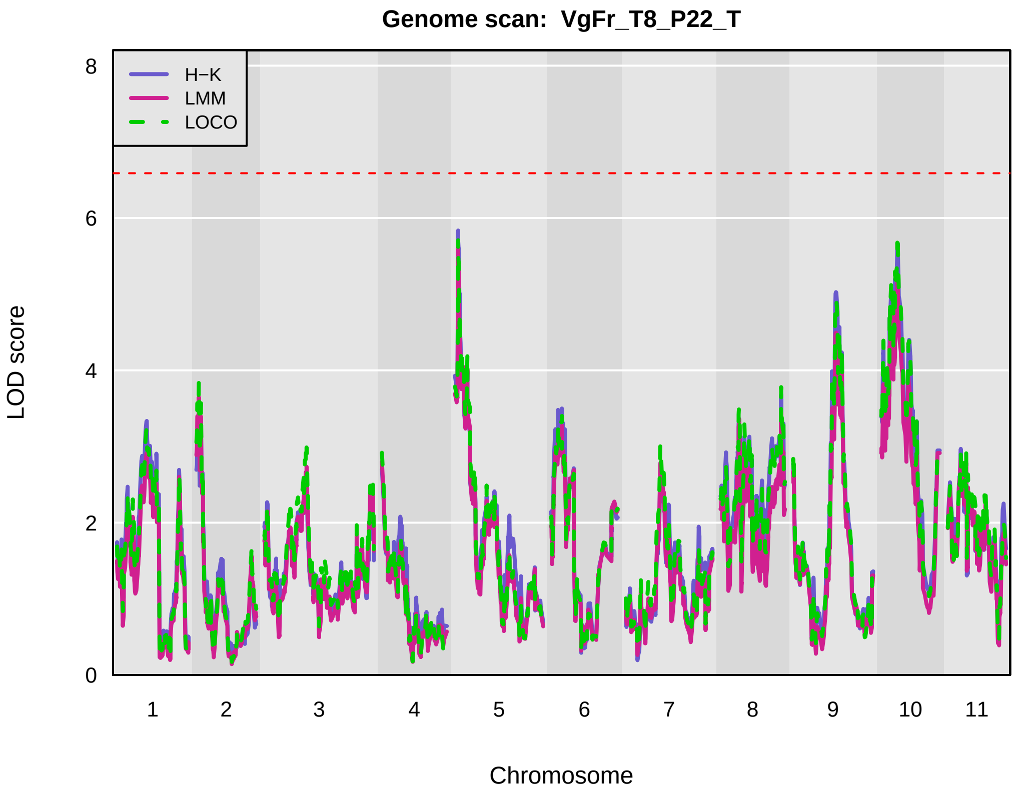
Genome scan: VgFr_T8_P22_T
H−K
LMM
LOCO
LOD score
Chromosome
0
2
4
6
8
1
2
3
4
5
6
7
8
9
10
11

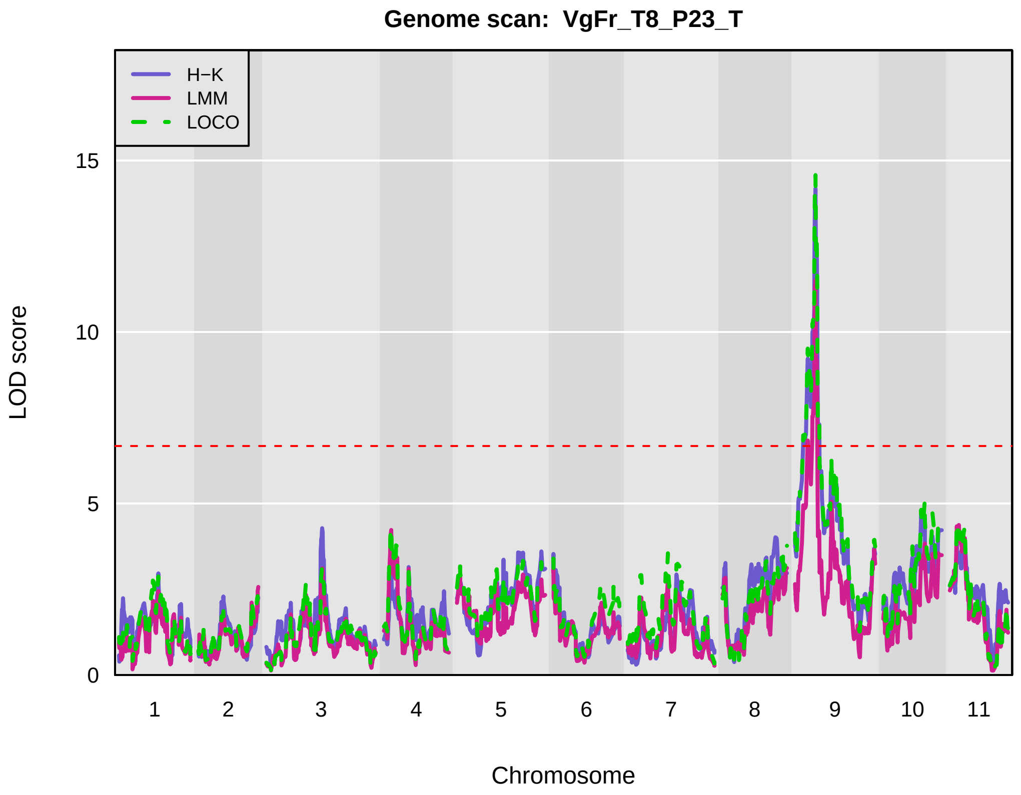
Genome scan: VgFr_T8_P23_T
H−K
LMM
LOCO
LOD score
Chromosome
0
5
10
15
1
2
3
4
5
6
7
8
9
10
11

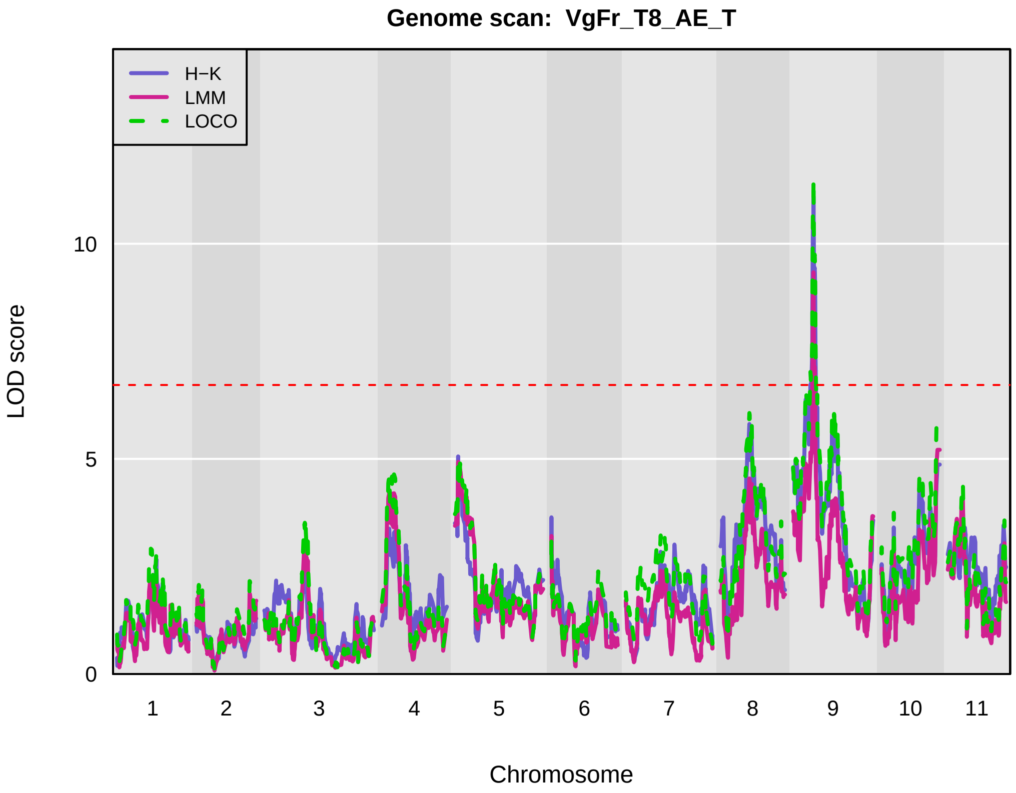
Genome scan: VgFr_T8_AE_T
H−K
LMM
LOCO
0
5
10
LOD score
1
2
3
4
5
6
7
8
9
10
11
Chromosome

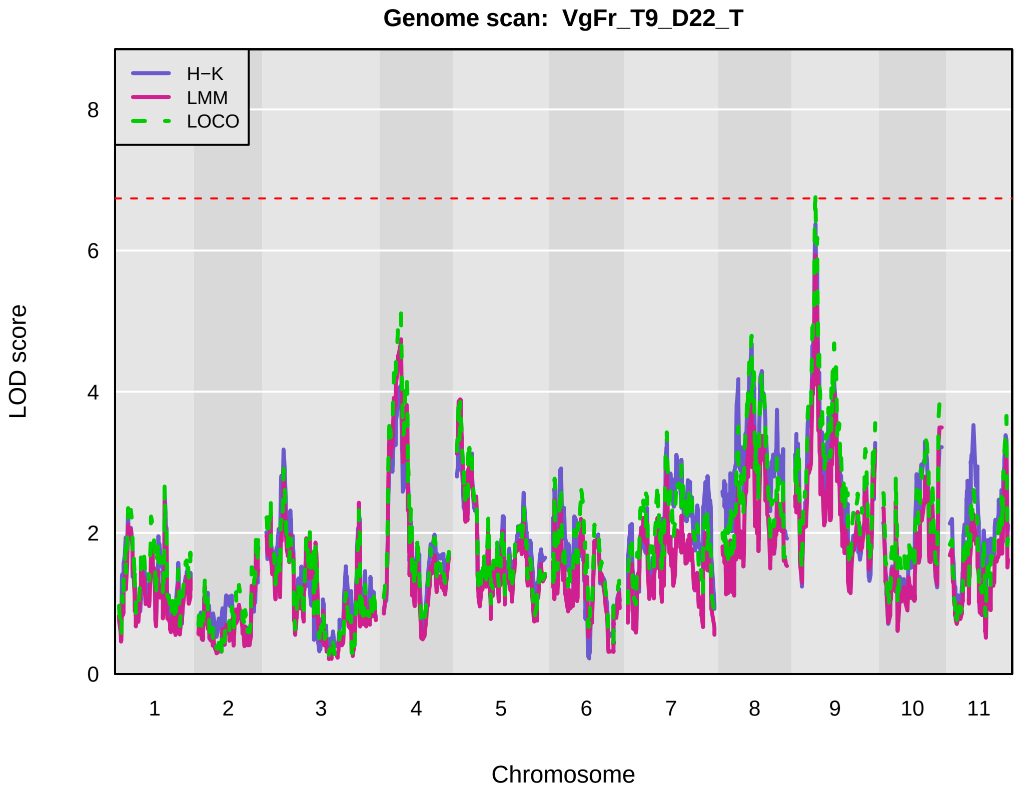
Genome scan: VgFr_T9_D22_T
H−K
LMM
LOCO
0
2
4
6
8
LOD score
1
2
3
4
5
6
7
8
9
10
11
Chromosome

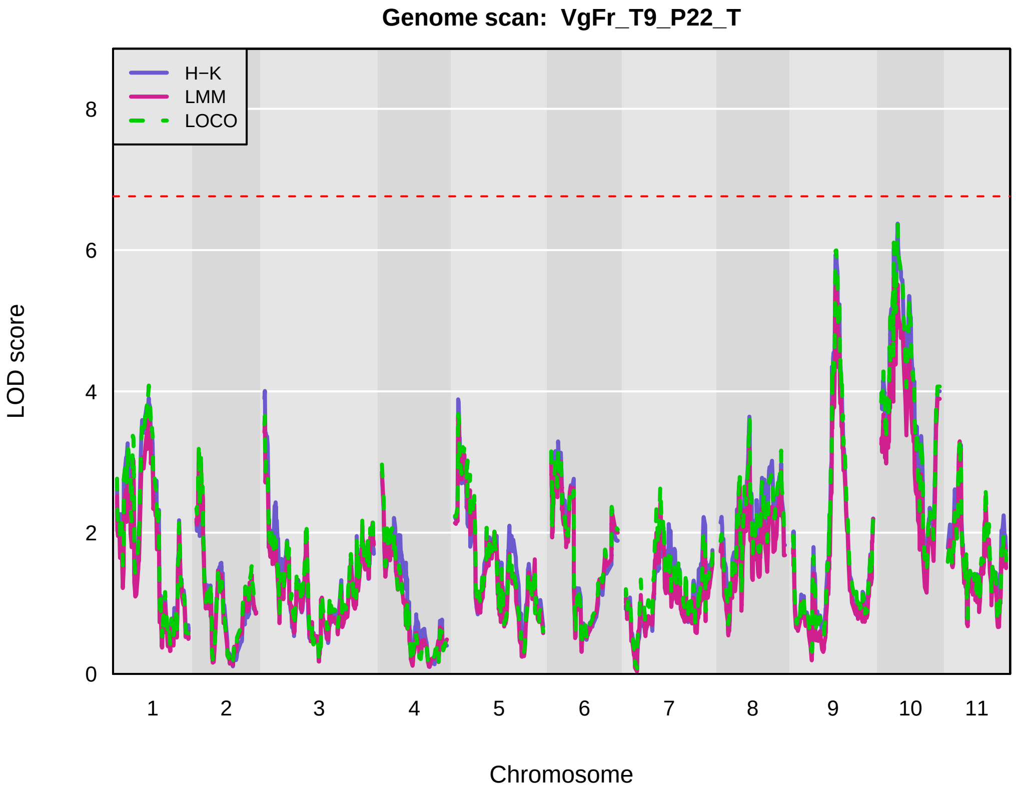
Genome scan: VgFr_T9_P22_T
H−K
LMM
LOCO
0
2
4
6
8
LOD score
1
2
3
4
5
6
7
8
9
10
11
Chromosome

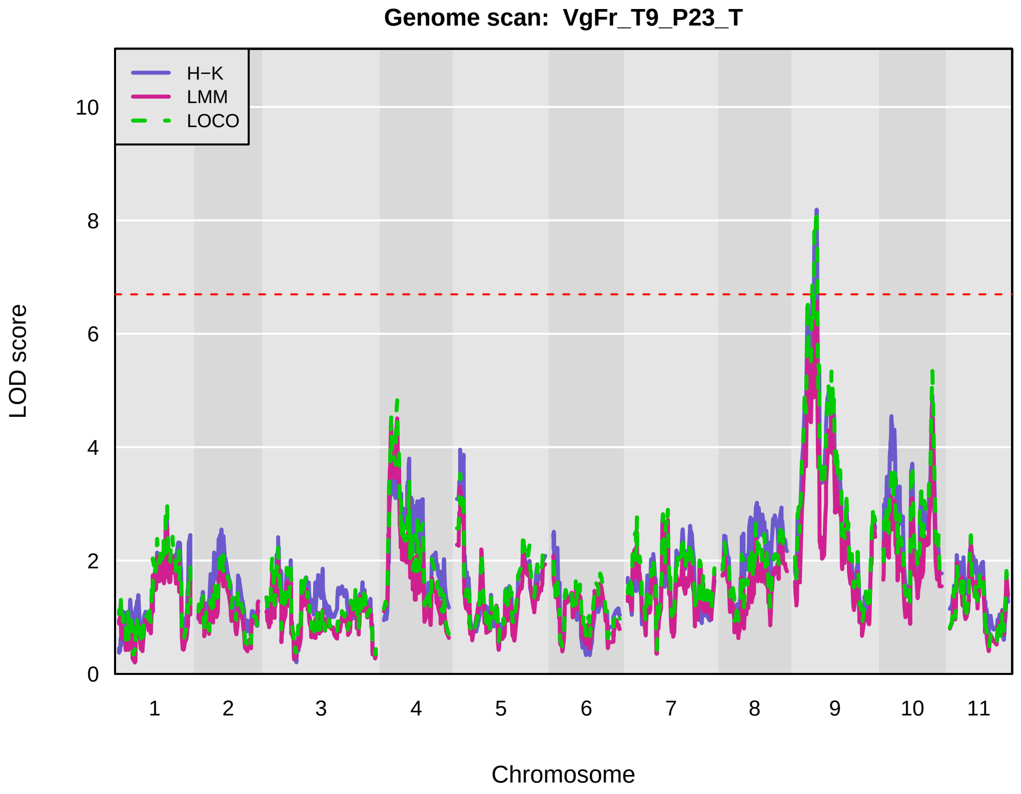
Genome scan: VgFr_T9_P23_T
H−K
LMM
LOCO
0
2
4
6
8
10
LOD score
1
2
3
4
5
6
7
8
9
10
11
Chromosome

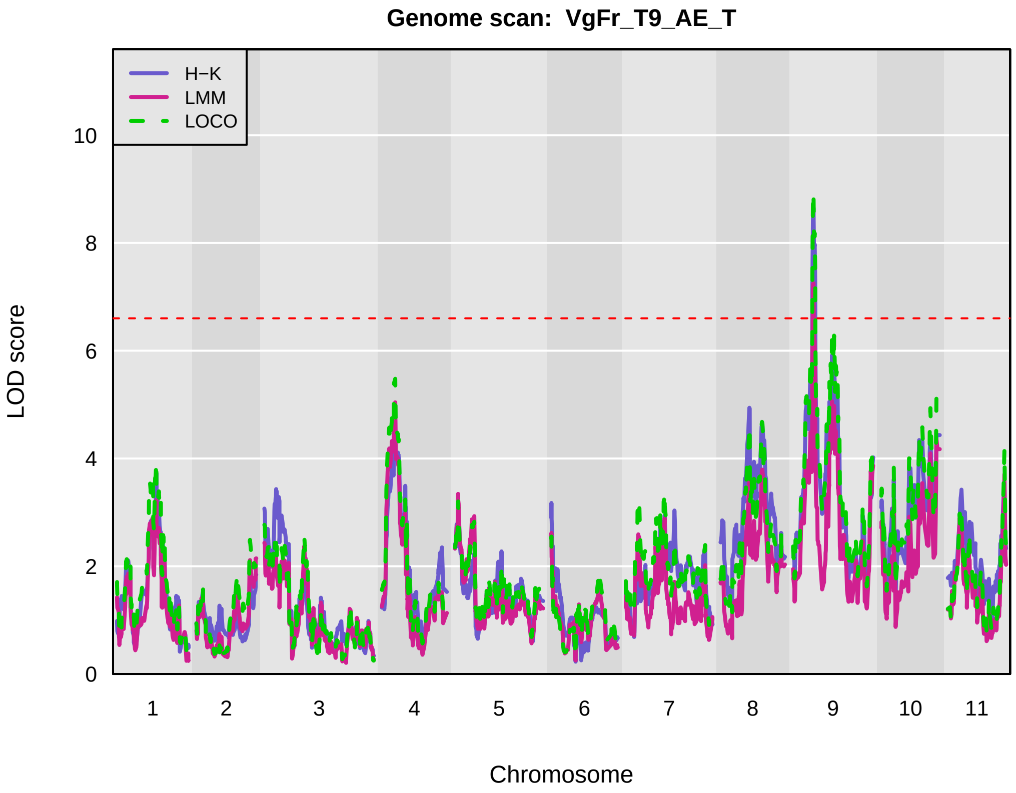
Genome scan: VgFr_T9_AE_T
H−K
LMM
LOCO
LOD score
0
2
4
6
8
10
1
2
3
4
5
6
7
8
9
10
11
Chromosome

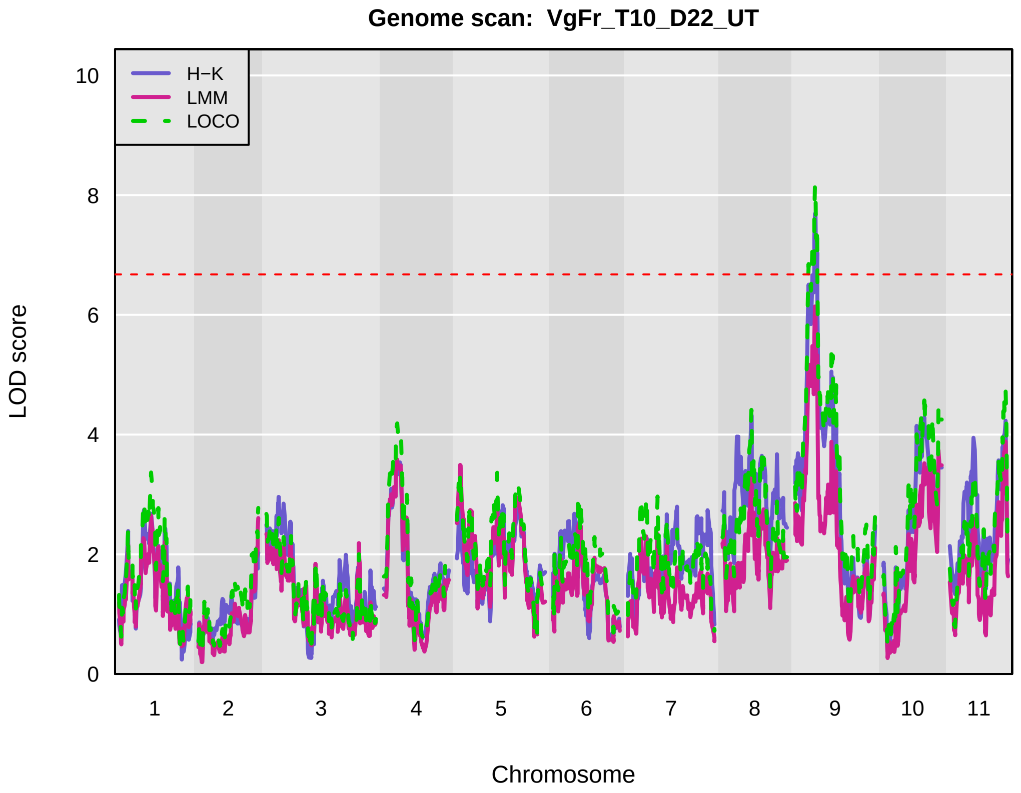
Genome scan: VgFr_T10_D22_UT
H−K
LMM
LOCO
LOD score
0
2
4
6
8
10
1
2
3
4
5
6
7
8
9
10
11
Chromosome

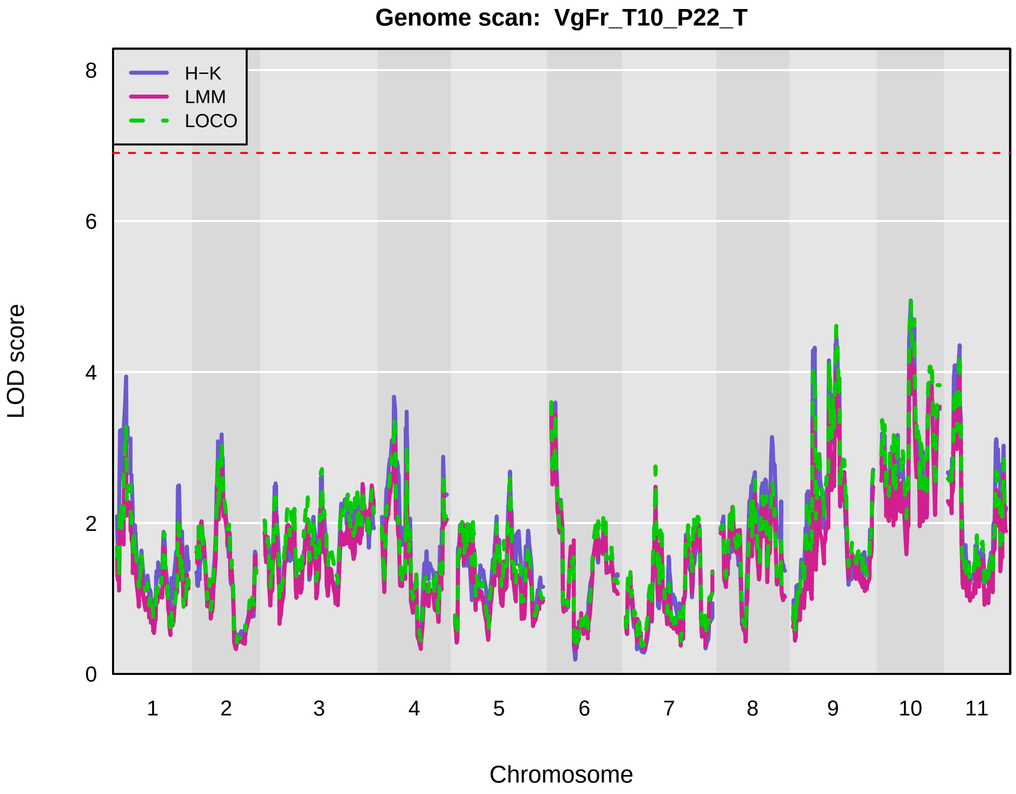
Genome scan: VgFr_T10_P22_T
H−K
LMM
LOCO
LOD score
0
2
4
6
8
1
2
3
4
5
6
7
8
9
10
11
Chromosome
