## Supplementary figures and images for "Genetic mapping and genomic prediction for agronomic, grain compositional, and sensing-enabled traits in a cowpea MAGIC population along an environmental gradient"

### SuppFig_04_AcrossEnv_QTLprofiles_No_BT_Filtered_Renamed_4perPage_RightLegend 2.pdf

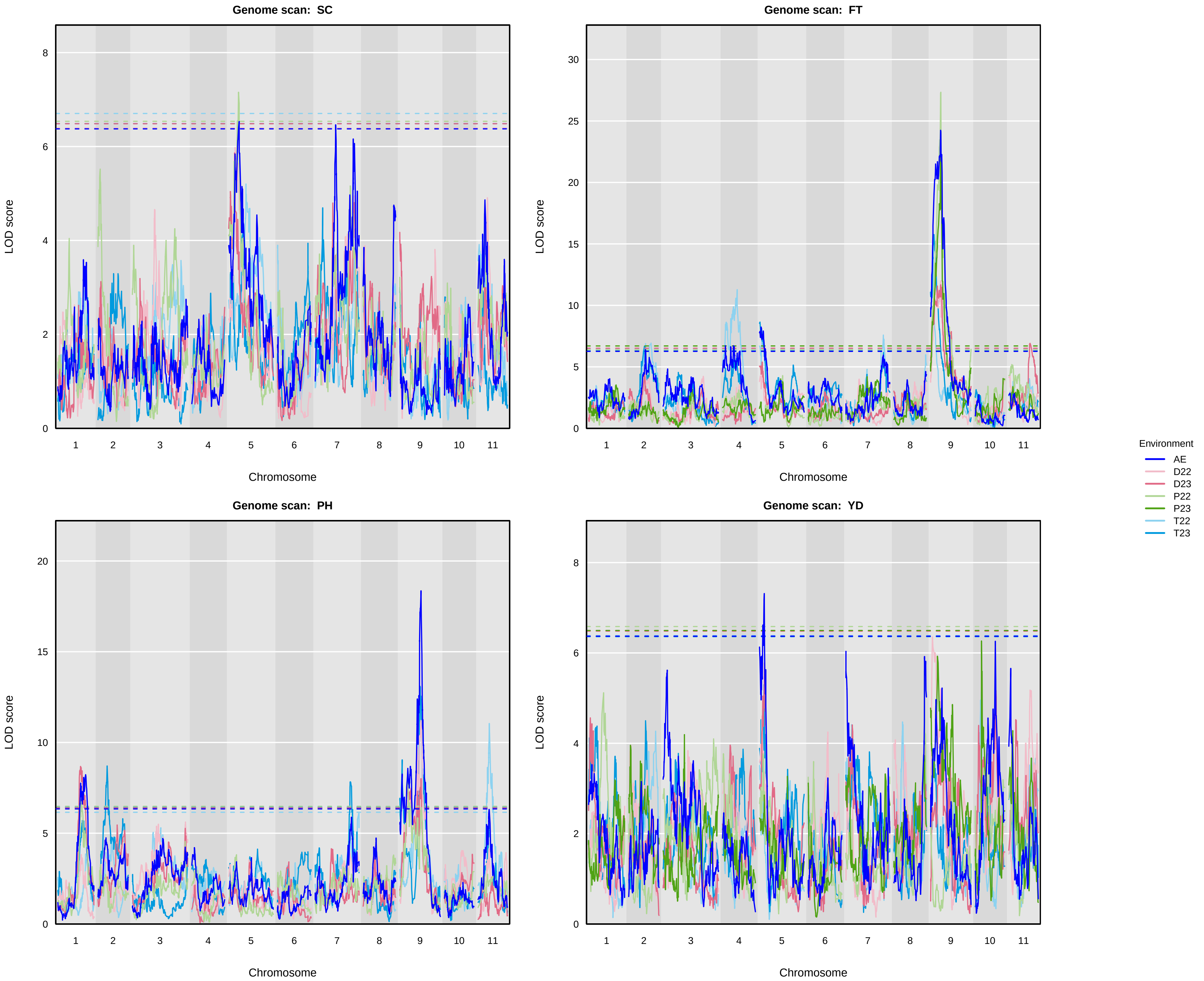

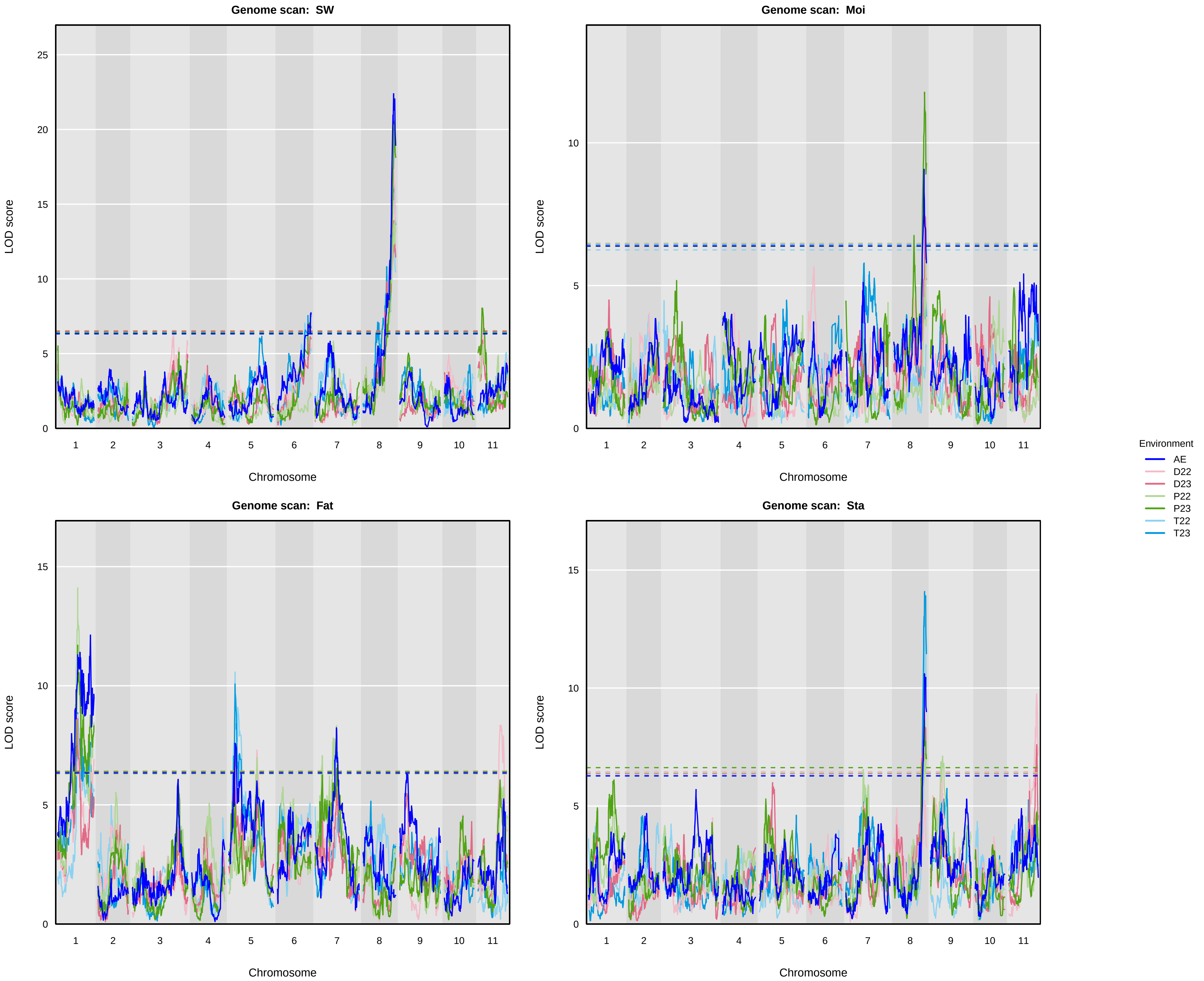

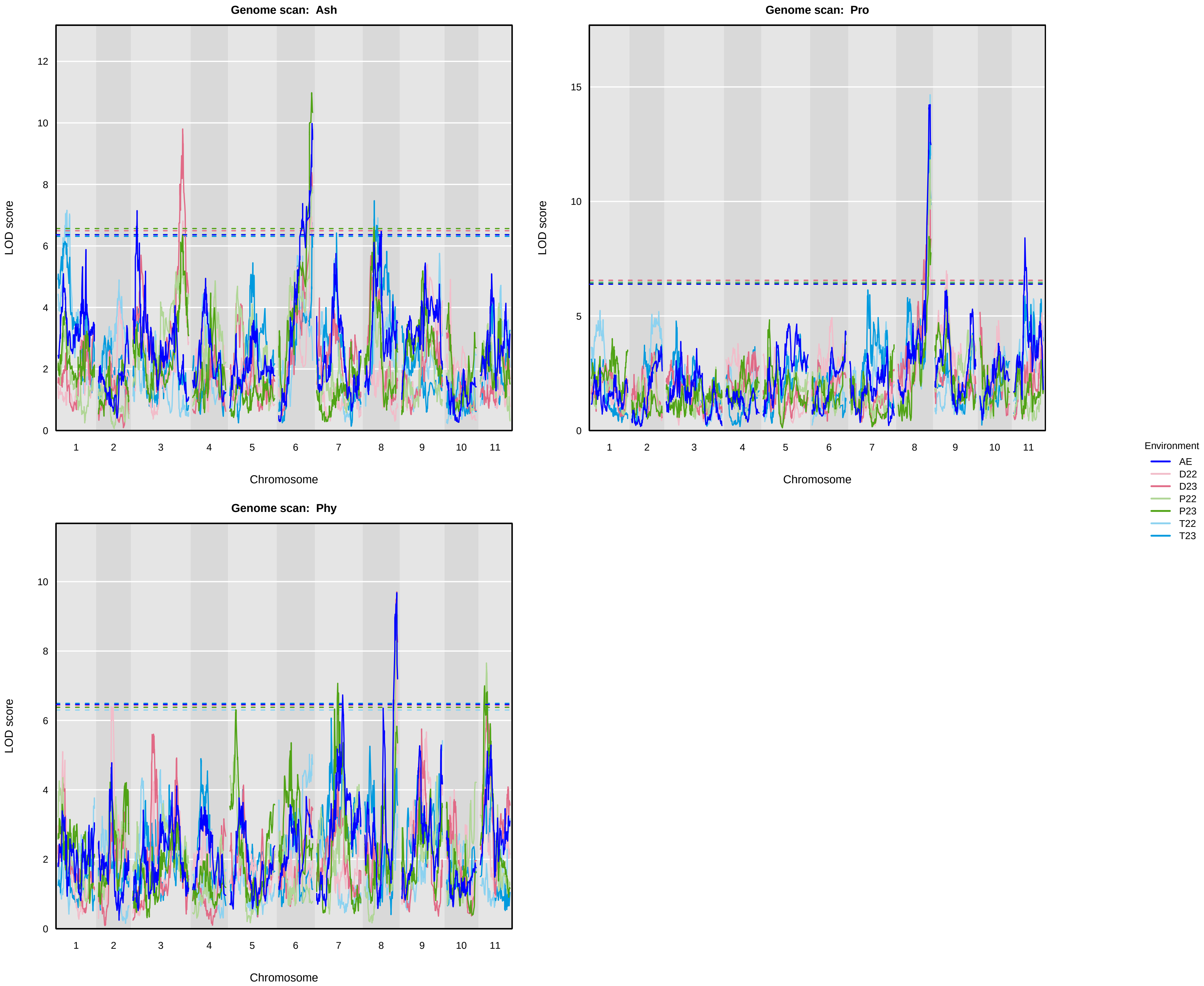

### SuppFig_06_QTLprofiles_Sensing_Traits_Across_Timepoints_Filtered_Renamed_4perPage_RightLegend.pdf

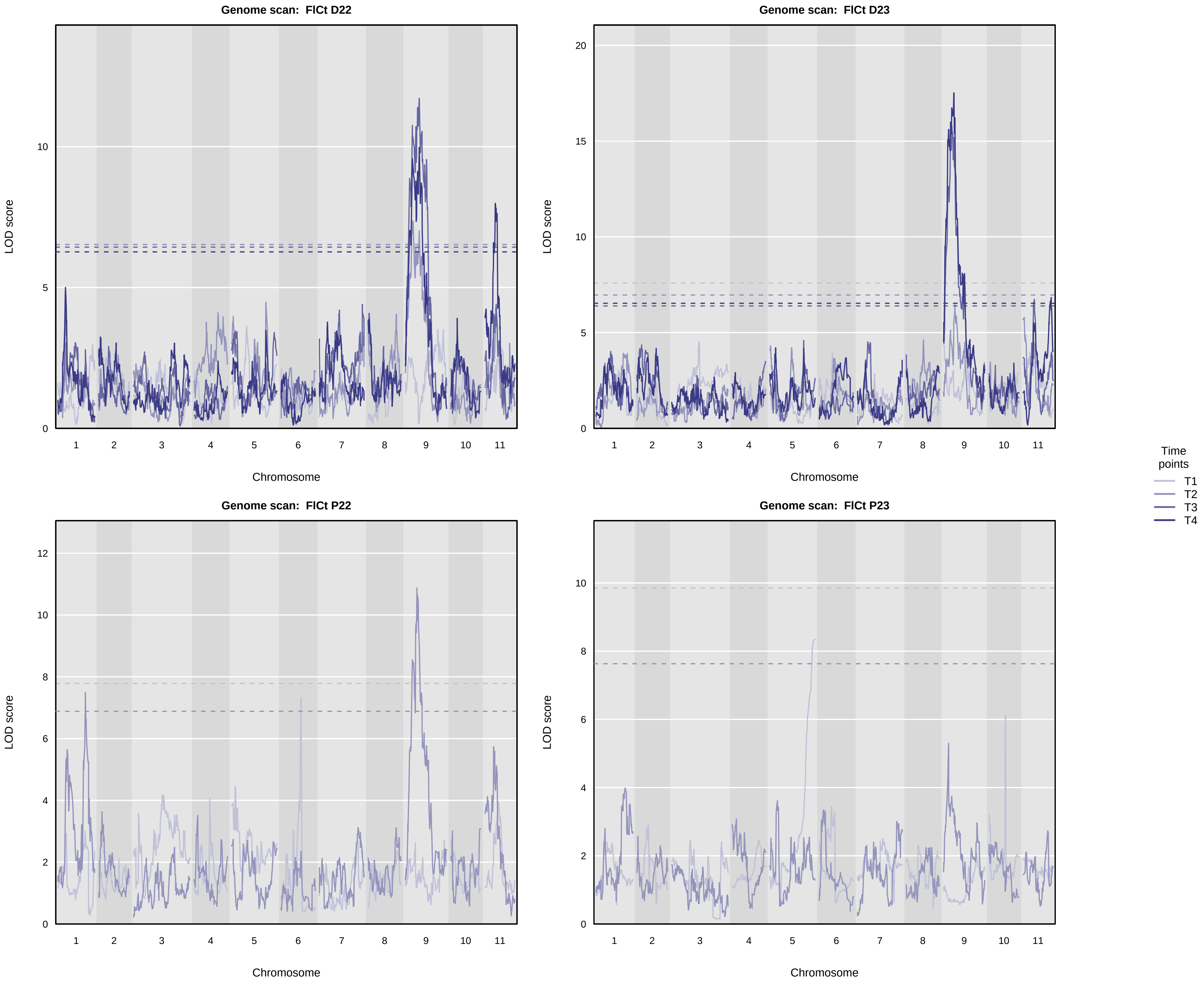

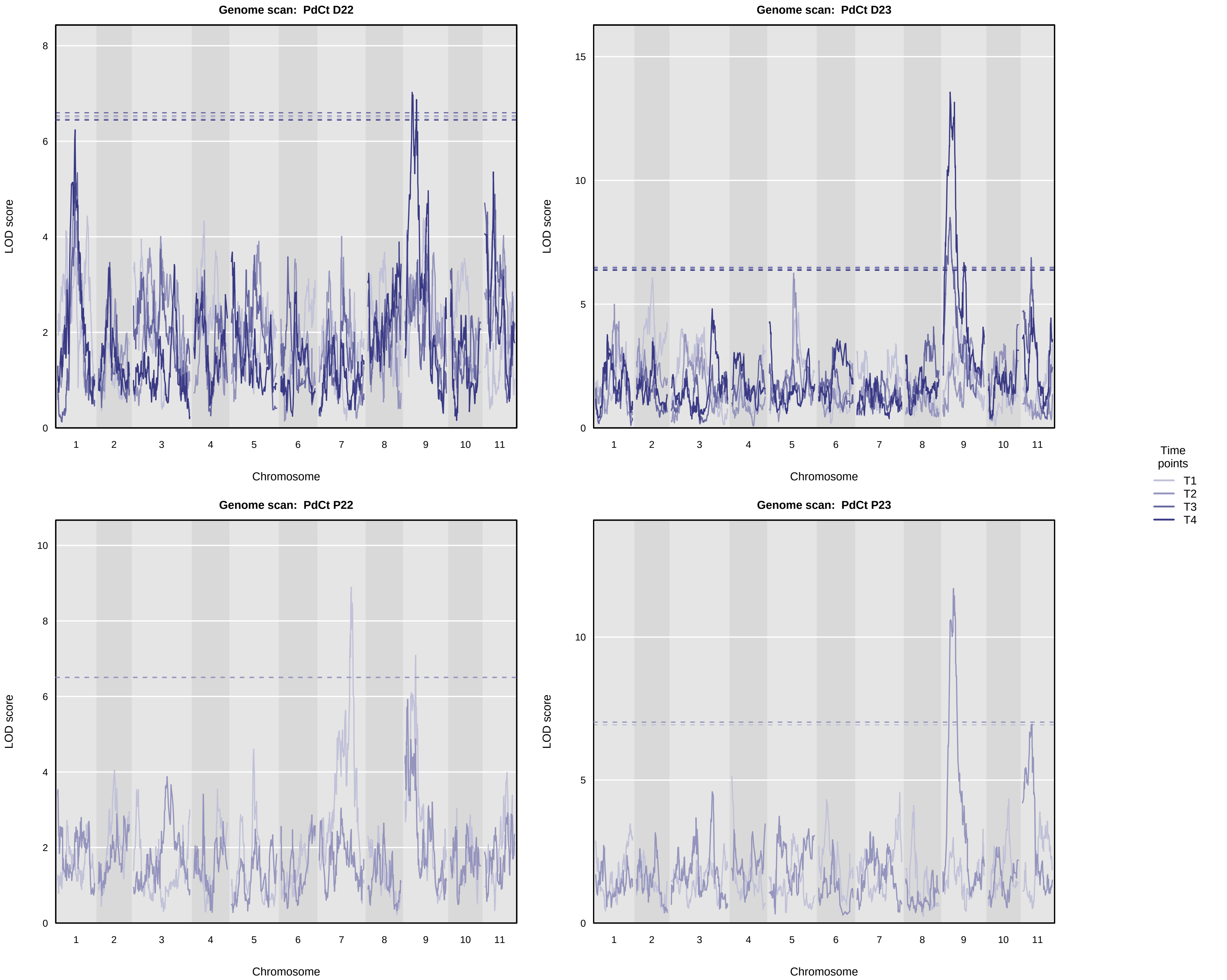

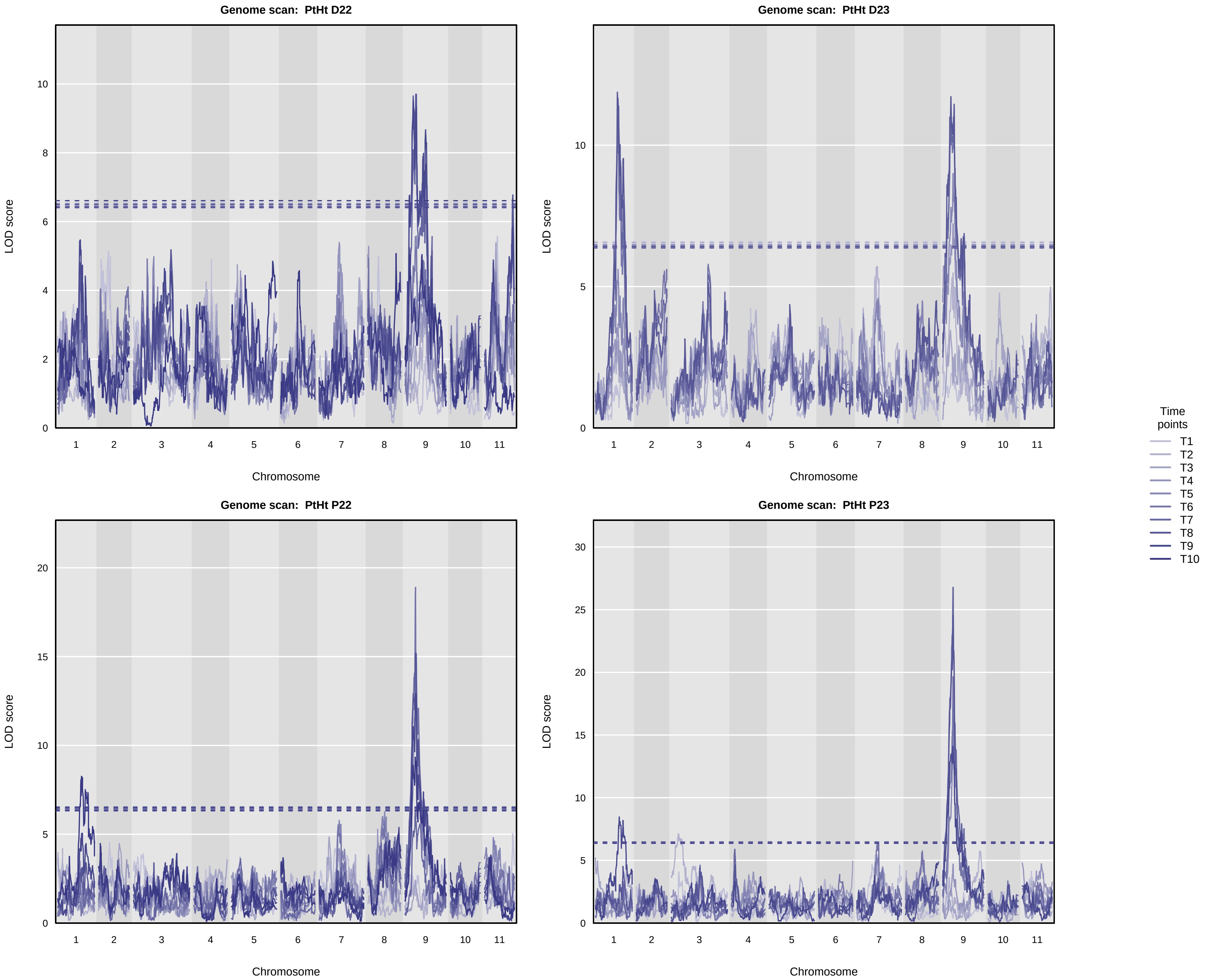

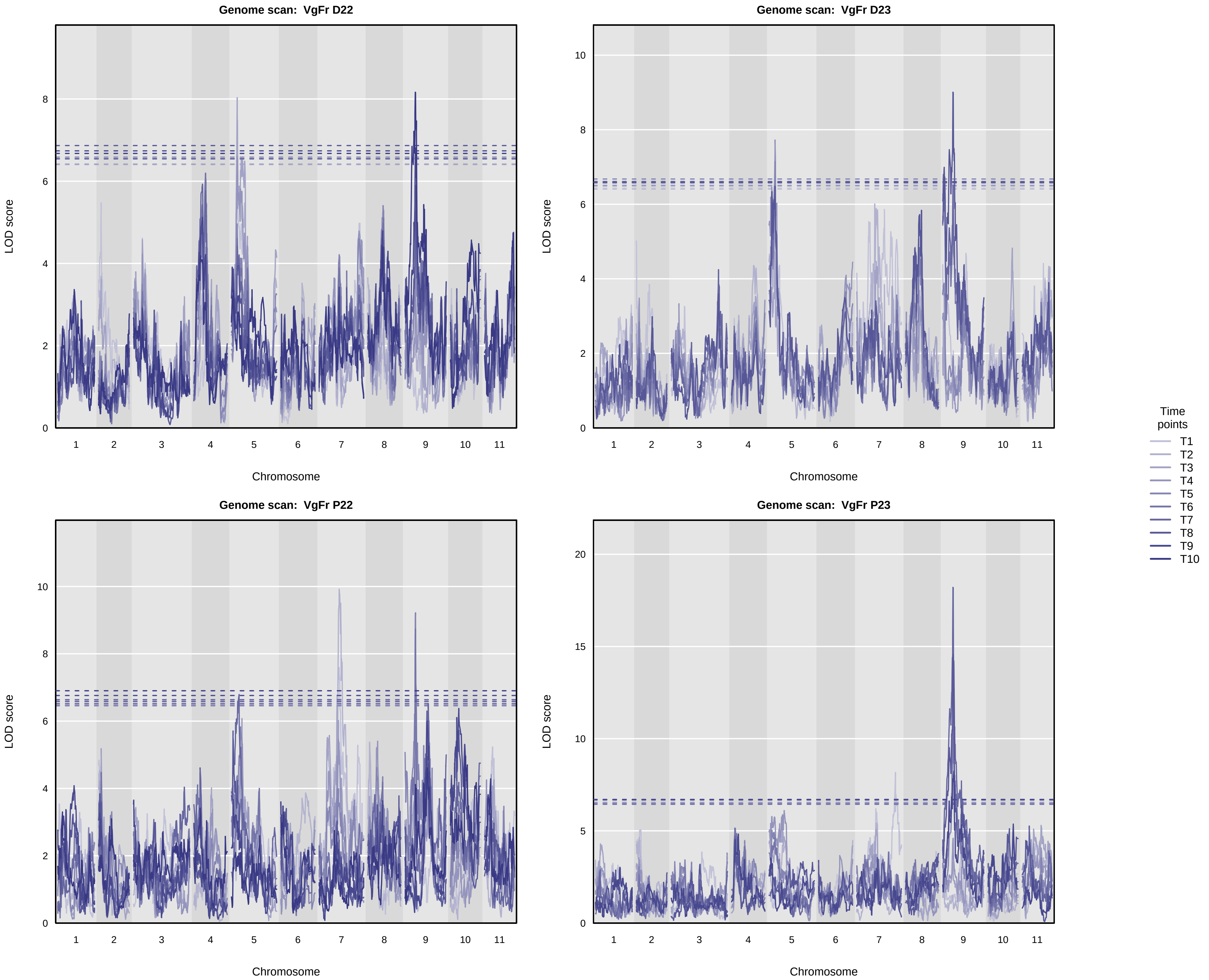

### SuppFig_07_AcrossEnv_QTLprofiles_SWconditional_Chr8.pdf

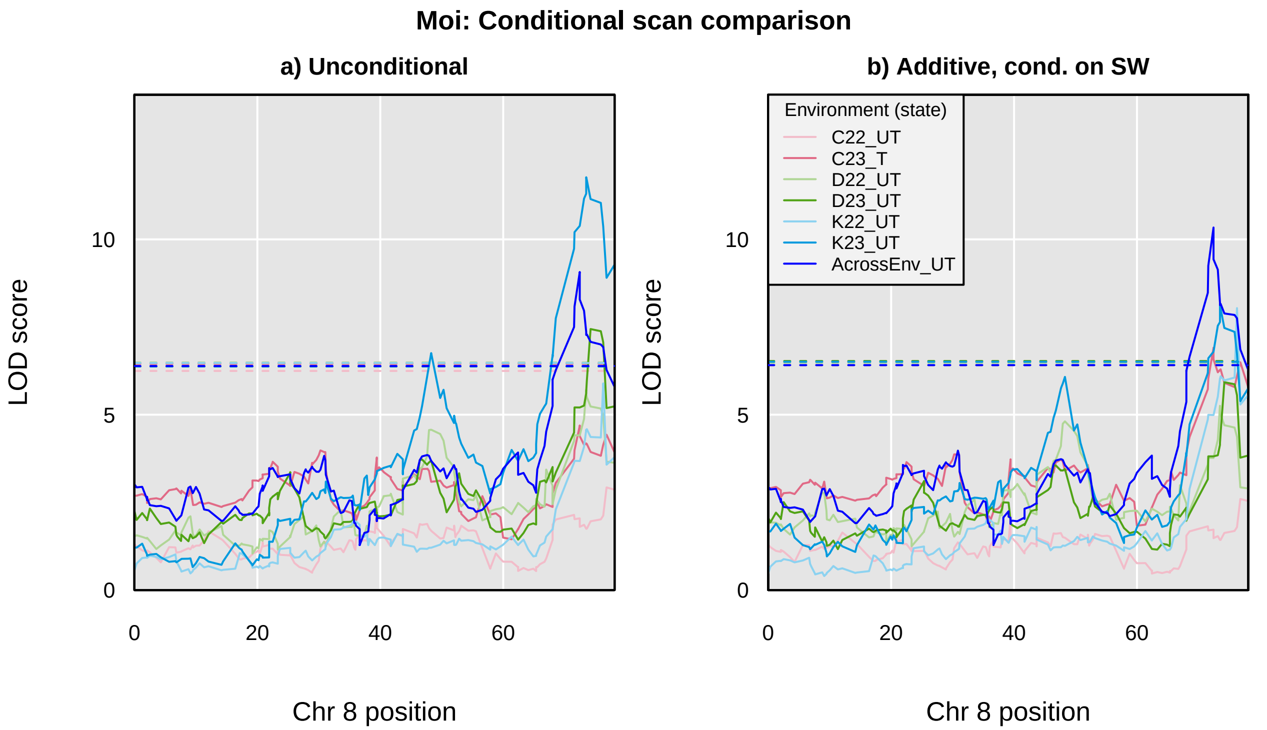

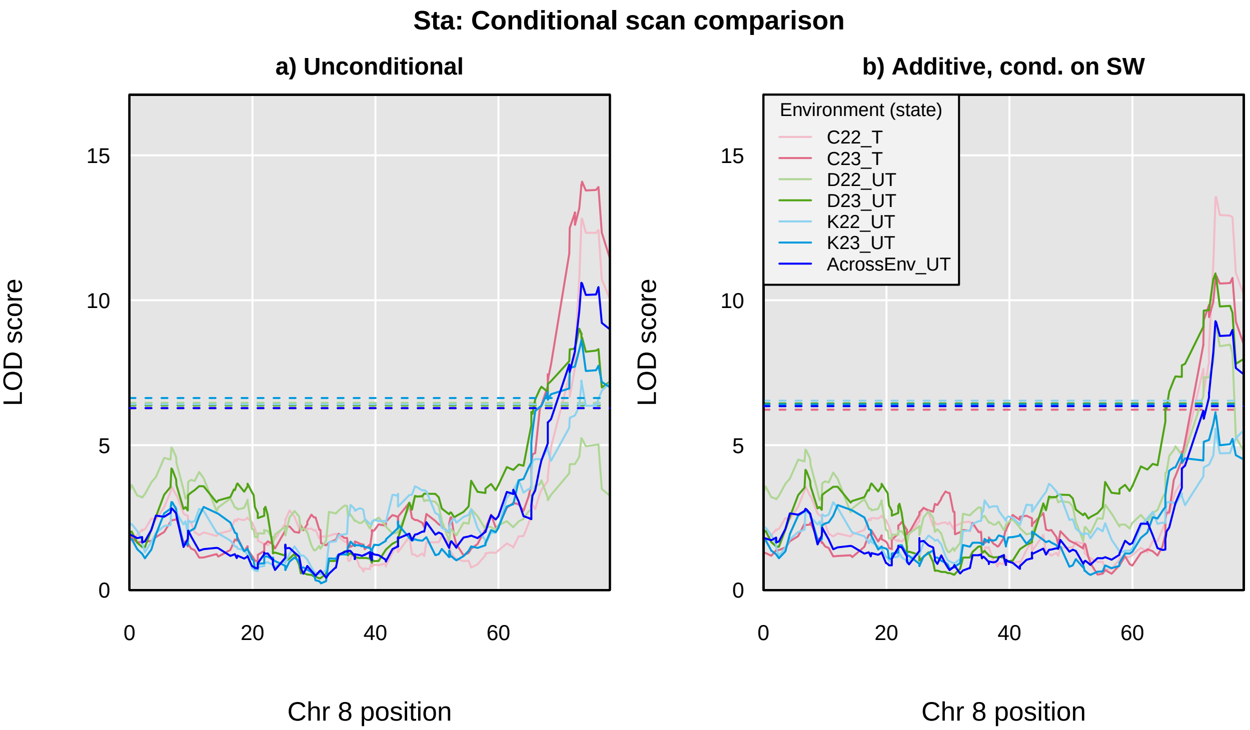

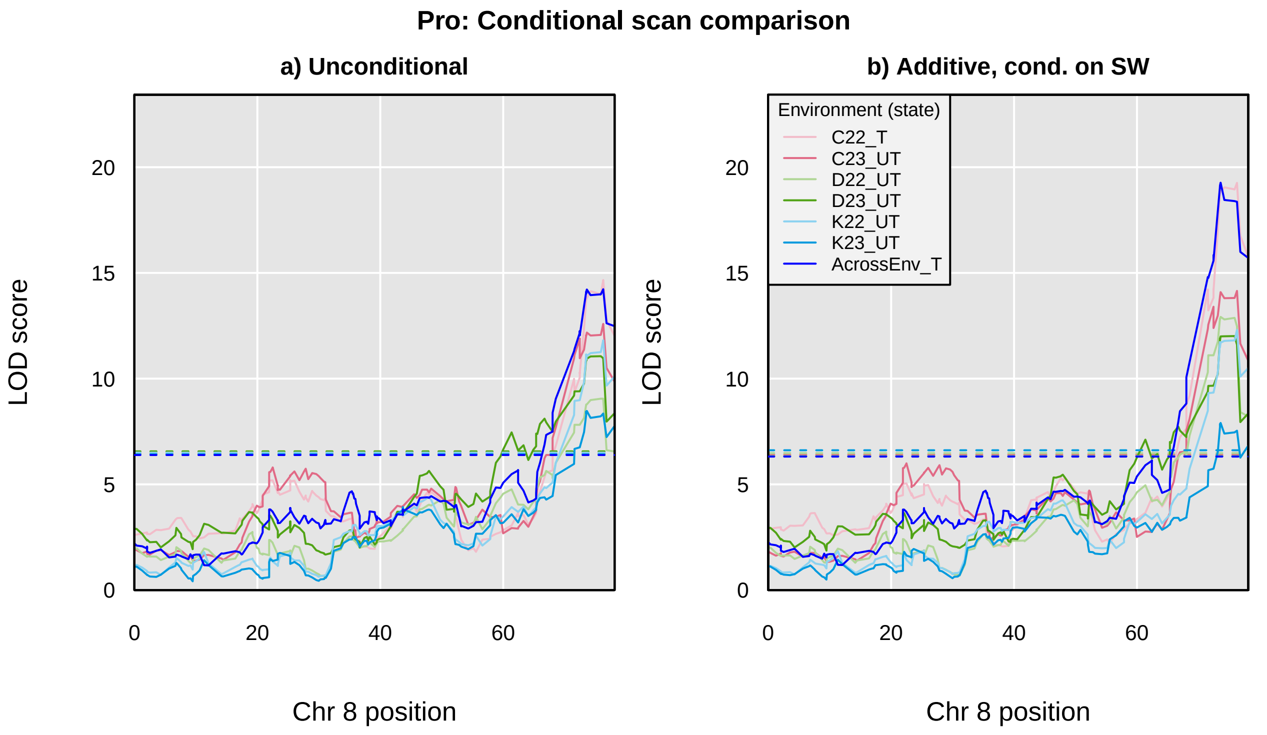

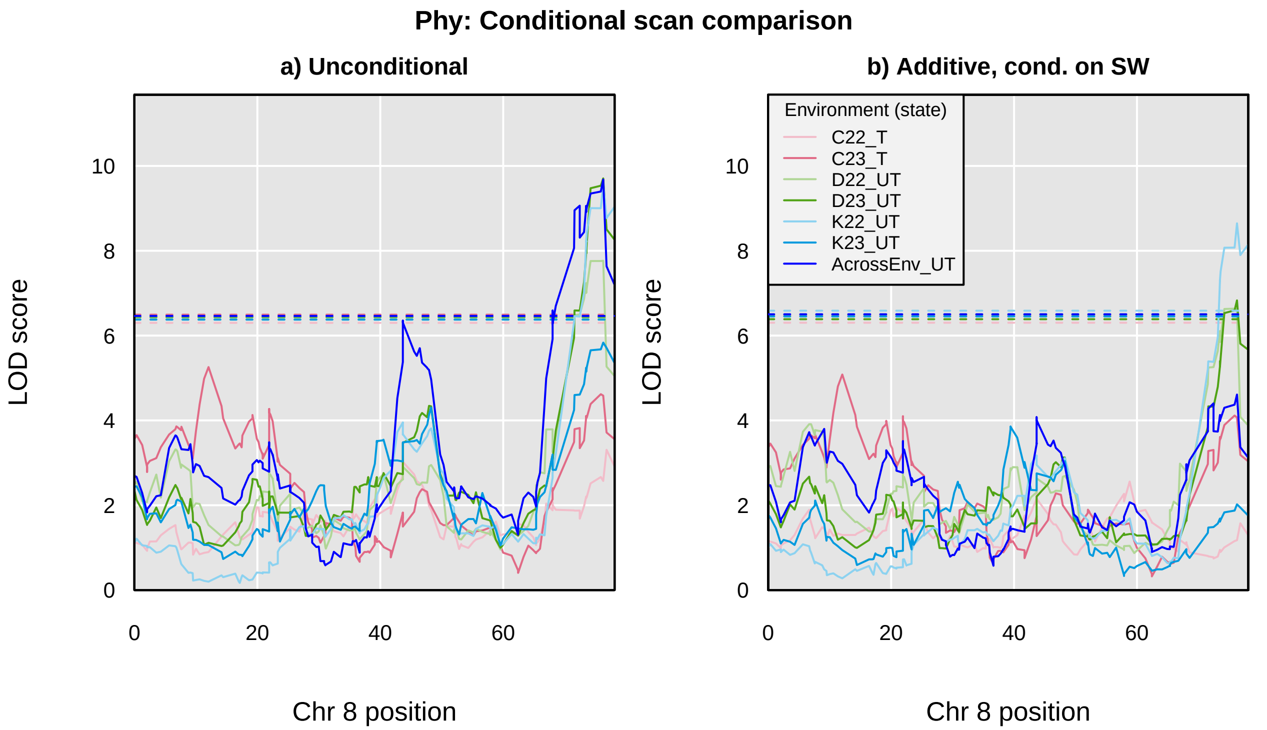

### SuppFig_08_AcrossEnv_QTLprofiles_SWconditional.pdf

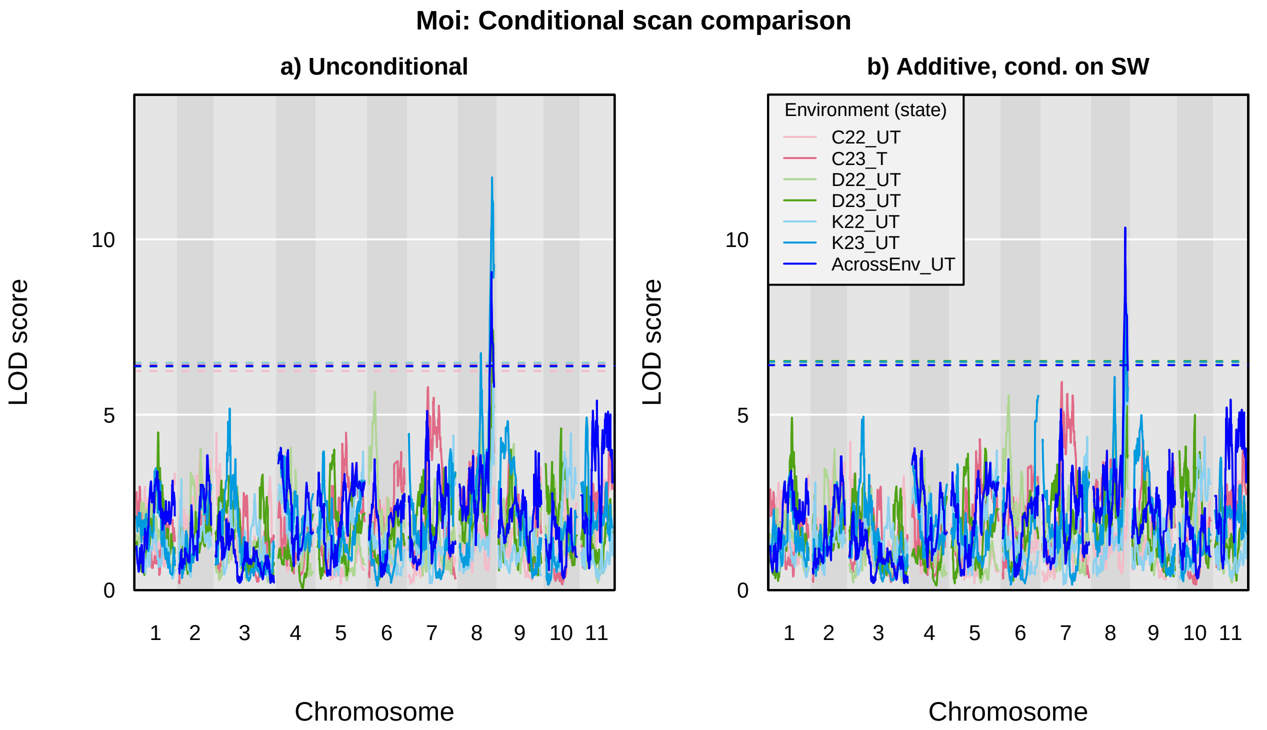

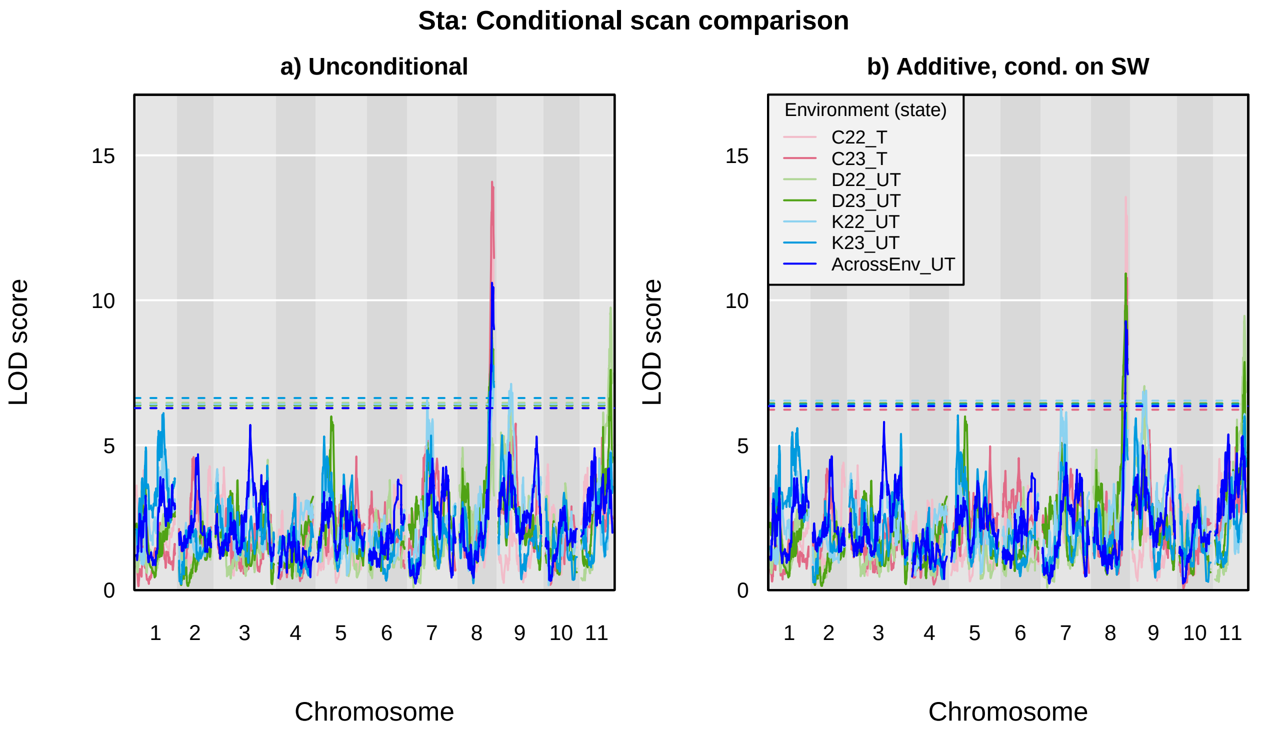

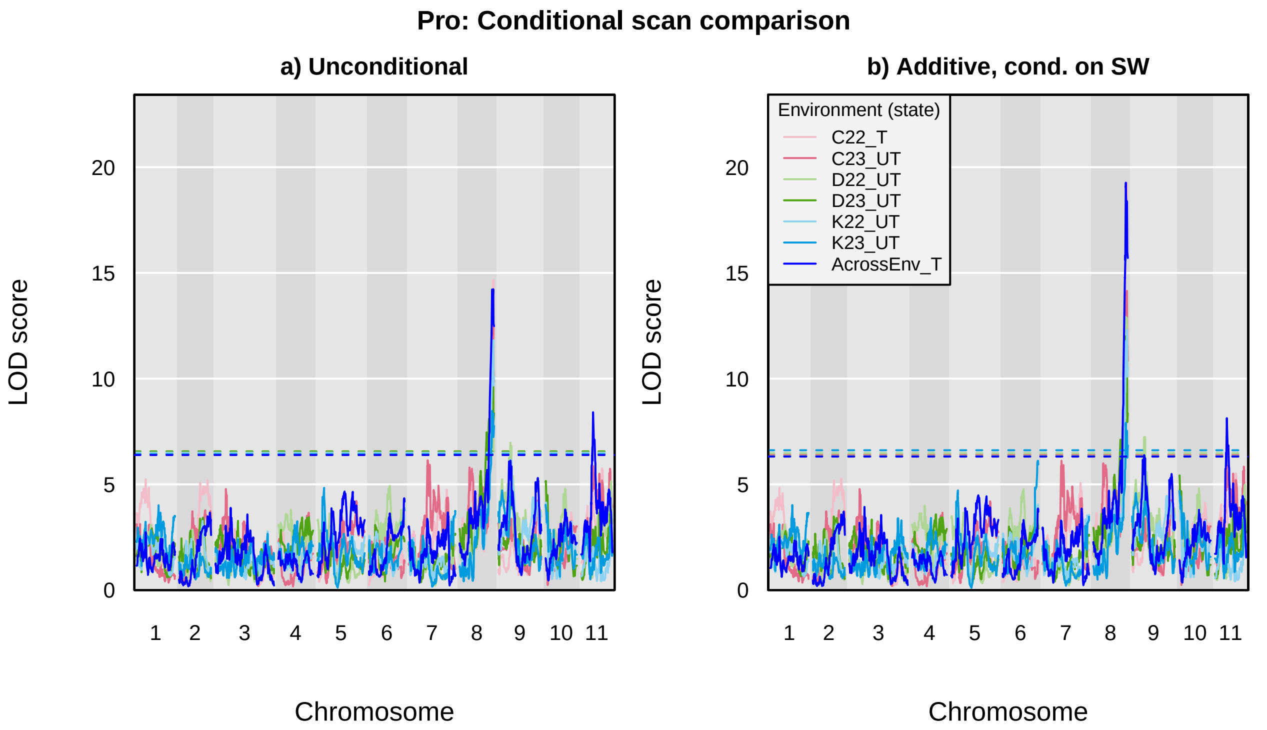

**Phy: Conditional scan comparison**

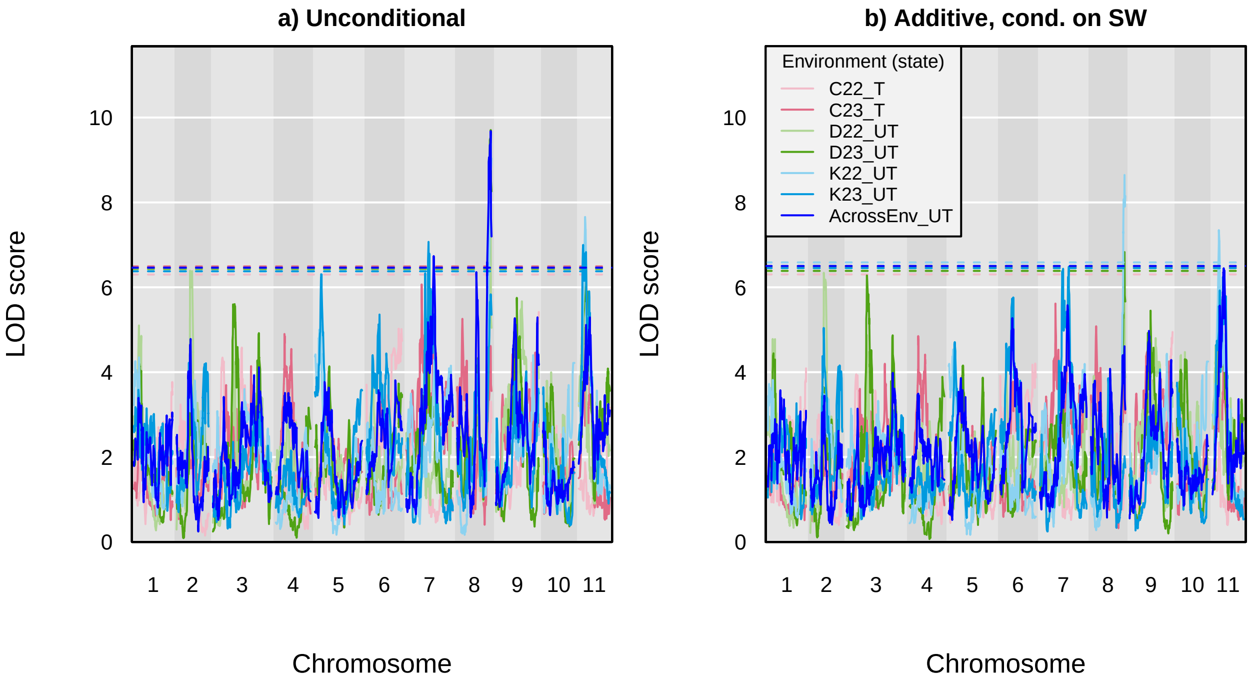
